# Asymmetrical Bi-antennary Glycans Prepared by a Stop-and-Go Strategy Reveal Receptor Binding Evolution of Human Influenza A Viruses

**DOI:** 10.1101/2023.11.08.566285

**Authors:** Shengzhou Ma, Lin Liu, Dirk Eggink, Sander Herfst, Ron A.M. Fouchier, Robert P. de Vries, Geert-Jan Boons

## Abstract

Glycan binding properties of respiratory viruses have been difficult to probe due to a lack of biological relevant glycans for binding studies. Here, a stop-and-go chemoenzymatic methodology is presented that gave access to a panel of 32 asymmetrical bi-antennary *N*-glycans having various numbers of *N*-acetyl lactosamine (LacNAc) repeating units capped by α2,3- or α2,6-sialosides resembling structures found in airway tissues. It exploits that the branching enzymes MGAT1 and MGAT2 can utilize unnatural UDP-2-deoxy-2-trifluoro-*N*-acetamido-glucose (UDP-GlcNTFA) as donor. The TFA moiety of the resulting glycans can be hydrolyzed to give GlcNH_2_ at one of the antennae that temporarily blocks extension by glycosyl transferases. The *N*-glycans were printed as a microarray that was probed for receptor binding specificities of evolutionary distinct human A(H3N2) and A(H1N1)pdm09 viruses. It was found that not only the sialoside type but also the length of the LacNAc chain and presentation at the α1,3-antenna of *N*-glycans is critical for binding. Early A(H3N2) viruses bound to 2,6-sialosides at a single LacNAc moiety at the α1,3-antenna whereas later viruses required the sialoside to be presented at a tri-LacNAc moiety. Surprisingly, most of the A(H3N2) viruses that appeared after 2021 regained binding capacity to sialosides presented at a di-LacNAc moiety. As a result, these viruses agglutinate erythrocytes again, commonly employed for antigenic characterization of influenza viruses. Human A(H1N1)pdm09 viruses have similar receptor binding properties as recent A(H3N2) viruses. The data indicates that an asymmetric *N*-glycan having 2,6-sialoside at a di-LacNAc moiety is a commonly employed receptor by human influenza A viruses.

## INTRODUCTION

Respiratory viruses, which cause enormous disease burden,^1,2^ often employ glycans as receptor for cell attachment and/or entry. The relentless pressure of viral infections at the mucosal interface has driven the evolution of host and pathogen.^1^ It has shaped the glycomes of the host and even close related species can express substantially different collections of glycans.^3^ In turn, pathogens evolved glycan receptor specificities that determine host range and tissue tropism. Furthermore, immunogenic pressure can cause substitutions in receptor binding domains which in turn can influence receptor specificities. Despite the importance, glycan binding properties of respiratory viruses have been difficult to probe due to a lack of panels of biological relevant glycans for structure-binding studies.

Glycomic analyses of respiratory tissues of several animal species and humans have shown the abundant presence of bi-antennary *N*-glycans having poly-*N*-acetyl-lactosamine (poly-LacNAc) extensions that can be modified by terminal α2,3-or α2,6-sialosides.^6,7^ In human lung tissue, α2,3-linked sialosides are mainly presented on elongated LacNAc chains whereas α2,6-sialosides are more often found on structures that have a single LacNAc moiety.^8-10^ Lectin staining has demonstrated that upper airway tissues of humans are rich in α2,6-linked sialosides whereas duck enteric and chicken upper respiratory tract tissues display ample quantities of α2,3-linked sialosides.^3-5^ These differences in expression of sialosides represent a species barrier because human influenza A viruses (IAVs) recognize sialosides that are α2,6-linked to galactoside (Gal), whereas ancestorial avian IAVs prefer α2,3-linked isomers. A notion is emerging that α2,3-*vs*. α2,6-selectivity of avian and human IAVs is an oversimplification and further structural elements of glycans can determine receptor specificity.^7,11-18^ Comprehensive panels of biological relevant glycans are needed to adequately uncover such binding properties.^4,5^

We report here a stop-and-go strategy that made it possible to conveniently prepare a large panel of asymmetrical bi-antennary *N*-glycans having various numbers of LacNAc repeating units capped by α2,3- or α2,6-sialosides. The resulting collection of glycans, which resemble structures found in airway tissues, was printed as a microarray that was probed for binding of evolutionary distinct A(H3N2) viruses ranging from 1968 to 2023. Also, receptor binding properties were examined for recent human A(H1N1)pdm09 viruses and an avian virus that can infect but not transmit between humans. It was found that for human viruses, not only the sialoside type but also the length of the LacNAc chain and presentation at a specific antenna determines receptor binding properties. Initially, A(H3N2) viruses evolved to have restricted binding patterns and only recognize 2,6-sialosides presented on a tri-LacNAc moiety on the α1,3-antenna of *N*-glycans. However, recently this trend reversed and most strains that appeared after 2021 also exhibit some affinity for 2,6-sialosides presented on a di-LacNAc moiety. We hypothesize that this binding pattern relates to a balancing act between antigenic divergence and glycan binding properties to the more abundant of 2,6-sialylated di-LacNAc structures.^19^ Our data shows that this is not only a property of A(H3N2) but also for A(H1N1)pdm09 viruses.

The new glycan microarray provides an attractive platform to assess receptor binding properties of influenza viruses. It has provided new insight into evolution of viruses during epidemics, where there is an interplay between virus-receptor interactions on the one hand and virus-antibody interactions on the other hand. Mutations that affect one of these properties may also affect the other.

## RESULTS AND DISCUSSION

### Chemoenzymatic Synthesis of Asymmetrical Bi-Antennary Glycans

Recently, we introduced chemoenzymatic methodologies that was coined “Chemoenzymatic Glycosylations by a Stop-and-Go Strategy”.^20^ It employs symmetric bi-antennary glycan **1** as the key intermediate (Scheme. 1a) that could readily be prepared from a sialoglycopeptide isolated from egg yolk powder.^21^ Next, we took advantage of recombinant α-1,3-mannosyl-glycoprotein 4-?-N-acetyl-glucosaminyltransferase (MGAT4) and ?-1,6-mannosylglycoprotein 6-?-N-acetyl-glucosaminyltransferase (MGAT5), and unnatural UDP-2-deoxy-2-trifluoro-*N*-acetamido-glucose (UDP-GlcNTFA) to transform **1** into tetra-antennary glycan **3** in only four steps. A key strategic principle is that after the installation of a GlcNTFA moiety, the trifluoro-*N*-acetamido (TFA) moiety can be removed under mild basic conditions to give glucosamine (GlcNH_2_) that can be further transformed into 2-deoxy-2-azido-glucose (GlcN_3_). GlcNH_2_ and GlcN_3_ are inert (<u>stop</u>) to modifications by our panel of mammalian glycosyltransferases. Selective elaboration of the natural GlcNAc residues at the MGAT1 or 2 antenna was possible by exploiting inherent branch selectivities of ST6Gal1 and *E. coli* galactosidase for the α3-antenna. At the next stage of synthesis, the GlcNH_2_ and GlcN_3_ can be sequentially “unmasked” (go) to give natural GlcNAc termini for selective enzymatic elaboration into complex appendages to give compounds such as **4**. Intermediates such as **2b** were also employed for the preparation of tri-antennary glycans.

**Scheme 1.**
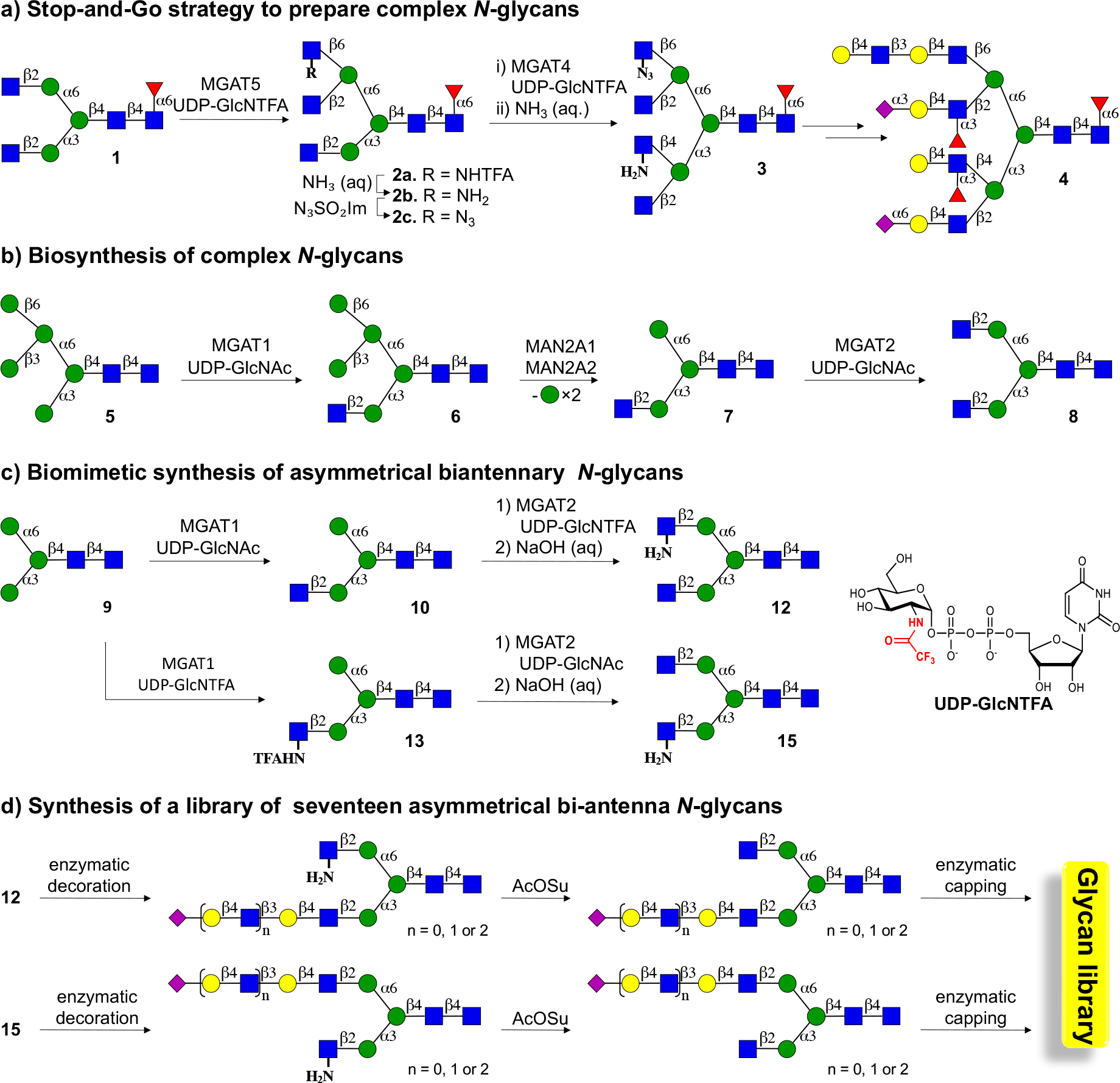
A stop-and-go strategy for the synthesis of asymmetrical bi-antennary *N*-glycans. a) Modification of symmetrical glycan **1**, which was derived from egg yolk powder, with MAGT4 and 5 using the unnatural donor UDP-GlcNTFA, followed by removal of the TFA moiety and chemical functionalization of the resulting amines gave **3**, which is an appropriate starting material to prepare asymmetrical glycans such as **4**. b) The biosynthesis of *N*-glycan involves trimming of high mannose *N*-glycans to **5** that can be modified by MGAT1 to provide **6** that after further mannoside trimming provides a substrate MGAT2. c) Compound **9**, which is assessable from **1**, was expected to be an appropriate substrate for MGAT1 and 2 and in combination with the natural and unnatural donor UDP-GlcNAc and UDP-GlcNTFA should give access to asymmetrical glycans **12** and **15**. d) Asymmetrical glycans **12** and **15** were expected to be appropriate starting materials for the preparation of asymmetrical bi-antennary *N*-glycans having extended LacNAc moieties typical of the respiratory glycome.

The stop-and-go strategy exploits the preference of ST6Gal1^22^ and *E. coli* galactosidase^23^ for the α3-antenna of a G2 structure to selectively elaborate the α3- and α6-antenna with specific appendages. The reaction conditions need careful controlling to achieve selectivity, and even when controlled, a regio-isomer (∼10%) is formed that needs to be removed by time-consuming HPLC purification.

In the biosynthesis of *N*-glycans, the α3- and α6-antenna are introduced by the branching enzymes α-1,3-mannosyl-glycoprotein 2-β-N-acetylglucosaminyltransferase (MGAT1) and 2-β-*N*-acetylglucosaminyltransferase (MGAT2), respectively. To achieve full control over the extension of the α3- and α6-antennae, we explored whether the stop-and-go strategy can be adapted to MGAT1 and MGAT2 to install a GlcNH_2_ moiety at one of these antennae to temporarily blocks extension by galactosyl transferases. MGAT1 and MGAT2 act early in biosynthetic pathway, and MGAT1 utilizes a Man5 structure as substrate (**5**) to produce hybrid Man5GlcNAc1 (**6**, Scheme 1b).^24^ Further processing of this compound by mannosidases creates Man3GlcNAc1 (**7**) that can be modified by MGAT2 to install a β1,2GlcNAc to the α1,6Man antenna to give Man3GlcNAc2 (**8**). It is, however, known that MGAT1 can also modify Man3^25^ thereby providing a substrate that potentially is adaptable to a stop-and-go strategy to prepare, in a controlled manner, asymmetrical bi-antennary *N*-glycans. It was found that treatment of Man3 (**9**) (Scheme 1c), which was readily obtained from a sialoglycopeptide isolated from egg yolk powder,^21,26^ can be modified by MGAT1 in the presence of UDP-GlcNAc or UDP-GlcNTFA to give glycans **10** and **13**. The latter two compounds could be further modified by MGAT2 in the presence of UDP-GlcNTFA or UDP-GlcNAc to give, after treatment with base, asymmetrical glycans **12** and **15**. The MGAT1 and 2 antenna of these compounds can then be selectively extended by various numbers of *N*-acetyl lactosamine moieties, which in turn can be capped by 2,3- and 2,6-linked sialosides to give structurally diverse asymmetrical *N*-glycans.

The Man3 glycosylated amino acid **9**, which at its anomeric center has a benzyloxycarbamante (Cbz) protected asparagine, could readily be prepared from a sialoglycopeptide (SGP) isolated from egg yolk powder (Scheme 2a).^21,26^ Thus, SGP was subsequently treated with a neuraminidase from *Clostridium perfringens* (→**16**) and galactosidase from *Aspergillus niger* to remove the sialosides and galactosides to give a glycopeptide **17** having a G0 structure. Next, the peptide moiety was hydrolyzed by pronase to leave a single Asn residue to afford **18**. The α-amine of Asn provides a convenient handle for immobilization on microarray slides having *N*-hydroxysuccinimide (NHS). It was, however, important to temporary protect this residue because in the subsequent synthesis, GlcNH_2_ residues are introduced at the MGAT1 or 2 antenna, and it was critical to differentiate the amines of GlcN and Asn. The latter could easily be accomplished by treatment of **18** with benzyloxycarbonyl chloride (CbzCl) in the presence of K_2_CO_3_ to give benzyloxycarbamate (Cbz) protected **19**. Finally, the terminal GlcNAc residues of **19** were removed by treatment with β-*N*-acetylglucosaminidase S from *Streptococcus pneumoniae* to give the targeted M3 glycosylated amino acid **9**, which was purified by size-exclusion and Hypercarb™ solid phase extraction (SPE) column chromatography and fully characterized by NMR and MS.

**Scheme 2.**
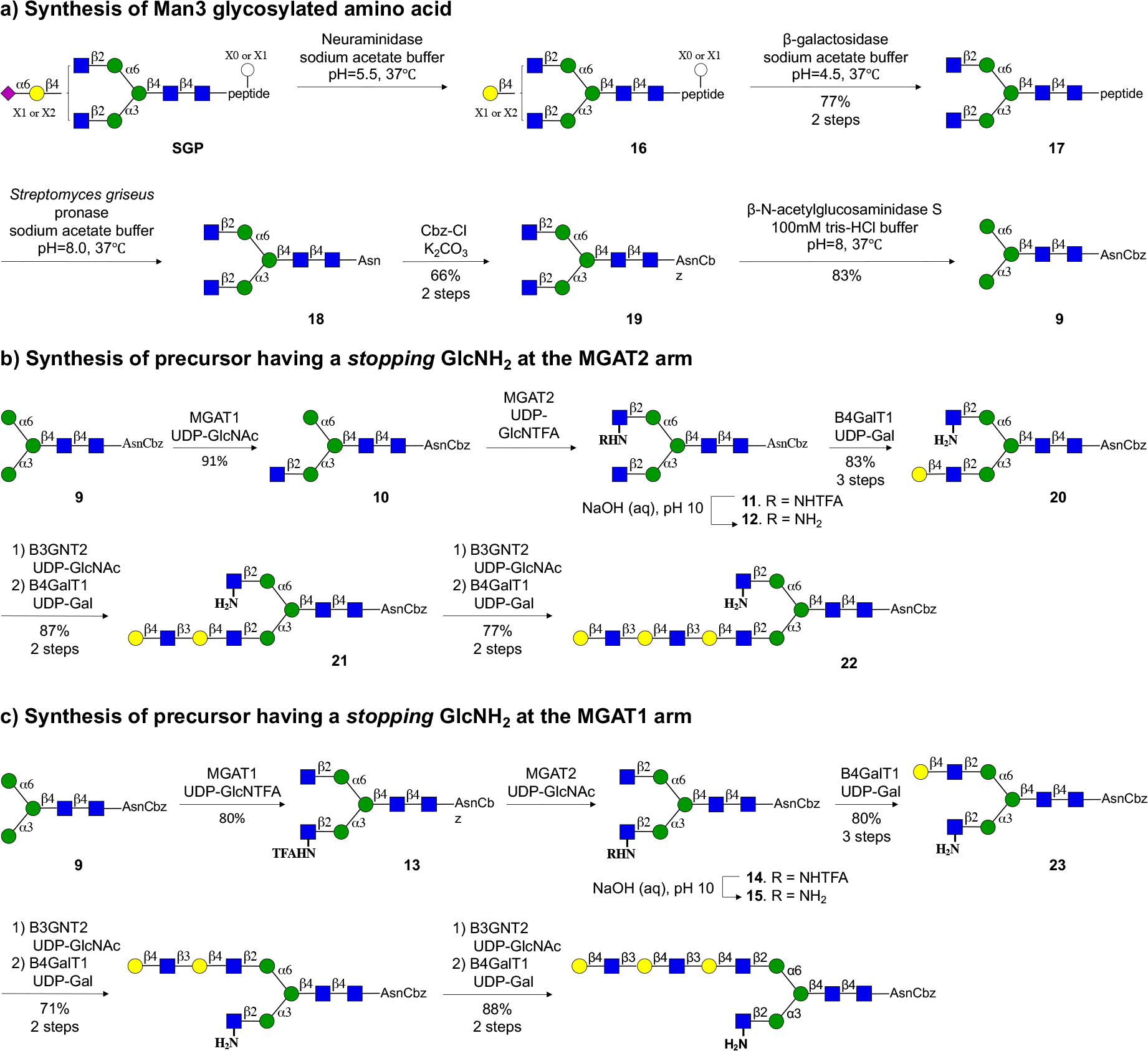
Chemoenzymatic synthesis of asymmetrical *N*-glycans having an extended LacNAc moiety at the MGAT1 or MGAT2 antenna. a) Preparation of tri-mannoside **9** from a sialoglycopeptide isolated from egg yok powder. b) Preparation of asymmetrical N-glycans having an extended LacNAc moiety at the MGAT1 antenna. c) Preparation of asymmetrical N-glycans having an extended LacNAc moiety at the MGAT1 antenna.

Next, attention was focused on the preparation of asymmetrical glycan **12**, which has a natural GlcNAc and an unnatural GlcNH_2_ moiety at the MGAT1 and MGAT2 antennae, respectively (Scheme 2b). Thus, treatment of Man3 derivative **9** with UDP-GlcNAc in the presence of recombinant MGAT1^27^ resulted in the formation of **10**, which was further treated with UDP-GlcNTFA in the presence recombinant MGAT2 to afford **11**, which was subjected aqueous sodium hydroxide (pH = 10) to provide target compound **12**. These result highlight that MGAT2 can utilize an unnatural donor such as UDP-GlcNTFA, which agrees with the finding that metabolic labeling of cells with diazirine modified GlcNAc results in low incorporation of the modified GlcN moiety at the MGAT2 position of *N*-glycans.^28^ The GlcNH_2_ residue of **12** is not a substrate for B4GalT1, thereby temporarily stopping all enzymatic modifications at this antenna. It allowed the MGAT1 antenna to be selectively extended by various LacNAc moieties by employing recombinant B4GalT1 and B3GnT2. Thus, treatment of **12** with B4GalT1 in the presence of UDP-GlcNAc introduced a Gal residue to give LacNAc containing derivative **20**. Additional LacNAc moieties could be introduced by one or two cycles of B3GnT2 and GalT1 catalyzed extensions to give compounds **21** and **22**, respectively. The intermediate and final compounds were purified by solid phase extraction using porous graphitized carbon which was followed by P-2 Bio-gel size exclusion column chromatography. All compounds were fully characterized by nuclear magnetic resonance (NMR) including Homonuclear Correlation Spectroscopy (COSY) and Heteronuclear Single Quantum Coherence (HSQC) experiments. Further confirmation of structural identity and purity came from analysis by liquid chromatography-mass spectrometry (LC-MS) using hydrophilic interaction liquid chromatography (Waters XBridge BEH, Amide column) (see Supporting Information for details).

We found that MGAT1 can also accept unnatural glycosyl donors and treatment of **9** with UDP-GlcNTFA in the presence of MGAT1 resulted in a quantitative conversion into **13** (Scheme 3). Despite the presence of the NTFA moiety, the latter compound is a proper acceptor for MGAT2 and could readily be converted into **14**, which after treatment with aqueous sodium hydroxide (pH = 10), gave target derivative **15**. In this case, the GlcNH_2_ at the MGAT1 stops temporary modification by glycosyl transferases, and thus it was possible to selectively extend the MGAT2 antenna by various LacNAc moieties to give access to compounds **23, 24**, and **25** having a mono-, di-or tri-LacNAc moiety.

**Scheme 3.**
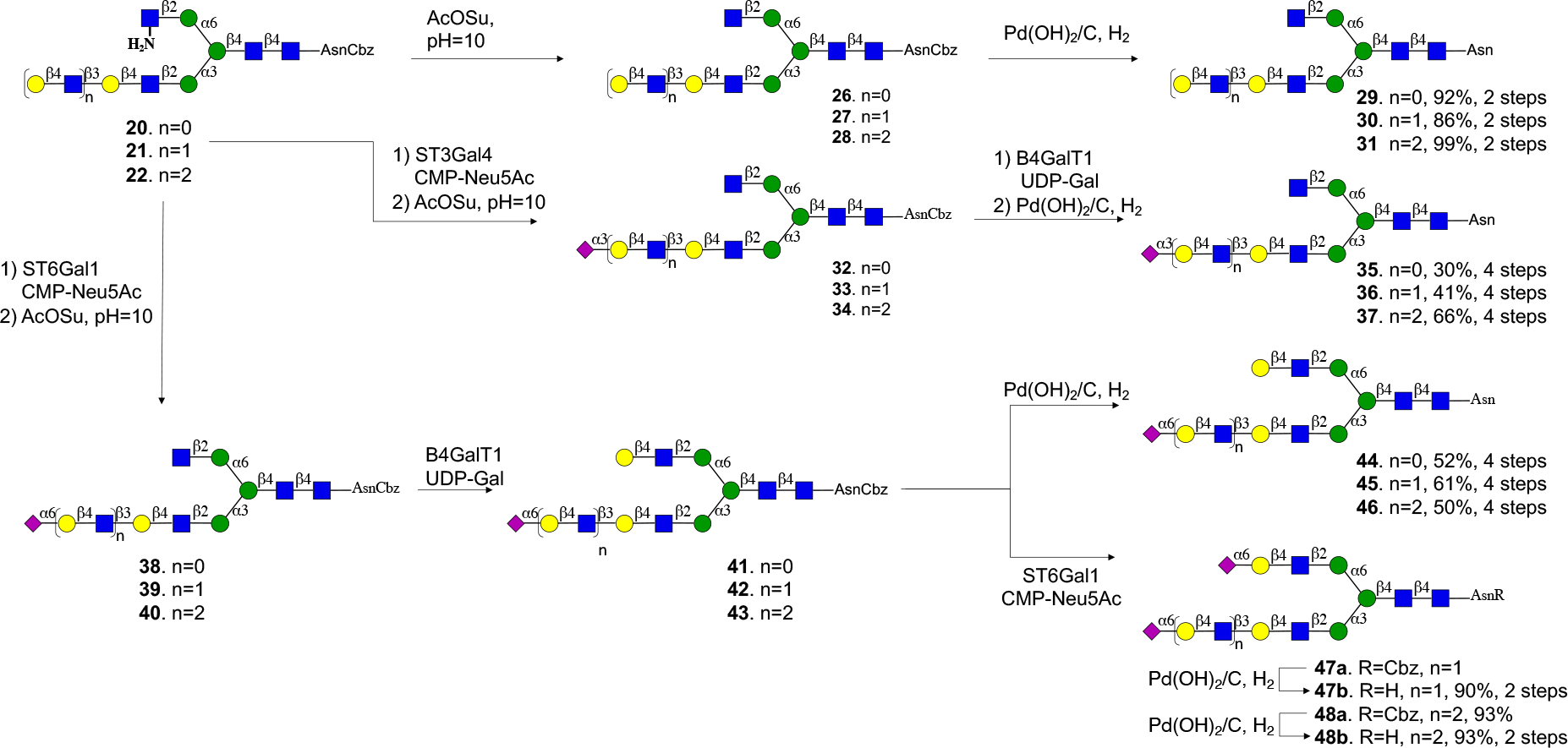
Selective modification of termini of the MGAT1 and MGAT2 antennae of glycans **20**-**22**.

Compounds **20**-**25** are ideally suited to prepare a panel of asymmetrical *N*-glycans having different patterns of sialosides at LacNAc moieties of different length. For example, the terminal galactoside of compounds of **20**-**22** could be modified by an α2,6-sialoside by ST6Gal1 in the presence of CMP-Neu5Ac to give, after acylation of the amine (→**38**-**40**) and galactosylation of the resulting terminal GlcNAc moiety with B4GalT1, compounds **41**-**43** (Scheme 3). The Cbz protecting group at the α-amine of the asparagine residue of the latter compounds was removed by hydrogenation over Pd/C to give sialosides **44**-**46**. The terminal Gal moiety of **41**-**43** could be further sialylated with ST6Gal1 (→**47a, 48a)** to give, after hydrogenation, di-sialosides **47b** and **48b**. Alternatively, the terminal galactoside of **20**-**22** could be modified by a 2,3-sialoside by treatment with ST3Gal4 to provide compounds **32**-**34**, which were subjected to hydrogenation to give target compounds **35**-**37**. Reference compounds **29**-**31** were prepared by acylation of the amine of **20**-**22** to give **26**-**28** followed by hydrogenation over Pd/C.

Compounds **23**-**25** were subjected to a similar sequence of enzymatic and chemical modifications to give a complementary panel of eleven asymmetrical glycans (**52**-**60, 67**-**71**, Scheme S1) that have an extended LacNAc moiety at the MGAT2 antenna. *N*-Glycans have been observed in which the MGAT1 is extended by various numbers of LacNAc moieties and lack a GlcNAc moiety at the MGAT2 antenna.^29^ Such compounds can also be prepared by the methodology described here by employing compound **10** as the starting material which can be galactosylated by B4GalT1 to install a LacNAc moiety. Various cycles of modification by B3GnT2 and B4GalT1 could install additional LacNAc moieties that could be capped by sialosides by using an appropriate sialyl transferases to give compounds **75, 77, 79, 82, 87**, and **89** (Scheme S2).

### Receptor Specificities of Influenza A Viruses

A(H3N2) and A(H1N1)pdm09 influenza virus subtypes that originate from the pandemics of 1968 and 2009, respectively are major causes of current seasonal influenza.^30^ Due to immunity caused by natural infection and vaccination, influenza A viruses continuously evolve to escape antibody-mediated neutralization. Protective antibody responses are mainly directed to the globular head of the hemagglutinin protein where binding occurs with sialic acid receptors of host cells.^31^ These mutational changes results in antigenic divergence from employed vaccine strains and circulating epidemic viruses, resulting in immune escape by the virus and poor protection.^32^ It can also alter receptor binding properties by orchestrated mutations of several amino acids in the globular head that allow immune evasion.^16^ The set of asymmetrical *N*-glycans developed here, offers unique opportunities to investigate receptor binding determinants of evolutionary distinct influenza A viruses. It can determine the importance of valency and sialoside-linkage type, the length of a LacNAc chain and placement of the binding epitope at the α1,3-(MGAT1) or α1,6-(MGAT2) antenna of bi-antennary *N*-glycans.

The glycans, which have an α-amine at the anomeric asparagine moiety, were printed on amine reactive NHS activated glass slides using a non-contact microarray printer (S3, Scienion *Inc*.). Quality control was performed using biotin-labeled plant lectins such as SNA, ECA and MAL. SNA binds 2,6-linked sialosides and as expected all compounds having such a moiety (**Fig. 1, M-W**) were recognized by approximate equal intensity (Fig. S40). ECA, which binds to terminal LacNAc moieties, bound to all compound having such a residue (Fig. 1, A-P, S and **T**) and as expected only the di-sialosides (Fig. 1, **Q, R, U, V**, and **W**) did not show responsiveness. As expected, the MAL proteins specifically bound to 2,3-linked sialosides (Fig. 1, **G-L**).

**Figure 1.**
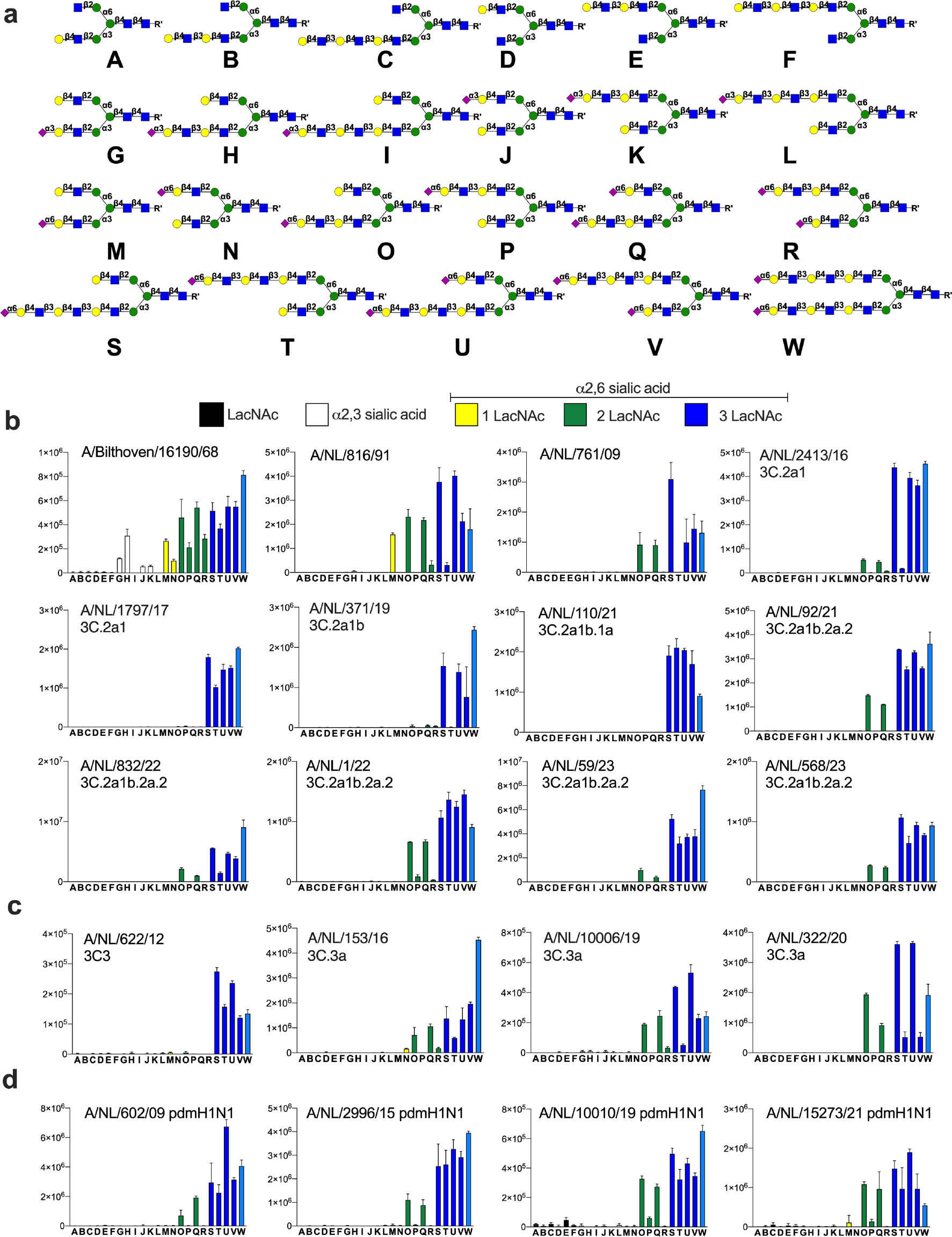
Probing receptor binding specificities of A(H3N2) and A(H1N1)pdm09. **a**) Collection of glycans printed on succinimide reactive microarray slides. Glycan binding data of **b**) early A(H3N2) and A(H3N2) 3C.2 viruses; **c**) A(H3N2) 3C.3 viruses; and **d**) A(H1N1)pdm09 viruses. Whole viruses were exposed to glycan microarray and binding was visualized anti-stalk antibodies. Bars represent the average relative fluorescence units (RFU) of four replicates ± SD.

Next, receptor binding properties were examined of evolutionary distinct A/H3N2 viruses. These viruses entered the human population during the 1968 pandemic and split into different antigenic clades such as those designated as 3C.2 and 3C.3 earlier this century, which further evolved into sub-clades. We analyzed receptor preferences of viruses isolated from the pandemic period to current epidemics, including the most recent 3C.2 and 3C.3 clades (Fig. 1). Also, receptor binding properties of several recent A(H1N1)pdm09 viruses and an avian A/H5N1 virus that can infect humans were examined. Viral isolates were applied to the array and detection of binding was accomplished using an appropriate anti-stalk antibody and a goat anti-human IgG antibody labeled with AlexaFluor-647.^18^ The array studies were performed in the presence of a neuramidase inhibitor to avoid interference of this protein.

An A(H3N2) virus that was isolated during the pandemic (A/Bilthoven/68) showed promiscuous binding behavior and bound to 2,3-as well as 2,6-linked sialosides (Fig. 1a). Interestingly, very little binding was observed when the 2,3-linked sialoside is presented on a tri-LacNAc moiety (**I** and **L**), whereas proper recognition is observed when such a sialoside is placed on a di-LacNAc structure (**G** and **H**). An A(H3N2) virus isolated in 1991 (A/NL/861/91), lost all binding to 2,3-linked sialosides and only recognized 2,6-linked sialosides. It displays a strong preference for the presentation of this epitope at the α1,3-antenna (MGAT1 extension). For example, compound **M**, in which the 2,6-sialoside is presented at a single LacNAc moiety at the α1,3-antenna, was bound strongly by the virus whereas this was not the case for isomeric glycan **N** that has the epitope at the α1,6-antenna (MGAT2 extension). *N*-glycans that have extended 2,6-sialyl-LacNAc moieties were also recognized and in these cases presentation of the epitope at the α1,3-antenna was also strongly preferred (**O** *vs*. **P, Q** *vs*. **R** and **S** *vs*. **T**). A virus isolated in 2009 (A/NL/761/NL09) became even more selective, recognizing fewer glycans and did not bind to structures having an α2,6-sialosides presented at a mono-LacNAc moiety (**M** and **N**). These viruses required a 2,6-sialoside to be presented at a di-or longer LacNAc moiety (**O** and **Q**) at the α1,3-antenna. Isomeric compounds in which these epitopes are presented at the α1,6-antennae (MGAT2 extension) (**P** and **T**) were bound poorly. A/NL/2413/16, which belongs to the 3C.2a clade, showed similar structure-binding properties, however, in this case the tri-LacNAc containing compounds bound much more robustly (**O** *vs*. **S** and **Q** *vs*. **U**). The further evolved A/NL/1797/17, A/NL/371/19 and A/NL/110/21 viruses (3C.2a clade) showed the most restrictive binding pattern and had an obligatory requirement for presentation of the α2,6-sialoside at a tri-LacNAc structure. A/NL/371/19 exhibited an antenna preference and for mono-sialosides, the sialyl-tri-LacNAc epitope needs to be displayed at the α1,3-arm (**S** *vs*. **T**) whereas this is not the case for the bis-sialosides (**Q, U** and **V**). A/NL/1797/17 and A/NL/110/21 did not exhibit such an antenna preference and compounds **S** and **T** were strongly bound. Subsequent 3C.2a1b.2a.2 viruses (A/NL/92/21, A/NL/832/22, A/NL/1/22, A/NL/59/23, A/NL/568/23) exhibited broader receptor binding properties compared to early 3C.2a viruses and also bound to **O** and **Q** that have the 2,6-sialoside at a di-LacNAc moiety at the α1,3-antenna only.

An early 3C3 viruses (A/NL/622/12) required a tri-LacNAc moiety modified by a 2,6-sialoside (**S**-**V**) for binding whereas later viruses also recognized compounds having the sialoside presented at a di-LacNAc moiety (**O** and **Q**) and here asymmetry is also noted (**S** *vs*. **T** and **U** *vs*. **V**. (Fig. 1c). Interestingly, several A(H1N1)pdm09 subtype viruses that re-entered the human population during the 2009 pandemic, showed a preference for 2,6-linked sialosides at di-LacNAc moieties presented at the α1,3-antenna (Fig. 1d) (**O** *vs*. **P**). In cases of tri-LacNAc derivatives, this antennae preference was not observed, and compounds **S**-**V** bound robustly.

Previously, it was proposed that A(H3N2) viruses bind host *N*-glycans through a bidentate binding mode in which *N*-glycan having two sialic acid moieties bind to a protomer of the same HA trimer thereby increasing the binding avidity.^15^ Modeling studies indicated that the sialosides need to be displayed at a tri-LacNAc moiety to be able to bind to two HA protomers. We observed that compound **W**, which has 2,6-sialylated tri-LacNAc moieties at the α1,3- and α1,6-antenna, showed similar responsiveness compared to compound **S** that only has one such moiety at the α1,3-antennae. Thus, it is unlikely that these viruses establish a bidentate binding mode, and it is probable that the extended LacNAc moiety makes interactions with the HA protein to compensate for reduced contacts with sialic acid caused by substitutions in the receptor-binding site arisen from antigenic pressure.^31^

Computational studies have indicated that A(H3N2) 3C2a viruses that have an obligatory requirement for a 2,6-sialylated tri-LacNAc moiety. This was acquired with mutational changes distal to the receptor binding domain that reoriented the Y159 side chain, resulting in an extended receptor binding site that can accommodate the tri-LacNAc moiety.^18^ Sequence alignment showed that contemporary 3C.2a1b.2a.2 viruses that can bind sialylated di-LacNAc containing structures have aspartic acid at position 159 (Table S3). Such a mutation was also observed in 3C3 viruses that adapted to binding to 2,6-sialylated di-LacNAc moiety. Moreover, the adjacent T160I abrogates an *N*-glycosylation site that may affect receptor binding properties.^33^ Importantly, the 190-helix, which can make interactions with the LacNAc moiety, has been under heavy antigenic pressure as several positions have changed including G186D, D190N, F193S and Y196F. Positions 186,^34^ 190,^35^ and 193^36^ are hallmark mutations for different receptor binding properties in A/H3 and other subtypes, however, their individual contributions to receptor binding specificity and antigenicity remains to be determined. Thus, it appears that recent A(H3N2) viruses have made compensatory mutations for loss of interactions with Y159. Furthermore, for most of the recent subtypes it has resulted in some ability to bind di-LacNAc containing sialosides.

Receptor binding properties were also investigated of an avian A/H5N1 virus that can infect but not transmit between humans (A/Indonesia/05/05). In this case, all 2,3-linked sialosides were bound without differentiation of presentation of the epitope on the α1,3- or α1,6-antenna and for example isomeric compounds **G** and **J** gave similar responsiveness (Fig. S40).

### Hemagglutination properties of contemporary A(H3N2) viruses

For antigenic surveillance and vaccine strain selection of influenza A viruses, the hemagglutination inhibition (HI) assay is used, in which the ability of serum antibodies to prevent virus-mediated agglutination of erythrocytes is measured.^37^ HI titers provide a correlate of protection and make it possible to select virus strains that are antigenically representative of circulating viruses for vaccine development.^38^ 3C2a viruses, which have an obligatory requirement for a 2,6-sialylated tri-LacNAc moiety, lost the ability to agglutinate commonly employed chicken and turkey erythrocytes.^16^ Glycomic analysis of the latter erythrocytes have shown these erythrocytes do not express *N*-glycans having such residues thereby providing a rationale for a lack .^18^ The microarray studies presented here demonstrate that recent A(H3N2) strains of C2 and C3 clades regained an ability to bind di-LacNAc containing sialosides. Turkey erythrocytes express low levels of these glycans and thus we were compelled to investigate whether the recent C2 and C3 strains can agglutinate turkey erythrocytes. Previously, we introduced a glycan remodeling approach for turkey erythrocytes to elongate the LacNAc moieties at which sialosides are presented.^18^ It is based on treatment with a sialidase to remove sialosides to reveal terminal galactosides that can be extended by additional LacNAc moieties by treatment with B4GalT1 and B3GnT2 followed by sialylation of terminal galactosides by the sialyltransferase ST6Gal1. Agglutination properties of these glycan remodeled erythrocytes (2,6-Sia-poly-LN cells) were also examined. To investigate whether an increase in abundance of α2,6-sialosides on mono- and di-LacNAc containing structures can improve agglutination, control erythrocytes were prepared by treatment with neuraminidase followed by resialylation with ST6Gal1(2,6-Sia cells). In this way, 2,3-sialosides and terminal galactosides are converted in 2,6-sialosides resulting in a higher receptor density of human viruses.

Table 1 summarizes hemagglutination titers for the A(H3N2) viruses for which the receptor binding properties were investigated using the new glycan microarray. Regular as well as glycan remodeled turkey erythrocytes (2,6-Sia- and 2,6-Sia-poly-LN cells) were employed. The early viruses (1968 and 1991), which require for binding a single LacNAc moiety, could readily agglutinate the three erythrocyte types. A/NL/761/09 did not agglutinate regular erythrocytes while the 2,6-Sia and 2,6-Sia-poly-LN cells were robustly agglutinated with similar titers. The glycan array data for this virus showed it can bind di- and tri-LacNAc containing sialoglycans with similar affinities (Fig. 1b), and thus for this virus increasing the abundance of α2,6-linked sialosides having two consecutive LacNAc repeating units is sufficient to induce agglutination. The further evolved A/NL/2413/16 did bind 2,6-sialosides presented on a di-LacNAc moiety but with substantial lower responsiveness compared the tri-LacNAc containing counterparts. The hemagglutination properties agree with this binding pattern and no agglutination was observed for regular erythrocytes while the 2,6-Sia and 2,6-Sia-poly-LN cells exhibited agglutination with the latter giving a substantial higher titer. The 3C.2 viruses that appeared in 2017 and 2019 have a strict requirement for a 2,6-linked sialoside to be displayed at a tri-LacNAc moiety, and as expected they did not agglutinate the regular and 2,6-Sia erythrocytes and required the *N*-glycans to be extended with additional LacNAc moieties (2,6-Sia-poly-LN cells) to achieve agglutination.

**Table 1.**
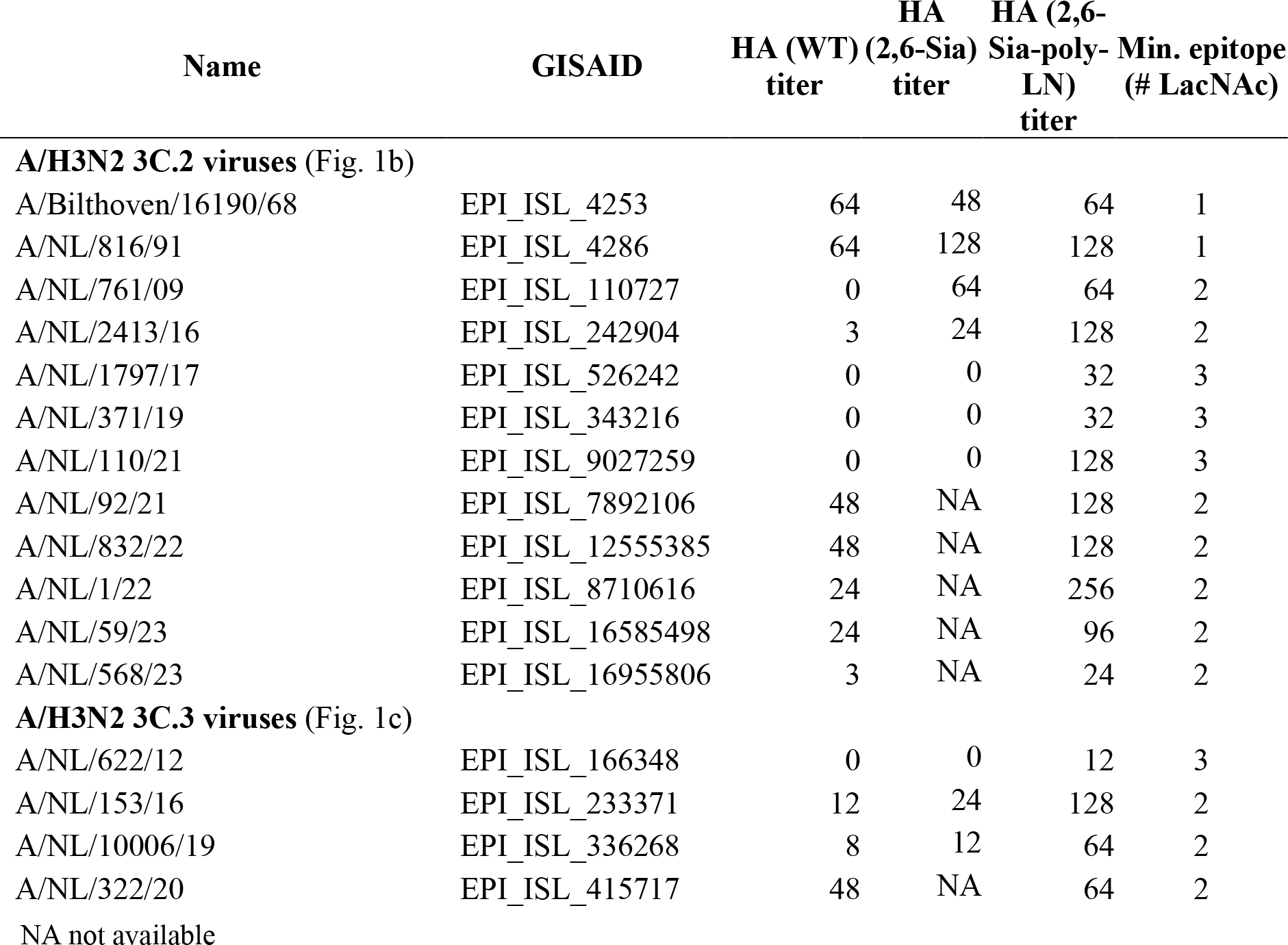
Hemagglutination titers of A(H3N2) viruses for wild type (WT) and glyco-remodeled (Mod) turkey erythrocytes and minimal requirement of the length of the LacNAc moiety for sialoglycan binding.

Most of the 3C.2 viruses that appeared after 2021 regained the ability to agglutinate regular erythrocytes and importantly these viruses reverted to bind to di-LacNAc-containing structures (Fig. 1b,c). Unlike A/NL/2413/16, these viruses do not require an increase in receptor density to achieve agglutination. This indicates that in addition to the length of the LacNAc chain, other factors such as avidity of binding and the balance between HA/NA activity may contribute the ability to agglutinate regular erythrocytes. These recent viruses did give higher agglutination titers for the 2,6-Sia-poly-LN erythrocytes, which agrees with the glycan array data that showed substantially higher responsiveness for tri-LacNAc containing structures.

A/NL/110/2021 is a recent 3.C2 viruses that could not agglutinate regular erythrocytes. Interestingly, it has an obligatory requirement for a sialylated tri-LacNAc containing *N*-glycans (Fig. 1b) thereby providing a rationale for this behavior. A/Netherlands/568/23 has some binding capacity to bind di-LacNAc containing sialosides, however, it agglutinated regular erythrocytes with a very low titer, and its behavior and is more akin to A/NL/2413/16. Hemagglutination titers of a broader range of A/H3N2 viruses isolated between 2021 and 2023 were determined and almost all gave good titers for regular erythrocytes (Table S2).

The agglutination properties of the 2C.2 viruses agreed with the glycan microarray data and a virus that only bound tri-LacNAc containing compounds only agglutinated the 2,6-Sia-poly-LN erythrocytes.

## CONCLUSION

Glycan-binding proteins often recognize relatively small oligosaccharide motifs found at termini of complex glycans and glycoconjugates.^1,39,40^ There are, however, indications that the topology of a complex glycan can modulate recognition of terminal glycan epitopes.^41-43^ This can be due to an extended binding site, unfavorable interactions by a glycan moiety at which a minimal epitope is appended, conformational changes of larger glycan moieties and multivalency. Panels of well-defined glycans are needed to probe the importance of glycan topology on glycan binding properties. Here, we describe a synthetic methodology to prepare asymmetrical bi-antennary glycans that have extended LacNAc moieties at the α1,3- (MGAT1) or α1,6- (MGAT2) antenna. It exploits the finding that MGAT1 and MGAT2 can utilize the unnatural sugar donor UDP-GlcNTFA. The TFA moiety of the resulting glycans can be selectively hydrolyzed to give compounds having a GlcNH_2_ moiety at one of the antennae to temporarily block extension by glycosyl transferases. It made it possible to conveniently prepare a library of asymmetrical *N*-glycans that resemble structures found in the respiratory tract.^6,7^ The compounds were printed as a glycan microarray that was used to examine receptor binding properties of evolutionary distinct A(H3N2) influenza viruses ranging from a pandemic strain of 1968 to recent isolates. Earlier this century, A(H3N2) viruses split into two different antigenic clades designated as 3C.2 and 3C.3, which further evolved in sub-clades. Binding studies with the collection of compounds described here, revealed that not only the length of the LacNAc chain but also presentation on a specific antenna is critical for receptor binding. For many of the examined viruses, a preference for presentation of the epitope on the α1,3-antenna was observed. The data also reveals that a single sialylated LacNAc moiety is sufficient for binding and thus it is unlikely that these viruses engage in a previously proposed bident binding mode.^44^

3C.2 viruses isolated between 2017 and 2019 displayed the most restricted binding pattern and only recognized glycans having a 2,6-linked sialoside presented at a tri-LacNAc structure on the α1,3-antenna (Fig. 1b). This specificity was also observed for a 3C.2a1b.1a virus isolated in 2021. However, most virus isolates at that time until now belong to the 2a.2 subclade and most of these viruses regained some binding capacity to sialylated di-LacNAc structures.^45^ These di-LacNAc-containing structures need to be displayed on the mannose of the α3-antenna for recognition (**O** *vs*. **P** and **Q** *vs*. **R**). This observation was further strengthened by analyzing A(H3N2) viruses of the 3C.3 antigenic clade that already in 2016 regained binding of these di-LacNAc containing structures (Fig. 1c).

ST6Gal1, which is the only sialyl transferase that installs terminal 2,6-sialosides at terminal LacNAc moieties, preferentially modifies the LacNAc moiety displayed at the α1,3-antenna,^44^ and thus early H3 viruses such as NL91 likely evolved to bind such glycans (Fig. 1b). The requirement for presentation of an extended sialylated LacNAc moiety on the MGAT1 antenna may be due to a similar antennae preference of ST6Gal1. It is also possible that due to steric reasons the α1,3-antenna is less assessable for binding. Future studies The molecular mechanism for α1,3-antenna preference needs further examination.

A(H3N2) influenza viruses that appeared after the turn of the centenary lost the ability to agglutinate commonly employed fowl erythrocytes.^46^ This greatly complicated antigenic surveillance and vaccine strain selection, which relies on the hemagglutination inhibition assay, in which the ability of serum antibodies to block receptor binding by the influenza virus HA protein is quantified. We found that all viruses that lost the ability to agglutinate regular and 2,6-resiaylated erythrocytes require as minimal epitope a 2,6-sialoside presented at a tri-LacNAc moiety. Surprisingly, most of the A(H3N2) that appeared after 2021 regained an ability to agglutinate common erythrocytes, and these viruses had reverted to the use of a sialoside presented a di-LacNAc containing structures as minimal epitope. Earlier A(H3N2) viruses, such as A/NL/761/09 and A/NL/2413/16, employ similar structures as minimal epitope, however, these viruses cannot agglutinate regular erythrocytes and require a higher density of receptors as found on the 2,6-Sia cells. This indicates that properties such as the avidity of binding and HA/NA balance contributes to agglutination properties of these viruses.

We also analyzed receptor binding properties of A(H1N1)pdm09 viruses that were reintroduced during the 2009 pandemic (Fig. 1d). All viruses tested had similar receptor binding properties as contemporary A(H3N2) viruses. Thus, a 2,6-sialoside at a di-LacNAc moiety presented at the α1,3-antenna appears to be a commonly employed receptor by human influenza A viruses. It is likely that the di-LacNAc moiety allows for additional interactions with HA for sufficient high affinity of binding to compensate for reduced binding of sialic acid due to mutational changes caused by antigenic pressure.^19^

## ASSOCIATED CONTENT

### Supporting Information

The Supporting Information is available free of charge at https://pubs.acs.org.

Synthetic protocols, compound characterization, experimental procedure for microarray screening, NMR spectra (PDF).

### Notes

The authors declare no competing financial interest.

## ACKNOWLEDGMENTS

Research reported in this publication was supported by the National Institute of Allergy And Infectious Diseases of the National Institutes of Health under Award Number R01 AI165692 (to G.J.B) and R01AI165818 (to R.A.M.F). The content is solely the responsibility of the authors and does not necessarily represent the official views of the National Institutes of Health. R.P.dV is supported by the European Commission (802780). S. van Leeuwen, K. de Haan, C. Russell, and Z. Felix-Garza (Amsterdam UMC, The Netherlands) have contributed to the production of A(H1N1)pdm09 viruses. M. Pronk and P. Lexmond (Erasmus MC, The Netherlands) assisted with hemagglutination studies. R. Liang (Utrecht University, The Netherlands) prepared the glycan remodeled erythrocytes.

## TOC graphic

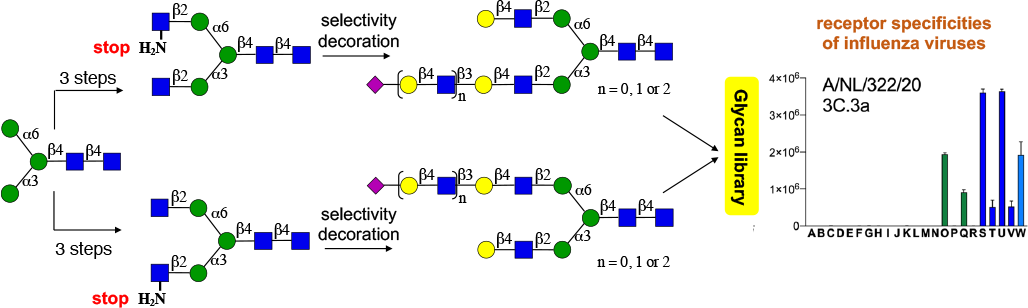

## Notes

### Competing Interest Statement

The authors have declared no competing interest.

