## Supporting information for "Asymmetrical Bi-antennary Glycans Prepared by a Stop-and-Go Strategy Reveal Receptor Binding Evolution of Human Influenza A Viruses"

Table of Contents

| Table of Contents | Page |
|---|---|
| 1. Preparation of compound **9** | S2 |
| 1.1 Materials and methods | S2 |
| 1.2 Sialylglycopeptide (SGP) isolation | S2 |
| 1.3 Trimming and modification of SGP to prepare compound **9** | S2 |
| 2. Enzyme mediated synthesis | S4 |
| 2.1 General methods | S4 |
| 2.2 General protocols for enzyme mediated reactions and glycan purification | S4 |
| 2.3 Schemes S1 and S2 | S7 |
| 2.4 NMR nomenclature | S9 |
| 2.5 Enzymatic reactions | S10 |
| 3. Microarray | S91 |
| 3.1 Materials | S91 |
| 3.2 Glycan printing surfaces | S91 |
| 3.3 Glycan microarray | S91 |
| Fig. S40. Validation of the microarray | S92 |
| 4. Hemaglutination and sequence alignment | S93 |
| 4.1 Erythrocyes preparation | S93 |
| 4.2 Enzymatic remodeling | S93 |
| 4.3 Hemagglutination assay | S93 |
| Table S2. Sequence alignment of H3 proteins | S94 |
| Table S3. Hemagglutination titers of A(H3N2) viruses | S95 |
| 5. References | S96 |
| 6. NMR spectra | S97 |

### 1. Preparation of compound 9

##### 1.1 Materials and methods

Chemicals and solvents were purchased from Sigma-Aldrich unless noted otherwise. Pronase from *Streptomyces griseus* (#P5147) was also purchased from Sigma-Aldrich. Egg yolk powder was purchased from Natural Foods Inc, Toledo, OH (#40504) and stored at 4 °C before used. Active carbon, NORIT™ SA 2, decolorizing grade, catalog 40403 and Celite® 545 was purchased from Acros. *Clostridium perfringens* Neuraminidase (#P0720), *Flavobacterium meningosepticum* PNGase F (#P0704) and *Streptococcus pneumoniae* β-N-Acetylglucosaminidase S (#P0744S) were obtained from New England Biolabs. *Aspergillus niger* Galactosidase (#E-BGLAN) was purchased from Megazyme.

##### 1.2 Sialyl glycopeptide (SGP) preparation

SGP was extracted, isolated and purified according to our previously reported method.[1,2] Briefly, commercialized egg yolk powder (2.27 Kg) was suspended twice in 95% ethanol (4 L) and mechanically stirred for 2 h at room temperature to remove lipids and other organic soluble components. The filtrate was discarded and the insoluble powder was suspended twice in aqueous ethanol (40% *v/v* ethanol, 3 L). The insoluble material was discarded and the filtrate was concentrated under reduced pressure at 40 °C. The collected liquid was concentrated *in vacuo* and purified using an active carbon/celite column (200 g of active carbon and 600 g celite). Impurities were removed by flushing the column with 3 L of water (trifluoroacetic acid (TFA) 0.1% *v/v*), 3 L of 5% acetonitrile in water (TFA, 0.1% *v/v*), and 3 L 10% acetonitrile in water (TFA, 0.1% *v/v*). The desired SGP was released from the column using a solution of 25% acetonitrile in water (TFA, 0.1% *v/v*), and fractions containing the product were pooled and concentraded *in vacuo*. The resulting crude SGP was subjected to size-exclusion chromatography (Bio-Rad® P-2, fine particle size 45 – 90 µm, column dimensions 5.0 cm x 80 cm, 250 mL fractions) eluting with 0.1 M ammonium bicarbonate to yield pure SGP (**5**) as a fluffy, white powder after be evaporated and lyophilized (1.6 g, or 0.7 mg SGP/g egg yolk powder). Additional P-2 bio-gel column purification was applied when necessary to obtain highly pure SGP.

##### 1.3 Trimming and modification of SGP to prepare compound 9

SGP (500 mg) was dissolved in 50 mM sodium acetate buffer (5 mL, pH 5.5). 5 mM $CaCl_2$ and 40 µL *Clostridium perfringens* neuraminidase were added to reaction mixture solution and then the reaction was incubated overnight at 37 °C with shaking. When ESI-MS indicated all the sialic acid residues had been removed, the pH of the reaction mixture was adjusted to 4.5 with acetic acid. After that, 5 mg BSA and 200 µL β-galactosidase from *Aspergillus niger* were successively added. The resulting reaction mixture was incubated overnight with shaking at 37 °C. Additional 100 µL β-galactosidase will be needed if a spot of substrates were observed via ESI-MS. There are no any substrates in reaction solution and full galactose removal was monitored by ESI-MS, after which an equal volume of cooled alcohol was added to precipitate protein residuals. The result solution was concentrated *in vacuo* and loaded into P-2 Bio-gel size-exclusion chromatography eluting with 0.1 M ammonium bicarbonate buffer. The fractions containing product were collected and lyophilized. Finally, compound **17** was obtained as white amorphous powder (265 mg, 77% over 2 steps).

Compound **17** (265 mg, 0.14 mmol) was dissolved in 100 mM tris buffer with 5 mM $CaCl_2$. Pronase (130 mg) was added keeping a 1 mg/mL final concentration. The reaction was incubated overnight at 37 °C with shaking. The reaction was monitored by ESI-MS and an additional 50 mg of pronase was added an incubation continued for 8 h if partial product formation was observed. Finally, an equal volume of cooled (0 ºC) alcohol was added to precipitate residual protein. The resulting solution was concentrated *in vacuo* and loaded into P-2 Bio-gel size-exclusion chromatography which was eluting with 0.1 M ammonium bicarbonate buffer. The fractions containing product were collected, lyophilized and dissolved in water (5 mL). Next, $K_2CO_3$ (1.5 eq), and Cbz-Cl (1.5 eq) were added successively. Subsequently, the mixed solution was incubated at 37 °C with shaking until ESI-MS indicated complete installation of the Cbz group. The reaction mixture was diluted using water (20 mL) and extracted twice with ethyl acetate (30 mL). The aqueous phase was concentrated *in vacuo* and loaded into P-2 Bio-gel size-exclusion chromatography, which was eluted with 0.1 M ammonium bicarbonate buffer. The fractions containing product were collected and lyophilized giving 140 mg of **19** as a white powder (66% over 2 steps).

Compound **19** (140 mg, 0.09 mmol) was dissolved in 5 mL of 100 mM Tris-HCl buffer (pH = 8.0). 5 mM $CaCl_2$ and 50 U β-Acetylglucosaminidase S were added to reaction solution. The resulting reaction mixture was incubated overnight with shaking at 37 °C. The progress of the reaction was monitored by ESI-MS until full conversion. As a result, an equal volume of cooled alcohol was added to precipitate protein residuals. The result solution was concentrated *in vacuo* and loaded into P-2 Bio-gel size-exclusion chromatography eluting with 0.1 M ammonium bicarbonate buffer. The fractions containing product were collected and lyophilized. Compound **9** was obtained as white powder (86 mg, 83%).

#### 2. Enzymatic synthesis

##### 2.1 General methods

All enzymatic reactions were performed in aqueous buffers at an appropriate pH for each enzyme. Recombinant human glycosyl Transferases including α-1,3-mannosyl-glycoprotein 2-β-*N*-acetylglucosaminyltransferase (MGAT1), α-1,6-mannosyl-glycoprotein 2-β-*N*-acetylglucosaminyltransferase (MGAT2), β-1,3-*N*-acetylglucosaminyltransferase 2 (B3GNT2), β-1,4-galactosyltransferase 1 (B4GalT 1), β-galactoside-α-2,6-sialyltransferase 1 (ST6Gal1) and β-galactoside-α-2, 3-sialyltransferase 4 (ST3Gal4) were provided by Dr. K. W. Moremen (Complex Carbohydrate Research Center, Athens, GA, USA) which were expressed according to the published protocols.[3-5] Alkaline phosphatase from calf intestine (CIAP) and Bovine serum albumin (BSA) were purchased from Sigma-Aldrich. Uridine 5'-diphospho-N-acetylglucosamine (UDP-GlcNAc) was purchased from Sigma-Aldrich. Uridine 5'-diphosphogalactose diphosphate galactose (UDP-Gal) and cytidine 5'monophospho-N-acetylneuraminc acid (CMP-Neu5Ac) were both purchased from Roche. Uridine 5'-diphospho-N- trifluoroglucosamine (UDP-GlcNTFA) was synthesized utilizing a one-pot three-enzyme combination as previously reported.[6] All enzymatic reactions were monitored using a Shimadzu 20AD UFLC LCMS-IT-TOF a Shimadzu 20AD UFLC LCMS-IT-TOF. To facilitate isolation and purification of products, all enzymatic reactions were driven to full completion by adding sufficient glycosyl transferase until all starting material was consumed. Glycan products were purified by a two-step purification approach (in details on **2.2 j**). All nuclear magnetic resonance (NMR) spectra were acquired on a 600 MHz Varian Inova operating at 25 °C. Samples were dissolved in 99.96% $D_2O$ and chemical shifts were referenced to residual HDO signal at 4.79 ppm. Data were collected using standard pulse programs.

##### 2.2 General protocols for enzymatic reactions and glycan purification

**a) General procedure for the installation of β1,2-GlcNAc or β1,2-GlcNTFA using MGAT1**

Compound **9** (1.0 eq) and UDP-GlcNAc or UDP-GlcNTFA (1.5 eq) were dissolved in a HEPES buffer solution (100 mM, pH 7.0) containing $MnCl_2$ (2 mM) and BSA (1% total volume, stock solution = 10 mg/mL) keeping a final acceptor concentration of 10 mM. Calf intestine alkaline phosphatase (CIAP, 1% total volume, stock solution = 1kUmL) and recombinant MGAT1 (50 µg/µmol acceptor) were successively added, and the reaction mixture was incubated overnight at 37 °C with shaking. The reaction was monitored by LC-ESI-IT-TOF MS, and additional MGAT1 and donor were added to consume material if the reaction is incomplete. The product was purified by Hypercarb™ SPE followed P-2 Bio-gel size-exclusion column chromatography (see **2.2 J** for details).

**b) General procedure for the installation of β1,2-GlcNAc or β1,2-GlcNTFA using MGAT2**

Acceptor (1.0 eq) and UDP-GlcNAc or UDP-GlcNTFA (1.5 eq) were dissolved in a HEPES buffer solution (100 mM, pH 7.0) containing $MnCl_2$ (2 mM), and BSA (1% total volume, stock solution = 10 mg/mL) to provide a final acceptor concentration of 10 mM. Calf intestine alkaline phosphatase (CIAP, 1% total volume, stock solution = 1kU/mL) and MGAT2 (50 µg/µmol acceptor) were successively added, and the mixture solution was incubated overnight at 37 °C with shaking. The reaction was monitored by LC-ESI-IT-TOF MS, and additional MGAT2 and donor were added if the reaction was incomplete. The product was purified by Hypercarb™ SPE followed P-2 Bio-gel size-exclusion column chromatography (see **2.2 J** for details).

**c) General procedure for the installation of β1,4 Gal using B4GALT1**

Acceptor (1.0 eq) and UDP-Gal (1.5 eq) were dissolved in a Tris-HCl buffer solution (100 mM, pH 7.5) containing $MnCl_2$ (2 mM), and BSA (1% total volume, stock solution = 10 mg/mL) to provide a final acceptor concentration of 5 mM. Calf intestine alkaline phosphatase (CIAP, 1% total volume, stock solution = 1kU/mL) and B4GalT 1 (1% *wt/wt* relative to acceptor) were successively added, and the mixture solution was incubated overnight at 37 °C with shaking. The reaction was monitored by LC-ESI-IT-TOF MS, and extra B4GalT 1 and donor were added to consume material if the reaction is incomplete. The product was purified by Hypercarb™ SPE followed P-2 Bio-gel size-exclusion column chromatography (see **2.2 J** for details).

**d) General procedure for the installation of β1,3 GlcNAc using B3GNT2**

Acceptor (1.0 eq) and UDP-GlcNAc (1.5 eq) were dissolved at a final acceptor concentration of 10 mM in a HEPES buffer solution (100 mM, pH 7.3) containing $MnCl_2$ (2 mM), KCl (25 mM), $MgCl_2$ (2 mM), DTT (1 mM) and BSA (1% total volume, stock solution = 10 mg/mL). Calf intestine alkaline phosphatase (CIAP, 1% total volume, stock solution = 1kU/mL) and B3GNT2 (1% *wt/wt* relative to acceptor) were successively added, and the reaction mixture was incubated overnight at 37 °C with shaking. The reaction was monitored by LC-ESI-IT-TOF MS, and additional B3GNT2 and donor were starting material was remaining. The product was purified by Hypercarb™ SPE followed P-2 Bio-gel size-exclusion column chromatography (see **2.2 J** for details).

**e) General procedure for the installation of terminal α2,6-Neu5Ac using ST6GAL1**

Acceptor (1 eq) and CMP-Neu5Ac (1.5 eq) were dissolved at a final acceptor concentration of 5 mM in a sodium cacodylate buffer (100 mM, pH 6.5) containing BSA (1% volume total, stock solution = 10 mg/mL). CIAP (1% volume total, stock solution = 1kU/mL) and ST6GAL1 (1% *wt/wt* relative to acceptor) were added. The reaction mixture was incubated with gentle shaking overnight at 37 °C. The progress of the reaction was monitored by LC-ESI-IT-TOF MS, and additional ST6Gal1 and donor were added if partial conversion was observed. The product was purified by Hypercarb™ SPE followed P-2 Bio-gel size-exclusion column chromatography (see **2.2 J** for details).

**f) General procedure for the installation of terminal α2,3-Neu5Ac using ST3GAL4**

Acceptor substrate (1 eq) and CMP-Neu5Ac (1.5 eq) were dissolved at a final acceptor concentration of 5 mM in a sodium cacodylate buffer (100 mM, pH 7.2) containing BSA (1% volume total, stock solution = 10 mg/mL). CIAP (1% volume total, stock solution = 1kU/mL) and ST3GAL4 (1% *wt/wt* relative to acceptor) were added. The reaction mixture was incubated with gentle shaking overnight at 37 °C. The progress of the reaction was monitored by LC-ESI-IT-TOF MS, and additional ST3Gal 4 and donor were if the reaction has not gone to completion. The product was purified by Hypercarb™ SPE followed P-2 Bio-gel size-exclusion column chromatography (see **2.2 J** for details).

**g) General procedure for removal of TFA group of synthesized *N*-glycans**

The substrate was dissolved in deionized water providing a final substrate concentration of 10 mM. The pH of reaction solution was adjusted to 10 using 1 M of NaOH solution, and the mixture solution was incubated 1-2 h at 37 °C with shaking. The reaction progress was monitored by LC-ESI-IT-TOF MS, and once had gone to completion, it was neutralized by 1 M aqueous acetic acid. The product was purified by Hypercarb™ SPE followed P-2 Bio-gel size-exclusion column chromatography (see **2.2 J** for details).

**h) General procedure for amine acetylation of synthesized *N*-glycans**

The amine containing *N*-glycans was dissolved (1 eq) in 50 mM of sodium acetate (NaOAc) buffered solution (pH = 8.5) with a final substrate concentration of 10 mM. Solids of AcOSu (10 eq) were added to convert the $NH_2$ moiety into NHAc moiety, and the reaction mixture was vigorously vortexed until all solids were dissolved. The reaction mixture was incubated with gentle shaking 1-2 h at 37 °C. The reaction progress was monitored by LC-ESI-IT-TOF MS, and additional AcOSu was added if starting material was remaining. The product was purified by P-2 Bio-gel size-exclusion column chromatography and lyophilized to yield the desired product.

**i) General procedure for removal of Cbz group of synthesized *N*-glycans**

The Cbz containg starting material was dissolved in a 10% *t*-butanol/water solution (1 mg/mL of substrate concentration). To the solution was added a palladium hydroxide on carbon (20% *wt* suspended in 100 µL *t*-butanol/water solution) with 1 mg/mL final concentration of 20% $Pd(OH)_2/C$ solid. The reaction was stirred vigorously under an atmosphere of hydrogen (1 atm) and monitored by LC-ESI-IT-TOF MS. Once the reaction had gon to completion, it was filtered through a Whatman ® syringe filter (0.2 µm) to remove the catalyst and the filtrate was lyophilized to provide the final product.

**j) General procedure for purification of synthesized *N*-glycans**

A combined purification system was constructed from pretreatment of reaction mixture, size-exclusion chromatography and Hypercarb™ SPE column to purify all of synthesized *N*-glycans.
*Pretreatment*: all of reaction mixtures were heated 10 min at 80 °C to precipitate the enzyme. The Suspension was centrifugated for a perio of short-time, and the decanted solution was lyophilized to yield a solid powder.
*Purification*: To purify the base-sensitive products, Hypercarb™ SPE column was used in the first step using 50 mM of buffered ammonium formate as eluant (pH = 7, including 0%-50% (*v/v*) of acetonitrile). Fractions containing product were collected and lyophilized. The lyophilized product was dissolved and loaded on a P-2 Bio-gel size-exclusion column which was eluted with a 5% (*v/v*) *n*-butanol/water, and the product fractions were collected and lyophilized to give the desired product as a fluffy white powder. To purify all of the other

products, the same procedures was applied but different buffered elution solution were used. The 50 mM ammonium bicarbonate buffer was replace the 50 mM of ammonium formate buffer for eluting the SPE column, and the 0.1 M ammonium bicarbonate buffer was used for eluting of the P-2 Bio-gel column rather than the 5% (*v/v*) *n*-butanol/water solution. Residues sugar nucleotides were eluted using 0%-10% (*v/v*) acetonitrile/salt solution and the different *N*-glycan were eluted and collected using 11%-18% (*v/v*) acetonitrile/salt solution.

**k) General procedure for performing the LC-MS spectra of final compounds**

LC-MS procedure was performed on a Shimadzu LC-ESI-IT-TOF with a Waters XBridge BEH, Amide column, 2.5 µm, 130 Å, 2.1×150 mm at a flow rate of 0.15 mL/min. Mobile phase A was 10 mM ammonium formate in water (pH 4.5); mobile phase B was aceteonitrile (LC-MS grade). The final compounds were purified using a linear gradient with the following conditions (**1** and/or **2**):

| | Condition 1 | Condition 2 |
|---|---|---|
| **Time (min)** | **B (%)** | **B (%)** |
| 0 | 65 | 85 |
| 18 | 50 | 60 |
| 20 | 25 | 25 |
| 24 | 25 | 25 |

#### 2.3 Schemes S1 and S2

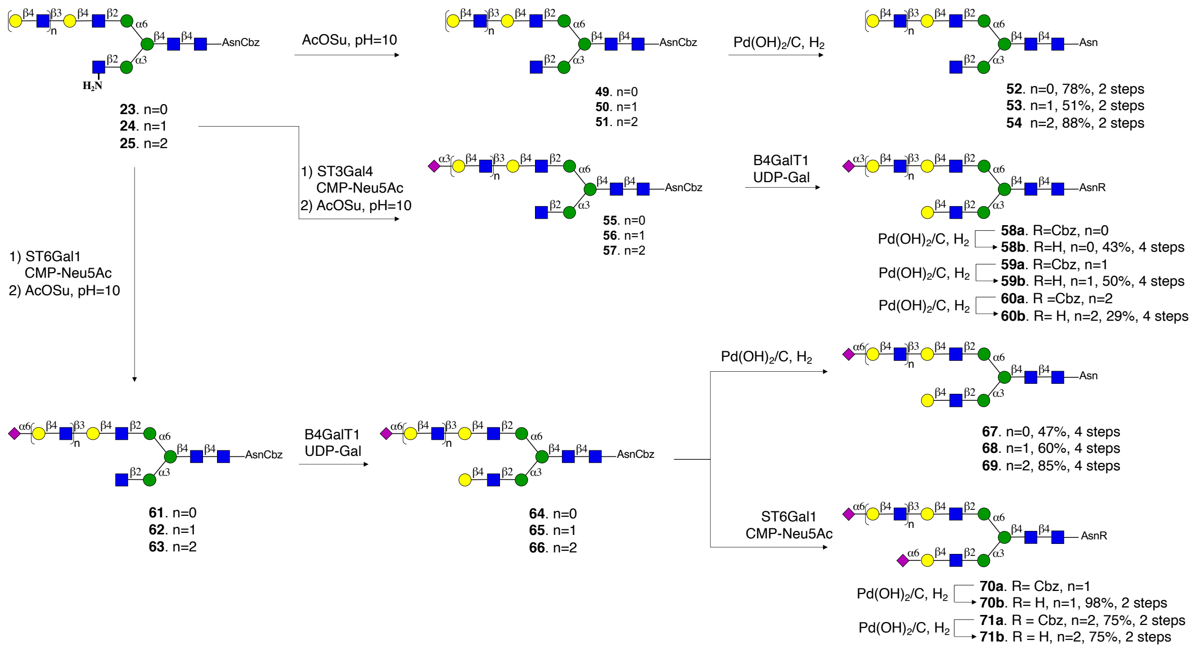


**Scheme S1.** Selective modification of termini at the α(1,3)- and α(1,6)-antennae starting for compounds **23-25**.

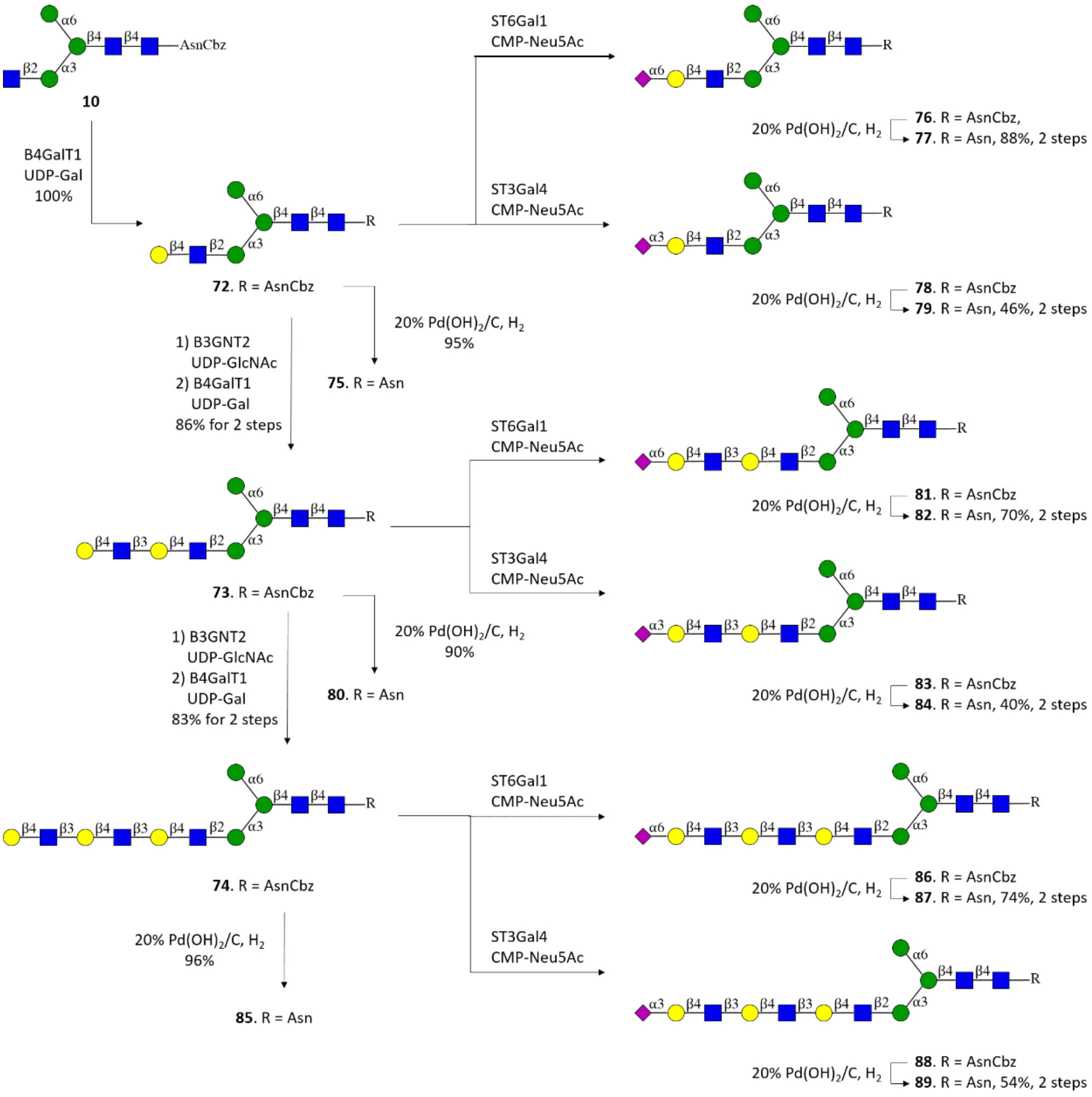


**Scheme S2.** Selective modification of termini of the α(1,3)-arm starting from **72**, **73** and **74**.

#### 2.4 NMR nomenclature

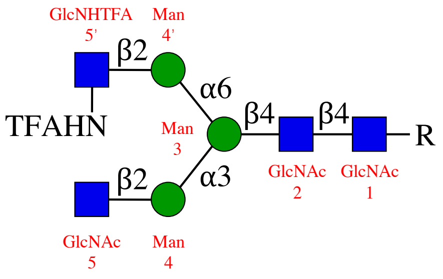


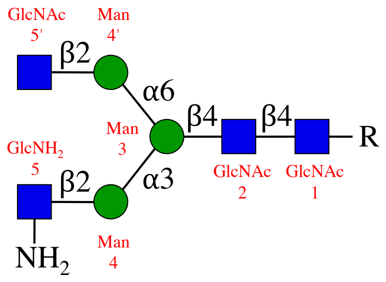


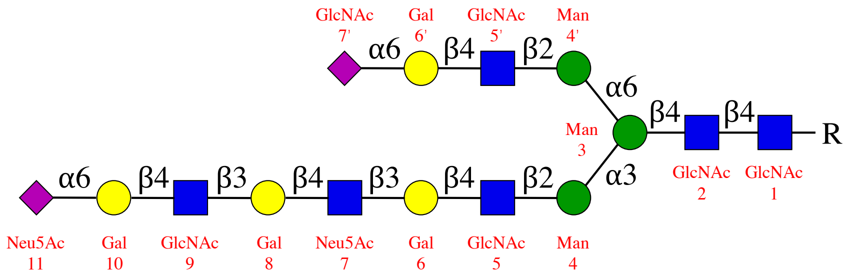


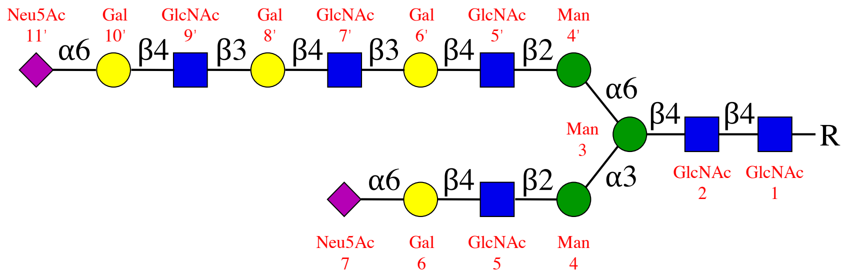


R = Asn or AsnCbz

#### 2.5 Enzymatic reactions

##### Compound 9[1,2]

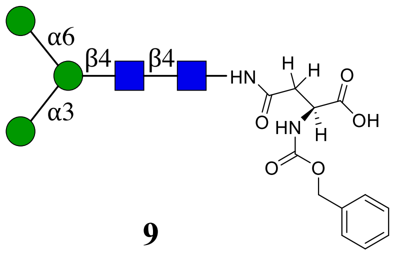


**9**

**$^{1}H$ (600 MHz, $D_2O$): δ (ppm)**

| | H1 | H2 | H3 | H4 | H5 | H6 |
|---|---|---|---|---|---|---|
| **GlcNAc1** | 5.05 (d, *J* = 9.7 Hz, 1H) | 3.82 | 3.73 | 3.64 | 3.54 | 3.80, 3.62 |
| **GlcNAc2** | 4.60 (d, *J* = 8.0 Hz, 1H) | 3.79 | 3.79 | 3.72 | 3.60 | 3.87, 3.74 |
| **Man3** | 4.78 (s, 1H) | 4.26 (d, *J* = 2.4 Hz, 1H) | 3.76 | 3.76 | 3.65 | 3.92, 3.80 |
| **Man4** | 5.10 | 4.07 (dd, *J* = 3.4, 1.7 Hz, 1H) | 3.88 | 3.63 | 3.80 | 3.91, 3.73 |
| **Man4'** | 4.91 (d, *J* = 1.7 Hz, 1H) | 3.97 (dd, *J* = 3.5, 1.7 Hz, 1H) | 3.88 | 3.64 | 3.64 | 3.88, 3.76 |

**$^{13}C$ (150 MHz, $D_2O$): δ (ppm)**

| | C1 | C2 | C3 | C4 | C5 | C6 |
|---|---|---|---|---|---|---|
| **GlcNAc1** | 78.03 | 53.63 | 72.66 | 78.49 | 76.10 | 59.75 |
| **GlcNAc2** | 101.14 | 54.78 | 65.72 | 79.56 | 74.29 | 59.90 |
| **Man3** | 100.28 | 70.06 | 80.41 | 71.87 | 74.07 | 65.72 |
| **Man4** | 102.44 | 69.91 | 70.23 | 66.77 | 73.35 | 61.04 |
| **Man4'** | 99.51 | 69.78 | 70.31 | 66.69 | 72.58 | 60.86 |

| Signal | Proton | Carbon |
|---|---|---|
| **NH<u>C</u>(O)CH3** | -[a] | 174.68, 174.60 |
| **NHC(O)<u>CH3</u>** | 2.07 (s, 3H), 1.91 (s, 3H) | 22.09, 21.89 |
| **Aromatic** | 7.51 – 7.34 (m, 5H) | 136.21, 128.67, 128.28, 127.73, 127.56 |
| **<u>CH2</u>-Ph** | 5.19 – 5.08 | 66.93 |
| **NH-<u>C</u>OO-** | - | 157.70 |
| **NH-<u>CH</u>-COOH** | 4.38 (dd, *J* = 9.0, 4.3 Hz, 1H) | 52.38 |
| **NH-CH-<u>C</u>OOH** | - | 176.86 |
| **C(O)-<u>CH2</u>-CH** | 2.82 (dd, *J* = 15.6, 4.3 Hz, 1H), 2.63 (dd, *J* = 15.6, 9.1 Hz, 1H) | 38.17 |
| **<u>C</u>(O)-CH2-CH** | - | 173.36 |

[a] Not applicable

ESI TOF-MS *m/z* calculated for $C_{46}H_{69}N_4O_{30}$, $[M-H]^-$: 1157.4002, found 1157.3935.

**Characterization of compound 9**

Initial examination of the $^{13}C$-$^{1}H$ HSQC (Fig. S1) illustrated the proton signals of the anomeric region, the $H_2$ of Man 3, 4 & 4', the Asn and the Cbz group. Especially, five unique signals in the anomeric region could clearly be assigned, which corresponded to five monosaccharides (Fig. S2). After the $H_1$ of Man 3, 4 and 4' were confirmed, the COSY data led us to distinguish each $H_2$ of three mannosides. As shown in Fig. S3, the $H_1$ of Man 3 have a neighboring correlation with the $H_2$ from the Man 3 spin system. The cross peak at 4.36 ppm correspond to the $H_2$ of Man 3. Similarly, the $H_2$ of Man 4 and 4' are observed at 4.07ppm and 3.97 ppm, respectively.

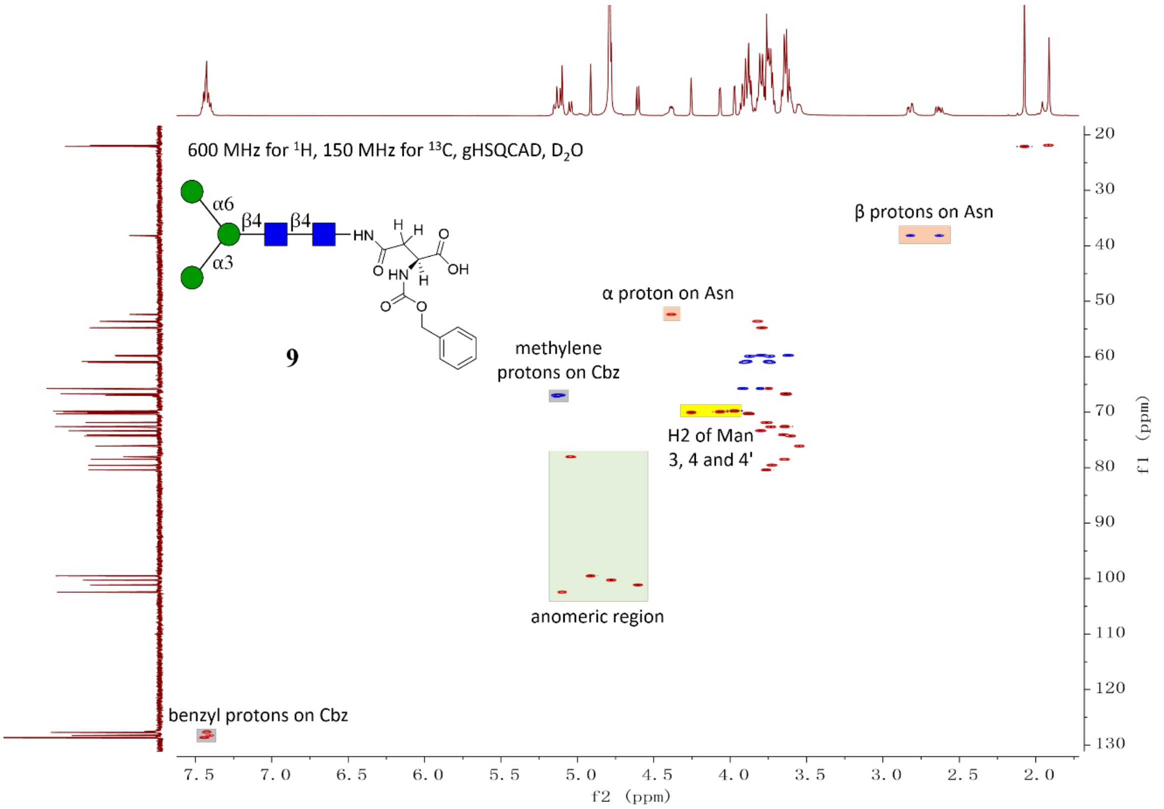


**Figure S1.** HSQC for compound **9**.

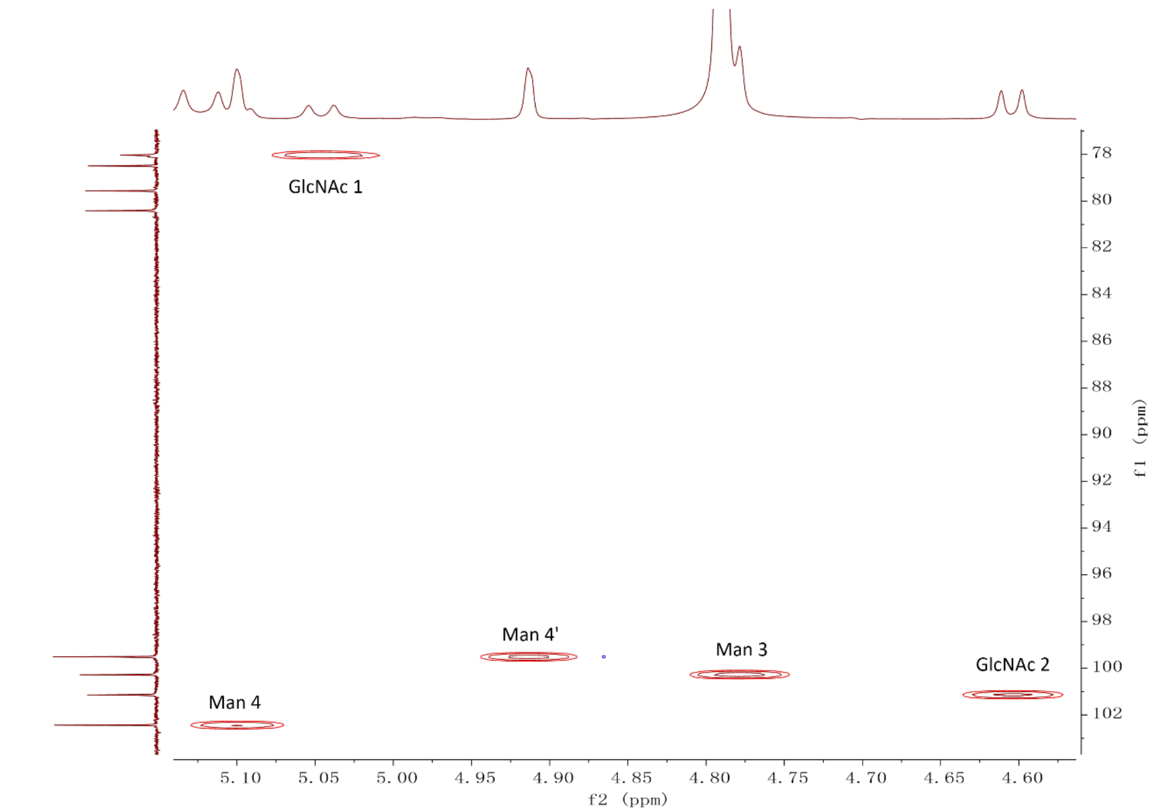


**Figure S2.** Expanded HSQC for anomeric region of compound **9**.

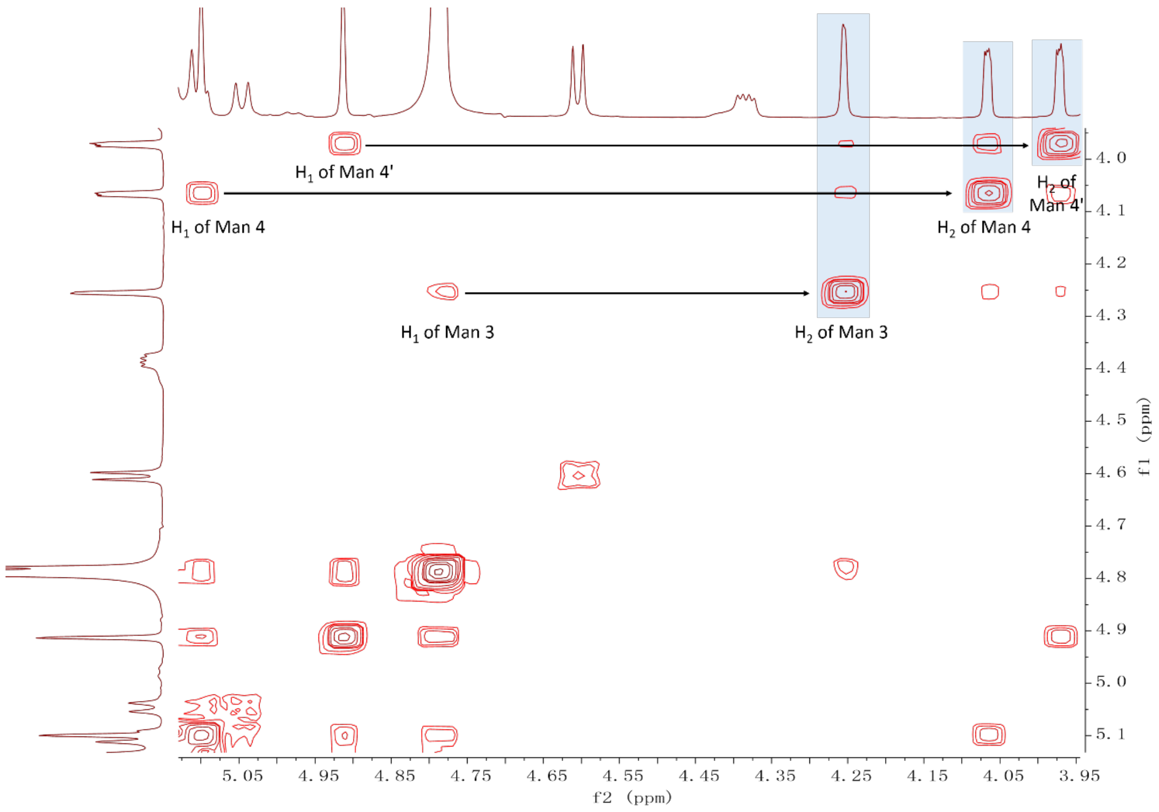


**Figure S3.** Expanded COSY for Mannose $H_{1\&2}$ region of compound **9**.

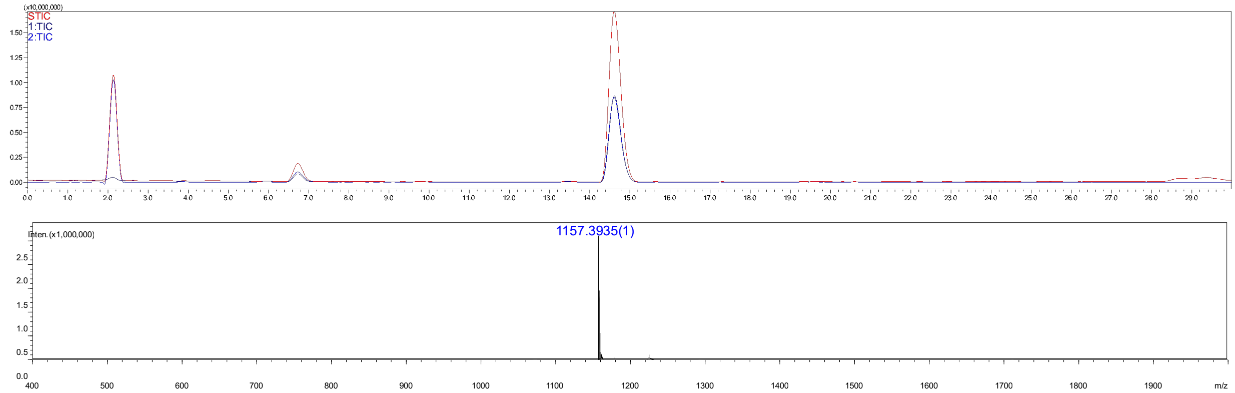


**Figure S4.** Analytical HPLC-MS chromatogram of compound **9**. The retention time = 14.6 min.

#### Compound 10

**10** was synthesized from starting material **9** (10 mg) according to the general procedure **2.2 a** for the installation of GlcNAc moiety with UDP-GlcNAc and recombinant MGAT 1. The product was purified using a combined purification system **(2.2 j)** providing **10** as a white fluffy solid (10.7 mg, 91%).

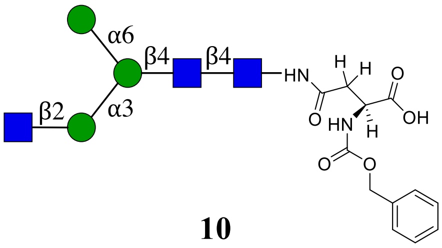


**10**

**$^1$H (600 MHz, $D_2O$): $\delta$ (ppm)**

| | H1 | H2 | H3 | H4 | H5 | H6 |
|---|---|---|---|---|---|---|
| **GlcNAc1** | 5.05 (d, $J$ = 9.7 Hz, 1H) | 3.82 | 3.74 | 3.65 | 3.55 | 3.80, 3.62 |
| **GlcNAc2** | 4.61 (d, $J$ = 8.0 Hz, 1H) | 3.79 | 3.75 | 3.74 | 3.60 | 3.88, 3.75 |
| **Man3** | 4.78 (s, 1H) | 4.25 (d, $J$ = 2.5 Hz, 1H) | 3.77 | 3.76 | 3.66 | 3.92, 3.80 |
| **Man4** | 5.17 – 5.09 | 4.19 (dd, $J$ = 3.3, 1.6 Hz, 1H) | 3.90 | 3.50 | 3.74 | 3.93, 3.62 |
| **Man4'** | 4.92 (d, $J$ = 1.8 Hz, 1H) | 3.97 (dd, $J$ = 3.4, 1.7 Hz, 1H) | 3.88 | 3.64 | 3.64 | 3.89, 3.75 |
| **GlcNAc5** | 4.55 (d, $J$ = 8.5 Hz, 1H) | 3.70 | 3.56 | 3.45 | 3.44 | 3.91, 3.76 |

$^{13}$C (150 MHz, $D_2O$): $\delta$ (ppm)

| | C1 | C2 | C3 | C4 | C5 | C6 |
|---|---|---|---|---|---|---|
| **GlcNAc1** | 78.05 | 53.65 | 72.63 | 78.48 | 76.11 | 59.75 |
| **GlcNAc2** | 101.15 | 54.77 | 65.80 | 79.58 | 74.26 | 59.86 |
| **Man3** | 100.31 | 70.10 | 80.29 | 71.87 | 74.06 | 65.75 |
| **Man4** | 99.53 | 76.32 | 69.30 | 67.19 | 73.46 | 61.61 |
| **Man4'** | 99.52 | 69.78 | 70.30 | 66.67 | 72.59 | 60.86 |
| **GlcNAc5** | 99.50 | 55.22 | 73.17 | 69.81 | 75.72 | 60.53 |

| Signal | Proton | Carbon |
|---|---|---|
| **NH<u>C</u>(O)CH₃** | -[a] | 174.67, 174.61 |
| **NHC(O)<u>CH₃</u>** | 2.08 (s, 3H), 2.05 (s, 3H), 1.92 (s, 3H) | 22.23, 22.09, 21.88 |
| **Aromatic** | 7.57 – 7.29 (m, 5H) | 136.18, 128.67, 128.29, 127.69 |
| **<u>CH₂</u>-Ph** | 5.17 – 5.09 (m, 2H) | 67.00 |
| **NH-<u>C</u>OO-** | - | 157.72 |
| **NH-<u>CH</u>-COOH** | 4.45 (dd, $J$ = 8.6, 4.6 Hz, 1H) | 51.79 |
| **NH-CH-<u>C</u>OOH** | - | 176.07 |
| **C(O)-<u>CH₂</u>-CH** | 2.84 (dd, $J$ = 15.7, 4.5 Hz, 1H),<br>2.69 (dd, $J$ = 15.7, 8.6 Hz, 1H) | 37.84 |
| **<u>C</u>(O)-CH₂-CH** | - | 173.11 |

[a] Not applicable

ESI TOF-MS $m/z$ calculated for $C_{54}H_{82}N_5O_{35}$ $[M-H]^-$: 1360.4796, found 1360.4693.

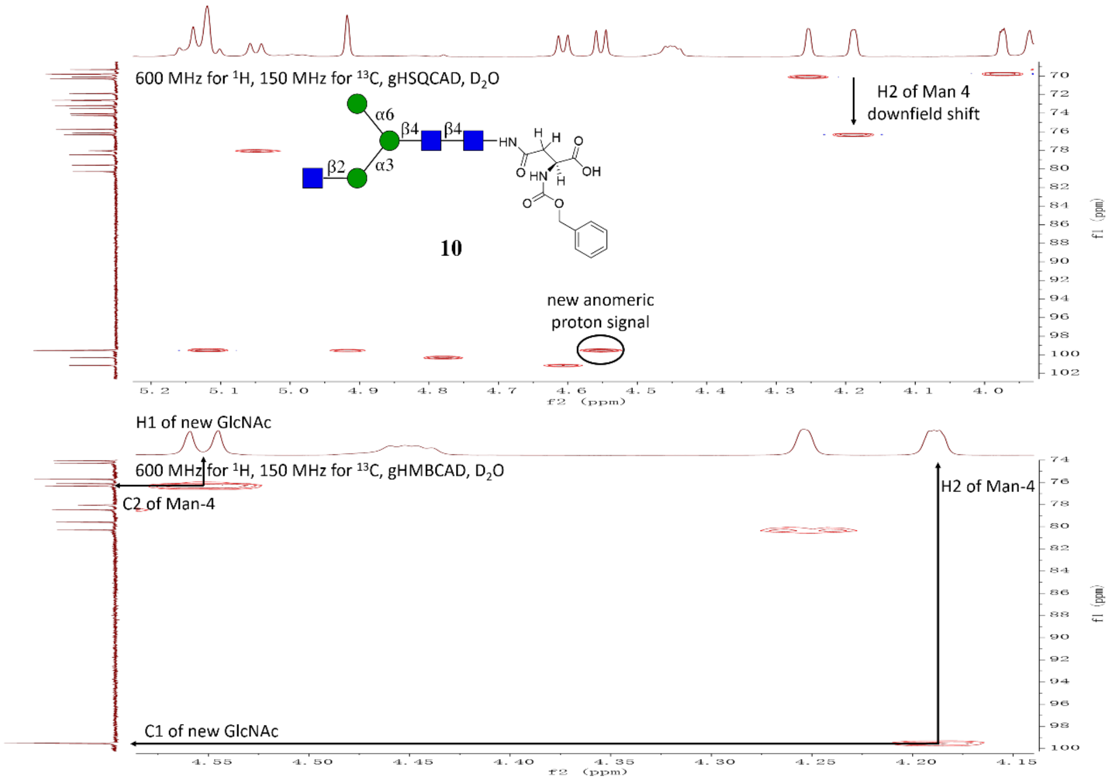


**Figure S5.** Expanded HSQC and HMBC of compound **10**.

#### Compound 13

**13** was synthesized from starting material **9** (10 mg, 1.0 eq) according to the general procedure **2.2 a** for the installation of GlcNTFA moiety with UDP-GlcNTFA and MGAT 1. The product was purified using the combined purification system **(2.2 j)** providing **13** as a white fluffy solid (9.8 mg, 80%).

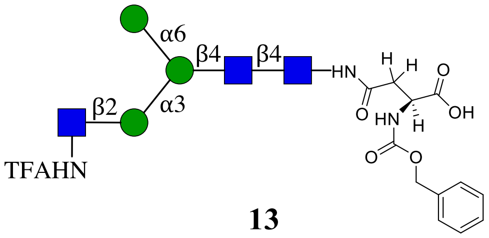


**13**

**$^{1}$H (600 MHz, $D_2O$): δ (ppm)**

| | H1 | H2 | H3 | H4 | H5 | H6 |
|---|---|---|---|---|---|---|
| **GlcNAc1** | 5.04 (d, $J$ = 9.7 Hz, 1H) | 3.81 | 3.72 | 3.64 | 3.54 | 3.79, 3.61 |
| **GlcNAc2** | 4.60 (d, $J$ = 8.0 Hz, 1H) | 3.78 | 3.74 | 3.73 | 3.59 | 3.87, 3.74 |
| **Man3** | 4.77 (s, 1H) | 4.24 (s, 1H) | 3.75 | 3.76 | 3.65 | 3.91, 3.79 |
| **Man4** | 5.07 (s, 1H) | 4.21 (m, 1H) | 3.90 | 3.45 | 3.72 | 3.89, 3.56 |
| **Man4'** | 4.77 (s, 1H) | 3.96 (m, 1H) | 3.87 | 3.63 | 3.64 | 3.88, 3.74 |
| **GlcTFA5** | 4.67 (d, $J$ = 8.3 Hz, 1H) | 3.80 | 3.67 | 3.47 | 3.47 | 3.92, 3.76 |

**$^{13}$C (150 MHz, $D_2O$): δ (ppm)**

| | C1 | C2 | C3 | C4 | C5 | C6 |
|---|---|---|---|---|---|---|
| **GlcNAc1** | 78.01 | 54.77 | 72.67 | 78.47 | 76.09 | 59.75 |
| **GlcNAc2** | 101.15 | 54.77 | 65.78 | 79.59 | 74.25 | 59.81 |
| **Man3** | 100.33 | 70.05 | 80.27 | 71.87 | 74.03 | 65.73 |
| **Man4** | 99.36 | 76.27 | 69.16 | 67.31 | 73.51 | 61.52 |
| **Man4'** | 99.53 | 69.77 | 70.30 | 66.66 | 72.59 | 60.85 |
| **GlcTFA5** | 98.71 | 55.80 | 72.51 | 69.77 | 75.75 | 60.47 |

| Signal | Proton | Carbon |
|---|---|---|
| **NH<u>C</u>(O)CH₃** | -[a] | 174.68, 174.60 |
| **NHC(O)<u>CH₃</u>** | 2.07 (s, 3H), 1.90 (s, 3H) | 22.08, 21.89 |
| **NH<u>C</u>(O)CF₃** | - | 159.63, 159.38 (d) |
| **NHC(O)<u>CF₃</u>** | - | 116.69-111.38 (m) |
| **Aromatic** | 7.48 – 7.36 (m, 5H) | 136.23, 128.66, 128.27, 127.73 |

| | | |
|---|---|---|
| **CH2-Ph** | 5.16 – 5.09 (m, 2H) | 66.60 |
| **NH-COO-** | - | 157.69 |
| **NH-CH-COOH** | 4.35 (dd, *J* = 8.1, 4.5 Hz, 1H) | 52.66 |
| **NH-CH-COOH** | - | 177.23 |
| **C(O)-CH2-CH** | 2.81 (dd, *J* = 15.3, 3.8 Hz, 1H),<br>2.60 (dd, *J* = 15.4, 9.4 Hz, 1H) | 38.32 |
| **C(O)-CH2-CH** | - | 173.49 |

[a] Not applicable

ESI TOF-MS *m/z* calculated for $C_{54}H_{79}F_3N_5O_{35}$, $[M-H]^-$: 1414.4513, found 1414.4509.

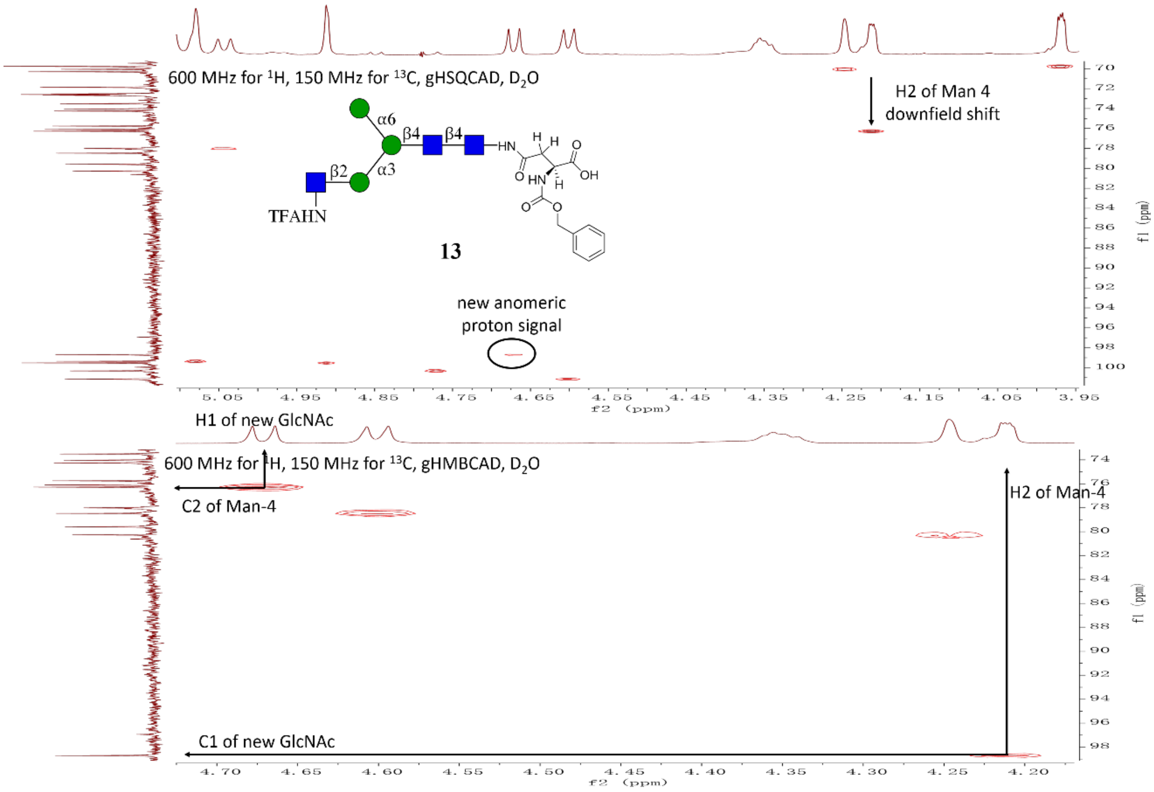


**Figure S6.** Expanded HSQC and HMBC of compound **13**.

#### Compound 20

Compound **10** (10.7 mg) was subjected procedures **2.2 b** to install GlcNTFA at the MGAT2 arm to provide intermediate **11** which was subjected to the purification protocol **2.2 j**. Compound **11** was subjected to the general procedures **2.2 g** and **c** for removal of the TFA moiety to give $GlcNH_2$ and galactosylation by B4GalT1. The product was purified using the described purification system **(2.2 j)** providing **20** as a white fluffy solid (11.0 mg, 83% yield over three steps).

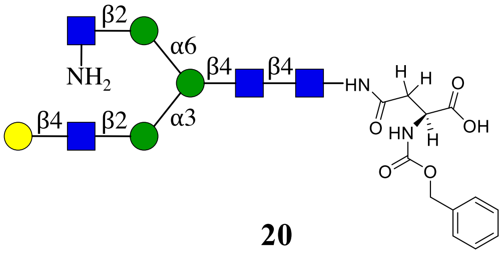


**20**

NMR and MS analysis of compound **11**

**$^{1}$H (600 MHz, $D_2O$): δ (ppm)**

| | H1 | H2 | H3 | H4 | H5 | H6 |
|---|---|---|---|---|---|---|
| **GlcNAc1** | 5.04 (d, *J* = 9.7 Hz, 1H) | 3.81 | 3.73 | 3.64 | 3.55 | 3.79, 3.62 |
| **GlcNAc2** | 4.60 (d, *J* = 7.9 Hz, 1H) | 3.78 | 3.77 | 3.72 | 3.60 | 3.86, 3.73 |
| **Man3** | 4.76 (s, 1H) | 4.24 (d, *J* = 2.5 Hz, 1H) | 3.76 | 3.75 | 3.73 | 3.94, 3.77 |
| **Man4** | 5.11 (s, 1H) | 4.20 – 4.16 (m, 1H) | 3.89 | 3.49 | 3.73 | 3.91, 3.60 |
| **Man4'** | 4.88 (s, 1H) | 4.15 – 4.11 (m, 1H) | 3.89 | 3.43 | 3.59 | 3.87, 3.56 |
| **GlcNAc5** | 4.55 (d, *J* = 8.4 Hz, 1H) | 3.69 | 3.55 | 3.45 | 3.44 | 3.90, 3.75 |
| **GlcTFA5'** | 4.67 (d, *J* = 8.4 Hz, 1H) | 3.80 | 3.65 | 3.49 | 3.44 | 3.92, 3.77 |

**$^{13}$C (150 MHz, $D_2O$): δ (ppm)**

| | C1 | C2 | C3 | C4 | C5 | C6 |
|---|---|---|---|---|---|---|
| **GlcNAc1** | 78.03 | 53.64 | 72.64 | 78.46 | 76.10 | 59.75 |
| **GlcNAc2** | 101.16 | 54.82 | 65.59 | 79.47 | 74.29 | 59.82 |
| **Man3** | 100.35 | 70.09 | 80.35 | 71.89 | 74.15 | 65.80 |
| **Man4** | 99.51 | 76.32 | 69.30 | 67.20 | 73.46 | 61.60 |
| **Man4'** | 96.81 | 76.24 | 69.24 | 67.39 | 72.80 | 61.48 |
| **GlcNAc5** | 99.51 | 55.23 | 73.17 | 69.81 | 75.72 | 60.53 |
| **GlcTFA5'** | 98.74 | 55.83 | 72.64 | 69.73 | 75.75 | 60.45 |

| Signal | Proton | Carbon |
|---|---|---|
| **NH<u>C</u>(O)CH3** | -[a] | 174.67, 174.55 |
| **NHC(O)<u>CH3</u>** | 2.24 – 1.82 (m, 9H) | 22.23, 22.09, 21.89 |
| **NH<u>C</u>(O)CF3** | - | 159.62, 159.37 |
| **NHC(O)<u>CF3</u>** | - | 116.69, 114.80 |
| **Aromatic** | 7.51 – 7.32 (m, 5H) | 128.67, 128.28, 127.71 |
| **<u>CH2</u>-Ph** | 5.16 – 5.07 (m, 2H) | 66.95 |

| | | |
|---|---|---|
| **NH-C̲OO-** | - | 157.70 |
| **NH-C̲H̲-COOH** | 4.42 – 4.38 (m, 1H) | 52.20 |
| **NH-CH-C̲OOH** | - | 176.56 |
| **C(O)-C̲H̲₂-CH** | 2.82 (dd, $J$ = 15.6, 4.4 Hz, 1H),<br>2.65 (dd, $J$ = 15.5, 9.1 Hz, 1H) | 38.06 |
| **C̲(O)-CH₂-CH** | - | 173.29 |

[a] Not applicable

ESI TOF-MS $m/z$ calculated for $C_{62}H_{91}F_3N_6O_{40}$, $[M-2H]^{2-}$: 808.2617, found 808.2627.

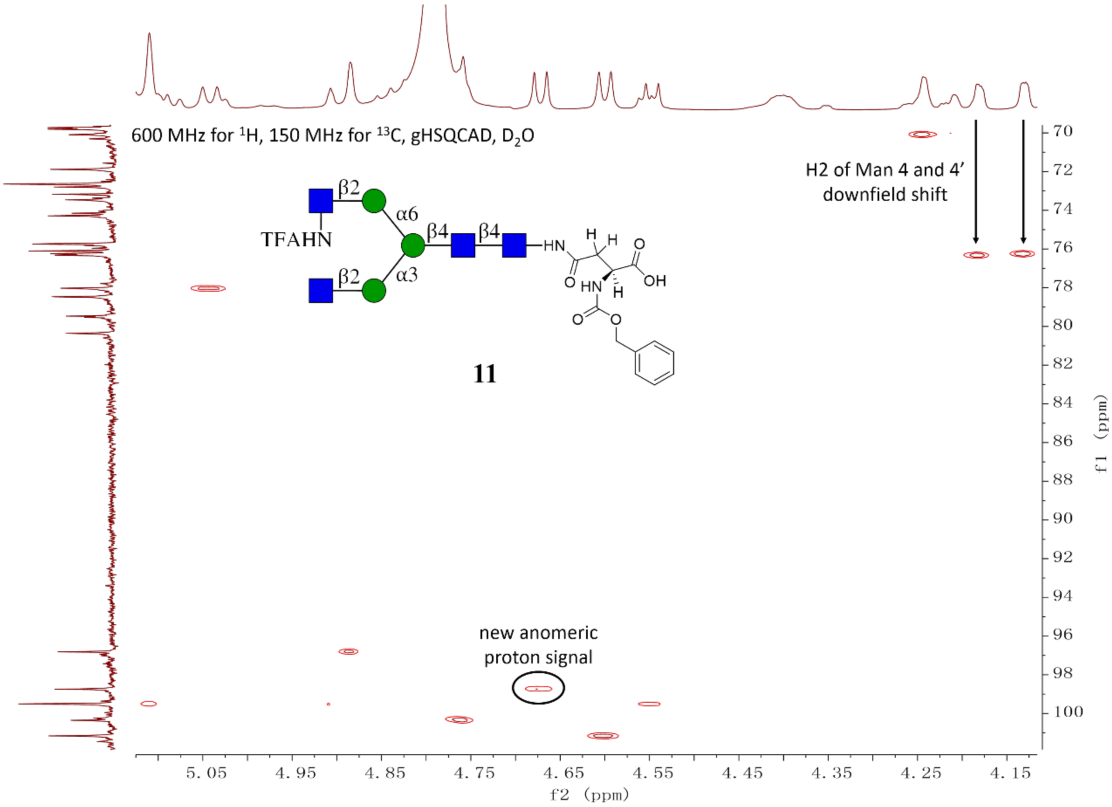


**Figure S7.** Expanded HSQC and HMBC of compound **11**.

NMR and MS analysis of compound **20**:

**$^1$H (600 MHz, $D_2O$): δ (ppm)**

| | H1 | H2 | H3 | H4 | H5 | H6 |
|---|---|---|---|---|---|---|
| **GlcNAc1** | 5.04 | 3.82 | 3.73 | 3.64 | 3.55 | 3.79, 3.62 |
| **GlcNAc2** | 4.60 | 3.79 | 3.77 | 3.74 | 3.60 | 3.88, 3.75 |
| **Man3** | 4.77 | 4.26 | 3.77 | 3.77 | 3.64 | 3.96, 3.80 |

| | | | | | | |
|---|---|---|---|---|---|---|
| **Man4** | 5.11 | 4.19 (d, *J* = 3.3 Hz, 1H) | 3.90 | 3.50 | 3.75 | 3.93, 3.61 |
| **Man4'** | 5.03 | 4.26 | 3.96 | 3.70 | 3.66 | 3.87, 3.83 |
| **GlcNAc5** | 4.58 | 3.73 | 3.73 | 3.73 | 3.57 | 3.98, 3.85 |
| **GlcNH$_2$5'** | 4.79 | 3.09 (t, *J* = 9.5 Hz, 1H) | 3.62 | 3.49 | 3.48 | 3.92, 3.77 |
| **Gal6** | 4.47 (d, *J* = 7.8 Hz, 1H) | 3.54 | 3.66 | 3.92 | 3.73 | 3.77, 3.75 |

**$^{13}$C (150 MHz, D$_2$O): δ (ppm)**

| | C1 | C2 | C3 | C4 | C5 | C6 |
|---|---|---|---|---|---|---|
| **GlcNAc1** | 78.00 | 53.63 | 72.70 | 78.47 | 76.10 | 59.77 |
| **GlcNAc2** | 101.15 | 54.81 | 65.62 | 79.39 | 74.27 | 59.81 |
| **Man3** | 100.29 | 70.08 | 80.36 | 71.88 | 74.07 | 65.92 |
| **Man4** | 99.45 | 76.24 | 69.30 | 67.18 | 73.47 | 61.60 |
| **Man4'** | 97.44 | 76.07 | 69.26 | 66.48 | 72.44 | 60.08 |
| **GlcNAc5** | 99.32 | 54.79 | 71.85 | 78.40 | 74.65 | 59.90 |
| **GlcNH$_2$5'** | 97.58 | 55.23 | 72.29 | 69.45 | 76.24 | 60.26 |
| **Gal6** | 102.83 | 70.85 | 72.41 | 68.44 | 75.26 | 60.93 |

| Signal | Proton | Carbon |
|---|---|---|
| **NHC(O)CH$_3$** | -[a] | 174.61, 174.56 |
| **NHC(O)CH$_3$** | 2.08 (s, 3H), 2.05 (s, 3H), 1.91 (s, 3H) | 22.25, 22.12, 21. 91 |
| **Aromatic** | 7.53 – 7.33 (m, 5H) | 136.25, 128.67, 128.27, 127.75 |
| **CH$_2$-Ph** | 5.17 – 5.07 (m, 2H) | 66.88 |
| **NH-COO-** | - | 157.68 |
| **NH-CH-COOH** | 4.33 (dd, *J* = 9.5, 4.1 Hz, 1H) | 52.94 |
| **NH-CH-COOH** | - | 177.57 |
| **C(O)-CH$_2$-CH** | 2.81 (dd, *J* = 15.5, 4.2 Hz, 1H)<br>2.59 (dd, *J* = 15.4, 9.4 Hz, 1H) | 38.47 |
| **C(O)-CH$_2$-CH** | - | 173.62 |

[a] Not applicable

ESI TOF-MS *m/z* calculated for $C_{66}H_{102}N_6O_{44}$, $[M-2H]^{2-}$: 841.2970, found 841.3034.

#### Compound 21

**21** was prepared from **20** (11.0 mg, 1.0 eq) using the general procedures **2.2 d** and **c** for the installation of GlcNAc and Gal moieties. The product was purified using the described purification system **(2.2 j)** providing **21** as a white fluffy solid (11.6 mg, 87% yield over two steps).

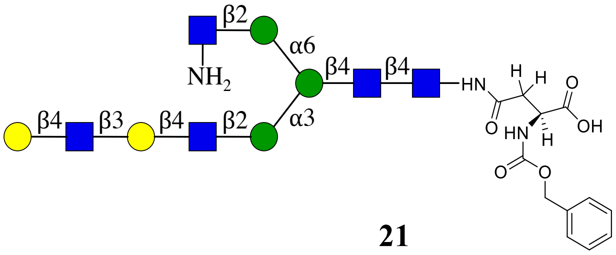


**21**

**$^{1}$H (600 MHz, $D_2O$): δ (ppm)**

| | H1 | H2 | H3 | H4 | H5 | H6 |
|---|---|---|---|---|---|---|
| **GlcNAc1** | 5.04 | 3.82 | 3.73 | 3.64 | 3.54 | 2.79, 3.62 |
| **GlcNAc2** | 4.60 (d, *J* = 7.8 Hz, 1H) | 3.79 | 3.77 | 3.74 | 3.60 | N/R[b] |
| **Man3** | 4.77 | 4.25 | 3.77 | 3.77 | 3.64 | 3.96, 3.80 |
| **Man4** | 5.11 | 4.19 (d, *J* = 3.4 Hz, 1H) | 3.90 | 3.50 | 3.74 | 3.92, 3.61 |
| **Man4'** | 5.04 | 4.24 | 3.96 | 3.69 | N/R | N/R |
| **GlcNAc5** | 4.58 (d, *J* = 7.0 Hz, 1H) | 3.73 | 3.73 | 3.72 | 3.57 | N/R |
| **GlcNH$_2$5'** | 4.73 (d, *J* = 8.1 Hz, 1H) | 3.02 (t, *J* = 9.4 Hz, 1H) | 3.58 | 3.48 | 3.48 | 3.92, 3.77 |
| **Gal6** | 4.45 | 3.58 | 3.72 | 4.16 (d, *J* = 3.1 Hz, 1H) | 3.71 | N/R |
| **GlcNAc7** | 4.70 (d, *J* = 8.4 Hz, 1H) | 3.81 | N/R | 3.74 | N/R | N/R |
| **Gal8** | 4.48 (d, *J* = 7.8 Hz, 1H) | 3.54 | 3.67 | 3.92 | 3.73 | 3.78, 3.74 |

**$^{13}$C (150 MHz, $D_2O$): δ (ppm)**

| | C1 | C2 | C3 | C4 | C5 | C6 |
|---|---|---|---|---|---|---|
| **GlcNAc1** | 78.00 | 53.63 | 72.69 | 78.47 | 76.09 | 59.76 |
| **GlcNAc2** | 101.15 | 54.82 | 65.64 | 79.42 | 74.27 | 59.81 |
| **Man3** | 100.29 | 70.08 | 80.33 | 71.87 | 74.08 | 65.91 |
| **Man4** | 99.45 | 76.27 | 69.29 | 67.18 | 73.46 | 61.60 |
| **Man4'** | 97.44 | 76.19 | 69.29 | 66.57 | N/R | N/R |
| **GlcNAc5** | 99.34 | 54.73 | 71.84 | 78.47 | 74.63 | 59.89 |
| **GlcNH$_2$5'** | 98.37 | 55.36 | 72.84 | 69.46 | 76.19 | 60.32 |
| **Gal6** | 102.86 | 69.85 | 82.01 | 68.19 | 74.78 | N/R |
| **GlcNAc7** | 102.68 | 55.09 | N/R | 78.04 | N/R | N/R |
| **Gal8** | 102.76 | 70.87 | 72.41 | 68.45 | 75.26 | 60.94 |

| Signal | Proton | Carbon |
|---|---|---|
| **NH<u>C</u>(O)CH3** | -[a] | 174.81, 174.60, 174.55 |
| **NHC(O)<u>CH3</u>** | 2.08 (s, 3H), 2.05 (s, 3H), 2.03 (s, 3H), 1.91 (s, 3H) | 22.24, 22.11, 22.09, 21.90 |
| **Aromatic** | 7.49 – 7.36 (m, 5H) | 136.25, 128.66, 128.26, 127.75 |
| **<u>CH2</u>-Ph** | 5.17 – 5.07 (m, 2H) | 66.87 |
| **NH-<u>C</u>OO-** | - | 157.68 |
| **NH-<u>CH</u>-COOH** | 4.35 – 4.30 (m, 3H) | 52.95 |
| **NH-CH-<u>C</u>OOH** | - | N/R |
| **C(O)-<u>CH2</u>-CH** | 2.83 – 2.78 (m, 1H)<br>2.58 (dd, *J* = 15.4, 9.3 Hz, 1H) | 38.47 |
| **<u>C</u>(O)-CH2-CH** | - | 173.62 |

[a] Not applicable

[b] Not reported

ESI TOF-MS *m/z* calculated for $C_{80}H_{125}N_7O_{54}$, $[M-2H]^{2-}$: 1023.8631, found 1023.8725.

#### Compound 22

**22** was synthesized from **21** (11.6 mg) using the general procedures **2.2 d** and **c** for the installation of GlcNAc and Gal moieties. The product was purified using the described purification system **(2.2 j)** providing **22** as a white fluffy solid (11.5 mg, 77% yield over two steps).

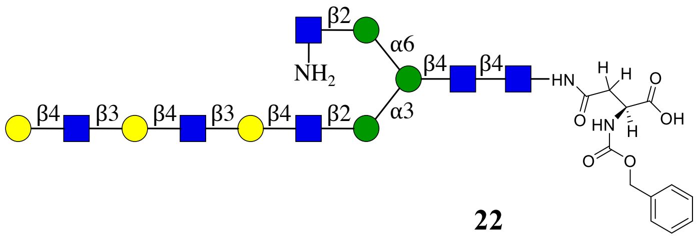


**22**

**¹H (600 MHz, D2O): δ (ppm)**

| | H1 | H2 | H3 | H4 | H5 | H6 |
|---|---|---|---|---|---|---|
| **GlcNAc1** | 5.04 | 3.81 | 3.73 | 3.64 | 3.54 | N/R[b] |
| **GlcNAc2** | 4.60 (d, *J* = 8.2 Hz, 1H) | 3.80 | 3.77 | 3.74 | N/R | N/R |
| **Man3** | 4.77 | 4.25 | 3.77 | 3.77 | N/R | 3.96, 3.79 |
| **Man4** | 5.11 | 4.19 (d, *J* = 3.4 Hz, 1H) | 3.89 | 3.50 | 3.74 | 3.92, 3.61 |
| **Man4'** | 5.03 | 4.25 | 3.96 | 3.69 | N/R | N/R |
| **GlcNAc5** | 4.58 (d, *J* = 7.1 Hz, 1H) | 3.73 | 3.72 | 3.71 | N/R | N/R |
| **GlcNH25'** | | 3.08 (t, *J* = 9.5 Hz, 1H) | N/R | 3.49 | N/R | N/R |
| **Gal6** | 4.45 | 3.58 | N/R | 4.15 | N/R | N/R |
| **GlcNAc7** | 4.70 | 3.80 | N/R | N/R | N/R | N/R |

| | | | | | | |
|---|---|---|---|---|---|---|
| **Gal8** | 4.47 | 3.58 | N/R | 4.15 | N/R | N/R |
| **GlcNAc9** | 4.70 | 3.80 | N/R | N/R | N/R | N/R |
| **Gal10** | 4.48 | 3.54 | 3.67 | 3.92 | 3.72 | N/R |

**$^{13}C$ (150 MHz, $D_2O$): δ (ppm)**

| | **C1** | **C2** | **C3** | **C4** | **C5** | **C6** |
|---|---|---|---|---|---|---|
| **GlcNAc1** | 78.00 | 53.62 | 72.69 | 78.45 | 76.09 | N/R |
| **GlcNAc2** | 101.14 | 54.81 | 65.63 | 79.39 | N/R | N/R |
| **Man3** | 100.28 | 70.07 | 80.33 | 71.88 | N/R | 65.92 |
| **Man4** | 99.45 | 76.22 | 69.29 | 67.17 | 73.46 | 61.59 |
| **Man4'** | 97.43 | 76.09 | 69.26 | 66.48 | N/R | N/R |
| **GlcNAc5** | 99.33 | 54.73 | 71.84 | 78.45 | N/R | N/R |
| **$GlcNH_25'$** | N/R | 55.25 | N/R | 69.45 | N/R | N/R |
| **Gal6** | 102.87 | 69.85 | N/R | 68.21 | N/R | N/R |
| **GlcNAc7** | 102.67 | 55.04 | N/R | N/R | N/R | N/R |
| **Gal8** | 102.76 | 69.85 | N/R | 68.21 | N/R | N/R |
| **GlcNAc9** | 102.67 | 55.09 | N/R | N/R | N/R | N/R |
| **Gal10** | 102.76 | 70.87 | 72.40 | 68.45 | 75.26 | 60.94 |

| **Signal** | **Proton** | **Carbon** |
|---|---|---|
| **$NH\underline{C}(O)CH_3$** | -[a] | 174.80, 174.69, 174.60, 174.55, 174.50 |
| **$NHC(O)\underline{CH_3}$** | 2.07 (s, 2H), 2.06 – 2.02 (m, 8H), 1.91 (s, 2H) | 22.23, 22.08, 21.89 |
| **Aromatic** | 7.52 – 7.31 (m, 5H) | 136.25, 128.66, 128.26, 127.75 |
| **$\underline{CH_2}$-Ph** | 5.17 – 5.07 (m, 2H) | 66.87 |
| **NH-$\underline{C}$OO-** | - | 157.67 |
| **NH-$\underline{CH}$-COOH** | 4.35 – 4.29 (m, 1H) | 52.93 |
| **NH-CH-$\underline{C}$OOH** | - | N/R |
| **C(O)-$\underline{CH_2}$-CH** | 2.81 (dd, $J$ = 15.7, 4.1 Hz, 1H), 2.58 (dd, $J$ = 15.3, 9.4 Hz, 1H) | 38.46 |
| **$\underline{C}$(O)-$CH_2$-CH** | - | 173.61 |

[a] Not applicable

[b] Not reported

ESI TOF-MS *m/z* calculated for $C_{94}H_{148}N_8O_{64}$, $[M-2H]^{2-}$: 1206.4292, found 1206.4386.

**Compound 23**

Compound **13** (9.8 mg) was subjected to MGAT2 and UDP-GlcNAc according to general procedure **2.2 b** to give **14**, which was purified according to **2.2 j**. Compound **14** was subjected to general procedures **2.2 g** and **c** to remove the TFA moiety and introduce Gal by B4GalT1 to provide after purification using general protocal **2.2 j** compound **23** as a white fluffy solid (10.3 mg, 88% yield for three steps).

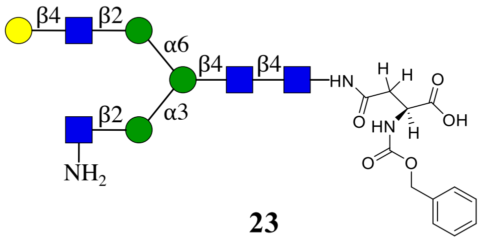


**23**

NMR and MS analysis of compound **14**:

**$^1$H (600 MHz, $D_2O$): δ (ppm)**

| | H1 | H2 | H3 | H4 | H5 | H6 |
|---|---|---|---|---|---|---|
| **GlcNAc1** | 5.05 (d, *J* = 9.7 Hz, 1H) | 3.83 | 3.74 | 3.65 | 3.55 | 3.81, 3.63 |
| **GlcNAc2** | 4.61 (d, *J* = 7.9 Hz, 1H) | 3.80 | 3.81 | 3.74 | 3.62 | 3.88, 3.74 |
| **Man3** | 4.76 (s, 1H) | 4.25 (d, *J* = 2.9 Hz, 1H) | 3.76 | 3.75 | 3.62 | 3.97, 3.78 |
| **Man4** | 5.08 (s, 2H) | 4.22 (dd, *J* = 3.3, 1.6 Hz, 1H) | 3.91 | 3.51 or 3.47 | 3.74 | 3.93, 3.62 |
| **Man4'** | 4.92 (s, 1H) | 4.11 (dd, *J* = 3.4, 1.7 Hz, 1H) | 3.91 | 3.51 or 3.47 | 3.62 | 3.90, 3.58 |
| **GlcTFA5** | 4.68 (d, *J* = 8.4 Hz, 1H) | 3.81 | 3.69 | 3.49 | 3.48 | 3.94, 3.79 |
| **GlcNAc5'** | 4.55 (d, *J* = 8.4 Hz, 1H) | 3.71 | 3.55 | 3.45 | 3.44 | 3.91, 3.75 |

**$^{13}$C (150 MHz, $D_2O$): δ (ppm)**

| | C1 | C2 | C3 | C4 | C5 | C6 |
|---|---|---|---|---|---|---|
| **GlcNAc1** | 78.03 | 53.64 | 72.65 | 78.47 | 76.11 | 59.75 |
| **GlcNAc2** | 101.17 | 54.83 | 65.56 | 79.44 | 74.29 | 59.79 |
| **Man3** | 100.36 | 70.06 | 80.34 | 71.89 | 74.19 | 65.72 |
| **Man4** | 99.36 | 76.30 | 69.17 | 67.32 | 73.51 | 61.53 |
| **Man4'** | 96.92 | 76.23 | 69.37 | 67.24 | 72.75 | 61.53 |
| **GlcTFA5** | 98.72 | 55.81 | 72.52 | 69.76 | 75.77 | 60.48 |
| **GlcNAc5'** | 99.51 | 55.23 | 73.29 | 69.79 | 75.70 | 60.51 |

| Signal | Proton | Carbon |
|---|---|---|
| NHC(O)CH3 | -[a] | 174.69, 174.54 |
| NHC(O)CH3 | 2.08 (s, 3H), 2.05 (s, 3H), 1.92 (s, 3H) | 22.23, 22.11, 21.89 |
| NHC(O)CF3 | - | 159.63, 159.38 |
| NHC(O)CF3 | - | 116.70, 114.80 |
| Aromatic | 7.55 – 7.31 (m, 5H) | 136.20, 128.67, 128.28, 127.71 |
| CH2-Ph | 5.17 – 5.09 (m, 2H) | 66.96 |
| NH-COO- | - | 157.71 |
| NH-CH-COOH | 4.41 (dd, $J$ = 9.0, 4.5 Hz, 1H) | 52.16 |
| NH-CH-COOH | - | 176.66 |
| C(O)-CH2-CH | 2.83 (dd, $J$ = 15.7, 4.3 Hz, 1H),<br>2.66 (dd, $J$ = 15.3, 9.1 Hz, 3H) | 38.05 |
| C(O)-CH2-CH | - | 173.28 |

[a] Not applicable

ESI TOF-MS $m/z$ calculated for $C_{62}H_{91}F_3N_6O_{40}$, $[M-2H]^{2-}$: 808.2617, found 808.2579.

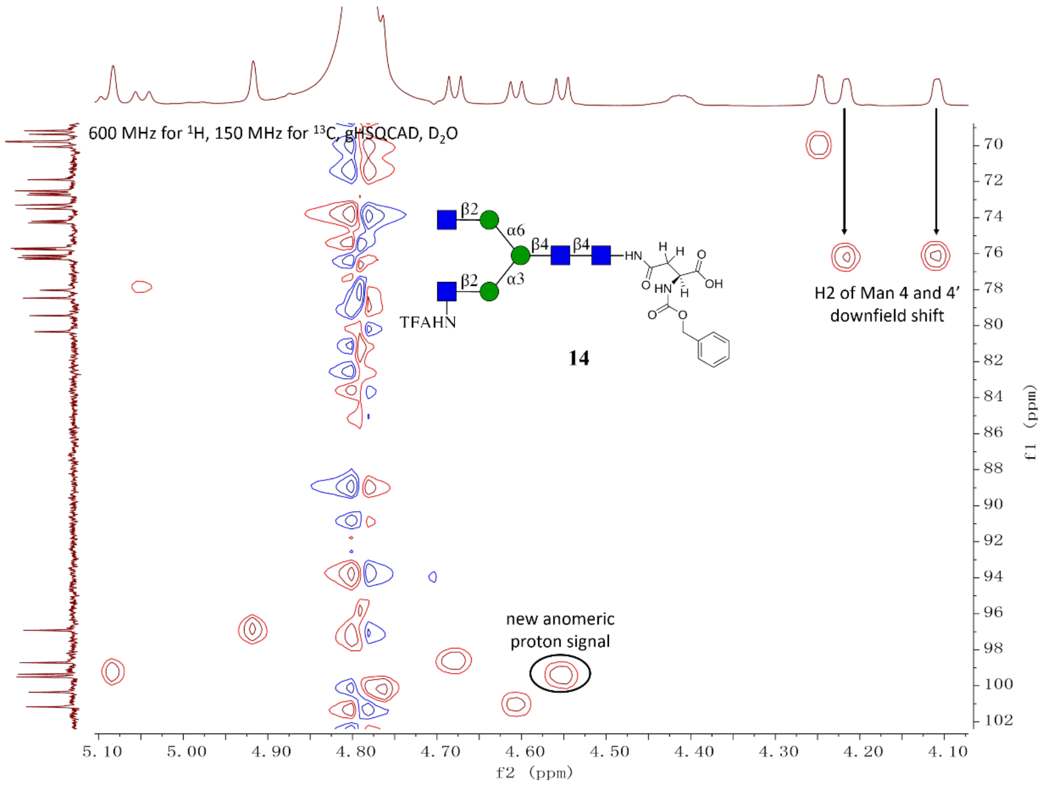


**Figure S8.** Expanded HSQC and HMBC of compound **14**.

NMR and MS analysis of compound **23**

**$^1$H (600 MHz, D$_2$O): δ (ppm)**

| | H1 | H2 | H3 | H4 | H5 | H6 |
|---|---|---|---|---|---|---|
| **GlcNAc1** | 5.04 (d, $J$ = 9.6 Hz, 1H) | 3.82 | 3.73 | 3.65 | 3.54 | N/R[b] |
| **GlcNAc2** | 4.60 (d, $J$ = 8.0 Hz, 1H) | 3.79 | 3.81 | N/R | N/R | N/R |
| **Man3** | 4.76 | 4.23 (d, $J$ = 2.7 Hz, 1H) | 3.80 | N/R | N/R | N/R |
| **Man4** | 5.22 (s, 1H) | 4.33 (d, $J$ = 3.5 Hz, 1H) | 3.97 | N/R | N/R | N/R |
| **Man4'** | 4.92 (s, 1H) | 4.11 (d, $J$ = 3.7 Hz, 1H) | 3.88 | N/R | N/R | N/R |
| **GlcNH$_2$5** | 4.76 | 3.04 (t, $J$ = 9.4 Hz, 1H) | 3.59 | N/R | 3.48 | N/R |
| **GlcNAc5'** | 4.58 (d, $J$ = 8.1 Hz, 1H) | 3.74 | N/R | 3.73 | N/R | N/R |
| **Gal6'** | 4.47 (d, $J$ = 7.8 Hz, 1H) | 3.54 | 3.67 | 3.92 | 3.73 | 3.78, 3.74 |

**$^{13}$C (150 MHz, $D_2O$): δ (ppm)**

| | C1 | C2 | C3 | C4 | C5 | C6 |
|---|---|---|---|---|---|---|
| **GlcNAc1** | 77.98 | 53.62 | 72.69 | 78.41 | 76.08 | N/R |
| **GlcNAc2** | 101.13 | 54.88 | 65.40 | N/R | N/R | N/R |
| **Man3** | 100.26 | 70.08 | 80.70 | N/R | N/R | 65.55 |
| **Man4** | 100.00 | 76.13 | 69.18 | N/R | N/R | N/R |
| **Man4'** | 96.92 | 76.20 | 69.36 | N/R | N/R | N/R |
| **GlcNH$_2$5** | 98.04 | 55.26 | 72.74 | N/R | 76.20 | N/R |
| **GlcNAc5'** | 99.33 | 54.75 | N/R | 78.41 | N/R | N/R |
| **Gal6'** | 102.85 | 70.87 | 72.40 | 68.43 | 75.25 | 60.92 |

| Signal | Proton | Carbon |
|---|---|---|
| **NH<u>C</u>(O)CH$_3$** | -[a] | 174.61, 174.53 |
| **NHC(O)<u>CH$_3$</u>** | 2.07 (s, 1H), 2.04 (s, 1H), 1.91 (s, 1H) | 22.24, 22.12, 21.90 |
| **Aromatic** | 7.51 – 7.34 (m, 5H) | 136.24, 128.66, 128.26, 127.74 |
| **<u>CH$_2$</u>-Ph** | 5.12 (q, $J$ = 12.7, 12.3 Hz, 2H) | 66.87 |
| **NH-<u>C</u>OO-** | - | 157.68 |
| **NH-<u>CH</u>-COOH** | 4.33 | 52.89 |
| **NH-CH-<u>C</u>OOH** | - | N/R |
| **C(O)-<u>CH$_2$</u>-CH** | 2.81 (d, $J$ = 15.2 Hz, 1H)<br>2.58 (dd, $J$ = 15.4, 9.1 Hz, 1H) | 38.47 |
| **<u>C</u>(O)-CH$_2$-CH** | - | 173.61 |

[a] Not applicable

[b] Not reported

ESI TOF-MS *m/z* calculated for $C_{66}H_{102}N_6O_{44}$, $[M-2H]^{2-}$: 841.2970, found 841.3032.

#### Compound 24

**24** was prepared from **23** (10.5 mg) using the general procedures **2.2 d** and **c** for the installation of GlcNAc and Gal moieties. The product was purified using the described two-stage purification system **(2.2 j)** providing **24** as a white fluffy solid (9.1 mg, 71% yield for two steps).

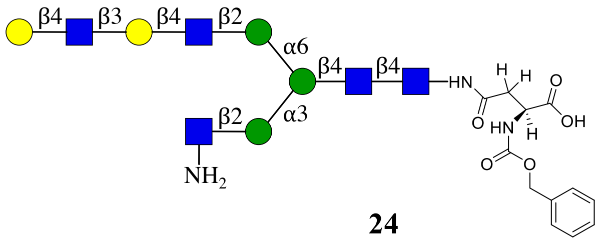


**24**

**$^1$H (600 MHz, $D_2O$): δ (ppm)**

| | H1 | H2 | H3 | H4 | H5 | H6 |
|---|---|---|---|---|---|---|
| **GlcNAc1** | 5.06 (d, *J* = 9.7 Hz, 1H) | 3.83 | 3.74 | 3.66 | 3.56 | N/R[b] |
| **GlcNAc2** | 4.62 (d, *J* = 8.0 Hz, 1H) | 3.81 | 3.82 | 3.75 | N/R | N/R |
| **Man3** | 4.78 | 4.24 (s, 1H) | 3.82 | N/R | N/R | 3.97 3.79 |
| **Man4** | 5.24 (d, *J* = 1.8 Hz, 1H) | 4.38 – 4.35 (m, 1H) | 3.99 | N/R | N/R | N/R |
| **Man4'** | 4.94 (d, *J* = 1.7 Hz, 1H) | 4.12 (d, *J* = 3.0 Hz, 1H) | 3.89 | N/R | N/R | N/R |
| **GlcNH$_2$5** | 4.85 (d, *J* = 8.5 Hz, 1H) | 3.14 | 3.62 | N/R | 3.51 | N/R |
| **GlcNAc5'** | 4.59 (d, *J* = 8.2 Hz, 1H) | 3.76 | N/R | 3.72 | N/R | N/R |
| **Gal6'** | 4.51 – 4.44 (m, 2H) | 3.60 | 3.74 | N/R | N/R | N/R |
| **GlcNAc7'** | 4.71 (d, *J* = 8.4 Hz, 1H) | 3.82 | N/R | N/R | N/R | N/R |
| **Gal8'** | 4.51 – 4.44 (m, 2H) | 3.55 | 3.68 | 3.94 | 3.74 | N/R |

**$^{13}$C (150 MHz, $D_2O$): δ (ppm)**

| | C1 | C2 | C3 | C4 | C5 | C6 |
|---|---|---|---|---|---|---|
| **GlcNAc1** | 78.01 | 53.64 | 72.70 | 78.43 | 76.10 | N/R |
| **GlcNAc2** | 101.14 | 54.89 | 65.45 | 79.32 | N/R | N/R |
| **Man3** | 100.28 | 70.09 | 80.71 | N/R | N/R | 65.57 |
| **Man4** | 99.98 | 76.01 | 69.16 | N/R | N/R | N/R |
| **Man4'** | 96.94 | 76.23 | 69.38 | N/R | N/R | N/R |
| **GlcNH$_2$5** | 97.19 | 55.12 | 72.76 | N/R | 76.27 | N/R |
| **GlcNAc5'** | 99.36 | 54.73 | N/R | 78.49 | N/R | N/R |
| **Gal6'** | 102.90 | 69.87 | 82.02 | N/R | N/R | N/R |
| **GlcNAc7'** | 102.68 | 55.12 | N/R | N/R | N/R | N/R |
| **Gal8'** | 102.78 | 70.89 | 72.43 | 68.47 | 75.25 | N/R |

| Signal | Proton | Carbon |
|---|---|---|
| NHC(O)CH3 | -[a] | 174.81, 174.70, 174.62, 174.54 |
| NHC(O)CH3 | 2.20 – 1.84 (m, 12H) | 22.26, 22.15, 22.12, 21.92 |
| Aromatic | 7.52 – 7.35 (m, 5H) | 136.26, 128.67, 128.27, 127.76 |
| CH2-Ph | 5.18 – 5.08 (m, 2H) | 66.89 |
| NH-COO- | - | 157.68 |
| NH-CH-COOH | 4.34 (dd, $J$ = 9.1, 3.8 Hz, 1H) | 52.94 |
| NH-CH-COOH | - | 177.54 |
| C(O)-CH2-CH | 2.82 (dd, $J$ = 15.4, 4.2 Hz, 1H)<br>2.60 (dd, $J$ = 15.4, 9.4 Hz, 1H) | 38.48 |
| C(O)-CH2-CH | - | 173.61 |

[a] Not applicable

[b] Not reported

ESI TOF-MS *m/z* calculated for $C_{80}H_{125}N_7O_{54}$, $[M-2H]^{2-}$: 1023.8631, found 1023.8719.

#### Compound 25

**25** was prepared from **24** (9.1 mg) using the general procedures **2.2 d** and **c** for the installation of GlcNAc and Gal moieties. The product was purified using the described two-stage purification system **(2.2 j)** providing **25** as a white fluffy solid (9.4 mg, 88% yield for two steps).

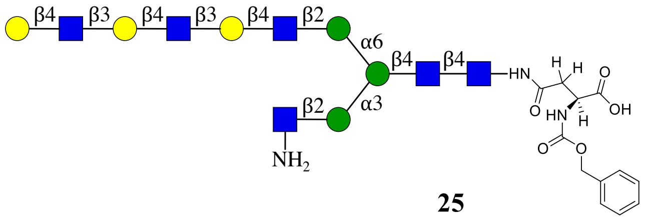


**25**

**$^1$H (600 MHz, D$_2$O): δ (ppm)**

| | H1 | H2 | H3 | H4 | H5 | H6 |
|---|---|---|---|---|---|---|
| GlcNAc1 | 5.08 (d, $J$ = 9.3 Hz, 1H) | 3.86 | 3.77 | 3.68 | 3.58 | N/R[b] |
| GlcNAc2 | 4.65 (d, $J$ = 7.8 Hz, 1H) | 3.81 | 3.84 | 3.77 | N/R | N/R |
| Man3 | 4.80 | 4.27 | 3.83 | N/R | N/R | 4.01<br>3.82 |
| Man4 | 5.27 (s, 1H) | 4.37 | 4.00 | N/R | N/R | N/R |
| Man4' | 4.96 (s, 1H) | 4.14 | 3.92 | N/R | N/R | N/R |
| GlcNH25 | 4.76 | 3.09 – 3.00 (m, 1H) | 3.61 | N/R | 3.51 | N/R |
| GlcNAc5' | 4.61 (d, $J$ = 8.0 Hz, 1H) | 3.78 | N/R | 3.75 | N/R | N/R |
| Gal6' | 4.49 | 3.62 | 3.76 | N/R | N/R | N/R |
| GlcNAc7' | 4.75 – 4.72 (m, 1H) | 3.84 | N/R | N/R | N/R | N/R |
| Gal8' | 4.50 | 3.62 | 3.76 | N/R | N/R | N/R |
| GlcNAc9' | 4.75 – 4.72 (m, 1H) | 3.78 | N/R | N/R | N/R | N/R |
| Gal10' | 4.52 | 3.58 | 3.70 | 3.96 | 3.76 | N/R |

**$^{13}C$ (150 MHz, $D_2O$): δ (ppm)**

| | C1 | C2 | C3 | C4 | C5 | C6 |
|---|---|---|---|---|---|---|
| **GlcNAc1** | 78.06 | 53.68 | 72.73 | 78.50 | 76.13 | N/R |
| **GlcNAc2** | 101.17 | 54.93 | 65.49 | 79.36 | N/R | N/R |
| **Man3** | 100.31 | 70.12 | 80.74 | N/R | N/R | 66.91 |
| **Man4** | 100.03 | 76.24 | 69.26 | N/R | N/R | N/R |
| **Man4'** | 96.99 | 76.28 | 69.42 | N/R | N/R | N/R |
| **GlcNH$_2$5** | 98.40 | 55.39 | 73.03 | N/R | 76.23 | N/R |
| **GlcNAc5'** | 99.41 | 54.76 | N/R | 78.54 | N/R | N/R |
| **Gal6'** | 102.94 | 69.91 | 82.03 | N/R | N/R | N/R |
| **GlcNAc7'** | 102.68 | 55.15 | N/R | N/R | N/R | N/R |
| **Gal8'** | 102.82 | 69.91 | 82.01 | N/R | N/R | N/R |
| **GlcNAc9'** | 102.68 | 55.11 | N/R | N/R | N/R | N/R |
| **Gal10'** | 102.84 | 70.92 | 72.47 | 68.50 | 75.30 | N/R |

| Signal | Proton | Carbon |
|---|---|---|
| **NH<u>C</u>(O)CH$_3$** | -[a] | 174.84, 174.73, 174.65, 174.56 |
| **NHC(O)<u>CH$_3$</u>** | 2.33 – 1.79 (m, 15H) | 22.29, 22.18, 22.14, 21.95 |
| **Aromatic** | 7.64 – 7.30 (m, 5H) | 136.30, 128.71, 128.30, 127.79 |
| **<u>CH$_2$</u>-Ph** | 7.64 – 7.30 (m, 2H) | 66.91 |
| **NH-<u>C</u>OO-** | - | 157.70 |
| **NH-<u>CH</u>-COOH** | 4.36 (s, 2H) | 53.00 |
| **NH-CH-<u>C</u>OOH** | - | 177.57 |
| **C(O)-<u>CH$_2$</u>-CH** | 2.85 (d, $J$ = 14.6 Hz, 1H),<br>2.62 (dd, $J$ = 15.4, 9.5 Hz, 1H) | 38.53 |
| **<u>C</u>(O)-CH$_2$-CH** | - | 173.65 |

[a] Not applicable

[b] Not reported

ESI TOF-MS *m/z* calculated for $C_{94}H_{148}N_8O_{64}$, $[M-2H]^{2-}$: 1206.4292, found 1206.4188.

**Table S1.** The anomeric proton and carbon chemical shift of key intermediate compounds.

| | **Compound 20** | | **Compound21** | | **Compound 22** | |
|---|---|---|---|---|---|---|
| | **H1** | **C1** | **H1** | **C1** | **H1** | **C1** |
| **GlcNAc1** | 5.04 | 78.00 | 5.04 | 78.00 | 5.04 | 78.00 |
| **GlcNAc2** | 4.60 | 101.15 | 4.60 | 101.15 | 4.60 | 101.14 |
| **Man3** | 4.77 | 100.29 | 4.77 | 100.29 | 4.77 | 100.28 |
| **Man4** | 5.11 | 99.45 | 5.11 | 99.45 | 5.11 | 99.45 |
| **Man4'** | 5.03 | 97.44 | 5.04 | 97.44 | 5.03 | 97.43 |
| **GlcNAc5** | 4.58 | 99.32 | 4.58 | 99.34 | 4.58 | 99.33 |
| **GlcNH$_2$5'** | 4.79 | 97.58 | 4.73 | 98.37 | N/R[b] | N/R |
| **Gal6** | 4.47 | 102.83 | 4.45 | 102.86 | 4.45 | 102.87 |
| **GlcNAc7** | --[a] | -- | 4.70 | 102.68 | 4.70 | 102.67 |
| **Gal8** | -- | -- | 4.48 | 102.76 | 4.47 | 102.76 |
| **GlcNAc9** | -- | -- | -- | -- | 4.70 | 102.67 |
| **Gal10** | -- | -- | -- | -- | 4.48 | 102.76 |
| | **Compound 23** | | **Compound24** | | **Compound 25** | |
| | **H1** | **C1** | **H1** | **C1** | **H1** | **C1** |
| **GlcNAc1** | 5.04 | 77.98 | 5.06 | 78.01 | 5.08 | 78.06 |
| **GlcNAc2** | 4.60 | 101.13 | 4.62 | 101.14 | 4.65 | 101.17 |
| **Man3** | 4.76 | 100.26 | 4.78 | 100.28 | 4.80 | 100.31 |
| **Man4** | 5.22 | 100.00 | 5.24 | 99.98 | 5.27 | 100.03 |
| **Man4'** | 4.92 | 96.92 | 4.94 | 96.94 | 4.96 | 96.99 |
| **GlcNH$_2$5** | 4.76 | 98.04 | 4.85 | 97.19 | 4.76 | 98.40 |
| **GlcNAc5'** | 4.58 | 99.33 | 4.59 | 99.36 | 4.61 | 99.41 |
| **Gal6'** | 4.47 | 102.85 | 4.51 – 4.44 | 102.90 | 4.49 | 102.94 |
| **GlcNAc7'** | -- | -- | 4.71 | 102.68 | 4.75 – 4.72 | 102.68 |
| **Gal8'** | -- | -- | 4.51 – 4.44 | 102.78 | 4.50 | 102.82 |
| **GlcNAc9'** | -- | -- | -- | -- | 4.75 – 4.72 | 102.68 |
| **Gal10'** | -- | -- | -- | -- | 4.52 | 102.84 |

[a] Not applicable

[b] Not reported

**Compound 29**

**29** was prepared by acetylation of **20** (1.0 mg) to give intermediate **26** using general procedure **2.2 h** that was purified using protocal **2.2 j**. The CBz protecting group was removed by general procedure **2.2 i** to give the final product **29** as a white fluffy solid (0.9 mg, 92% yield over two steps). ESI TOF-MS *m/z* calculated for $C_{60}H_{98}N_6O_{43}$, $[M-2H]^{2-}$: 795.2839, found 795.2831.

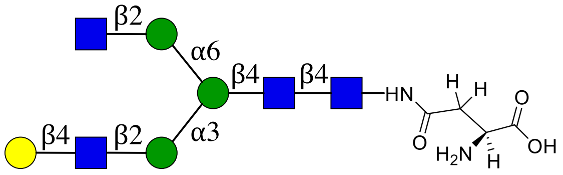


**29**

NMR and MS analysis of compound **26**

**$^1$H (600 MHz, $D_2O$): δ (ppm)**

| | H1 | H2 | H3 | H4 | H5 | H6 |
|---|---|---|---|---|---|---|
| **GlcNAc1** | 5.04 (d, *J* = 9.8 Hz, 1H) | 3.81 | 3.60 | 3.63 | N/R[b] | N/R |
| **GlcNAc2** | 4.59 (d, *J* = 7.8 Hz, 1H) | 3.77 | 3.77 | 3.72 | 3.59 | N/R |
| **Man3** | 4.76 | 4.24 (d, *J* = 2.7 Hz, 1H) | 3.76 | 3.76 | 3.61 | 3.95, 3.76 |
| **Man4** | 5.10 | 4.20 – 4.16 (m, 1H) | 3.88 | 3.48 | 3.73 | N/R |
| **Man4'** | 4.90 (s, 1H) | 4.11 – 4.08 (m, 1H) | 3.88 | 3.48 | N/R | N/R |
| **GlcNAc5** | 4.57 (d, *J* = 7.2 Hz, 1H) | 3.72 | 3.72 | 3.71 | 3.56 | N/R |
| **GlcNAc5'** | 4.54 (d, *J* = 8.5 Hz, 1H) | 3.69 | N/R | N/R | N/R | N/R |
| **Gal6** | 4.45 (d, *J* = 7.8 Hz, 1H) | 3.52 | 3.65 | 3.91 | 3.72 | N/R |

**$^{13}$C (150 MHz, $D_2O$): δ (ppm)**

| | C1 | C2 | C3 | C4 | C5 | C6 |
|---|---|---|---|---|---|---|
| **GlcNAc1** | 78.06 | 53.65 | 72.73 | 78.45 | N/R | N/R |
| **GlcNAc2** | 101.16 | 54.81 | 65.58 | 79.43 | 74.27 | N/R |
| **Man3** | 100.33 | 70.08 | 80.23 | 71.86 | 74.20 | 65.75 |
| **Man4** | 99.44 | 76.28 | 69.35 | 67.23 | 73.44 | N/R |
| **Man4'** | 96.91 | 76.19 | 69.27 | 67.17 | N/R | N/R |
| **GlcNAc5** | 99.34 | 54.76 | 71.84 | 78.35 | 74.62 | N/R |
| **GlcNAc5'** | 99.48 | 55.22 | N/R | N/R | N/R | N/R |
| **Gal6** | 102.81 | 70.85 | 72.38 | 68.43 | 75.24 | N/R |

| Signal | Proton | Carbon |
|---|---|---|
| **NH<u>C</u>(O)CH3** | -[a] | 174.67, 174.60 |
| **NHC(O)<u>CH3</u>** | 2.09 – 1.85 (m, 12H) | 22.23, 22.10, 21.86 |
| **Aromatic** | 7.50 – 7.32 (m, 5H) | 136.12, 128.66, 128.30, 127.64 |
| **<u>CH2</u>-Ph** | 5.16 – 5.08 (m, 2H) | 67.07 |
| **NH-<u>C</u>OO-** | - | 157.75 |
| **NH-<u>CH</u>-COOH** | 4.52 (m, 1H) | 50.99 |
| **NH-CH-<u>C</u>OOH** | - | 177.09 |

| | | |
|---|---|---|
| **C(O)-CH$_2$-CH** | 2.84 (dd, *J* = 14.7, 3.0 Hz, 1H), 2.74 (dd, *J* = 15.9, 7.8 Hz, 1H) | 37.40 |
| **C(O)-CH$_2$-CH** | - | 172.78 |

[a] Not applicable

[b] Not reported

ESI TOF-MS *m/z* calculated for $C_{68}H_{104}N_6O_{45}$, $[M-2H]^{2-}$: 862.3023, found 862.3026.

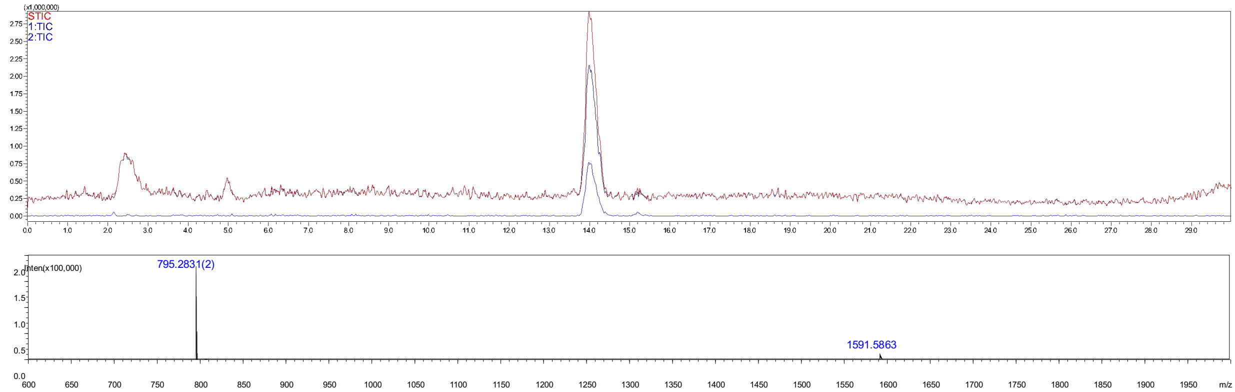


**Figure S9.** Analytical HPLC-MS chromatogram of compound **29**. The retention time = 14.0 min.

#### Compound 30

**30** was prepared by acetylation of **21** (1.0 mg) by general procedures **2.2 h** to give intermediate **27**, which was purified using the combined purification approach **(2.2 j)**. The Cbz protecting group of **27** was removed by hydrogenaton according to **2.2 i** to give final product **30** as a white fluffy solid (0.8 mg, 86% yield over two steps). ESI TOF-MS *m/z* calculated for $C_{74}H_{121}N_7O_{53}$, $[M-2H]^{2-}$: 977.8500, found 977.8472.

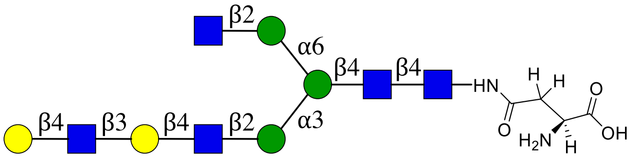


**30**

NMR and MS analysis of compound **27**

**$^1$H (600 MHz, D$_2$O): δ (ppm)**

| | **H1** | **H2** | **H3** | **H4** | **H5** | **H6** |
|---|---|---|---|---|---|---|
| **GlcNAc1** | 5.04 (d, *J* = 9.7 Hz, 1H) | 3.81 | 3.61 | 3.64 | N/R[b] | N/R |
| **GlcNAc2** | 4.60 (d, *J* = 7.9 Hz, 1H) | 3.77 | 3.78 | 3.73 | 3.61 | N/R |
| **Man3** | 4.76 (s, 1H) | 4.25 (s, 1H) | 3.76 | 3.77 | 3.62 | 3.96, 3.77 |
| **Man4** | 5.11 | 4.20 – 4.17 (m, 1H) | 3.88 | 3.49 | 3.74 | N/R |
| **Man4'** | 4.91 (s, 1H) | 4.10 | 3.89 | 3.49 | N/R | N/R |

| | | | | | | |
|---|---|---|---|---|---|---|
| | | (d, *J* = 3.0 Hz, 1H) | | | | |
| **GlcNAc5** | 4.57 (d, *J* = 7.5 Hz, 1H) | 3.73 | 3.72 | 3.71 | 3.57 | N/R |
| **GlcNAc5'** | 4.55 (d, *J* = 8.4 Hz, 1H) | 3.70 | N/R | N/R | N/R | N/R |
| **Gal6** | 4.45 (d, *J* = 7.8 Hz, 1H) | 3.57 | 3.72 | 4.15 (d, *J* = 2.6 Hz, 1H) | N/R | N/R |
| **GlcNAc7** | 4.69 (d, *J* = 8.3 Hz, 1H) | 3.80 | N/R | N/R | N/R | N/R |
| **Gal8** | 4.47 (d, *J* = 7.8 Hz, 1H) | 3.53 | 3.66 | 3.92 | 3.72 | N/R |

**$^{13}$C (150 MHz, $D_2O$): δ (ppm)**

| | **C1** | **C2** | **C3** | **C4** | **C5** | **C6** |
|---|---|---|---|---|---|---|
| **GlcNAc1** | 78.03 | 51.78 | 72.74 | 78.45 | N/R | N/R |
| **GlcNAc2** | 101.16 | 54.82 | 65.58 | 79.42 | 74.28 | N/R |
| **Man3** | 100.33 | 70.10 | 80.34 | 71.88 | 74.20 | 65.73 |
| **Man4** | 99.45 | 76.28 | 69.36 | 67.24 | 73.45 | N/R |
| **Man4'** | 96.92 | 76.20 | 69.28 | 67.18 | N/R | N/R |
| **GlcNAc5** | 99.35 | 54.72 | 71.83 | 78.39 | 74.62 | N/R |
| **GlcNAc5'** | 99.49 | 55.23 | N/R | N/R | N/R | N/R |
| **Gal6** | 102.85 | 69.85 | 81.97 | 68.21 | N/R | N/R |
| **GlcNAc7** | 102.67 | 55.08 | N/R | N/R | N/R | N/R |
| **Gal8** | 102.76 | 70.86 | 72.40 | 68.45 | 75.25 | N/R |

| **Signal** | **Proton** | **Carbon** |
|---|---|---|
| **NH<u>C</u>(O)CH$_3$** | -[a] | 174.80, 174.68, 174.60, 174.54 |
| **NHC(O)<u>CH$_3$</u>** | 2.10 – 1.88 (m, 15H) | 22.23, 22.10, 22.08, 21.87 |
| **Aromatic** | 7.52 – 7.31 (m, 5H) | 136.17, 128.66, 128.29, 127.69 |
| **<u>CH$_2$</u>-Ph** | 5.18 – 5.08 (m, 2H) | 67.00 |
| **NH-<u>C</u>OO-** | - | 157.72 |
| **NH-<u>CH</u>-COOH** | 4.44 | 51.78 |
| **NH-CH-<u>C</u>OOH** | - | N/R |
| **C(O)-<u>CH$_2$</u>-CH** | 2.87 – 2.80 (m, 1H), 2.68 (dd, *J* = 15.4, 8.9 Hz, 1H) | 37.83 |
| **<u>C</u>(O)-CH$_2$-CH** | - | 173.12 |

[a] Not applicable

[b] Not reported

ESI TOF-MS *m/z* calculated for $C_{82}H_{127}N_7O_{55}$, $[M-2H]^{2-}$: 1044.8684, found 1044.8703.

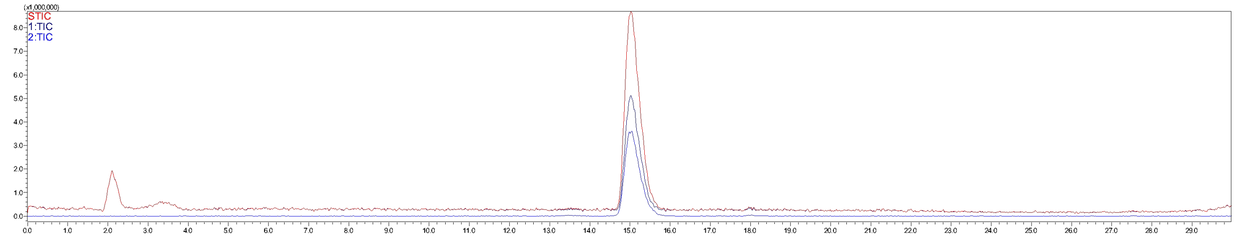

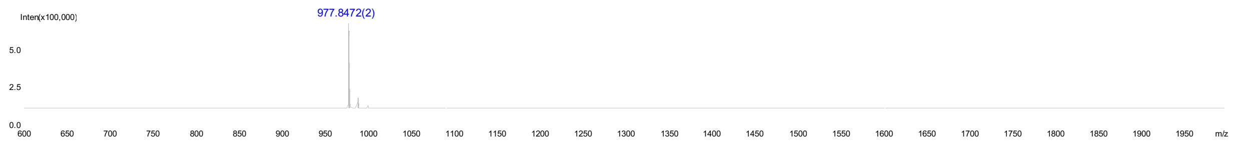


**Figure S10.** Analytical HPLC-MS chromatogram of compound **30**. The retention time = 15.0 min.

#### Compound 31

**31** was prepared by acetylation of **22** (1.0 mg) according to the general procedure **2.2 h** to give, after purified using the described two-stage purification protocal **(2.2 j)**, intermediate **28**. The Cbz protecting group of **28** was removed by hydrogenation (**2.2 i**) to give the final product **31** as a white fluffy solid (1.0 mg, 99% yield over two steps). ESI TOF-MS *m/z* calculated for $C_{88}H_{144}N_8O_{63}$, $[M-2H]^{2-}$: 1160.4161, found 1160.4127.

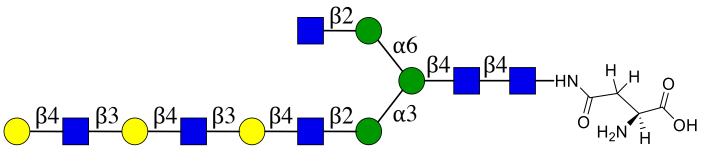


**31**

NMR and MS analysis of compound **28**

**$^{1}$H (600 MHz, $D_2O$): δ (ppm)**

| | H1 | H2 | H3 | H4 | H5 | H6 |
|---|---|---|---|---|---|---|
| **GlcNAc1** | 5.04 (d, *J* = 10.0 Hz, 1H) | 3.81 | 3.61 | 3.64 | N/R[b] | N/R |
| **GlcNAc2** | 4.60 (d, *J* = 7.8 Hz, 1H) | 3.78 | 3.78 | 3.72 | 3.61 | N/R |
| **Man3** | 4.76 | 4.24 (s, 1H) | 3.77 | 3.77 | 3.61 | 3.95, 3.77 |
| **Man4** | 5.18 – 5.08 (m, 3H) | 4.18 (s, 1H) | 3.88 | 3.49 | 3.73 | N/R |
| **Man4'** | 4.91 (s, 1H) | 4.10 (s, 1H) | 3.88 | 3.49 | N/R | N/R |
| **GlcNAc5** | 4.57 (d, *J* = 6.6 Hz, 1H) | 3.72 | 3.72 | 3.71 | 3.57 | N/R |
| **GlcNAc5'** | 4.54 (d, *J* = 8.4 Hz, 1H) | 3.69 | N/R | N/R | N/R | N/R |
| **Gal6** | 4.51 – 4.41 (m, 4H) | 3.57 | 3.71 | 4.15 (s, 2H) | N/R | N/R |
| **GlcNAc7** | 4.69 (d, *J* = 8.4 Hz, 2H) | 3.73 | N/R | N/R | N/R | N/R |
| **Gal8** | 4.51 – 4.41 (m, 4H) | 3.57 | 3.71 | 4.15 (s, 2H) | N/R | N/R |
| **GlcNAc9** | 4.69 (d, *J* = 8.4 Hz, 2H) | 3.79 | N/R | N/R | N/R | N/R |
| **Gal10** | 4.51 – 4.41 (m, 4H) | 3.53 | 3.66 | 3.92 | 3.72 | N/R |

**$^{13}C$ (150 MHz, $D_2O$): δ (ppm)**

| | C1 | C2 | C3 | C4 | C5 | C6 |
|---|---|---|---|---|---|---|
| **GlcNAc1** | 78.03 | 53.65 | 72.74 | 78.45 | N/R | N/R |
| **GlcNAc2** | 101.16 | 54.81 | 65.58 | 79.43 | 74.28 | N/R |
| **Man3** | 100.33 | 70.09 | 80.34 | 71.87 | 74.20 | 65.74 |
| **Man4** | 99.45 | 76.28 | 69.36 | 67.23 | 73.44 | N/R |
| **Man4'** | 96.91 | 76.20 | 69.27 | 67.18 | N/R | N/R |
| **GlcNAc5** | 99.35 | 54.72 | 71.83 | 78.37 | 74.61 | N/R |
| **GlcNAc5'** | 99.49 | 55.22 | N/R | N/R | N/R | N/R |
| **Gal6** | 102.76 | 69.85 | 81.96 | 68.21 | N/R | N/R |
| **GlcNAc7** | 102.66 | 55.09 | N/R | N/R | N/R | N/R |
| **Gal8** | 102.84 | 69.85 | 81.96 | 68.21 | N/R | N/R |
| **GlcNAc9** | 102.66 | 55.04 | N/R | N/R | N/R | N/R |
| **Gal10** | 102.76 | 70.86 | 72.40 | 68.44 | 75.25 | N/R |

| Signal | Proton | Carbon |
|---|---|---|
| **NH<u>C</u>(O)CH₃** | -[a] | 174.80, 174.68, 174,60, 174.54 |
| **NHC(O)<u>CH₃</u>** | 2.10 – 1.87 (m, 18H) | 22.23, 22.08, 21.87 |
| **Aromatic** | 7.52 – 7.32 (m, 5H) | 136.15, 128.66, 128.29, 127.66 |
| **<u>CH₂</u>-Ph** | 5.18 – 5.08 (m, 3H) | 67.03 |
| **NH-<u>C</u>OO-** | - | 157.73 |
| **NH-<u>CH</u>-COOH** | 4.51 – 4.41 (m, 4H) | 51.44 |
| **NH-CH-<u>C</u>OOH** | - | 177.29 |
| **C(O)-<u>CH₂</u>-CH** | 2.84 (dd, $J$ = 15.4, 3.6 Hz, 1H), 2.71 (dd, $J$ = 16.1, 8.5 Hz, 1H) | 37.65 |
| **<u>C</u>(O)-CH₂-CH** | - | 172.96 |

[a] Not applicable

[b] Not reported

ESI TOF-MS *m/z* calculated for $C_{96}H_{150}N_8O_{65}$, $[M-2H]^{2-}$: 1227.4344, found 1227.4352.

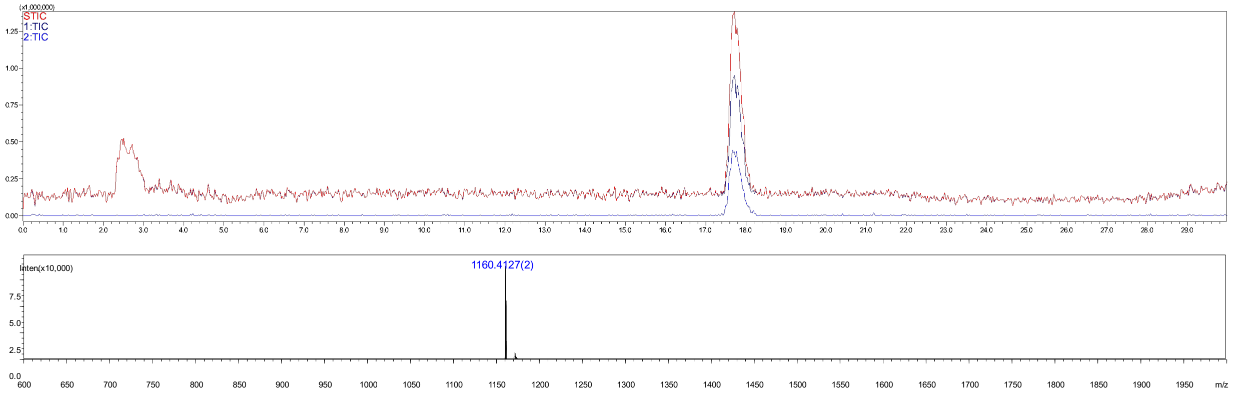

**Figure S11.** Analytical HPLC-MS chromatogram of compound **31**. The retention time = 17.8 min.

#### Compound 35

**35** was synthesized from **20** (3 mg, 1.0 eq) using the general procedures **2.2 f**, **h**, **c** and **i** for the installation of α2,3-sialic acid with ST3Gal4, converting $GlcNH_2$ to GlcNAc with AcOSu, installation of β1,4-Gal with B4GalT1 to give **32**, which was was purified using the described two-stage purification system **(2.2 j)**. The Cbz protecting group of **32** was removed using method **2.2 i** to give final product **35** as a white fluffy solid (1.1 mg, 30% yield over four steps). ESI TOF-MS *m/z* calculated for $C_{77}H_{125}N_7O_{56}$, $[M-2H]^{2-}$: 1021.8580, found 1021.8613.

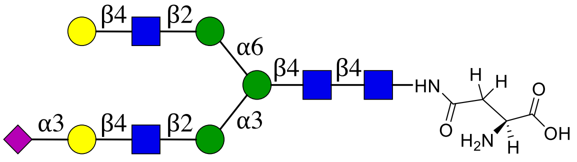

**35**

NMR and MS analysis of compound **32**

**$^1$H (600 MHz, $D_2O$): δ (ppm)**

| | H1 | H2 | H3 | H4 | H5 | H6 |
|---|---|---|---|---|---|---|
| **GlcNAc1** | 5.05 (d, *J* = 9.8 Hz, 1H) | 3.82 | 3.74 | 3.65 | 3.56 | N/R[b] |
| **GlcNAc2** | 4.61 (d, *J* = 8.0 Hz, 1H) | 3.80 | 3.79 | N/R | N/R | N/R |
| **Man3** | 4.77 | 4.25 (d, *J* = 2.7 Hz, 1H) | 3.77 | N/R | N/R | 3.97, 3.80 |
| **Man4** | 5.13 | 4.20 (d, *J* = 3.5 Hz, 1H) | 3.90 | N/R | N/R | N/R |
| **Man4'** | 4.93 (d, *J* = 7.4 Hz, 1H) | 4.15 – 4.08 (m, 2H) | 3.90 | N/R | N/R | N/R |
| **GlcNAc5** | 4.60 – 4.57 (m, 2H) | 3.75 | N/R | N/R | N/R | N/R |
| **GlcNAc5'** | 4.60 – 4.57 (m, 2H) | 3.75 | N/R | N/R | N/R | N/R |
| **Gal6** | 4.48 (d, *J* = 7.8 Hz, 1H) | 3.55 | N/R | 4.15 – 4.08 (m, 2H) | N/R | N/R |
| **Gal6'** | 4.55 (d, *J* = 7.8 Hz, 1H) | 3.57 | N/R | N/R | N/R | N/R |

| | H1 | H2 | H3 | H4 | H5 | H6 | H7 | H8 | H9 |
|---|---|---|---|---|---|---|---|---|---|
| **Neu5Ac7** | -[a] | - | 2.76 (dd, $J$ = 12.5, 4.6 Hz, 1H), 1.81 (t, $J$ = 12.1 Hz, 1H) | 3.70 | 3.85 | N/R | N/R | N/R | N/R |

**$^{13}$C (150 MHz, $D_2O$): δ (ppm)**

| | C1 |
|---|---|
| **GlcNAc1** | 78.07 |
| **GlcNAc2** | 101.18 |
| **Man3** | 100.34 |
| **Man4** | 99.47 |
| **Man4'** | 96.94 |
| **GlcNAc5** | 99.33 |
| **GlcNAc5'** | 99.41 |
| **Gal6** | 102.86 |
| **Gal6'** | 102.50 |

| | C1 | C2 | C3 | C4 | C5 | C6 | C7 | C8 | C9 |
|---|---|---|---|---|---|---|---|---|---|
| **Neu5Ac7** | N/R | N/R | 39.54 | 68.26 | 51.59 | N/R | N/R | N/R | N/R |

| Signal | Proton | Carbon |
|---|---|---|
| **NH<u>C</u>(O)CH3** | - | 174.92, 174.63, 174.55, 173.76 |
| **NHC(O)<u>CH3</u>** | 2.19 – 1.87 (m, 15H) | 22.25, 22.14, 21.95, 21.89 |
| **Aromatic** | 7.52 – 7.34 (m, 5H) | 136.20, 128.68, 128.29, 127.71 |
| **<u>CH2</u>-Ph** | 5.14 (q, $J$ = 11.9 Hz, 2H) | 66.99 |
| **NH-<u>C</u>OO-** | - | 157.73 |
| **NH-<u>CH</u>-COOH** | 4.44 (dd, $J$ = 8.7, 4.6 Hz, 1H) | 51.95 |
| **NH-CH-<u>C</u>OOH** | - | N/R |
| **C(O)-<u>CH2</u>-CH** | 2.84 (dd, $J$ = 15.6, 4.4 Hz, 1H), 2.68 (dd, $J$ = 15.7, 8.8 Hz, 1H) | 37.93 |
| **<u>C</u>(O)-CH2-CH** | - | 173.20 |

[a] Not applicable

[b] Not reported

ESI TOF-MS $m/z$ calculated for $C_{85}H_{131}N_7O_{58}$, $[M-2H]^{2-}$: 1088.8764, found 1088.8828.

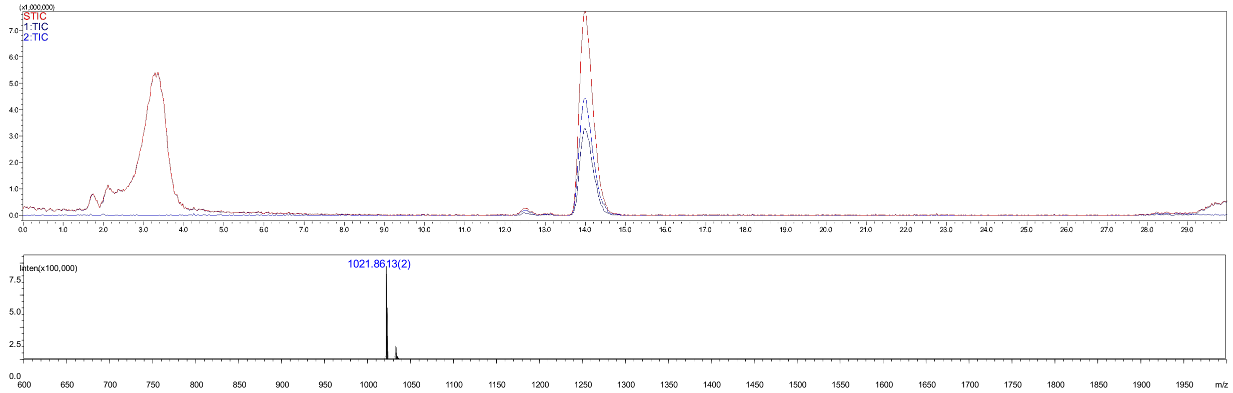

**Figure S12.** Analytical HPLC-MS chromatogram of compound **35**. The retention time = 14.0 min.

#### Compound 36

**36** was synthesized from **21** (3.0 mg) using the general procedures **2.2 f**, **h**, **c** for the installation of α2,3-sialic acid with ST3Gal4, converting $GlcNH_2$ to GlcNAc with AcOSu, installation of β1,4-Gal with B4GalT1 to give product **33**, which was purified using the described two-stage purification system **(2.2 j)**. The Cbz protecting group of **33** was removed by hydrogenation (**2.2 i**) to give final product **36** as a white fluffy solid (1.4 mg, 41% yield for four steps). ESI TOF-MS *m/z* calculated for $C_{91}H_{148}N_8O_{66}$, $[M-2H]^{2-}$: 1204.4241, found 1204.4245.

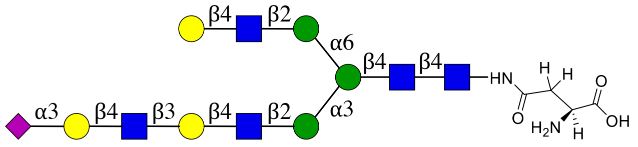

**36**

NMR and MS analysis of compound **33**:

**$^1$H (600 MHz, $D_2O$): δ (ppm)**

| | H1 | H2 | H3 | H4 | H5 | H6 |
|---|---|---|---|---|---|---|
| **GlcNAc1** | 4.99 (d, *J* = 9.7 Hz, 1H) | 3.76 | N/R[b] | N/R | N/R | N/R |
| **GlcNAc2** | 4.54 (d, *J* = 8.0 Hz, 1H) | 3.73 | 3.72 | N/R | N/R | N/R |
| **Man3** | 4.71 (d, *J* = 6.4 Hz, 1H) | 4.19 (d, *J* = 2.6 Hz, 1H) | 3.71 | N/R | N/R | 3.90, 3.73 |
| **Man4** | 5.05 | 4.13 (d, *J* = 3.5 Hz, 1H) | 3.83 | N/R | N/R | N/R |
| **Man4'** | 4.87 (s, 1H) | 4.05 | 3.89 | N/R | N/R | N/R |
| **GlcNAc5** | 4.53 – 4.47 (m, 3H) | 3.68 | N/R | N/R | N/R | N/R |
| **GlcNAc5'** | 4.53 – 4.47 (m, 3H) | 3.68 | N/R | N/R | N/R | N/R |
| **Gal6** | 4.44 – 4.36 (m, 3H) | 3.47 | 3.66 | 4.10 | N/R | N/R |
| **Gal6'** | 4.53 – 4.47 (m, 3H) | 3.51 | N/R | N/R | N/R | N/R |
| **GlcNAc7** | 4.62 (d, *J* = 8.3 Hz, 1H) | 4.13 (d, *J* = 3.5 Hz, 1H) | N/R | N/R | N/R | N/R |
| **Gal8** | 4.44 – 4.36 (m, 3H) | 3.51 | N/R | 4.06 | N/R | N/R |

|  | H1 | H2 | H3 | H4 | H5 | H6 | H7 | H8 | H9 |
|---|---|---|---|---|---|---|---|---|---|
| **Neu5Ac9** | -[a] | - | 2.69 (dd, *J* = 12.5, 4.6 Hz, 1H), 1.74 (t, *J* = 12.2 Hz, 1H) | 3.62 | 3.79 | N/R | N/R | N/R | N/R |

**$^{13}C$ (150 MHz, $D_2O$): δ (ppm)**

|  | C1 |
|---|---|
| **GlcNAc1** | 78.01 |
| **GlcNAc2** | 101.09 |
| **Man3** | 100.28 |
| **Man4** | 99.40 |
| **Man4'** | 96.84 |
| **GlcNAc5** | 99.26 |
| **GlcNAc5'** | 99.26 |
| **Gal6** | 102.75 |
| **Gal6'** | 102.33 |
| **GlcNAc7** | 102.71 |
| **Gal8** | 102.75 |

|  | C1 | C2 | C3 | C4 | C5 | C6 | C7 | C8 | C9 |
|---|---|---|---|---|---|---|---|---|---|
| **Neu5Ac9** | N/R | N/R | 39.44 | 68.21 | 51.48 | N/R | N/R | N/R | N/R |

| Signal | Proton | Carbon |
|---|---|---|
| **NH<u>C</u>(O)$CH_3$** | - | 174.81, 174.73, 174.57, 174.48 |
| **NHC(O)<u>$CH_3$</u>** | 2.12 – 1.79 (m, 18H) | 22.16, 22.04, 21.99, 21.86, 21.79 |
| **Aromatic** | 7.49 – 7.25 (m, 5H) | 136.08, 128.59, 128.22, 127.60 |
| **<u>$CH_2$</u>-Ph** | 5.11 – 5.02 (m, 2H) | 66.94 |
| **NH-<u>C</u>OO-** | - | 157.69 |
| **NH-<u>CH</u>-COOH** | 4.44 – 4.36 (m, 3H) | 51.54 |
| **NH-CH-<u>C</u>OOH** | - | N/R |
| **C(O)-<u>$CH_2$</u>-CH** | 2.81 – 2.75 (m, 1H), 2.65 – 2.60 (m, 1H) | 37.66 |
| **<u>C</u>(O)-$CH_2$-CH** | - | 173.69 |

[a] Not applicable

[b] Not reported

ESI TOF-MS *m/z* calculated for $C_{99}H_{154}N_8O_{68}$, $[M-2H]^{2-}$: 1271.4425, found 1271.4403.

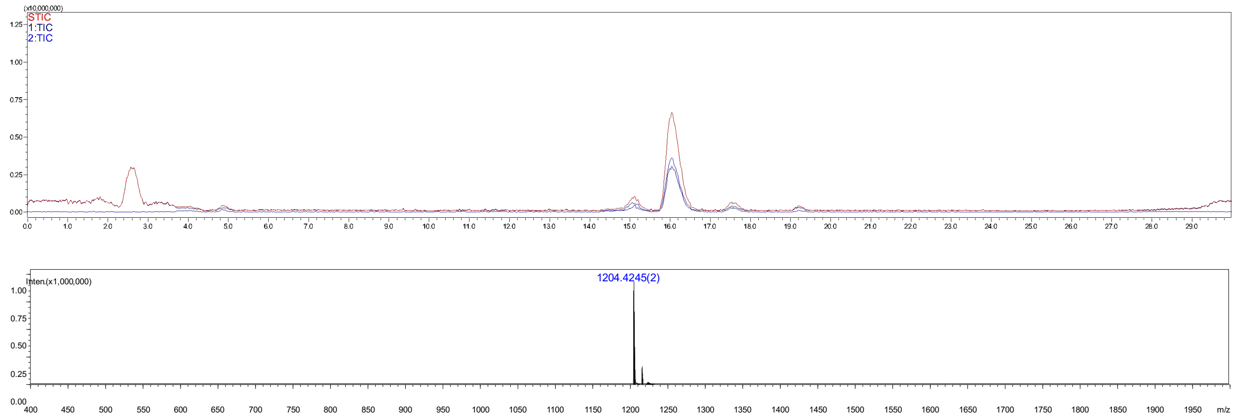

**Figure S13.** Analytical HPLC-MS chromatogram of compound **36**. The retention time = 16.0 min.

#### Compound 37

**37** was synthesized from **22** (3 mg, 1.0 eq) using the general procedures **2.2 f**, **h**, **c** for the installation of α2,3-sialic acid with ST3Gal4, converting $GlcNH_2$ to GlcNAc with AcOSu, installation of β1,4-Gal with B4GalT1 to give intermediate product **34**, which was purified using the described two-stage purification system **(2.2 j)**. The Cbz group protecting group of **34** was removed by hydrogenation to give final product **37** as a white fluffy solid (2.3 mg, 66% yield over four steps). ESI TOF-MS *m/z* calculated for $C_{105}H_{171}N_9O_{76}$, $[M-2H]^{2-}$: 1386.9902, found 1386.9860.

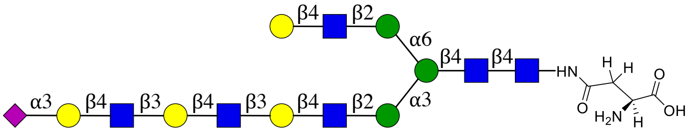

**37**

NMR and MS analysis of compound **34**

$^1$H (600 MHz, $D_2O$): δ (ppm)

| | H1 | H2 | H3 | H4 | H5 | H6 |
|---|---|---|---|---|---|---|
| GlcNAc1 | 5.05 (d, *J* = 9.7 Hz, 1H) | 3.82 | 3.75 | 3.65 | 3.56 | N/R[b] |
| GlcNAc2 | 4.61 (d, *J* = 7.9 Hz, 1H) | 3.79 | 3.80 | N/R | N/R | N/R |
| Man3 | 4.47 | 4.25 (d, *J* = 2.8 Hz, 1H) | 3.77 | N/R | N/R | 3.97, 3.80 |
| Man4 | 5.12 | 4.19 (d, *J* = 3.5 Hz, 1H) | 3.91 | N/R | N/R | N/R |
| Man4' | 4.93 (s, 1H) | 4.14 – 4.08 (m, 2H) | 3.89 | N/R | N/R | N/R |
| GlcNAc5 | 4.60 – 4.53 (m, 3H) | 3.75 | N/R | N/R | N/R | N/R |
| GlcNAc5' | 4.60 – 4.53 (m, 3H) | 3.75 | N/R | N/R | N/R | N/R |
| Gal6 | 4.51 – 4.41 (m, 4H) | 3.58 | 3.73 | 4.16 (s, 2H) | N/R | N/R |
| Gal6' | 4.60 – 4.53 (m, 3H) | 3.75 | N/R | N/R | N/R | N/R |
| GlcNAc7 | 4.70 (d, *J* = 8.4 Hz, 1H) | 3.81 | N/R | N/R | N/R | N/R |
| Gal8 | 4.51 – 4.41 (m, 4H) | 3.59 | 3.73 | 4.16 (s, 2H) | N/R | N/R |

| | | | | | | |
|---|---|---|---|---|---|---|
| **GlcNAc9** | 4.70 (d, $J$ = 8.4 Hz, 1H) | 3.81 | N/R | N/R | N/R | N/R |
| **Gal11** | 4.51 – 4.41 (m, 4H) | 3.55 | N/R | 4.14 – 4.08 (m, 2H) | N/R | N/R |

| | **H1** | **H2** | **H3** | **H4** | **H5** | **H6** | **H7** | **H8** | **H9** |
|---|---|---|---|---|---|---|---|---|---|
| **Neu5Ac9** | -[a] | - | 2.76 (dd, $J$ = 12.5, 4.6 Hz, 1H), 1.80 (t, $J$ = 12.1 Hz, 1H) | 3.70 | 3.86 | N/R | N/R | N/R | N/R |

**$^{13}$C (150 MHz, $D_2O$): δ (ppm)**

| | **C1** |
|---|---|
| **GlcNAc1** | 78.05 |
| **GlcNAc2** | 101.18 |
| **Man3** | 100.33 |
| **Man4** | 99.47 |
| **Man4'** | 96.95 |
| **GlcNAc5** | 99.35 |
| **GlcNAc5'** | 99.35 |
| **Gal6** | 102.86 |
| **Gal6'** | 102.46 |
| **GlcNAc7** | 102.68 |
| **Gal8** | 102.86 |
| **GlcNAc9** | 102.71 |
| **Gal10** | 102.79 |

| | **C1** | **C2** | **C3** | **C4** | **C5** | **C6** | **C7** | **C8** | **C9** |
|---|---|---|---|---|---|---|---|---|---|
| **Neu5Ac11** | N/R | N/R | 39.55 | 68.25 | 51.60 | N/R | N/R | N/R | N/R |

| **Signal** | **Proton** | **Carbon** |
|---|---|---|
| **NH<u>C</u>(O)CH3** | - | 174.92, 174.81, 174.72, 174.62, 174.54, 173.78 |
| **NHC(O)<u>CH3</u>** | 2.25 – 1.86 (m, 21H) | 22.26, 22.14, 22.10, 21.96, 21.89 |
| **Aromatic** | 7.52 – 7.32 (m, 5H) | 136.20, 128.68, 128.29, 127.71 |
| **<u>CH2</u>-Ph** | 5.18 – 5.09 (m, 2H) | 66.99 |
| **NH-<u>C</u>OO-** | - | 157.73 |
| **NH-<u>CH</u>-COOH** | 4.51 – 4.41 (m, 4H) | 51.95 |
| **NH-CH-<u>C</u>OOH** | - | N/R |
| **C(O)-<u>CH2</u>-CH** | 2.84 (dd, $J$ = 15.6, 4.3 Hz, 1H), 2.68 (dd, $J$ = 15.3, 8.6 Hz, 1H) | 37.94 |
| **<u>C</u>(O)-CH2-CH** | - | 173.19 |

[a] Not applicable

[b] Not reported

ESI TOF-MS *m/z* calculated for $C_{113}H_{177}N_9O_{78}$, $[M-2H]^{2-}$: 1454.0086, found 1453.9947.

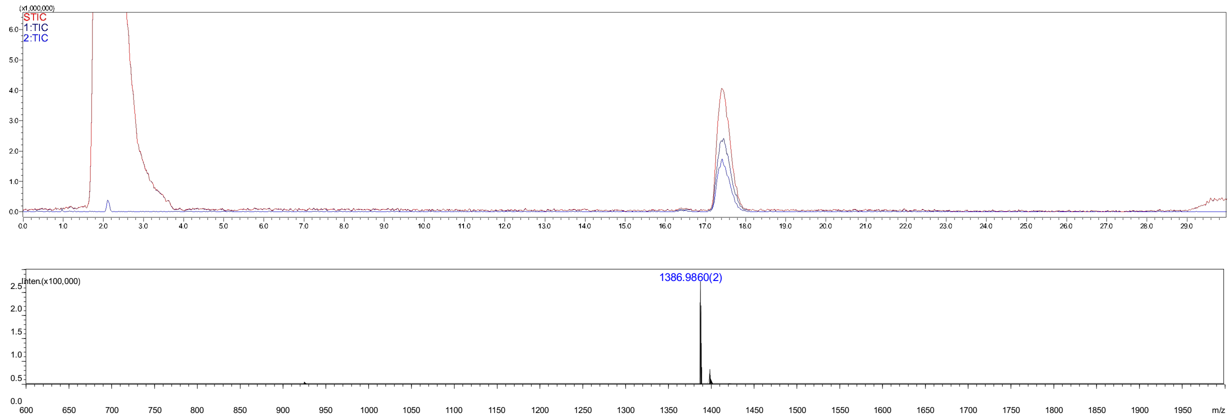

**Figure S14.** Analytical HPLC-MS chromatogram of compound **37**. The retention time = 17.5 min.

#### Compound 44

**44** was synthesized from **20** (3.0 mg, 1.0 eq) using the general procedures **2.2 e**, **h**, **c** for the installation of α2,6-sialic acid with ST6Gal1, converting $GlcNH_2$ to GlcNAc with AcOSu, installation of β1,4-Gal with B4GalT1 to give intermediate **41** which was purified using the described two-stage purification system **(2.2 j)**. The Cbz protecrting group of **41** was removed by hydrogentation (**2.2 i**) to give final product **44** as a white fluffy solid (1.9 mg, 52% yield over four steps). ESI TOF-MS *m/z* calculated for $C_{77}H_{125}N_7O_{56}$, $[M-2H]^{2-}$: 1021.8580, found 1021.8526.

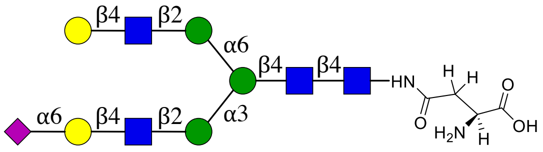

**44**

NMR and MS analysis of compound **41:**

**$^1$H (600 MHz, $D_2O$): δ (ppm)**

| | H1 | H2 | H3 | H4 | H5 | H6 |
|---|---|---|---|---|---|---|
| **GlcNAc1** | 5.02 (d, *J* = 9.9 Hz, 1H) | 3.79 | 3.70 | 3.61 | 3.52 | N/R[b] |
| **GlcNAc2** | 4.61 – 4.52 (m, 3H) | 3.74 | 3.76 | N/R | N/R | N/R |
| **Man3** | 4.74 | 4.23 (s, 1H) | 3.75 | N/R | N/R | 3.93, 3.77 |
| **Man4** | 5.11 | 4.17 (s, 1H) | 3.87 | N/R | N/R | N/R |
| **Man4'** | 4.90 (s, 1H) | 4.11 – 4.06 (m, 1H) | 3.87 | N/R | N/R | N/R |
| **GlcNAc5** | 4.61 – 4.52 (m, 3H) | 3.74 | N/R | N/R | N/R | N/R |
| **GlcNAc5'** | 4.61 – 4.52 (m, 3H) | 3.74 | N/R | N/R | N/R | N/R |
| **Gal6** | 4.42 (dd, *J* = 7.8, 1.7 Hz, 1H) | 3.51 | N/R | N/R | N/R | N/R |
| **Gal6'** | 4.44 (d, *J* = 7.8 Hz, 1H) | 3.51 | N/R | N/R | N/R | N/R |

| | H1 | H2 | H3 | H4 | H5 | H6 | H7 | H8 | H9 |
|---|---|---|---|---|---|---|---|---|---|
| **Neu5Ac7** | -[a] | - | 2.64 (dd, *J* = 12.4, 4.6 Hz, 1H), 1.69 (t, *J* = 12.2 Hz, 1H) | 3.62 | 3.78 | N/R | N/R | N/R | N/R |

**$^{13}C$ (150 MHz, $D_2O$): δ (ppm)**

| | C1 |
|---|---|
| **GlcNAc1** | 78.03 |
| **GlcNAc2** | 101.18 |
| **Man3** | 100.32 |
| **Man4** | 99.46 |
| **Man4'** | 96.94 |
| **GlcNAc5** | 99.23 |
| **GlcNAc5'** | 99.34 |
| **Gal6** | 103.45 |
| **Gal6'** | 102.85 |

| | C1 | C2 | C3 | C4 | C5 | C6 | C7 | C8 | C9 |
|---|---|---|---|---|---|---|---|---|---|
| **Neu5Ac7** | N/R | 100.03 | 39.96 | 68.13 | 51.79 | N/R | N/R | N/R | N/R |

| Signal | Proton | Carbon |
|---|---|---|
| **NH<u>C</u>(O)$CH_3$** | - | 174.81, 174.69, 174.63, 174.53 |
| **NHC(O)<u>$CH_3$</u>** | 2.26 – 1.77 (m, 15H) | 22.34, 22.25, 22.13, 21.95, 21.89 |
| **Aromatic** | 7.52 – 7.27 (m, 5H) | 136.22, 128.66, 128.27, 127.73 |
| **<u>$CH_2$</u>-Ph** | 5.10 (d, *J* = 13.8 Hz, 2H) | 66.92 |
| **NH-<u>C</u>OO-** | - | 157.70 |
| **NH-<u>CH</u>-COOH** | 4.34 (dd, *J* = 9.1, 4.2 Hz, 1H) | 52.50 |
| **NH-CH-<u>C</u>OOH** | - | N/R |
| **C(O)-<u>$CH_2$</u>-CH** | 2.82 – 2.76 (m, 1H), 2.61 – 2.56 (m, 1H) | 38.23 |
| **<u>C</u>(O)-$CH_2$-CH** | - | 173.43 |

[a] Not applicable

[b] Not reported

ESI TOF-MS *m/z* calculated for $C_{85}H_{131}N_7O_{58}$, $[M-2H]^{2-}$: 1088.8764, found 1088.8744.

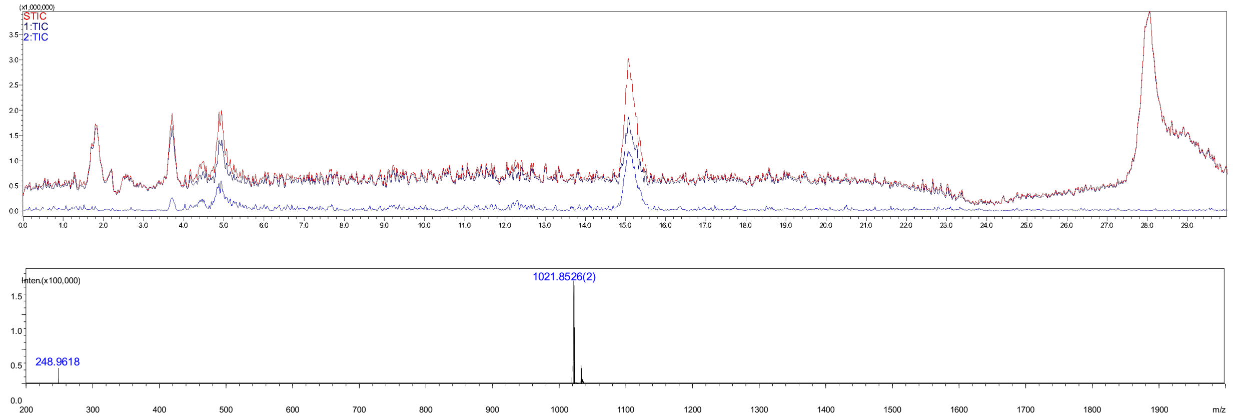

**Figure S15.** Analytical HPLC-MS chromatogram of compound **44**. The retention time = 15.2 min.

#### Compound 45

**45** was synthesized from **21** (3 mg) using the general procedures **2.2 e**, **h**, **c** and **i** for the installation of α2,6-sialic acid with ST6Gal1, converting $GlcNH_2$ to GlcNAc with AcOSu, installation of β1,4-Gal with B4GalT1 to give compound **42** which was purified using the described two-stage purification system **(2.2 j)**. The Cbz protecrting group of **42** was removed by hydrogentation (**2.2 i**) to give final product **45** as a white fluffy solid (2.2 mg, 61% yield over four steps). ESI TOF-MS *m/z* calculated for $C_{91}H_{148}N_8O_{66}$, $[M-2H]^{2-}$: 1204.4241, found 1204.4341.

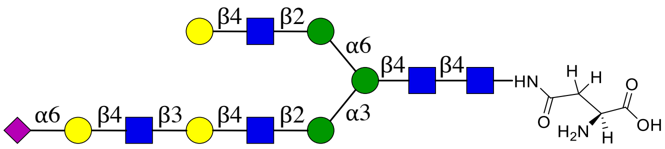

**45**

NMR and MS analysis of compound **42**

**$^1H$ (600 MHz, $D_2O$): δ (ppm)**

| | H1 | H2 | H3 | H4 | H5 | H6 |
|---|---|---|---|---|---|---|
| **GlcNAc1** | 5.05 (d, *J* = 9.9 Hz, 1H) | 3.82 | 3.73 | 3.65 | 3.55 | N/R[b] |
| **GlcNAc2** | 4.64 – 4.53 (m, 3H) | 3.79 | 3.79 | N/R | N/R | N/R |
| **Man3** | 4.76 | 4.25 (d, *J* = 2.7 Hz, 1H) | 3.78 | N/R | N/R | 3.96, 3.79 |
| **Man4** | 5.12 | 4.19 (d, *J* = 3.3 Hz, 1H) | 3.91 | N/R | N/R | N/R |
| **Man4'** | 4.93 (s, 1H) | 4.13 – 4.09 (m, 1H) | 3.90 | N/R | N/R | N/R |
| **GlcNAc5** | 4.64 – 4.53 (m, 3H) | 3.74 | N/R | N/R | N/R | N/R |
| **GlcNAc5'** | 4.64 – 4.53 (m, 3H) | 3.74 | N/R | N/R | N/R | N/R |
| **Gal6** | 4.51 – 4.42 (m, 3H) | 3.56 | 3.73 | 4.16 (d, *J* = 3.1 Hz, 1H) | N/R | N/R |

| | | | | | | |
|---|---|---|---|---|---|---|
| **Gal6'** | 4.51 – 4.42 (m, 3H) | 3.56 | N/R | N/R | N/R | N/R |
| **GlcNAc7** | 4.73 (d, *J* = 7.7 Hz, 1H) | 3.80 | N/R | N/R | N/R | N/R |
| **Gal8** | 4.51 – 4.42 (m, 3H) | 3.56 | N/R | N/R | N/R | N/R |

| | **H1** | **H2** | **H3** | **H4** | **H5** | **H6** | **H7** | **H8** | **H9** |
|---|---|---|---|---|---|---|---|---|---|
| **Neu5Ac9** | -[a] | - | 2.67 (dd, *J* = 12.5, 4.6 Hz, 1H), 1.72 (t, *J* = 12.1 Hz, 1H) | 3.65 | 3.81 | N/R | N/R | N/R | N/R |

**$^{13}$C (150 MHz, $D_2O$): δ (ppm)**

| | **C1** |
|---|---|
| **GlcNAc1** | 78.01 |
| **GlcNAc2** | 101.17 |
| **Man3** | 100.34 |
| **Man4** | 99.47 |
| **Man4'** | 96.94 |
| **GlcNAc5** | 99.38 |
| **GlcNAc5'** | 99.38 |
| **Gal6** | 102.86 |
| **Gal6'** | 102.86 |
| **GlcNAc7** | 102.51 |
| **Gal8** | 103.38 |

| | **C1** | **C2** | **C3** | **C4** | **C5** | **C6** | **C7** | **C8** | **C9** |
|---|---|---|---|---|---|---|---|---|---|
| **Neu5Ac9** | N/R | N/R | 40.00 | 68.14 | 51.79 | N/R | N/R | N/R | N/R |

| **Signal** | **Proton** | **Carbon** |
|---|---|---|
| **$NH\underline{C}(O)CH_3$** | - | 174.82, 174.65, 174.54 |
| **$NHC(O)\underline{CH_3}$** | 2.36 – 1.79 (m, 18H) | 22.25, 22.20, 22.14, 21.94, 21.90 |
| **Aromatic** | 7.55 – 7.31 (m, 5H) | 136.24, 128.67, 128.27, 127.75 |
| **$\underline{CH_2}$-Ph** | 5.18 – 5.07 (m, 2H) | 66.87 |
| **NH-$\underline{C}$OO-** | - | N/R |
| **NH-$\underline{CH}$-COOH** | 4.34 (dd, *J* = 9.5, 3.9 Hz, 1H) | 52.76 |
| **NH-CH-$\underline{C}$OOH** | - | N/R |
| **C(O)-$\underline{CH_2}$-CH** | 2.84 – 2.78 (m, 1H), 2.60 (dd, *J* = 15.4, 9.3 Hz, 1H) | 38.40 |
| **$\underline{C}$(O)-$CH_2$-CH** | - | N/R |

[a] Not applicable

[b] Not reported

ESI TOF-MS *m/z* calculated for $C_{99}H_{154}N_8O_{68}$, $[M-2H]^{2-}$: 1271.4425, found 1271.4336.

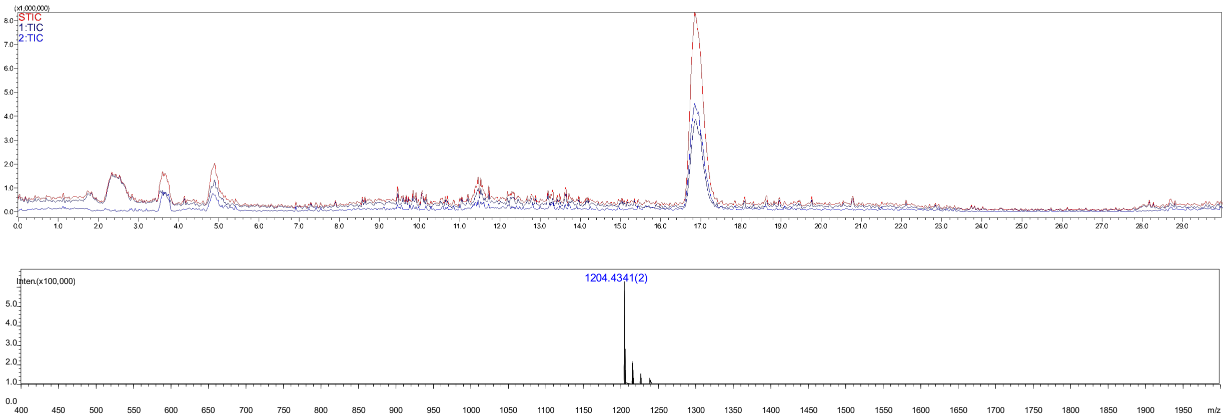

**Figure S16.** Analytical HPLC-MS chromatogram of compound **45**. The retention time = 17.0 min.

#### Compound 46

**46** was synthesized from **22** (3 mg, 1.0 eq) using the general procedures **2.2 e**, **h**, **c** for the installation of α2,6-sialic acid with ST6Gal1, converting $GlcNH_2$ to GlcNAc with AcOSu, installation of β1,4-Gal with B4GalT1 to give intermediate product **43** which was purified using the described two-stage purification system **(2.2 j)**. The Cbz protecrting group of **43** was removed by hydrogentation (**2.2 i**) to give final product **46** as a white fluffy solid (1.7 mg, 50% yield over four steps). ESI TOF-MS *m/z* calculated for $C_{125}H_{171}N_9O_{76}$, $[M-2H]^{2-}$: 1386.9902, found 1386.9802.

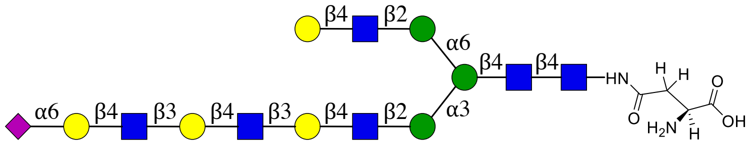

**46**

NMR and MS analysis of compound **43**

**$^1H$ (600 MHz, $D_2O$): δ (ppm)**

| | H1 | H2 | H3 | H4 | H5 | H6 |
|---|---|---|---|---|---|---|
| **GlcNAc1** | 5.04 (d, *J* = 9.6 Hz, 1H) | 3.81 | 3.72 | 3.64 | 3.54 | N/R[b] |
| **GlcNAc2** | 4.64 – 4.53 (m, 3H) | 3.78 | 3.79 | N/R | N/R | N/R |
| **Man3** | 4.76 | 4.24 (d, *J* = 2.8 Hz, 1H) | 3.76 | N/R | N/R | 3.96, 3.78 |
| **Man4** | 5.11 | 4.18 (d, *J* = 3.4 Hz, 1H) | 3.89 | N/R | N/R | N/R |
| **Man4'** | 4.92 (s, 1H) | 4.12 – 4.09 (m, 1H) | 3.88 | N/R | N/R | N/R |
| **GlcNAc5** | 4.64 – 4.53 (m, 3H) | 3.74 | N/R | N/R | N/R | N/R |
| **GlcNAc5'** | 4.64 – 4.53 (m, 3H) | 3.74 | N/R | N/R | N/R | N/R |
| **Gal6** | 4.51 – 4.40 (m, 4H) | 3.54 | 3.72 | 4.15 (d, *J* = 3.0 Hz, 2H) | N/R | N/R |

| | | | | | | |
|---|---|---|---|---|---|---|
| **Gal6'** | 4.51 – 4.40 (m, 4H) | 3.54 | N/R | N/R | N/R | N/R |
| **GlcNAc7** | 4.72 (d, *J* = 7.6 Hz, 1H) | 3.79 | N/R | N/R | N/R | N/R |
| **Gal8** | 4.51 – 4.40 (m, 4H) | 3.53 | 3.72 | 4.15 (d, *J* = 3.0 Hz, 2H) | N/R | N/R |
| **GlcNAc9** | 4.69 (d, *J* = 8.3 Hz, 1H) | 3.79 | N/R | N/R | N/R | N/R |
| **Gal10** | 4.51 – 4.40 (m, 4H) | 3.58 | N/R | N/R | N/R | N/R |

| | **H1** | **H2** | **H3** | **H4** | **H5** | **H6** | **H7** | **H8** | **H9** |
|---|---|---|---|---|---|---|---|---|---|
| **Neu5Ac11** | -[a] | - | 2.66 (dd, *J* = 12.5, 4.5 Hz, 1H), 1.71 (t, *J* = 12.1 Hz, 1H) | 3.64 | 3.80 | N/R | N/R | N/R | N/R |

**$^{13}$C (150 MHz, $D_2O$): δ (ppm)**

| | **C1** |
|---|---|
| **GlcNAc1** | 78.03 |
| **GlcNAc2** | 101.21 |
| **Man3** | 100.31 |
| **Man4** | 99.46 |
| **Man4'** | 96.91 |
| **GlcNAc5** | 99.40 |
| **GlcNAc5'** | 99.40 |
| **Gal6** | 102.83 |
| **Gal6'** | 102.83 |
| **GlcNAc7** | 102.56 |
| **Gal8** | 99.40 |
| **GlcNAc9** | 102.71 |
| **Gal10** | 103.36 |

| | **C1** | **C2** | **C3** | **C4** | **C5** | **C6** | **C7** | **C8** | **C9** |
|---|---|---|---|---|---|---|---|---|---|
| **Neu5Ac10** | N/R | N/R | 39.99 | 68.16 | 51.77 | N/R | N/R | N/R | N/R |

| **Signal** | **Proton** | **Carbon** |
|---|---|---|
| **NH<u>C</u>(O)CH₃** | - | 174.81, 174.62, 174.53 |
| **NHC(O)<u>CH₃</u>** | 2.42 – 1.77 (m, 21H) | 22.24, 22.19, 22.13, 22.08, 21.94, 21.90 |
| **Aromatic** | 7.51 – 7.33 (m, 5H) | 136.23, 128.66, 128.26, 127.74 |
| **<u>CH₂</u>-Ph** | 5.16 – 5.06 (m, 2H) | 66.89 |
| **NH-<u>C</u>OO-** | - | N/R |
| **NH-<u>CH</u>-COOH** | 4.32 (dd, *J* = 9.6, 4.2 Hz, 1H) | 52.91 |
| **NH-CH-<u>C</u>OOH** | - | N/R |
| **C(O)-<u>CH₂</u>-CH** | 2.80 (dd, *J* = 15.5, 4.2 Hz, 1H), 2.58 (dd, *J* = 15.4, 9.5 Hz, 1H) | 38.47 |
| **<u>C</u>(O)-CH₂-CH** | - | 173.45 |

[a] Not applicable

[b] Not reported

ESI TOF-MS *m/z* calculated for $C_{133}H_{177}N_9O_{78}$, $[M-2H]^{2-}$: 1454.0086, found 1454.0188.

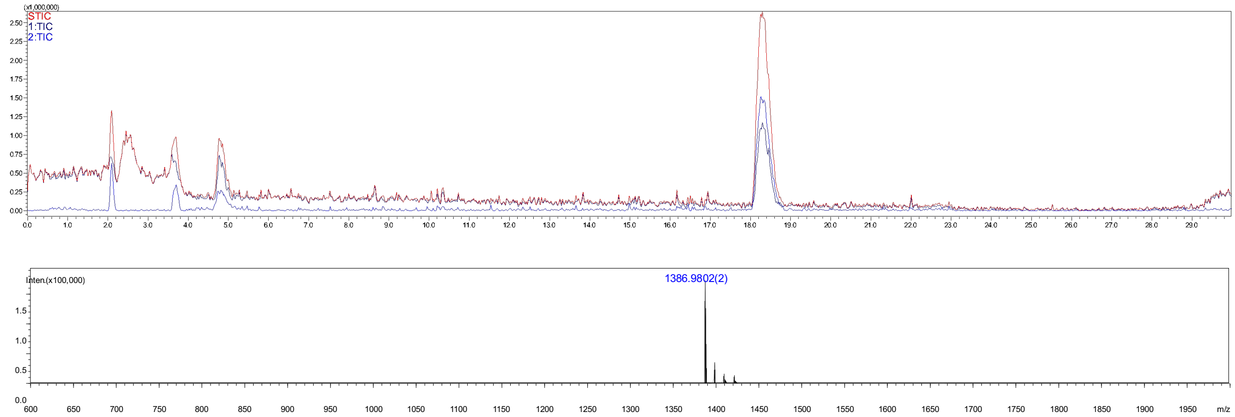


**Figure S17.** Analytical HPLC-MS chromatogram of compound **46**. The retention time = 18.3 min.

#### Compound 47b

**47b** was prepared from **42** (1.0 mg) using the general procedures **2.2 e** for the installation of α2,6-sialic acid with ST6Gal1 to give intermediate products **47a** which was purified using the described two-stage purification system **(2.2 j)**. The Cbz protecrting group of **47a** was removed by hydrogentation (**2.2 i**) to give final product **47b** as a white fluffy solid (1 mg, 90% yield for two steps). ESI TOF-MS *m/z* calculated for $C_{102}H_{165}N_9O_{74}$, $[M-2H]^{2-}$: 1349.9718, found 1349.9654.

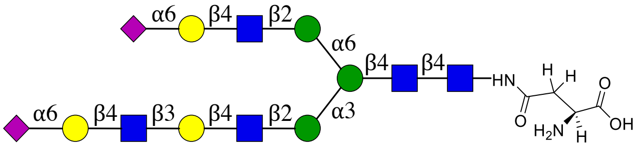


**47**

NMR and MS analysis of compound **47a**

**$^1$H (600 MHz, $D_2O$): δ (ppm)**

| | H1 | H2 | H3 | H4 | H5 | H6 |
|---|---|---|---|---|---|---|
| **GlcNAc1** | 5.09 (d, *J* = 9.8 Hz, 1H) | 3.86 | 3.77 | 3.68 | 3.59 | N/R[b] |
| **GlcNAc2** | 4.65 | 3.77 | 3.82 | N/R | N/R | N/R |
| **Man3** | 4.81 | 4.29 (s, 1H) | 3.80 | N/R | N/R | 4.0, 3.83 |
| **Man4** | 5.15 | 4.23 (d, *J* = 3.5 Hz, 1H) | 3.94 | N/R | N/R | N/R |

| | | | | | | |
|---|---|---|---|---|---|---|
| **Man4'** | 4.98 (s, 1H) | 4.17 – 4.13 (m, 1H) | 3.92 | N/R | N/R | N/R |
| **GlcNAc5** | 4.64 | 3.77 | N/R | N/R | N/R | N/R |
| **GlcNAc5'** | 4.62 | 3.77 | N/R | N/R | N/R | N/R |
| **Gal6** | 4.49 | 3.63 | 3.77 | 4.19 (d, $J$ = 3.2 Hz, 1H) | N/R | N/R |
| **Gal6'** | 4.49 | 3.57 | N/R | N/R | N/R | N/R |
| **GlcNAc7** | 4.77 | 3.84 | N/R | N/R | N/R | N/R |
| **Gal8** | 4.49 | 3.57 | N/R | N/R | N/R | N/R |

| | **H1** | **H2** | **H3** | **H4** | **H5** | **H6** | **H7** | **H8** | **H9** |
|---|---|---|---|---|---|---|---|---|---|
| **Neu5Ac7'** | -[a] | - | 2.71 (dd, $J$ = 12.5, 4.7 Hz, 2H), 1.76 (t, $J$ = 12.2 Hz, 2H) | 3.70 | 51.84 | N/R | N/R | N/R | N/R |
| **Neu5Ac9** | - | - | 2.71 (dd, $J$ = 12.5, 4.7 Hz, 2H), 1.76 (t, $J$ = 12.2 Hz, 2H) | 3.70 | 51.84 | N/R | N/R | N/R | N/R |

**$^{13}$C (150 MHz, $D_2O$): δ (ppm)**

| | **C1** |
|---|---|
| **GlcNAc1** | 78.06 |
| **GlcNAc2** | 101.18 |
| **Man3** | 100.37 |
| **Man4** | 99.54 |
| **Man4'** | 96.88 |
| **GlcNAc5** | 99.21 |
| **GlcNAc5'** | 99.39 |
| **Gal6** | 102.87 |
| **Gal6'** | 103.43 |
| **GlcNAc7** | 102.50 |
| **Gal8** | 103.43 |

| | **C1** | **C2** | **C3** | **C4** | **C5** | **C6** | **C7** | **C8** | **C9** |
|---|---|---|---|---|---|---|---|---|---|
| **Neu5Ac7'** | N/R | N/R | 40.00 | 68.16 | 51.82 | N/R | N/R | N/R | N/R |
| **Neu5Ac9** | N/R | N/R | 40.00 | 68.16 | 51.82 | N/R | N/R | N/R | N/R |

| **Signal** | **Proton** | **Carbon** |
|---|---|---|
| **NH<u>C</u>(O)$CH_3$** | - | 174.87, 174.64 |
| **NHC(O)<u>$CH_3$</u>** | 2.15 – 1.93 (m, 21H) | 22.25, 22.20 |
| **Aromatic** | 7.57 – 7.35 (m, 5H) | 128.71, 128.33, 127.72 |
| **<u>$CH_2$</u>-Ph** | 5.21 – 5.12 (m, 2H) | 67.04 |
| **NH-<u>C</u>OO-** | - | N/R |
| **NH-<u>CH</u>-COOH** | 4.52 | 51.48 |
| **NH-CH-<u>C</u>OOH** | - | N/R |
| **C(O)-<u>$CH_2$</u>-CH** | 2.88 (d, $J$ = 16.3 Hz, 1H), 2.78 – 2.73 (m, 1H) | 37.75 |

| | | |
|---|---|---|
| **C(O)-CH$_2$-CH** | - | N/R |

[a] Not applicable

[b] Not reported

ESI TOF-MS *m/z* calculated for $C_{110}H_{171}N_9O_{76}$, $[M-2H]^{2-}$: 1416.9902, found 1416.9864.

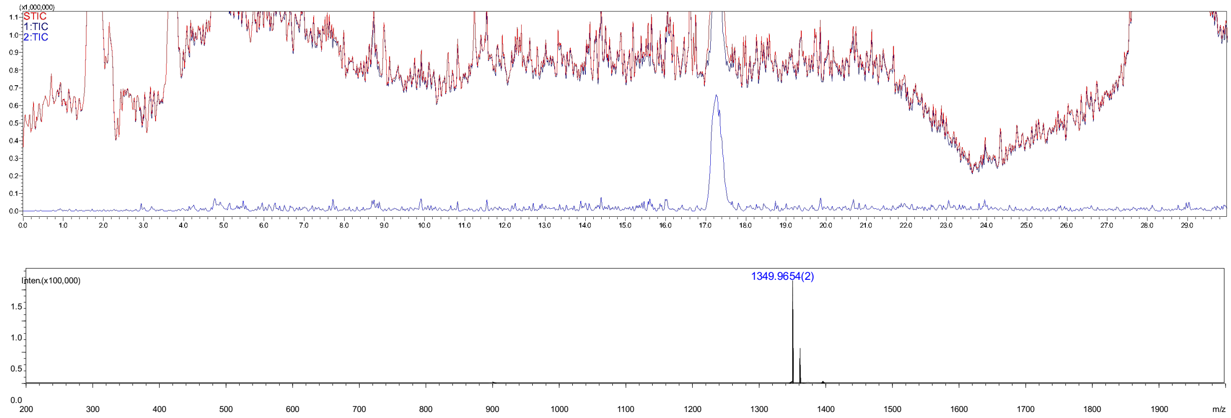

**Figure S18.** Analytical HPLC-MS chromatogram of compound **47b**. The retention time = 17.3 min.

#### Compound 48b

**48b** was prepared from **43** (1.0 mg) using the general procedures **2.2 e** for the installation of α2,6-sialic acid with ST6Gal1 to give intermediate product **48a**, which was purified using the described two-stage purification system **(2.2 j)**. The Cbz protecting group of **48a** was removed by hydrogentation (**2.2 i**) to give final product **48b** as a white fluffy solid (1.0 mg, 93% yield for two steps). ESI TOF-MS *m/z* calculated for $C_{116}H_{191}N_{10}O_{84}$, $[M-2H]^{2-}$: 1532.5379, found 1532.5301.

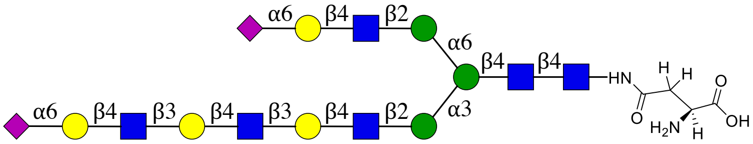


**48**

NMR and MS analysis of compound **48a**:

**$^1$H (600 MHz, $D_2O$): δ (ppm)**

| | H1 | H2 | H3 | H4 | H5 | H6 |
|---|---|---|---|---|---|---|
| **GlcNAc1** | 5.08 (d, *J* = 9.7 Hz, 1H) | 3.85 | 3.77 | 3.68 | 3.59 | N/R[b] |
| **GlcNAc2** | 4.64 | 3.78 | 3.82 | N/R | N/R | N/R |
| **Man3** | 4.80 | 4.28 (d, *J* = 2.6 Hz, 1H) | N/R | N/R | N/R | 4.00, 3.82 |
| **Man4** | 5.51 | 4.22 (d, *J* = 3.5 Hz, 1H) | N/R | N/R | N/R | N/R |

| | | | | | | |
|---|---|---|---|---|---|---|
| **Man4'** | 4.98 (s, 1H) | 4.17 – 4.13 (m, 1H) | N/R | N/R | N/R | N/R |
| **GlcNAc5** | 4.64 | 3.78 | N/R | N/R | N/R | N/R |
| **GlcNAc5'** | 4.61 | 3.78 | N/R | N/R | N/R | N/R |
| **Gal6** | 4.50 | 3.62 | 3.76 | 4.19 (d, $J$ = 2.9 Hz, 3H) | N/R | N/R |
| **Gal6'** | 4.48 | 3.57 | N/R | N/R | N/R | N/R |
| **GlcNAc7** | 4.76 | 3.83 | N/R | N/R | N/R | N/R |
| **Gal8** | 4.50 | 3.62 | 3.76 | 4.19 (d, $J$ = 2.9 Hz, 3H) | N/R | N/R |
| **GlcNAc9** | 4.73 | 3.83 | N/R | N/R | N/R | N/R |
| **Gal10** | 4.48 | 3.57 | N/R | N/R | N/R | N/R |

| | **H1** | **H2** | **H3** | **H4** | **H5** | **H6** | **H7** | **H8** | **H9** |
|---|---|---|---|---|---|---|---|---|---|
| **Neu5Ac7'** | -[a] | - | 2.75 – 2.66 (m, 3H), 1.75 (t, $J$ = 12.2 Hz, 2H) | 3.70 | 3.84 | N/R | N/R | N/R | N/R |
| **Neu5Ac11** | - | - | 2.75 – 2.66 (m, 3H), 1.75 (t, $J$ = 12.2 Hz, 2H) | 3.70 | 3.84 | N/R | N/R | N/R | N/R |

**$^{13}$C (150 MHz, $D_2O$): δ (ppm)**

| | **C1** |
|---|---|
| **GlcNAc1** | 78.12 |
| **GlcNAc2** | 101.20 |
| **Man3** | 100.43 |
| **Man4** | 99.51 |
| **Man4'** | 96.89 |
| **GlcNAc5** | 99.23 |
| **GlcNAc5'** | 99.41 |
| **Gal6** | 102.88 |
| **Gal6'** | 103.46 |
| **GlcNAc7** | 102.54 |
| **Gal8** | 102.88 |
| **GlcNAc9** | 102.65 |
| **Gal10** | 103.46 |

| | **C1** | **C2** | **C3** | **C4** | **C5** | **C6** | **C7** | **C8** | **C9** |
|---|---|---|---|---|---|---|---|---|---|
| **Neu5Ac7'** | N/R | N/R | 40.02 | 68.15 | 51.84 | N/R | N/R | N/R | N/R |
| **Neu5Ac11** | N/R | N/R | 40.02 | 68.15 | 51.84 | N/R | N/R | N/R | N/R |

| **Signal** | **Proton** | **Carbon** |
|---|---|---|
| **NHC(O)$CH_3$** | - | 174.86, 174.78, 174.64 |
| **NHC(O)$CH_3$** | 2.15 – 1.91 (m, 24H) | 22.37, 22.29, 22.24, 22.20, 22.14, 22.00, 21.93 |
| **Aromatic** | 7.55 – 7.36 (m, 5H) | 136.24, 128.71, 128.32, 127.75 |
| **$CH_2$-Ph** | 5.21 – 5.11 (m, 2H) | 67.01 |

| | | |
|---|---|---|
| **NH-COO-** | - | 157.74 |
| **NH-CH-COOH** | 4.46 | 52.09 |
| **NH-CH-COOH** | - | N/R |
| **C(O)-$CH_2$-CH** | 2.90 – 2.84 (m, 1H), 2.75 – 2.66 (m, 3H) | 38.04 |
| **C(O)-$CH_2$-CH** | - | N/R |

[a] Not applicable

[b] Not reported

ESI TOF-MS *m/z* calculated for $C_{124}H_{197}N_{10}O_{86}$, $[M-2H]^{2-}$: 1599.5563, found 1599.5423.

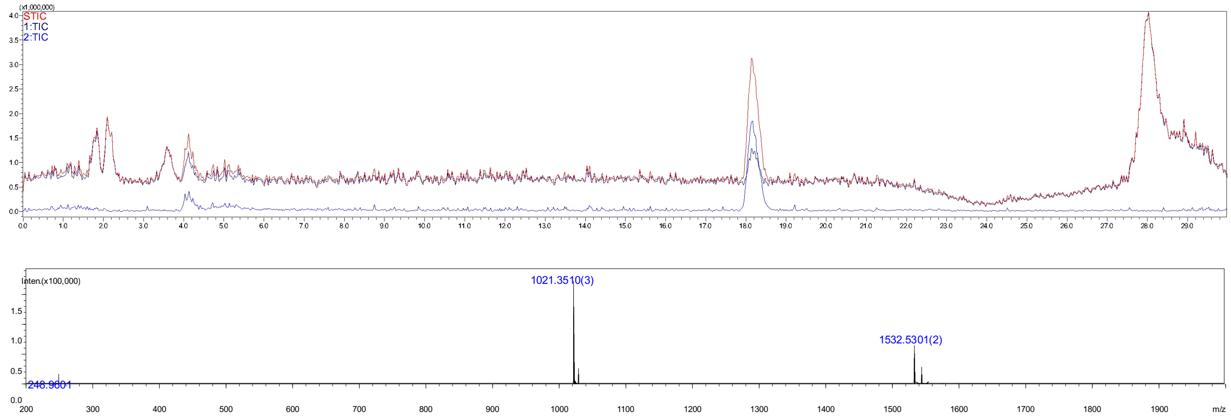


**Figure S19.** Analytical HPLC-MS chromatogram of compound **48b**. The retention time = 18.2 min.

#### Compound 52

**52** was prepared by acetylation of **23** (1 mg, 1.0 eq) according to the general procedures **2.2 h** to give intermediate product **49**, which was purified using the described two-stage purification system **(2.2 j)**. The Cbz protecting group of **49** was removed by hydrogentation (**2.2 i**) to give final product **52** as a white fluffy solid (0.7 mg, 78% yield for two steps). ESI TOF-MS *m/z* calculated for $C_{60}H_{98}N_6O_{43}$, $[M-2H]^{2-}$: 795.2839, found 795.2879.

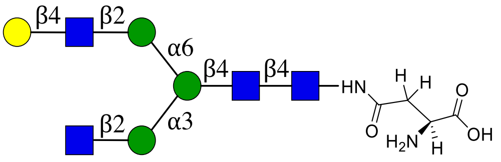


**52**

NMR and MS analysis of compound **49**

**$^1$H (600 MHz, $D_2O$): δ (ppm)**

| | H1 | H2 | H3 | H4 | H5 | H6 |
|---|---|---|---|---|---|---|
| **GlcNAc1** | 5.05 (d, $J$ = 9.7 Hz, 1H) | 3.83 | 3.74 | 3.65 | 3.56 | 3.81<br>3.63 |
| **GlcNAc2** | 4.61 (d, $J$ = 8.0 Hz, 1H) | 3.79 | 3.80 | 3.74 | 3.61 | N/R[b] |
| **Man3** | 4.77 | 4.25 (s, 1H) | 3.78 | N/R | 3.62 | 3.97<br>3.79 |
| **Man4** | 5.12 | 4.19 (s, 1H) | 3.90 | 3.51 | 3.75 | 3.92<br>3.63 |
| **Man4'** | 4.93 (s, 1H) | 4.12 (s, 1H) | 3.89 | 3.50 | 3.62 | N/R |
| **GlcNAc5** | 4.56 (d, $J$ = 8.4 Hz, 1H) | 3.70 | N/R | N/R | N/R | N/R |
| **GlcNAc5'** | 4.58 (d, $J$ = 8.1 Hz, 1H) | 3.75 | N/R | 3.73 | N/R | N/R |
| **Gal6'** | 4.48 (d, $J$ = 7.7 Hz, 2H) | 3.54 | 3.67 | 3.93 | 3.74 | 3.77 |

**$^{13}$C (150 MHz, $D_2O$): δ (ppm)**

| | C1 | C2 | C3 | C4 | C5 | C6 |
|---|---|---|---|---|---|---|
| **GlcNAc1** | 78.06 | 53.66 | 72.74 | 78.48 | 76.12 | 59.76 |
| **GlcNAc2** | 101.17 | 54.87 | 65.55 | 79.38 | 74.31 | N/R |
| **Man3** | 100.35 | 70.10 | 80.35 | N/R | 74.28 | 65.55 |
| **Man4** | 99.52 | 76.34 | 69.31 | 67.21 | 73.47 | 61.55 |
| **Man4'** | 96.95 | 76.22 | 69.36 | 67.25 | 72.76 | N/R |
| **GlcNAc5** | 99.52 | 55.24 | N/R | N/R | N/R | N/R |
| **GlcNAc5'** | 99.35 | 54.77 | N/R | 78.42 | N/R | N/R |
| **Gal6'** | 102.86 | 70.87 | 72.42 | 68.44 | 75.26 | 60.92 |

| Signal | Proton | Carbon |
|---|---|---|
| **NHC(O)CH3** | -[a] | 174.67, 174.63, 174.54 |
| **NHC(O)CH3** | 2.17 – 1.88 (m, 12H) | 22.25, 22.24, 22.14, 21.89 |
| **Aromatic** | 7.53 – 7.32 (m, 5H) | 136.18, 128.68, 128.30, 127.69 |
| **CH2-Ph** | 5.18 – 5.10 (m, 2H) | 67.02 |
| **NH-COO-** | - | 157.73 |
| **NH-CH-COOH** | 4.48 (d, $J$ = 7.7 Hz, 2H) | 51.62 |
| **NH-CH-COOH** | - | 175.88 |
| **C(O)-CH2-CH** | 2.87 – 2.81 (m, 1H),<br>2.71 (dd, $J$ = 15.7, 8.4 Hz, 1H) | 37.76 |
| **C(O)-CH2-CH** | - | 173.06 |

[a] Not applicable

[b] Not reported

ESI TOF-MS *m/z* calculated for $C_{68}H_{104}N_6O_{45}$, $[M-2H]^{2-}$: 862.3023, found 862.2961.

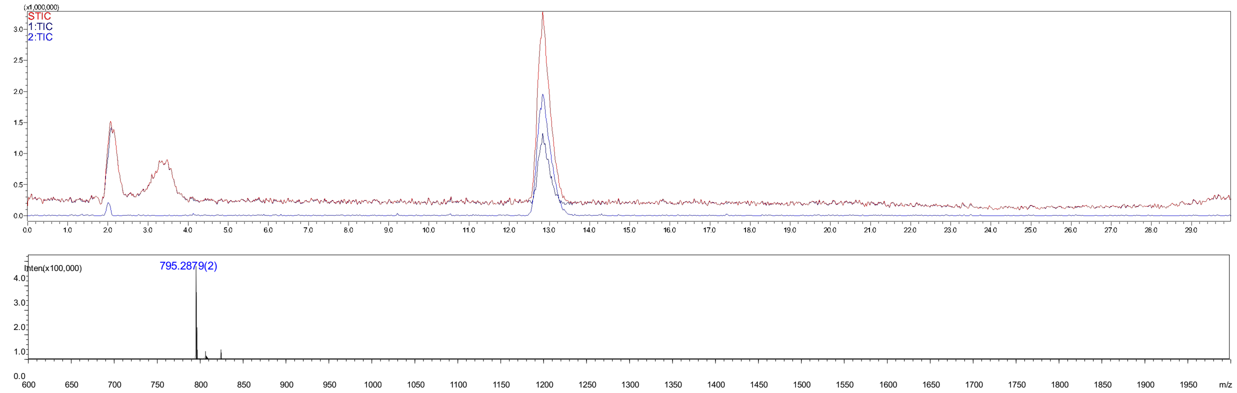

**Figure S20.** Analytical HPLC-MS chromatogram of compound **52**. The retention time = 12.8 min.

#### Compound 53

**53** was prepared by acetylation of **24** (1 mg, 1.0 eq) by the general procedures **2.2 h** to give intermediate product **50**, which was purified using the described two-stage purification system **(2.2 j)**. The Cbz protecrting group of **50** was removed by hydrogentation (**2.2 i**) to give final product **53** as a white fluffy solid (0.5 mg, 51% yield over two steps). ESI TOF-MS *m/z* calculated for $C_{74}H_{121}N_7O_{53}$, $[M-2H]^{2-}$: 977.8500, found 977.8415.

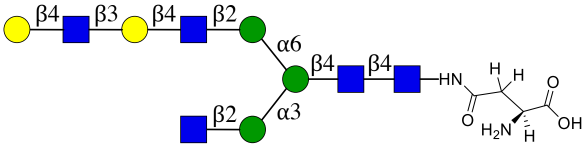


**53**

NMR and MS analysis of compound **50**

**$^1$H (600 MHz, $D_2O$): δ (ppm)**

| | H1 | H2 | H3 | H4 | H5 | H6 |
|---|---|---|---|---|---|---|
| **GlcNAc1** | 5.06 (d, *J* = 9.7 Hz, 1H) | 3.83 | 3.74 | 3.66 | 3.55 | N/R[b] |
| **GlcNAc2** | 4.62 (d, *J* = 8.1 Hz, 1H) | 3.80 | 3.80 | 3.75 | N/R | N/R |
| **Man3** | 4.77 | 4.26 (d, *J* = 2.7 Hz, 1H) | 3.78 | N/R | N/R | 3.97, 3.80 |
| **Man4** | 5.12 | 4.20 (d, *J* = 3.6 Hz, 1H) | 3.90 | 3.51 | N/R | N/R |
| **Man4'** | 4.93 (s, 1H) | 4.11 (d, *J* = 3.6 Hz, 1H) | 3.90 | 3.50 | N/R | N/R |
| **GlcNAc5** | 4.56 (d, *J* = 8.4 Hz, 1H) | 3.70 | N/R | N/R | N/R | N/R |
| **GlcNAc5'** | 4.59 (d, *J* = 8.2 Hz, 1H) | 3.75 | N/R | 3.72 | N/R | N/R |
| **Gal6'** | 4.47 (d, *J* = 8.0 Hz, 1H) | 3.59 | 3.73 | N/R | N/R | N/R |
| **GlcNAc7'** | 4.71 (d, *J* = 8.4 Hz, 1H) | 3.81 | N/R | N/R | N/R | N/R |
| **Gal8'** | 4.49 (d, *J* = 7.9 Hz, 1H) | 3.55 | 3.67 | 3.93 | 3.74 | 3.76 |

**$^{13}$C (150 MHz, $D_2O$): δ (ppm)**

| | C1 | C2 | C3 | C4 | C5 | C6 |
|---|---|---|---|---|---|---|
| **GlcNAc1** | 78.06 | 53.65 | 72.66 | 78.46 | 76.11 | N/R |
| **GlcNAc2** | 101.16 | 54.86 | 65.56 | 79.38 | N/R | N/R |
| **Man3** | 100.35 | 70.10 | 80.38 | N/R | N/R | 65.66 |
| **Man4** | 99.51 | 76.33 | 69.30 | 67.21 | N/R | N/R |
| **Man4'** | 96.95 | 76.22 | 69.36 | 67.25 | N/R | N/R |
| **GlcNH$_2$5** | 99.51 | 55.24 | N/R | N/R | N/R | N/R |
| **GlcNAc5'** | 99.36 | 54.72 | N/R | 78.46 | N/R | N/R |
| **Gal6'** | 102.89 | 69.87 | 81.98 | N/R | N/R | N/R |
| **GlcNAc7'** | 102.67 | 55.10 | N/R | N/R | N/R | N/R |
| **Gal8'** | 102.77 | 70.88 | 72.42 | 68.46 | 75.26 | 60.94 |

| Signal | Proton | Carbon |
|---|---|---|
| **NH<u>C</u>(O)CH$_3$** | -[a] | 174.81, 174.67, 174.62, 174.54 |
| **NHC(O)<u>CH$_3$</u>** | 2.23 – 1.83 (m, 15H) | 22.25, 22.14, 22.10, 21.89 |
| **Aromatic** | 7.54 – 7.32 (m, 5H) | 136.21, 128.67, 128.28, 127.72 |
| **<u>CH$_2$</u>-Ph** | 5.19 – 5.08 (m, 2H) | 66.95 |
| **NH-<u>C</u>OO-** | - | 157.71 |
| **NH-<u>CH</u>-COOH** | 4.41 (dd, *J* = 8.9, 4.4 Hz, 1H) | 52.26 |
| **NH-CH-<u>C</u>OOH** | - | 176.58 |
| **C(O)-<u>CH$_2$</u>-CH** | 2.84 (dd, *J* = 15.7, 4.4 Hz, 1H)<br>2.66 (dd, *J* = 12.3, 6.6 Hz, 1H) | 38.10 |
| **<u>C</u>(O)-CH$_2$-CH** | - | 173.31 |

[a] Not applicable

[b] Not reported

ESI TOF-MS *m/z* calculated for $C_{82}H_{127}N_7O_{55}$, $[M-2H]^{2-}$: 1044.8684, found 1044.8757.

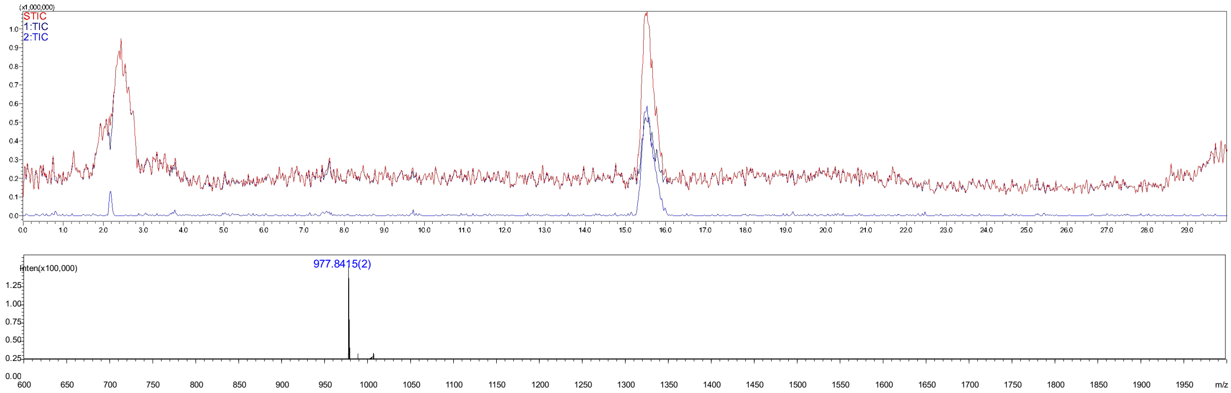

**Figure S21.** Analytical HPLC-MS chromatogram of compound **53**. The retention time = 15.6 min.

#### Compound 54

**54** was prepared by acetylation of **25** (1.0 mg) according to the general procedures **2.2 h** to give intermediate product **51** which was purified using the described two-stage purification system **(2.2 j)**. The Cbz protecrting group of **51** was removed by hydrogentation (**2.2 i**) to give final product **54** as a white fluffy solid (0.8 mg, 88% yield over two steps). ESI TOF-MS *m/z* calculated for $C_{88}H_{144}N_8O_{63}$, $[M-2H]^{2-}$: 1160.4161, found 1160.4119.

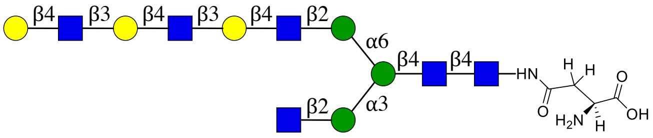

**54**

NMR and MS analysis of compound **51**

**$^1$H (600 MHz, $D_2O$): δ (ppm)**

| | H1 | H2 | H3 | H4 | H5 | H6 |
|---|---|---|---|---|---|---|
| **GlcNAc1** | 5.05 (d, *J* = 9.7 Hz, 1H) | 3.83 | 3.74 | 3.65 | 3.55 | N/R[b] |
| **GlcNAc2** | 4.61 (d, *J* = 8.2 Hz, 1H) | 3.81 | 3.80 | 3.74 | N/R | N/R |
| **Man3** | 4.77 | 4.25 (d, *J* = 2.7 Hz, 1H) | 3.78 | N/R | N/R | 3.97, 3.80 |
| **Man4** | 5.12 | 4.21 – 4.17 (m, 1H) | 3.90 | 3.50 | N/R | N/R |
| **Man4'** | 4.93 (s, 1H) | 4.11 (d, *J* = 3.6 Hz, 1H) | 3.90 | 3.50 | N/R | N/R |
| **GlcNAc5** | 4.56 (d, *J* = 8.3 Hz, 1H) | 3.70 | N/R | N/R | N/R | N/R |
| **GlcNAc5'** | 4.58 (d, *J* = 8.3 Hz, 1H) | 3.75 | N/R | 3.72 | N/R | N/R |
| **Gal6'** | 4.52 – 4.42 (m, 3H) | N/R | 3.73 | N/R | N/R | N/R |
| **GlcNAc7'** | 4.71 (d, *J* = 8.1 Hz, 2H) | 3.80 | N/R | N/R | N/R | N/R |
| **Gal8'** | 4.52 – 4.42 (m, 3H) | N/R | 3.73 | N/R | N/R | N/R |
| **GlcNAc9'** | 4.71 (d, *J* = 8.1 Hz, 2H) | 3.80 | N/R | N/R | N/R | N/R |
| **Gal10'** | 4.52 – 4.42 (m, 3H) | 3.55 | 3.67 | 3.93 | 3.73 | 3.75 |

**$^{13}$C (150 MHz, D$_2$O): δ (ppm)**

| | C1 | C2 | C3 | C4 | C5 | C6 |
|---|---|---|---|---|---|---|
| **GlcNAc1** | 78.06 | 53.66 | 72.64 | 78.47 | 76.12 | N/R |
| **GlcNAc2** | 101.17 | 54.87 | 65.56 | 79.38 | N/R | N/R |
| **Man3** | 100.35 | 70.10 | 80.37 | N/R | N/R | 65.65 |
| **Man4** | 99.51 | 76.33 | 69.31 | 67.21 | N/R | N/R |
| **Man4'** | 96.95 | 76.22 | 69.36 | 67.25 | N/R | N/R |
| **GlcNH$_2$5** | 99.51 | 55.24 | N/R | N/R | N/R | N/R |
| **GlcNAc5'** | 99.35 | 54.72 | N/R | 78.45 | N/R | N/R |
| **Gal6'** | 102.89 | N/R | 81.99 | N/R | N/R | N/R |
| **GlcNAc7'** | 102.67 | 55.10 | N/R | N/R | N/R | N/R |
| **Gal8'** | 102.80 | N/R | 81.99 | N/R | N/R | N/R |
| **GlcNAc9'** | 102.67 | 55.06 | N/R | N/R | N/R | N/R |
| **Gal10'** | 102.77 | 70.88 | 72.42 | 68.46 | 75.26 | 60.94 |

| Signal | Proton | Carbon |
|---|---|---|
| **NHC(O)CH$_3$** | -[a] | 174.81, 174.67, 174.62, 174.53 |
| **NHC(O)CH$_3$** | 2.11 – 1.87 (m, 18H) | 22.25, 22.14, 22.10, 21.89 |
| **Aromatic** | 7.54 – 7.34 (m, 5H) | 136.19, 128.68, 128.30, 127.70 |
| **CH$_2$-Ph** | 5.18 – 5.09 (m, 2H) | 67.01 |
| **NH-COO-** | - | 157.72 |
| **NH-CH-COOH** | 4.46 | 51.78 |
| **NH-CH-COOH** | - | 176.07 |
| **C(O)-CH$_2$-CH** | 2.84 (dd, $J$ = 15.7, 4.5 Hz, 1H), 2.69 (dd, $J$ = 15.9, 8.7 Hz, 1H) | 37.85 |
| **C(O)-CH$_2$-CH** | - | 173.11 |

[a] Not applicable

[b] Not reported

ESI TOF-MS *m/z* calculated for $C_{96}H_{150}N_8O_{65}$, $[M-2H]^{2-}$: 1227.4344, found 1227.4397.

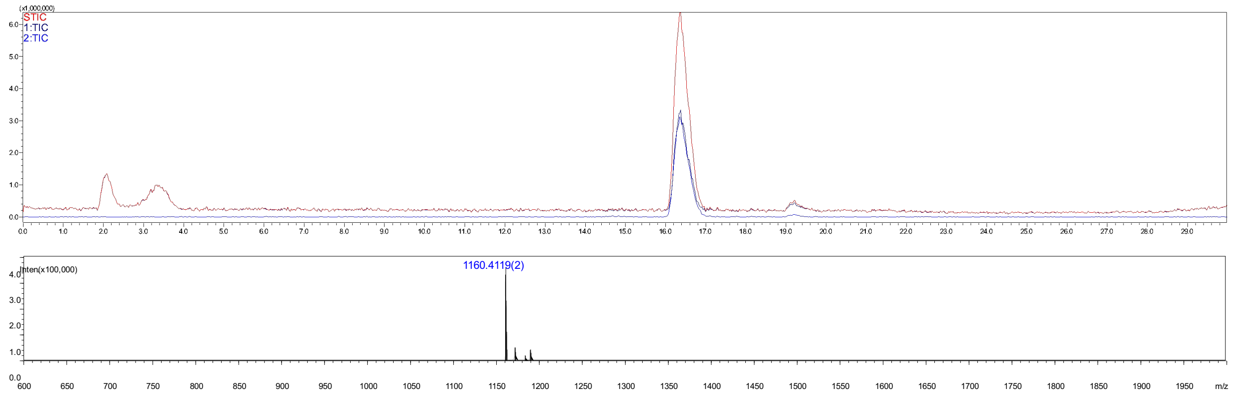


**Figure S22.** Analytical HPLC-MS chromatogram of compound **54**. The retention time = 16.4 min.

#### Compound 58b

**58b** was synthesized from **23** (3 mg, 1.0 eq) using the general procedures **2.2 f**, **h**, **c** for the installation of α2,3-sialic acid with ST3Gal4, converting $GlcNH_2$ to GlcNAc with AcOSu, installation of β1,4-Gal with B4GalT1 to give the intermediate product **58a**, which was purified using the described two-stage purification system **(2.2 j)**. The Cbz protecting group of **58a** was removed by hydrogentation (**2.2 i**) to give final product **58b** as a white fluffy solid (1.6 mg, 43% yield over four steps). ESI TOF-MS *m/z* calculated for $C_{77}H_{125}N_7O_{56}$, $[M-2H]^{2-}$: 1021.8580, found 1021.8615.

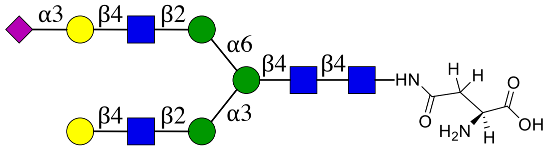


**58**

NMR and MS analysis of compound **58a**

**$^1$H (600 MHz, $D_2O$): δ (ppm)**

| | H1 | H2 | H3 | H4 | H5 | H6 |
|---|---|---|---|---|---|---|
| **GlcNAc1** | 5.06 (d, *J* = 9.8 Hz, 1H) | 3.82 | 3.74 | 3.65 | 3.57 | N/R[b] |
| **GlcNAc2** | 4.62 (d, *J* = 8.1 Hz, 1H) | 3.80 | 3.80 | N/R | N/R | N/R |
| **Man3** | 4.77 | 4.26 (d, *J* = 2.9 Hz, 1H) | 3.77 | N/R | N/R | 3.98, 3.79 |
| **Man4** | 5.13 | 4.20 (dd, *J* = 3.3, 1.6 Hz, 1H) | 3.91 | N/R | N/R | N/R |
| **Man4'** | 4.93 (d, *J* = 1.7 Hz, 1H) | 4.16 – 4.09 (m, 2H) | 3.97 | N/R | N/R | N/R |
| **GlcNAc5** | 4.58 (dd, *J* = 7.8, 3.1 Hz, 2H) | 3.76 | N/R | N/R | N/R | N/R |
| **GlcNAc5'** | 4.58 (dd, *J* = 7.8, 3.1 Hz, 2H) | 3.76 | N/R | N/R | N/R | N/R |
| **Gal6** | 4.56 (d, *J* = 7.9 Hz, 1H) | 3.55 | N/R | N/R | N/R | N/R |
| **Gal6'** | 4.47 (d, *J* = 7.7 Hz, 1H) | 3.58 | 3.68 | 4.16 – 4.09 (m, 2H) | N/R | N/R |

| | H1 | H2 | H3 | H4 | H5 | H6 | H7 | H8 | H9 |
|---|---|---|---|---|---|---|---|---|---|
| **Neu5Ac7'** | -[a] | - | 2.77 (dd, *J* = 12.5, 4.6 Hz, 1H), 1.81 (t, *J* = 12.1 Hz, 1H) | 3.70 | 3,86 | N/R | N/R | N/R | N/R |

**$^{13}$C (150 MHz, $D_2O$): δ (ppm)**

| | C1 |
|---|---|
| **GlcNAc1** | 78.04 |
| **GlcNAc2** | 101.17 |
| **Man3** | 100.35 |
| **Man4** | 99.46 |

| | |
|---|---|
| **Man4'** | 97.02 |
| **GlcNAc5** | 99.36 |
| **GlcNAc5'** | 99.36 |
| **Gal6** | 102.53 |
| **Gal6'** | 102.83 |

| | **C1** | **C2** | **C3** | **C4** | **C5** | **C6** | **C7** | **C8** | **C9** |
|---|---|---|---|---|---|---|---|---|---|
| **Neu5Ac7'** | N/R | 99.71 | 39.54 | 68.26 | 51.60 | N/R | N/R | N/R | N/R |

| **Signal** | **Proton** | **Carbon** |
|---|---|---|
| **NH<u>C</u>(O)CH3** | - | 174.92, 174.67, 174.62, 174.55, 173.75 |
| **NHC(O)<u>CH3</u>** | 2.21 – 1.88 (m, 15H) | 22.25, 22.15, 21.95, 21.89 |
| **Aromatic** | 7.55 – 7.33 (m, 5H) | 136.19, 128.68, 128.30, 127.70 |
| **<u>CH2</u>-Ph** | 5.19 – 5.09 (m, 2H) | 67.01 |
| **NH-<u>C</u>OO-** | - | 157.74 |
| **NH-<u>CH</u>-COOH** | 4.45 | 51.76 |
| **NH-CH-<u>C</u>OOH** | - | N/R |
| **C(O)-<u>CH2</u>-CH** | 2.85 (dd, $J$ = 15.6, 4.5 Hz, 1H), 2.70 (dd, $J$ = 15.6, 8.6 Hz, 1H) | 47.84 |
| **<u>C</u>(O)-CH2-CH** | - | 173.11 |

[a] Not applicable

[b] Not reported

ESI TOF-MS *m/z* calculated for $C_{85}H_{131}N_7O_{58}$, $[M-2H]^{2-}$: 1088.8764, found 1088.8769.

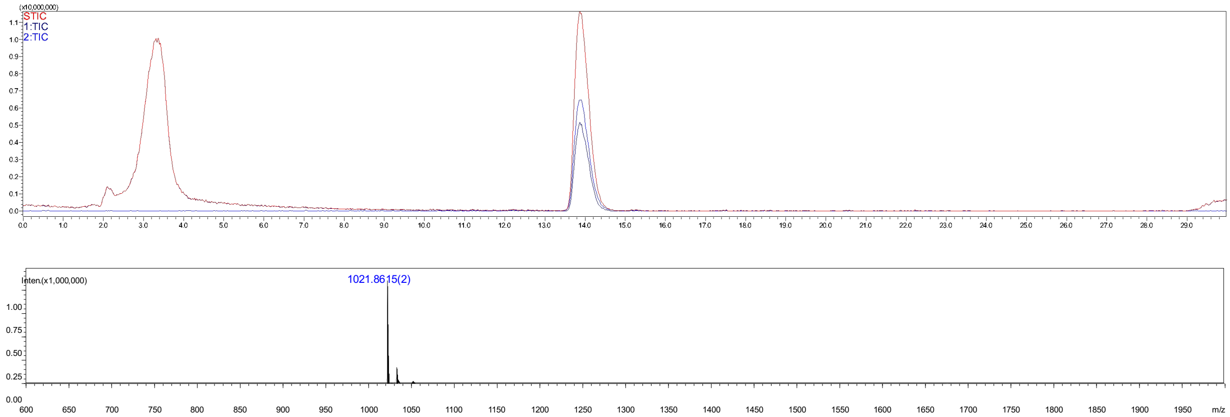


**Figure S23.** Analytical HPLC-MS chromatogram of compound **58b**. The retention time = 14.0 min.

**Compound 59b**

**59b** was synthesized from **24** (3 mg, 1.0 eq) using the general procedures **2.2 f**, **h**, **c** and **i** for the installation of α-2,3-sialic acid with ST3Gal4, converting $GlcNH_2$ to GlcNAc with AcOSu, installation of β1,4-Gal with B4GalT1 to give **59a** that was purified using the described two-stage purification system **(2.2 j)**, The Cbz protecrting group of **59a** was removed by hydrogentation (**2.2 i**) to give final product the final product **59b** as a white fluffy solid (1.8 mg, 50% over four steps). ESI TOF-MS *m/z* calculated for $C_{91}H_{148}N_8O_{66}$, $[M-2H]^{2-}$: 1204.4241, found 1204.4299.

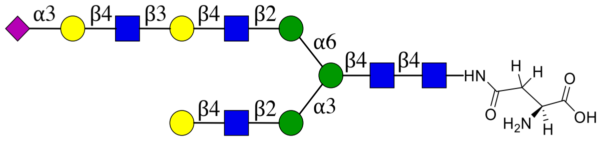


**59**

NMR and MS analysis of compound **59a** attaching Cbz group

**$^1$H (600 MHz, $D_2O$): δ (ppm)**

| | H1 | H2 | H3 | H4 | H5 | H6 |
|---|---|---|---|---|---|---|
| **GlcNAc1** | 5.03 (d, *J* = 9.7 Hz, 1H) | 3.79 | 3.72 | 3.62 | 3.53 | N/R[b] |
| **GlcNAc2** | 4.58 (d, *J* = 8.1 Hz, 1H) | 3.77 | 3.77 | N/R | N/R | N/R |
| **Man3** | 4.74 | 4.22 (d, *J* = 2.8 Hz, 1H) | 3.75 | N/R | N/R | 3.94, 3.76 |
| **Man4** | 5.09 | 4.19 – 4.15 (m, 1H) | 3.87 | N/R | N/R | N/R |
| **Man4'** | 4.90 (s, 1H) | 4.12 – 4.05 (m, 2H) | 3.86 | N/R | N/R | N/R |
| **GlcNAc5** | 4.57 – 4.52 (m, 3H) | 3.72 | N/R | N/R | N/R | N/R |
| **GlcNAc5'** | 4.57 – 4.52 (m, 3H) | 3.72 | N/R | N/R | N/R | N/R |
| **Gal6** | 4.57 – 4.52 (m, 3H) | 3.55 | N/R | N/R | N/R | N/R |
| **Gal6'** | 4.47 – 4.40 (m, 2H) | 3.56 | 3.70 | 4.14 (d, *J* = 3.2 Hz, 1H) | N/R | N/R |
| **GlcNAc7'** | 4.67 (d, *J* = 8.4 Hz, 1H) | 3.78 | N/R | N/R | N/R | N/R |
| **Gal8'** | 4.47 – 4.40 (m, 2H) | 3.51 | 3.93 | 4.12 – 4.05 (m, 2H) | N/R | N/R |

| | H1 | H2 | H3 | H4 | H5 | H6 | H7 | H8 | H9 |
|---|---|---|---|---|---|---|---|---|---|
| **Neu5Ac9'** | -[a] | - | 2.77 – 2.68 (m, 2H), 1.78 (t, *J* = 12.1 Hz, 1H) | 3.67 | 3.83 | N/R | N/R | N/R | N/R |

**$^{13}$C (150 MHz, $D_2O$): δ (ppm)**

| | C1 |
|---|---|
| **GlcNAc1** | 78.06 |
| **GlcNAc2** | 101.17 |
| **Man3** | 100.36 |

| | |
|---|---|
| **Man4** | 99.46 |
| **Man4'** | 96.96 |
| **GlcNAc5** | 99.36 |
| **GlcNAc5'** | 99.36 |
| **Gal6** | 102.45 |
| **Gal6'** | 102.83 |
| **GlcNAc7'** | 102.72 |
| **Gal8'** | 102.88 |

| | **C1** | **C2** | **C3** | **C4** | **C5** | **C6** | **C7** | **C8** | **C9** |
|---|---|---|---|---|---|---|---|---|---|
| **Neu5Ac9'** | N/R | 99.66 | 39.52 | 68.21 | 51.59 | N/R | N/R | N/R | N/R |

| **Signal** | **Proton** | **Carbon** |
|---|---|---|
| **NHC(O)CH$_3$** | - | 175.16, 174.92, 174.79, 174.65, 174.61, 174.54 |
| **NHC(O)CH$_3$** | 2.19 – 1.85 (m, 18H) | 22.25, 22.14, 22.11, 21.95, 21.88 |
| **Aromatic** | 7.54 – 7.27 (m, 5H) | 136.14, 128.68, 128.32, 127.67 |
| **CH$_2$-Ph** | 5.16 – 5.06 (m, 2H) | 67.07 |
| **NH-COO-** | - | 157.76 |
| **NH-CH-COOH** | 4.50 (dd, $J$ = 7.9, 4.8 Hz, 1H) | 51.05 |
| **NH-CH-COOH** | - | N/R |
| **C(O)-CH$_2$-CH** | 2.83 (dd, $J$ = 15.9, 4.7 Hz, 1H), 2.77 – 2.68 (m, 2H) | 37.45 |
| **C(O)-CH$_2$-CH** | - | 172.81 |

[a] Not applicable

[b] Not reported

ESI TOF-MS *m/z* calculated for $C_{99}H_{154}N_8O_{68}$, $[M-2H]^{2-}$: 1271.4425, found 1271.4428.

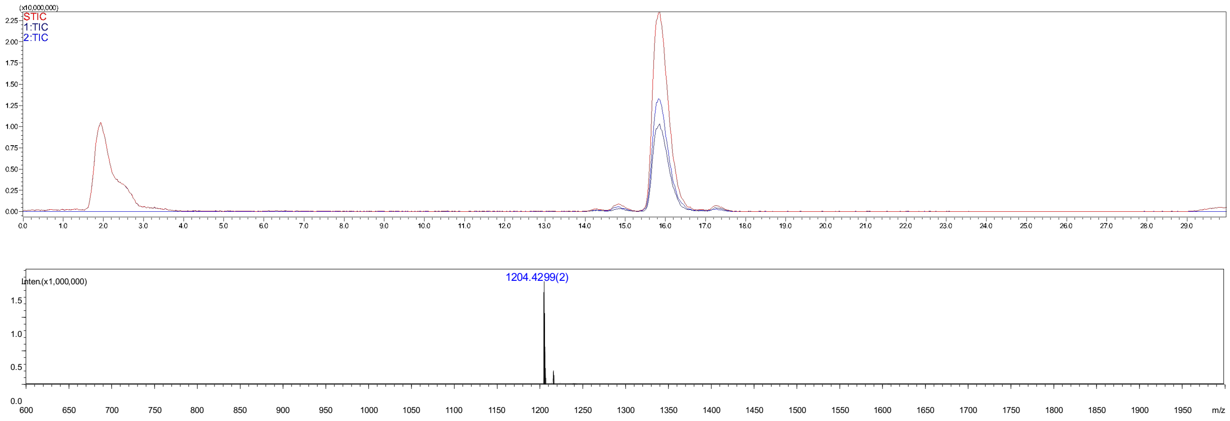


**Figure S24.** Analytical HPLC-MS chromatogram of compound **59b**. The retention time = 15.8 min.

**Compound 60b**

**60a** was synthesized from **25** (3 mg, 1.0 eq) using the general procedures **2.2 f**, **h**, **c** for the installation of α2,3-sialic acid with ST3Gal4, converting $GlcNH_2$ to GlcNAc with AcOSu, installation of β1,4-Gal with B4GalT1 to give product **60a** which purified using the described two-stage purification system **(2.2 j)**. The Cbz protecrting group of **60a** was removed by hydrogentation (**2.2 i**) to give final product the final product **60b** as a white fluffy solid (1 mg, 29% yield over four steps). ESI TOF-MS *m/z* calculated for $C_{105}H_{171}N_9O_{76}$, $[M-2H]^{2-}$: 1386.9902, found 1386.9792.

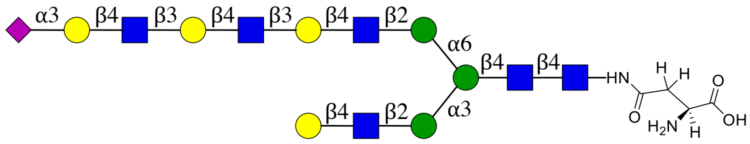


**60**

NMR and MS analysis of compound **60a**

**$^1$H (600 MHz, $D_2O$): δ (ppm)**

| | H1 | H2 | H3 | H4 | H5 | H6 |
|---|---|---|---|---|---|---|
| **GlcNAc1** | 5.06 (d, *J* = 9.7 Hz, 1H) | 3.84 | 3.75 | 3.66 | 3.57 | N/R[b] |
| **GlcNAc2** | 4.62 (d, *J* = 8.0 Hz, 1H) | 3.76 | 3.80 | N/R | N/R | N/R |
| **Man3** | 4.77 | 4.26 (d, *J* = 2.7 Hz, 1H) | 3.78 | N/R | N/R | 3.97, 3.80 |
| **Man4** | 5.13 | 4.20 (d, *J* = 3.6 Hz, 1H) | 3.91 | N/R | N/R | N/R |
| **Man4'** | 4.93 (s, 1H) | 4.15 – 4.09 (m, 2H) | 3.90 | N/R | N/R | N/R |
| **GlcNAc5** | 4.60 – 4.52 (m, 4H) | 3.74 | N/R | N/R | N/R | N/R |
| **GlcNAc5'** | 4.60 – 4.52 (m, 4H) | 3.74 | N/R | N/R | N/R | N/R |
| **Gal6** | 4.51 – 4.43 (m, 3H) | 3.59 | N/R | N/R | N/R | N/R |
| **Gal6'** | 4.51 – 4.43 (m, 3H) | 3.55 | 3.73 | 4.17 (d, *J* = 3.1 Hz, 2H) | N/R | N/R |
| **GlcNAc7'** | 4.70 (d, *J* = 8.3 Hz, 2H) | 3.76 | N/R | N/R | N/R | N/R |
| **Gal8'** | 4.51 – 4.43 (m, 3H) | 3.55 | 3.73 | 4.17 (d, *J* = 3.1 Hz, 2H) | N/R | N/R |
| **GlcNAc9'** | 4.70 (d, *J* = 8.3 Hz, 2H) | 3.76 | N/R | N/R | N/R | N/R |
| **Gal10'** | 4.60 – 4.52 (m, 4H) | 3.58 | 3.97 | 4.15 – 4.09 (m, 2H) | N/R | N/R |

| | H1 | H2 | H3 | H4 | H5 | H6 | H7 | H8 | H9 |
|---|---|---|---|---|---|---|---|---|---|
| **Neu5Ac11'** | -[a] | - | 2.81 – 2.73 (m, 2H), 1.81 (t, *J* = 12.1 Hz, 2H) | 3.70 | 3.86 | N/R | N/R | N/R | N/R |

**$^{13}$C (150 MHz, $D_2O$): δ (ppm)**

| | C1 |
|---|---|
| **GlcNAc1** | 78.07 |
| **GlcNAc2** | 101.17 |
| **Man3** | 100.35 |
| **Man4** | 99.45 |
| **Man4'** | 96.96 |
| **GlcNAc5** | 99.35 |
| **GlcNAc5'** | 99.35 |
| **Gal6** | 102.83 |
| **Gal6'** | 102.83 |
| **GlcNAc7'** | 102.72 |
| **Gal8'** | 102.83 |
| **GlcNAc9'** | 102.72 |
| **Gal10'** | 102.45 |

| | C1 | C2 | C3 | C4 | C5 | C6 | C7 | C8 | C9 |
|---|---|---|---|---|---|---|---|---|---|
| **Neu5Ac11'** | N/R | 99.63 | 39.51 | 68.21 | 51.60 | N/R | N/R | N/R | N/R |

| Signal | Proton | Carbon |
|---|---|---|
| **$NH\underline{C}(O)CH_3$** | - | 175.00, 174.92, 174.80, 174.62, 174.54 |
| **$NHC(O)\underline{CH_3}$** | 2.30 – 1.90 (m, 21H) | 22.25, 22.14, 22.10, 21.96, 21.88 |
| **Aromatic** | 7.53 – 7.34 (m, 5H) | 136.14, 128.69, 128.32, 127.67 |
| **$\underline{CH_2}$-Ph** | 5.19 – 5.10 (m, 2H) | 67.10 |
| **NH-$\underline{C}$OO-** | - | 157.77 |
| **NH-$\underline{CH}$-COOH** | 4.60 – 4.52 (m, 4H) | 50.95 |
| **NH-CH-$\underline{C}$OOH** | - | N/R |
| **C(O)-$\underline{CH_2}$-CH** | 2.87 (dd, $J$ = 15.9, 4.8 Hz, 1H),<br>2.81 – 2.73 (m, 2H) | 3739 |
| **$\underline{C}$(O)-$CH_2$-CH** | - | 172.76 |

[a] Not applicable

[b] Not reported

ESI TOF-MS *m/z* calculated for $C_{113}H_{177}N_9O_{78}$, $[M-2H]^{2-}$: 1454.0086, found 1453.9978.

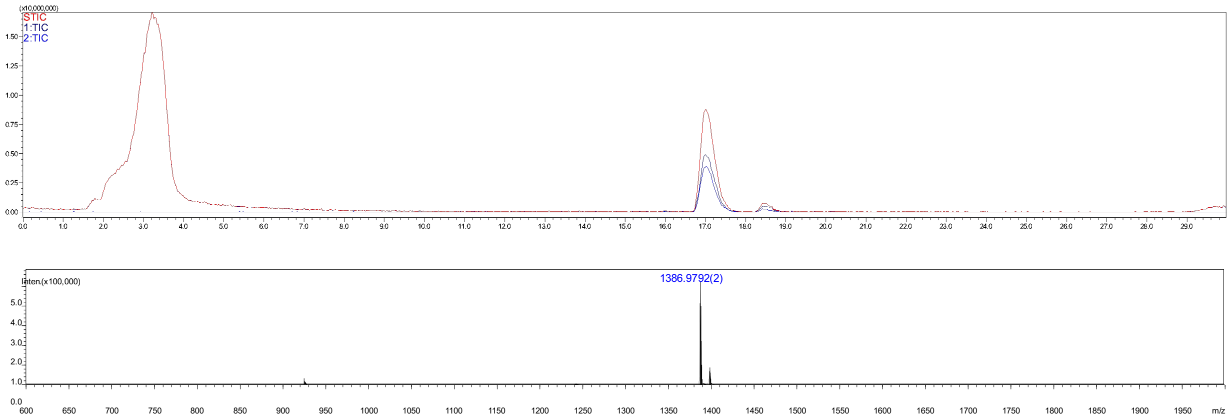


**Figure S25.** Analytical HPLC-MS chromatogram of compound **60b**. The retention time = 17.0 min.

#### Compound 67

**67** was synthesized from **23** (3 mg, 1.0 eq) using the general procedures **2.2 e**, **h**, **c** for the installation of α2,6-sialic acid with ST6Gal1, converting $GlcNH_2$ to GlcNAc with AcOSu, installation of β1,4-Gal with B4GalT1 to give the product **64** which was purified using the described two-stage purification system **(2.2 j)**. The Cbz protecrting group of **64** was removed by hydrogentation (**2.2 i**) to give final product **67** as a white fluffy solid (0.6 mg, 47% yield for four steps). ESI TOF-MS *m/z* calculated for $C_{77}H_{125}N_7O_{56}$, $[M-2H]^{2-}$: 1021.8580, found 1021.8581.

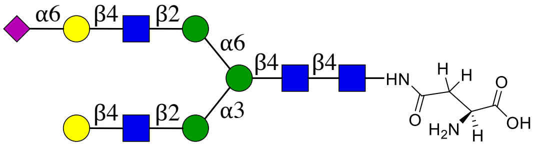


**67**

NMR and MS analysis of compound **64**

**$^1$H (600 MHz, $D_2O$): δ (ppm)**

| | H1 | H2 | H3 | H4 | H5 | H6 |
|---|---|---|---|---|---|---|
| **GlcNAc1** | 5.06 (d, $J$ = 9.8 Hz, 1H) | 3.83 | 3.74 | 3.65 | 3.56 | N/R[b] |
| **GlcNAc2** | 4.62 | 3.78 | 3.79 | N/R | N/R | N/R |
| **Man3** | 4.78 | 4.26 (s, 1H) | 3.79 | N/R | N/R | 3.97, 3.80 |
| **Man4** | 5.13 | 4.20 (d, $J$ = 3.4 Hz, 1H) | 3.92 | N/R | N/R | N/R |
| **Man4'** | 4.95 (s, 1H) | 4.12 (d, $J$ = 3.4 Hz, 1H) | 3.91 | N/R | N/R | N/R |
| **GlcNAc5** | 4.59 | 3.78 | N/R | N/R | N/R | N/R |
| **GlcNAc5'** | 4.61 | 3.78 | N/R | N/R | N/R | N/R |
| **Gal6** | 4.47 | 3.55 | N/R | N/R | N/R | N/R |
| **Gal6'** | 4.46 | 3.55 | N/R | N/R | N/R | N/R |

| | H1 | H2 | H3 | H4 | H5 | H6 | H7 | H8 | H9 |
|---|---|---|---|---|---|---|---|---|---|
| **Neu5Ac7'** | -[a] | - | 2.72 – 2.60 (m, 2H), 1.73 (t, $J$ = 12.1 Hz, 1H) | 3.67 | 3.82 | N/R | N/R | N/R | N/R |

**$^{13}$C (150 MHz, $D_2O$): δ (ppm)**

| | C1 |
|---|---|
| **GlcNAc1** | 78.05 |
| **GlcNAc2** | 101.17 |
| **Man3** | 100.42 |
| **Man4** | 99.46 |
| **Man4'** | 99.37 |
| **GlcNAc5** | 99.20 |
| **GlcNAc5'** | 102.83 |
| **Gal6** | 103.48 |
| **Gal6'** | 100.07 |

| | C1 | C2 | C3 | C4 | C5 | C6 | C7 | C8 | C9 |
|---|---|---|---|---|---|---|---|---|---|
| **Neu5Ac7'** | N/R | 100.07 | 39.99 | 68.12 | 51.80 | N/R | N/R | N/R | N/R |

| Signal | Proton | Carbon |
|---|---|---|
| **NH<u>C</u>(O)CH3** | - | 174.83, 174.62, 174.52, 173.40 |
| **NHC(O)<u>CH3</u>** | 2.28 – 1.83 (m, 15H) | 22.34, 22.26, 22.16, 21.97, 21.90 |
| **Aromatic** | 7.59 – 7.29 (m, 5H) | 136.21, 128.68, 128.29, 127.73 |
| **<u>CH2</u>-Ph** | 5.19 – 5.09 (m, 2H) | 66.95 |
| **NH-<u>C</u>OO-** | - | 157.72 |
| **NH-<u>CH</u>-COOH** | 4.41 (d, $J$ = 8.1 Hz, 1H) | 52.29 |
| **NH-CH-<u>C</u>OOH** | - | 176.36 |
| **C(O)-<u>CH2</u>-CH** | 2.87 – 2.80 (m, 1H), 2.72 – 2.60 (m, 2H) | 38.10 |
| **<u>C</u>(O)-CH2-CH** | - | 173.30 |

[a] Not applicable

[b] Not reported

ESI TOF-MS *m/z* calculated for $C_{85}H_{131}N_7O_{58}$, $[M-2H]^{2-}$: 1088.8764, found 1088.8733.

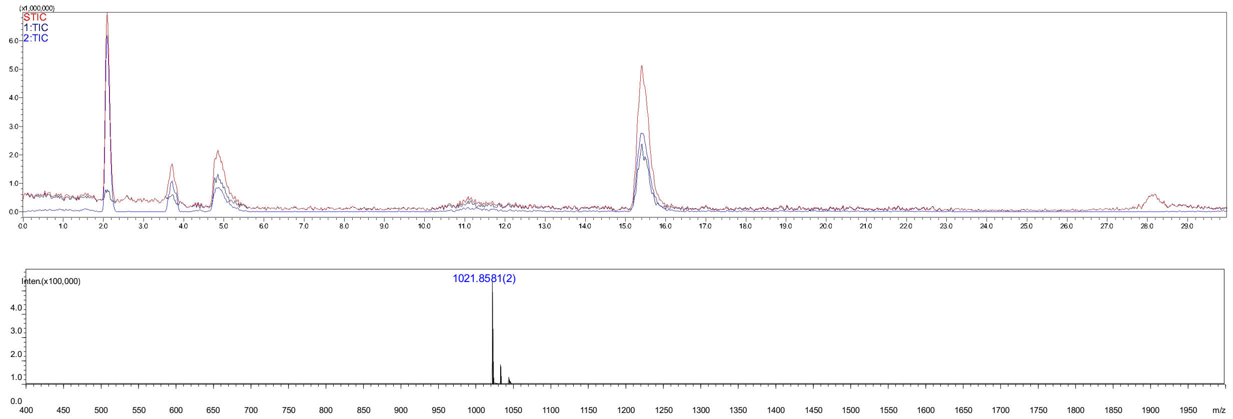


**Figure S26.** Analytical HPLC-MS chromatogram of compound **67**. The retention time = 15.4 min.

#### Compound 68

**68** was synthesized from **24** (3.0 mg) using the general procedures **2.2 e**, **h**, **c** for the installation of α2,6-sialic acid with ST6Gal1, converting $GlcNH_2$ to GlcNAc with AcOSu, installation of β1,4-Gal with B4GalT1 to give **65** which was purified using the described two-stage purification system **(2.2 j)**. The Cbz protecting group of **65** was removed by hydrogentation (**2.2 i**) to give final product **68** as a white fluffy solid (0.7 mg, 60% yield for four steps). ESI TOF-MS *m/z* calculated for $C_{91}H_{148}N_8O_{66}$, $[M-2H]^{2-}$: 1204.4241, found 1204.4337.

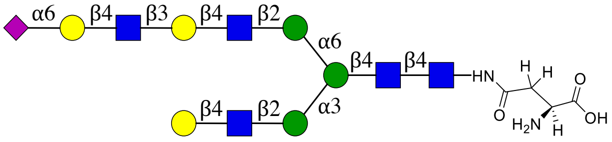


**68**

NMR and MS analysis of compound **65**

**$^1$H (600 MHz, $D_2O$): δ (ppm)**

| | H1 | H2 | H3 | H4 | H5 | H6 |
|---|---|---|---|---|---|---|
| **GlcNAc1** | 5.09 (d, *J* = 9.8 Hz, 1H) | 3.86 | 3.77 | 3.69 | 3.59 | N/R[b] |
| **GlcNAc2** | 4.65 (d, *J* = 8.1 Hz, 1H) | 3.78 | 3.83 | N/R | N/R | N/R |
| **Man3** | 4.80 | 4.29 (d, *J* = 2.7 Hz, 1H) | 3.81 | N/R | N/R | 4.00, 3.83 |
| **Man4** | 5.16 | 4.25 – 4.22 (m, 1H) | 3.94 | N/R | N/R | N/R |
| **Man4'** | 4.97 (s, 1H) | 4.17 – 4.13 (m, 1H) | 3.92 | N/R | N/R | N/R |
| **GlcNAc5** | 4.62 (d, *J* = 7.7 Hz, 2H) | 3.78 | N/R | N/R | N/R | N/R |
| **GlcNAc5'** | 4.62 (d, *J* = 7.7 Hz, 2H) | 3.78 | N/R | N/R | N/R | N/R |
| **Gal6** | 4.53 – 4.45 (m, 3H) | 3.58 | N/R | N/R | N/R | N/R |

| | | | | | | |
|---|---|---|---|---|---|---|
| **Gal6'** | 4.53 – 4.45 (m, 3H) | 3.58 | 3.77 | 4.20 (d, $J$ = 3.2 Hz, 1H) | | |
| **GlcNAc7'** | 4.77 (s, 1H) | 3.84 | N/R | N/R | N/R | N/R |
| **Gal8'** | 4.53 – 4.45 (m, 3H) | 3.64 | N/R | N/R | N/R | N/R |

| | **H1** | **H2** | **H3** | **H4** | **H5** | **H6** | **H7** | **H8** | **H9** |
|---|---|---|---|---|---|---|---|---|---|
| **Neu5Ac9'** | -[a] | - | 2.77 – 2.67 (m, 2H), 1.76 (t, $J$ = 12.2 Hz, 1H) | 3.70 | 3.84 | N/R | N/R | N/R | N/R |

**$^{13}C$ (150 MHz, $D_2O$): δ (ppm)**

| | **C1** |
|---|---|
| **GlcNAc1** | 78.10 |
| **GlcNAc2** | 101.20 |
| **Man3** | 100.39 |
| **Man4** | 99.49 |
| **Man4'** | 97.02 |
| **GlcNAc5** | 99.40 |
| **GlcNAc5'** | 99.40 |
| **Gal6** | 102.92 |
| **Gal6'** | 102.87 |
| **GlcNAc7'** | 102.52 |
| **Gal8'** | 103.40 |

| | **C1** | **C2** | **C3** | **C4** | **C5** | **C6** | **C7** | **C8** | **C9** |
|---|---|---|---|---|---|---|---|---|---|
| **Neu5Ac9'** | N/R | 100.07 | 40.02 | 68.15 | 51.85 | N/R | N/R | N/R | N/R |

| **Signal** | **Proton** | **Carbon** |
|---|---|---|
| **NH<u>C</u>(O)CH$_3$** | - | 174.86, 174.70, 174.64, 174.57 |
| **NHC(O)<u>CH$_3$</u>** | 2.32 – 1.89 (m, 18H) | 22.29, 22.26, 22.18, 21.99, 21.93 |
| **Aromatic** | 7.63 – 7.34 (m, 5H) | 136.23, 128.71, 128.33, 127.73 |
| **<u>CH$_2$</u>-Ph** | 5.21 – 5.12 (m, 2H) | 67.04 |
| **NH-<u>C</u>OO-** | - | 157.74 |
| **NH-<u>CH</u>-COOH** | 4.48 | 51.85 |
| **NH-CH-<u>C</u>OOH** | - | N/R |
| **C(O)-<u>CH$_2$</u>-CH** | 2.88 (dd, $J$ = 15.8, 4.4 Hz, 1H), 2.77 – 2.67 (m, 2H) | 37.92 |
| **<u>C</u>(O)-CH$_2$-CH** | - | 173.42 |

[a] Not applicable

[b] Not reported

ESI TOF-MS *m/z* calculated for $C_{99}H_{154}N_8O_{68}$, $[M-2H]^{2-}$: 1271.4425, found 1271.439.

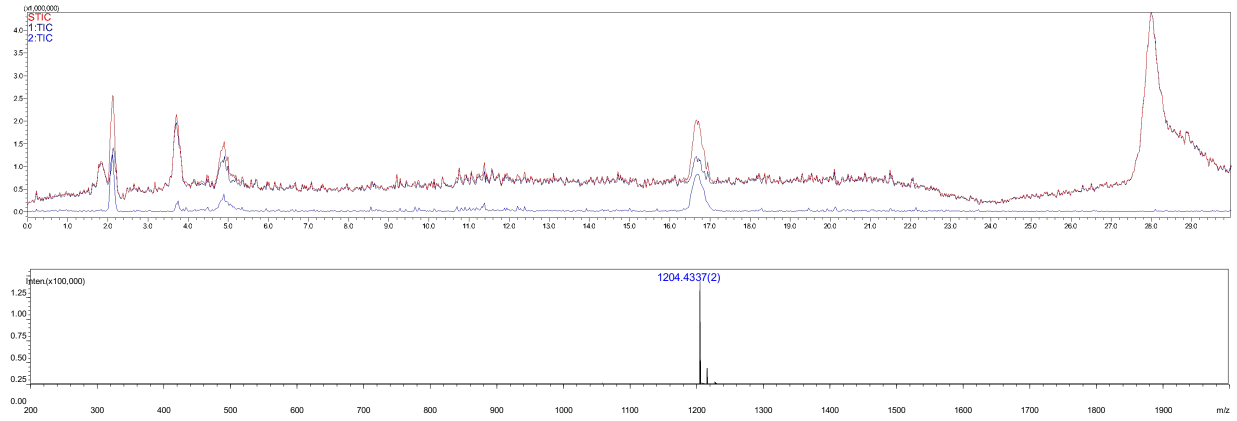

**Figure S27.** Analytical HPLC-MS chromatogram of compound **68**. The retention time = 16.7 min.

#### Compound 69

**69** was synthesized from **25** (3 mg, 1.0 eq) using the general procedures **2.2 e**, **h**, **c** for the installation of α2,6-sialic acid with ST6Gal1, converting $GlcNH_2$ to GlcNAc with AcOSu, installation of β1,4-Gal with B4GalT1 to give **66**, which was purified using the described two-stage purification system **(2.2 j)**. The Cbz protecting group of **66** was removed by hydrogentation (**2.2 i**) to give final product **69** as a white fluffy solid (1.0 mg, 85% yield over four steps). ESI TOF-MS *m/z* calculated for $C_{105}H_{171}N_9O_{76}$, $[M-2H]^{2-}$: 1386.9902, found 1386.9943.

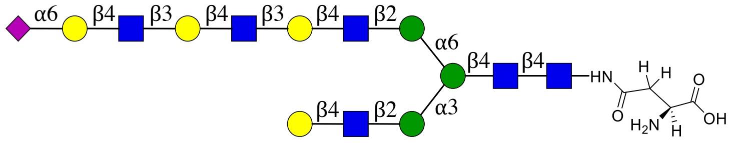

**69**

NMR and MS analysis of compound **66**

**$^1$H (600 MHz, $D_2O$): δ (ppm)**

| | H1 | H2 | H3 | H4 | H5 | H6 |
|---|---|---|---|---|---|---|
| **GlcNAc1** | 5.01 (d, *J* = 9.8 Hz, 1H) | 3.78 | 3.70 | 3.61 | 3.52 | N/R[b] |
| **GlcNAc2** | 4.57 (d, *J* = 8.2 Hz, 1H) | 3.76 | 3.77 | N/R | N/R | N/R |
| **Man3** | 4.73 (s, 2H) | 4.22 (d, *J* = 2.7 Hz, 1H) | 3.74 | N/R | N/R | 3.93, 3.75 |
| **Man4** | 5.08 | 4.18 – 4.14 (m, 1H) | 3.87 | N/R | N/R | N/R |
| **Man4'** | 4.89 (s, 1H) | 4.07 | 3.85 | N/R | N/R | N/R |
| **GlcNAc5** | 4.55 (d, *J* = 7.8 Hz, 2H) | 3.71 | N/R | N/R | N/R | N/R |
| **GlcNAc5'** | 4.55 (d, *J* = 7.8 Hz, 2H) | 3.71 | N/R | N/R | N/R | N/R |
| **Gal6** | 4.48 – 4.37 (m, 4H) | 3.56 | N/R | N/R | N/R | N/R |

| | | | | | | |
|---|---|---|---|---|---|---|
| **Gal6'** | 4.48 – 4.37 (m, 4H) | 3.50 | 3.70 | 4.13 (d, *J* = 3.1 Hz, 2H) | | N/R |
| **GlcNAc7'** | 4.70 (d, *J* = 7.6 Hz, 1H) | 3.77 | N/R | N/R | N/R | N/R |
| **Gal8'** | 4.48 – 4.37 (m, 4H) | 3.50 | 3.70 | 4.13 (d, *J* = 3.1 Hz, 2H) | | N/R |
| **GlcNAc9'** | 4.66 (d, *J* = 8.3 Hz, 1H) | 3.77 | N/R | N/R | N/R | N/R |
| **Gal10'** | 4.48 – 4.37 (m, 4H) | 3.56 | N/R | N/R | N/R | N/R |

| | **H1** | **H2** | **H3** | **H4** | **H5** | **H6** | **H7** | **H8** | **H9** |
|---|---|---|---|---|---|---|---|---|---|
| **Neu5Ac9'** | -[a] | - | 2.64 (dd, *J* = 12.4, 4.7 Hz, 1H), 1.69 (t, *J* = 12.2 Hz, 1H) | 3.62 | 3.77 | N/R | N/R | N/R | N/R |

**$^{13}C$ (150 MHz, $D_2O$): δ (ppm)**

| | **C1** |
|---|---|
| **GlcNAc1** | 78.02 |
| **GlcNAc2** | 101.15 |
| **Man3** | 100.34 |
| **Man4** | 99.45 |
| **Man4'** | 96.95 |
| **GlcNAc5** | 99.35 |
| **GlcNAc5'** | 96.35 |
| **Gal6** | 102.81 |
| **Gal6'** | 102.81 |
| **GlcNAc7'** | 102.50 |
| **Gal8'** | 102.81 |
| **GlcNAc9'** | 102.69 |
| **Gal10'** | 103.38 |

| | **C1** | **C2** | **C3** | **C4** | **C5** | **C6** | **C7** | **C8** | **C9** |
|---|---|---|---|---|---|---|---|---|---|
| **Neu5Ac9'** | N/R | N/R | 40.00 | 68.13 | 51.79 | N/R | N/R | N/R | N/R |

| **Signal** | **Proton** | **Carbon** |
|---|---|---|
| **NH<u>C</u>(O)CH3** | - | 174.81, 174.62, 174.54 |
| **NHC(O)<u>CH3</u>** | 2.17 – 1.81 (m, 33H) | 22.24, 22.19, 22.13, 22.10, 21.94, 21.90 |
| **Aromatic** | 7.48 – 7.31 (m, 5H) | 136.24, 128.66, 128.26, 127.75 |
| **<u>CH2</u>-Ph** | 5.14 – 5.04 (m, 2H) | 66.86 |
| **NH-<u>C</u>OO-** | - | N/R |
| **NH-<u>CH</u>-COOH** | 4.29 (dd, *J* = 9.2, 4.1 Hz, 1H) | 52.92 |
| **NH-CH-<u>C</u>OOH** | - | 177.55 |
| **C(O)-<u>CH2</u>-CH** | 2.78 (dd, *J* = 15.7, 4.2 Hz, 1H), 2.55 (dd, *J* = 15.4, 9.5 Hz, 1H) | 38.46 |

| | | |
|---|---|---|
| **C(O)-CH$_2$-CH** | - | 173.46 |

[a] Not applicable

[b] Not reported

ESI TOF-MS *m/z* calculated for $C_{113}H_{177}N_9O_{78}$, $[M-2H]^{2-}$: 1454.0086, found 1453.9998.

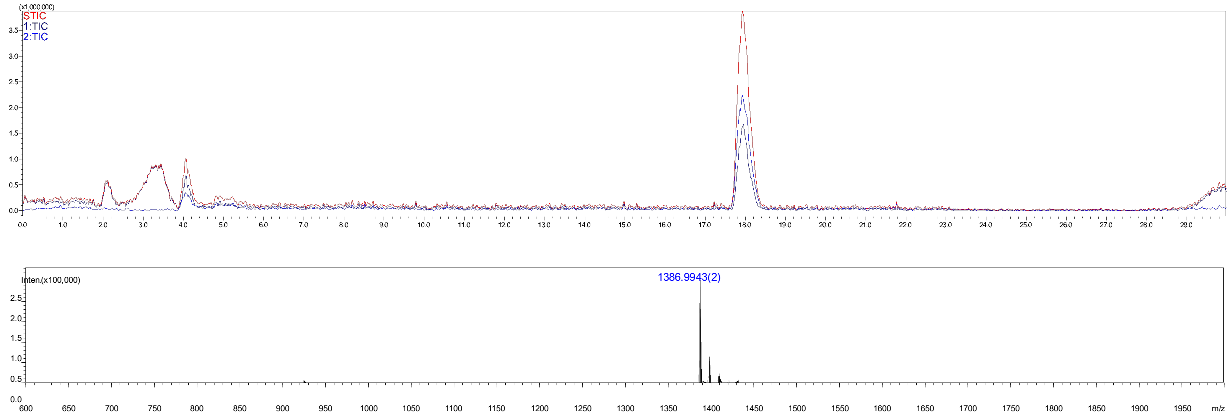


**Figure S28.** Analytical HPLC-MS chromatogram of compound **69**. The retention time = 18.0 min.

#### Compound 70b

**70b** was prepared from **65** (1.0 mg, 1.0 eq) using the general procedures **2.2 e** for the installation of α2, 6-sialic acid with ST6Gal1 to give the intermediate product **70a**, which was purified using the described two-stage purification system **(2.2 j)**. The Cbz protecrting group of **70a** was removed by hydrogentation (**2.2 i**) to give final product **70b** as a white fluffy solid (1.0 mg, 98% yield over two steps). ESI TOF-MS *m/z* calculated for $C_{102}H_{165}N_9O_{74}$, $[M-2H]^{2-}$: 1349.9718, found 1349.9790.

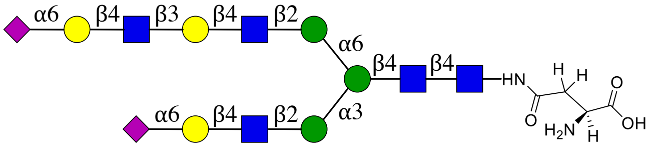


**70**

NMR and MS analysis of compound **70a**:

**$^1$H (600 MHz, D$_2$O): δ (ppm)**

| | H1 | H2 | H3 | H4 | H5 | H6 |
|---|---|---|---|---|---|---|
| **GlcNAc1** | 5.05 (d, *J* = 9.1 Hz, 1H) | 3.82 | 3.74 | 3.65 | 3.56 | N/R[b] |
| **GlcNAc2** | 4.63 – 4.55 (m, 3H) | 3.77 | 3.79 | N/R | N/R | N/R |
| **Man3** | 4.77 | 4.25 (s, 1H) | 3.78 | N/R | N/R | 3.96, 3.78 |
| **Man4** | 5.13 | 4.20 (s, 1H) | 3.90 | N/R | N/R | N/R |
| **Man4'** | 4.92 (s, 1H) | 4.11 (s, 1H) | 3.89 | N/R | N/R | N/R |

| | | | | | | |
|---|---|---|---|---|---|---|
| **GlcNAc5** | 4.63 – 4.55 (m, 3H) | 3.75 | N/R | N/R | N/R | N/R |
| **GlcNAc5'** | 4.63 – 4.55 (m, 3H) | 3.75 | N/R | N/R | N/R | N/R |
| **Gal6** | 4.50 – 4.38 (m, 3H) | 3.54 | N/R | N/R | N/R | N/R |
| **Gal6'** | 4.50 – 4.38 (m, 3H) | 3.59 | 3.73 | 4.16 (s, 1H) | N/R | N/R |
| **GlcNAc7'** | 4.73 | 3.80 | N/R | N/R | N/R | N/R |
| **Gal8'** | 4.50 – 4.38 (m, 3H) | 3.54 | N/R | N/R | N/R | N/R |

| | **H1** | **H2** | **H3** | **H4** | **H5** | **H6** | **H7** | **H8** | **H9** |
|---|---|---|---|---|---|---|---|---|---|
| **Neu5Ac7** | -[a] | - | 2.67 (d, $J$ = 12.4 Hz, 3H),<br>1.72 (t, $J$ = 12.0 Hz, 2H) | 3.65 | 3.81 | N/R | N/R | N/R | N/R |
| **Neu5Ac9'** | - | - | 2.67 (d, $J$ = 12.4 Hz, 3H),<br>1.72 (t, $J$ = 12.0 Hz, 2H) | 3.65 | 3.81 | N/R | N/R | N/R | N/R |

**$^{13}$C (150 MHz, $D_2O$): δ (ppm)**

| | **C1** |
|---|---|
| **GlcNAc1** | 78.10 |
| **GlcNAc2** | 101.22 |
| **Man3** | 100.35 |
| **Man4** | 99.48 |
| **Man4'** | 97.03 |
| **GlcNAc5** | 99.27 |
| **GlcNAc5'** | 99.44 |
| **Gal6** | 103.47 |
| **Gal6'** | 102.94 |
| **GlcNAc7'** | 102.52 |
| **Gal8'** | 103.40 |

| | **C1** | **C2** | **C3** | **C4** | **C5** | **C6** | **C7** | **C8** | **C9** |
|---|---|---|---|---|---|---|---|---|---|
| **Neu5Ac7** | N/R | N/R | 40.01 | 68.17 | 51.84 | N/R | N/R | N/R | N/R |
| **Neu5Ac9'** | N/R | N/R | 40.01 | 68.17 | 51.84 | N/R | N/R | N/R | N/R |

| **Signal** | **Proton** | **Carbon** |
|---|---|---|
| **NH<u>C</u>(O)CH3** | - | 174.81, 174.65, 174.62 |
| **NHC(O)<u>CH3</u>** | 2.25 – 1.86 (m, 21H) | 22.38, 22.29, 22.25, 22.17, 21.99, 21.93 |
| **Aromatic** | 7.56 – 7.30 (m, 5H) | 128.70, 128.32, 127.74 |
| **<u>CH2</u>-Ph** | 5.20 – 5.08 (m, 2H) | 67.01 |
| **NH-<u>C</u>OO-** | - | N/R |
| **NH-<u>CH</u>-COOH** | N/R | N/R |
| **NH-CH-<u>C</u>OOH** | - | N/R |
| **C(O)-<u>CH2</u>-CH** | 2.83 (s, 1H),<br>2.67 (d, $J$ = 12.4 Hz, 3H) | 67.07 |
| **<u>C</u>(O)-CH2-CH** | - | N/R |

[a] Not applicable

[b] Not reported

ESI TOF-MS *m/z* calculated for $C_{110}H_{171}N_9O_{76}$, $[M-2H]^{2-}$: 1416.9902, found 1416.9896.

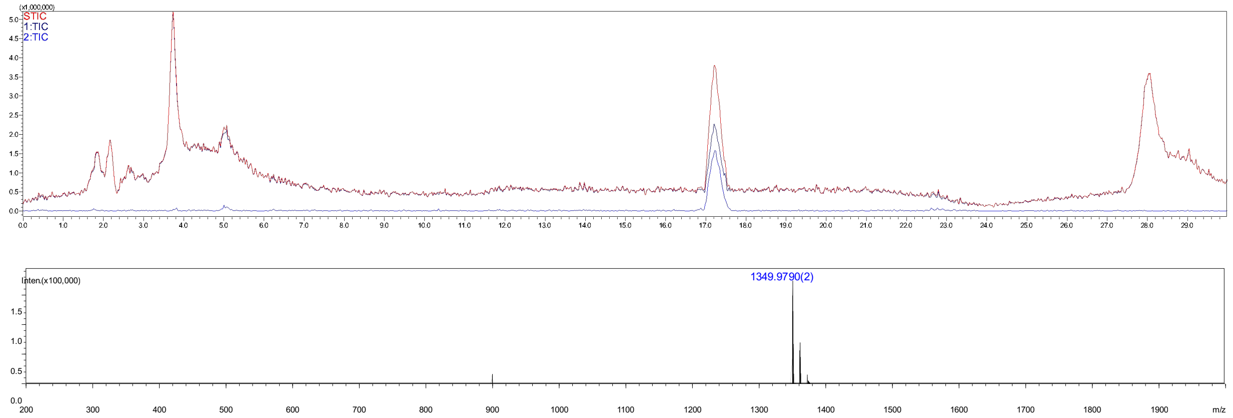


**Figure S29.** Analytical HPLC-MS chromatogram of compound **70b**. The retention time = 17.2 min.

#### Compound 71b

**71b** was prepared from **66** (1 mg, 1.0 eq) using the general procedures **2.2 e** for the installation of α2, 6-sialic acid with ST6Gal1 to give intermediate product **71a** which was purified using the described two-stage purification system **(2.2 j)**. The Cbz protecrting group of **71a** was removed by hydrogentation (**2.2 i**) to give final product **71b** as a white fluffy solid (0.8 mg, 75% yield over two steps). ESI TOF-MS *m/z* calculated for $C_{116}H_{188}N_{10}O_{84}$, $[M-2H]^{2-}$: 1532.5379, found 1532.5241.

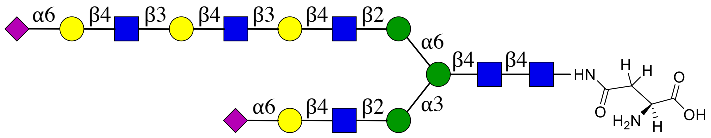


**71**

NMR and MS analysis of compound **71a**

**$^1$H (600 MHz, $D_2O$): δ (ppm)**

| | H1 | H2 | H3 | H4 | H5 | H6 |
|---|---|---|---|---|---|---|
| **GlcNAc1** | 5.08 (d, *J* = 9.7 Hz, 1H) | 3.86 | 3.77 | 3.68 | 3.59 | N/R[b] |
| **GlcNAc2** | 4.67 – 4.58 (m, 3H) | 3.78 | 3.82 | N/R | N/R | N/R |
| **Man3** | 4.80 | 4.29 (d, *J* = 2.5 Hz, 1H) | 3.82 | N/R | N/R | 3.99, 3.83 |
| **Man4** | 5.17 | 4.23 (d, *J* = 3.4 Hz, 1H) | 3.93 | N/R | N/R | N/R |
| **Man4'** | 4.96 (s, 1H) | 4.14 (d, *J* = 3.6 Hz, 1H) | 3.92 | N/R | N/R | N/R |

| | | | | | | |
|---|---|---|---|---|---|---|
| **GlcNAc5** | 4.67 – 4.58 (m, 3H) | 3.78 | N/R | N/R | N/R | N/R |
| **GlcNAc5'** | 4.67 – 4.58 (m, 3H) | 3.78 | N/R | N/R | N/R | N/R |
| **Gal6** | 4.57 – 4.44 (m, 5H) | 3.62 | N/R | N/R | N/R | N/R |
| **Gal6'** | 4.57 – 4.44 (m, 5H) | 3.57 | 3.77 | 4.19 (d, $J$ = 3.1 Hz, 2H) | N/R | N/R |
| **GlcNAc7'** | 4.75 – 4.71 (m, 2H) | 3.84 | N/R | N/R | N/R | N/R |
| **Gal8'** | 4.57 – 4.44 (m, 5H) | 3.57 | 3.77 | 4.19 (d, $J$ = 3.1 Hz, 2H) | N/R | N/R |
| **GlcNAc9'** | 4.75 – 4.71 (m, 2H) | 3.84 | N/R | N/R | N/R | N/R |
| **Gal10'** | 4.57 – 4.44 (m, 5H) | 3.62 | N/R | N/R | N/R | N/R |

| | **H1** | **H2** | **H3** | **H4** | **H5** | **H6** | **H7** | **H8** | **H9** |
|---|---|---|---|---|---|---|---|---|---|
| **Neu5Ac7** | -[a] | - | 2.70 (dd, $J$ = 13.3, 4.0 Hz, 2H), 1.76 (t, $J$ = 12.1 Hz, 2H) | 3.70 | 3.84 | N/R | N/R | N/R | N/R |
| **Neu5Ac11'** | - | - | 2.70 (dd, $J$ = 13.3, 4.0 Hz, 2H), 1.76 (t, $J$ = 12.1 Hz, 2H) | 3.70 | 3.84 | N/R | N/R | N/R | N/R |

**$^{13}$C (150 MHz, $D_2O$): δ (ppm)**

| | **C1** |
|---|---|
| **GlcNAc1** | 78.12 |
| **GlcNAc2** | 101.22 |
| **Man3** | 100.34 |
| **Man4** | 99.52 |
| **Man4'** | 97.03 |
| **GlcNAc5** | 99.31 |
| **GlcNAc5'** | 99.43 |
| **Gal6** | 103.48 |
| **Gal6'** | 102.91 |
| **GlcNAc7'** | 102.54 |
| **Gal8'** | 102.91 |
| **GlcNAc9'** | 102.70 |
| **Gal10'** | 103.48 |

| | **C1** | **C2** | **C3** | **C4** | **C5** | **C6** | **C7** | **C8** | **C9** |
|---|---|---|---|---|---|---|---|---|---|
| **Neu5Ac7** | N/R | N/R | 39.97 | 68.14 | 51.83 | N/R | N/R | N/R | N/R |
| **Neu5Ac11'** | N/R | N/R | 39.97 | 68.14 | 51.83 | N/R | N/R | N/R | N/R |

| **Signal** | **Proton** | **Carbon** |
|---|---|---|
| **NH<u>C</u>(O)CH3** | - | 174.89, 174.70, 174.61, 174.54 |
| **NHC(O)<u>CH3</u>** | 2.33 – 1.88 (m, 24H) | 22.39, 22.29, 22.24, 22.18, 22.15, 21.99, 21.92 |
| **Aromatic** | 7.56 – 7.37 (m, 5H) | 128.71, 128.34, 127.72 |
| **<u>CH2</u>-Ph** | 5.21 – 5.13 (m, 2H) | 67.13 |
| **NH-<u>C</u>OO-** | - | N/R |

| | | |
|---|---|---|
| **NH-<u>CH</u>-COOH** | 4.57 – 4.44 (m, 5H) | N/R |
| **NH-CH-<u>C</u>OOH** | - | N/R |
| **C(O)-<u>CH₂</u>-CH** | 2.88 (d, *J* = 16.4 Hz, 1H),<br>2.76 (dd, *J* = 15.5, 8.0 Hz, 1H) | 37.70 |
| **<u>C</u>(O)-CH₂-CH** | - | N/R |

[a] Not applicable

[b] Not reported

ESI TOF-MS *m/z* calculated for $C_{124}H_{194}N_{10}O_{86}$, $[M-2H]^{2-}$: 1599.5563, found 1599.5561.

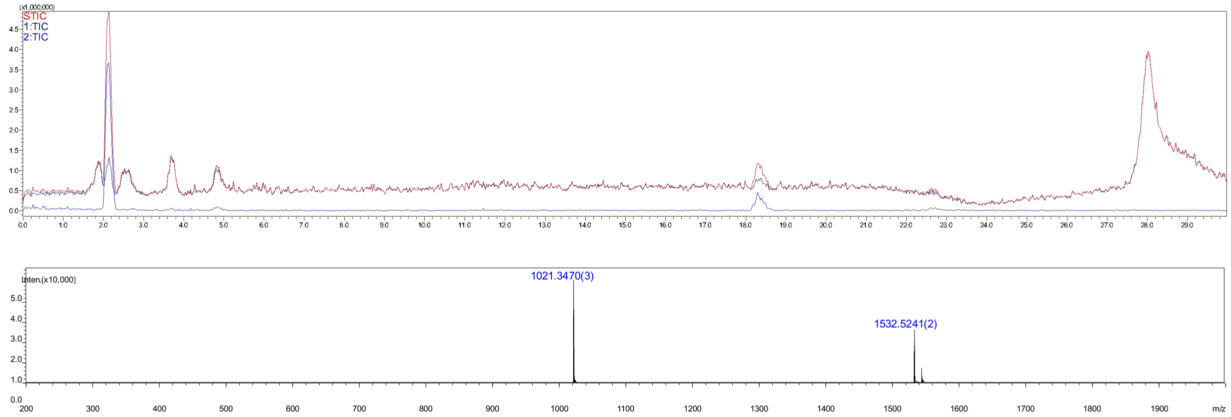


**Figure S30.** Analytical HPLC-MS chromatogram of compound **71b**. The retention time = 18.5 min.

#### Compound 72

**72** was prepared by install a β1, 4-Gal in the extended arm of **10** (10 mg, 1.0 eq) according to the general procedure **2.2 c** with B4GalT1. The product was was purified using the described two-stage purification system **(2.2 j)** providing **72** as a white fluffy solid (12 mg, 100%).

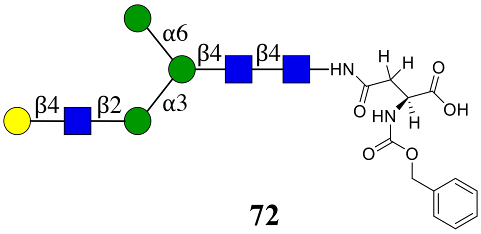


**72**

**$^1$H (600 MHz, D₂O): δ (ppm)**

| | H1 | H2 | H3 | H4 | H5 | H6 |
|---|---|---|---|---|---|---|
| **GlcNAc1** | 5.06 (d, *J* = 9.5 Hz, 1H) | 3.83 | 3.74 | 3.66 | 3.56 | 3.81, 3.63 |
| **GlcNAc2** | 4.62 (d, *J* = 7.9 Hz, 1H) | 3.79 | 3.75 | 3.75 | 3.62 | 3.89, 3.76 |

| | | | | | | |
|---|---|---|---|---|---|---|
| **Man3** | 4.79 | 4.27 (s, 1H) | 3.78 | 3.78 | 3.67 | 3.93, 3.81 |
| **Man4** | 5.13 | 4.20 (s, 1H) | 3.91 | 3.51 | 3.75 | 3.94, 3.63 |
| **Man4'** | 4.93 (s, 1H) | 3.98 | 3.88 | 3.66 | 3.66 | 3.90, 3.77 |
| **GlcNAc5** | 4.59 (d, *J* = 7.3 Hz, 1H) | 3.75 | 3.74 | 3.74 | 3.59 | 3.98, 3.85 |
| **Gal6** | 4.48 (d, *J* = 7.8 Hz, 1H) | 3.55 | 3.68 | 3.94 | 3.74 | 3.79, 3.75 |

**$^{13}C$ (150 MHz, $D_2O$): δ (ppm)**

| | **C1** | **C2** | **C3** | **C4** | **C5** | **C6** |
|---|---|---|---|---|---|---|
| **GlcNAc1** | 78.04 | 53.64 | 72.68 | 78.50 | 76.11 | 59.77 |
| **GlcNAc2** | 101.16 | 54.78 | 65.82 | 79.59 | 74.28 | 59.89 |
| **Man3** | 100.32 | 70.11 | 80.30 | 71.87 | 74.07 | 65.76 |
| **Man4** | 99.46 | 76.29 | 69.30 | 67.20 | 73.47 | 61.62 |
| **Man4'** | 99.55 | 69.79 | 70.32 | 66.69 | 72.61 | 60.88 |
| **GlcNAc5** | 99.36 | 54.78 | 71.87 | 78.39 | 74.65 | 59.89 |
| **Gal6** | 102.83 | 70.88 | 72.42 | 68.45 | 75.26 | 60.93 |

| **Signal** | **Proton** | **Carbon** |
|---|---|---|
| **NHC(O)CH3** | -[a] | 174.62 |
| **NHC(O)CH3** | 2.09 (s, 3H), 2.06 (s, 3H), 1.93 (s, 3H) | 22.26, 22.10, 21.90 |
| **Aromatic** | 7.59 – 7.26 (m, 5H) | 136.23, 128.68, 128.28, 127.74 |
| **CH2-Ph** | 5.19 – 5.09 (m, 2H) | 66.94 |
| **NH-COO-** | - | 157.71 |
| **NH-CH-COOH** | 4.43 – 4.37 (m, 1H) | 52.42 |
| **NH-CH-COOH** | - | N/R[b] |
| **C(O)-CH2-CH** | 2.84 (d, *J* = 14.4 Hz, 2H)<br>2.64 (dd, *J* = 15.0, 9.5 Hz, 2H) | 38.21 |
| **C(O)-CH2-CH** | - | 173.39 |

[a] Not applicable

[b] Not reported

ESI TOF-MS *m/z* calculated for $C_{60}H_{91}N_5O_{40}$, $[M-2H]^{2-}$: 760.7626, found 760.7662.

##### Compound 73

**73** was prepared from **72** (8 mg, 1.0 eq) using the general procedures **2.2 d** and **c** for the installation of GlcNAc and Gal moieties. The product was purified using the described two-stage purification system **(2.2 j)** providing **73** as a white fluffy solid (8.5 mg, 86% yield over two steps).

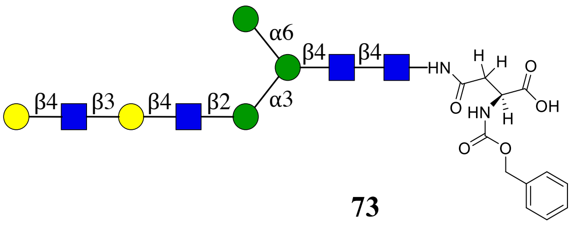


**73**

**$^1$H (600 MHz, $D_2O$): δ (ppm)**

| | H1 | H2 | H3 | H4 | H5 | H6 |
|---|---|---|---|---|---|---|
| **GlcNAc1** | 5.06 (d, *J* = 9.8 Hz, 1H) | 3.83 | 3.74 | 3.66 | 3.56 | N/R[b] |
| **GlcNAc2** | 4.62 (d, *J* = 8.0 Hz, 1H) | 3.80 | 3.77 | 3.75 | 3.62 | N/R |
| **Man3** | 4.79 | 4.26 (d, *J* = 2.4 Hz, 1H) | 3.78 | 3.78 | 3.67 | 3.93, 3.81 |
| **Man4** | 5.13 | 4.20 (d, *J* = 3.4 Hz, 1H) | 3.89 | 3.51 | 3.75 | 3.93, 3.63 |
| **Man4'** | 4.93 (s, 1H) | 3.98 | 3.89 | 3.65 | 3.65 | N/R |
| **GlcNAc5** | 4.58 (d, *J* = 7.4 Hz, 1H) | 3.74 | 3.74 | 3.73 | 3.59 | N/R |
| **Gal6** | 4.46 (d, *J* = 7.9 Hz, 1H) | 3.59 | 3.73 | 4.17 (d, *J* = 3.2 Hz, 1H) | N/R | N/R |
| **GlcNAc7** | 4.71 (d, *J* = 8.3 Hz, 1H) | 3.82 | N/R | 3.75 | N/R | N/R |
| **Gal8** | 4.49 (d, *J* = 7.9 Hz, 1H) | 3.55 | 3.68 | 3.94 | 3.74 | N/R |

**$^{13}$C (150 MHz, $D_2O$): δ (ppm)**

| | C1 | C2 | C3 | C4 | C5 | C6 |
|---|---|---|---|---|---|---|
| **GlcNAc1** | 78.03 | 53.64 | 72.69 | 78.50 | 76.11 | N/R |
| **GlcNAc2** | 101.16 | 54.79 | 65.83 | 79.59 | 74.28 | N/R |
| **Man3** | 100.32 | 70.11 | 80.30 | 71.86 | 74.07 | 65.76 |
| **Man4** | 99.46 | 76.29 | 69.30 | 67.20 | 73.47 | 61.62 |
| **Man4'** | 99.55 | 69.79 | 70.32 | 66.69 | 72.61 | N/R |
| **GlcNAc5** | 99.38 | 54.74 | 71.89 | 78.42 | 74.64 | N/R |
| **Gal6** | 102.87 | 66.87 | 81.97 | 68.23 | N/R | N/R |
| **GlcNAc7** | 1.2.67 | 55.11 | N/R | 78.08 | N/R | N/R |
| **Gal8** | 102.78 | 70.89 | 72.43 | 68.47 | 75.27 | N/R |

| Signal | Proton | Carbon |
|---|---|---|
| **NH<u>C</u>(O)$CH_3$** | -[a] | 174.83, 174.62 |
| **NHC(O)<u>$CH_3$</u>** | 2.22 – 1.83 (m, 12H) | 22.25, 22.10, 21.91 |
| **Aromatic** | 7.53 – 7.35 (m, 5H) | 136.24, 128.68, 128.28, 127.75 |
| **<u>$CH_2$</u>-Ph** | 5.18 – 5.09 (m, 2H) | 66.92 |
| **NH-<u>C</u>OO-** | - | 157.72 |
| **NH-<u>CH</u>-COOH** | 4.39 (s, 1H) | N/R |
| **NH-CH-<u>C</u>OOH** | - | N/R |
| **C(O)-<u>$CH_2$</u>-CH** | 2.83 (d, *J* = 15.2 Hz, 1H)<br>2.63 (d, *J* = 6.8 Hz, 1H) | 38.31 |

| C(O)-CH2-CH | - | N/R |
|---|---|---|

[a] Not applicable

[b] Not reported

ESI TOF-MS *m/z* calculated for $C_{74}H_{114}N_6O_{50}$, $[M-2H]^{2-}$: 943.3287, found 943.3307.

##### Compound 74

**74** was prepared from **73** (4 mg) using the general procedures **2.2 d** and **c** for the installation of GlcNAc and Gal moieties. The product was purified using the described two-stage purification system **(2.2 j)** providing **74** as a white fluffy solid (4 mg, 83% yield for two steps).

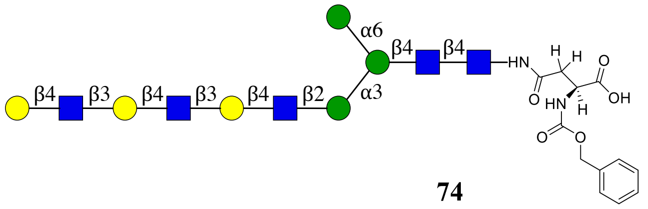


**74**

**$^1$H (600 MHz, $D_2O$): δ (ppm)**

| | H1 | H2 | H3 | H4 | H5 | H6 |
|---|---|---|---|---|---|---|
| **GlcNAc1** | 5.04 (d, *J* = 9.8 Hz, 1H) | 3.82 | 3.73 | 3.65 | 3.55 | N/R[b] |
| **GlcNAc2** | 4.60 (d, *J* = 8.1 Hz, 1H) | 3.80 | 3.76 | 3.73 | 3.61 | N/R |
| **Man3** | 4.78 | 4.25 (s, 1H) | 3.76 | 3.77 | 3.66 | 3.92, 3.80 |
| **Man4** | 5.12 | 4.19 (d, *J* = 3.3 Hz, 1H) | 3.90 | 3.50 | 3.74 | 3.92, 3.61 |
| **Man4'** | 4.92 (s, 1H) | 3.97 | 3.88 | 3.64 | 3.64 | N/R |
| **GlcNAc5** | 4.57 (d, *J* = 7.3 Hz, 1H) | 3.73 | 3.73 | 3.72 | 3.57 | N/R |
| **Gal6** | 4.50 – 4.43 (m, 3H) | 3.58 | 3.72 | 4.17 – 4.13 (m, 2H) | N/R | N/R |
| **GlcNAc7** | 4.72 – 4.67 (m, 2H) | 3.80 | | 3.74 | N/R | N/R |
| **Gal8** | 4.50 – 4.43 (m, 3H) | 3.58 | 3.72 | 4.17 – 4.13 (m, 2H) | N/R | N/R |
| **GlcNAc9** | 4.72 – 4.67 (m, 2H) | 3.80 | | 3.74 | N/R | N/R |
| **Gal10** | 4.50 – 4.43 (m, 3H) | 3.54 | 3.67 | 3.92 | 3.73 | N/R |

**$^{13}$C (150 MHz, $D_2O$): δ (ppm)**

| | C1 | C2 | C3 | C4 | C5 | C6 |
|---|---|---|---|---|---|---|
| **GlcNAc1** | 78.04 | 53.62 | 72.68 | 78.47 | 76.09 | N/R |
| **GlcNAc2** | 101.14 | 54.76 | 65.81 | 79.57 | 74.26 | N/R |
| **Man3** | 100.31 | 70.09 | 80.28 | 71.84 | 74.05 | 65.74 |
| **Man4** | 99.45 | 76.27 | 69.28 | 67.18 | 73.45 | 61.60 |
| **Man4'** | 99.53 | 69.77 | 70.30 | 66.67 | 72.59 | N/R |
| **GlcNAc5** | 99.34 | 54.72 | 71.87 | 78.37 | 74.62 | N/R |
| **Gal6** | 102.92 – 102.59 (m) | 69.86 | 81.96 | 68.22 | N/R | N/R |

| | | | | | | |
|---|---|---|---|---|---|---|
| **GlcNAc7** | 102.67 | 55.05 | N/R | 78.04 | N/R | N/R |
| **Gal8** | 102.92 – 102.59 (m) | 69.86 | 81.96 | 68.22 | N/R | N/R |
| **GlcNAc9** | 102.67 | 55.09 | N/R | 78.04 | N/R | N/R |
| **Gal10** | 102.92 – 102.59 (m) | 70.87 | 72.40 | 68.45 | 75.26 | N/R |

| Signal | Proton | Carbon |
|---|---|---|
| **NHC(O)$CH_3$** | -[a] | 174.81, 174.69, 174.61 |
| **NHC(O)$CH_3$** | 2.24 – 1.77 (m, 15H) | 22.24, 22.08, 21.89 |
| **Aromatic** | 2.24 – 1.77 (m, 5H) | 136.24, 128.66, 128,26, 127.75 |
| **$CH_2$-Ph** | 5.18 – 5.07 (m, 2H) | 66.87 |
| **NH-COO-** | - | 157.68 |
| **NH-CH-COOH** | 4.32 (dd, *J* = 9.5, 4.1 Hz, 1H) | 52.92 |
| **NH-CH-COOH** | - | 177.57 |
| **C(O)-$CH_2$-CH** | 2.81 (dd, *J* = 15.8, 4.1 Hz, 1H), 2.58 (dd, *J* = 15.4, 9.5 Hz, 1H) | 38.47 |
| **C(O)-$CH_2$-CH** | - | 173.60 |

[a] Not applicable

[b] Not reported

ESI TOF-MS *m/z* calculated for $C_{88}H_{137}N_7O_{60}$, $[M-2H]^{2-}$: 1125.8948, found 1125.8906.

#### Compound 75

**75** was prepared from **72** (1 mg) using the general procedure **2.2 i** for the removing of Cbz group with 20% $Pd(OH)_2/C$ and $H_2$. The final product **75** was obtained as a white fluffy solid (0.9 mg, 95%). ESI TOF-MS *m/z* calculated for $C_{52}H_{86}N_5O_{38}$, $[M-H]^-$: 1388.4956, found 1388.4899.

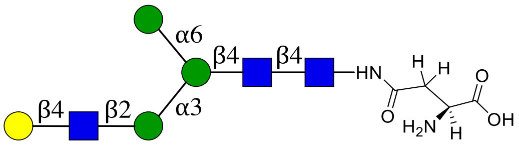


**75**

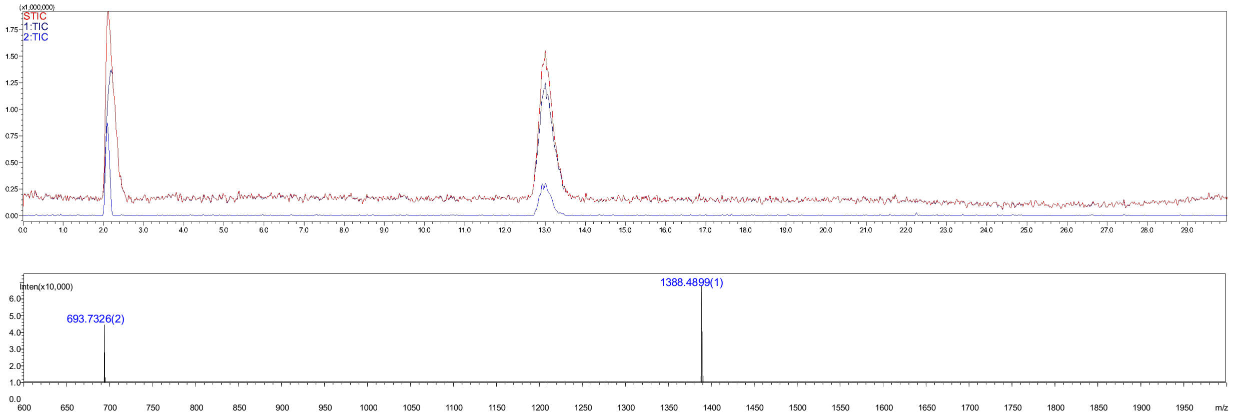

**Figure S31.** Analytical HPLC-MS chromatogram of compound **75**. The retention time = 13.0 min.

#### Compound 77

**77** was synthesized from **72** (1 mg, 1.0 eq) through the installation of α2, 6-Neu5Ac using the general procedure **2.2 e** to give intermediate product **76** which was purified using the described two-stage purification system **(2.2 j)**. The Cbz protecrting group of **76** was removed by hydrogentation (**2.2 i**) to give final product **77** as a white fluffy solid (1 mg, 88% yield over two steps). ESI TOF-MS *m/z* calculated for $C_{63}H_{102}N_6O_{46}$, $[M-2H]^{2-}$: 839.2919, found 839.2950.

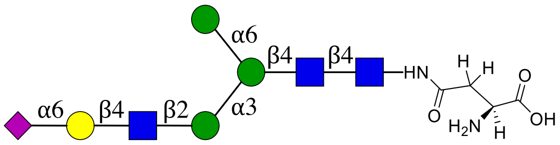

**77**

NMR and MS analysis of compound **76**

**$^1$H (600 MHz, $D_2O$): δ (ppm)**

| | H1 | H2 | H3 | H4 | H5 | H6 |
|---|---|---|---|---|---|---|
| **GlcNAc1** | 5.08 (d, *J* = 9.7 Hz, 1H) | 3.86 | 3.77 | 3.68 | 3.59 | N/R[b] |
| **GlcNAc2** | 4.64 (dd, *J* = 7.9, 2.6 Hz, 4H) | 3.82 | 3.79 | 3.77 | N/R | N/R |
| **Man3** | 4.82 (s, 1H) | 4.29 (d, *J* = 2.8 Hz, 1H) | 3.81 | N/R | N/R | 3.95, 3.84 |
| **Man4** | 5.17 | 4.23 (dd, *J* = 3.3, 1.6 Hz, 1H) | 3.93 | 3.55 | N/R | N/R |
| **Man4'** | 4.95 (d, *J* = 1.7 Hz, 1H) | 4.01 | 3.91 | 3.68 | N/R | N/R |
| **GlcNAc5** | 4.64 (dd, *J* = 7.9, 2.6 Hz, 4H) | 3.79 | | 3.69 | N/R | N/R |
| **Gal6** | 4.48 (d, *J* = 7.9 Hz, 1H) | 3.57 | 3.69 | 3.96 | N/R | N/R |

| | H1 | H2 | H3 | H4 | H5 | H6 | H7 | H8 | H9 |
|---|---|---|---|---|---|---|---|---|---|
| **Neu5Ac7** | -[a] | - | 2.70 (dd, *J* = 12.4, 4.7 Hz, 1H), 1.76 (t, *J* = 12.1 Hz, 1H) | 3.70 | 3.84 | N/R | N/R | N/R | N/R |

**$^{13}C$ (150 MHz, $D_2O$): δ (ppm)**

| | C1 | C2 | C3 | C4 | C5 | C6 |
|---|---|---|---|---|---|---|
| **GlcNAc1** | 78.11 | 53.69 | 72.67 | 78.58 | 76.16 | N/R |
| **GlcNAc2** | 101.21 | 54.57 | 65.86 | 79.60 | N/R | N/R |
| **Man3** | 100.31 | 70.14 | 80.35 | N/R | N/R | 65.82 |
| **Man4** | 99.50 | 76.36 | 69.38 | 67.23 | N/R | N/R |
| **Man4'** | 99.59 | 69.83 | 70.35 | 66.72 | N/R | N/R |
| **GlcNAc5** | 99.29 | 54.84 | N/R | 80.66 | N/R | N/R |
| **Gal6** | 103.47 | 70.69 | N/R | 68.31 | N/R | N/R |

| | C1 | C2 | C3 | C4 | C5 | C6 | C7 | C8 | C9 |
|---|---|---|---|---|---|---|---|---|---|
| **Neu5Ac7** | N/R | 100.08 | 39.99 | 68.13 | 51.83 | N/R | N/R | N/R | N/R |

| Signal | Proton | Carbon |
|---|---|---|
| **NHC(O)CH3** | - | 174.86, 174.71, 174.63 |
| **NHC(O)CH3** | 2.22 – 1.86 (m, 12H) | 22.38, 22.14, 22.00, 21.92 |
| **Aromatic** | 7.54 – 7.37 (m, 5H) | 136.21, 128.71, 128.34, 127.72 |
| **CH2-Ph** | 5.22 – 5.13 (m, 2H) | 67.06 |
| **NH-COO-** | - | 157.76 |
| **NH-CH-COOH** | 4.51 | 51.71 |
| **NH-CH-COOH** | - | N/R |
| **C(O)-CH2-CH** | 2.88 (dd, *J* = 15.7, 4.4 Hz, 1H), 2.77 – 2.73 (m, 1H) | 37.82 |
| **C(O)-CH2-CH** | - | 173.08 |

[a] Not applicable

[b] Not reported

ESI TOF-MS *m/z* calculated for $C_{71}H_{108}N_6O_{48}$, $[M-2H]^{2-}$: 906.3103, found 906.3120.

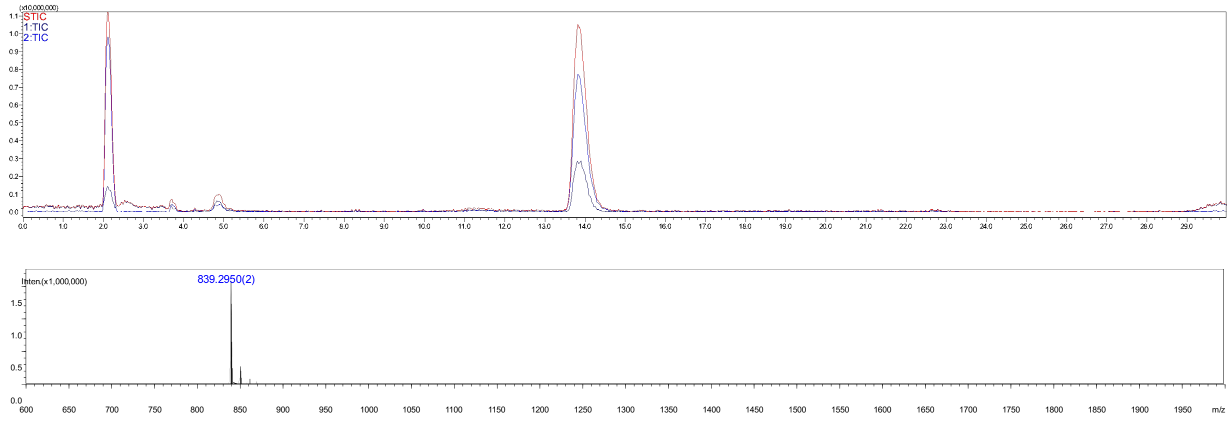


**Figure S32.** Analytical HPLC-MS chromatogram of compound **77**. The retention time = 13.8 min.

**Compound 79**

**79** was synthesized from **72** (2 mg) through the installation of α2,3-Neu5Ac using the general procedure **2.2 f** to give intermediate product **78** after purification using the described two-stage purification system **(2.2 j)**. The Cbz protecrting group of **78** was removed by hydrogentation (**2.2 i**) to give final product **79** as a white fluffy solid (0.5 mg, 46% yield over two steps). ESI TOF-MS *m/z* calculated for $C_{63}H_{102}N_6O_{46}$, $[M-2H]^{2-}$: 839.2919, found 839.2884.

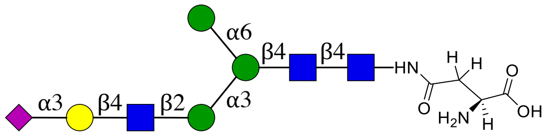


**79**

NMR and MS analysis of compound **78**

**$^1$H (600 MHz, $D_2O$): δ (ppm)**

| | H1 | H2 | H3 | H4 | H5 | H6 |
|---|---|---|---|---|---|---|
| **GlcNAc1** | 5.05 (d, $J$ = 9.8 Hz, 1H) | 3.83 | N/R[b] | 3.65 | 3.56 | N/R |
| **GlcNAc2** | 4.61 (d, $J$ = 8.0 Hz, 1H) | 3.80 | 3.77 | 3.75 | N/R | N/R |
| **Man3** | 4.78 | 4.26 (d, $J$ = 2.4 Hz, 1H) | 3.77 | N/R | N/R | 3.92, 3.81 |
| **Man4** | 5.13 | 4.20 (d, $J$ = 3.5 Hz, 1H) | 3.90 | 3.51 | N/R | N/R |
| **Man4'** | 4.92 (s, 1H) | 3.98 | 3.88 | 3.65 | N/R | N/R |
| **GlcNAc5** | 4.58 (d, $J$ = 7.6 Hz, 1H) | 3.75 | N/R | 3.72 | N/R | N/R |
| **Gal6** | 4.55 (d, $J$ = 7.8 Hz, 1H) | 3.57 | 3.96 | 4.12 | N/R | N/R |

| | H1 | H2 | H3 | H4 | H5 | H6 | H7 | H8 | H9 |
|---|---|---|---|---|---|---|---|---|---|
| **Neu5Ac7** | -[a] | - | 2.76 (dd, $J$ = 12.5, 4.5 Hz, 1H), 1.81 (t, $J$ = 12.1 Hz, 1H) | 3.70 | 3.85 | N/R | N/R | N/R | N/R |

**$^{13}$C (150 MHz, $D_2O$): δ (ppm)**

| | C1 | C2 | C3 | C4 | C5 | C6 |
|---|---|---|---|---|---|---|
| **GlcNAc1** | 78.05 | 53.64 | N/R | 78.51 | 76.12 | N/R |
| **GlcNAc2** | 101.17 | 54.78 | 65.82 | 79.58 | N/R | N/R |
| **Man3** | 100.30 | 70.10 | 80.26 | N/R | N/R | 65.77 |
| **Man4** | 99.46 | 76.30 | 69.30 | 67.19 | N/R | N/R |
| **Man4'** | 99.55 | 69.78 | 70.31 | 66.68 | N/R | N/R |
| **GlcNAc5** | 99.42 | 54.78 | N/R | 75.08 | N/R | N/R |
| **Gal6** | 102.50 | 69.30 | 67.38 | 75.39 | N/R | N/R |

| | C1 | C2 | C3 | C4 | C5 | C6 | C7 | C8 | C9 |
|---|---|---|---|---|---|---|---|---|---|
| **Neu5Ac7** | N/R | 99.74 | 39.54 | 68.26 | 51.59 | N/R | N/R | N/R | N/R |

| Signal | Proton | Carbon |
|---|---|---|
| **NH$\underline{C}$(O)CH$_3$** | - | 174.88, 174.78, 174.60 |
| **NHC(O)$\underline{CH_3}$** | 2.10 – 1.89 (m, 12H) | 22.25, 22.10, 21.95, 21.89 |
| **Aromatic** | 7.49 – 7.37 (m, 5H) | 136.20, 128.67, 128.29, 127.71 |
| **$\underline{CH_2}$-Ph** | 5.17 – 5.10 (m, 2H) | 66.98 |
| **NH-$\underline{C}$OO-** | - | 157.73 |
| **NH-$\underline{CH}$-COOH** | 4.44 | 52.08 |
| **NH-CH-$\underline{C}$OOH** | - | N/R |
| **C(O)-$\underline{CH_2}$-CH** | 2.84 (d, $J$ = 15.4 Hz, 1H), 2.67 (d, $J$ = 14.3 Hz, 1H) | 37.97 |
| **$\underline{C}$(O)-CH$_2$-CH** | - | N/R |

[a] Not applicable

[b] Not reported

ESI TOF-MS *m/z* calculated for $C_{71}H_{108}N_6O_{48}$, $[M-2H]^{2-}$: 906.3103, found 906.3124.

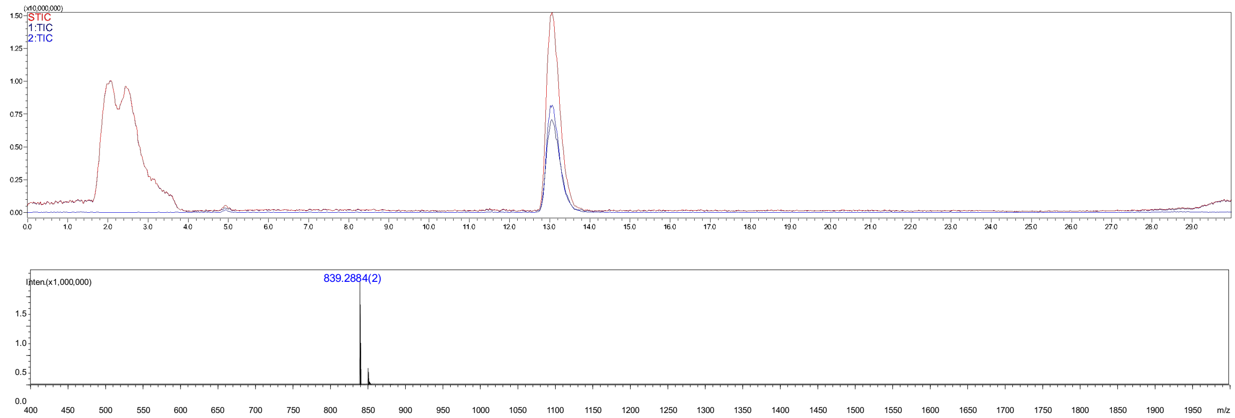


**Figure S33.** Analytical HPLC-MS chromatogram of compound **79**. The retention time = 13.0 min.

#### Compound 80

**80** was prepared from **73** (1.0 mg) using the general procedure **2.2 i** for the removing of Cbz group with 20% $Pd(OH)_2/C$ and $H_2$. The final product **80** was obtained as a white fluffy solid (0.8 mg, 90%). ESI TOF-MS *m/z* calculated for $C_{66}H_{108}N_6O_{48}$, $[M-2H]^{2-}$: 876.3103, found 876.3092.

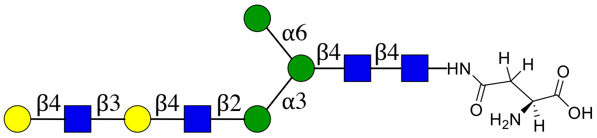


**80**

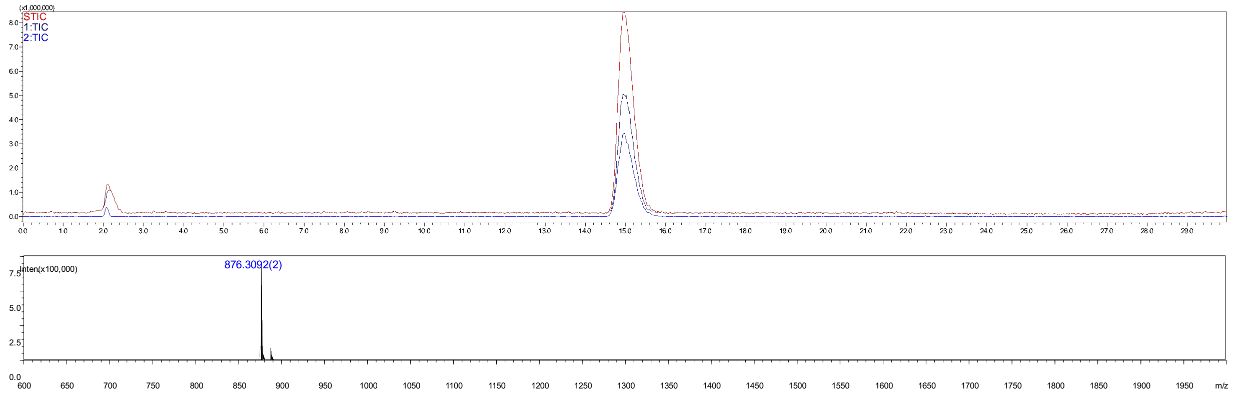

**Figure S34.** Analytical HPLC-MS chromatogram of compound **80**. The retention time = 15.0 min.

#### Compound 82

**82** was synthesized from **73** (1.0 mg) through the installation of α2 6-Neu5Ac using the general procedure **2.2** to give the intermediate product **81** which was purified using the described two-stage purification system **(2.2 j)**. The Cbz protecrting group of **81** was removed by hydrogentation (**2.2 i**) to give final product **82** as a white fluffy solid (0.8 mg, 70% yield over two steps). ESI TOF-MS *m/z* calculated for $C_{77}H_{125}N_7O_{56}$, $[M-2H]^{2-}$: 1021.8580, found 1021.8574.

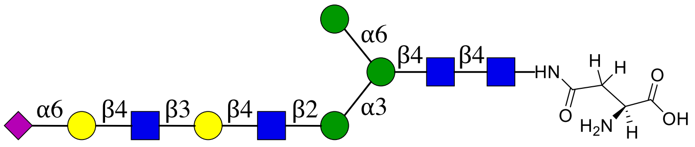


**82**

NMR and MS analysis of compound **81**

**$^{1}$H (600 MHz, $D_2O$): δ (ppm)**

| | H1 | H2 | H3 | H4 | H5 | H6 |
|---|---|---|---|---|---|---|
| **GlcNAc1** | 5.08 (d, *J* = 9.8 Hz, 1H) | 3.86 | | 3.68 | 3.58 | N/R[b] |
| **GlcNAc2** | 4.64 | 3.82 | 3.79 | 3.77 | N/R | N/R |
| **Man3** | 4.81 (s, 1H) | 4.28 (d, *J* = 2.5 Hz, 1H) | 3.80 | N/R | N/R | 3.95, 3.84 |
| **Man4** | 5.15 | 4.24 – 4.21 (m, 1H) | | 3.54 | N/R | N/R |
| **Man4'** | 4.95 (d, *J* = 1.8 Hz, 1H) | 4.01 | N/R | 3.67 | N/R | N/R |
| **GlcNAc5** | 4.61 (d, *J* = 7.4 Hz, 1H) | 3.77 | N/R | 3.75 | N/R | N/R |
| **Gal6** | 4.51 – 4.47 (m, 2H) | 3.62 | 3.77 | 4.19 (d, *J* = 3.2 Hz, 1H) | N/R | N/R |
| **GlcNAc7** | 4.76 | 3.82 | N/R | N/R | N/R | N/R |
| **Gal8** | 4.51 – 4.47 (m, 2H) | 3.57 | N/R | N/R | N/R | N/R |

| | H1 | H2 | H3 | H4 | H5 | H6 | H7 | H8 | H9 |
|---|---|---|---|---|---|---|---|---|---|
| **Neu5Ac9** | -[a] | - | 2.70 (dd, *J* = 12.3, 4.6 Hz, 2H), 1.75 (t, *J* = 12.2 Hz, 1H) | 3.68 | 3.84 | N/R | N/R | N/R | N/R |

**$^{13}C$ (150 MHz, $D_2O$): δ (ppm)**

| | C1 | C2 | C3 | C4 | C5 | C6 |
|---|---|---|---|---|---|---|
| **GlcNAc1** | 78.09 | 53.68 | N/R | 78.56 | 76.15 | N/R |
| **GlcNAc2** | 101.19 | 54.83 | 65.86 | 79.61 | N/R | N/R |
| **Man3** | 100.33 | 70.14 | 80.31 | N/R | N/R | 65.80 |
| **Man4** | 99.50 | 76.36 | N/R | 67.23 | N/R | N/R |
| **Man4'** | 99.58 | 69.82 | N/R | 66.73 | N/R | N/R |
| **GlcNAc5** | 99.42 | 54.76 | N/R | 78.48 | N/R | N/R |
| **Gal6** | 103.40 | 69.92 | 81.95 | 68.24 | N/R | N/R |
| **GlcNAc7** | 102.52 | 54.90 | N/R | N/R | N/R | N/R |
| **Gal8** | 102.90 | 70.69 | N/R | N/R | N/R | N/R |

| | C1 | C2 | C3 | C4 | C5 | C6 | C7 | C8 | C9 |
|---|---|---|---|---|---|---|---|---|---|
| **Neu5Ac9** | N/R | 100.08 | 40.02 | 68.15 | 51.84 | N/R | N/R | N/R | N/R |

| Signal | Proton | Carbon |
|---|---|---|
| **NH<u>C</u>(O)CH3** | - | 174.86, 174.64 |
| **NHC(O)<u>CH3</u>** | 2.25 – 1.86 (m, 15H) | 22.28, 22.24, 22.13, 21.98, 21.92 |
| **Aromatic** | 7.55 – 7.37 (m, 5H) | 136.24, 128.71, 128.32, 127.74 |
| **<u>CH2</u>-Ph** | 5.21 – 5.11 (m, 1H) | 67.01 |
| **NH-<u>C</u>OO-** | - | 157.75 |
| **NH-<u>CH</u>-COOH** | 4.45 (dd, *J* = 8.9, 4.8 Hz, 1H) | 52.11 |
| **NH-CH-<u>C</u>OOH** | - | 176.47 |
| **C(O)-<u>CH2</u>-CH** | 2.87 (dd, *J* = 15.6, 4.3 Hz, 1H), 2.70 (dd, *J* = 12.3, 4.6 Hz, 2H) | 38.05 |
| **<u>C</u>(O)-CH2-CH** | - | 173.44 |

[a] Not applicable

[b] Not reported

ESI TOF-MS *m/z* calculated for $C_{85}H_{131}N_7O_{58}$, $[M-2H]^{2-}$: 1088.8764, found 1088.8674.

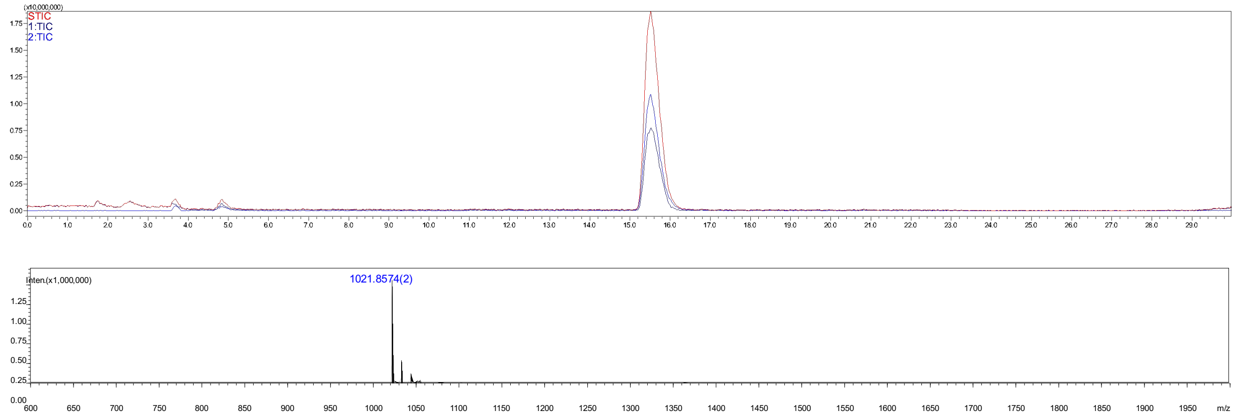

**Figure S35.** Analytical HPLC-MS chromatogram of compound **82**. The retention time = 15.6 min.

#### Compound 84

**84** was synthesized from **73** (2 mg) through the installation of α2,3-Neu5Ac using the general procedure **2.2 f** to give the intermediate product **83** which was purified using the described two-stage purification system **(2.2 j)**. The Cbz protecrting group of **83** was removed by hydrogentation (**2.2 i**) to give final product product **84** as a white fluffy solid (0.4 mg, 44% yield over two steps). ESI TOF-MS *m/z* calculated for $C_{77}H_{125}N_7O_{56}$, $[M-2H]^{2-}$: 1021.8580, found 1021.8552.

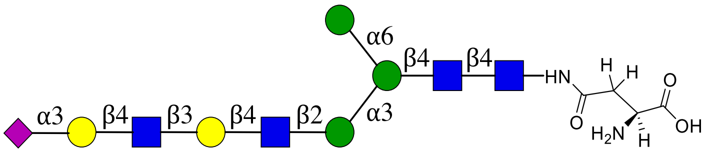


**84**

NMR and MS analysis of compound **83**

**$^1$H (600 MHz, D$_2$O): δ (ppm)**

| | H1 | H2 | H3 | H4 | H5 | H6 |
|---|---|---|---|---|---|---|
| **GlcNAc1** | 5.06 (d, *J* = 9.8 Hz, 1H) | 3.83 | 3.75 | 3.66 | 3.57 | N/R[b] |
| **GlcNAc2** | 4.61 (d, *J* = 8.0 Hz, 1H) | 3.80 | 3.76 | 3.74 | N/R | N/R |
| **Man3** | 4.79 | 4.26 (d, *J* = 2.5 Hz, 1H) | 3.77 | N/R | N/R | 3.92, 3.81 |
| **Man4** | 5.12 | 4.20 (d, *J* = 3.6 Hz, 1H) | 3.90 | 3.51 | N/R | N/R |
| **Man4'** | 4.92 (d, *J* = 1.8 Hz, 1H) | 3.98 | 3.88 | 3.65 | N/R | N/R |
| **GlcNAc5** | 4.60 – 4.55 (m, 2H) | 3.75 | | 3.72 | N/R | N/R |
| **Gal6** | 4.46 (d, *J* = 7.9 Hz, 1H) | 3.59 | 3.73 | 4.17 (d, *J* = 3.2 Hz, 1H) | N/R | N/R |
| **GlcNAc7** | 4.70 (d, *J* = 8.3 Hz, 1H) | 3.81 | N/R | N/R | N/R | N/R |
| **Gal8** | 4.60 – 4.55 (m, 2H) | 3.58 | 3.96 | 4.12 | N/R | N/R |

| | H1 | H2 | H3 | H4 | H5 | H6 | H7 | H8 | H9 |
|---|---|---|---|---|---|---|---|---|---|
| **Neu5Ac9** | -[a] | - | 2.79 – 2.71 (m, 2H),<br>1.81 (t, $J$ = 12.2 Hz, 1H) | 3.70 | 3.86 | N/R | N/R | N/R | N/R |

**$^{13}$C (150 MHz, $D_2O$): δ (ppm)**

| | C1 | C2 | C3 | C4 | C5 | C6 |
|---|---|---|---|---|---|---|
| **GlcNAc1** | 78.08 | 53.67 | N/R | 78.50 | 76.13 | N/R |
| **GlcNAc2** | 101.17 | 54.73 | 65.82 | 79.59 | N/R | N/R |
| **Man3** | 100.31 | 70.11 | 80.28 | N/R | N/R | 65.77 |
| **Man4** | 99.46 | 76.30 | 69.30 | 67.20 | N/R | N/R |
| **Man4'** | 99.55 | 69.78 | 70.31 | 66.69 | N/R | N/R |
| **GlcNAc5** | 99.38 | 54.79 | N/R | 75.09 | N/R | N/R |
| **Gal6** | 102.86 | 69.87 | 81.99 | 68.23 | N/R | N/R |
| **GlcNAc7** | 102.73 | 55.10 | N/R | N/R | N/R | N/R |
| **Gal8** | 102.45 | 69.30 | 67.39 | 75.40 | N/R | N/R |

| | C1 | C2 | C3 | C4 | C5 | C6 | C7 | C8 | C9 |
|---|---|---|---|---|---|---|---|---|---|
| **Neu5Ac9** | N/R | 99.67 | 39.53 | 68.23 | 51.59 | N/R | N/R | N/R | N/R |

| Signal | Proton | Carbon |
|---|---|---|
| **NH<u>C</u>(O)CH3** | - | 174.92, 174.80, 174.61 |
| **NHC(O)<u>CH3</u>** | 2.24 – 1.88 (m, 15H) | 22.25, 22.09, 21.95, 21.88 |
| **Aromatic** | 7.58 – 7.30 (m, 5H) | 136.15, 128.68, 128.32, 127.68 |
| **<u>CH2</u>-Ph** | 5.20 – 5.09 (m, 2H) | 67.08 |
| **NH-<u>C</u>OO-** | - | 157.76 |
| **NH-<u>CH</u>-COOH** | 4.52 (dd, $J$ = 7.9, 4.7 Hz, 1H) | 51.19 |
| **NH-CH-<u>C</u>OOH** | - | N/R |
| **C(O)-<u>CH2</u>-CH** | 2.86 (dd, $J$ = 15.7, 4.6 Hz, 1H),<br>2.79 – 2.71 (m, 2H) | 37.52 |
| **<u>C</u>(O)-CH2-CH** | - | N/R |

[a] Not applicable

[b] Not reported

ESI TOF-MS *m/z* calculated for $C_{85}H_{131}N_7O_{58}$, $[M-2H]^{2-}$: 1088.8764, found 1088.8734.

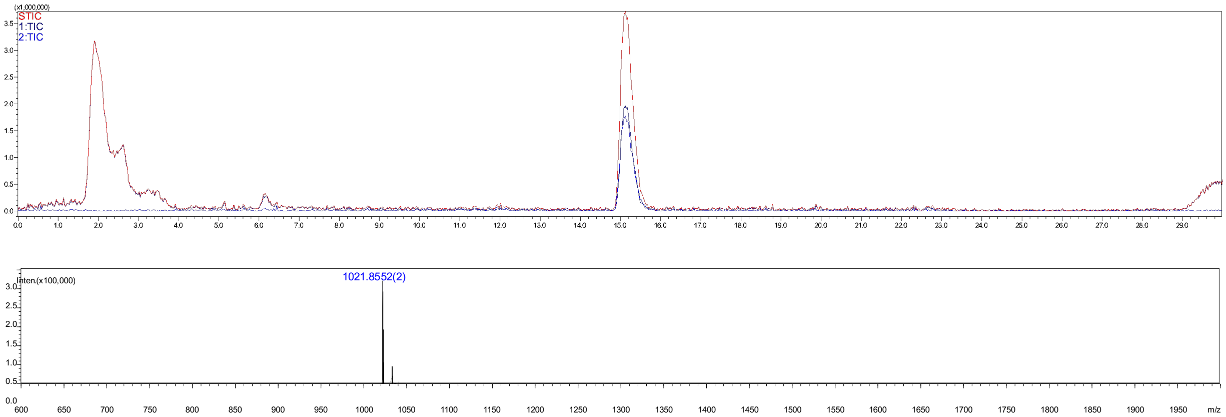


**Figure S36.** Analytical HPLC-MS chromatogram of compound **84**. The retention time = 15.2 min.

#### Compound 85

**85** was prepared from **74** (1 mg) using the general procedure **2.2 i** for the removing of Cbz group with 20% $Pd(OH)_2/C$ and $H_2$. The final product **85** was obtained as a white fluffy solid (0.9 mg, 96%). ESI TOF-MS *m/z* calculated for $C_{80}H_{131}N_7O_{58}$, $[M-2H]^{2-}$: 1058.8764, found 1058.8682.

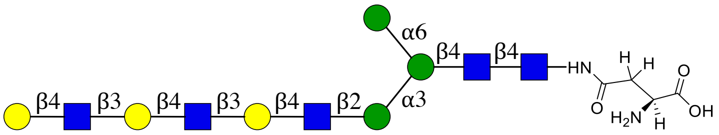


**85**

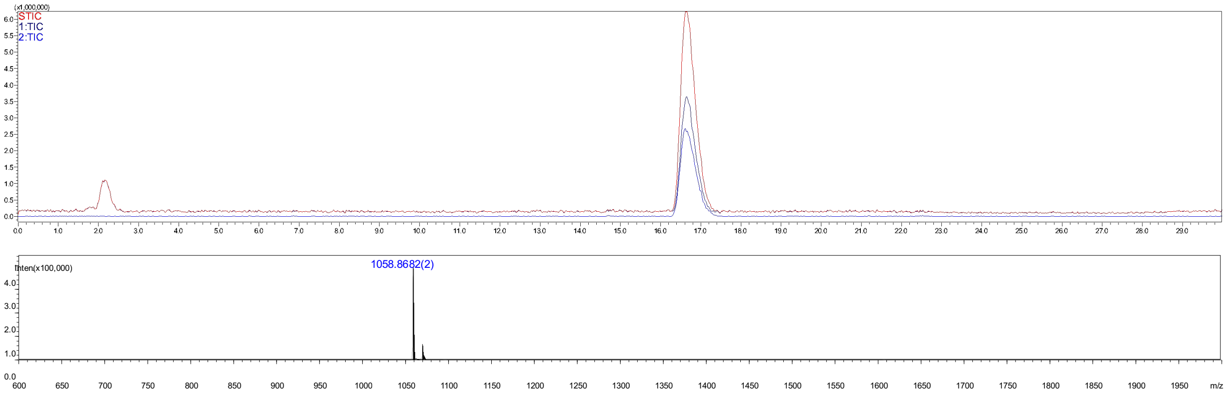


**Figure S37.** Analytical HPLC-MS chromatogram of compound **85**. The retention time = 14.0 min.

**Compound 87**

**87** was synthesized from **74** (1 mg, 1.0 eq) through the installation of α2, 6-Neu5Ac using the general procedure **2.2 e** to give the intermediate product **86**, which was purified using the described two-stage purification system **(2.2 j)**. The Cbz protecting group of **86** was removed by hydrogentation (**2.2 i**) to give final product **87** as a white fluffy solid (0.8 mg, 74% yield over two steps). ESI TOF-MS *m/z* calculated for $C_{91}H_{148}N_8O_{66}$, $[M-2H]^{2-}$: 1204.4241, found 1204.4175.

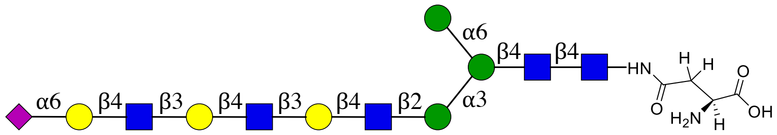


**87**

NMR and MS analysis of compound **86**

**$^1$H (600 MHz, D$_2$O): δ (ppm)**

| | H1 | H2 | H3 | H4 | H5 | H6 |
|---|---|---|---|---|---|---|
| **GlcNAc1** | 5.08 (d, *J* = 9.8 Hz, 1H) | 3.86 | N/R[b] | 3.68 | 3.59 | N/R |
| **GlcNAc2** | 4.64 (d, *J* = 7.9 Hz, 1H) | 3.83 | 3.80 | 3.76 | N/R | N/R |
| **Man3** | 4.81 | 4.28 (d, *J* = 2.3 Hz, 1H) | 3.80 | N/R | N/R | 3.95, 3.84 |
| **Man4** | 5.15 | 4.24 – 4.21 (m, 1H) | N/R | 3.54 | N/R | N/R |
| **Man4'** | 4.95 (d, *J* = 1.7 Hz, 1H) | 4.01 | N/R | 3.67 | N/R | N/R |
| **GlcNAc5** | 4.61 (d, *J* = 7.4 Hz, 1H) | 3.77 | N/R | 3.75 | N/R | N/R |
| **Gal6** | 4.52 – 4.45 (m, 3H) | 3.62 | 3.76 | 4.19 (s, 2H) | N/R | N/R |
| **GlcNAc7** | 4.76 (s, 1H) | 3.83 | N/R | N/R | N/R | N/R |
| **Gal8** | 4.52 – 4.45 (m, 3H) | N/R | 3.76 | 4.19 (s, 2H) | N/R | N/R |
| **GlcNAc9** | 4.73 (d, *J* = 8.4 Hz, 1H) | 3.83 | N/R | N/R | N/R | N/R |
| **Gal10** | 4.52 – 4.45 (m, 3H) | N/R | N/R | N/R | N/R | N/R |

| | H1 | H2 | H3 | H4 | H5 | H6 | H7 | H8 | H9 |
|---|---|---|---|---|---|---|---|---|---|
| **Neu5Ac9** | -[a] | - | 2.71 (dd, *J* = 12.4, 4.7 Hz, 1H), 1.76 (t, *J* = 12.2 Hz, 1H) | 3.70 | 3.84 | N/R | N/R | N/R | N/R |

**$^{13}$C (150 MHz, D$_2$O): δ (ppm)**

| | C1 | C2 | C3 | C4 | C5 | C6 |
|---|---|---|---|---|---|---|
| **GlcNAc1** | 78.15 | 53.70 | N/R | 78.56 | 76.16 | N/R |
| **GlcNAc2** | 101.19 | 54.83 | 65.86 | 79.61 | N/R | N/R |
| **Man3** | 100.33 | 70.13 | 80.31 | N/R | N/R | 65.80 |
| **Man4** | 99.49 | 76.34 | N/R | 67.23 | N/R | N/R |
| **Man4'** | 99.58 | 69.82 | N/R | 66.72 | N/R | N/R |
| **GlcNAc5** | 99.40 | 54.77 | N/R | 78.46 | N/R | N/R |
| **Gal6** | 102.89 | 69.92 | 81.96 | 68.25 | N/R | N/R |
| **GlcNAc7** | 102.51 | 55.08 | N/R | N/R | N/R | N/R |

| | | | | | | |
|---|---|---|---|---|---|---|
| **Gal8** | 102.83 | N/R | 82.00 | 68.25 | N/R | N/R |
| **GlcNAc9** | 102.69 | 54.90 | N/R | N/R | N/R | N/R |
| **Gal10** | 103.40 | N/R | N/R | N/R | N/R | N/R |

| | **C1** | **C2** | **C3** | **C4** | **C5** | **C6** | **C7** | **C8** | **C9** |
|---|---|---|---|---|---|---|---|---|---|
| **Neu5Ac9** | N/R | 100.02 | 39.98 | 68.12 | 51.83 | N/R | N/R | N/R | N/R |

| **Signal** | **Proton** | **Carbon** |
|---|---|---|
| **NHC(O)CH$_3$** | - | 174.85, 174.69, 174.63 |
| **NHC(O)CH$_3$** | 2.34 – 1.86 (m, 18H) | 22.28, 22.24, 22.13, 21.98, 21.91 |
| **Aromatic** | 7.56 – 7.36 (m, 5H) | 128.71, 128.34, 127.71 |
| **CH$_2$-Ph** | 5.21 – 5.13 (m, 2H) | 67.09 |
| **NH-COO-** | - | 157.75 |
| **NH-CH-COOH** | 4.54 (dd, *J* = 8.1, 4.8 Hz, 1H) | 51.29 |
| **NH-CH-COOH** | - | N/R |
| **C(O)-CH$_2$-CH** | 2.89 (dd, *J* = 15.9, 4.7 Hz, 1H), 2.77 (dd, *J* = 15.8, 8.3 Hz, 1H) | 37.60 |
| **C(O)-CH$_2$-CH** | - | 173.35 |

[a] Not applicable

[b] Not reported

ESI TOF-MS *m/z* calculated for $C_{99}H_{154}N_8O_{68}$, $[M-2H]^{2-}$: 1271.4425, found 1271.4315.

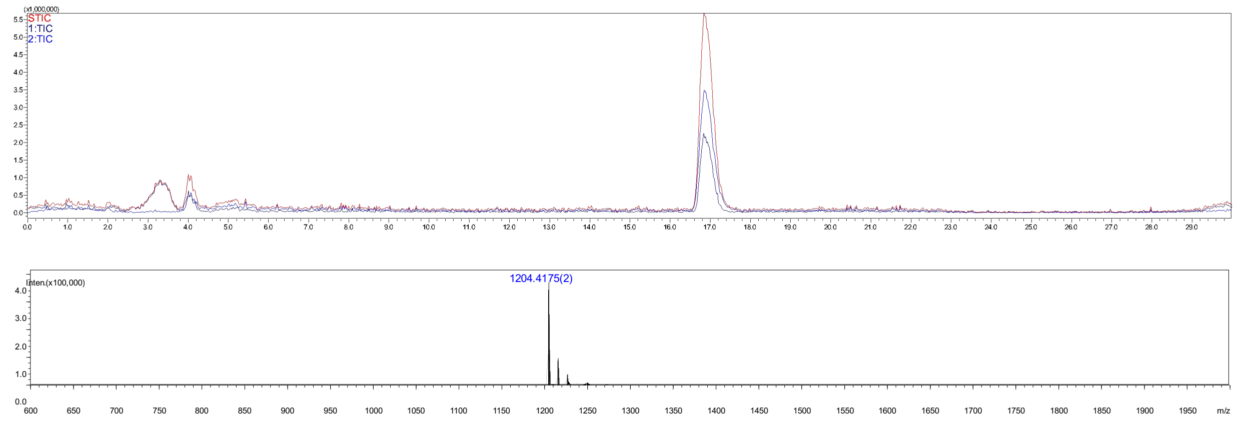


**Figure S38.** Analytical HPLC-MS chromatogram of compound **87**. The retention time = 17.0 min.

**Compound 89**

**89** was synthesized from **74** (2 mg, 1.0 eq) through the installation of α2,3-Neu5Ac using the general procedure **2.2 f** to give intermediate product **88** which was purified using the described two-stage purification system **(2.2 j)**. The Cbz protecrting group of **88** was removed by hydrogentation (**2.2 i**) to give final product **89** as a white fluffy solid (0.6 mg, 54% yield over two steps). ESI TOF-MS *m/z* calculated for $C_{91}H_{148}N_8O_{66}$, $[M-2H]^{2-}$: 1204.4241, found 1204.4201.

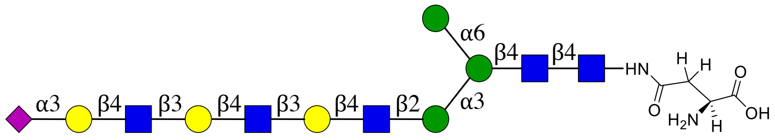


**89**

NMR and MS analysis of compound **88**

**$^1$H (600 MHz, $D_2O$): δ (ppm)**

| | H1 | H2 | H3 | H4 | H5 | H6 |
|---|---|---|---|---|---|---|
| **GlcNAc1** | 5.06 (d, *J* = 9.8 Hz, 1H) | 3.83 | N/R[b] | 3.66 | N/R | N/R |
| **GlcNAc2** | 4.61 (d, *J* = 8.0 Hz, 1H) | 3.80 | 3.77 | 3.74 | N/R | N/R |
| **Man3** | 4.79 | 4.26 (d, *J* = 2.5 Hz, 1H) | 3.77 | N/R | N/R | 3.92, 3.81 |
| **Man4** | 5.13 | 4.20 (d, *J* = 3.6 Hz, 1H) | 3.91 | 3.51 | N/R | N/R |
| **Man4'** | 4.93 (d, *J* = 1.7 Hz, 0H) | 3.98 | 3.89 | 3.65 | N/R | N/R |
| **GlcNAc5** | 4.60 – 4.53 (m, 1H) | 3.75 | N/R | 3.72 | N/R | N/R |
| **Gal6** | 4.50 – 4.41 (m, 2H) | 3.59 | 3.73 | 4.17 (d, *J* = 3.0 Hz, 1H) | N/R | N/R |
| **GlcNAc7** | 4.70 (d, *J* = 8.3 Hz, 2H) | 3.81 | N/R | N/R | N/R | N/R |
| **Gal8** | 4.50 – 4.41 (m, 2H) | 3.59 | 3.73 | 4.17 (d, *J* = 3.0 Hz, 1H) | N/R | N/R |
| **GlcNAc9** | 4.70 (d, *J* = 8.3 Hz, 2H) | 3.81 | N/R | N/R | N/R | N/R |
| **Gal10** | 4.60 – 4.53 (m, 1H) | 3.58 | 3.96 | 4.12 (dd, *J* = 9.9, 3.1 Hz, 1H) | N/R | N/R |

| | H1 | H2 | H3 | H4 | H5 | H6 | H7 | H8 | H9 |
|---|---|---|---|---|---|---|---|---|---|
| **Neu5Ac9** | -[a] | - | 2.77 (dd, *J* = 12.5, 4.6 Hz, 1H), 1.81 (t, *J* = 12.1 Hz, 1H) | 3.70 | 3.85 | N/R | N/R | N/R | N/R |

**$^{13}$C (150 MHz, $D_2O$): δ (ppm)**

| | C1 | C2 | C3 | C4 | C5 | C6 |
|---|---|---|---|---|---|---|
| **GlcNAc1** | 78.07 | 53.65 | N/R | 78.49 | 76.12 | N/R |

| | | | | | | |
|---|---|---|---|---|---|---|
| **GlcNAc2** | 101.16 | 54.79 | 65.82 | 79.59 | N/R | N/R |
| **Man3** | 100.31 | 70.11 | 80.28 | N/R | N/R | 65.76 |
| **Man4** | 99.47 | 76.29 | 69.30 | 67.19 | N/R | N/R |
| **Man4'** | 99.55 | 69.78 | 70.31 | 66.69 | N/R | N/R |
| **GlcNAc5** | 99.37 | 54.73 | N/R | 75.09 | N/R | N/R |
| **Gal6** | 102.86 | 69.88 | 81.98 | 68.25 | N/R | N/R |
| **GlcNAc7** | 102.72 | 55.10 | N/R | N/R | N/R | N/R |
| **Gal8** | 102.79 | 69.88 | 81.98 | 68.25 | N/R | N/R |
| **GlcNAc9** | 102.68 | 55.06 | N/R | N/R | N/R | N/R |
| **Gal10** | 102.45 | 69.30 | 67.39 | 75.41 | N/R | N/R |

| | **C1** | **C2** | **C3** | **C4** | **C5** | **C6** | **C7** | **C8** | **C9** |
|---|---|---|---|---|---|---|---|---|---|
| **Neu5Ac9** | N/R | 99.71 | 39.55 | 68.23 | 51.60 | N/R | N/R | N/R | N/R |

| **Signal** | **Proton** | **Carbon** |
|---|---|---|
| **NH<u>C</u>(O)CH3** | - | 174.92, 174.81, 174.69, 174.61 |
| **NHC(O)<u>CH3</u>** | 2.35 – 1.85 (m, 18H) | 22.25, 22.09, 21.95, 21.89 |
| **Aromatic** | 7.55 – 7.33 (m, 5H) | 136.19, 128.68, 128.30, 127.70 |
| **<u>CH2</u>-Ph** | 5.19 – 5.09 (m, 2H) | 65.76 |
| **NH-<u>C</u>OO-** | - | 157.73 |
| **NH-<u>CH</u>-COOH** | 4.45 | 51.94 |
| **NH-CH-<u>C</u>OOH** | - | N/R |
| **C(O)-<u>CH2</u>-CH** | 2.85 (dd, $J$ = 15.7, 4.5 Hz, 1H), 2.69 (dd, $J$ = 15.6, 8.7 Hz, 1H) | 37.89 |
| **<u>C</u>(O)-CH2-CH** | - | 173.73 |

[a] Not applicable

[b] Not reported

ESI TOF-MS *m/z* calculated for $C_{99}H_{154}N_8O_{68}$, $[M-2H]^{2-}$: 1271.4425, found 1271.4318.

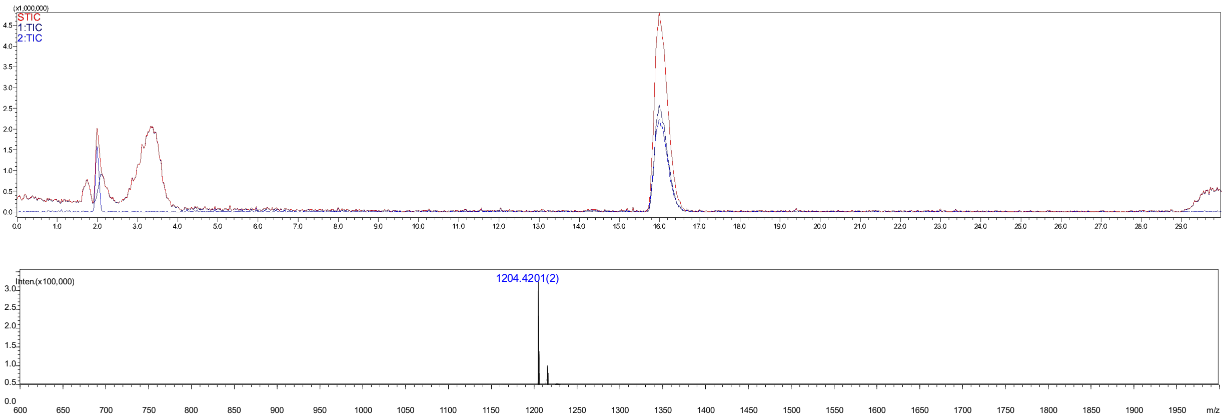


**Figure S39.** Analytical HPLC-MS chromatogram of compound **89**. The retention time = 1.0 min.

#### 3. Microarray

##### 3.1 Materials

Virus isolates were produced as described previously.[7] Oseltamivir was purchased from Sigma Aldrich [Cat# SML1606]. CR8020 group 2, and CR6261 group 1 stem antibodies provided by Dr. Dirk Eggink and expressed following previously published procedures.[8] Goat anti-human Alexa-647 [Cat# A21445] antibodies and streptavidin-AlexaFluor 635 [Cat# SA1011] were obtained from Thermo Fisher. Control lectins *Erythrina cristagalli* agglutinin (ECA) [Cat#B-1145], *Sambuca nigra* agglutinin (SNA) [Cat# B-1305], and *Maackia amurensis* lectin I (MAL-I) [Cat# B-1315] were purchased from Vector Labs.

##### 3.2 Glycan printing surfaces

Compounds were printed on amine reactive, N-hydroxy succinimide (NHS)-activated glass slides (NEXTERION ® Slide H, Schott Inc) employing the free amine of the anomeric asparagine moiety of the *N*-glycans for immobilization. The printing was performed using a Scienion sciFLEXARRAYER S3 non-contact microarray printer equipped with a Scienion PDC80 nozzle (Scienion Inc). Glycans were dissolved in sodium phosphate buffer (250 mM, pH 8.5) at a concentration of 100 μM and printed in replicates of six (spot volume of ~400 pL, at 20 °C and 50% humidity). Slides were blocked with 5 mM ethanolamine in Tris buffer (pH 9, 50 mM) for 1 h at 50 °C and rinsed with DI water after printing.

##### 3.3 Glycan microarray

Quality control was performed using the plant lectins SNA, ECA and MAL-I at 10 μg/mL precomplexed with 2 μg/mL Streptavidin-AlexaFluor 635.

Viral isolates (25 μL) diluted in PBS-T (PBS + 0.1% Tween, 25 μL) were applied to subarrays in the presence of oseltamivir (200 nM) in a humidified chamber for 1 h. Next, the microarray slide was rinsed with PBS-T (PBS + 0.1% Tween), PBS, and deionized water (2x) and dried by centrifugation. The slide was incubated for 1 h in the presence of the CR8020 A/H3N2 influenza hemaglutinin or CR6261 stem specific antibody (100 μL, 5 μg $mL^{-1}$ in PBS-T) and washed as described above. A secondary goat anti-human AlexaFluor-647 antibody (100 μL, 2 μg $mL^{-1}$ in PBS-T) was applied and the resulting slide was incubated for 1 h in a humidified chamber and then washed by the standard procedure. Slides were dried by centrifugation after the washing steps and scanned immediately using an Innopsys Innoscan 710 microarray scanner. Various gains and PMT values were used to ensure the signals were in the linear range and to avoid saturation of the signals. Images were analyzed with Mapix software (version 8.1.0 Innopsys) and processed with an Excel macro (https://github.com/enthalpyliu/carbohydrate-microarray-processing). The average fluorescence intensity and SD were determined for each compound after removal of the highest and lowest intensities from the spot replicates to give $n = 4$.

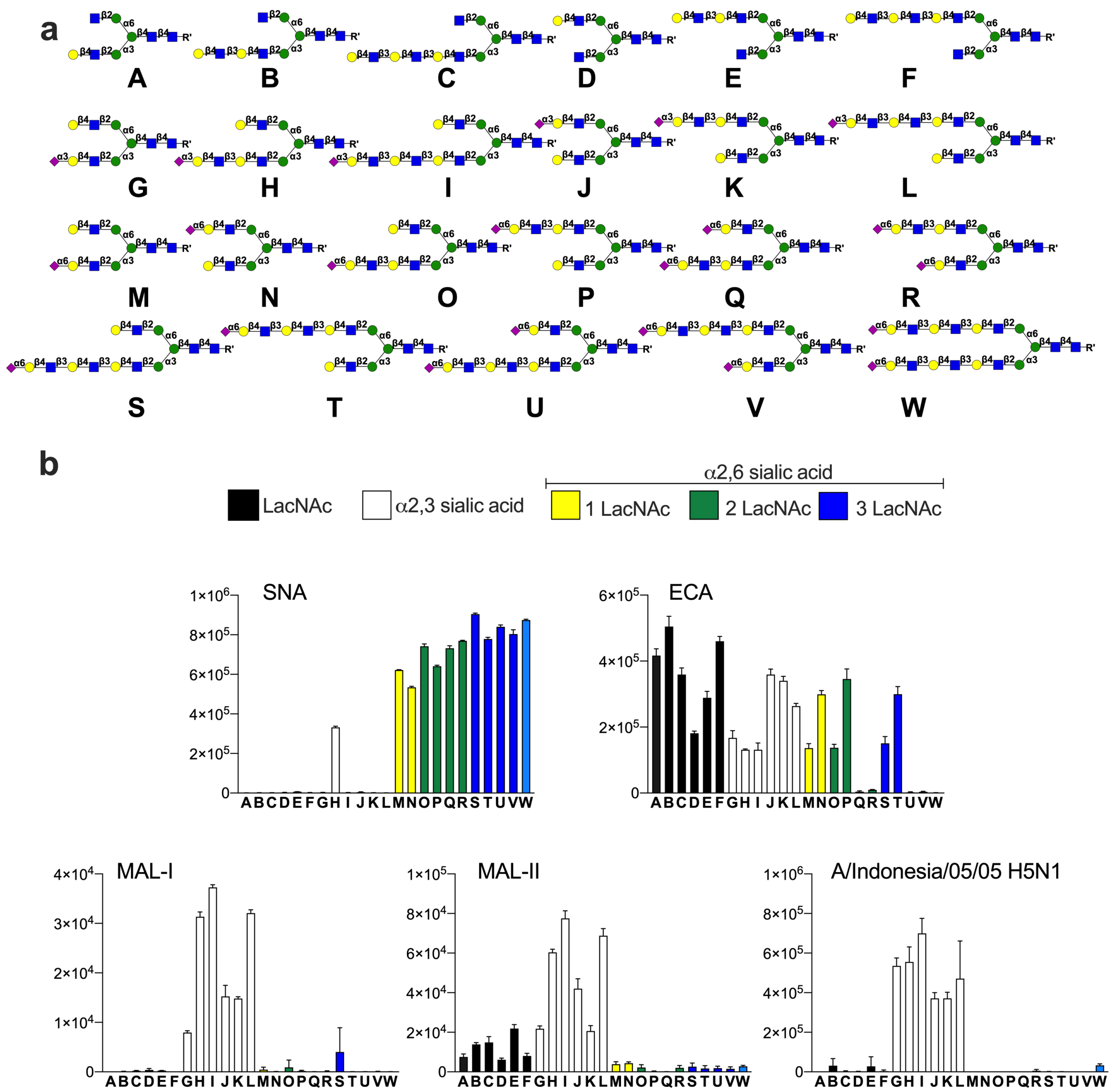


**Figure S40.** Validation of the microarray. **a**) Library of compounds. **b**) Microarray screening for lectins and and A/Indonesia/05/05 H5N1. Bars represent the background-subtracted average relative fluorescence units (RFU) of four replicates ±SD.

### 4. Hemagglutination and sequence alignment

##### 4.1 Erythrocyte preparation

Fresh turkey blood was centrifuged (430 rcf, 10 min) followed by removal of the supernatant. Pellets were washed with PBS three times with intermittent centrifugation (430 rcf, 10 min). Erythrocyte suspensions were stored a 50% solution in PBS.

##### 4.2 Enzymatic erythrocyte remodeling

*Arthrobacter ureafaciens* neuraminidase (12 U; New England Biolabs) was added to a suspension of turkey erythrocytes (250 µL, 50%) in PBS (900 µL) and incubated for 6 h at 37 °C with tilting. Next, recombinant B4GALT1 (37.5 µL, 1 mg/mL) and B3GnT2 (37.5 µL, 1 mg/mL), UDP-Gal (4.4 mM) and UDP-GlcNAc (4.4 mM), alkaline phosphatase (6 U), $MnCl_2$ (2 mM) and BSA (6 µL, 2 mg/mL) were added. This mixture was incubated for 18 h at 37 °C with gentle tilting. Next, the erythrocytes were washed with PBS (2x, 750 µL) and the pellet was reconstituted in PBS (900 µL). The erythrocytes were resialylated with ST6Gal1 (37 µL, 1 mg/mL) and CMP-Neu5Ac (4.5 mM) in the presence of alkaline phosphatase (6 U) and BSA (6 µL, 2 mg/mL) for 4 h at 37 °C with gentle tilting. The erythrocytes were washed with PBS (1x, 600 µL) and diluted to a 1% solution.

##### 4.3 Hemagglutination assay

Hemagglutination assays were performed according to a standard procedure.[9.] Briefly, virus stocks were two-fold serial diluted in the presence of oseltamivir (20 nM) in PBS. Turkey erythrocytes (1%, 25 µL) were mixed with the serial diluted viruses and incubated for 1 h at 4 °C. Titers were expressed as the highest dilution of virus stock that completely agglutinated the turkey erythrocytes.

**Table S2.** Amino acid alignment of H3 proteins.

| No. | Name | Clade | Year | 98 | 120 | 121 | 122 | 123 | 124 | 125 | 126 | 127 | 128 | 129 | 130 | 131 | 132 | 133 | 134 | 135 | 136 | 137 | 138 | 139 | 140 | 141 | 142 | 143 | 144 | 145 | 146 | 155 | 156 | 157 | 158 | 159 | 160 | 161 | 162 | 163 | 164 | 186 | 187 | 188 | 189 | 190 | 191 | 192 | 193 | 194 | 195 | 196 | 197 | 198 | 199 | 200 | 201 | 202 | 203 | 204 | 205 | 206 | 207 | 208 | 209 | 210 | 211 | 212 | 213 | 219 | 220 | 221 | 222 | 223 | 224 | 225 | 226 | 227 |
|---|---|---|---|---|---|---|---|---|---|---|---|---|---|---|---|---|---|---|---|---|---|---|---|---|---|---|---|---|---|---|---|---|---|---|---|---|---|---|---|---|---|---|---|---|---|---|---|---|---|---|---|---|---|---|---|---|---|---|---|---|---|---|---|---|---|---|---|---|---|---|---|---|---|---|---|---|---|---|
| 1 | A/Hong Kong/001/1968 | Unknown | 1968 | Y | F | I | T | E | G | F | T | W | T | G | V | T | Q | N | G | G | S | N | A | C | K | R | G | P | G | S | G | T | K | S | G | S | T | Y | P | V | L | S | T | N | Q | E | Q | T | S | L | Y | V | Q | A | S | G | R | V | T | V | S | T | R | R | S | Q | Q | T | I | S | R | P | W | V | R | G | L | S |
| 2 | A/Bilthoven/16190/1968 | Unknown | 1968 | Y | F | I | T | E | G | F | T | W | T | G | V | T | Q | N | G | G | S | N | A | C | K | R | G | P | G | S | G | T | K | S | G | S | T | Y | P | V | L | S | T | N | Q | E | Q | T | S | L | Y | V | Q | A | S | G | R | V | T | V | S | T | R | R | S | Q | Q | T | I | S | R | P | W | V | R | G | L | S |
| 3 | A/Beijing/353/1989 | Unknown | 1989 | Y | F | I | N | E | D | F | N | W | T | G | V | A | Q | S | G | E | S | Y | A | C | K | R | G | S | V | K | S | H | E | S | E | Y | K | Y | P | A | L | S | T | D | R | E | Q | T | K | L | Y | V | R | A | S | G | R | V | T | V | S | T | K | R | S | Q | Q | T | V | S | R | P | W | V | R | G | L | S |
| 4 | A/Netherlands/816/1991 | Unknown | 1991 | Y | F | I | N | E | D | F | N | W | T | G | V | A | Q | S | G | E | S | Y | A | C | K | R | G | S | V | K | S | H | E | S | E | Y | K | Y | P | A | L | S | T | D | R | E | Q | T | S | L | Y | V | R | A | S | G | R | V | T | V | S | T | K | R | S | Q | Q | T | V | S | R | P | W | V | R | G | L | S |
| 5 | A/Netherlands/109/2003 | Unknown | 2003 | Y | F | N | N | E | S | F | N | W | T | G | V | T | Q | N | G | T | S | S | A | C | K | R | R | S | N | K | S | T | H | L | K | Y | K | Y | P | A | L | G | T | D | S | D | Q | I | S | L | Y | A | Q | A | S | G | R | I | T | V | S | T | K | R | S | Q | Q | T | V | S | R | P | R | V | R | D | I | S |
| 6 | A/Wisconsin/67/2005 | Unknown | 2005 | Y | F | N | D | E | S | F | N | W | T | G | V | T | Q | N | G | T | S | S | S | C | K | R | R | S | N | N | S | T | H | L | K | F | K | Y | P | A | L | V | T | D | N | D | Q | I | F | L | Y | A | Q | A | S | G | R | I | T | V | S | T | K | R | S | Q | Q | T | V | S | R | P | R | I | R | N | I | P |
| 7 | A/Brisbane/10/2007 | Unknown | 2007 | Y | F | N | N | E | S | F | N | W | T | G | V | T | Q | N | G | T | S | S | A | C | I | R | R | S | N | N | S | T | H | L | K | F | K | Y | P | A | L | G | T | D | N | D | Q | I | F | P | Y | A | Q | A | S | G | R | I | T | V | S | T | K | R | S | Q | Q | T | V | S | R | P | R | V | R | N | I | P |
| 8 | A/Netherlands/761/2009 | Unknown | 2009 | Y | F | N | N | E | S | F | N | W | T | G | V | T | Q | N | G | T | S | S | A | C | I | R | R | S | N | N | S | T | H | L | R | F | K | Y | P | A | L | G | T | D | N | D | Q | I | F | L | Y | A | Q | A | S | G | R | I | T | V | S | T | K | R | S | Q | Q | T | V | S | R | P | R | V | R | N | I | P |
| 9 | A/Singapore/INFIMH-16-0019/2016 | 3c.2a1 | 2016 | Y | F | K | N | E | S | F | N | W | T | G | V | T | Q | N | G | T | S | S | A | C | I | R | G | S | S | S | S | T | H | L | N | Y | T | Y | P | A | L | G | T | D | K | D | Q | I | F | L | Y | A | Q | S | S | G | R | I | T | V | S | T | K | R | S | Q | Q | A | V | S | R | P | R | I | R | D | I | P |
| 10 | A/Netherlands/2413/16 | 3C.2a1 | 2016 | Y | F | K | N | E | S | F | N | W | T | G | V | T | Q | N | G | T | S | S | A | C | M | R | R | S | S | S | S | T | H | L | H | Y | T | Y | P | A | L | G | T | D | K | D | Q | I | F | L | Y | A | Q | S | S | G | R | I | T | V | S | T | K | R | S | Q | Q | A | V | S | R | P | R | I | R | D | I | P |
| 11 | A/Netherlands/1797/2017 | 3C.2a1 | 2017 | Y | F | K | N | E | S | F | N | W | A | G | V | T | Q | N | G | K | S | S | A | C | I | R | G | S | S | S | S | T | H | L | N | Y | T | Y | P | A | L | G | T | D | K | D | Q | I | F | L | Y | A | Q | S | S | G | R | I | T | V | S | T | K | R | S | Q | Q | A | V | S | R | P | R | I | R | D | I | P |
| 12 | A/Netherlands/371/2019 | 3C.2a1b | 2019 | Y | F | K | N | E | S | F | N | W | A | G | V | T | Q | N | G | K | S | S | A | C | I | R | G | S | S | S | S | T | H | L | N | Y | T | Y | P | A | L | G | T | D | K | D | Q | I | F | L | Y | A | Q | S | S | G | R | I | T | V | S | T | K | R | S | Q | Q | A | V | S | R | P | R | I | R | D | I | P |
| 13 | A/Netherlands/10/2019 | 3C.2a1b.1 | 2019 | Y | F | K | N | E | S | F | N | W | A | G | V | T | Q | N | G | K | S | S | A | C | I | R | G | S | S | S | S | T | H | L | N | Y | T | Y | P | A | L | G | T | D | K | D | Q | I | F | L | Y | A | Q | S | S | G | R | I | T | V | S | T | K | R | S | Q | Q | A | V | S | R | P | R | I | R | D | I | P |
| 14 | A/Netherlands/314/2019 | 3C.2a1b | 2019 | Y | F | K | N | E | S | F | N | W | A | G | V | T | Q | N | G | K | S | S | A | C | I | R | G | S | S | S | S | T | H | L | N | Y | T | Y | P | A | L | G | T | D | K | D | Q | I | F | L | Y | A | Q | S | S | G | R | I | T | V | S | T | K | R | S | Q | Q | A | V | S | R | P | R | I | R | D | I | P |
| 15 | A/Netherlands/751/2017 | 3C.2a1 | 2017 | Y | F | K | N | E | S | F | N | W | T | G | V | T | Q | N | G | T | S | S | A | C | M | R | R | S | S | S | S | T | H | L | N | Y | T | Y | P | A | L | G | T | D | K | D | Q | I | F | L | Y | A | Q | S | S | G | R | I | T | V | S | T | K | R | S | Q | Q | A | V | S | R | P | R | I | R | D | I | P |
| 16 | A/Switzerland/8060/2017 | 3C.2a2 | 2017 | Y | F | N | N | E | S | F | N | W | T | G | V | K | Q | N | G | T | S | S | A | C | I | R | K | S | S | S | S | T | H | L | N | Y | T | Y | P | A | L | G | T | D | K | D | Q | I | F | L | Y | A | Q | S | S | G | R | I | T | V | S | T | K | R | S | Q | Q | A | V | S | R | P | R | I | R | D | I | P |
| 17 | A/Netherlands/3466/2017 | 3C.2a | 2017 | Y | X | N | N | E | S | F | N | W | T | G | V | K | Q | N | G | T | S | S | A | C | I | R | K | S | S | S | S | T | H | L | N | Y | T | Y | P | A | L | G | T | D | K | D | Q | I | F | L | Y | A | Q | S | S | G | R | I | T | V | S | T | K | R | S | Q | Q | A | V | S | R | P | R | I | R | D | I | P |
| 18 | A/Netherlands/1802/2018 | 3C.2a2 | 2018 | Y | F | N | N | E | S | F | N | W | T | G | V | K | Q | N | G | T | S | S | A | C | I | R | K | S | R | S | S | T | H | L | N | Y | T | Y | P | A | L | G | T | D | K | D | Q | I | F | L | Y | A | Q | S | S | G | R | I | T | V | S | T | K | R | S | Q | Q | A | V | F | R | P | R | I | R | D | I | P |
| 19 | A/Netherlands/295/2019 | 3C.2a1b | 2019 | Y | F | K | N | E | S | F | N | W | T | G | V | K | Q | N | G | T | S | S | A | C | I | R | G | S | S | S | S | T | H | L | N | Y | T | Y | P | A | L | G | T | D | K | D | Q | I | F | L | Y | A | Q | S | S | G | R | I | T | V | S | T | K | R | S | Q | Q | A | V | F | R | P | R | I | R | D | I | P |
| 20 | A/Netherlands/10616/2019 | 3C.2a2 | 2019 | Y | F | N | N | E | N | F | N | W | T | G | V | K | Q | N | G | T | S | S | A | C | I | R | K | S | S | S | S | T | H | L | N | Y | T | Y | P | A | L | G | T | D | K | D | Q | I | F | L | Y | A | Q | S | S | G | R | I | T | V | S | T | K | R | S | Q | Q | A | V | S | R | P | R | I | R | D | I | P |
| 21 | A/Netherlands/10009/2019 | 3C.2a1b | 2019 | Y | F | K | N | E | S | F | N | W | T | G | V | K | Q | N | G | T | S | S | A | C | I | R | G | S | S | S | S | T | H | L | N | Y | T | Y | P | A | L | G | T | D | K | D | Q | I | F | L | Y | A | R | S | S | G | R | I | T | V | S | T | K | R | S | Q | Q | A | V | F | R | P | R | I | R | D | I | P |
| 22 | A/Netherlands/110/2021 | 3C.2a1b.1a | 2021 | Y | F | K | N | E | S | F | N | W | A | G | V | T | Q | N | G | K | S | S | S | C | I | R | G | S | S | S | S | T | H | L | N | Y | T | Y | P | A | L | D | T | D | K | N | Q | I | S | L | Y | A | Q | P | S | G | R | I | T | V | F | T | K | R | S | Q | Q | A | V | S | R | P | R | I | R | D | I | P |
| 23 | A/Netherlands/7/2021 | 3C.2a1b.2a.2 | 2021 | Y | F | K | N | E | S | F | N | W | T | G | V | K | Q | N | G | T | S | S | A | C | I | R | G | S | S | S | S | T | S | L | N | N | I | Y | P | A | Q | D | T | D | K | N | Q | I | S | L | F | A | Q | S | S | G | R | I | T | V | S | T | K | R | S | Q | Q | A | V | S | R | P | R | I | R | D | I | P |
| 24 | A/Netherlands/8/2021 | 3C.2a1b.2a.2 | 2021 | Y | F | K | N | E | S | F | N | W | T | G | V | K | Q | N | G | T | S | S | A | C | I | R | G | S | S | S | S | T | S | L | N | N | I | Y | P | A | Q | D | T | D | K | N | Q | F | S | L | F | A | Q | S | S | G | R | I | T | V | S | T | K | R | S | Q | Q | A | V | S | R | P | R | I | R | D | I | P |
| 25 | A/Netherlands/92/2021 | 3C.2a1b.2a.2 | 2021 | Y | F | K | N | E | S | F | N | W | T | G | V | K | Q | N | G | T | S | S | A | C | I | R | G | S | S | S | S | T | S | L | N | N | I | Y | P | A | Q | D | T | D | K | N | Q | I | S | L | F | A | Q | S | S | G | R | I | T | V | S | T | K | R | S | Q | Q | A | V | S | R | P | R | I | R | D | I | P |
| 26 | A/Netherlands/1/2022 | 3C.2a1b.2a.2 | 2022 | Y | F | K | N | E | S | F | N | W | T | G | V | K | Q | N | G | T | S | S | A | C | I | R | G | S | S | S | S | T | H | L | N | N | I | Y | P | A | Q | D | T | D | K | N | Q | I | S | L | F | A | Q | S | S | G | R | I | T | V | S | T | K | R | S | Q | Q | A | V | S | R | P | R | I | R | D | I | P |
| 27 | A/Netherlands/832/2022 | 3C.2a1b.2a.2 | 2022 | Y | F | K | N | E | S | F | N | W | T | G | V | K | Q | N | G | T | S | S | A | C | K | R | G | S | S | S | S | T | H | L | N | N | I | Y | P | A | Q | D | T | D | K | N | Q | I | S | L | F | A | Q | S | S | G | R | I | T | V | S | T | K | R | S | Q | Q | A | V | S | R | P | K | I | R | D | I | P |
| 28 | A/Netherlands/568/2023 | 3C.2a1b.2a.2 | 2023 | Y | F | K | N | E | S | F | N | W | A | G | V | T | Q | N | G | K | S | S | A | C | I | R | G | S | S | S | S | T | H | L | N | Y | T | Y | P | A | L | G | T | D | K | D | Q | I | F | L | Y | A | Q | S | S | G | R | I | T | V | S | T | K | R | S | Q | Q | A | V | S | R | P | R | I | R | D | I | P |
| 29 | A/Netherlands/59/2023 | 3C.2a1b.2a.2 | 2023 | Y | F | K | D | E | S | F | N | W | T | G | V | K | Q | N | G | T | S | S | A | C | K | R | G | S | S | S | S | T | S | L | N | N | I | Y | P | A | Q | D | T | D | K | N | Q | F | S | L | F | A | Q | S | S | G | R | I | T | V | S | T | K | R | S | Q | Q | A | V | S | R | P | R | V | R | D | I | P |
| 30 | A/Texas/50/2012 | 3C.1 | 2012 | Y | F | N | N | E | S | F | N | W | N | G | V | T | Q | N | G | T | S | S | A | C | I | R | R | S | N | N | S | T | H | L | N | F | K | Y | P | A | L | G | T | D | K | D | Q | I | F | L | Y | A | Q | P | S | G | R | I | T | V | S | T | K | R | S | Q | Q | A | V | S | R | P | R | I | R | N | I | P |
| 31 | A/Switzerland/9715293/2013 | 3c.3a | 2013 | Y | F | N | N | E | S | F | N | W | A | G | V | T | Q | N | G | T | S | S | S | C | I | R | G | S | N | S | S | T | H | L | N | S | K | Y | P | A | L | G | T | D | K | D | Q | I | F | L | Y | A | Q | S | S | G | R | I | T | V | S | T | K | R | S | Q | Q | A | V | S | R | P | R | I | R | D | I | P |
| 32 | A/Netherlands/622/2012 | 3C.3 | 2012 | Y | F | N | N | E | S | F | N | W | A | G | V | T | Q | N | G | T | S | S | A | C | I | R | G | S | N | S | S | T | H | L | N | F | K | Y | P | A | L | G | T | D | K | D | Q | I | F | L | Y | A | Q | S | S | G | R | I | T | V | S | T | K | R | S | Q | Q | A | V | S | R | P | R | I | R | N | I | P |
| 33 | A/Netherlands/153/2016+B38 | 3C.3a | 2016 | Y | F | N | N | E | S | F | N | W | A | G | V | T | Q | N | G | T | S | S | S | C | I | R | G | S | N | S | S | T | H | L | N | S | K | Y | P | A | L | G | T | D | K | D | Q | I | F | L | Y | A | Q | S | S | G | R | I | T | V | S | T | K | R | S | Q | Q | A | V | S | R | P | R | I | R | D | I | P |
| 34 | A/Kansas/14/2017 | 3C.3a | 2017 | Y | F | N | N | E | S | F | N | W | A | G | V | T | Q | N | G | T | S | S | S | C | I | R | G | S | K | S | S | T | H | L | N | S | K | Y | P | A | L | G | T | D | K | D | Q | I | S | L | Y | A | Q | S | S | G | R | I | T | V | S | T | K | R | S | Q | Q | A | V | S | R | P | R | I | R | D | I | P |
| 35 | A/Netherlands/384/2019 | 3C.3a | 2019 | Y | F | N | N | E | S | F | N | W | A | G | V | T | Q | N | G | T | S | S | S | C | I | R | G | S | N | S | S | T | H | L | N | S | K | Y | P | A | L | G | T | D | K | D | Q | I | F | L | Y | A | Q | S | S | G | R | I | T | V | S | T | K | R | S | Q | Q | A | V | S | R | P | R | I | R | D | I | P |
| 36 | A/Netherlands/10002/2019 | 3C.3a | 2019 | Y | F | N | N | E | S | F | N | W | A | G | V | T | Q | N | G | T | S | S | S | C | I | R | G | S | K | S | S | T | H | L | N | S | K | Y | P | A | L | G | T | D | K | D | Q | T | S | L | Y | A | Q | S | S | G | R | I | T | V | S | T | K | R | S | Q | Q | A | V | S | R | P | R | I | R | D | I | P |
| 37 | A/Netherlands/10006/2019 | 3C.3a | 2019 | Y | F | N | N | E | S | F | N | W | A | G | V | T | Q | N | G | T | S | S | S | C | I | R | G | S | K | S | S | T | H | L | N | S | K | Y | P | A | L | G | T | D | K | D | Q | T | S | L | Y | A | Q | S | S | G | R | I | T | V | S | T | K | R | S | Q | Q | A | V | S | R | P | R | I | R | D | I | P |
| 38 | A/Netherlands/322/2020 | 3C.3a1 | 2020 | Y | F | N | N | E | S | F | N | W | A | G | V | T | Q | N | G | T | S | S | S | C | I | R | G | S | K | S | S | T | H | L | N | S | K | Y | P | A | L | G | T | D | K | D | Q | I | S | L | Y | A | Q | S | S | G | R | I | T | V | S | T | K | R | S | Q | Q | A | V | S | R | P | R | I | R | D | I | P |

**Table S3.** Hemagglutination titers of additional A(H3N2) viruses isolated between 2021 and 2023 for wild type (WT) and glyco-remodeled (mod) turkey erythrocytes.

| Name | GISAID | HA (WT) titer | HA (mod) titer |
|---|---|---|---|
| **A/H3N2 viruses** | | | |
| A/NL/7/21 | EPI_ISL_3447260 | 48 | 64 |
| A/NL/8/21 | EPI_ISL_4551782 | 96 | 256 |
| A/NL/11832/22 | EPI_ISL_15895148 | 96 | 128 |
| A/NL/12136/22 | EPI_ISL_16494722 | 96 | 96 |
| A/NL/1431/22 | EPI_ISL_16147119 | 48 | 64 |
| A/NL/1867/22 | EPI_ISL_16334493 | 24 | 32 |
| A/NL/10162/23 | EPI_ISL_16812568 | 96 | 128 |
| A/NL/496/23 | EPI_ISL_16877488 | 48 | 32 |
| A/NL/10142/23 | EPI_ISL_16700790 | 48 | 64 |
| A/NL/10290/23 | EPI_ISL_17017548 | 96 | 128 |
| A/NL/10277/23 | EPI_ISL_17017534 | 192 | 256 |
| A/NL/10132/23 | EPI_ISL_16700787 | 96 | 256 |
| A/NL/10226/23 | EPI_ISL_16867389 | 192 | 128 |
| A/NL/10228/23 | EPI_ISL_16954700 | 192 | 192 |
| A/NL/652/23 | EPI_ISL_16955822 | 24 | 128 |
| A/NL/442/23 | EPI_ISL_16877479 | 24 | 192 |

#### 6. NMR spectra

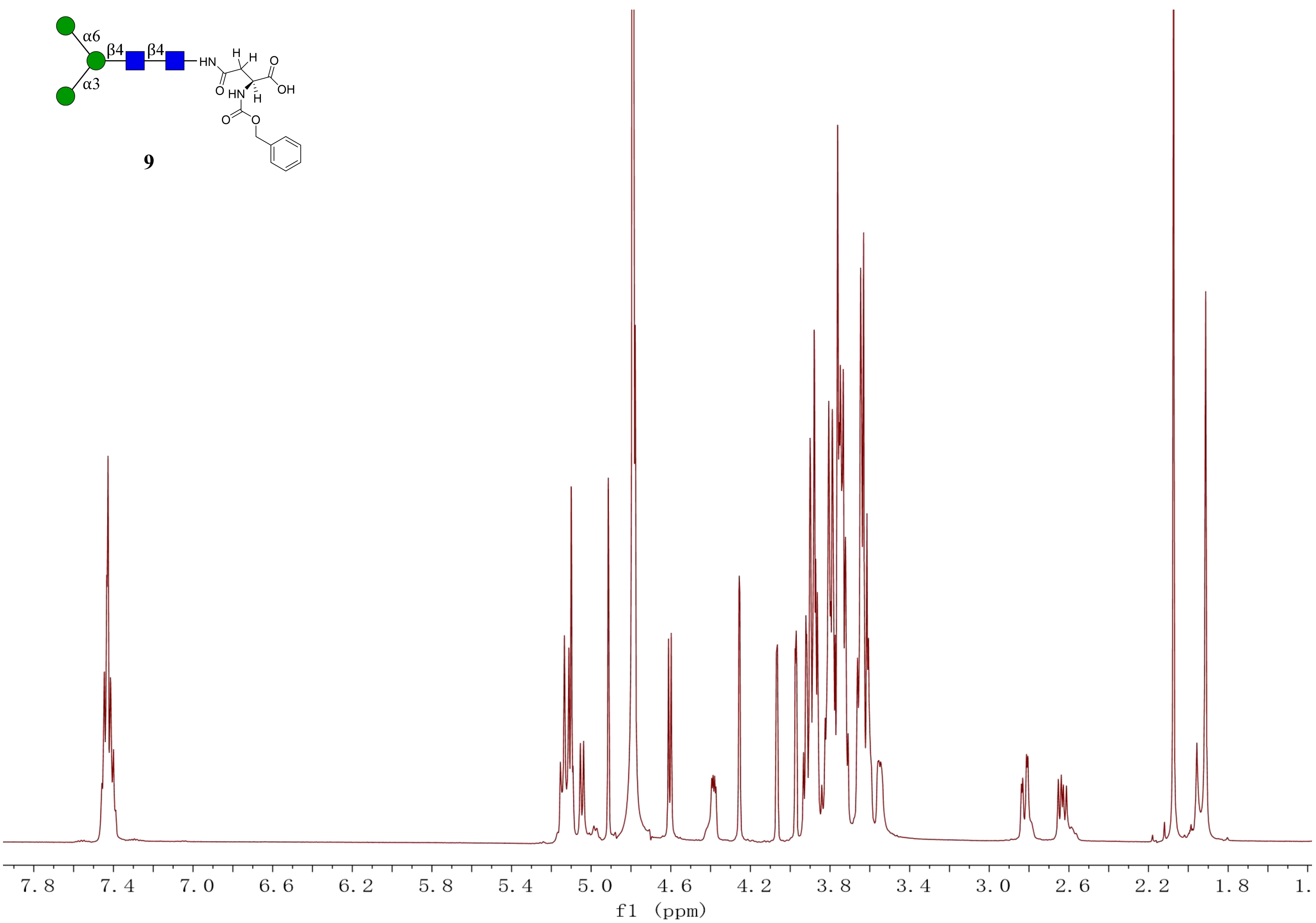

α6
β4
β4
α3
HN
H
H
O
OH
O
HN
H
O
O
9
7.8
7.4
7.0
6.6
6.2
5.8
5.4
5.0
4.6
4.2
3.8
3.4
3.0
2.6
2.2
1.8
1.
f1 (ppm)

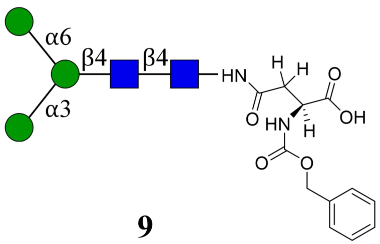

α6
β4
β4
α3
HN
H
H
O
O
HN
H
OH
O
O


**9**

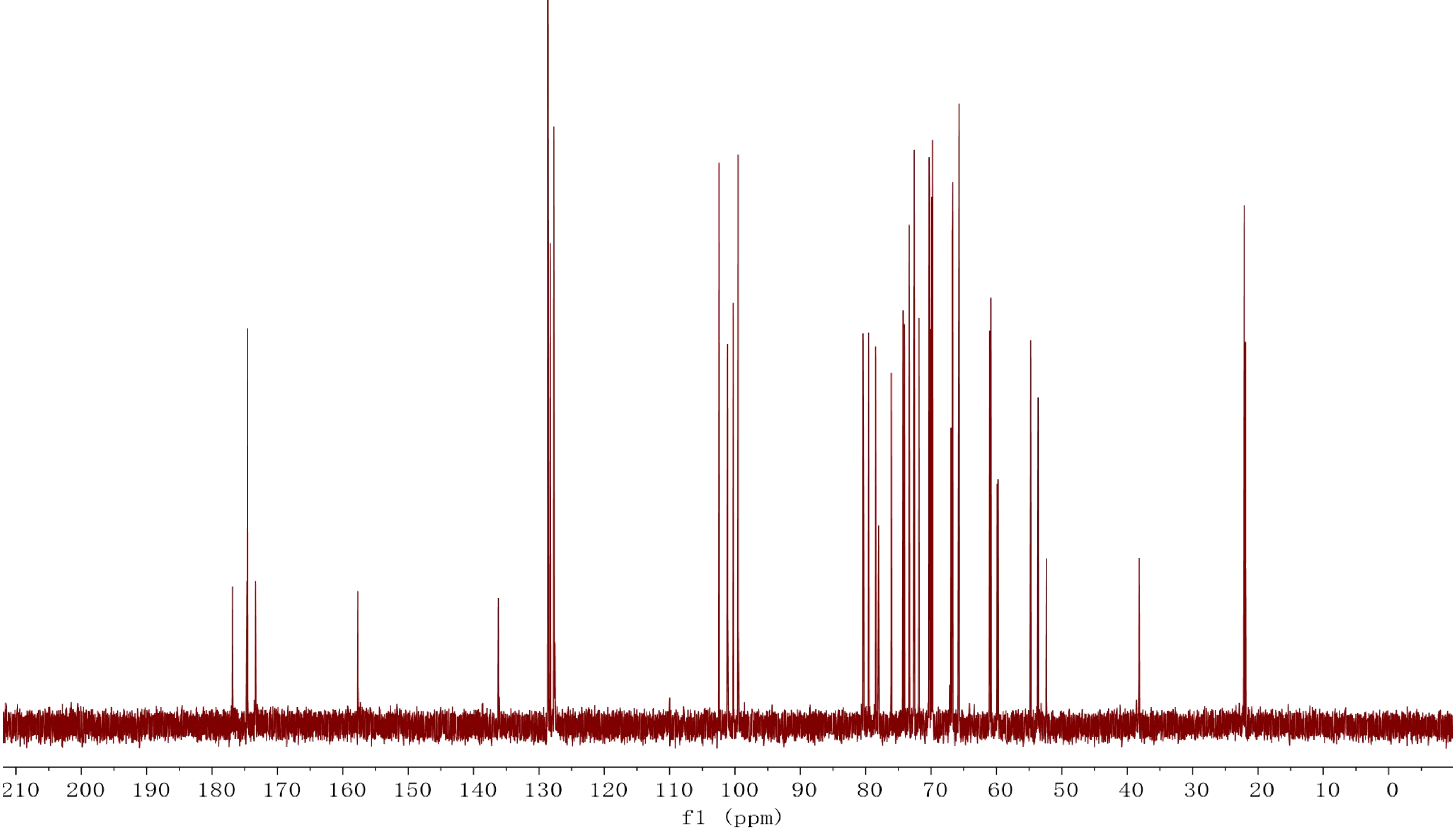

210 200 190 180 170 160 150 140 130 120 110 100 90 80 70 60 50 40 30 20 10 0
f1 (ppm)

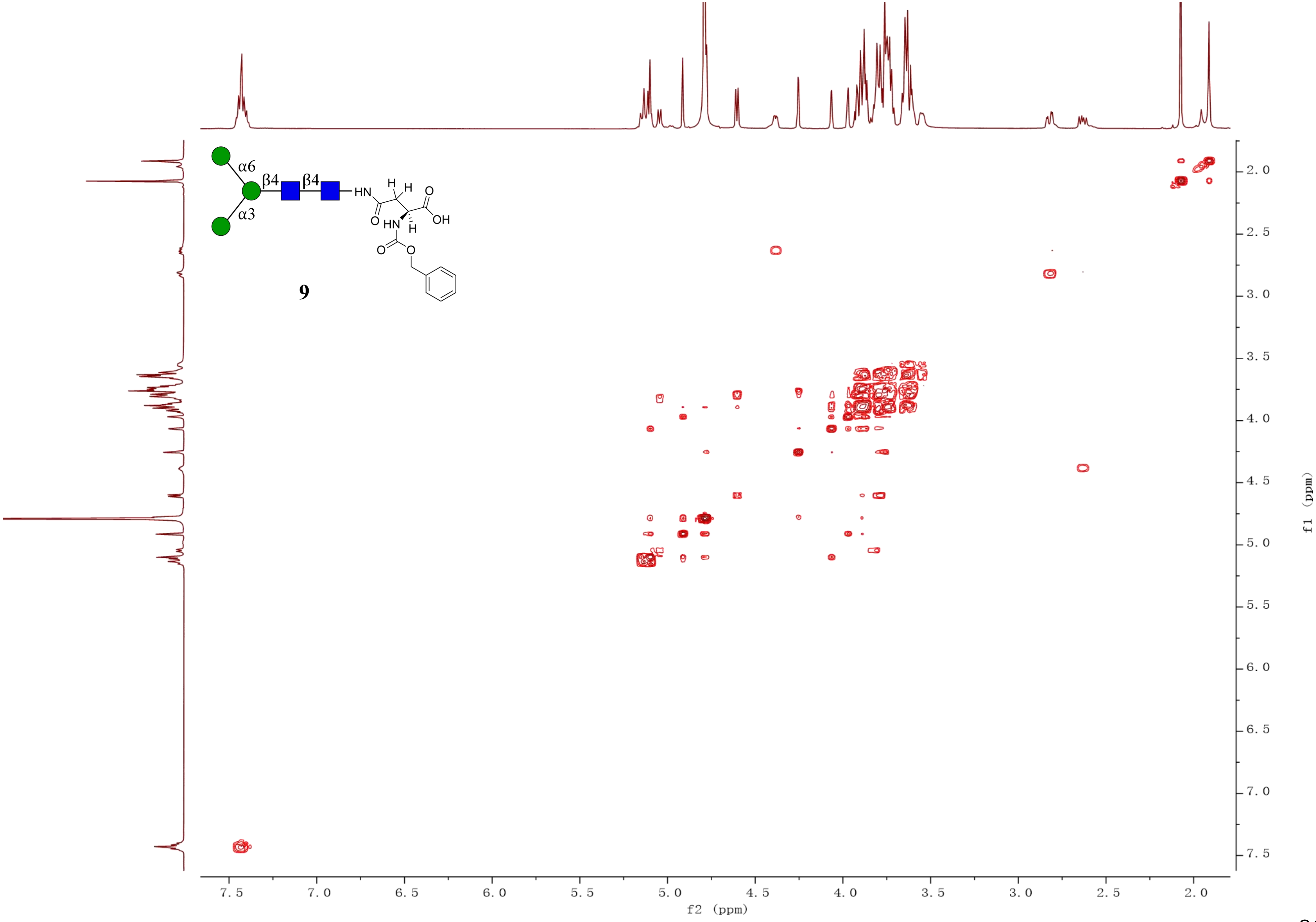
α6
β4
β4
α3
HN
H
H
O
O
OH
HN
H
O
O
9
f2 (ppm)
f1 (ppm)

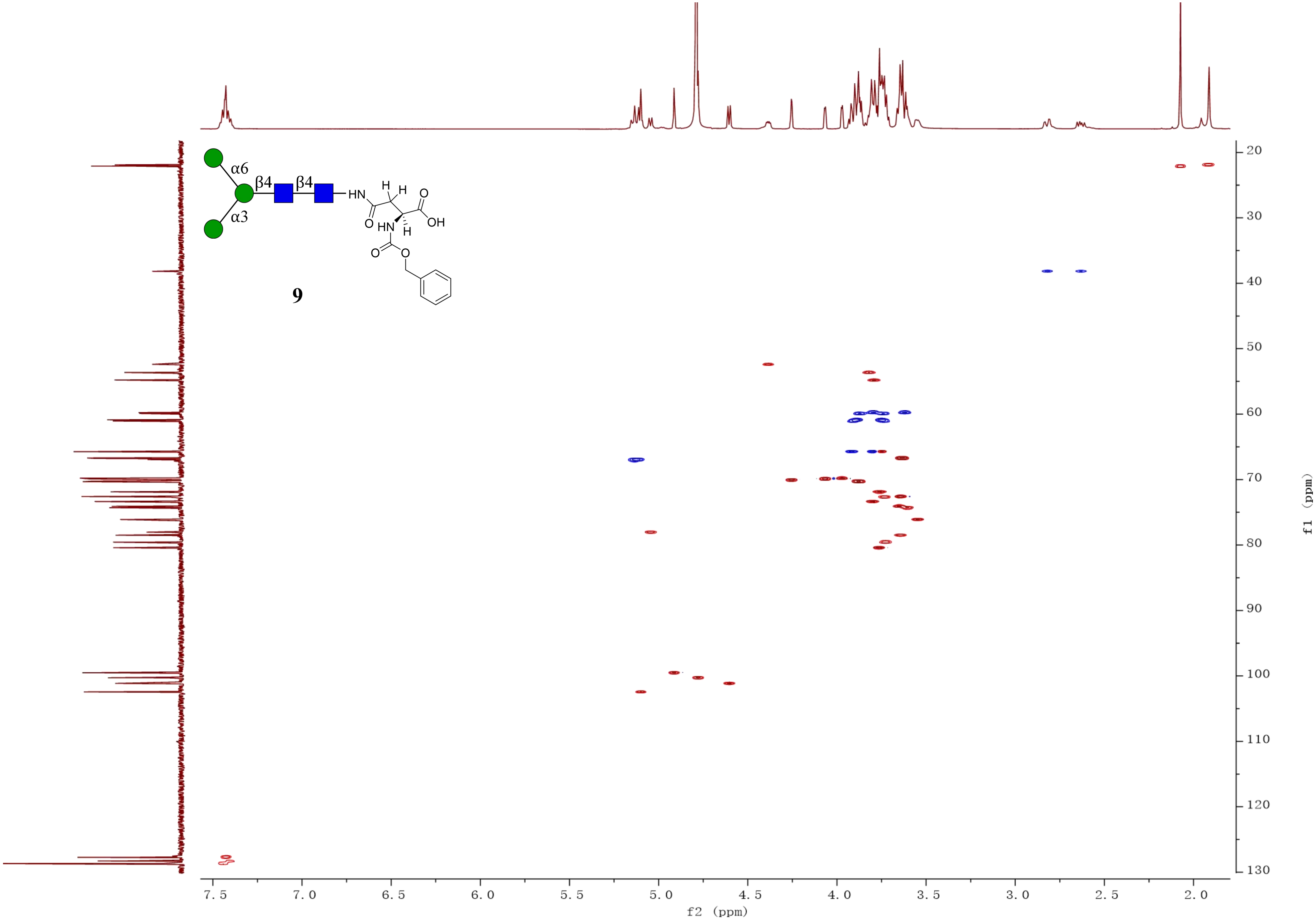
α6
β4
β4
α3
HN
H
H
O
O
OH
HN
H
O
O
9
f2 (ppm)
f1 (ppm)

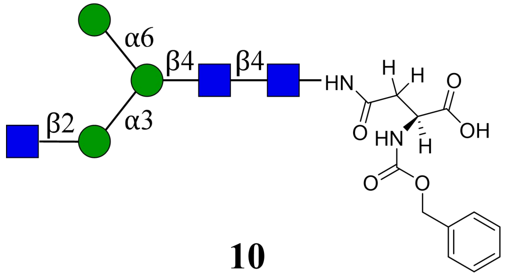

**10**

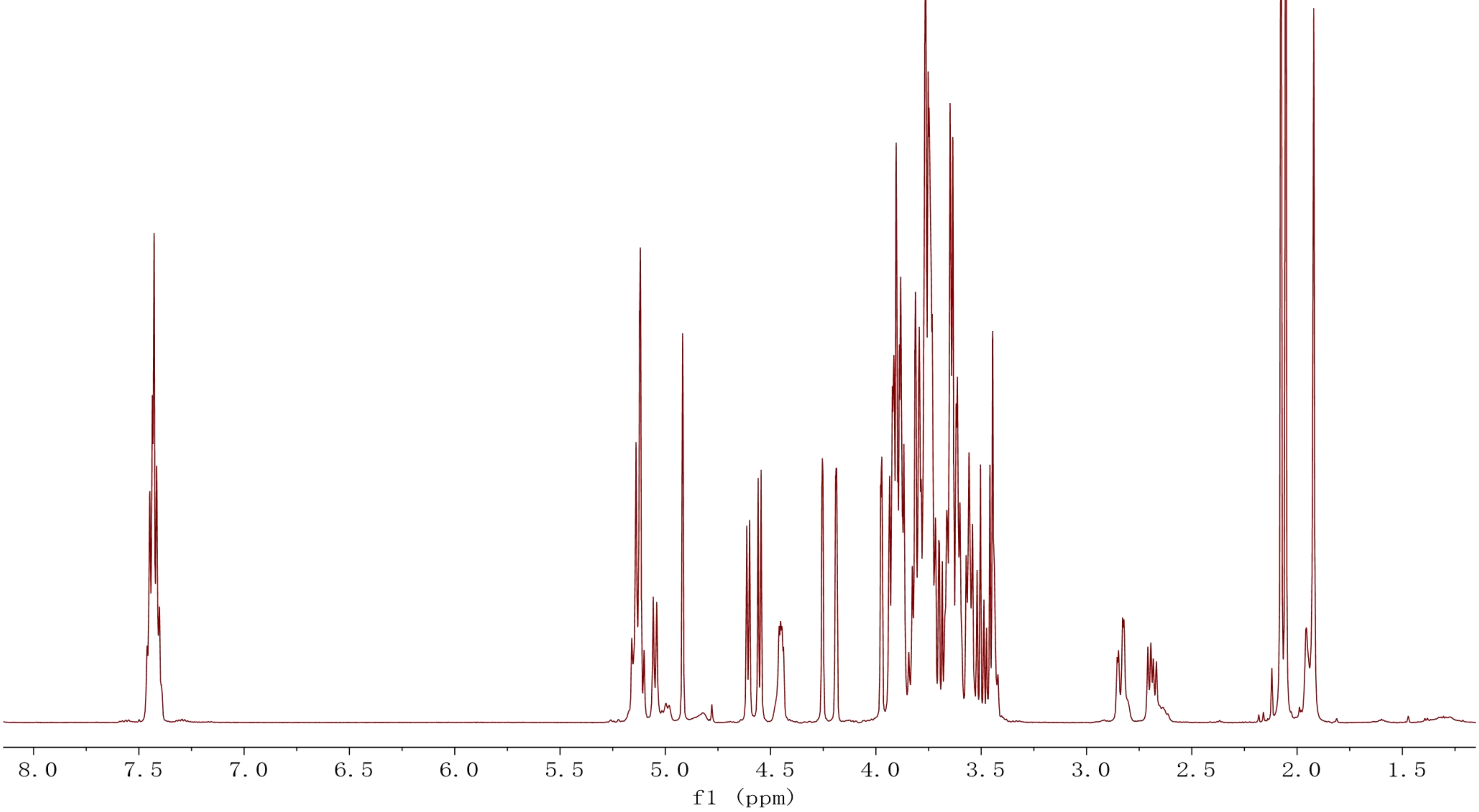

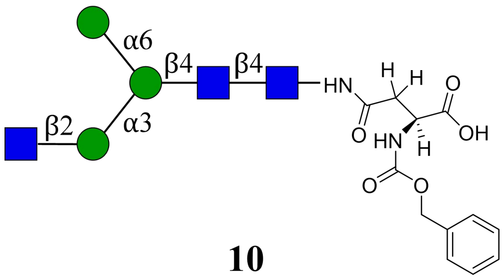
α6
β4
β4
α3
β2
HN
H
H
O
O
OH
HN
H
O
O

**10**

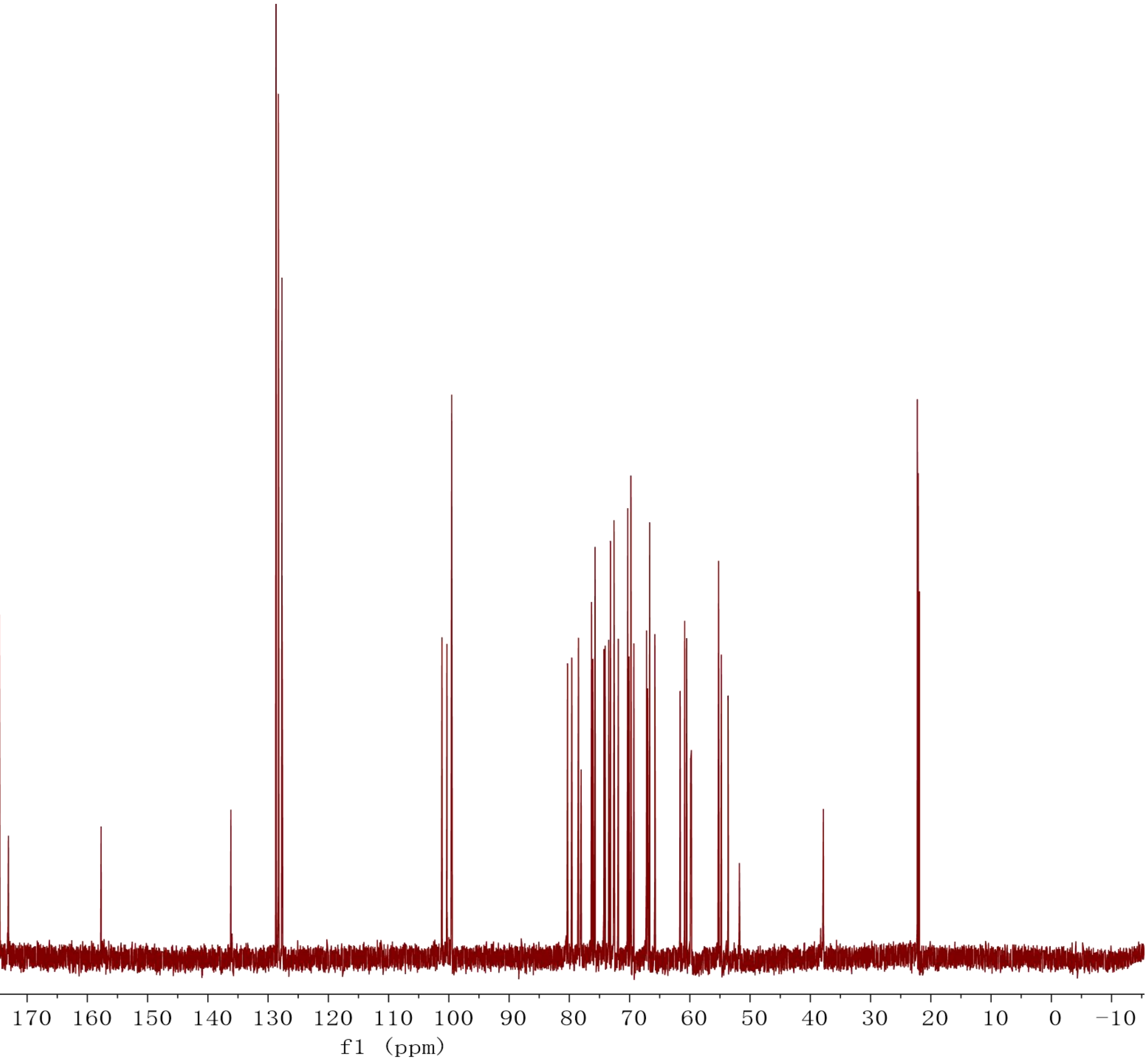
230 220 210 200 190 180 170 160 150 140 130 120 110 100 90 80 70 60 50 40 30 20 10 0 −10
f1 (ppm)

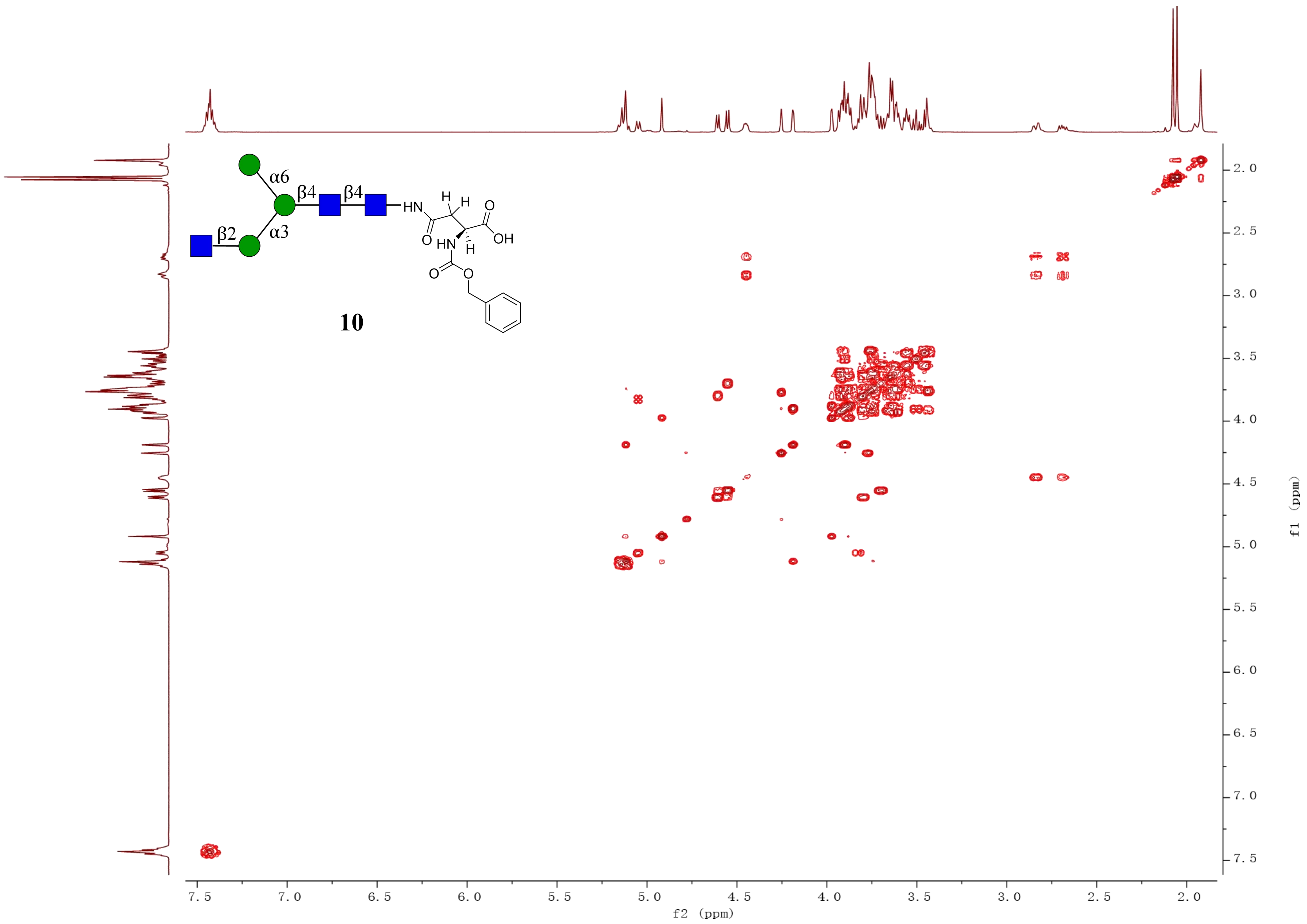
α6
β4
β4
β2
α3
HN
H
H
O
OH
O
HN
H
O
O
10
f2 (ppm)
f1 (ppm)
7.5
7.0
6.5
6.0
5.5
5.0
4.5
4.0
3.5
3.0
2.5
2.0

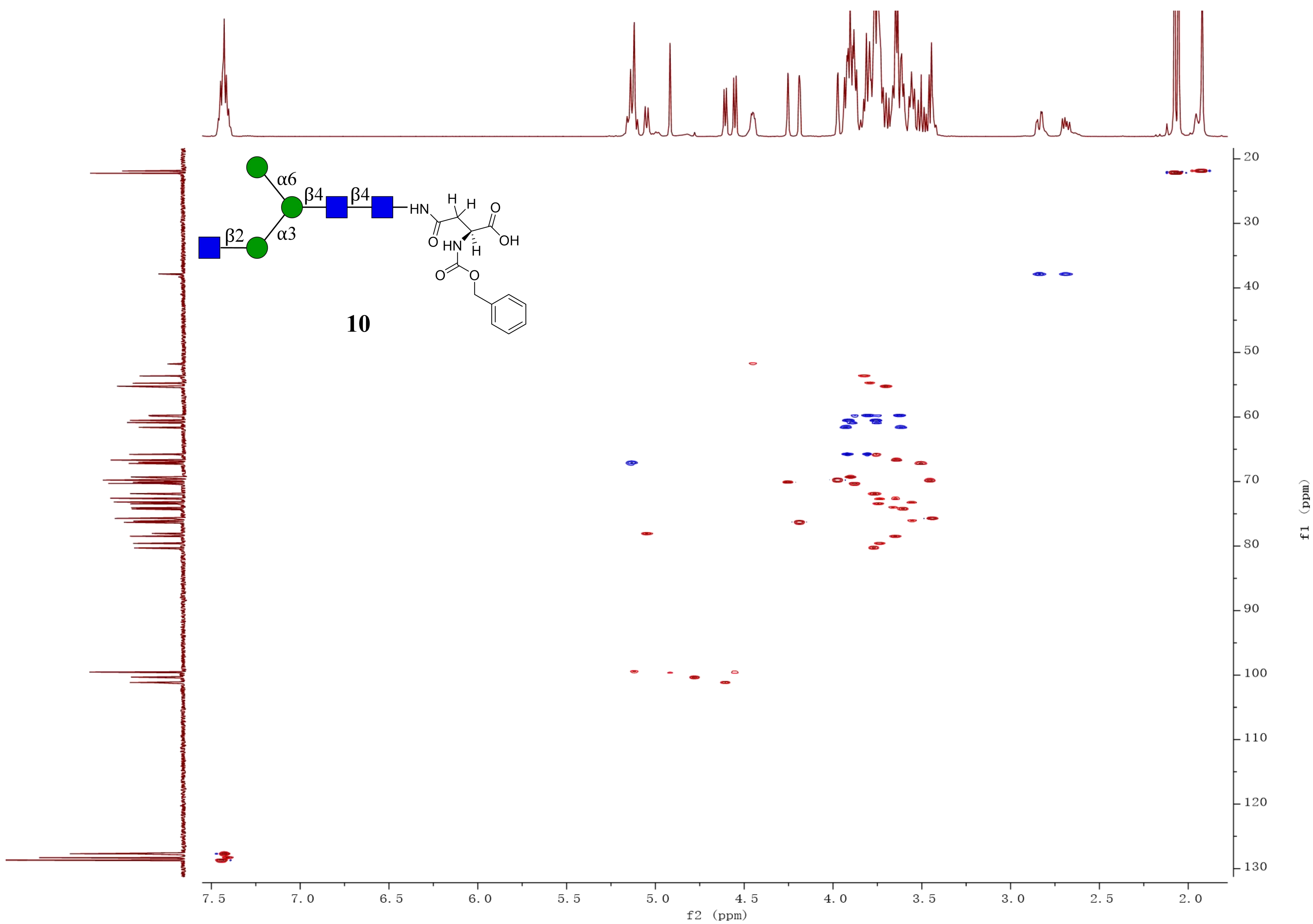

α6
β4
β4
β2
α3
HN
H
H
O
OH
O
HN
H
O
O
10
f2 (ppm)
f1 (ppm)

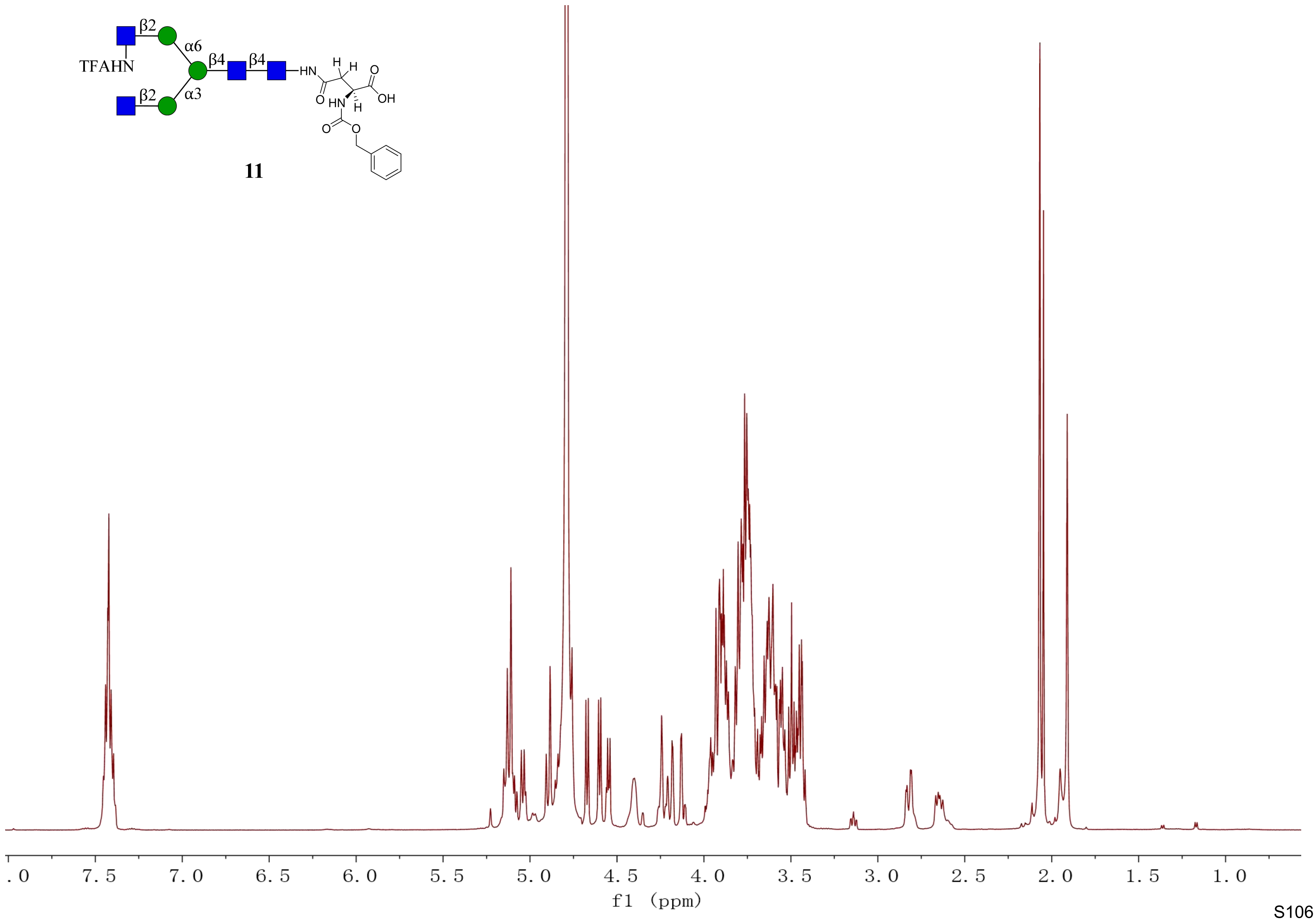
β2
α6
TFAHN
β4
β4
HN
H
H
O
OH
O
HN
H
β2
α3
O
O
11
. 0
7. 5
7. 0
6. 5
6. 0
5. 5
5. 0
4. 5
4. 0
3. 5
3. 0
2. 5
2. 0
1. 5
1. 0
f1 (ppm)

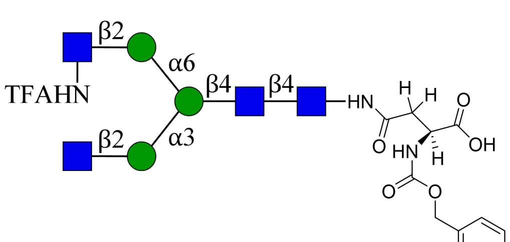
β2
α6
TFAHN
β4
β4
HN
H
H
O
O
OH
HN
H
O
O
α3
β2

**11**

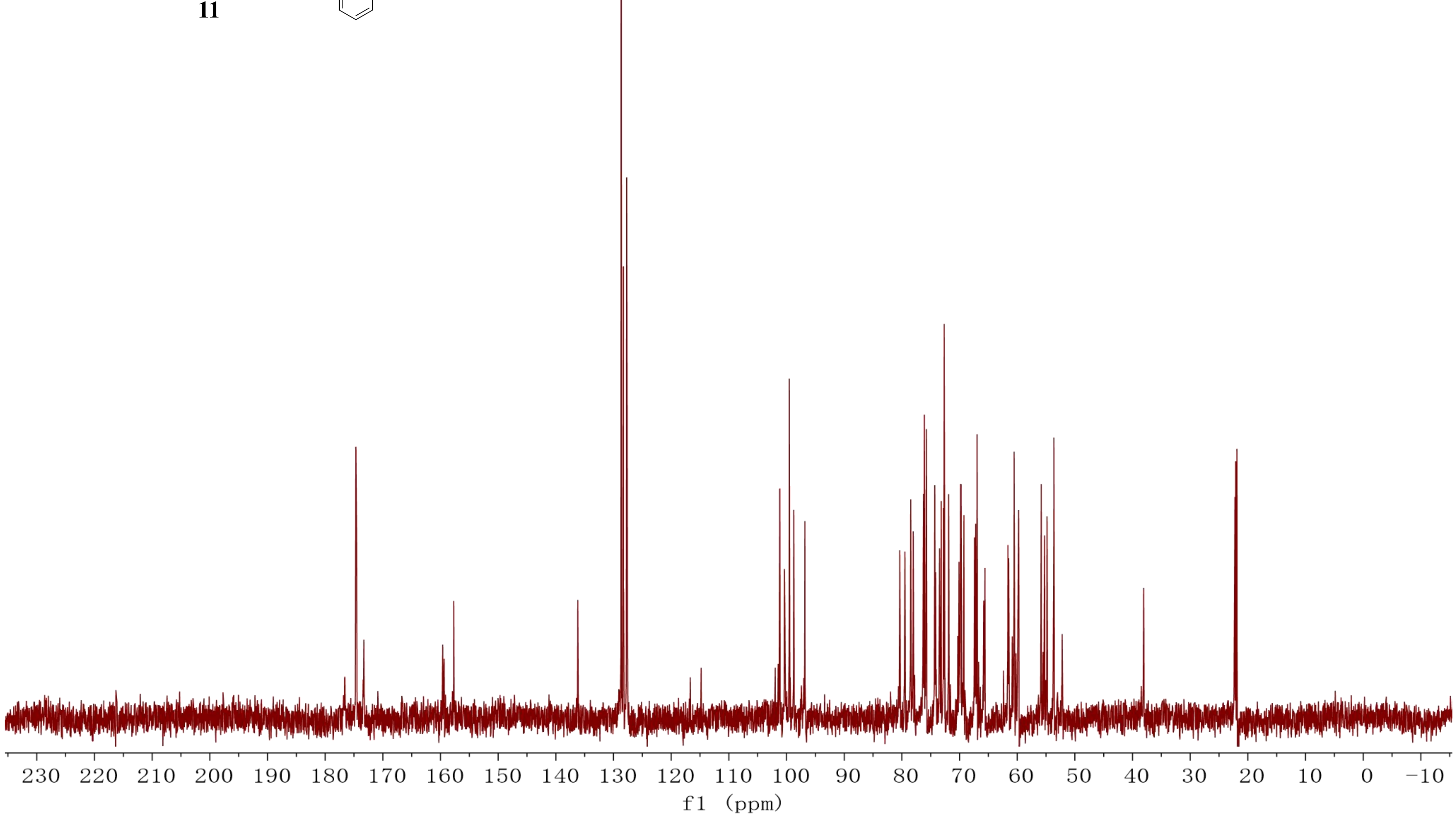
230 220 210 200 190 180 170 160 150 140 130 120 110 100 90 80 70 60 50 40 30 20 10 0 −10
f1 (ppm)

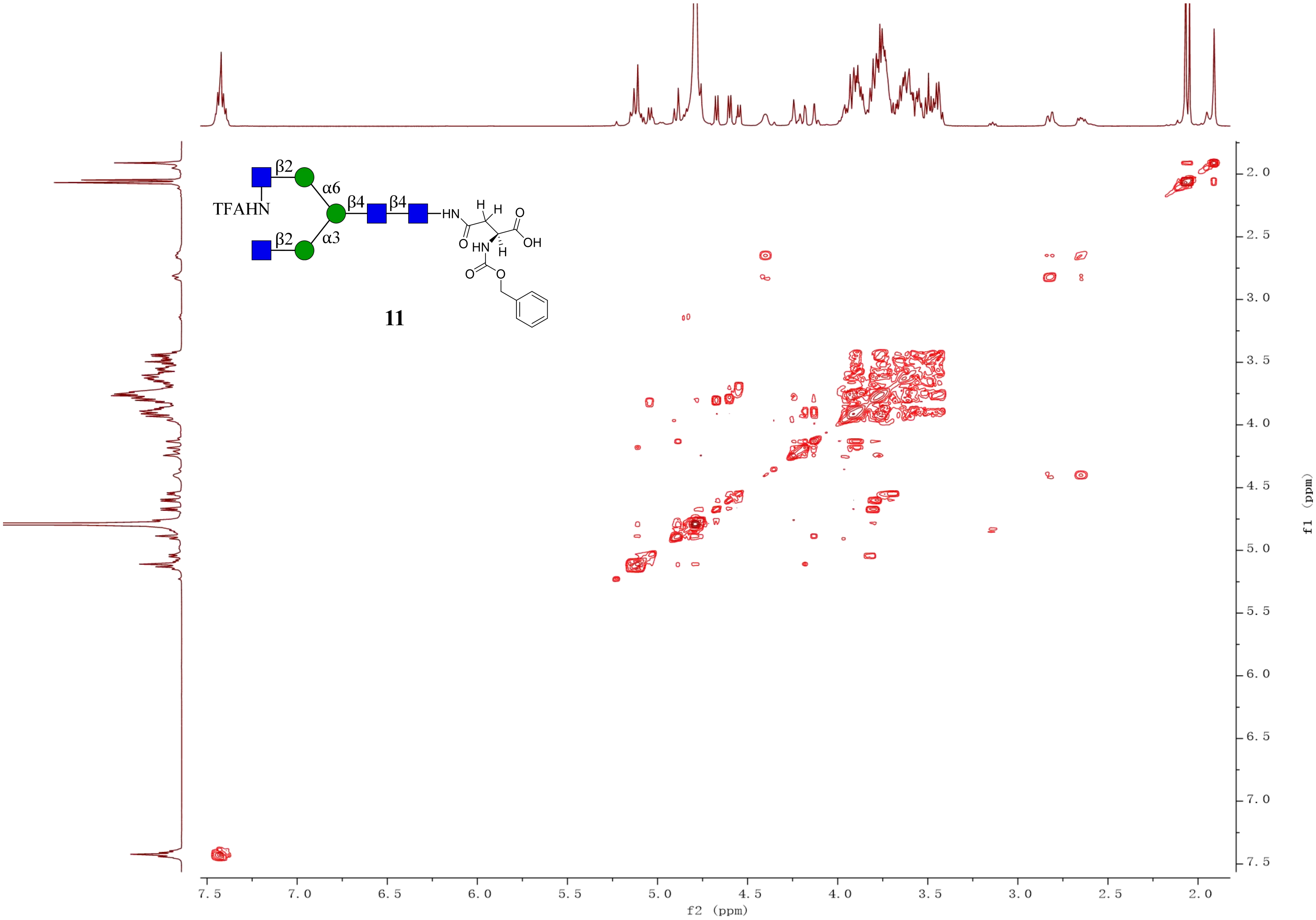
β2
α6
TFAHN
β4
β4
HN
H
H
O
OH
O
HN
H
β2
α3
O
O
11
7.5
7.0
6.5
6.0
5.5
5.0
4.5
4.0
3.5
3.0
2.5
2.0
f2 (ppm)
f1 (ppm)

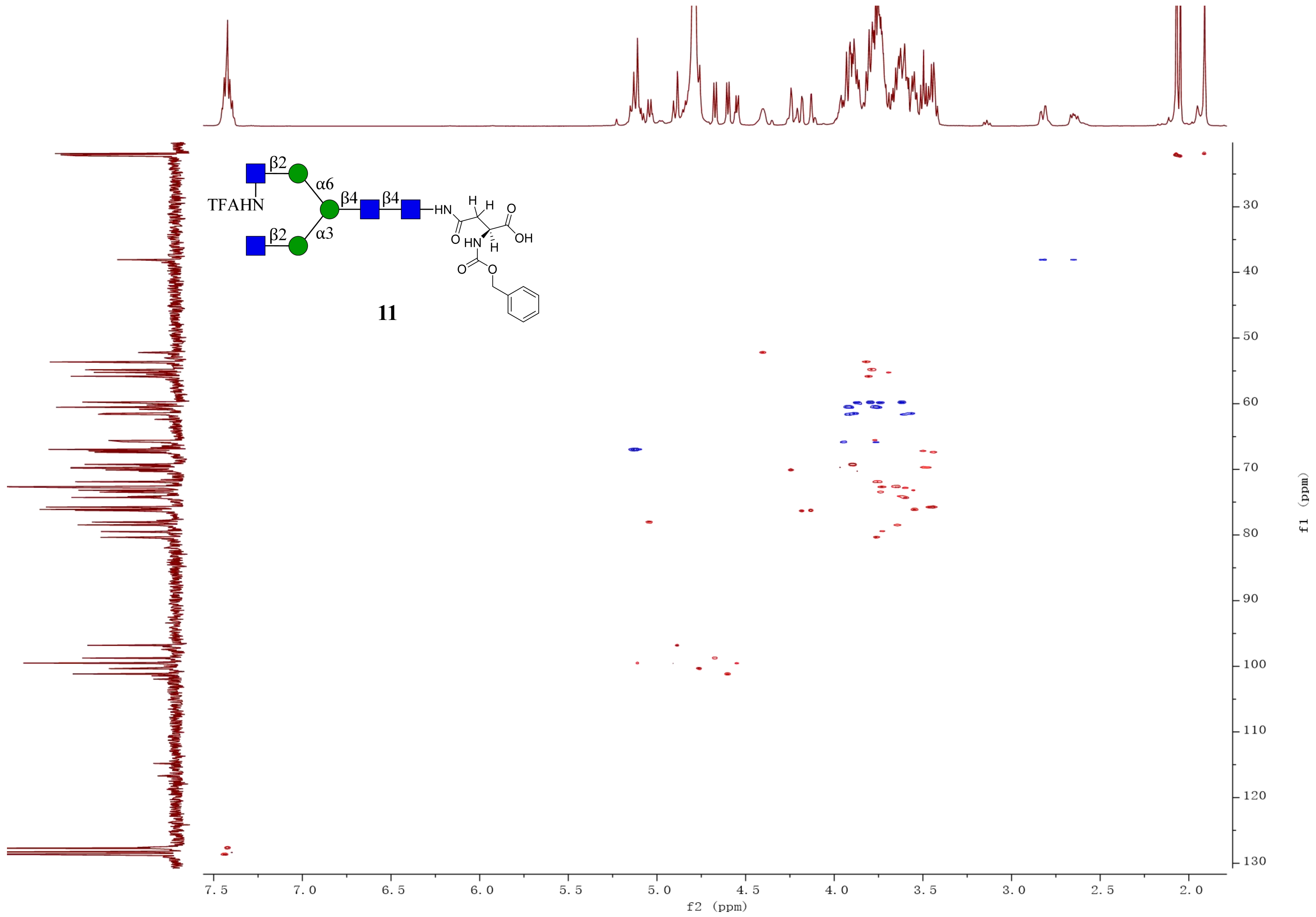

β2
α6
β4
β4
TFAHN
HN
H
H
O
OH
O
HN
H
β2
α3
O
O
11
f2 (ppm)
f1 (ppm)

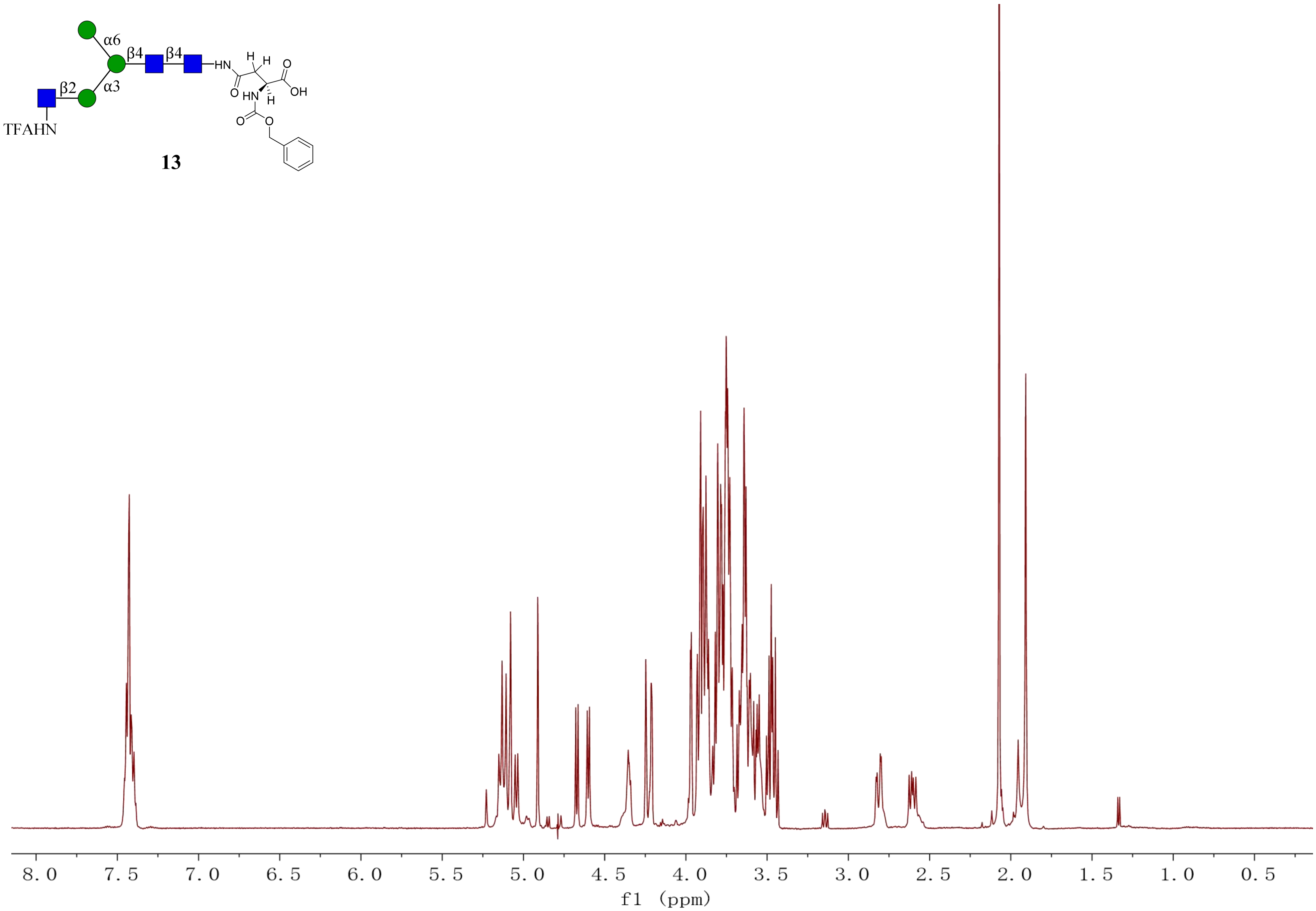
α6
β4
β4
α3
β2
TFAHN
HN
H
H
O
O
OH
HN
H
O
O
13
8.0
7.5
7.0
6.5
6.0
5.5
5.0
4.5
4.0
3.5
3.0
2.5
2.0
1.5
1.0
0.5
f1 (ppm)

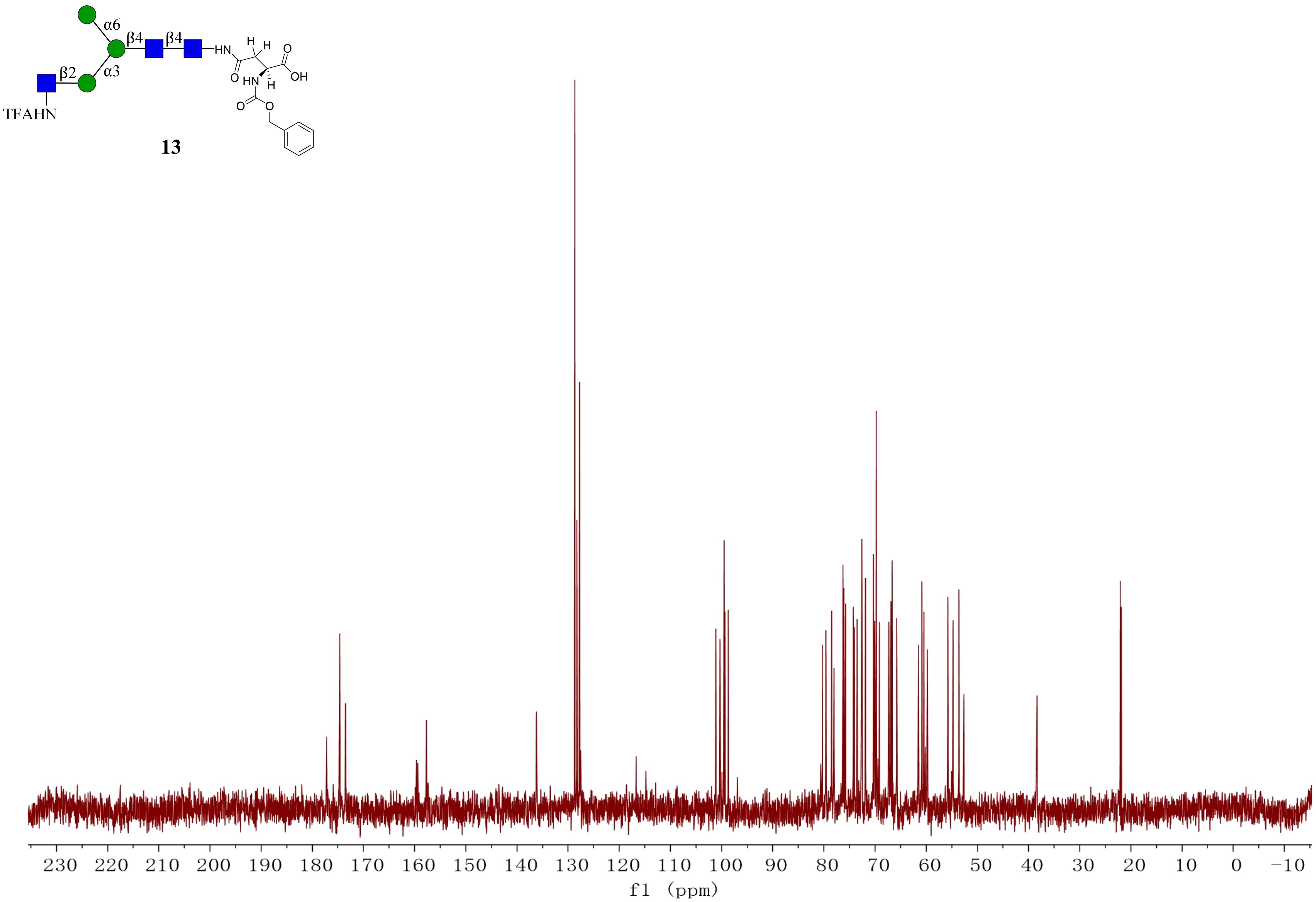
α6
β4
β4
β2
α3
TFAHN
HN
H
H
O
O
OH
HN
H
O
O
13
230 220 210 200 190 180 170 160 150 140 130 120 110 100 90 80 70 60 50 40 30 20 10 0 −10
f1 (ppm)

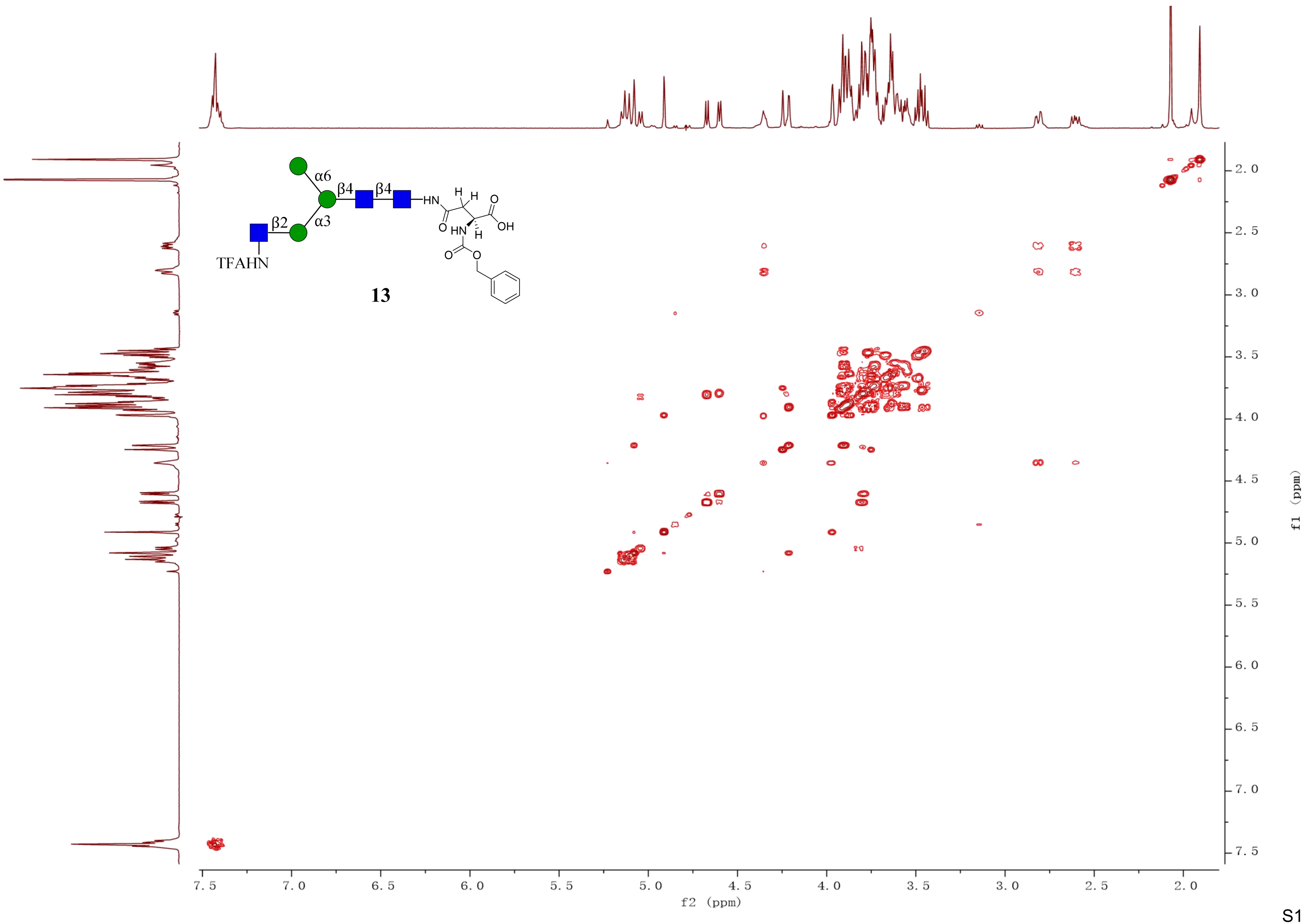
α6
β4
β4
α3
β2
TFAHN
HN
H
H
O
OH
O
HN
H
O
O
13
f2 (ppm)
f1 (ppm)

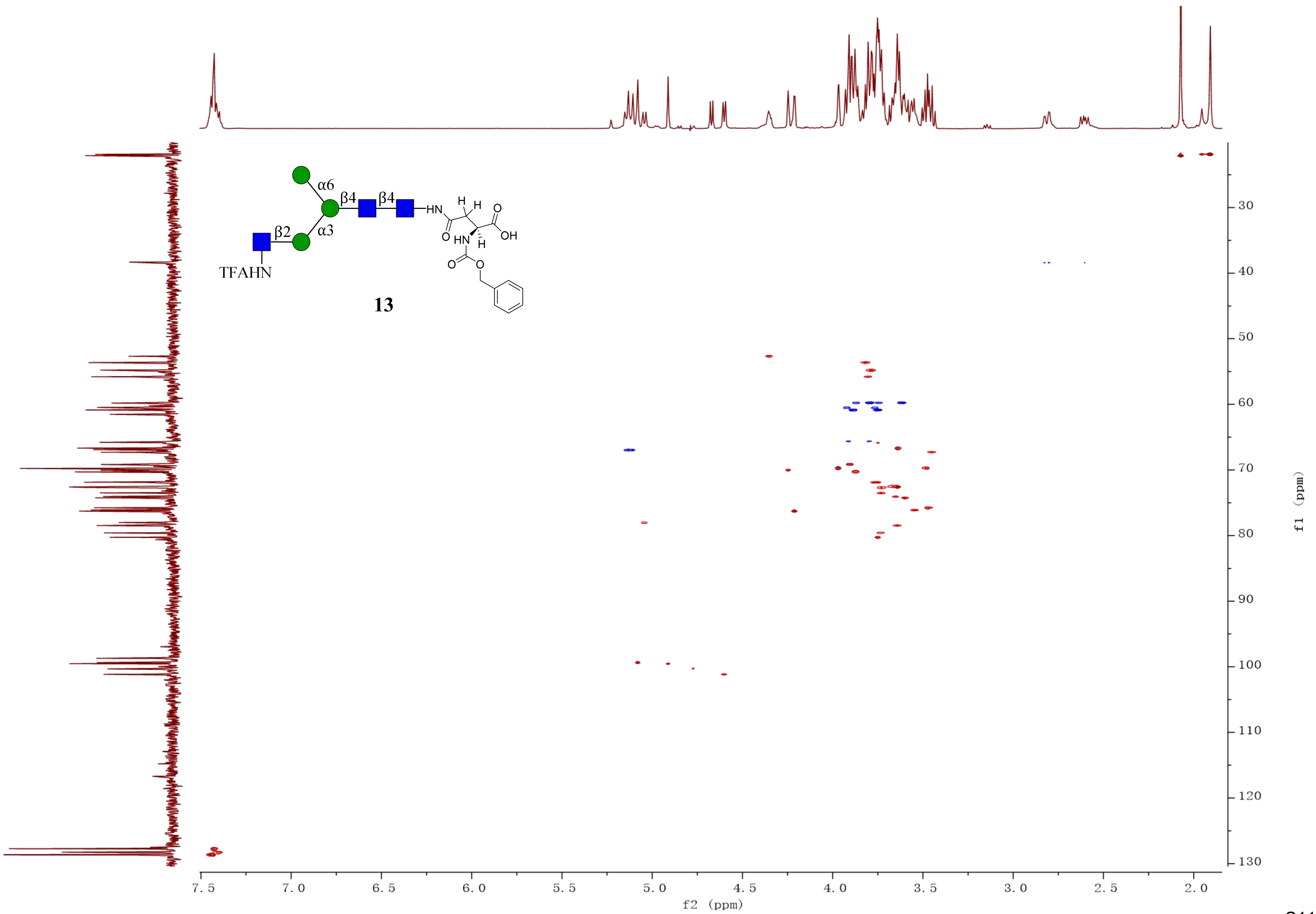

α6
β4
β4
β2
α3
HN
H
H
O
OH
O
HN
H
O
O
TFAHN
13
7.5
7.0
6.5
6.0
5.5
5.0
4.5
4.0
3.5
3.0
2.5
2.0
f2 (ppm)
30
40
50
60
70
80
90
100
110
120
130
f1 (ppm)

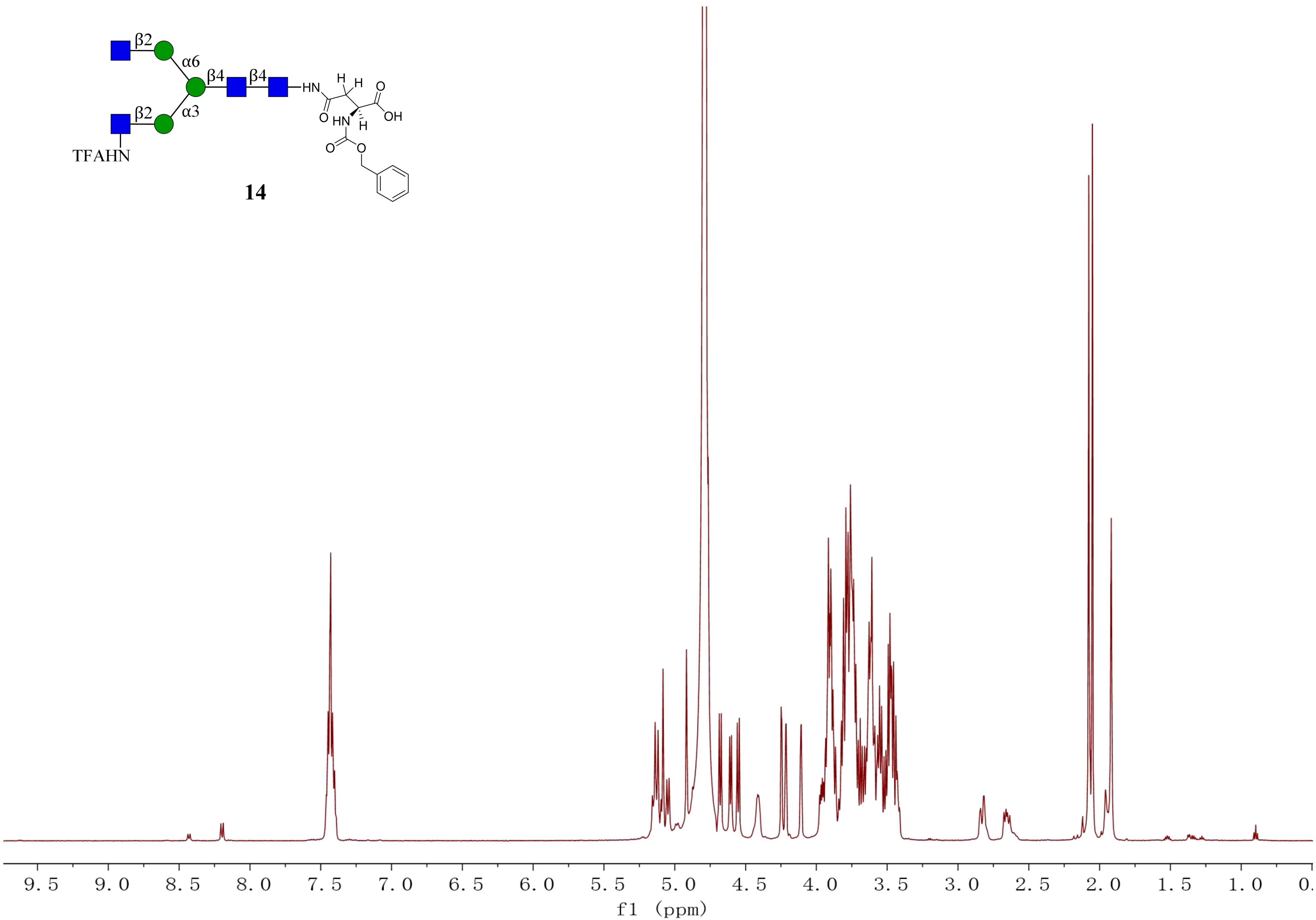
β2
α6
β4
β4
HN
H
H
O
OH
O
HN
H
α3
β2
TFAHN
O
O
9.5
9.0
8.5
8.0
7.5
7.0
6.5
6.0
5.5
5.0
4.5
4.0
3.5
3.0
2.5
2.0
1.5
1.0
f1 (ppm)

**14**

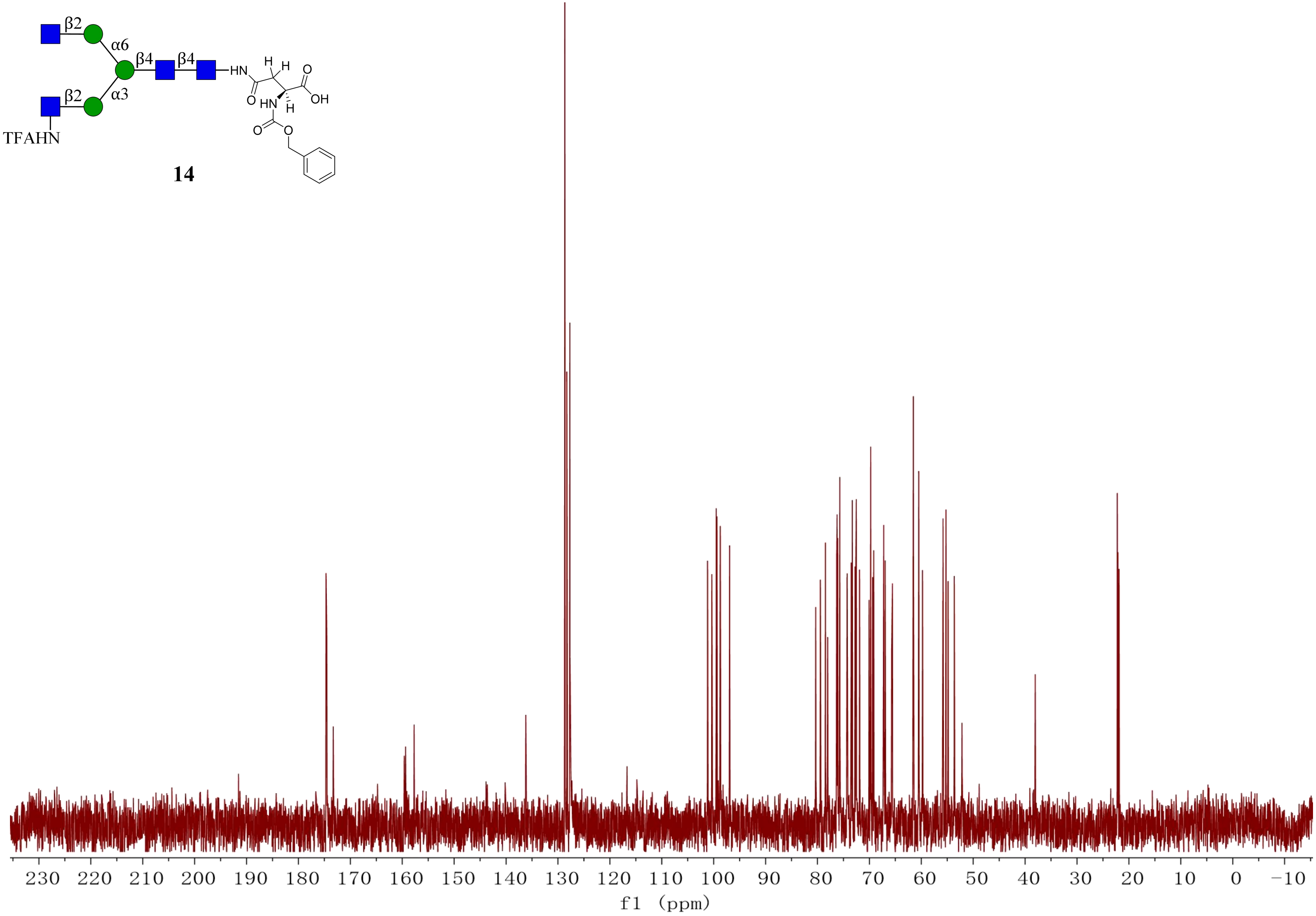
β2
α6
β4
β4
HN
H
H
O
OH
O
HN
H
β2
α3
O
O
TFAHN
230
220
210
200
190
180
170
160
150
140
130
120
110
100
90
80
70
60
50
40
30
20
10
0
−10
f1 (ppm)

**14**

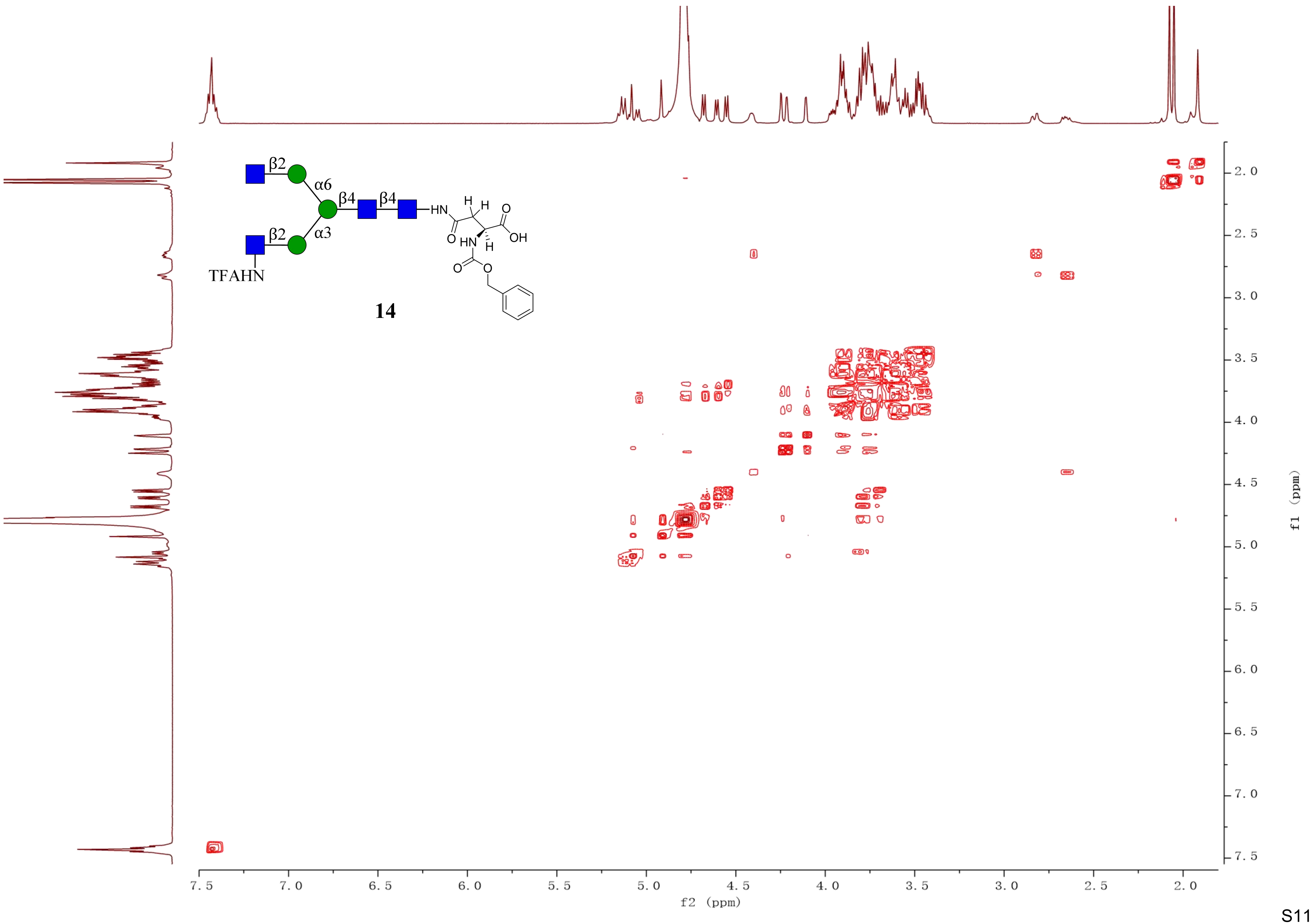
β2
α6
β4
β4
HN
H
H
O
OH
O
HN
H
β2
α3
TFAHN
O
O
14
7.5
7.0
6.5
6.0
5.5
5.0
4.5
4.0
3.5
3.0
2.5
2.0
f2 (ppm)
f1 (ppm)

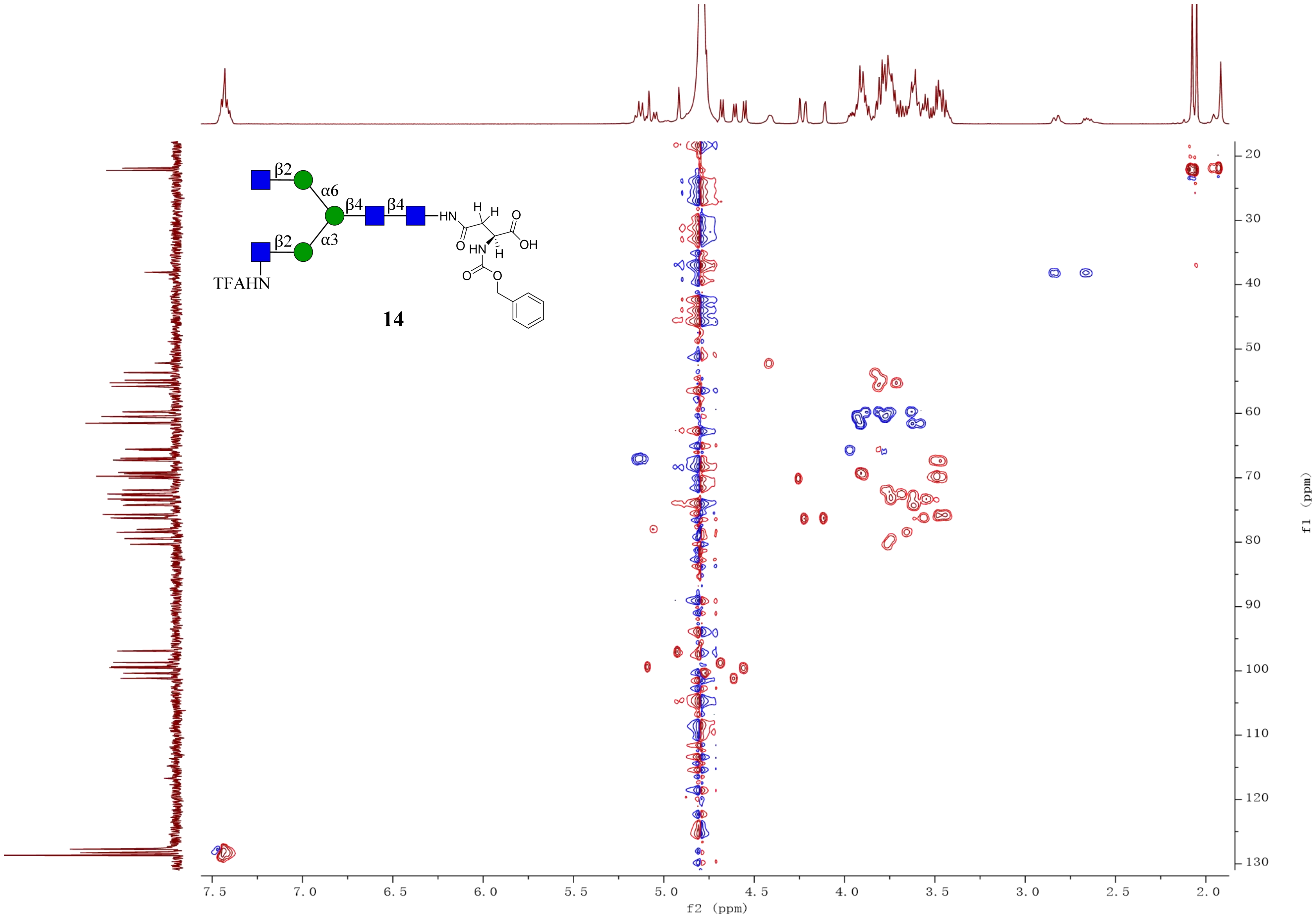
β2
α6
β4
β4
HN
H
H
O
OH
O
HN
H
O
O
β2
α3
TFAHN
14
f2 (ppm)
f1 (ppm)

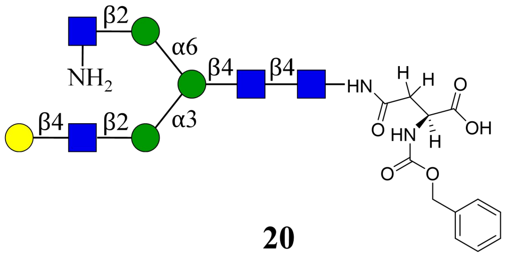
β2
α6
NH2
β4
β4
HN
H
H
O
OH
β4
β2
α3
O
HN
H
O
O

**20**

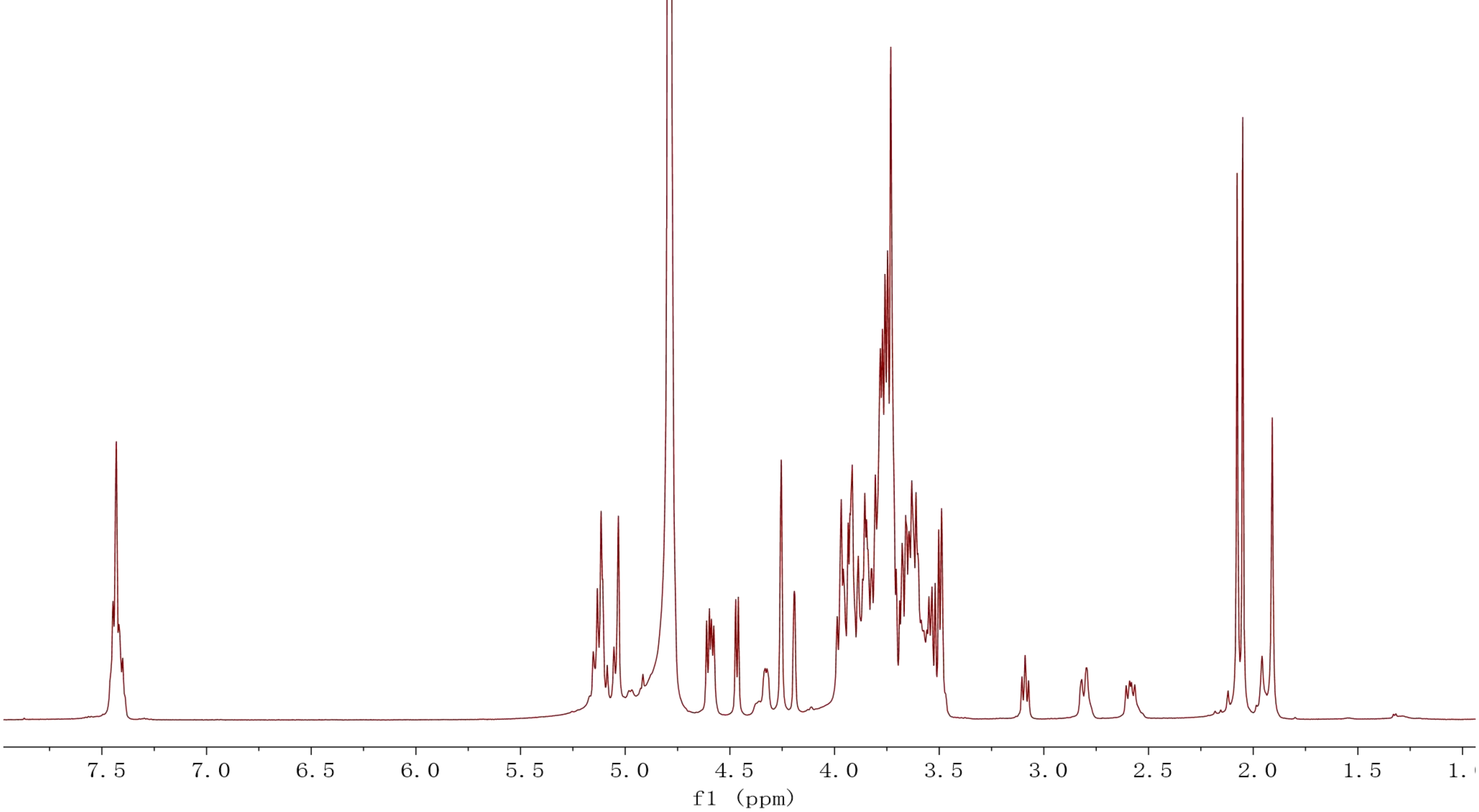
7.5
7.0
6.5
6.0
5.5
5.0
4.5
4.0
3.5
3.0
2.5
2.0
1.5
f1 (ppm)

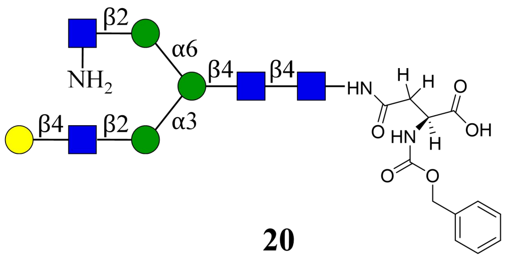
β2
α6
NH2
β4
β4
HN
H
H
O
O
OH
HN
H
β4
β2
α3
O
O

**20**

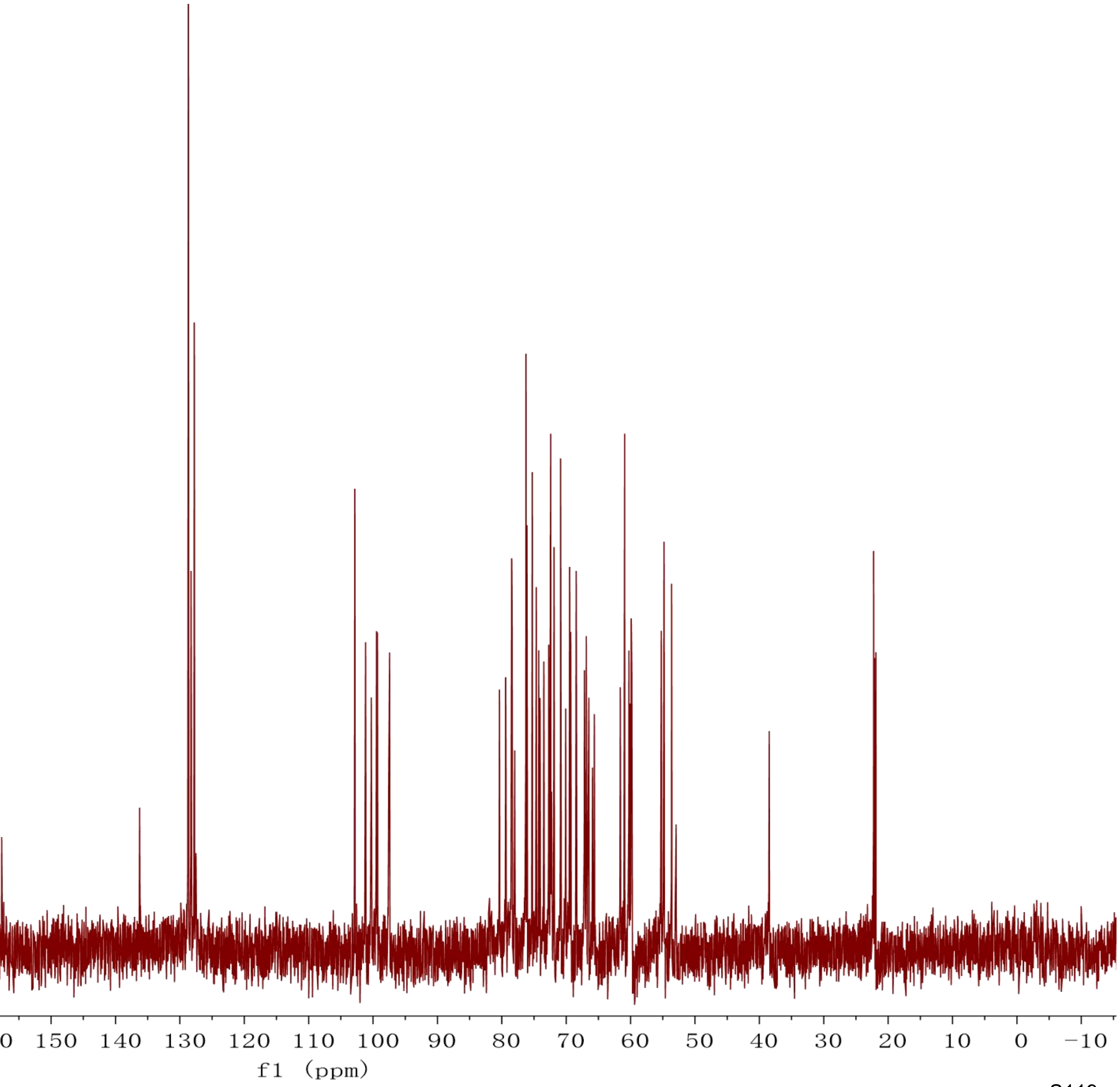
230 220 210 200 190 180 170 160 150 140 130 120 110 100 90 80 70 60 50 40 30 20 10 0 −10
f1 (ppm)

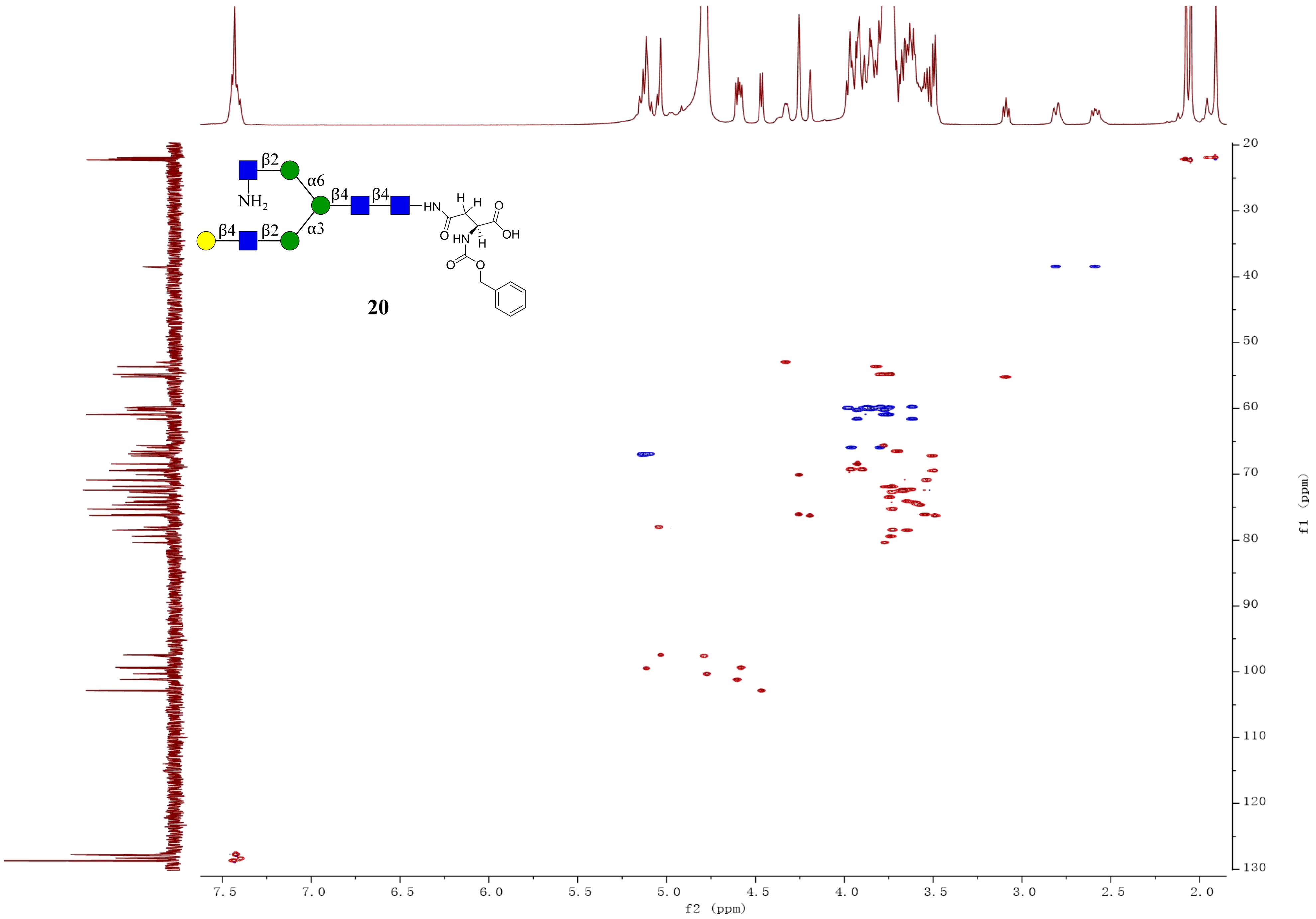
β2
α6
β4
β4
NH2
HN
H
H
O
OH
O
HN
H
O
O
β4
β2
α3
20
f2 (ppm)
f1 (ppm)

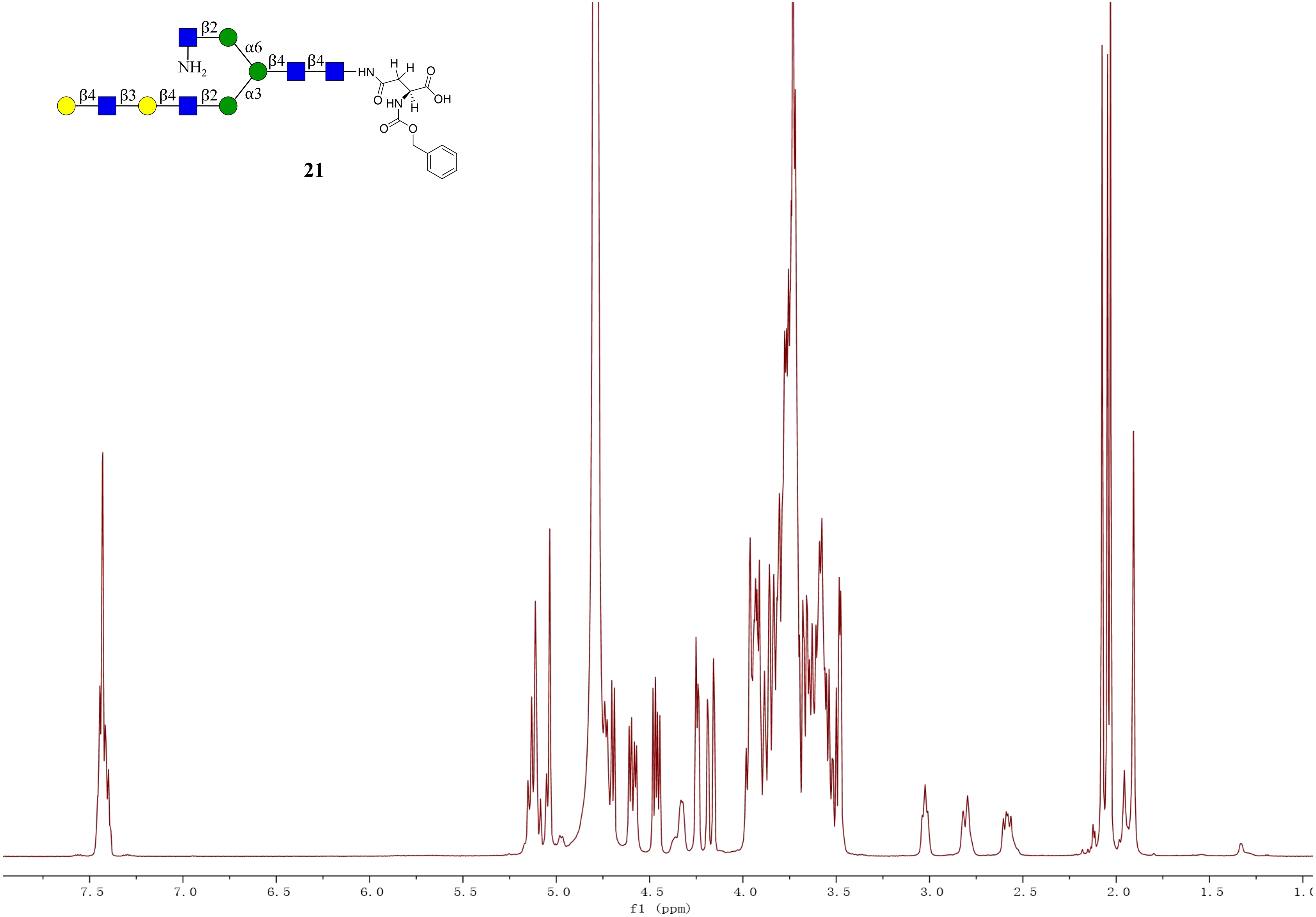
β2
α6
β4
β4
HN
H
H
O
NH$_2$
OH
β4
β3
β4
β2
α3
O
HN
H
O
O
21
7.5
7.0
6.5
6.0
5.5
5.0
4.5
4.0
3.5
3.0
2.5
2.0
1.5
f1 (ppm)

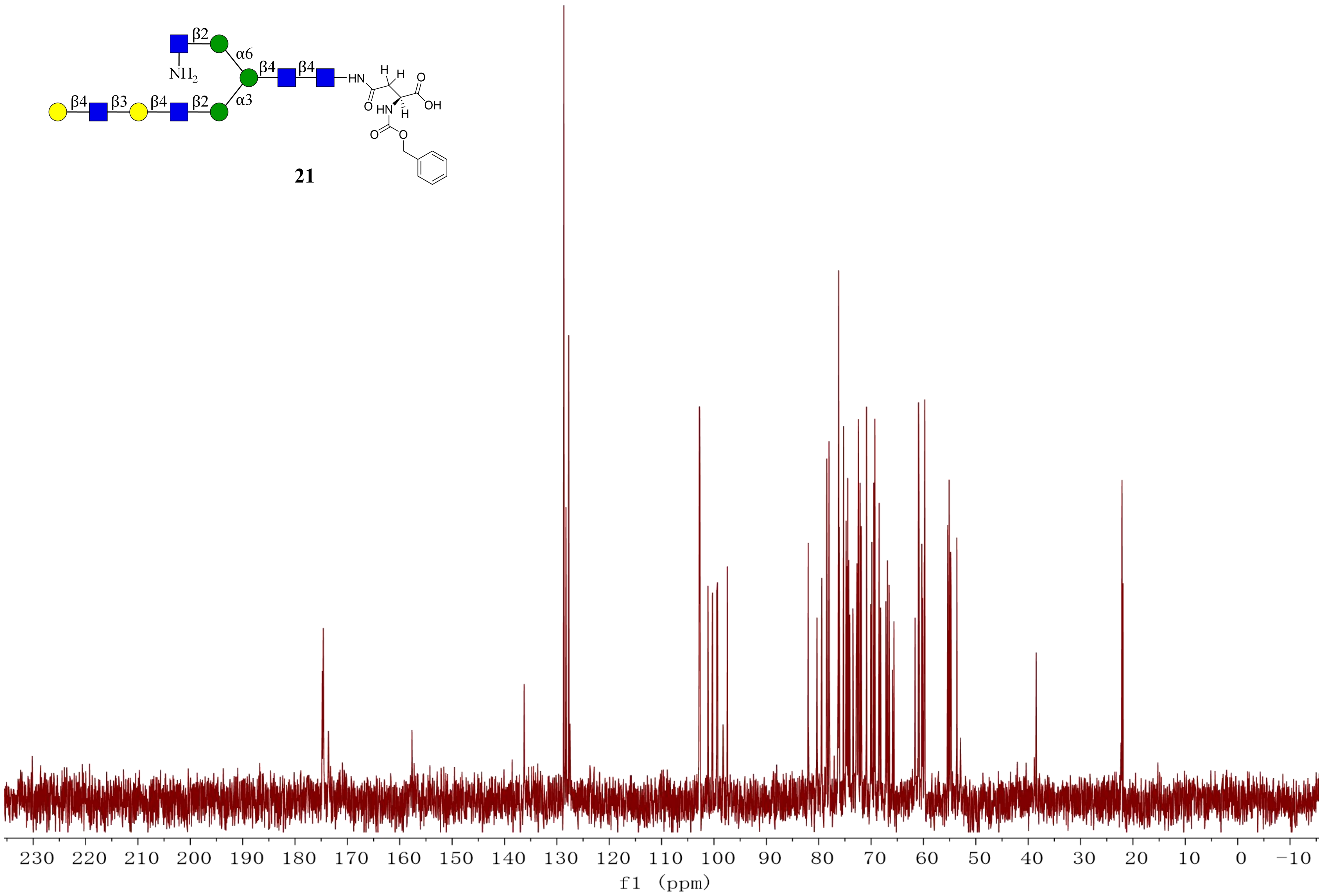

β2
α6
β4
β4
NH2
HN
H
H
O
OH
β4
β3
β4
β2
α3
O
HN
H
O
O
230 220 210 200 190 180 170 160 150 140 130 120 110 100 90 80 70 60 50 40 30 20 10 0 −10
f1 (ppm)


21

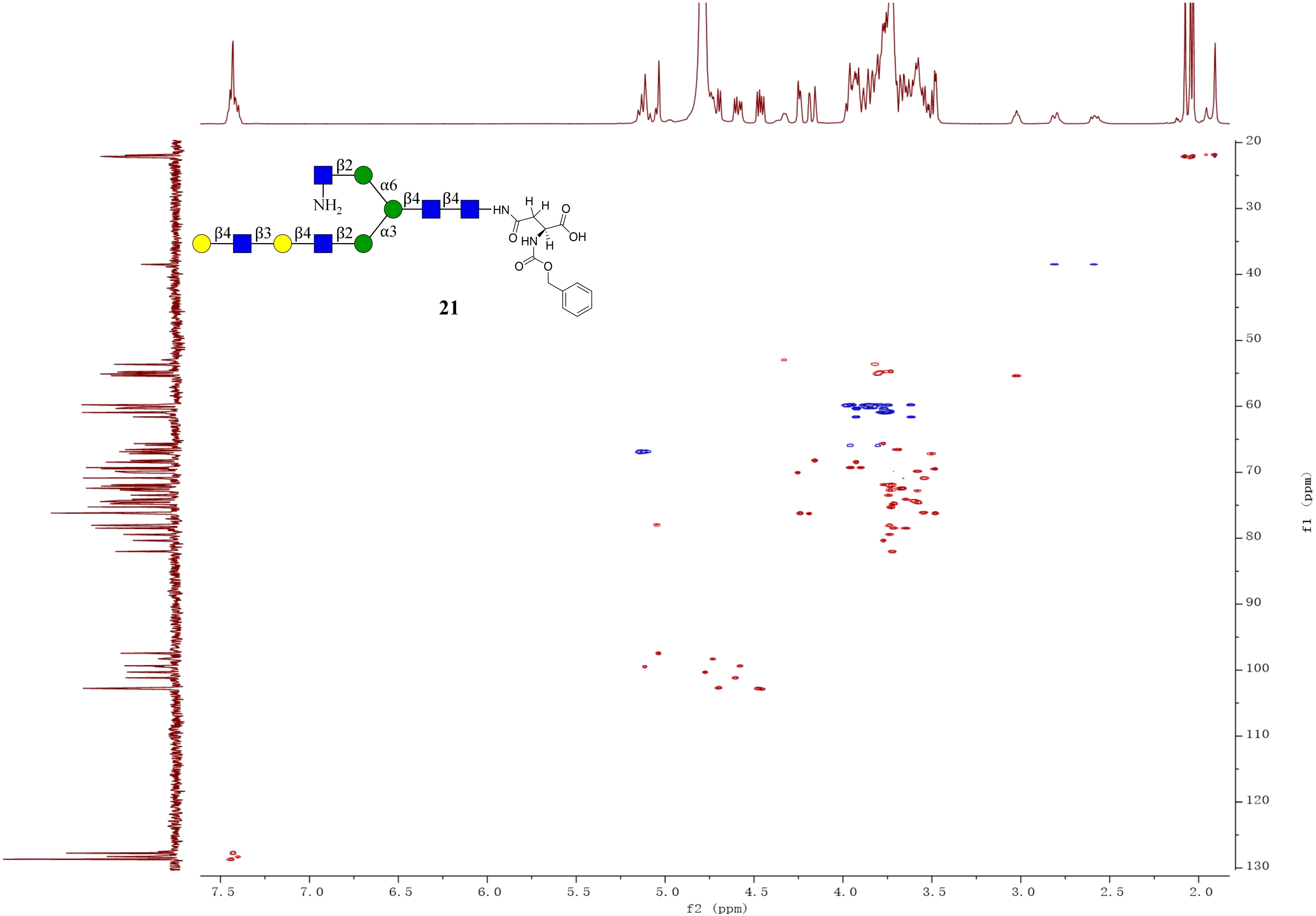
β2
α6
NH2
β4
β4
HN
H
H
O
OH
O
HN
H
O
O
β4
β3
β4
β2
α3
21
f2 (ppm)
f1 (ppm)
7.5
7.0
6.5
6.0
5.5
5.0
4.5
4.0
3.5
3.0
2.5
2.0
20
30
40
50
60
70
80
90
100
110
120
130

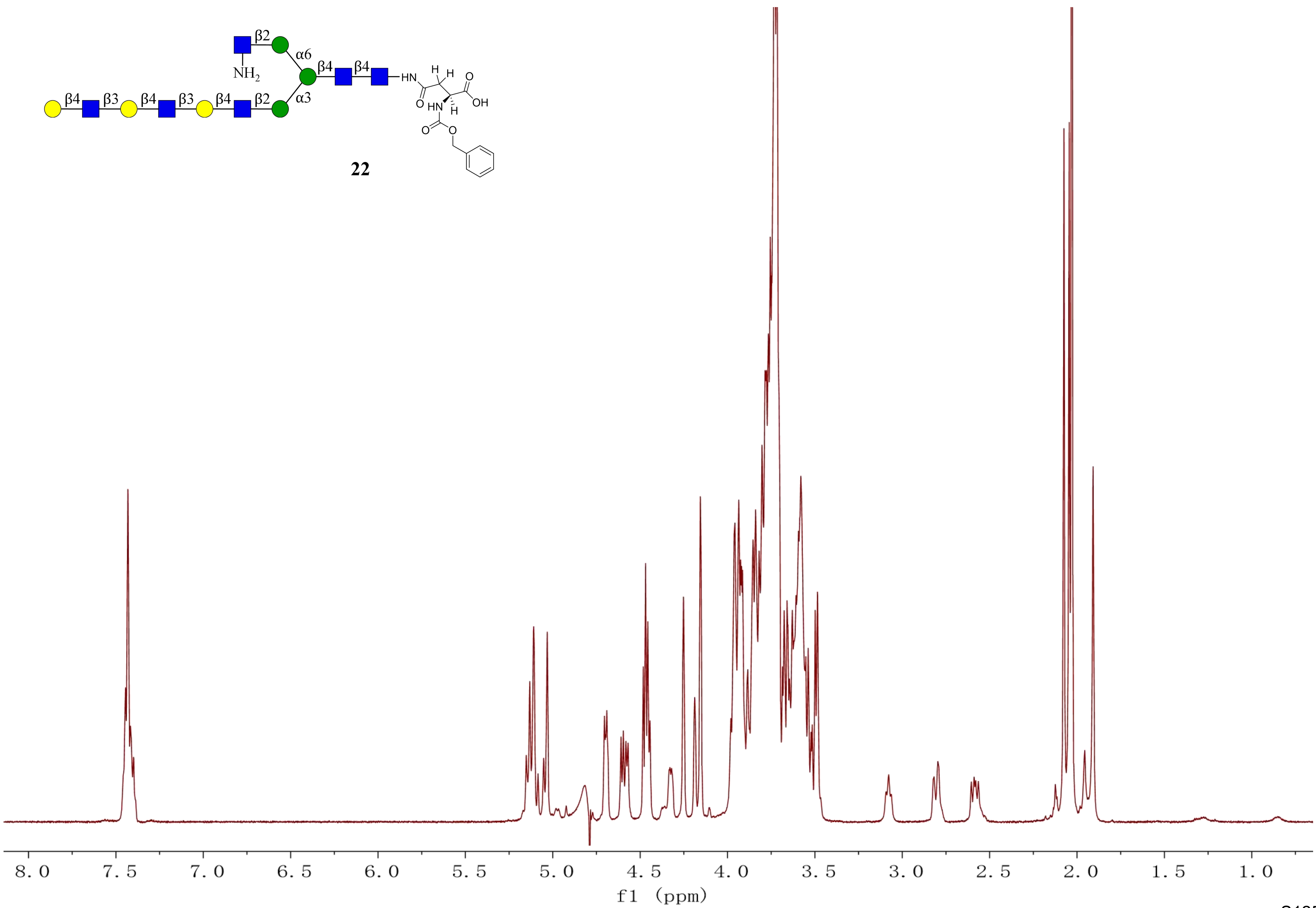
β2
α6
NH2
β4
β4
HN
H
H
O
β4
β3
β4
β3
β4
β2
α3
O
HN
H
OH
O
O
22
8.0
7.5
7.0
6.5
6.0
5.5
5.0
4.5
4.0
3.5
3.0
2.5
2.0
1.5
1.0
f1 (ppm)

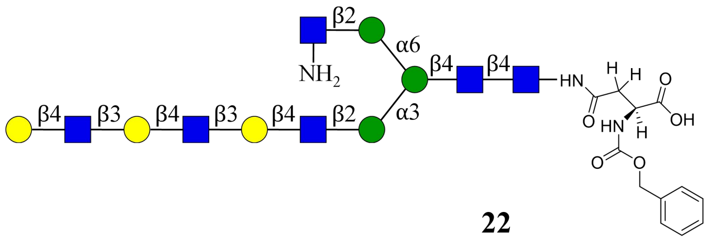
β2
α6
β4
β4
NH2
HN
H
H
O
O
OH
HN
H
O
O
β4
β3
β4
β3
β4
β2
α3

**22**

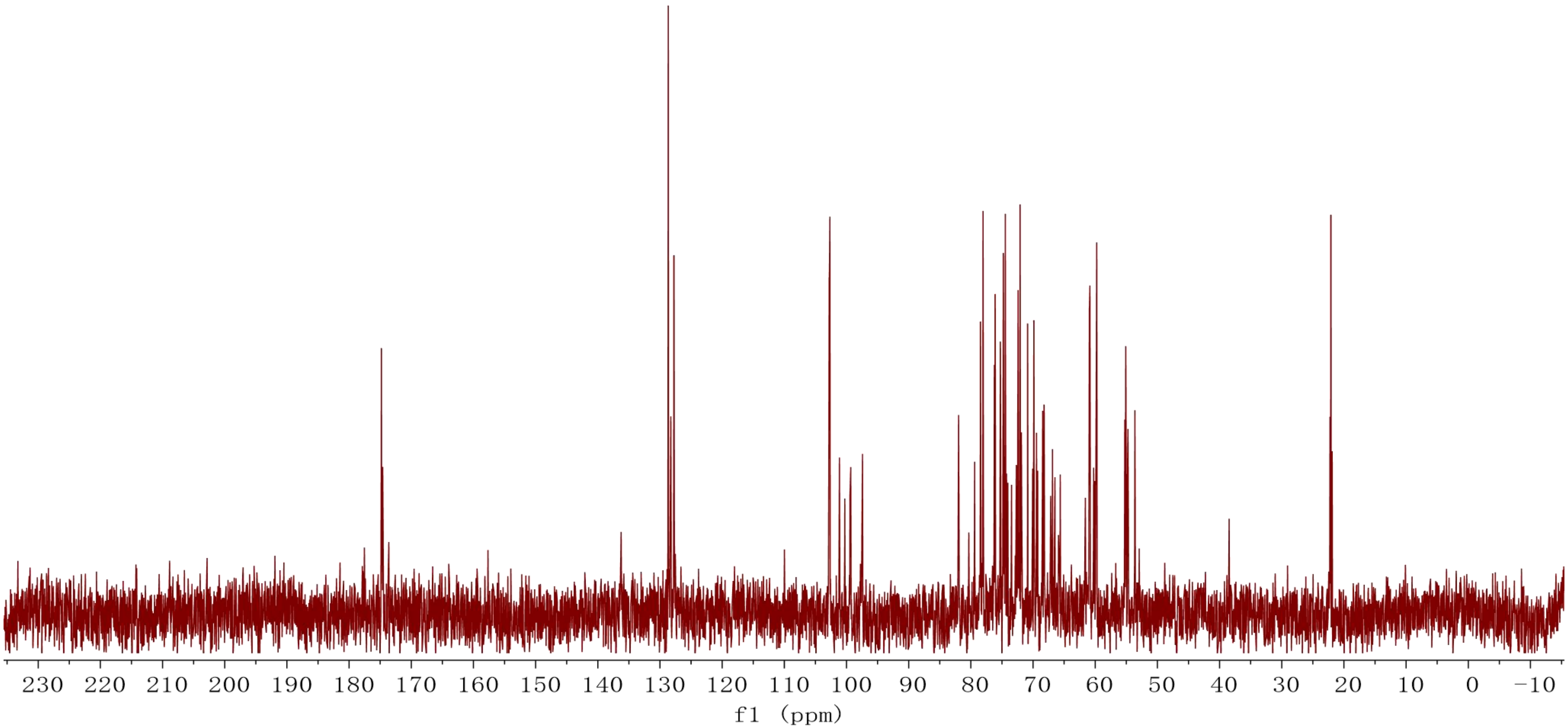
230 220 210 200 190 180 170 160 150 140 130 120 110 100 90 80 70 60 50 40 30 20 10 0 −10
f1 (ppm)

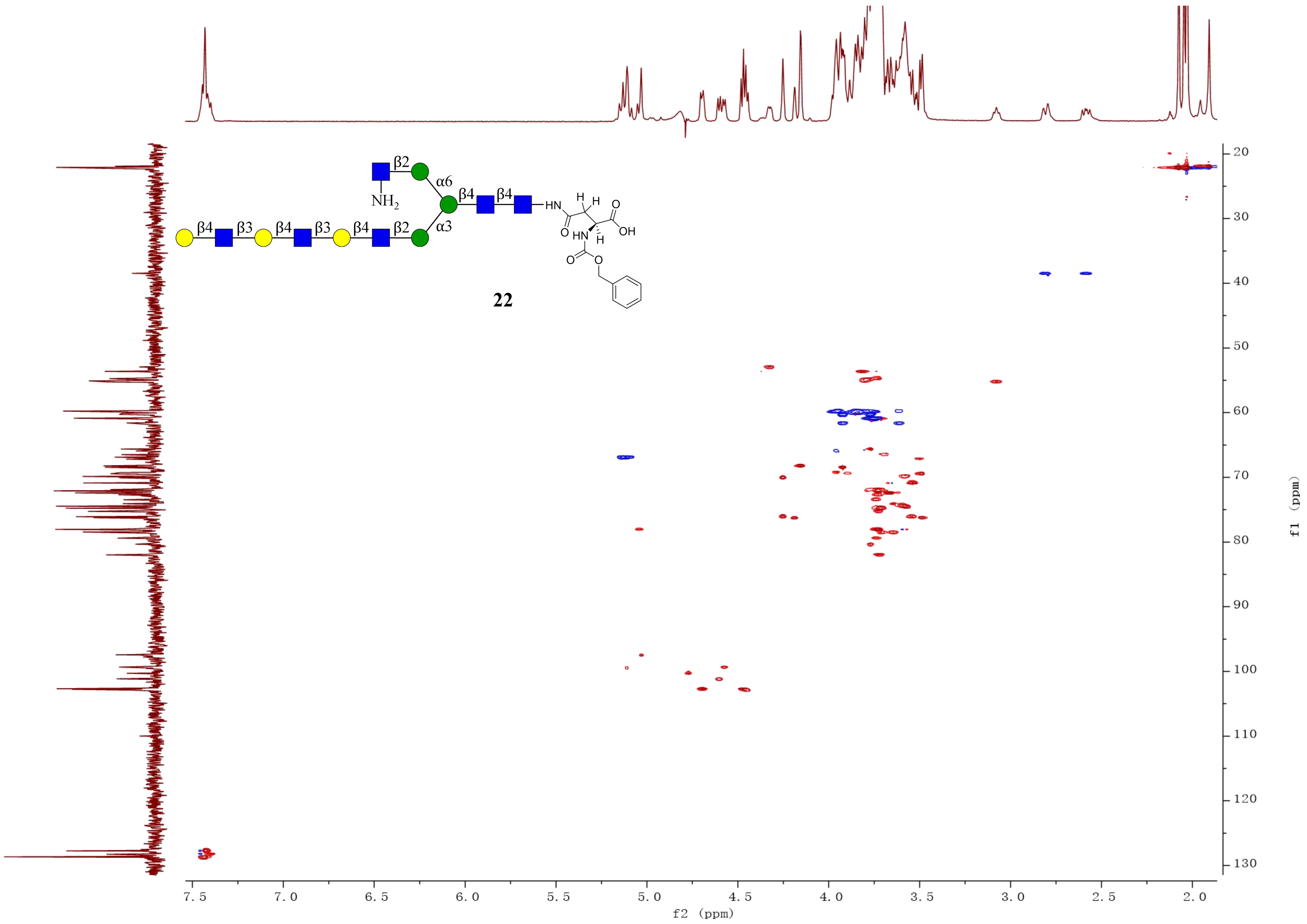

β2
α6
β4
β4
NH2
β4
β3
β4
β3
β4
β2
α3
HN
H
H
O
O
OH
HN
H
O
O
22
f2 (ppm)
f1 (ppm)

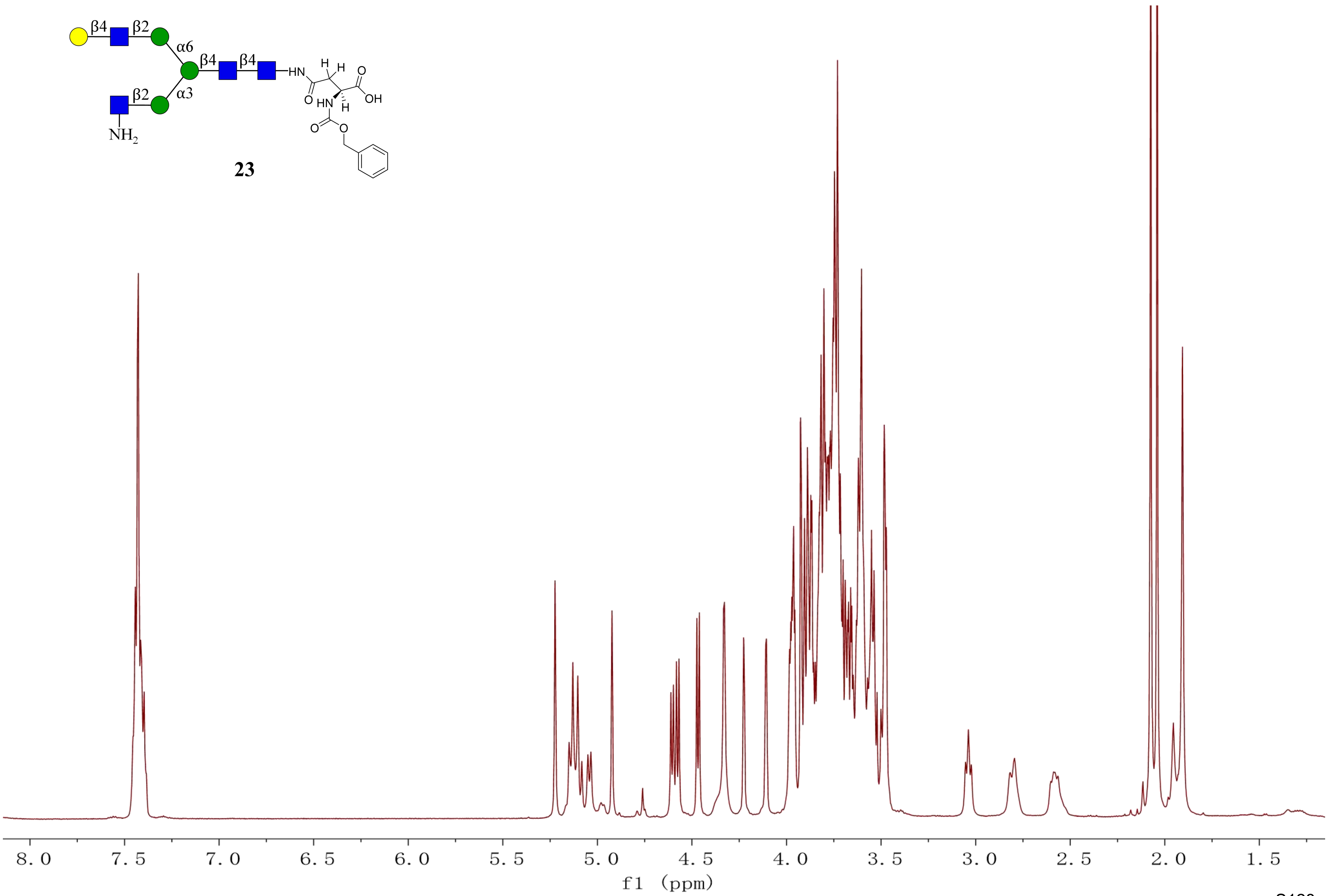
β4
β2
α6
β4
β4
α3
β2
HN
H
H
O
O
OH
HN
H
O
O
NH2
8.0
7.5
7.0
6.5
6.0
5.5
5.0
4.5
4.0
3.5
3.0
2.5
2.0
1.5
f1 (ppm)

**23**

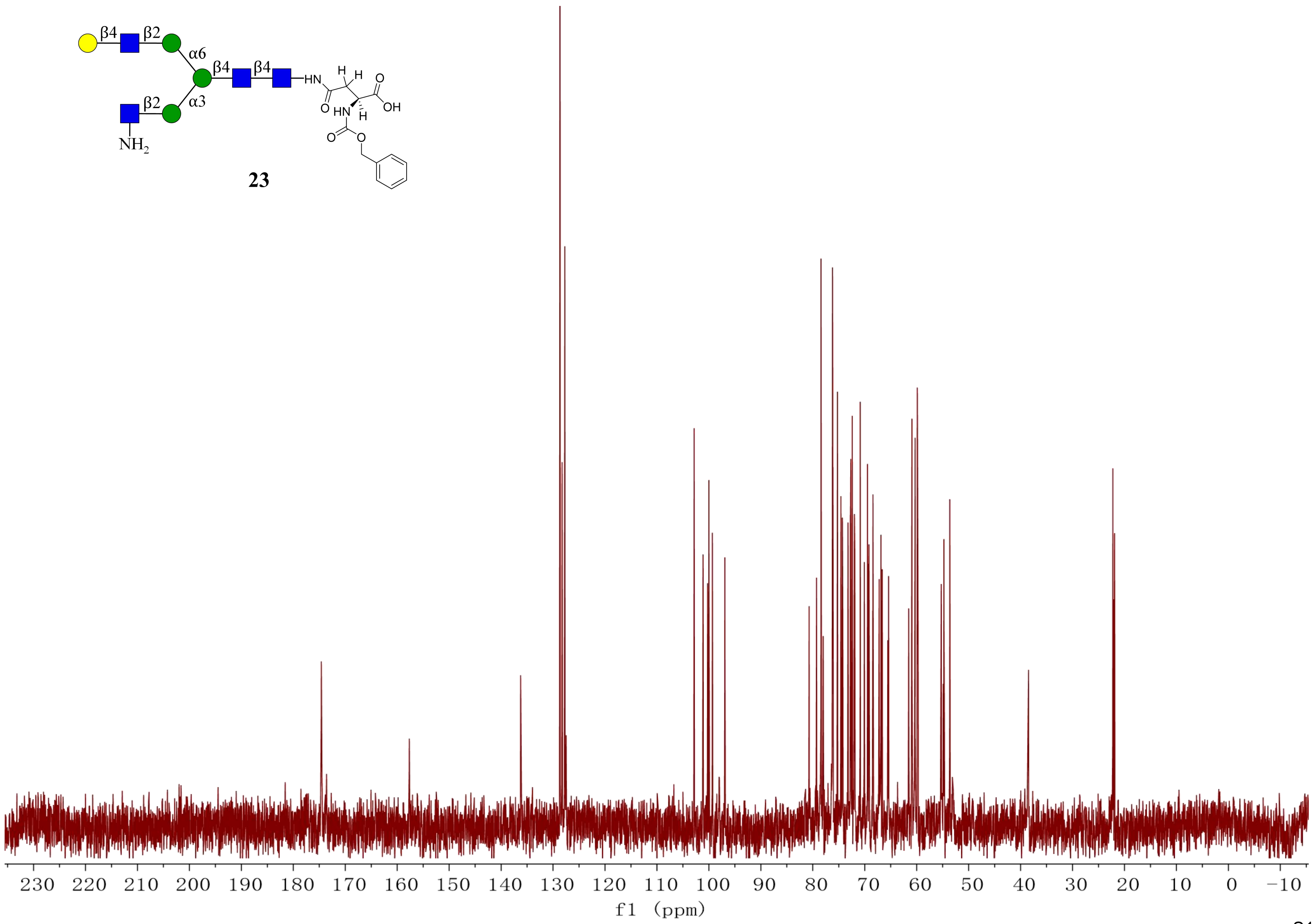

23
230 220 210 200 190 180 170 160 150 140 130 120 110 100 90 80 70 60 50 40 30 20 10 0 −10
f1 (ppm)

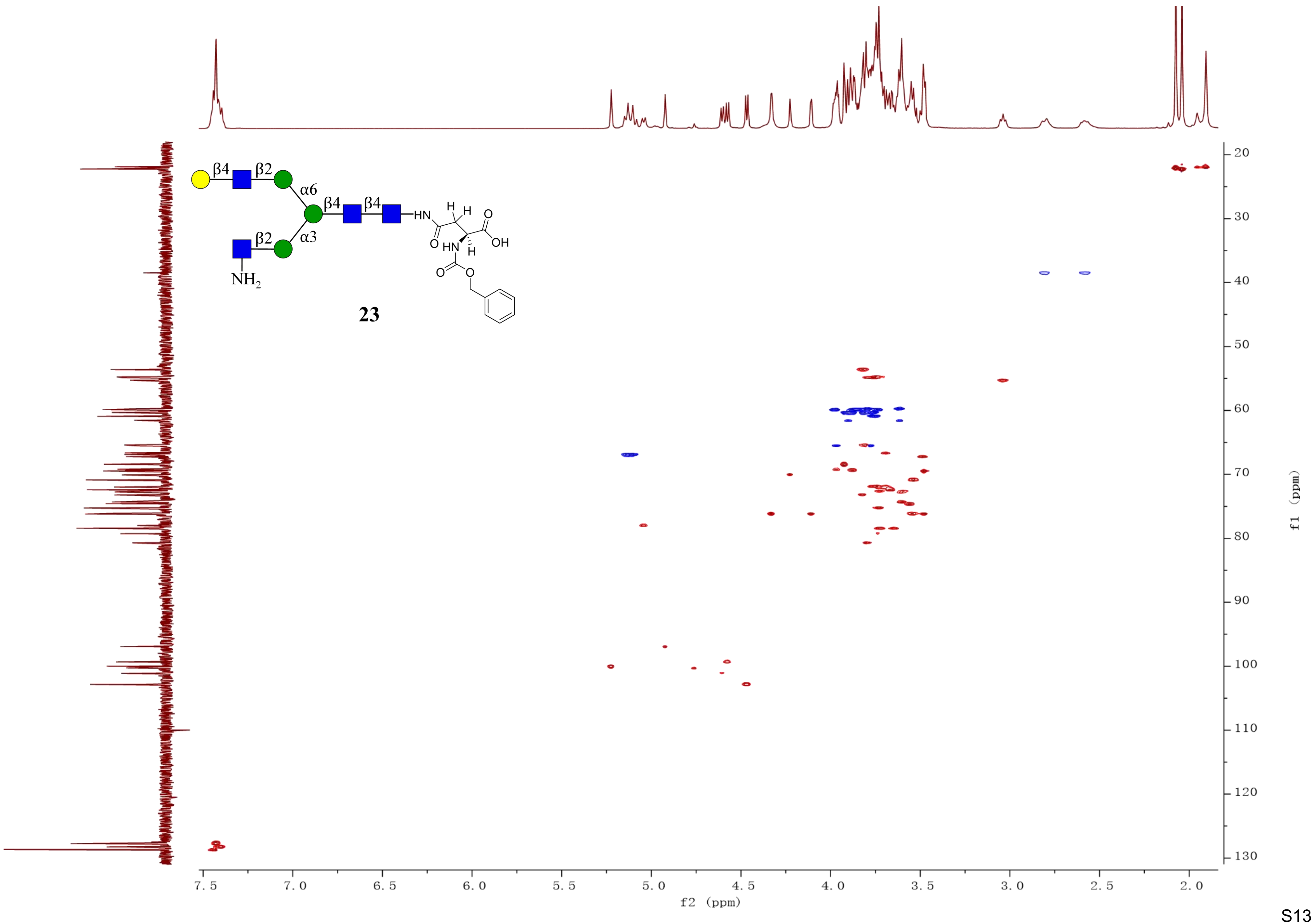
β4
β2
α6
β4
β4
α3
β2
NH2
HN
H
H
O
OH
O
HN
H
O
O
23
7.5
7.0
6.5
6.0
5.5
5.0
4.5
4.0
3.5
3.0
2.5
2.0
f2 (ppm)
20
30
40
50
60
70
80
90
100
110
120
130
f1 (ppm)

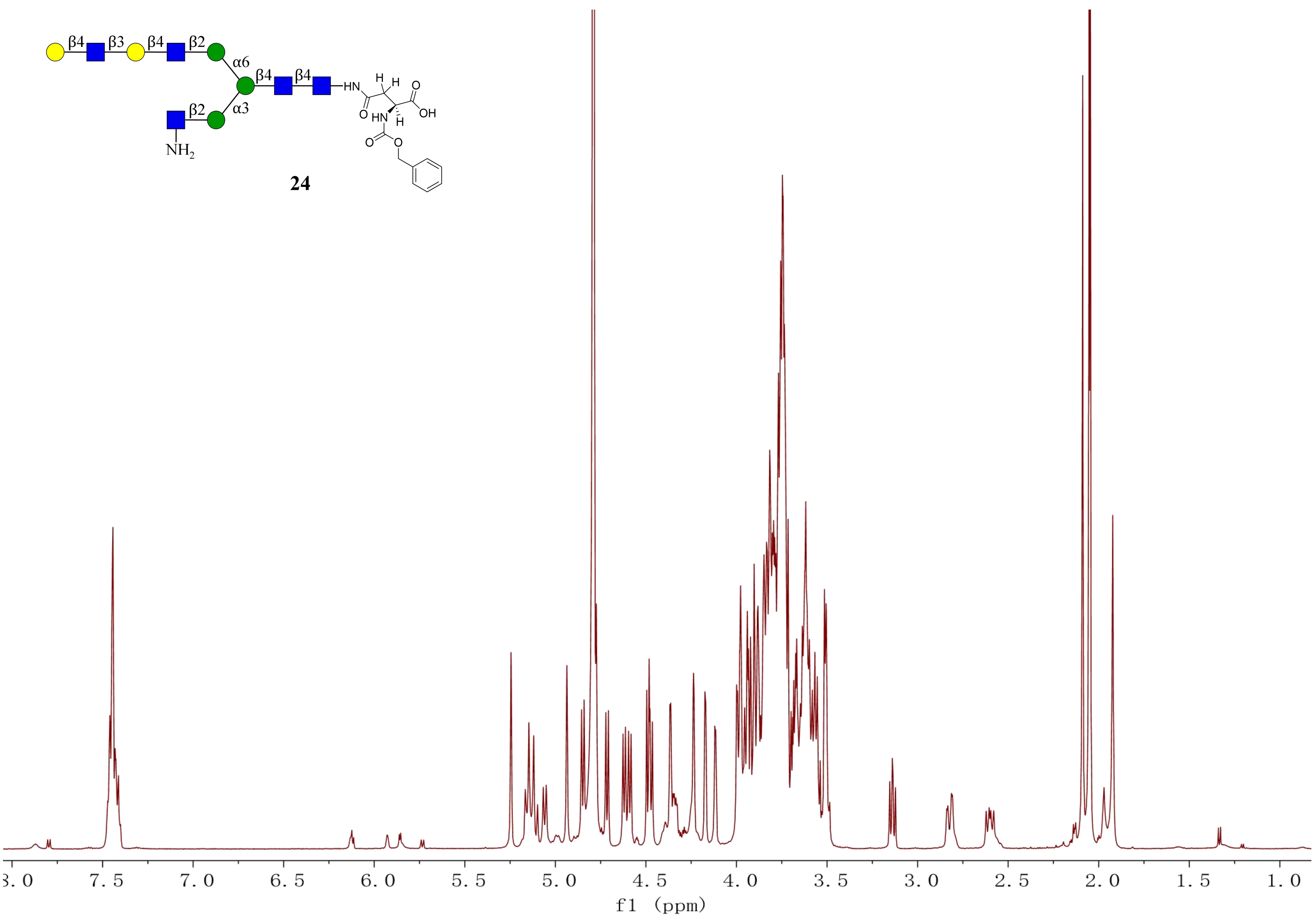


**24**

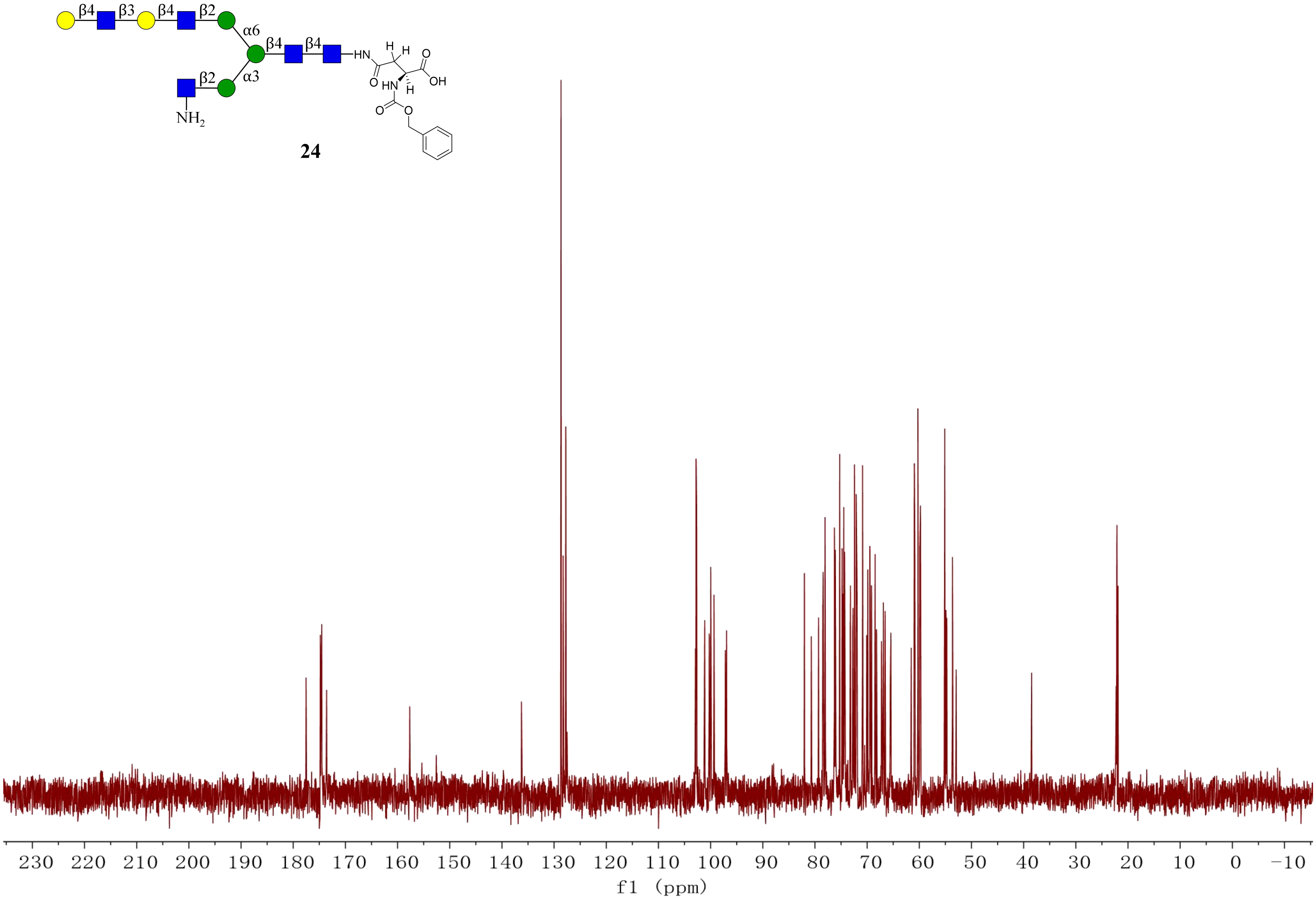
β4
β3
β4
β2
α6
β4
β4
α3
β2
NH2
HN
H
H
O
OH
O
HN
H
O
O
24
230 220 210 200 190 180 170 160 150 140 130 120 110 100 90 80 70 60 50 40 30 20 10 0 −10
f1 (ppm)

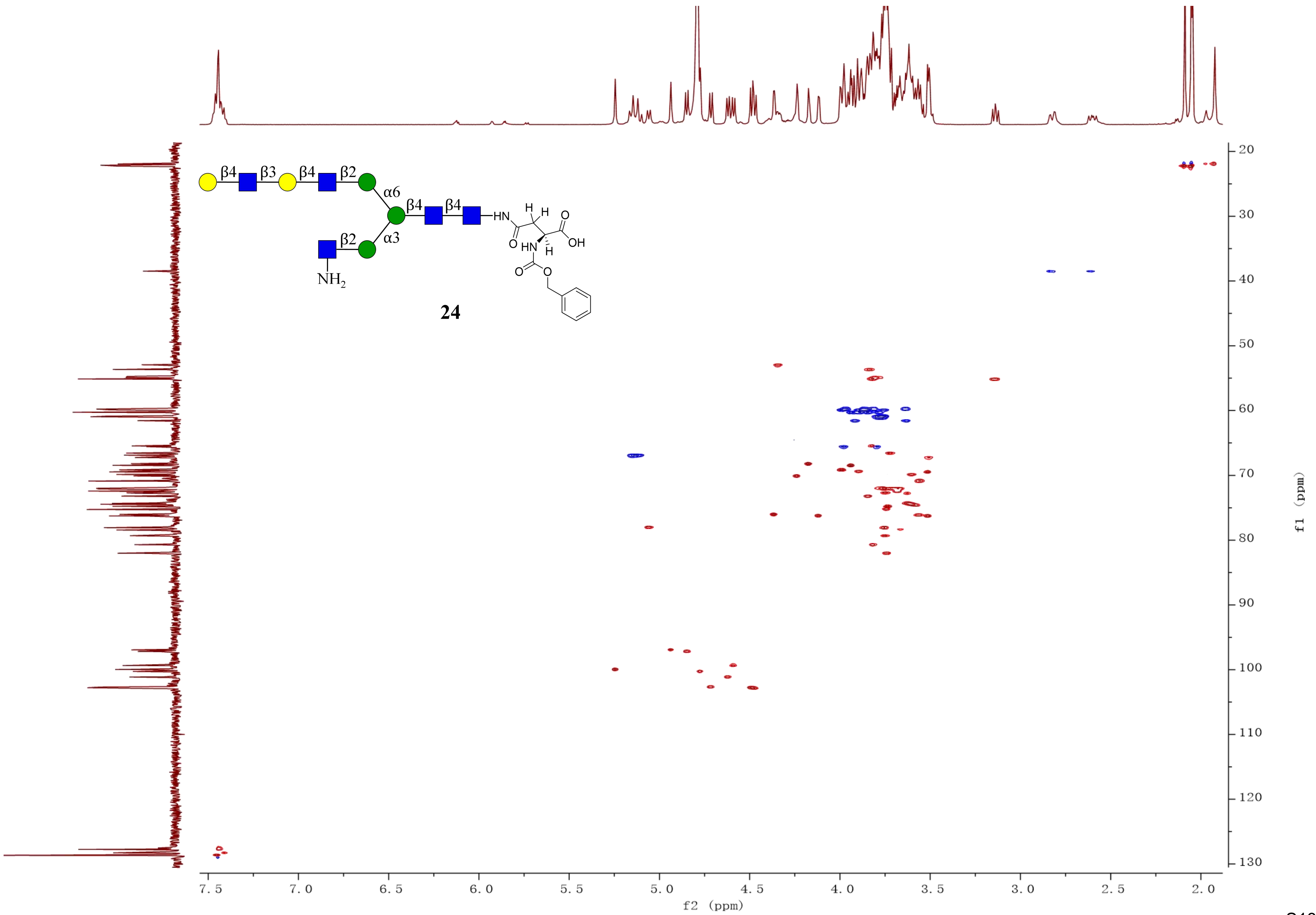
β4
β3
β4
β2
α6
β4
β4
β2
α3
HN
H
H
O
OH
O
HN
H
O
O
NH2
24
f2 (ppm)
f1 (ppm)
7.5
7.0
6.5
6.0
5.5
5.0
4.5
4.0
3.5
3.0
2.5
2.0
20
30
40
50
60
70
80
90
100
110
120
130

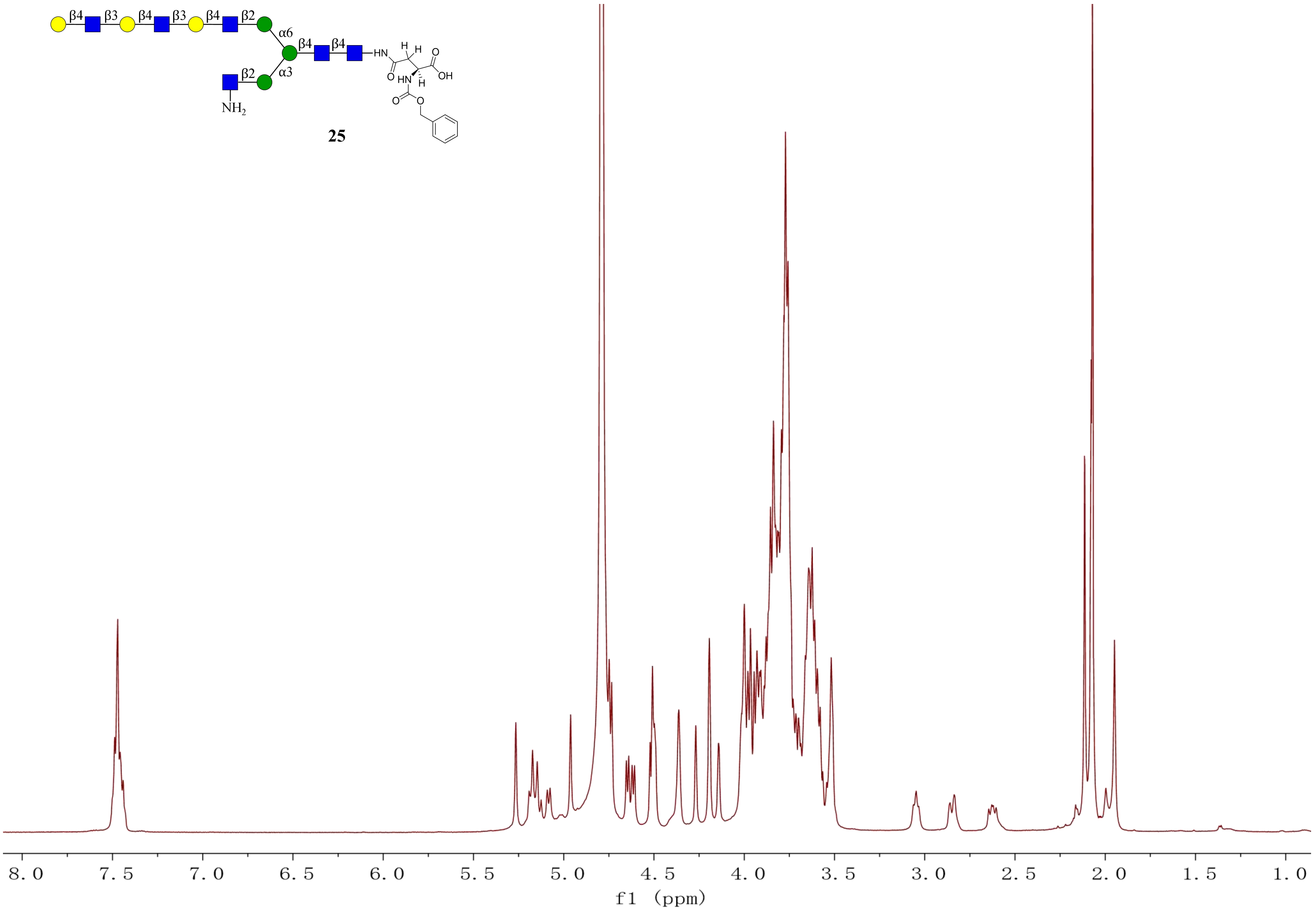
β4
β3
β4
β3
β4
β2
α6
β4
β4
β2
α3
NH2
HN
H
H
O
OH
O
HN
H
O
O
25
8.0
7.5
7.0
6.5
6.0
5.5
5.0
4.5
4.0
3.5
3.0
2.5
2.0
1.5
1.0
f1 (ppm)

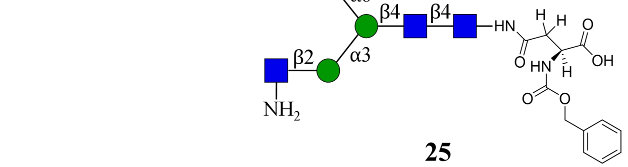
β4
β3
β4
β3
β4
β2
α6
β4
β4
HN
H
H
O
OH
O
HN
H
O
O
β2
α3
NH2

**25**

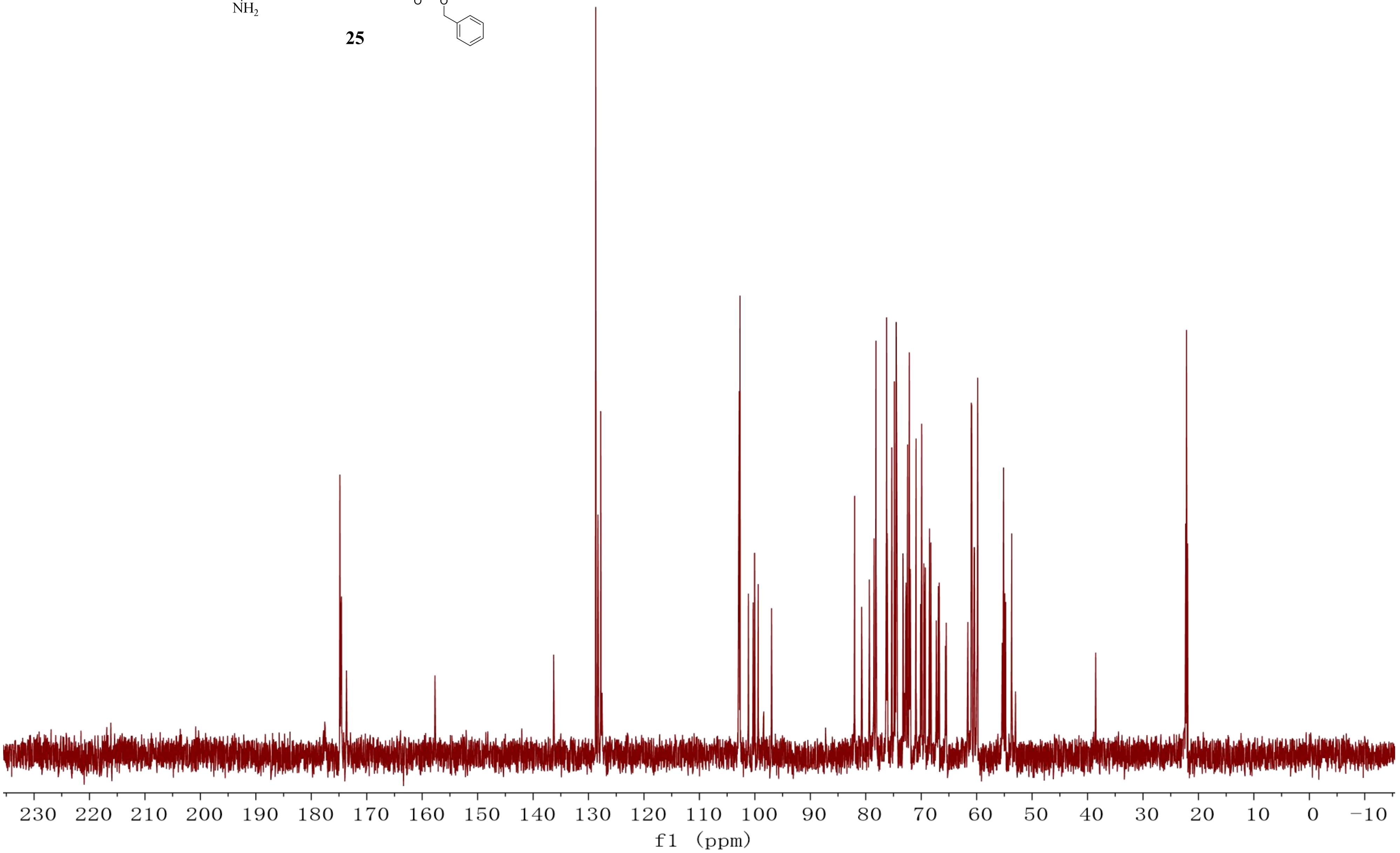
230
220
210
200
190
180
170
160
150
140
130
120
110
100
90
80
70
60
50
40
30
20
10
0
−10
f1 (ppm)

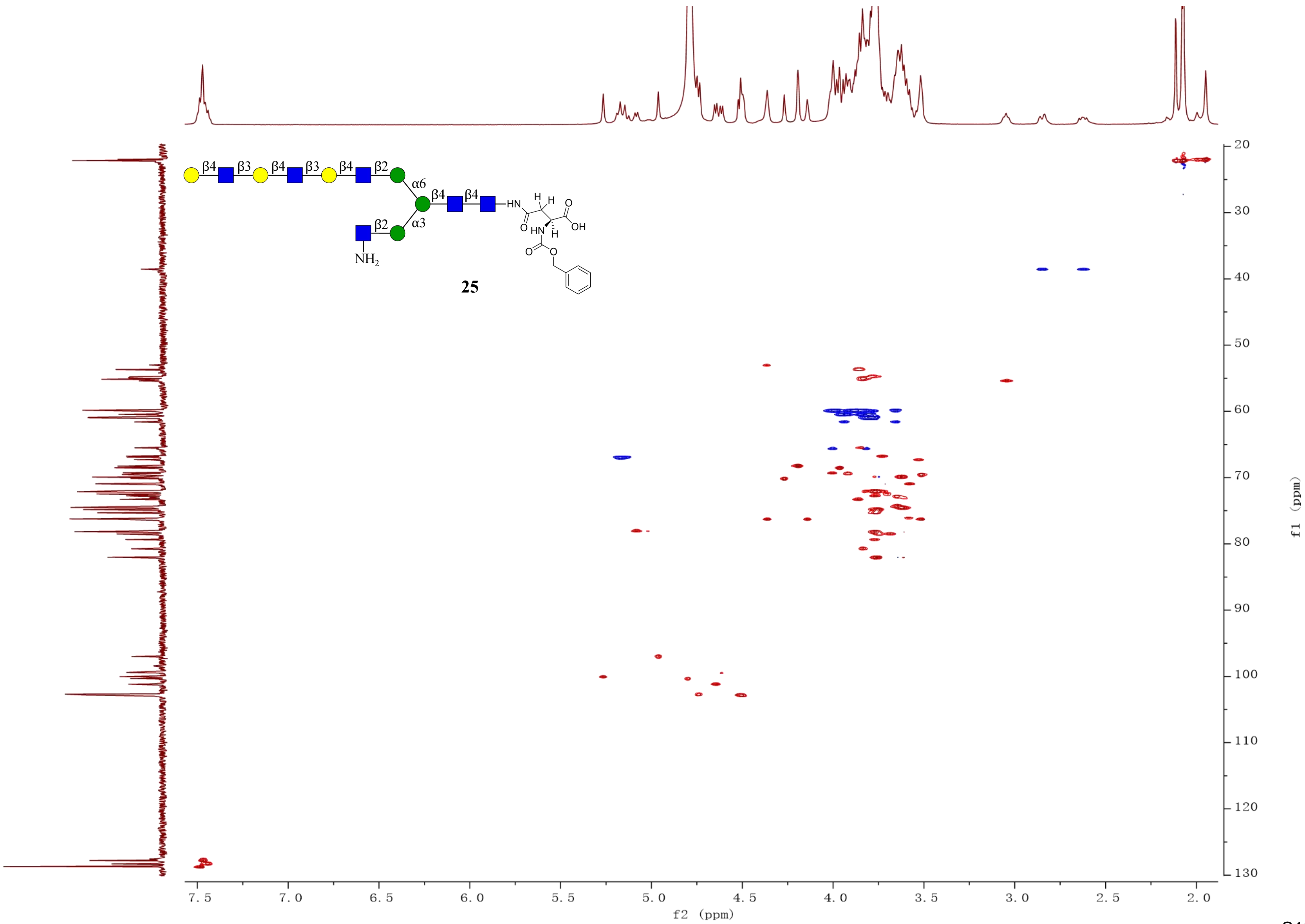
β4
β3
β4
β3
β4
β2
α6
β4
β4
α3
β2
NH2
HN
H
H
O
OH
O
HN
H
O
O
25
f2 (ppm)
f1 (ppm)

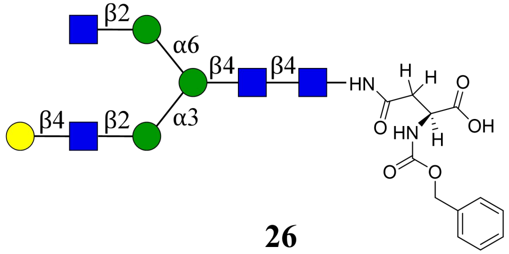

β2
α6
β4
β4
HN
H
H
O
β4
β2
α3
O
HN
H
OH
O
O


**26**

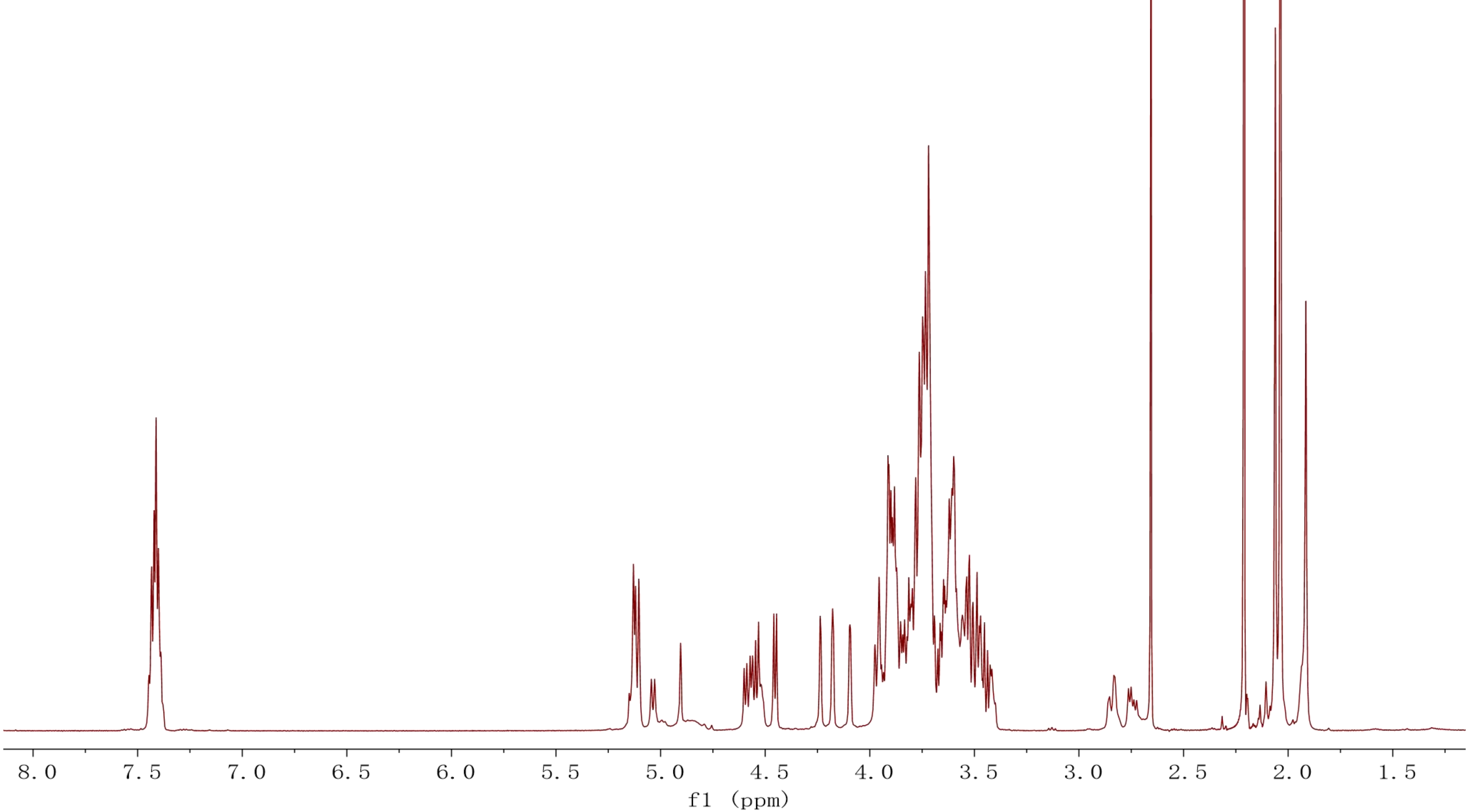

8.0
7.5
7.0
6.5
6.0
5.5
5.0
4.5
4.0
3.5
3.0
2.5
2.0
1.5
f1 (ppm)

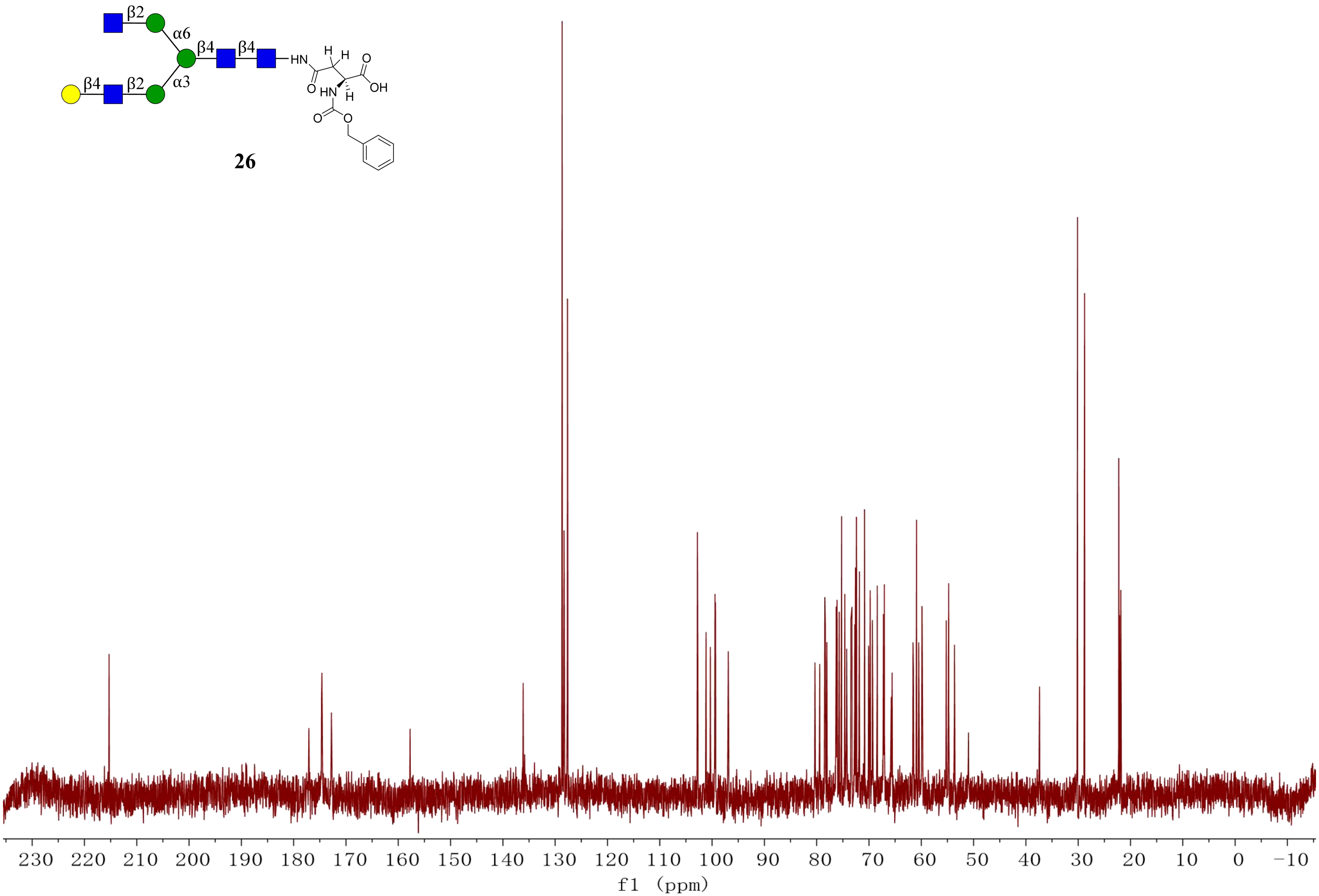
β2
α6
β4
β4
HN
H
H
O
OH
O
HN
H
O
O
β4
β2
α3
230 220 210 200 190 180 170 160 150 140 130 120 110 100 90 80 70 60 50 40 30 20 10 0 −10
f1 (ppm)

**26**

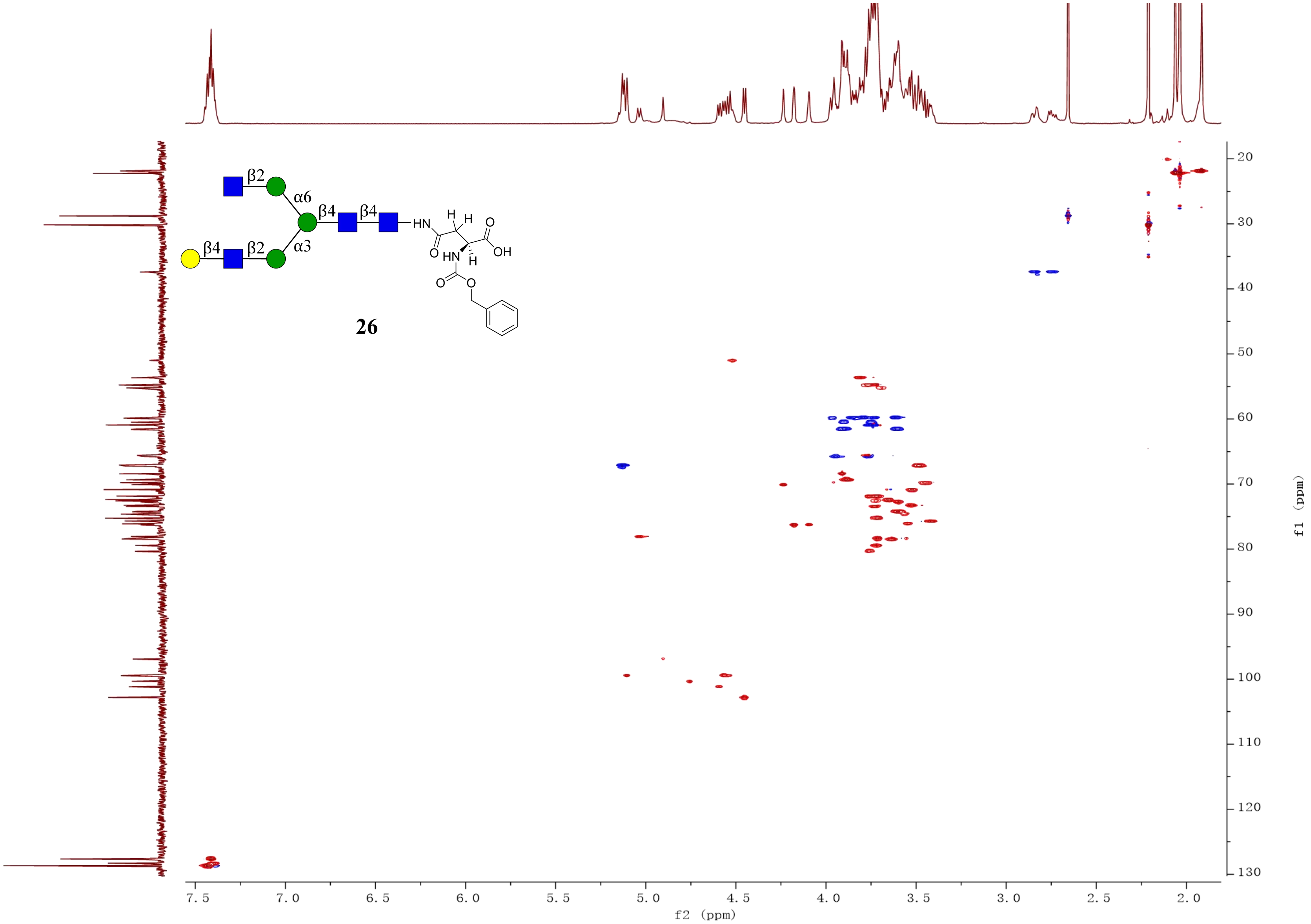
β2
α6
β4
β4
β4
β2
α3
HN
H
H
O
OH
O
HN
H
O
O
26
f2 (ppm)
f1 (ppm)
7.5
7.0
6.5
6.0
5.5
5.0
4.5
4.0
3.5
3.0
2.5
2.0
20
30
40
50
60
70
80
90
100
110
120
130

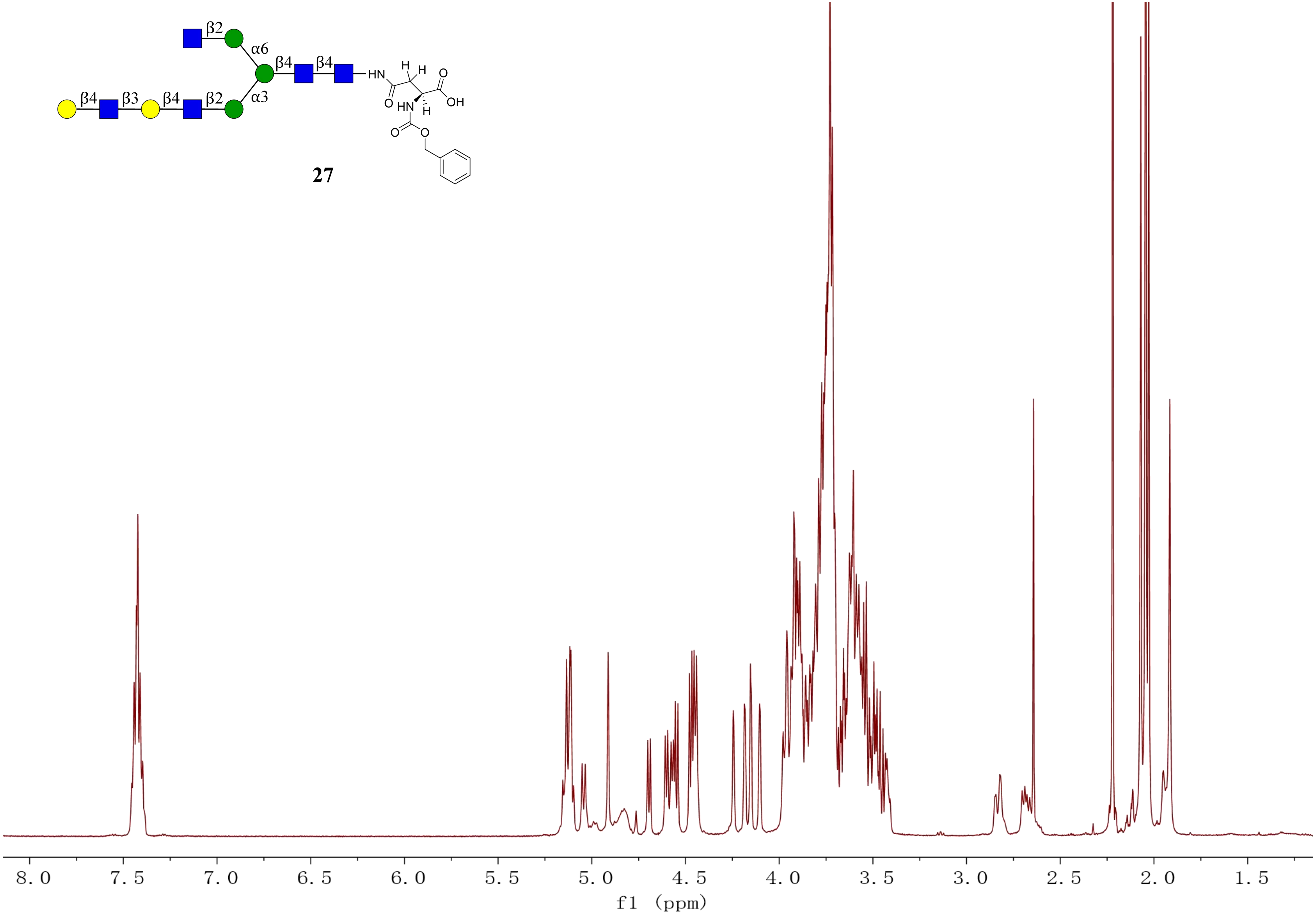
β2
α6
β4
β4
HN
H
H
O
OH
O
HN
H
O
O
β4
β3
β4
β2
α3
27
8.0
7.5
7.0
6.5
6.0
5.5
5.0
4.5
4.0
3.5
3.0
2.5
2.0
1.5
f1 (ppm)

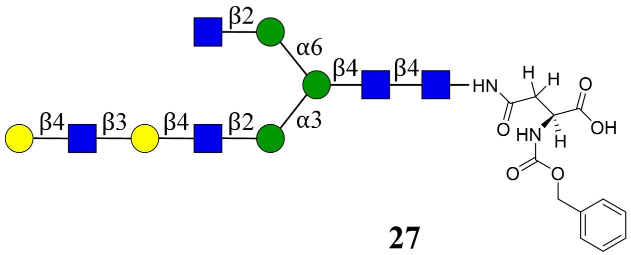
β2
α6
β4
β4
HN
H
H
O
OH
O
HN
H
O
O
β4
β3
β4
β2
α3

**27**

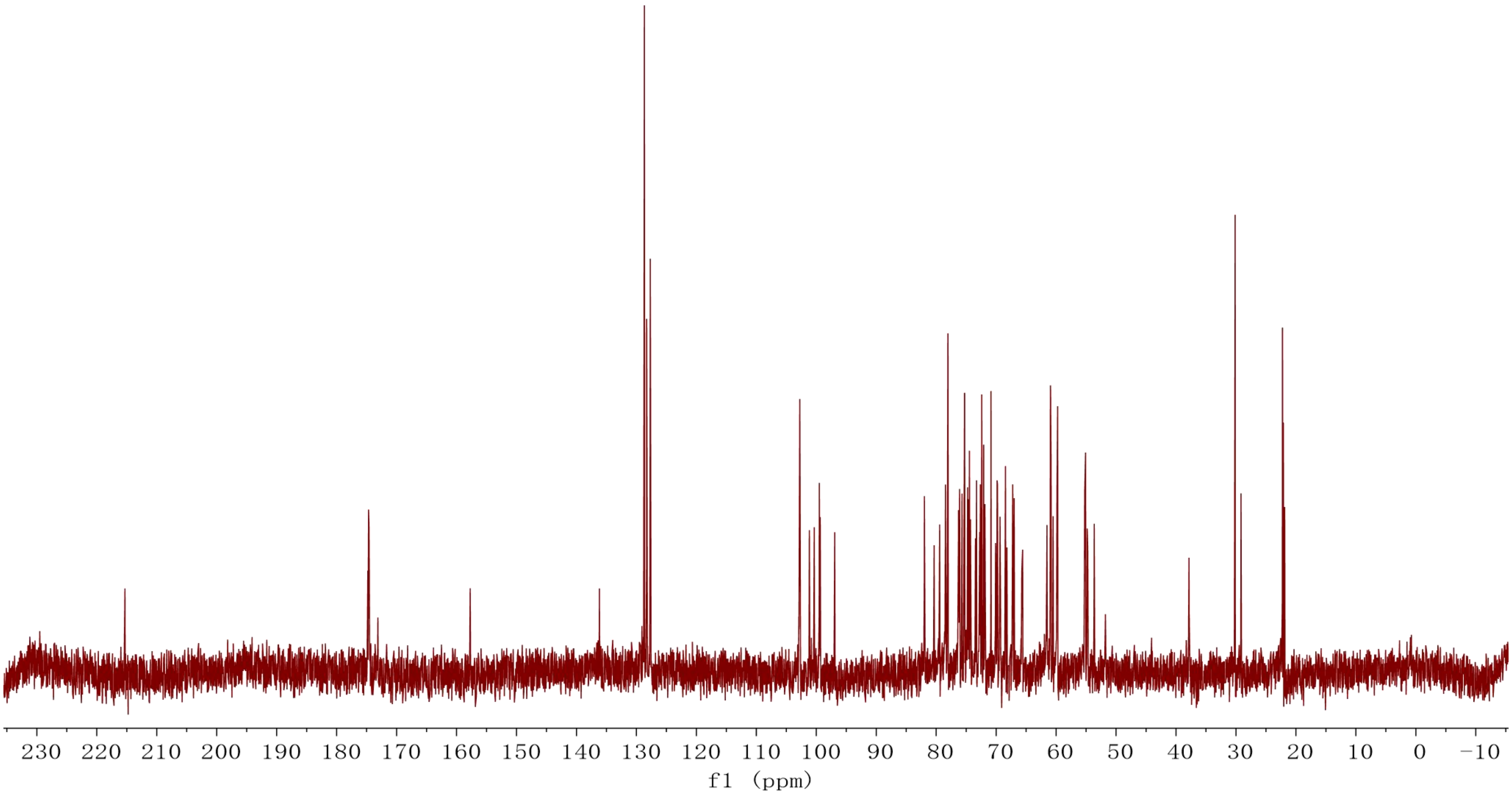
230 220 210 200 190 180 170 160 150 140 130 120 110 100 90 80 70 60 50 40 30 20 10 0 −10
f1 (ppm)

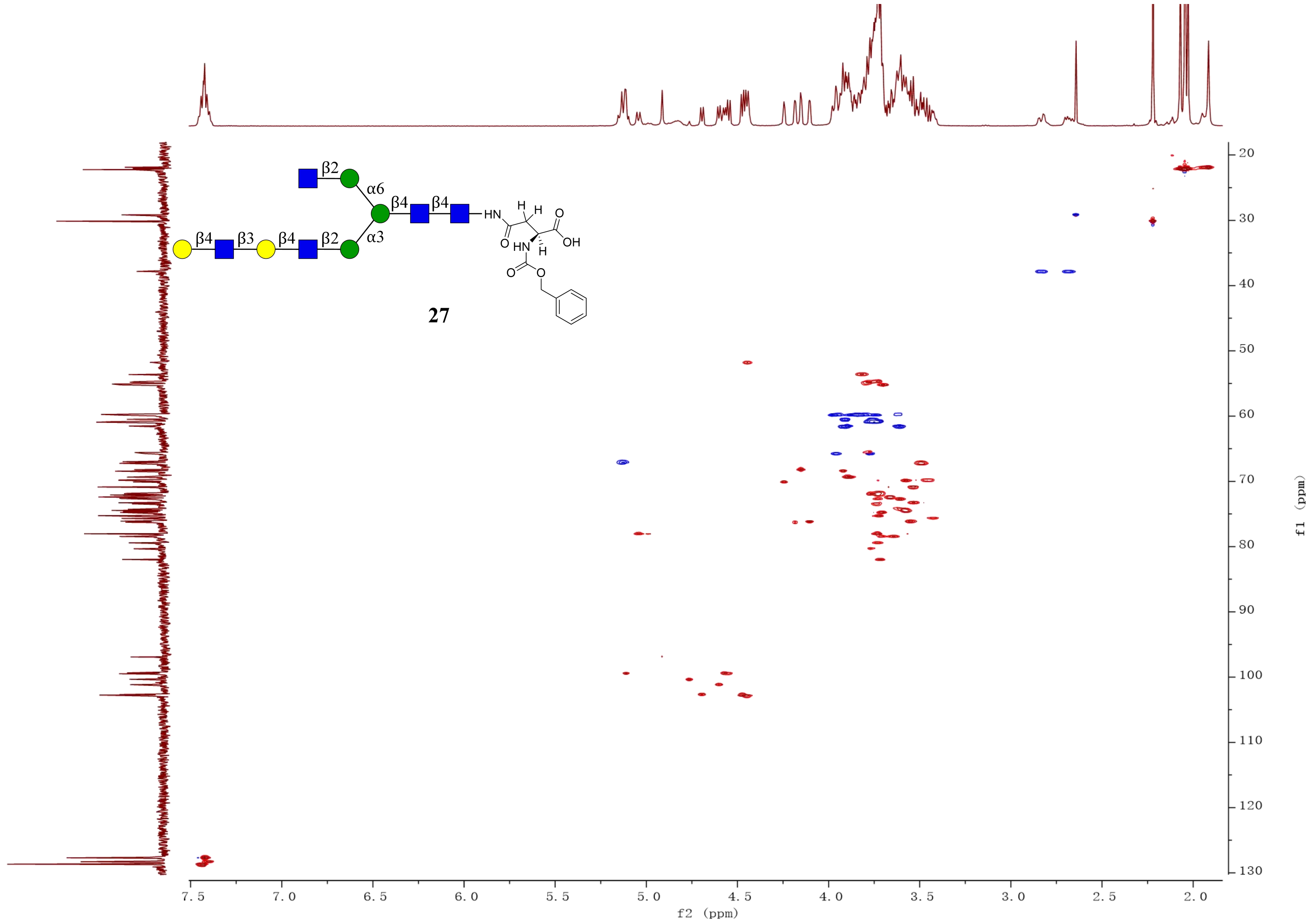
β2
α6
β4
β4
HN
β4
β3
β4
β2
α3
27
f2 (ppm)
f1 (ppm)

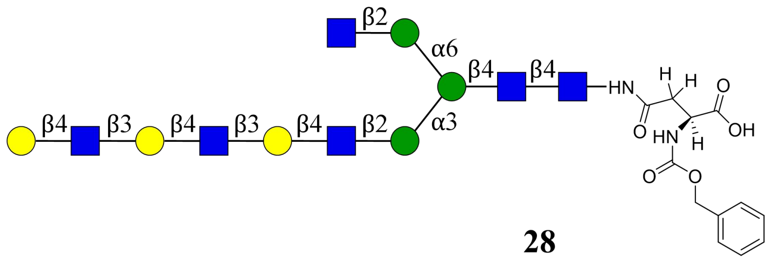

**28**

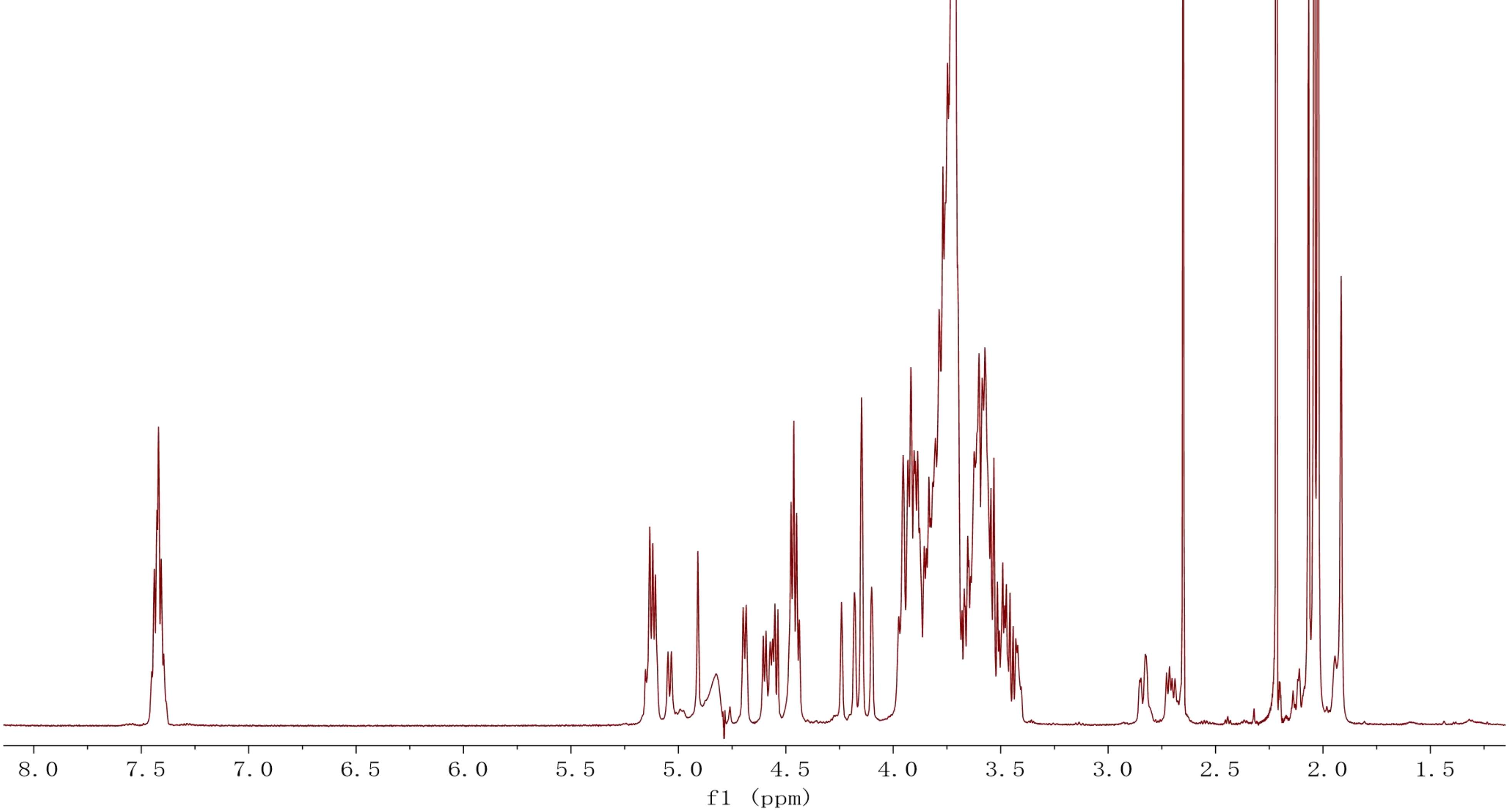

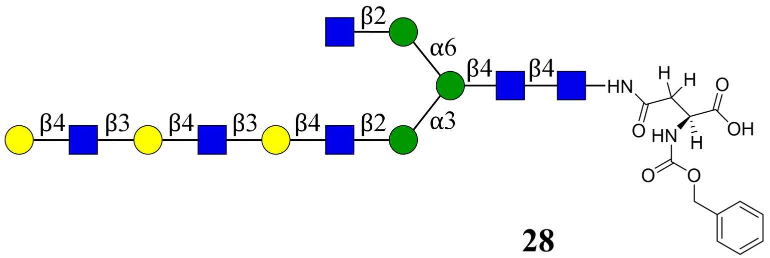

β2
α6
β4
β4
HN
H
H
O
O
OH
HN
H
O
O
β4
β3
β4
β3
β4
β2
α3


**28**

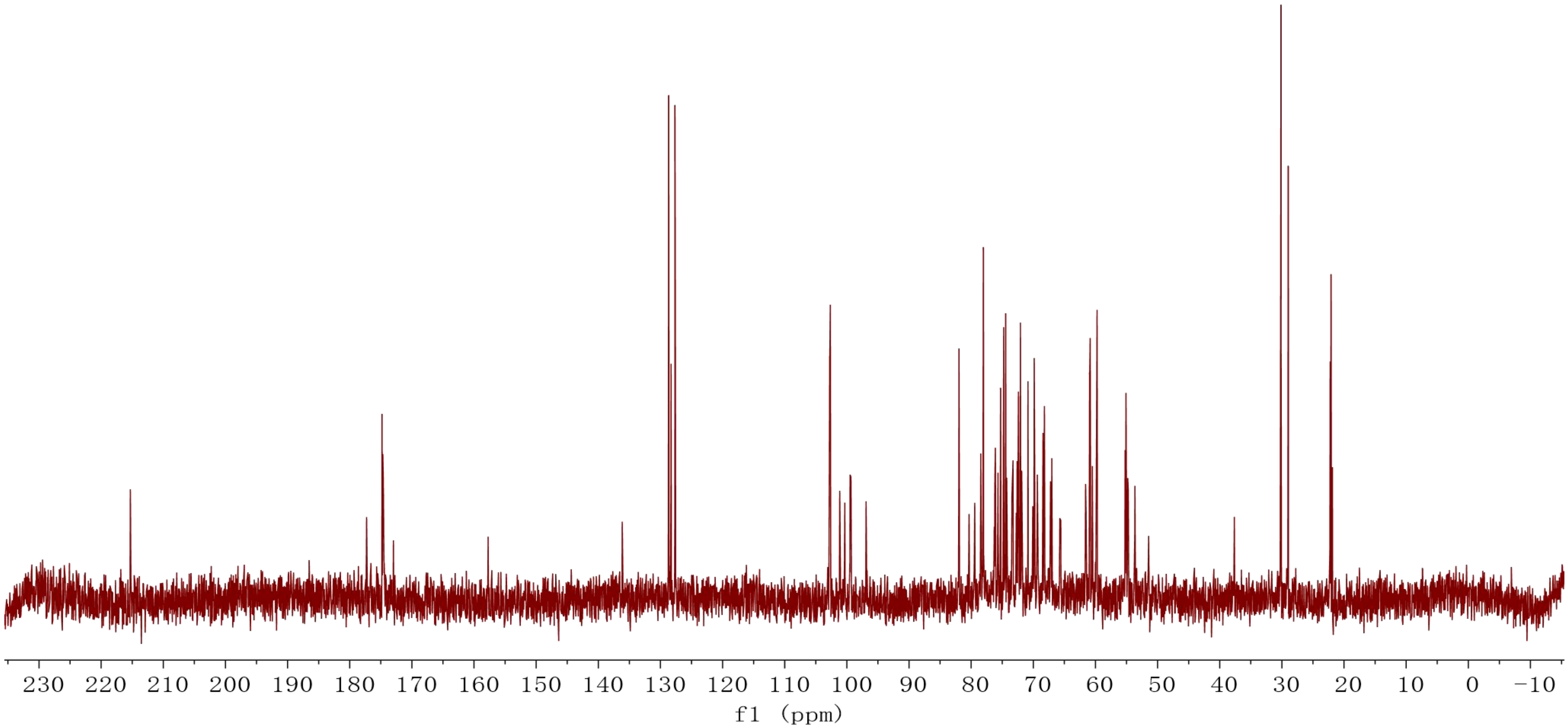

230 220 210 200 190 180 170 160 150 140 130 120 110 100 90 80 70 60 50 40 30 20 10 0 −10
f1 (ppm)

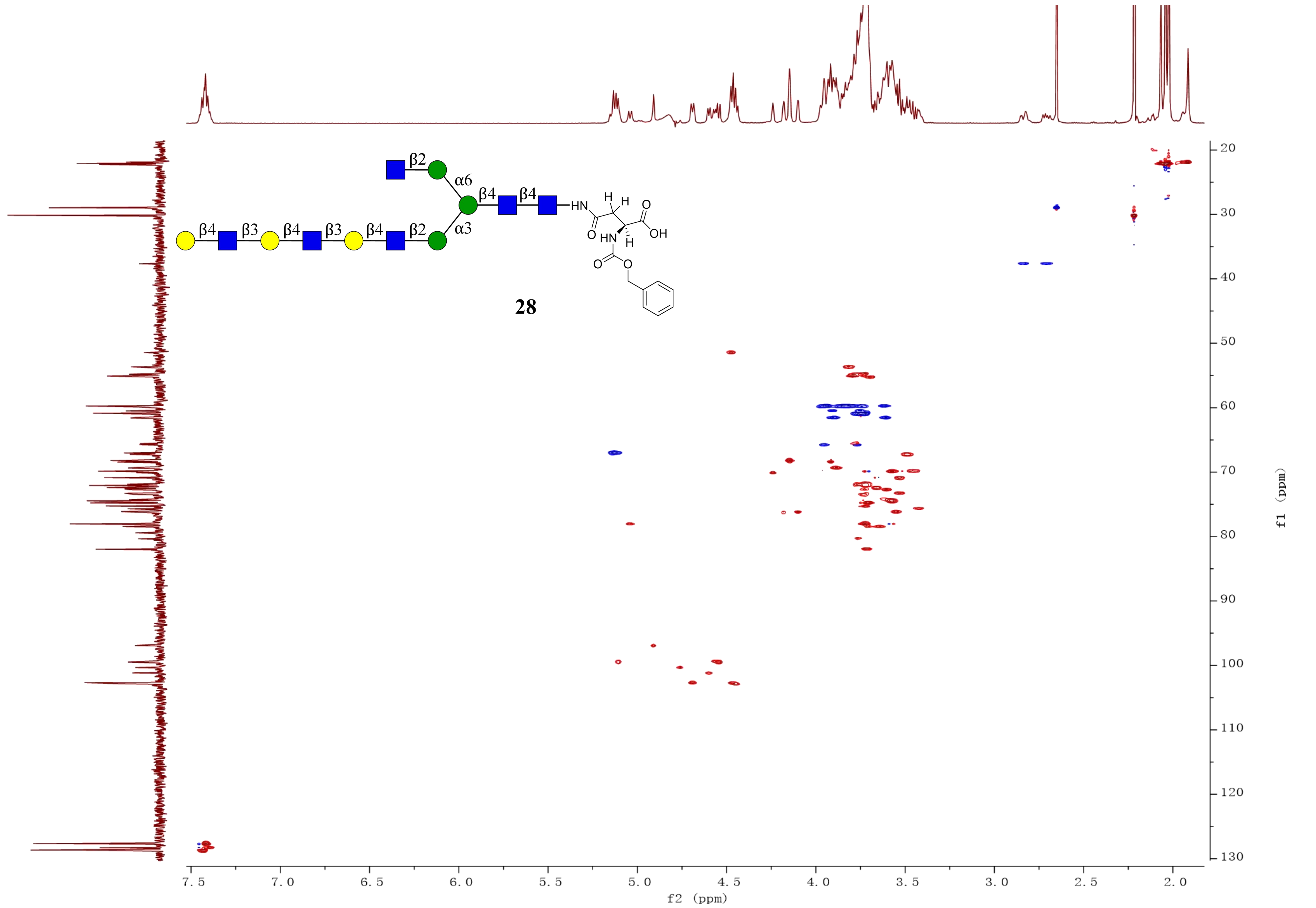
β2
α6
β4
β4
HN
H
H
O
OH
O
HN
H
O
O
β4
β3
β4
β3
β4
β2
α3
28
f2 (ppm)
f1 (ppm)

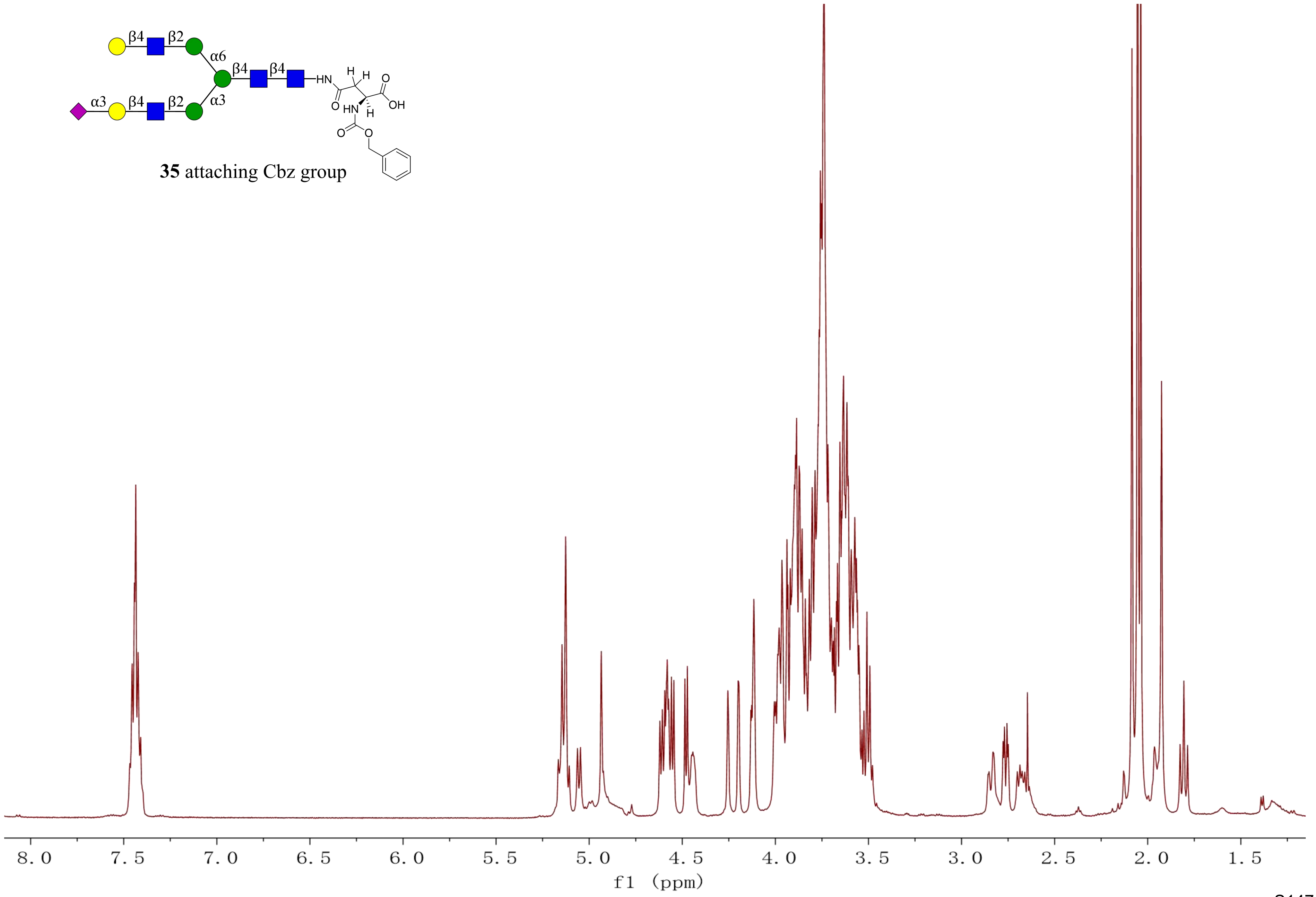

35 attaching Cbz group
8.0
7.5
7.0
6.5
6.0
5.5
5.0
4.5
4.0
3.5
3.0
2.5
2.0
1.5
f1 (ppm)

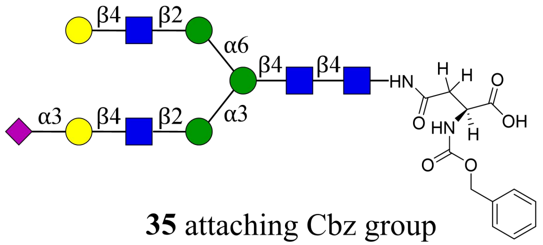


**35** attaching Cbz group

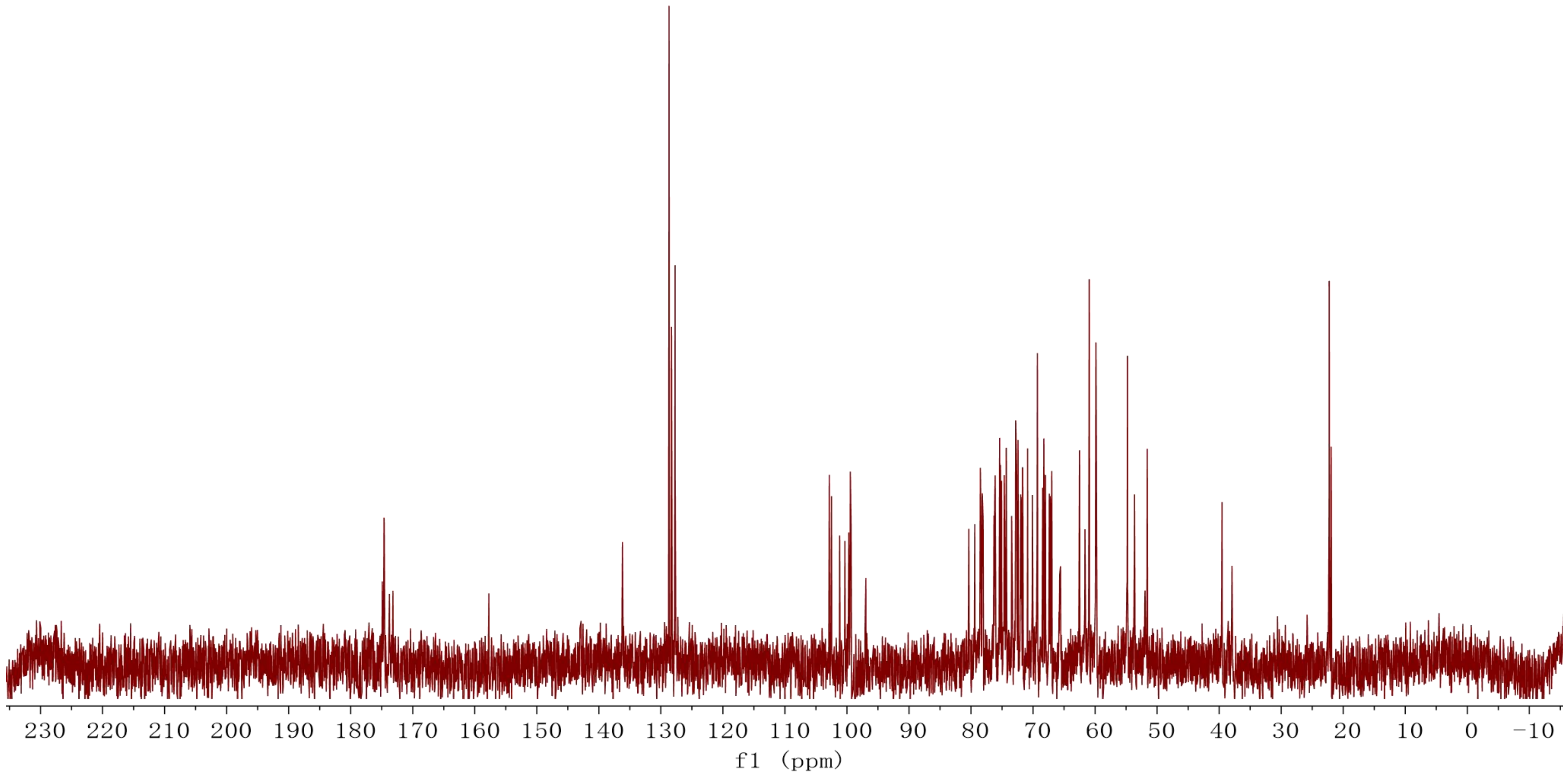

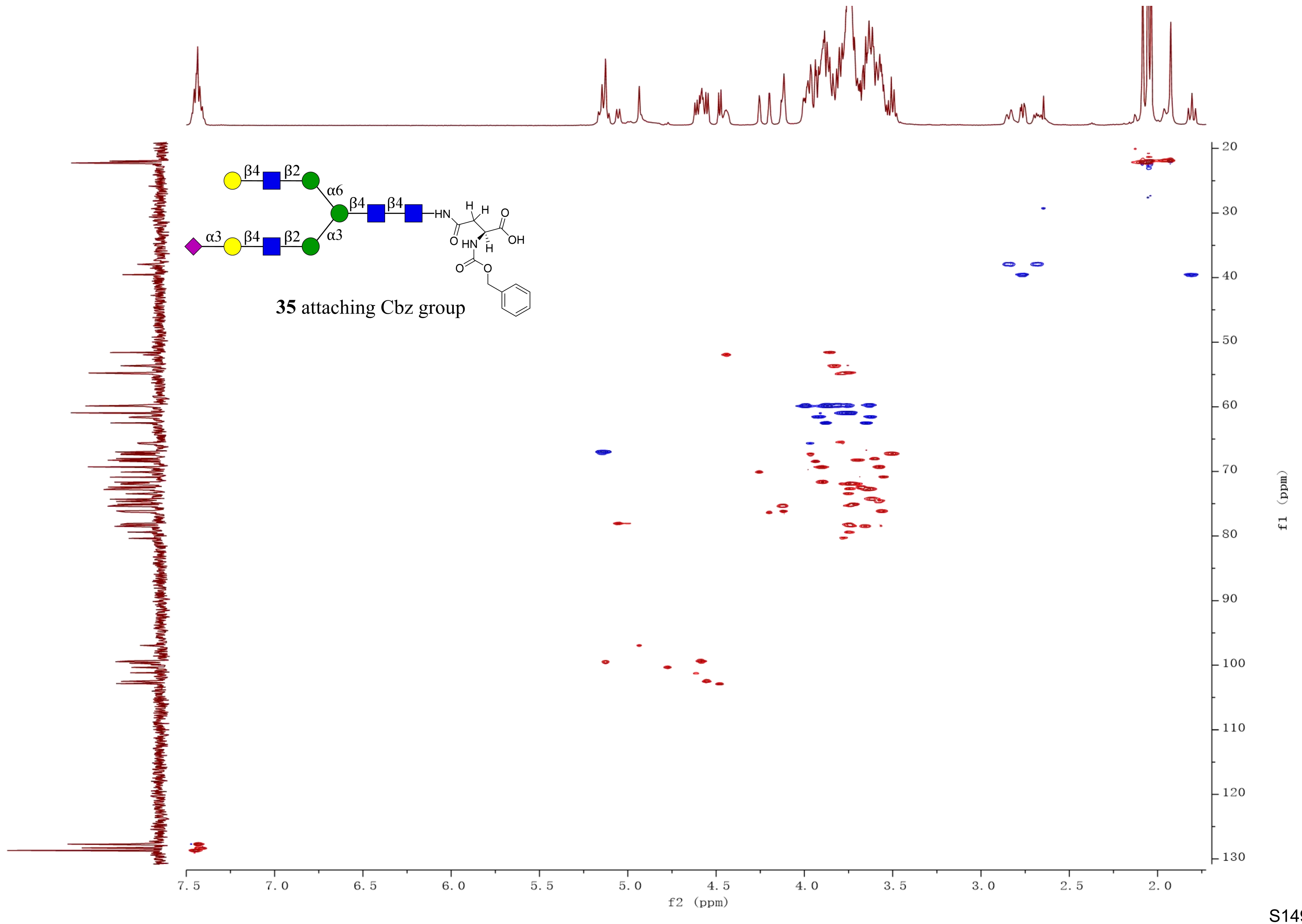

β4
β2
α6
β4
β4
α3
β4
β2
α3
HN
H
H
O
OH
O
HN
H
O
O
35 attaching Cbz group
20
30
40
50
60
70
80
90
100
110
120
130
f1 (ppm)
7.5
7.0
6.5
6.0
5.5
5.0
4.5
4.0
3.5
3.0
2.5
2.0
f2 (ppm)

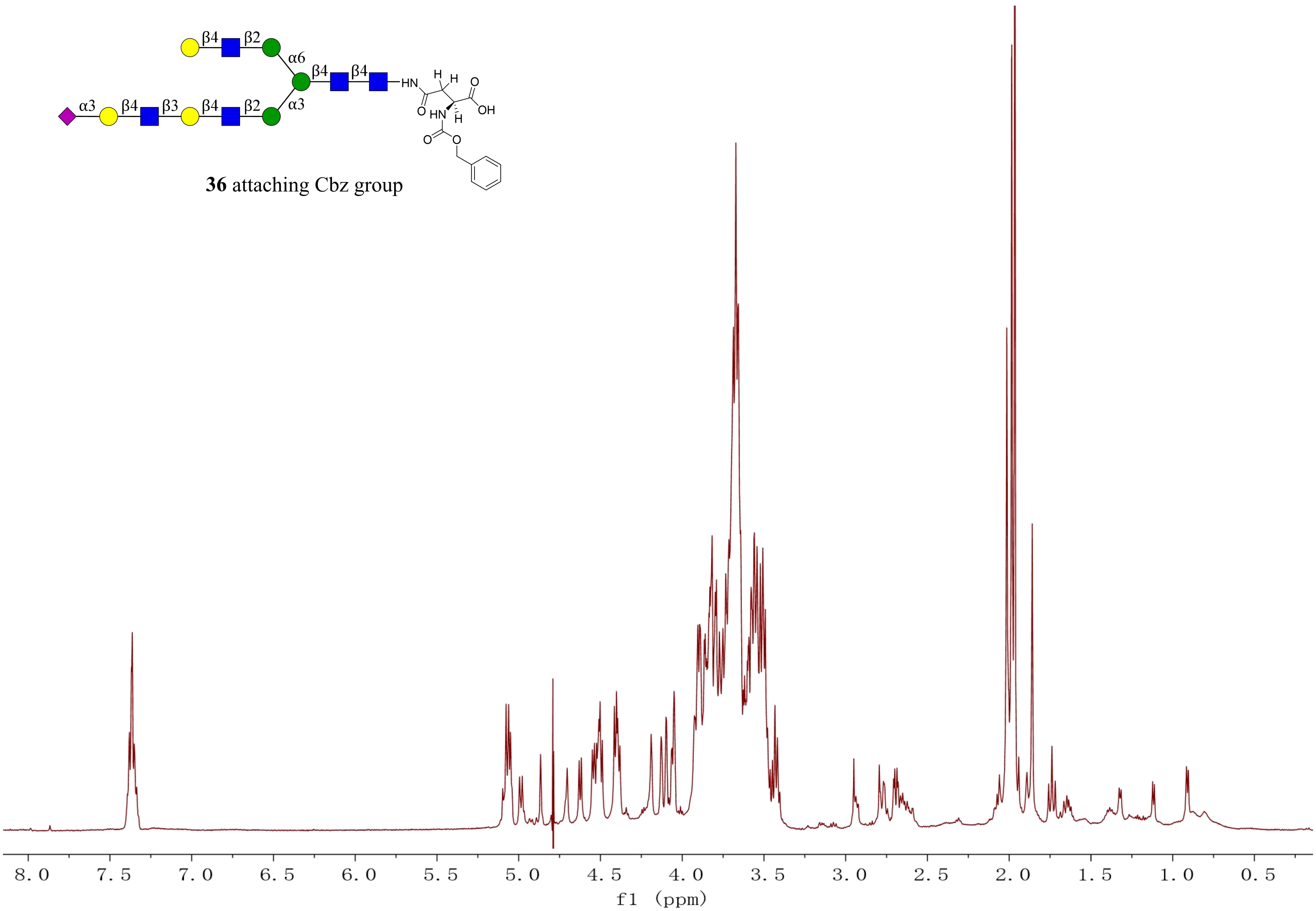


**36** attaching Cbz group

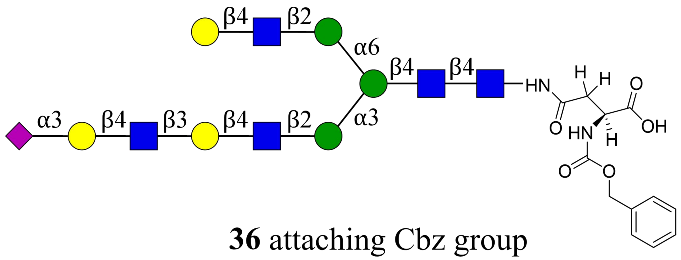

**36** attaching Cbz group

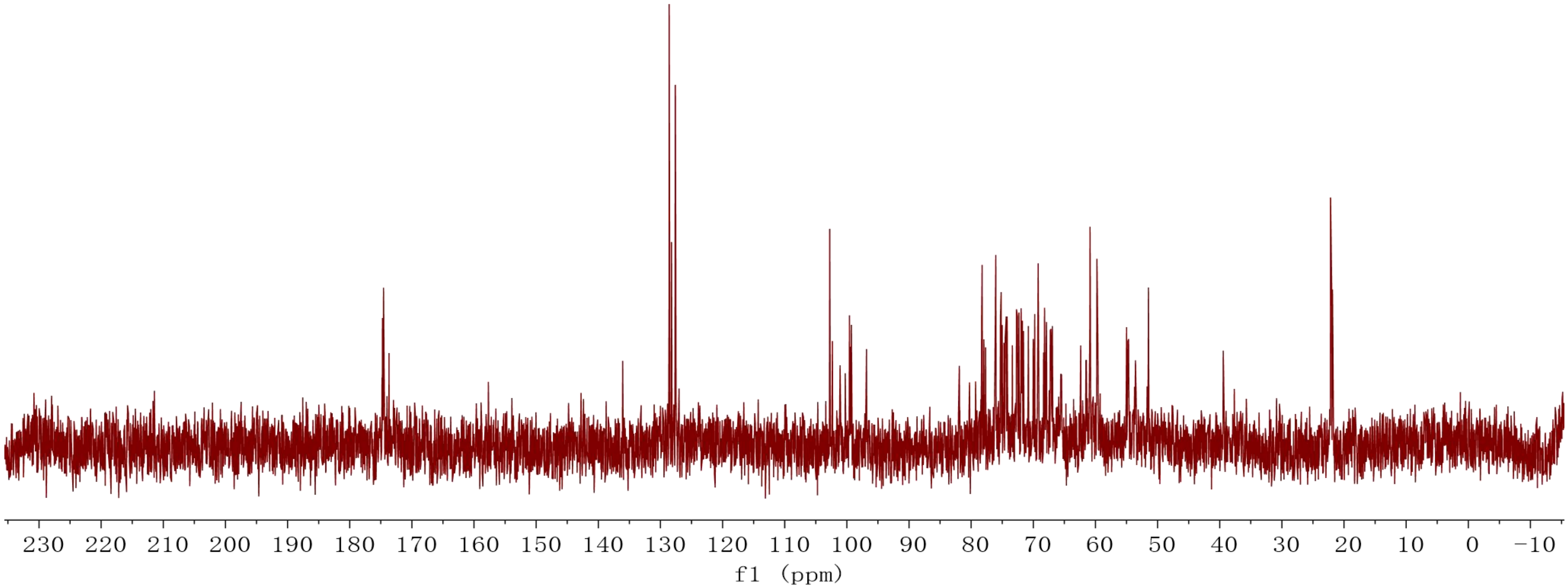

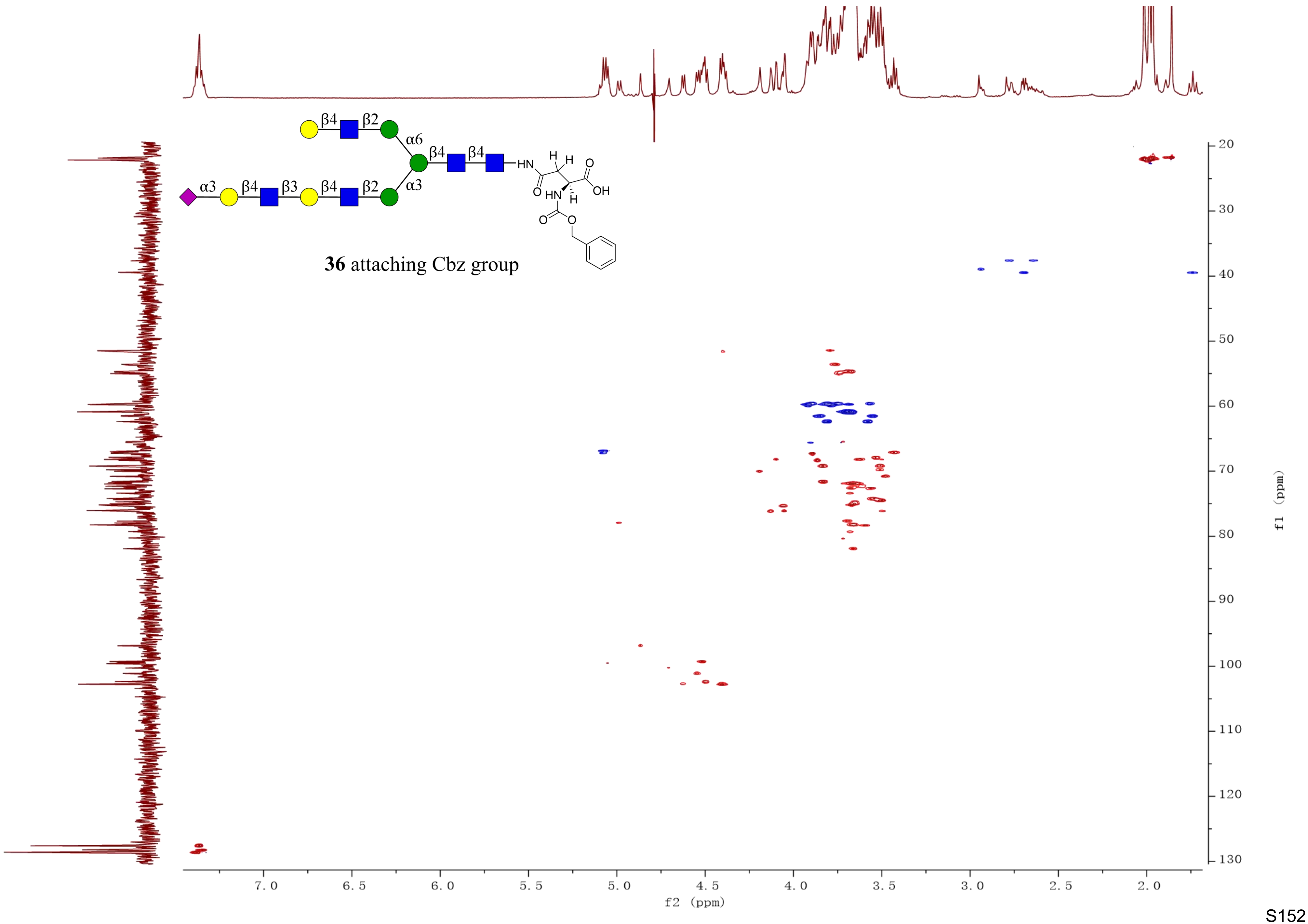

β4
β2
α6
β4
β4
α3
β4
β3
β4
β2
α3
HN
H
H
O
O
HN
H
OH
O
O
36 attaching Cbz group
f2 (ppm)
f1 (ppm)

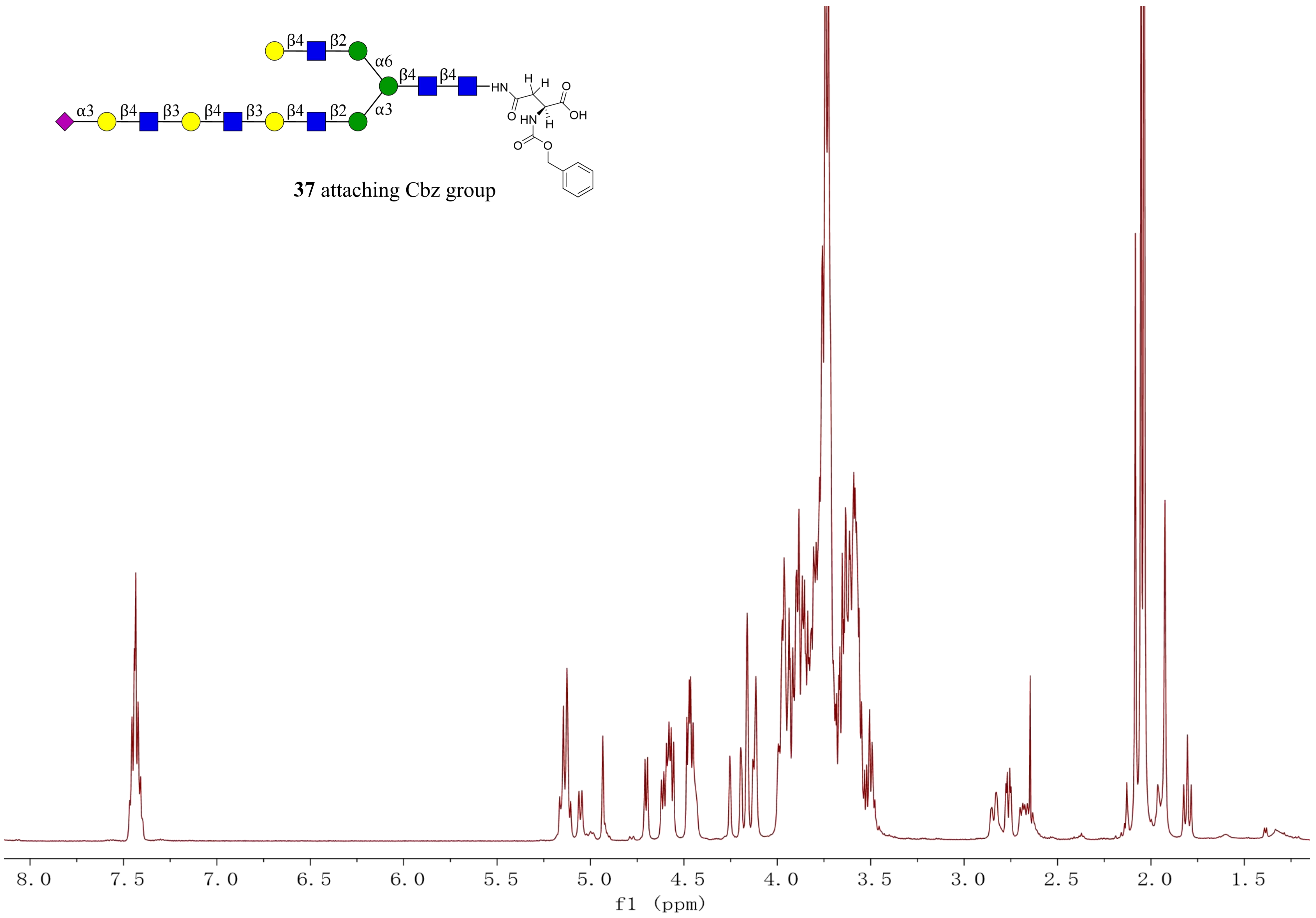
37 attaching Cbz group
f1 (ppm)

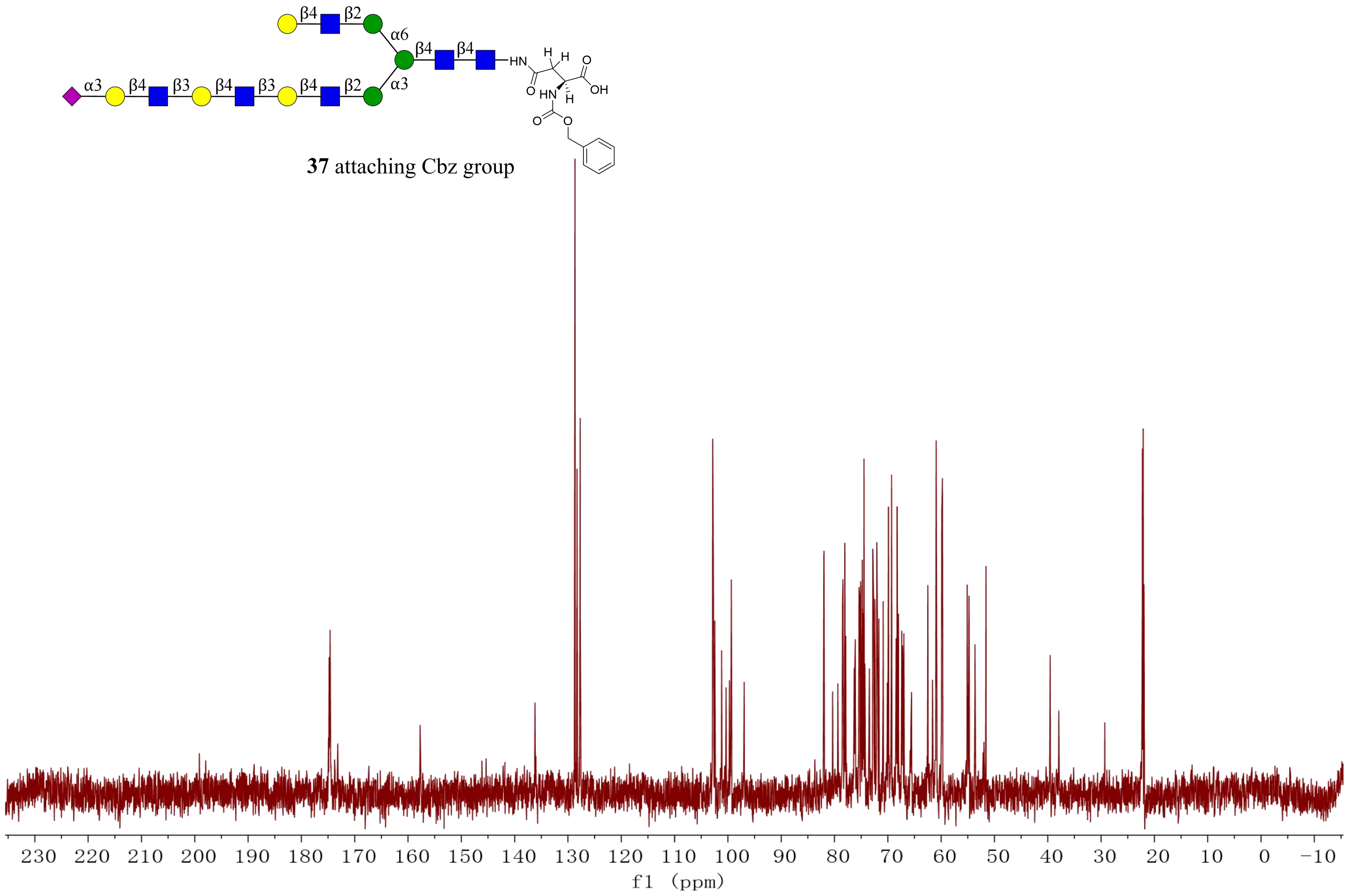


**37** attaching Cbz group

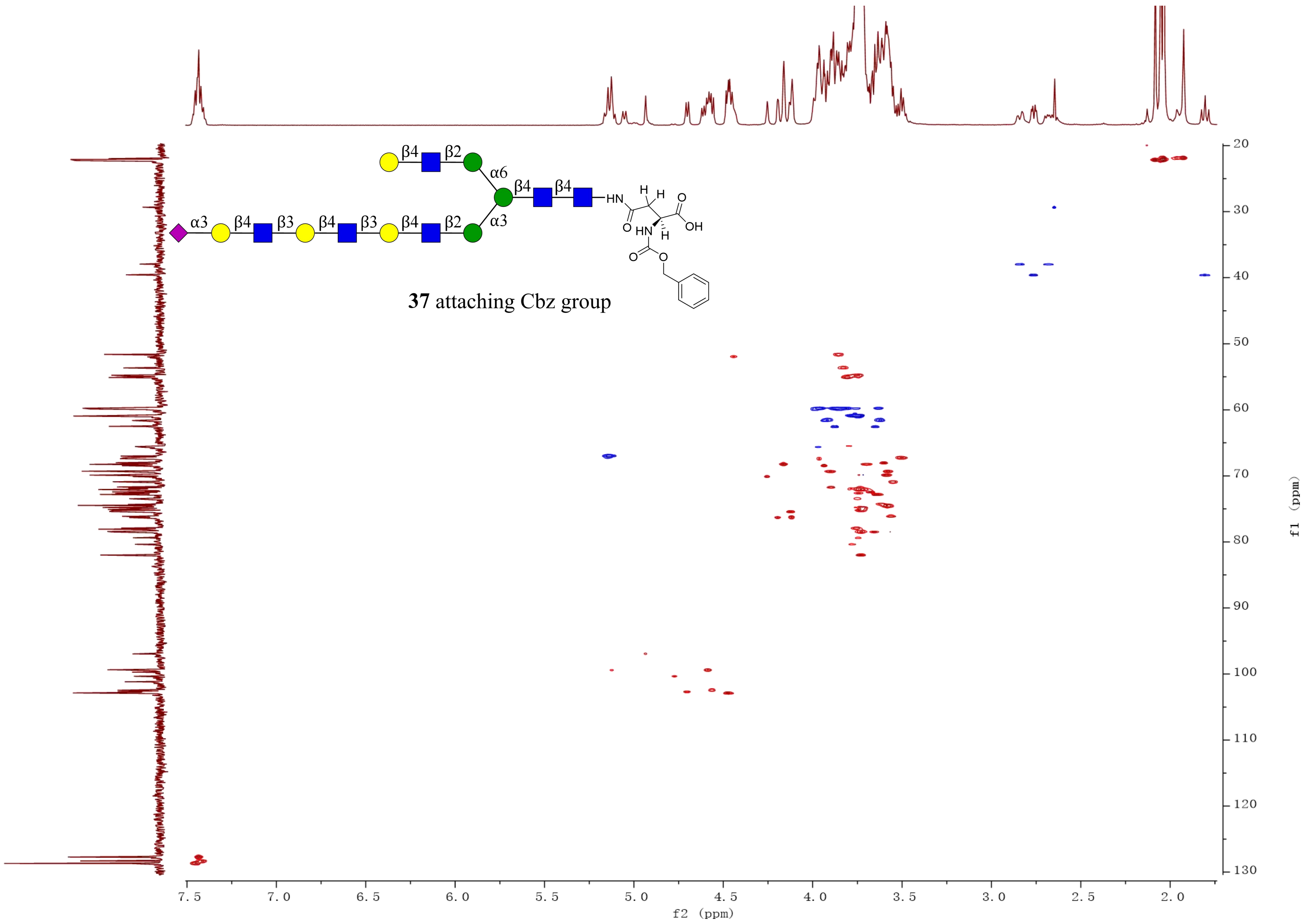
β4
β2
α6
β4
β4
HN
H
H
O
OH
O
HN
H
O
O
α3
β4
β3
β4
β3
β4
β2
α3
37 attaching Cbz group
f2 (ppm)
f1 (ppm)

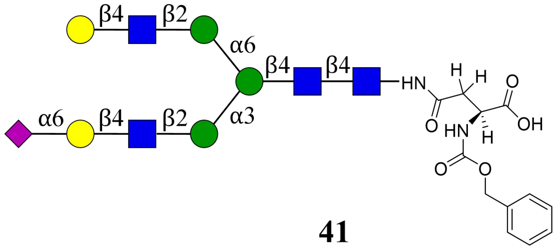
β4
β2
α6
β4
β4
HN
H
H
O
OH
O
HN
H
O
O
α6
β4
β2
α3

**41**

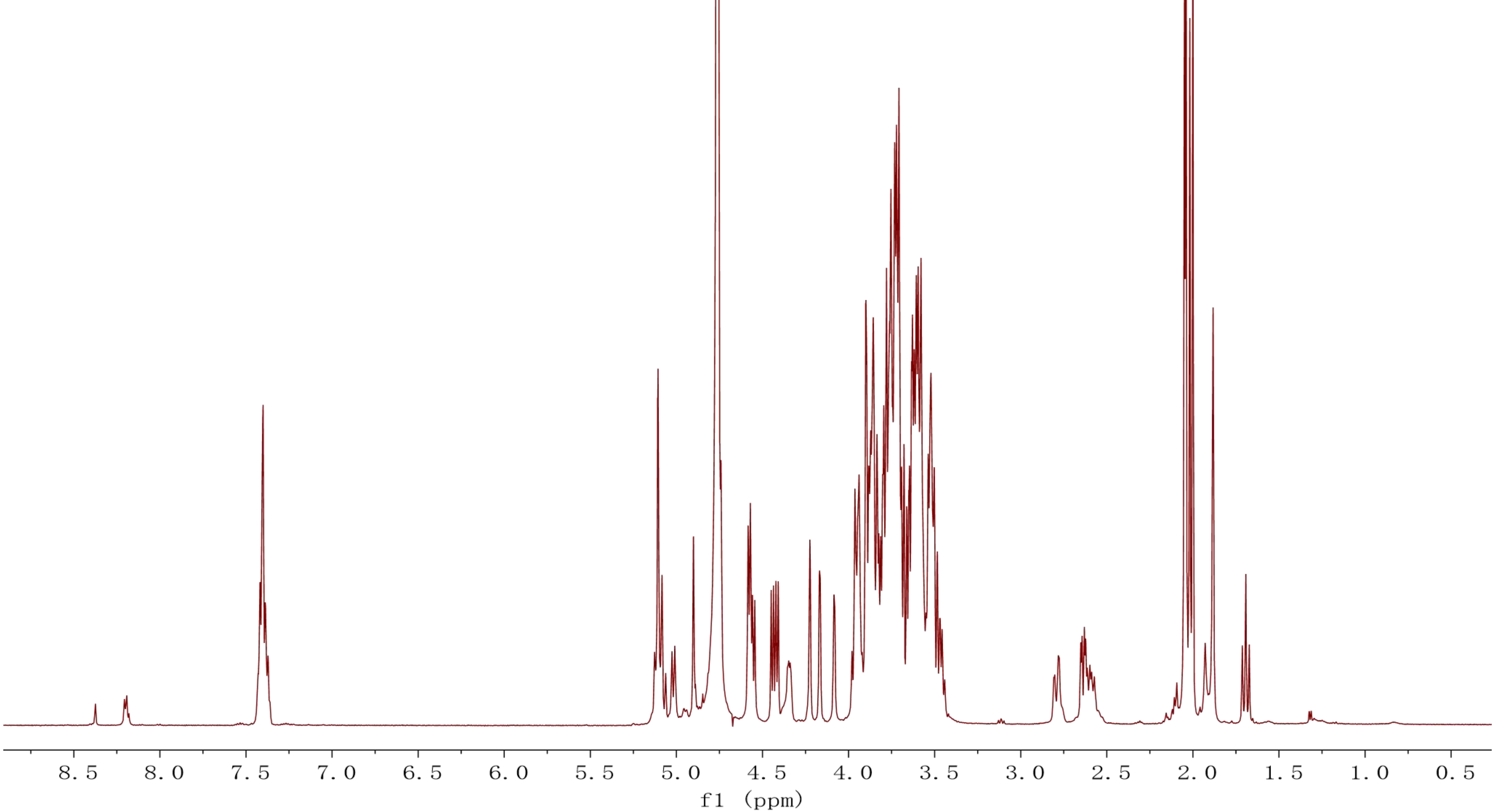
8.5
8.0
7.5
7.0
6.5
6.0
5.5
5.0
4.5
4.0
3.5
3.0
2.5
2.0
1.5
1.0
0.5
f1 (ppm)

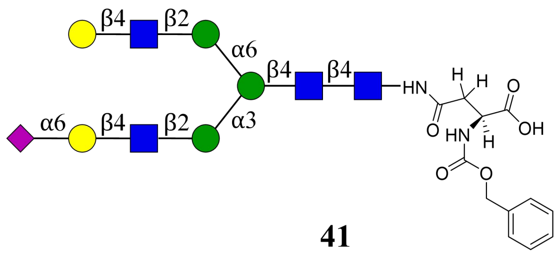
β4
β2
α6
β4
β4
HN
H
H
O
OH
O
HN
H
O
O
α6
β4
β2
α3

**41**

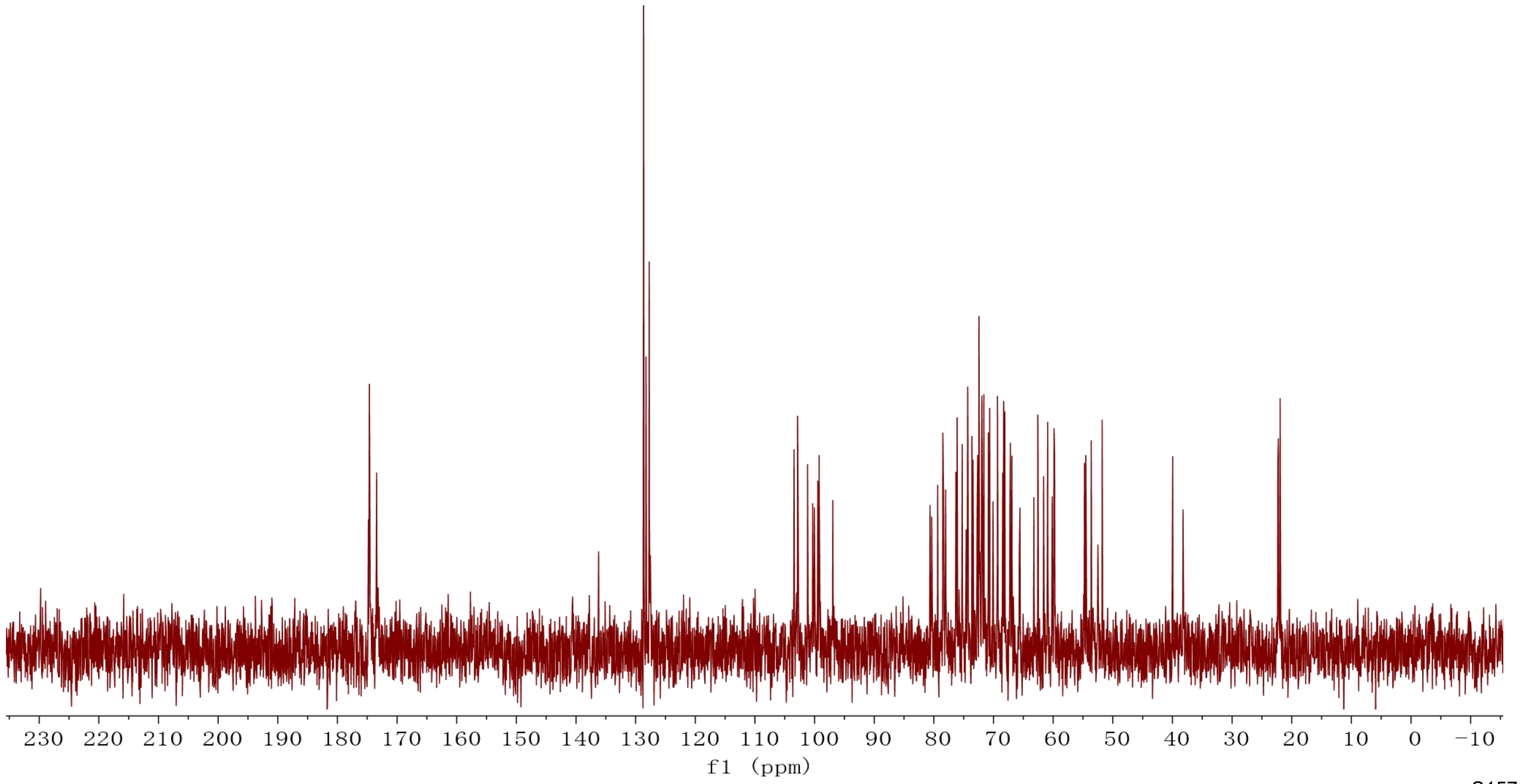
230 220 210 200 190 180 170 160 150 140 130 120 110 100 90 80 70 60 50 40 30 20 10 0 −10
f1 (ppm)

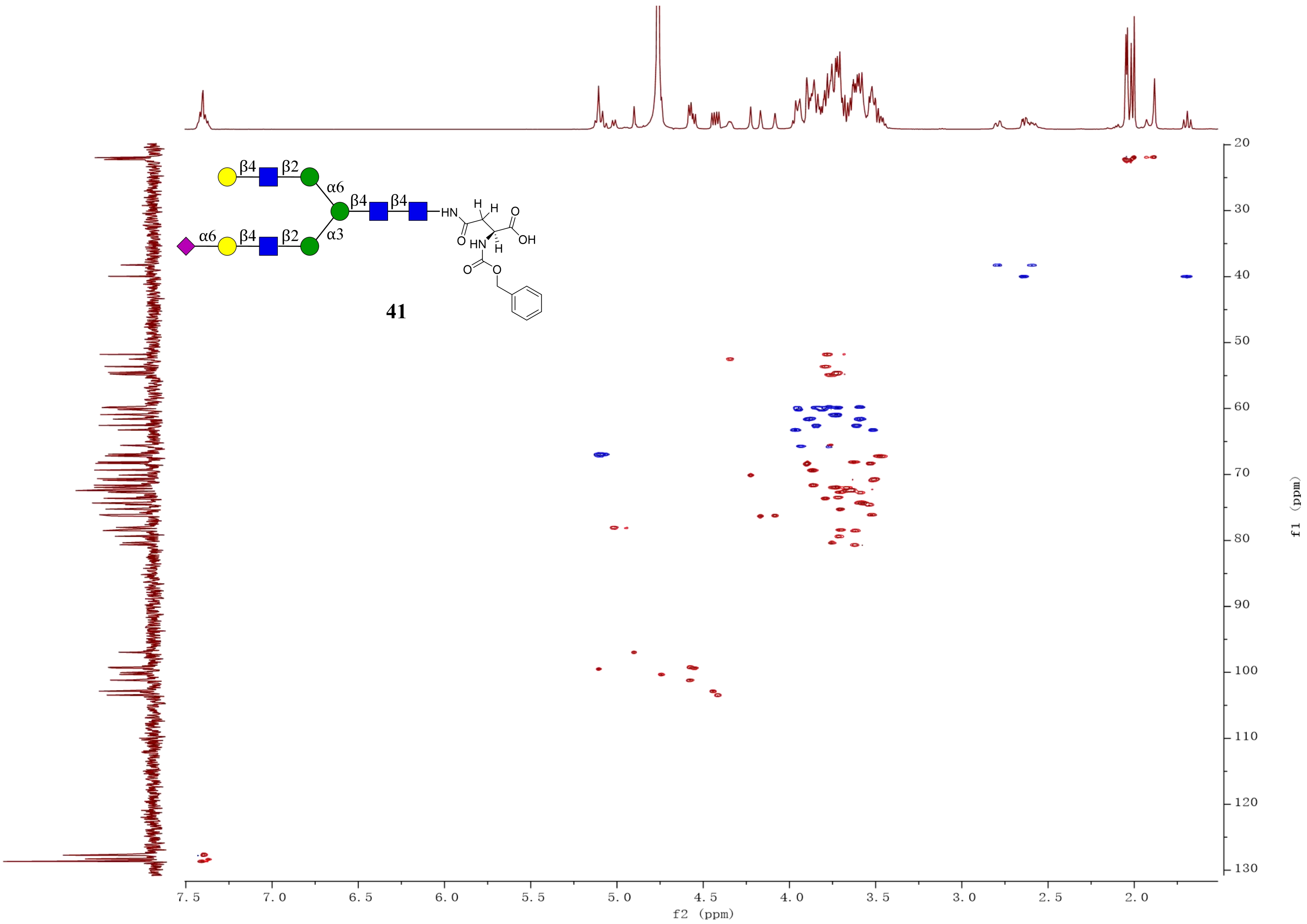
β4
β2
α6
β4
β4
HN
H
H
O
OH
α6
β4
β2
α3
O
HN
H
O
O
41
f2 (ppm)
f1 (ppm)

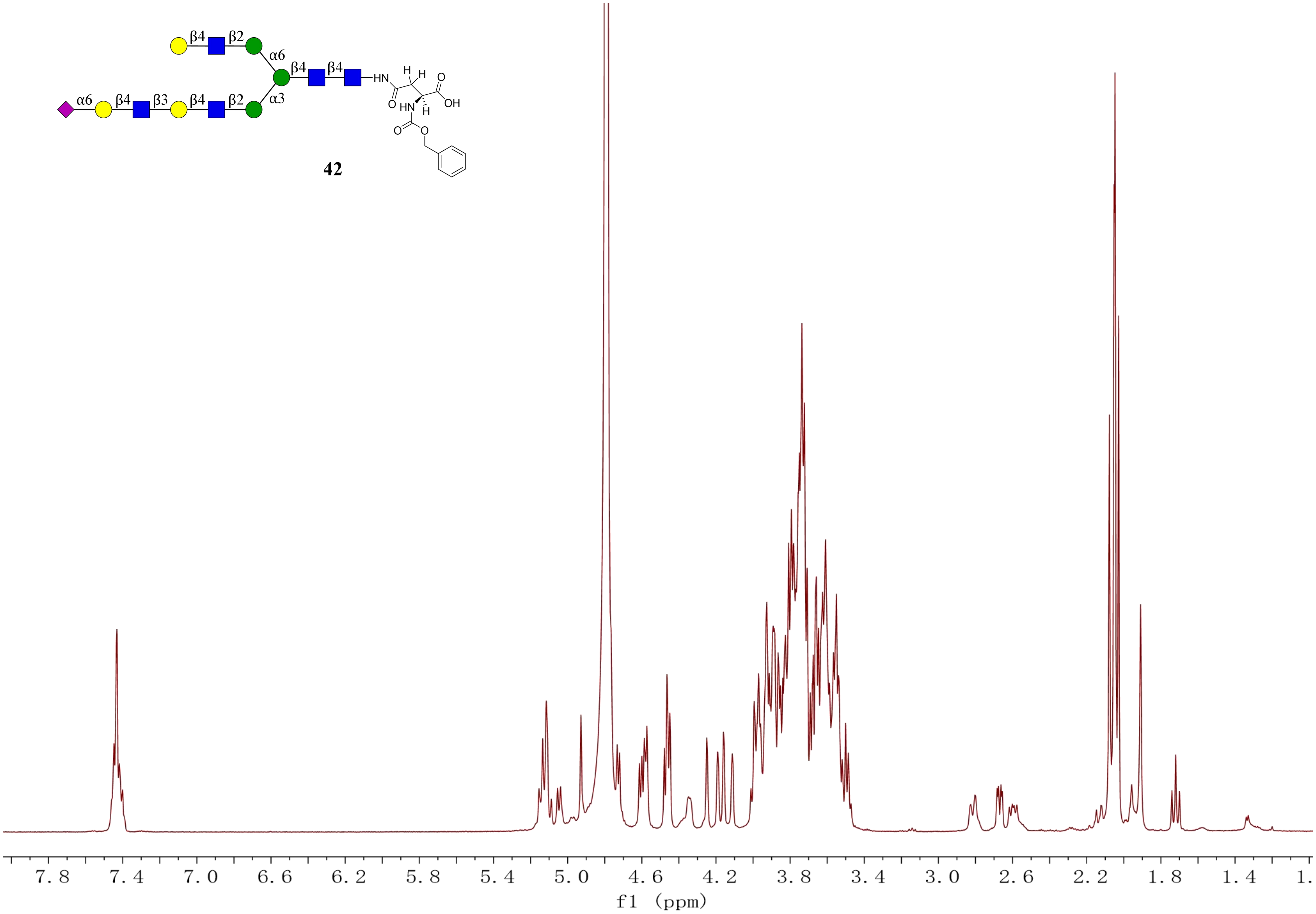

β4 β2 α6
β4 β4 HN
α6 β4 β3 β4 β2 α3
H H O OH
O HN H
O O
7.8 7.4 7.0 6.6 6.2 5.8 5.4 5.0 4.6 4.2 3.8 3.4 3.0 2.6 2.2 1.8 1.4 1.
f1 (ppm)


42

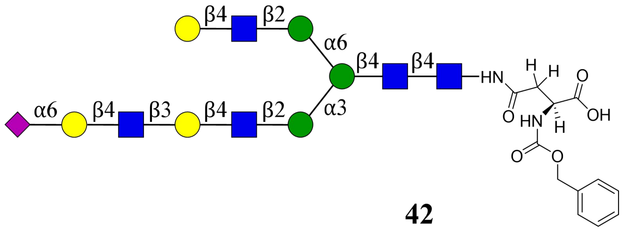
β4
β2
α6
β4
β4
HN
H
H
O
OH
O
HN
H
O
O
α6
β4
β3
β4
β2
α3

**42**

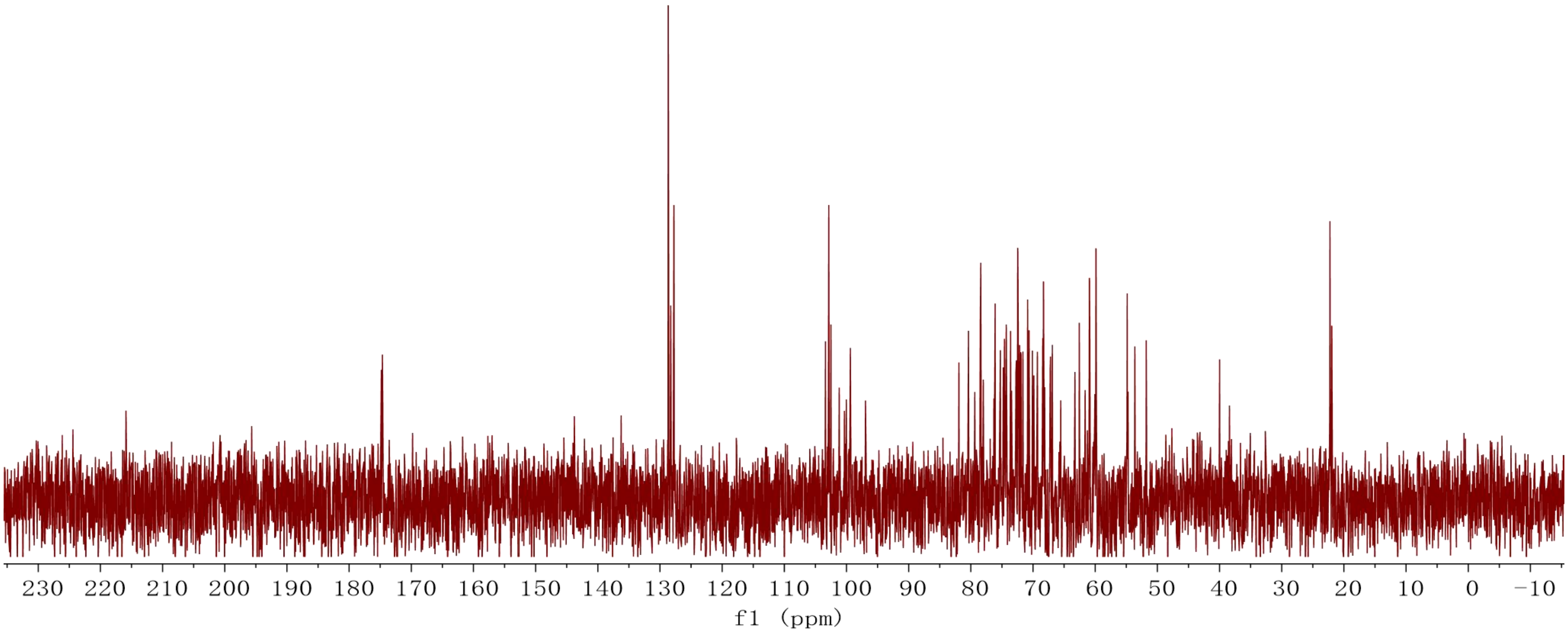
230 220 210 200 190 180 170 160 150 140 130 120 110 100 90 80 70 60 50 40 30 20 10 0 −10
f1 (ppm)

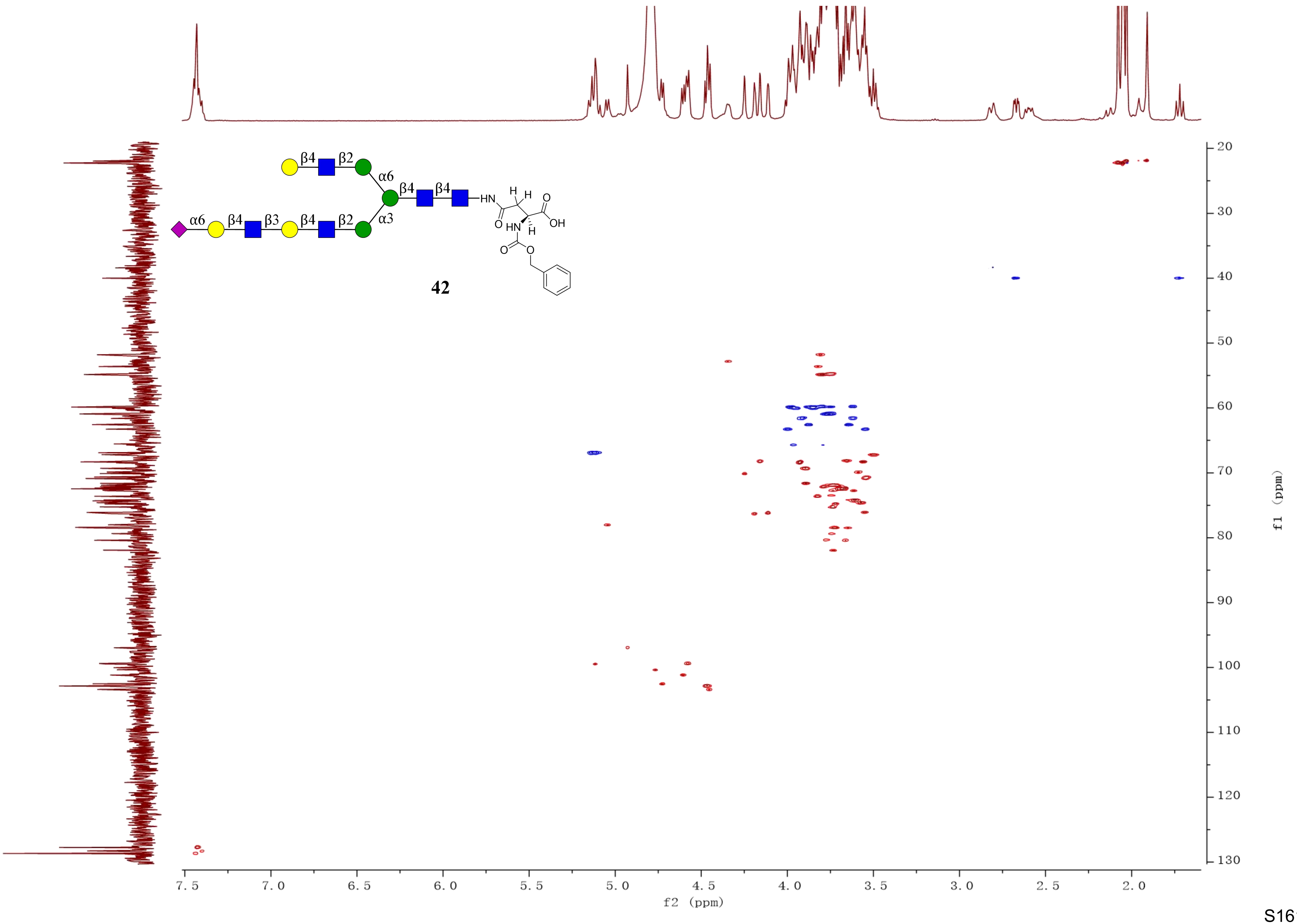

β4
β2
α6
β4
β4
HN
α6
β4
β3
β4
β2
α3
H
H
O
OH
O
HN
H
O
O
42
f2 (ppm)
f1 (ppm)

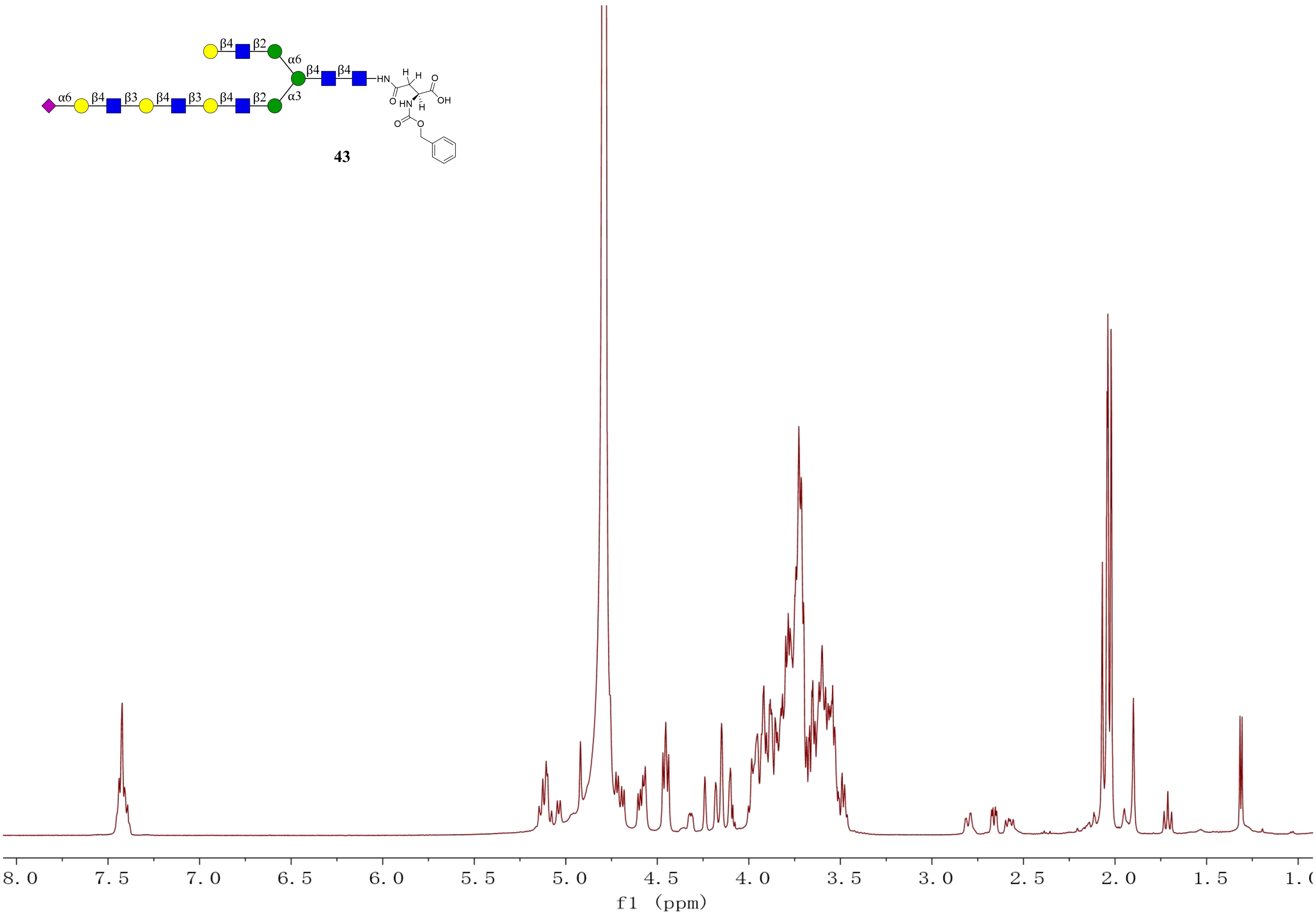
β4
β2
α6
β4
β4
HN
H
H
O
O
HN
H
OH
α6
β4
β3
β4
β3
β4
β2
α3
O
O
43
8.0
7.5
7.0
6.5
6.0
5.5
5.0
4.5
4.0
3.5
3.0
2.5
2.0
1.5
f1 (ppm)

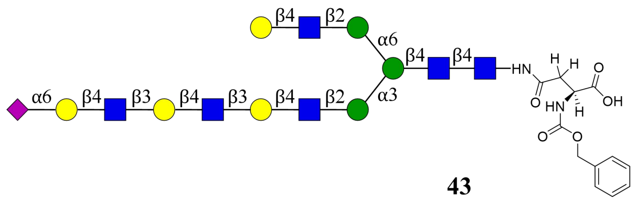
α6
β4
β3
β2
α3
HN
O
H
OH
O

**43**

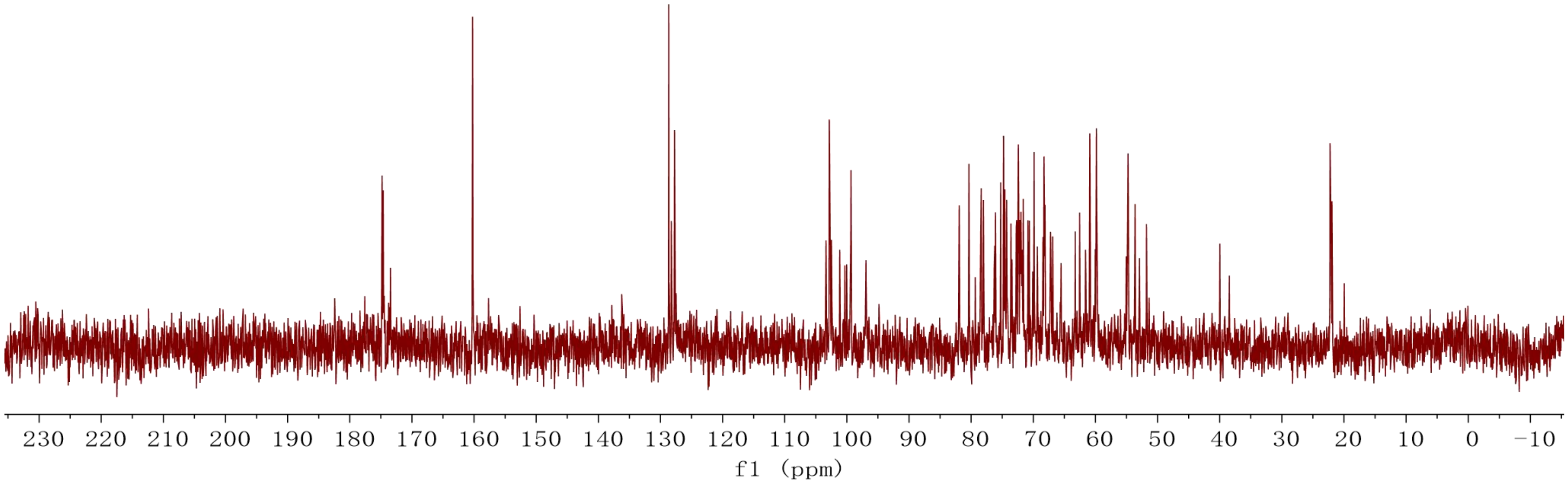
230 220 210 200 190 180 170 160 150 140 130 120 110 100 90 80 70 60 50 40 30 20 10 0 −10
f1 (ppm)

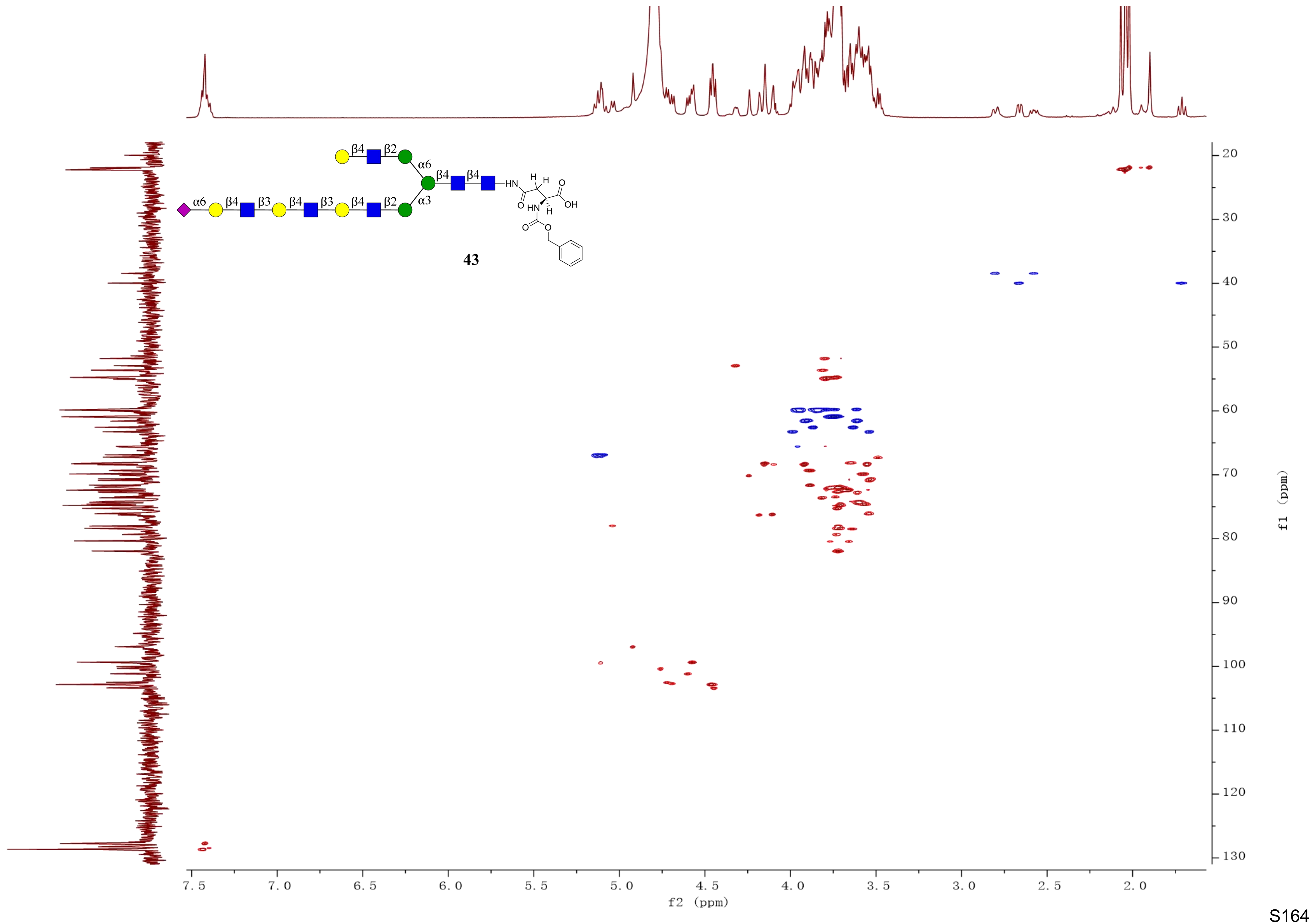

β4
β2
α6
β4
β4
α6
β4
β3
β4
β3
β4
β2
α3
HN
H
H
O
OH
O
HN
H
O
O
43
7.5
7.0
6.5
6.0
5.5
5.0
4.5
4.0
3.5
3.0
2.5
2.0
f2 (ppm)
20
30
40
50
60
70
80
90
100
110
120
130
f1 (ppm)

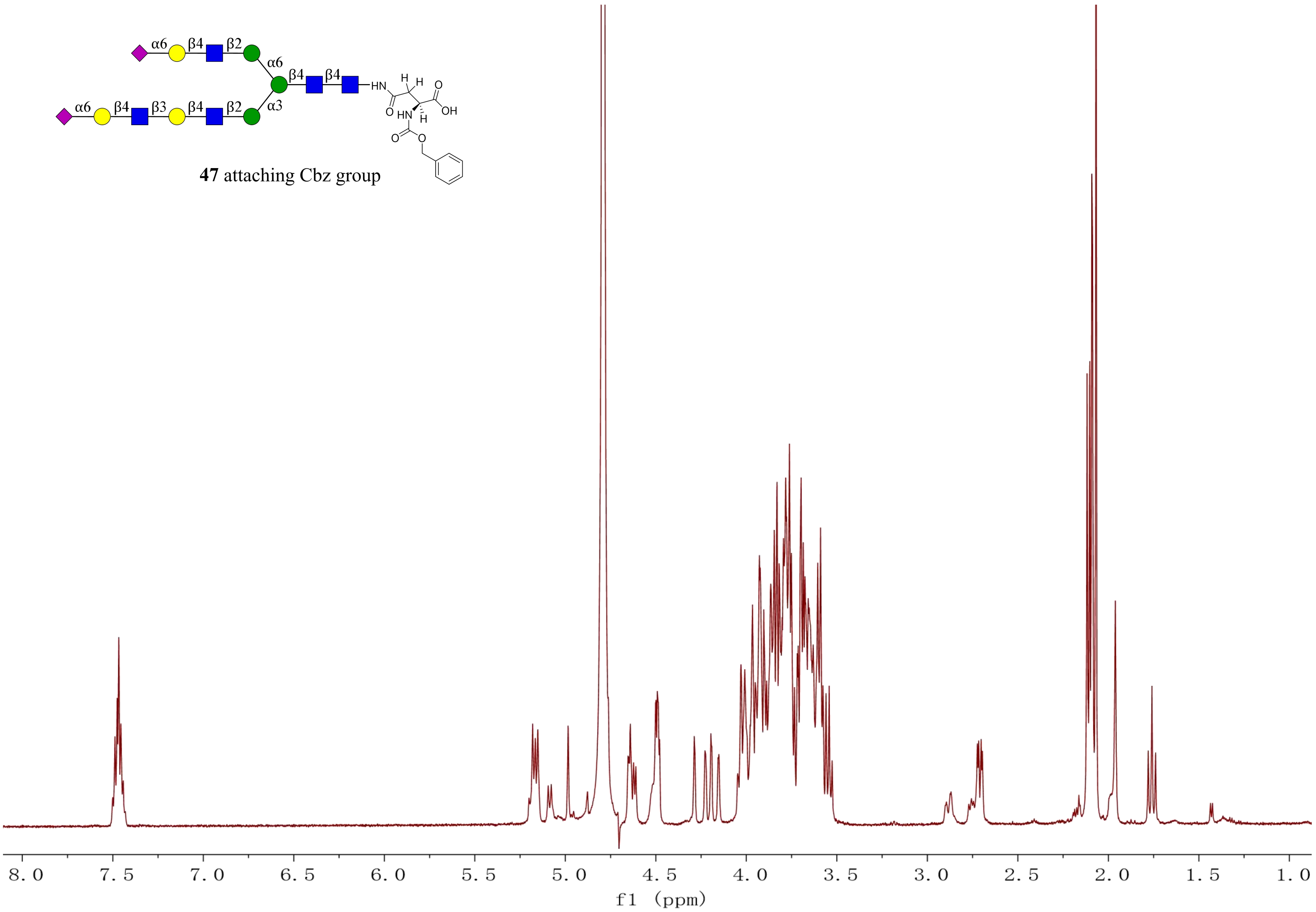


**47** attaching Cbz group

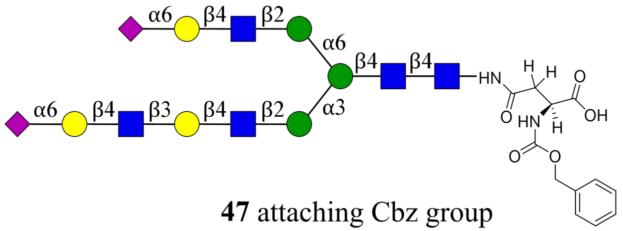


**47** attaching Cbz group

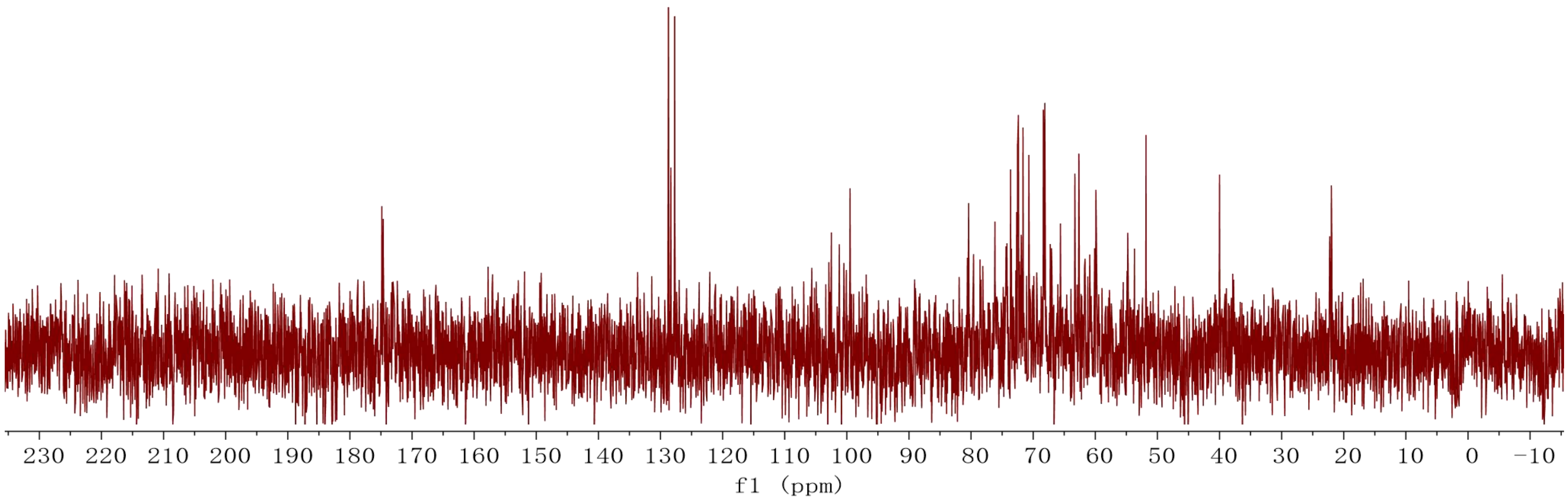

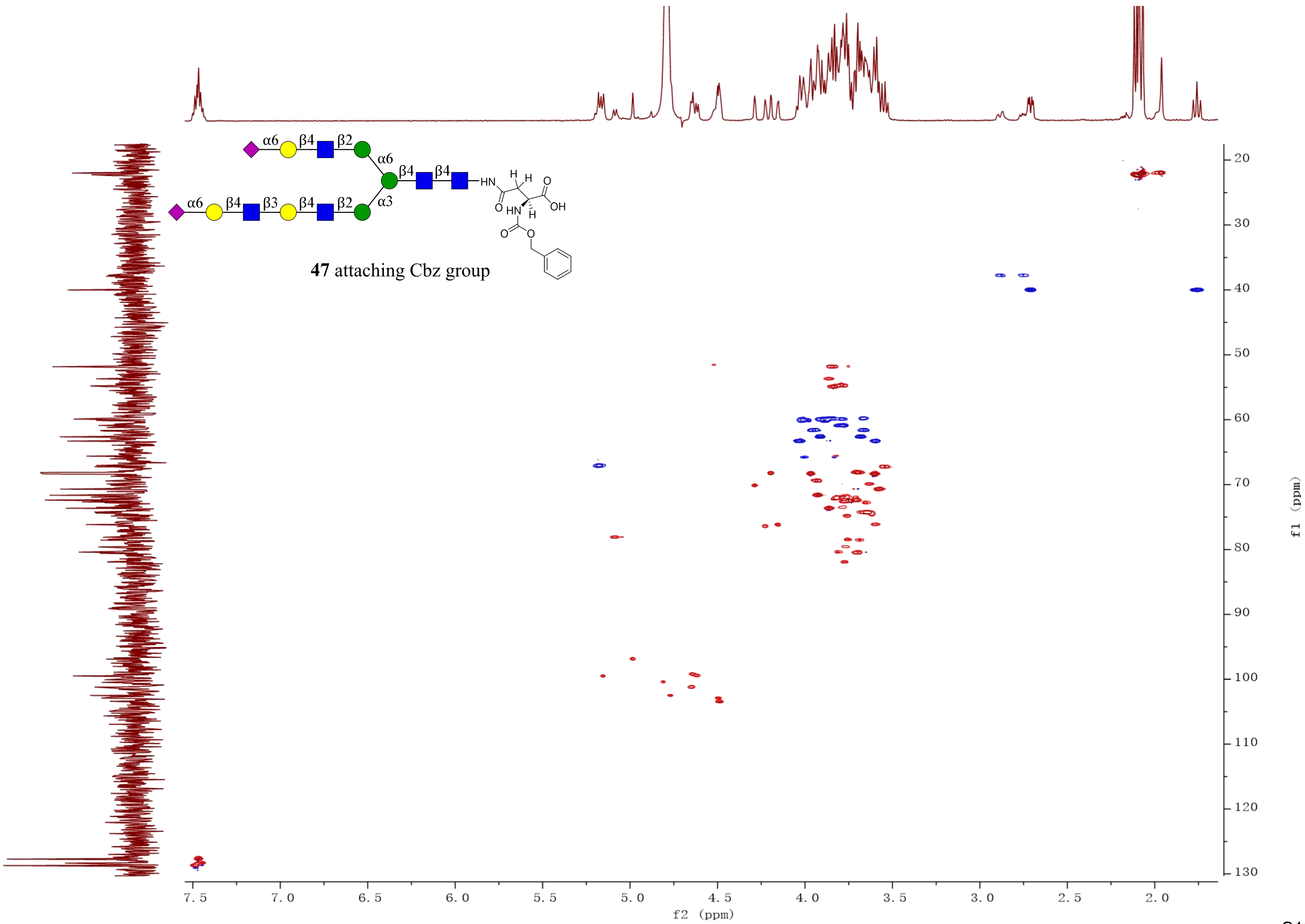

47 attaching Cbz group
f2 (ppm)
f1 (ppm)

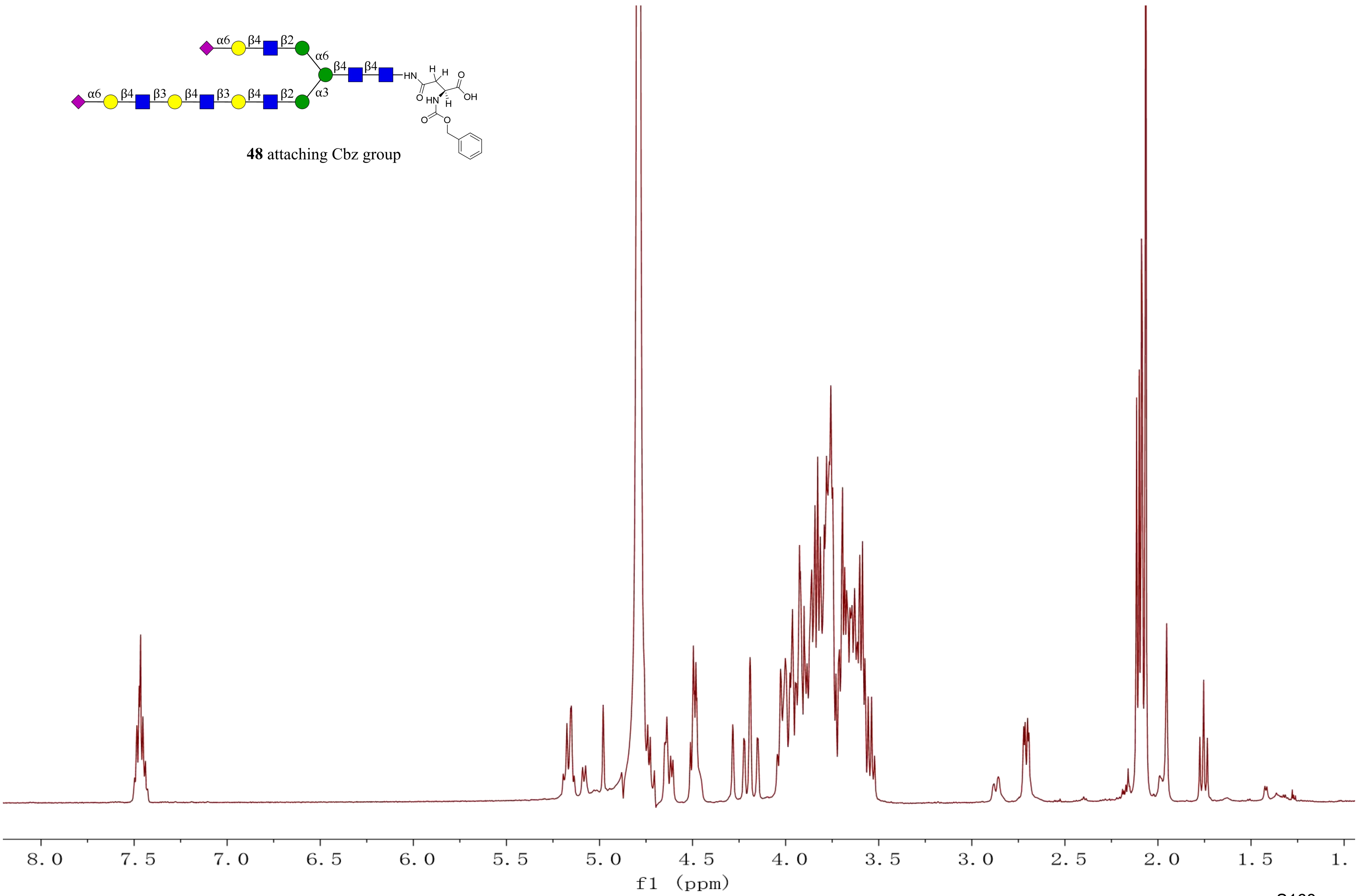


**48** attaching Cbz group

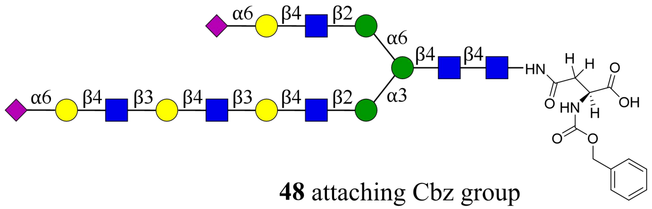


**48** attaching Cbz group

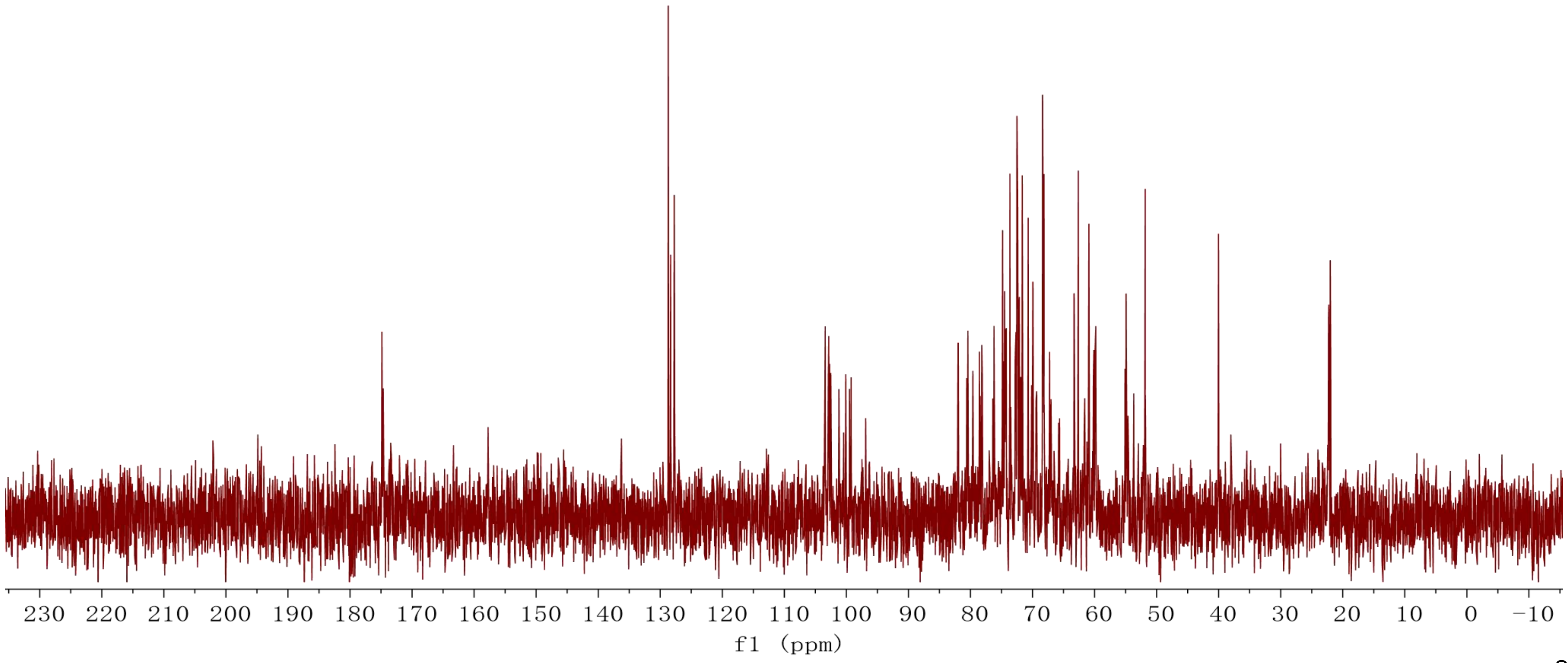

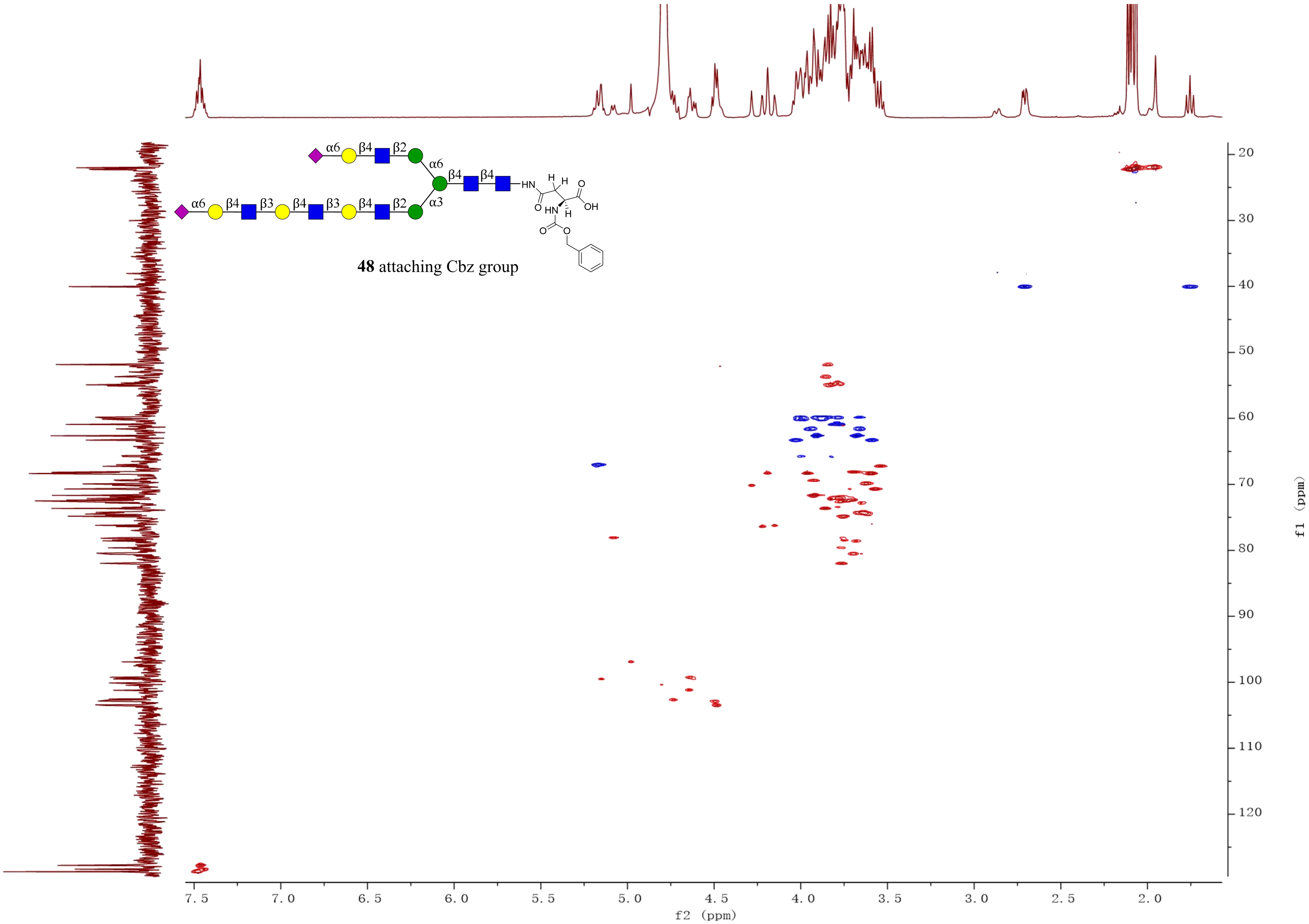


**48** attaching Cbz group

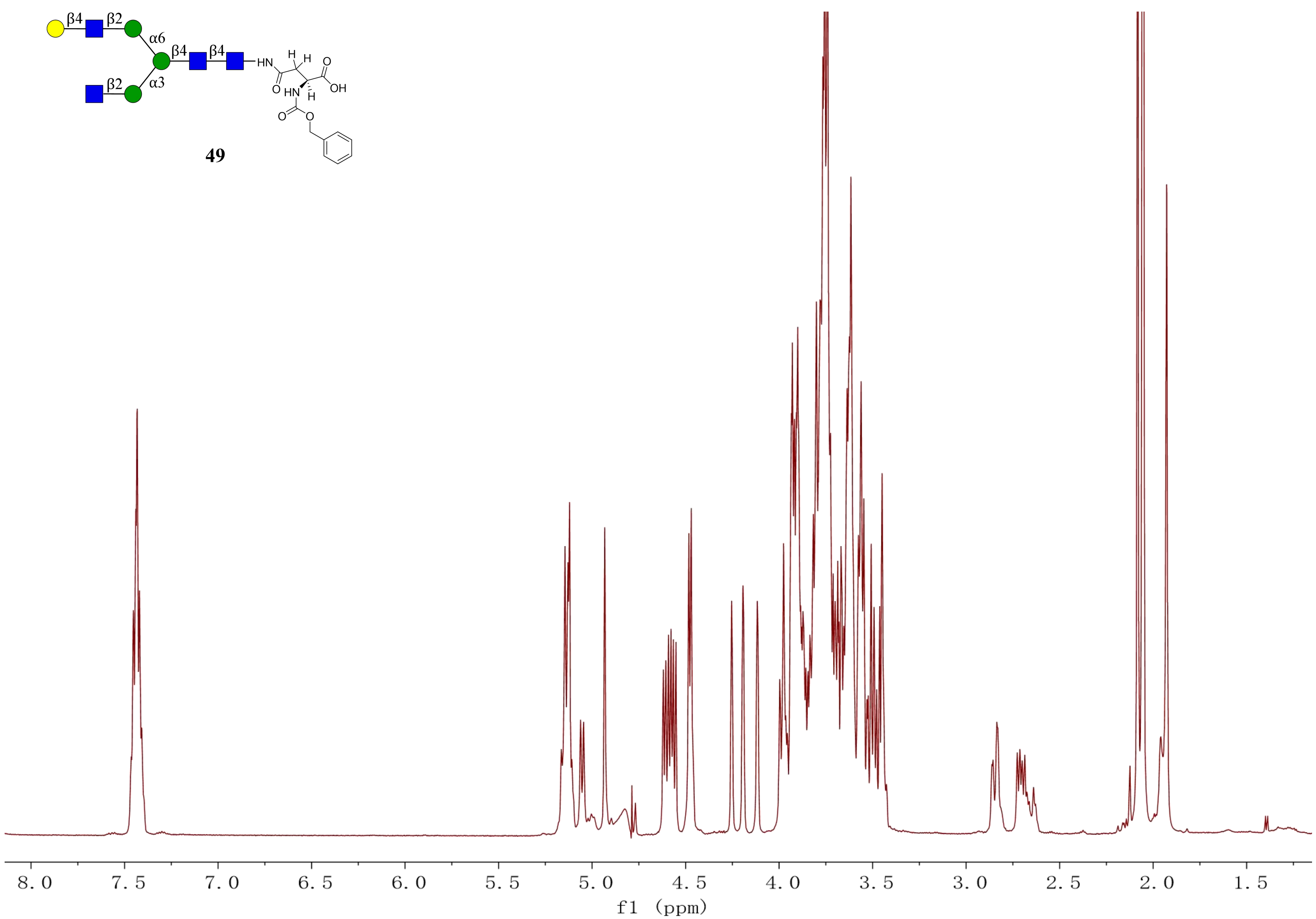
β4
β2
α6
β4
β4
β2
α3
HN
H
H
O
OH
O
HN
H
O
O
49
8.0
7.5
7.0
6.5
6.0
5.5
5.0
4.5
4.0
3.5
3.0
2.5
2.0
1.5
f1 (ppm)

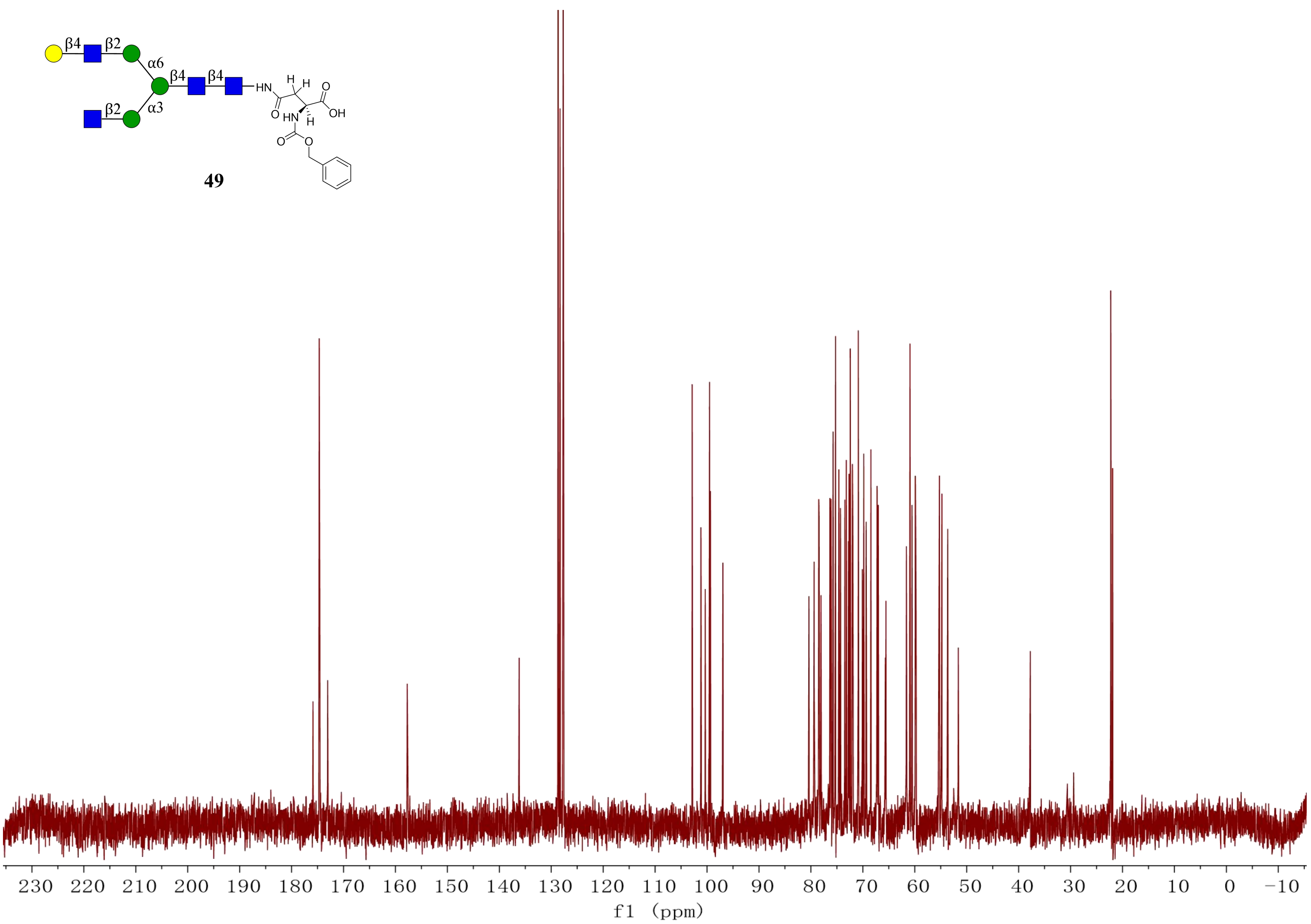

49
230 220 210 200 190 180 170 160 150 140 130 120 110 100 90 80 70 60 50 40 30 20 10 0 −10
f1 (ppm)

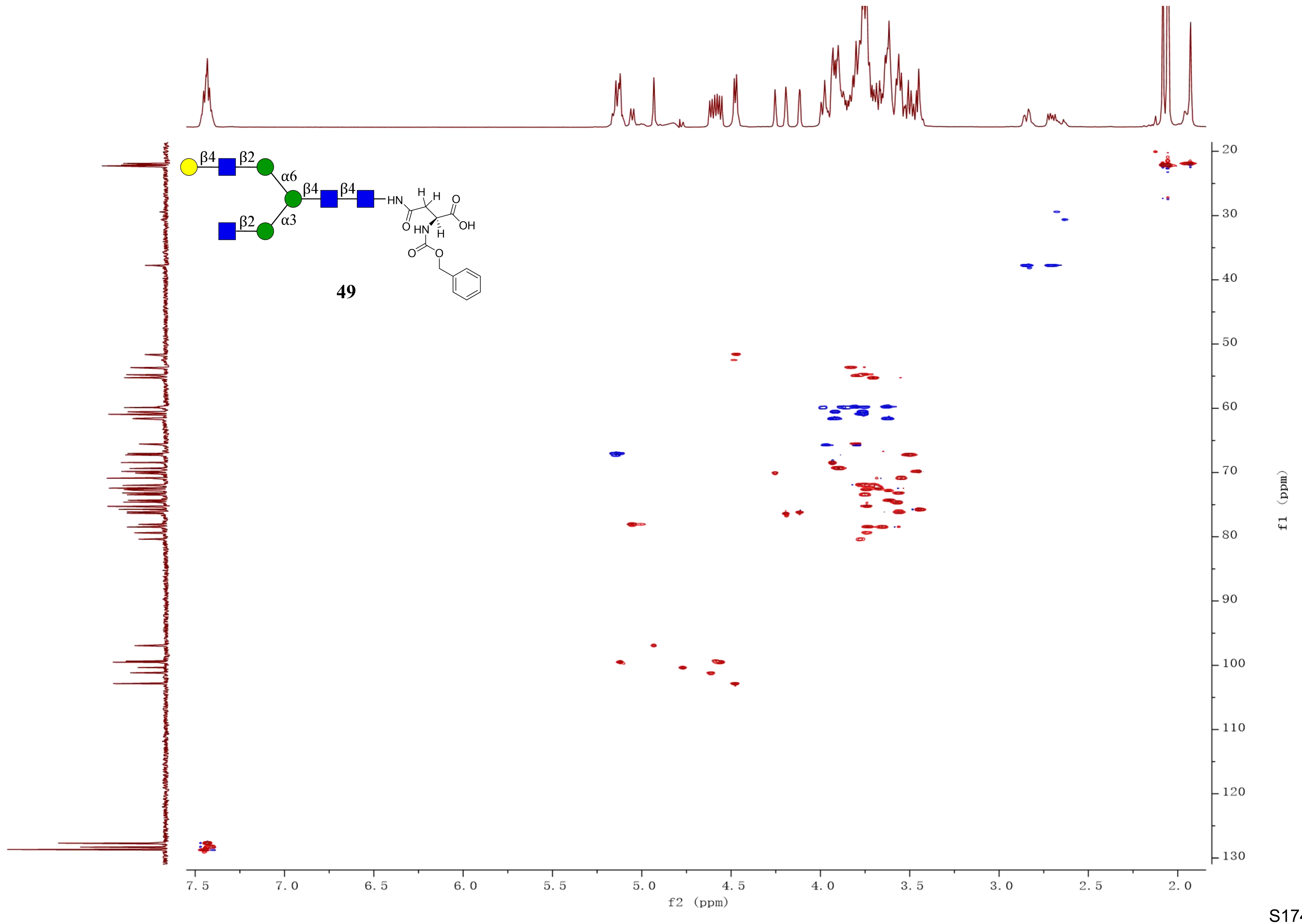
β4
β2
α6
β4
β4
HN
H
H
O
OH
O
HN
H
O
O
β2
α3
49
7.5
7.0
6.5
6.0
5.5
5.0
4.5
4.0
3.5
3.0
2.5
2.0
f2 (ppm)
20
30
40
50
60
70
80
90
100
110
120
130
f1 (ppm)

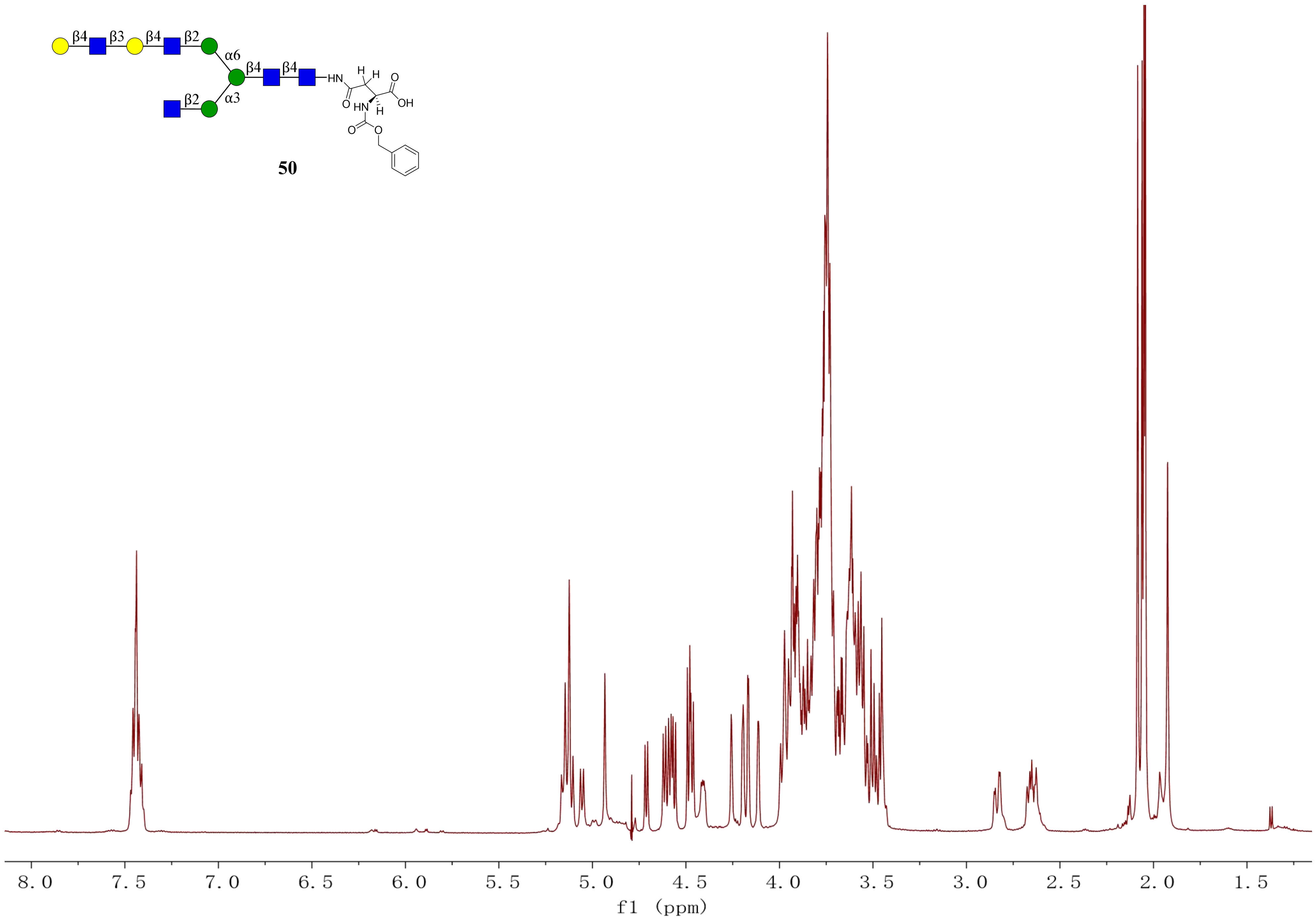
β4
β3
β4
β2
α6
β4
β4
α3
β2
HN
H
H
O
O
OH
HN
H
O
O

**50**

8.0
7.5
7.0
6.5
6.0
5.5
5.0
4.5
4.0
3.5
3.0
2.5
2.0
1.5
f1 (ppm)

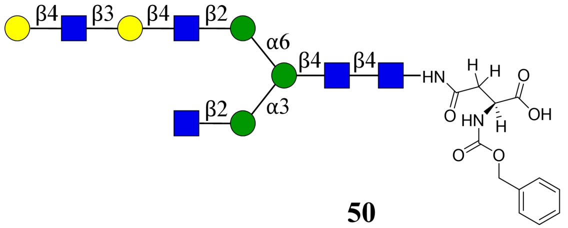
β4
β3
β4
β2
α6
β4
β4
HN
H
H
O
OH
O
HN
H
O
O
β2
α3

**50**

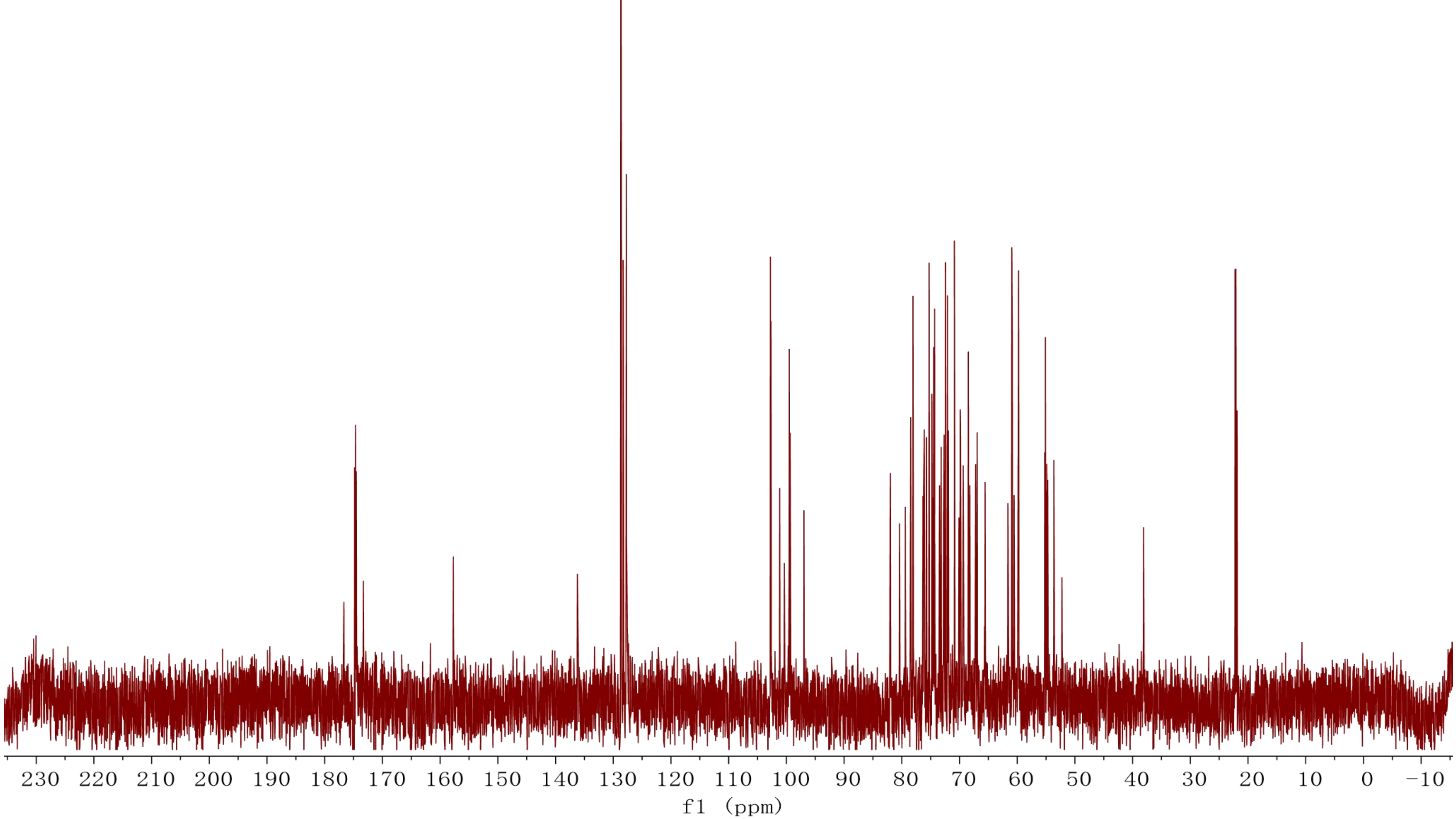
230 220 210 200 190 180 170 160 150 140 130 120 110 100 90 80 70 60 50 40 30 20 10 0 −10
f1 (ppm)

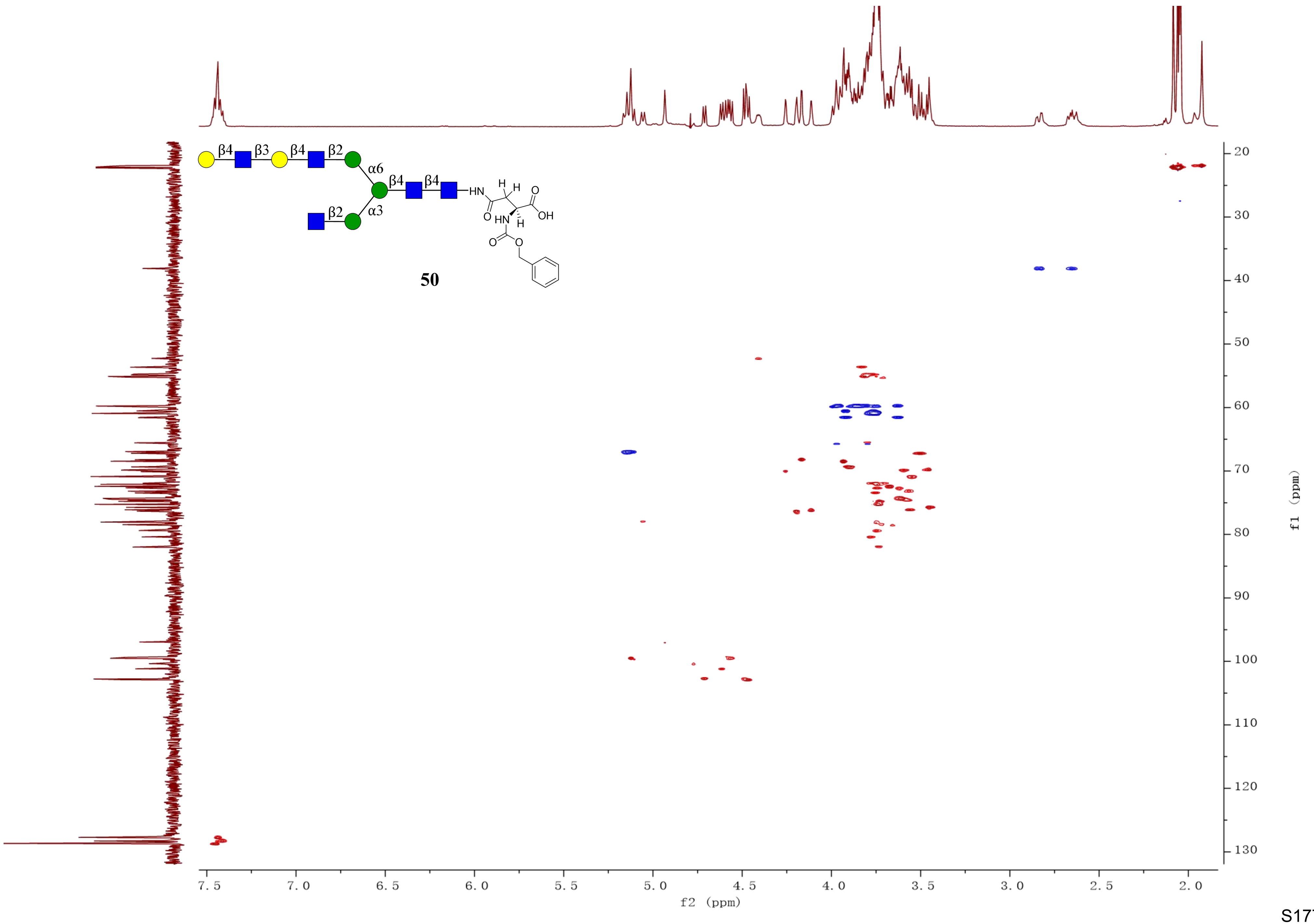
β4
β3
β4
β2
α6
β4
β4
β2
α3
HN
H
H
O
OH
O
HN
H
O
O
50
f2 (ppm)
f1 (ppm)

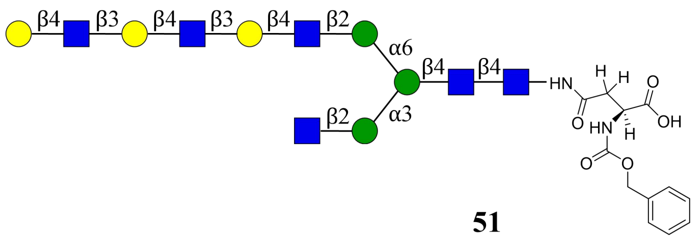
β4
β3
β4
β3
β4
β2
α6
β4
β4
HN
H
H
O
OH
O
HN
H
O
O
β2
α3

**51**

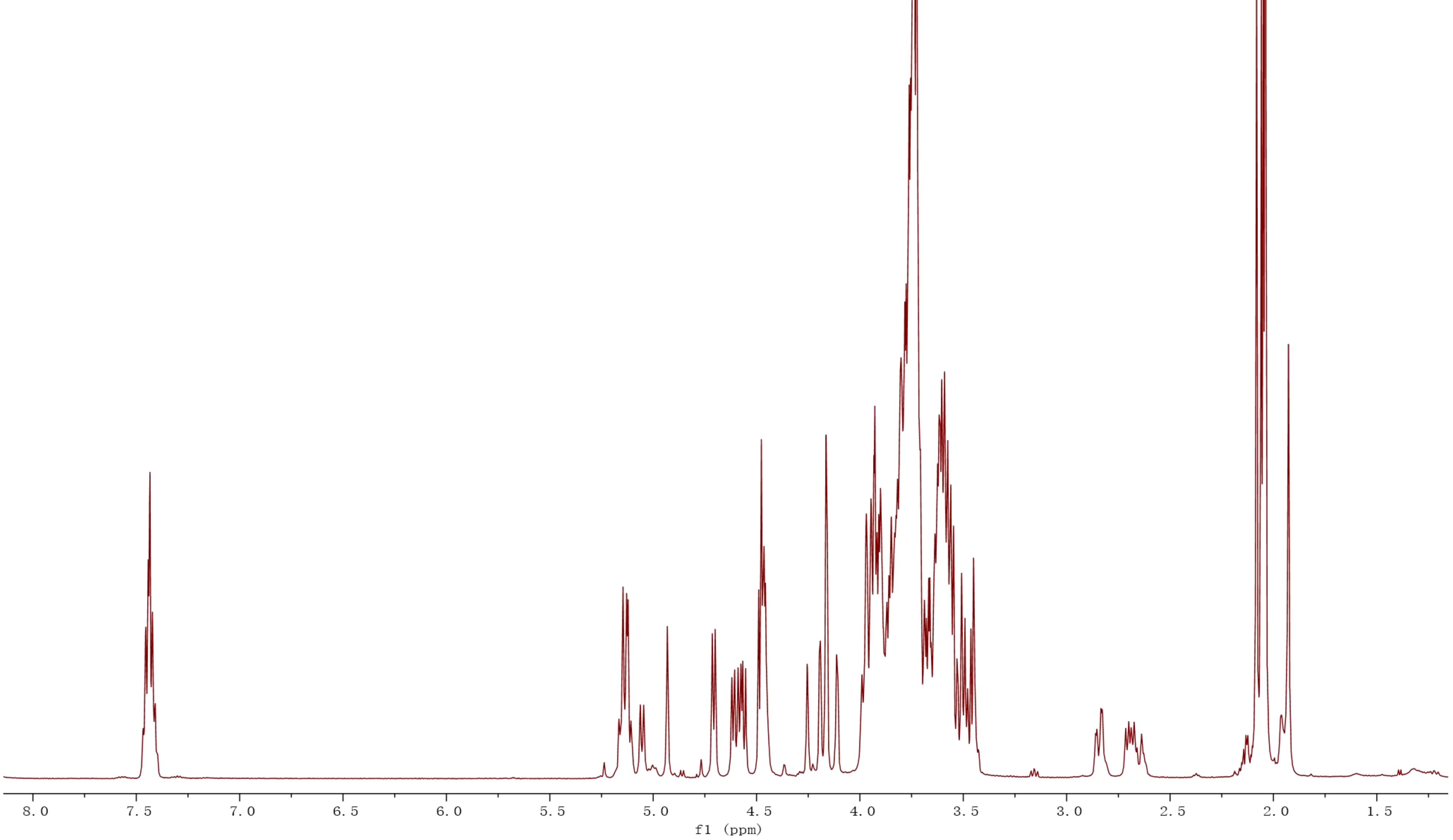
8.0
7.5
7.0
6.5
6.0
5.5
5.0
4.5
4.0
3.5
3.0
2.5
2.0
1.5
f1 (ppm)

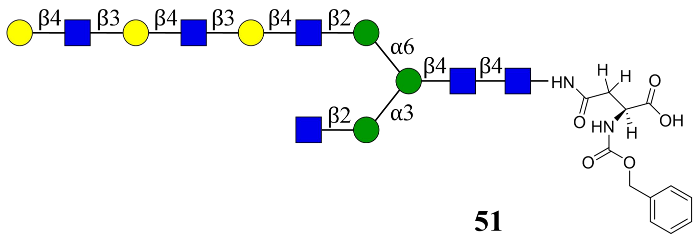

β4
β3
β4
β3
β4
β2
α6
β4
β4
HN
H
H
O
O
OH
HN
H
β2
α3
O
O


**51**

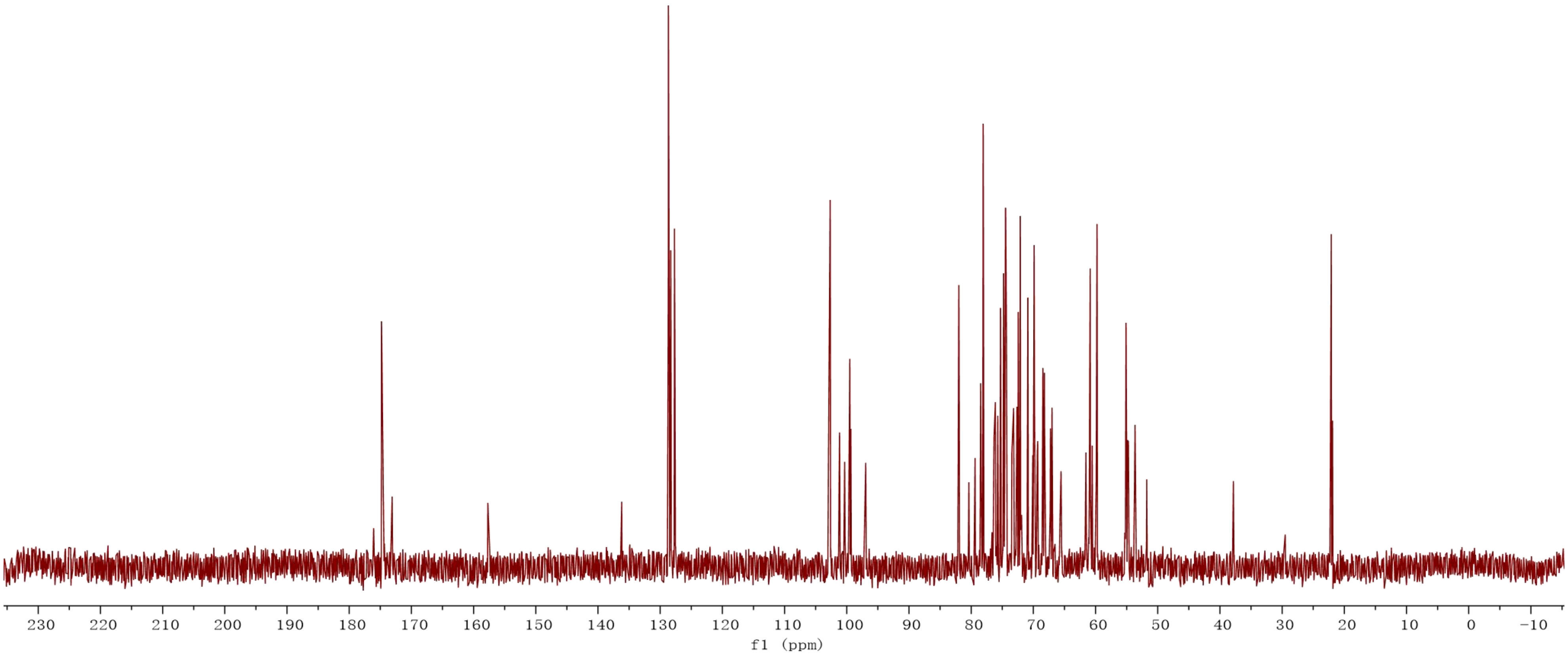

230
220
210
200
190
180
170
160
150
140
130
120
110
100
90
80
70
60
50
40
30
20
10
0
-10
f1 (ppm)

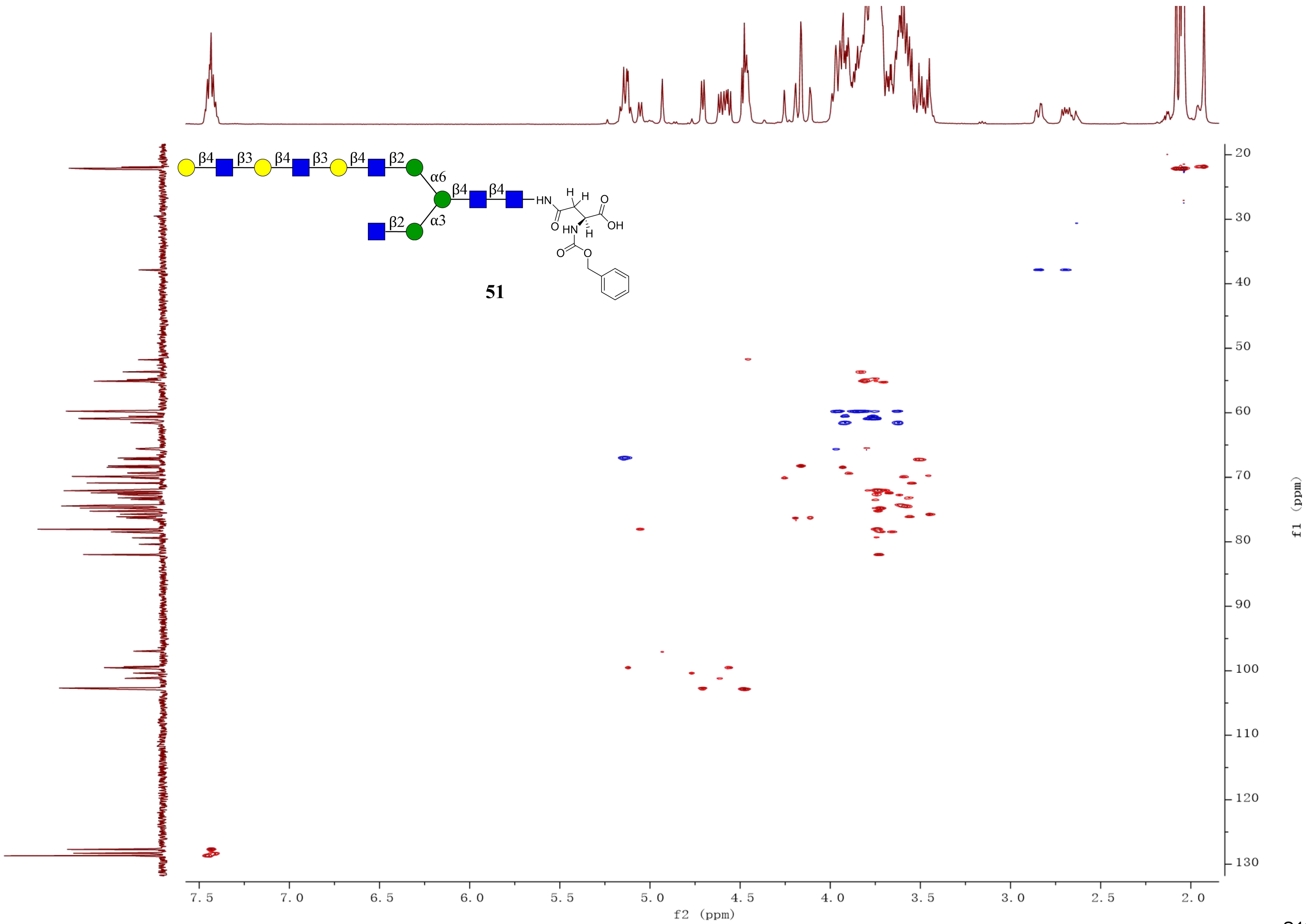

β4
β3
β4
β3
β4
β2
α6
β4
β4
β2
α3
HN
H
H
O
OH
O
HN
H
O
O
51
f2 (ppm)
f1 (ppm)

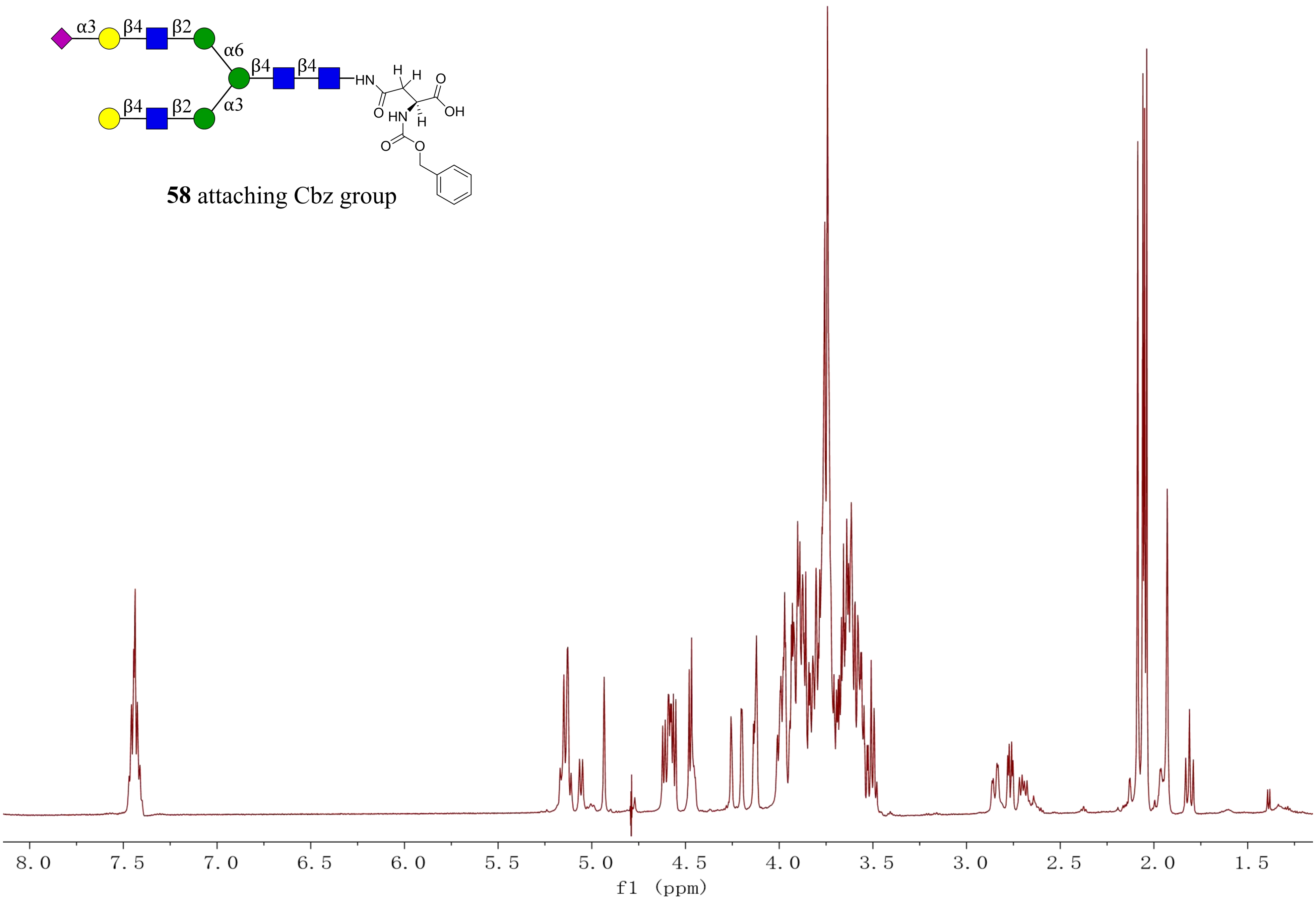
α3
β4
β2
α6
β4
β4
β4
β2
α3
HN
H
H
O
OH
O
HN
H
O
O
58 attaching Cbz group
8.0
7.5
7.0
6.5
6.0
5.5
5.0
4.5
4.0
3.5
3.0
2.5
2.0
1.5
f1 (ppm)

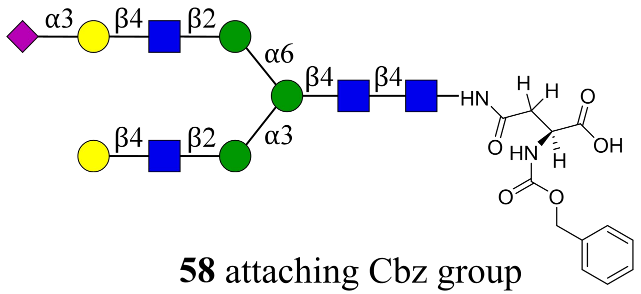


**58** attaching Cbz group

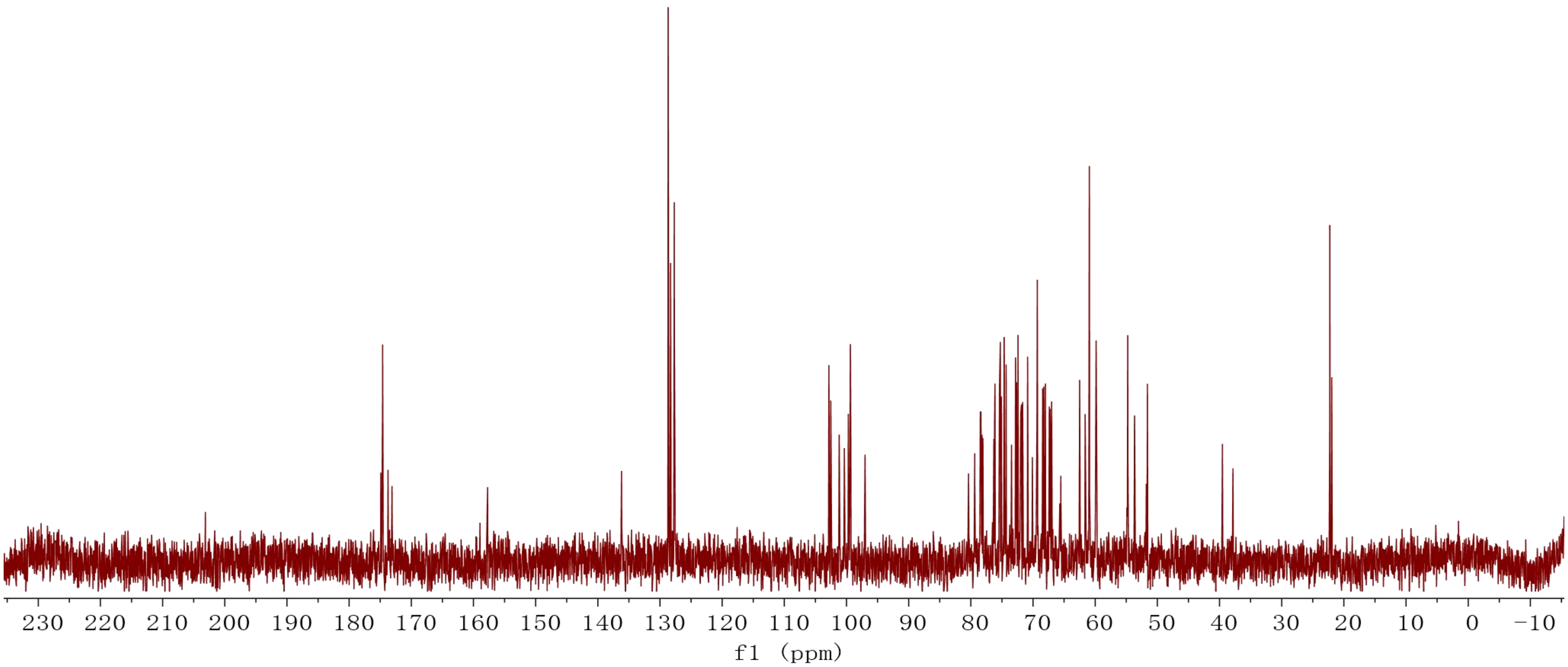

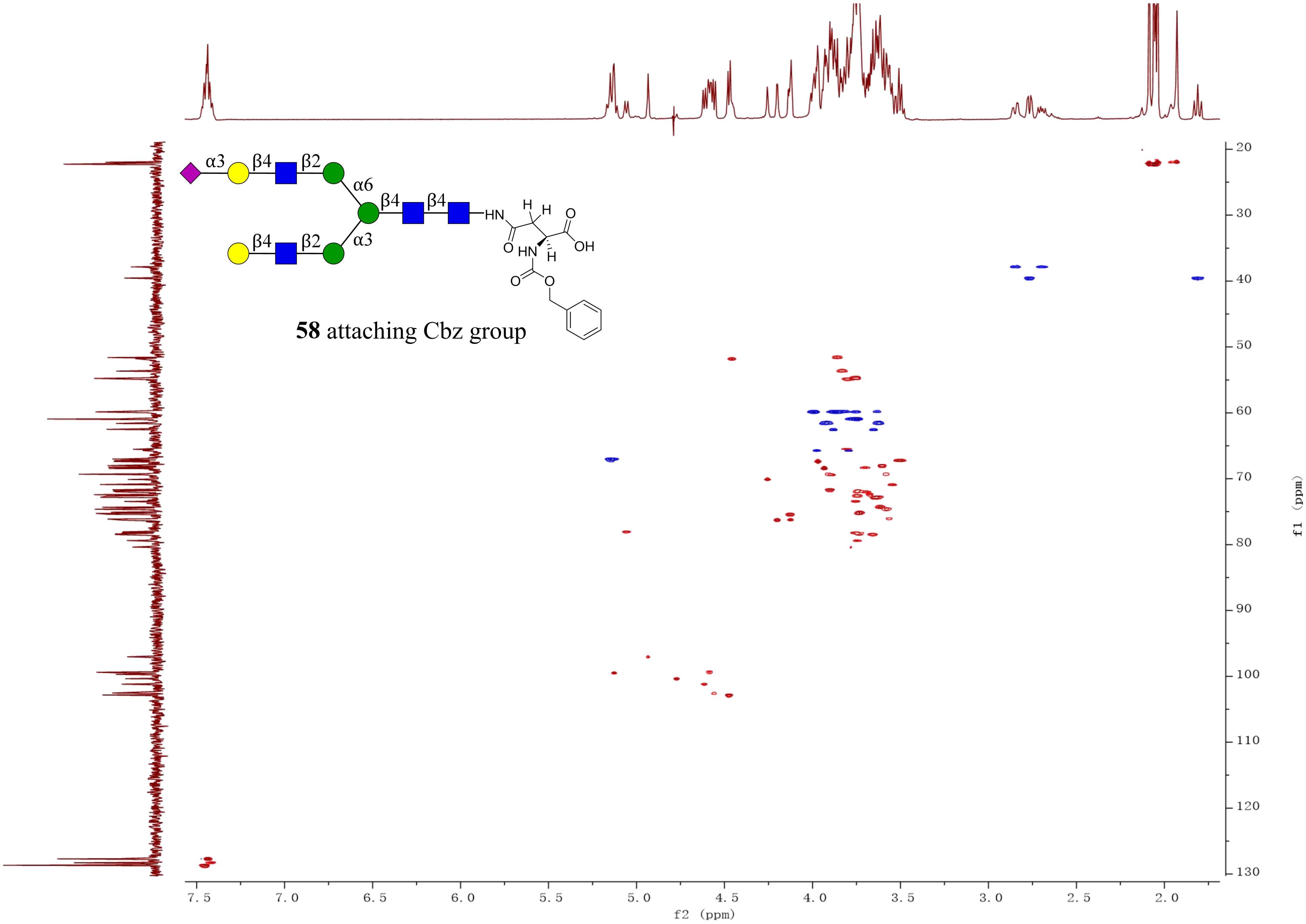
α3
β4
β2
α6
β4
β4
β4
β2
α3
HN
H
H
O
OH
O
HN
H
O
O
58 attaching Cbz group
f2 (ppm)
f1 (ppm)

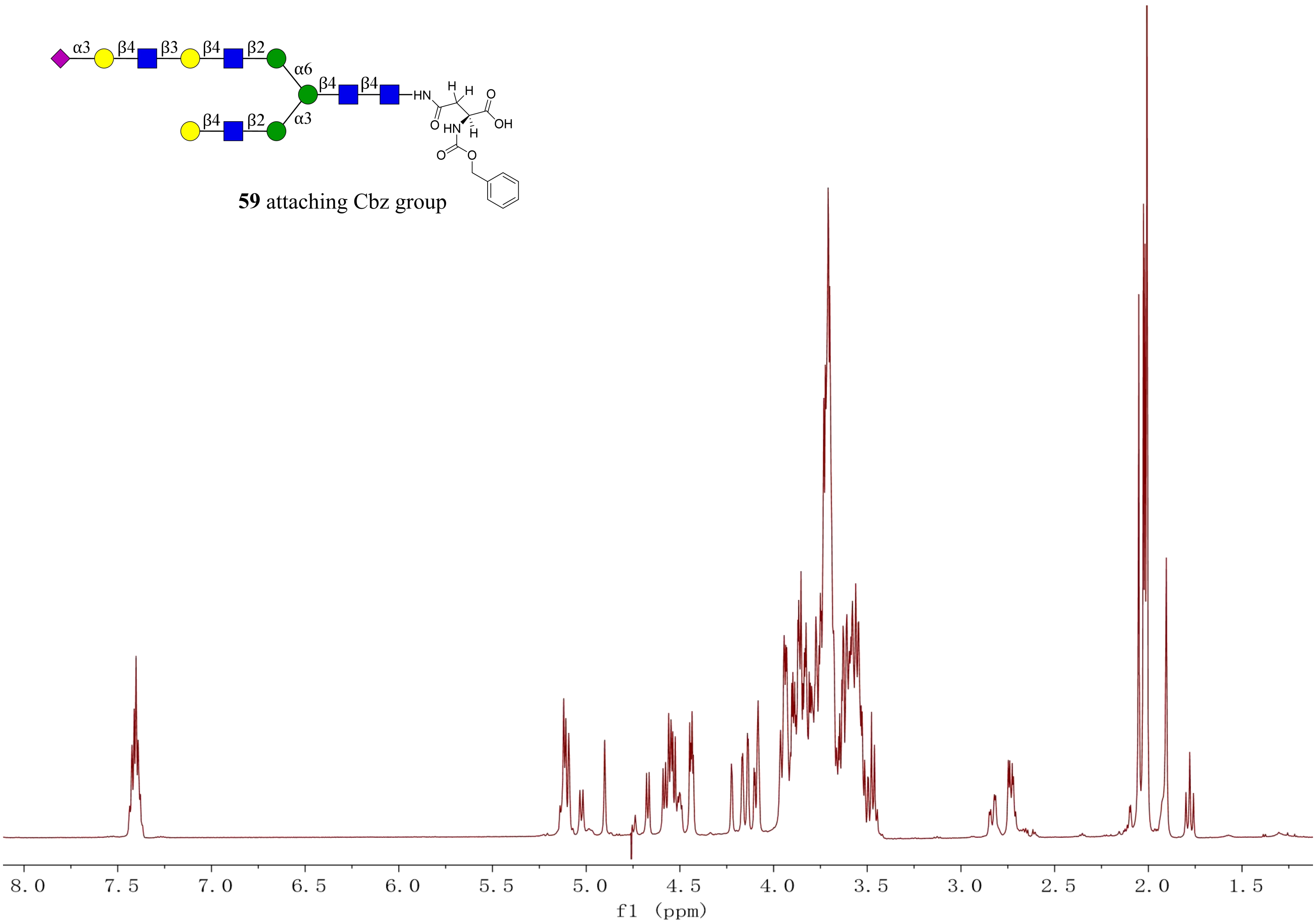


**59** attaching Cbz group

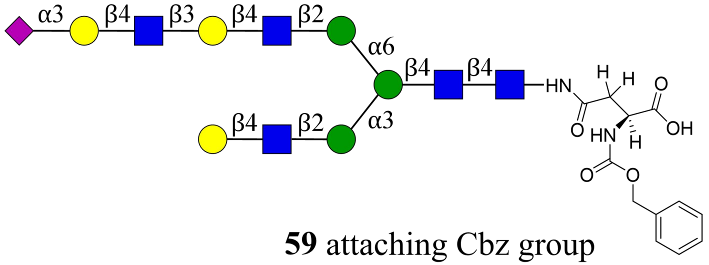


**59** attaching Cbz group

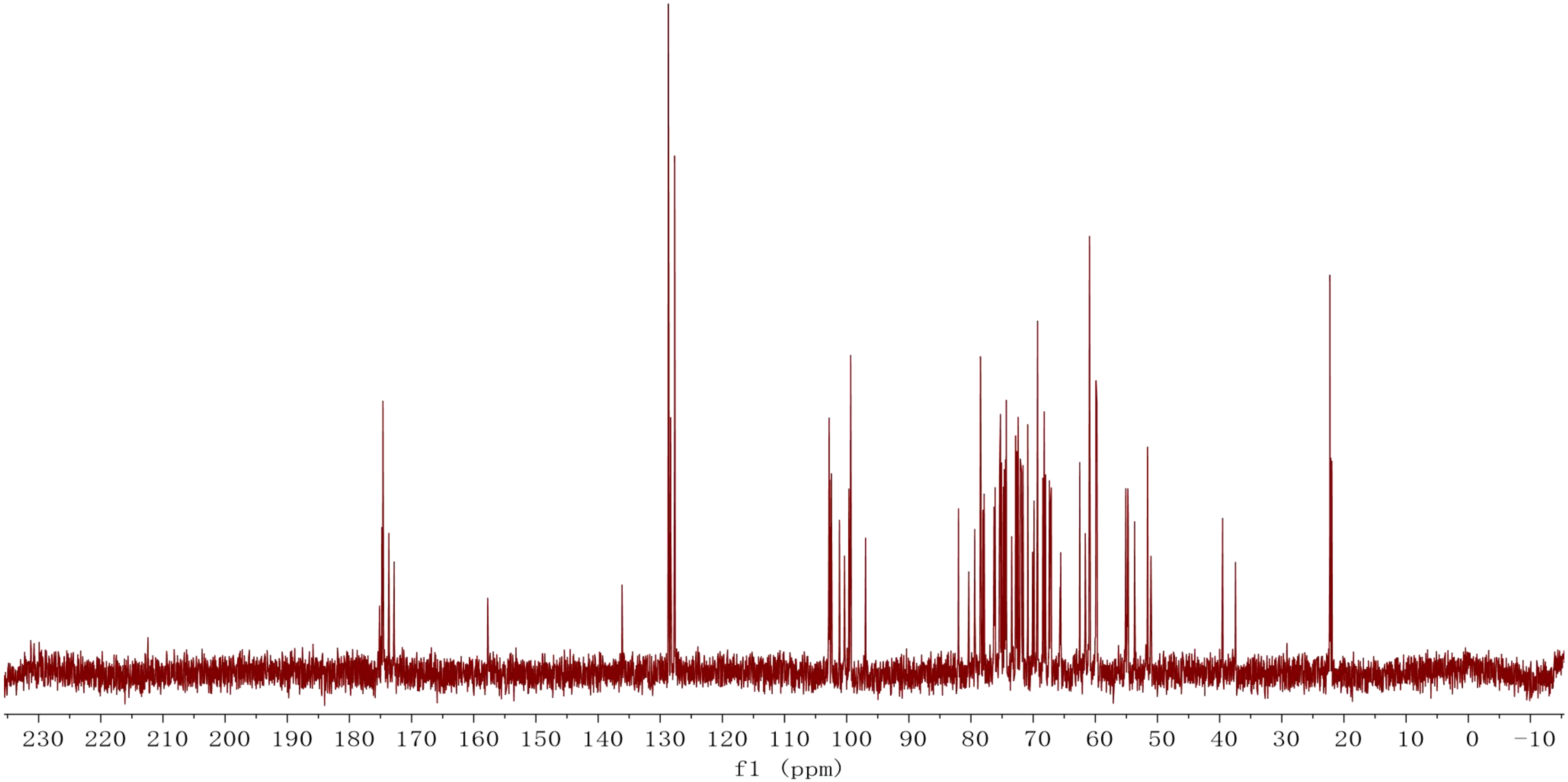

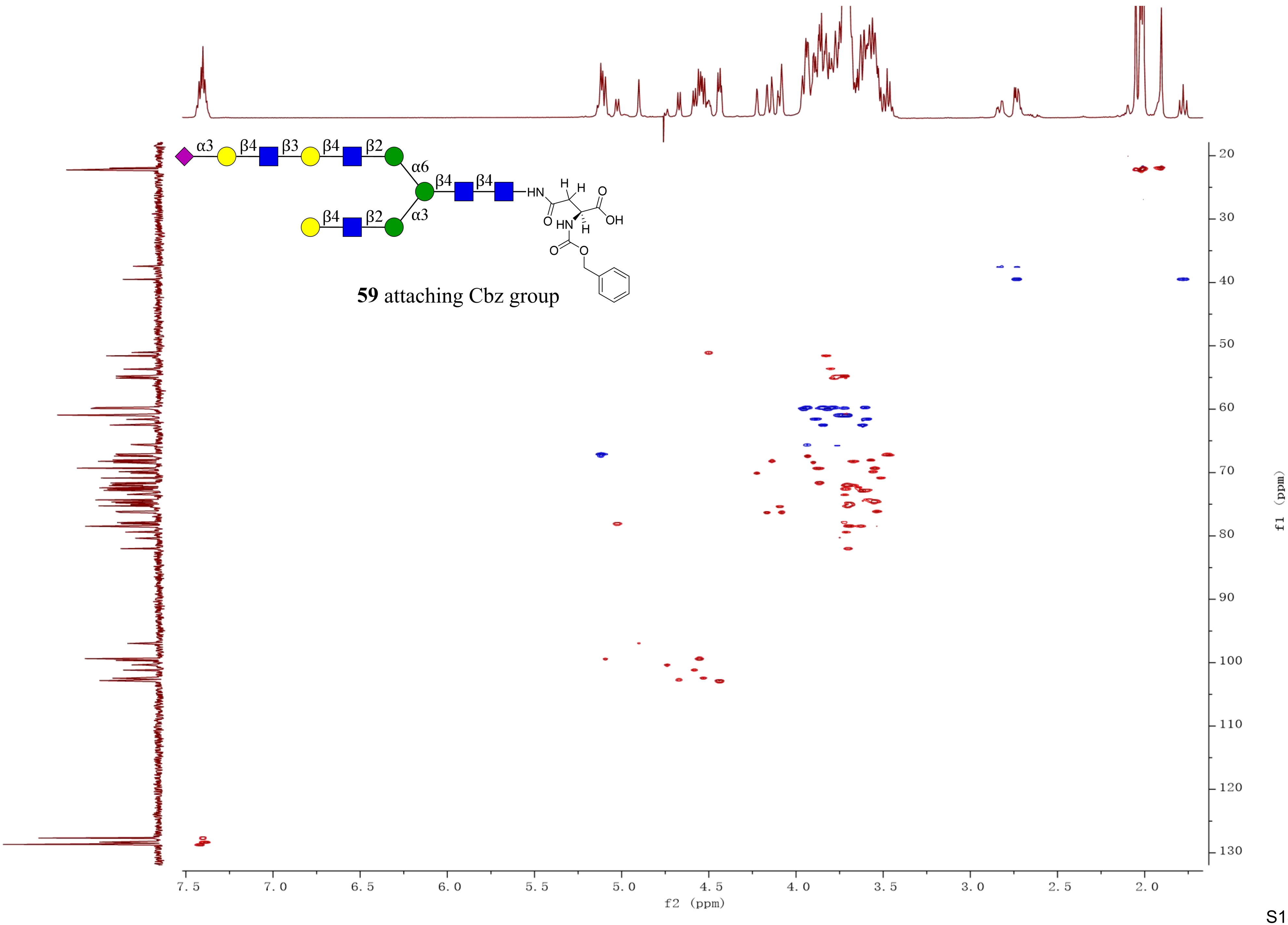
α3
β4
β3
β4
β2
α6
β4
β4
β4
β2
α3
HN
H
H
O
OH
O
HN
H
O
O
59 attaching Cbz group
f2 (ppm)
f1 (ppm)

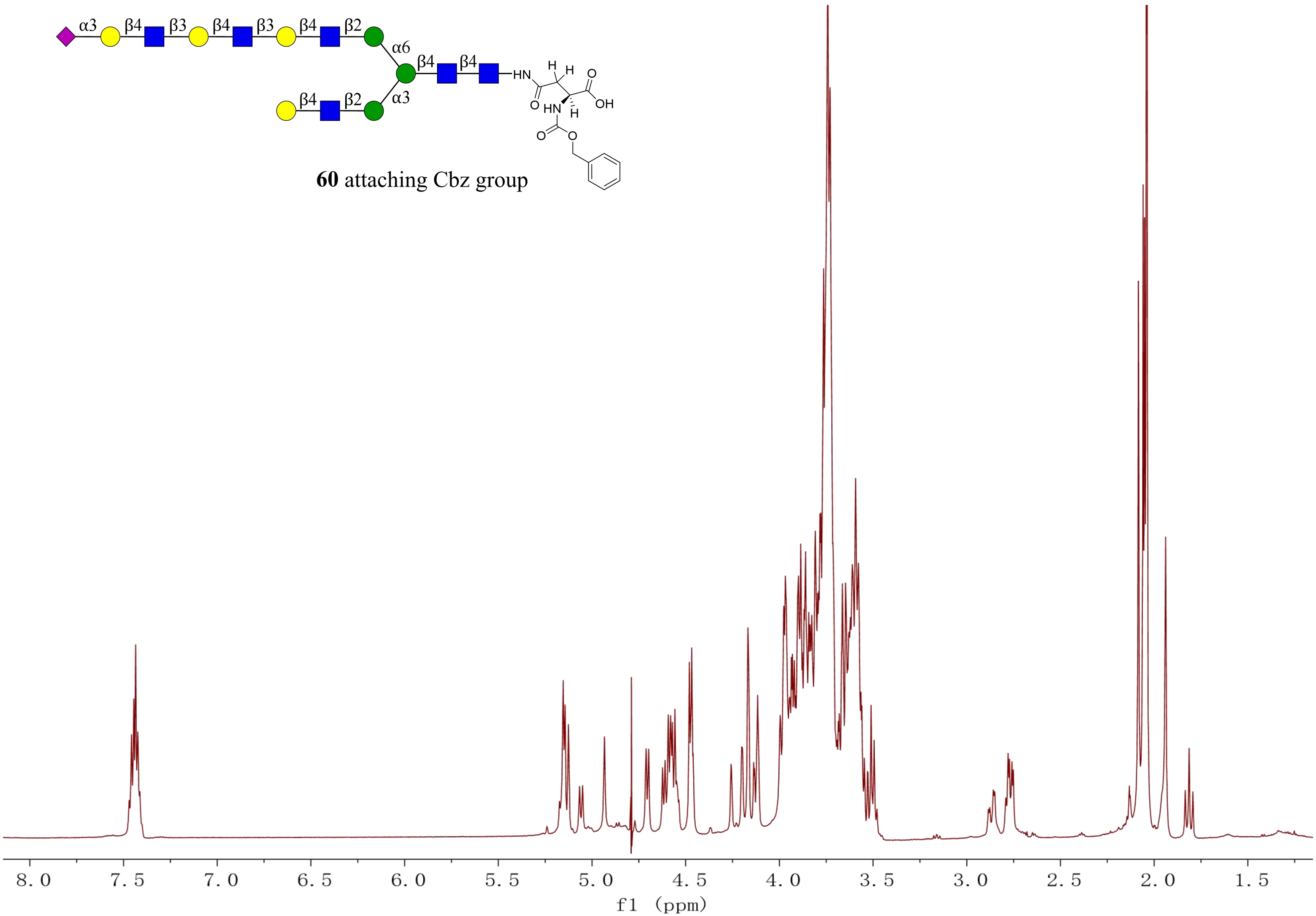


**60** attaching Cbz group

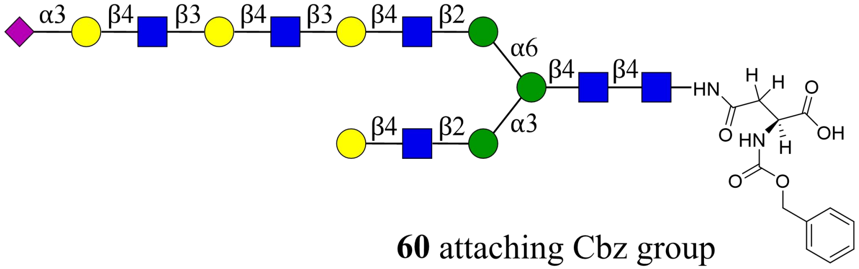


**60** attaching Cbz group

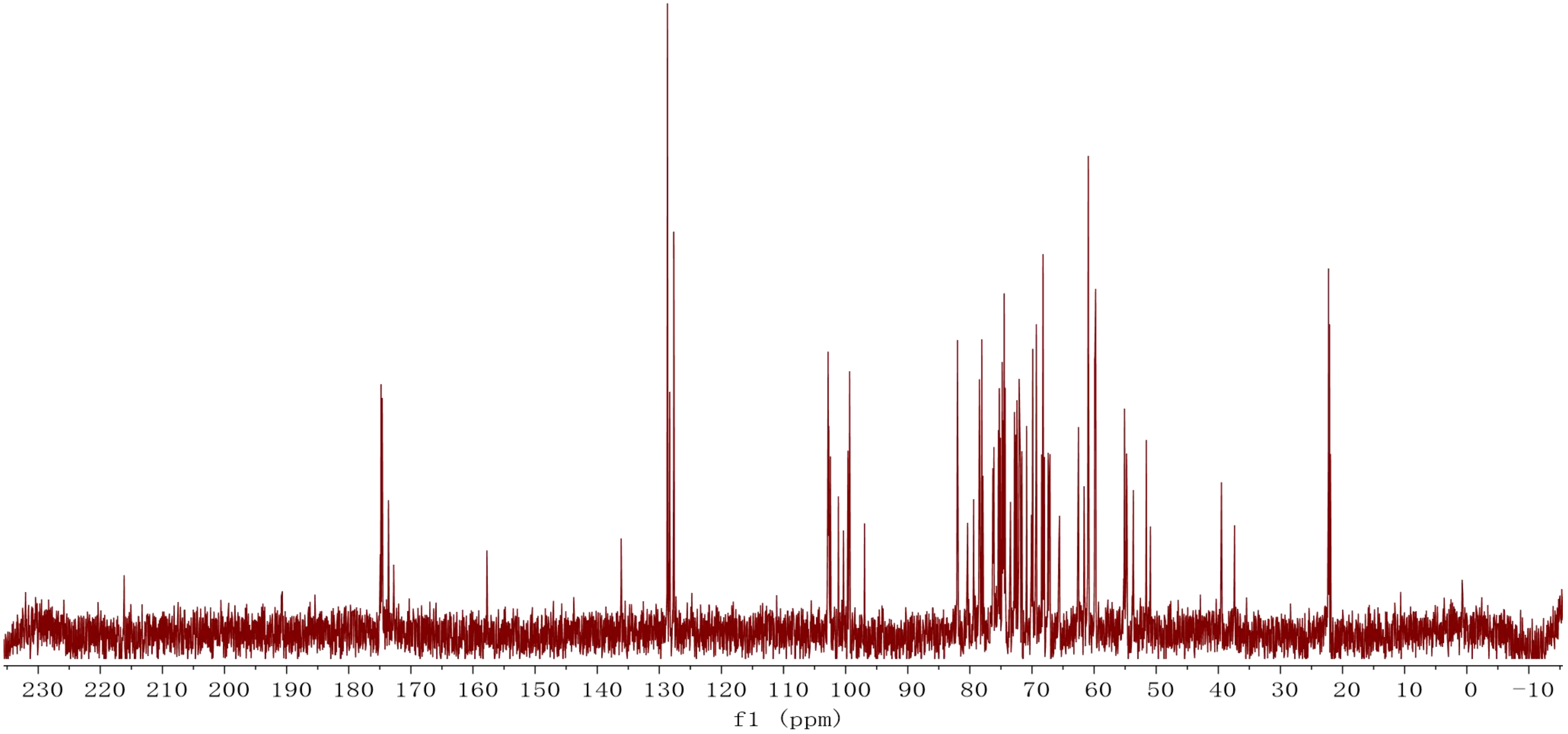

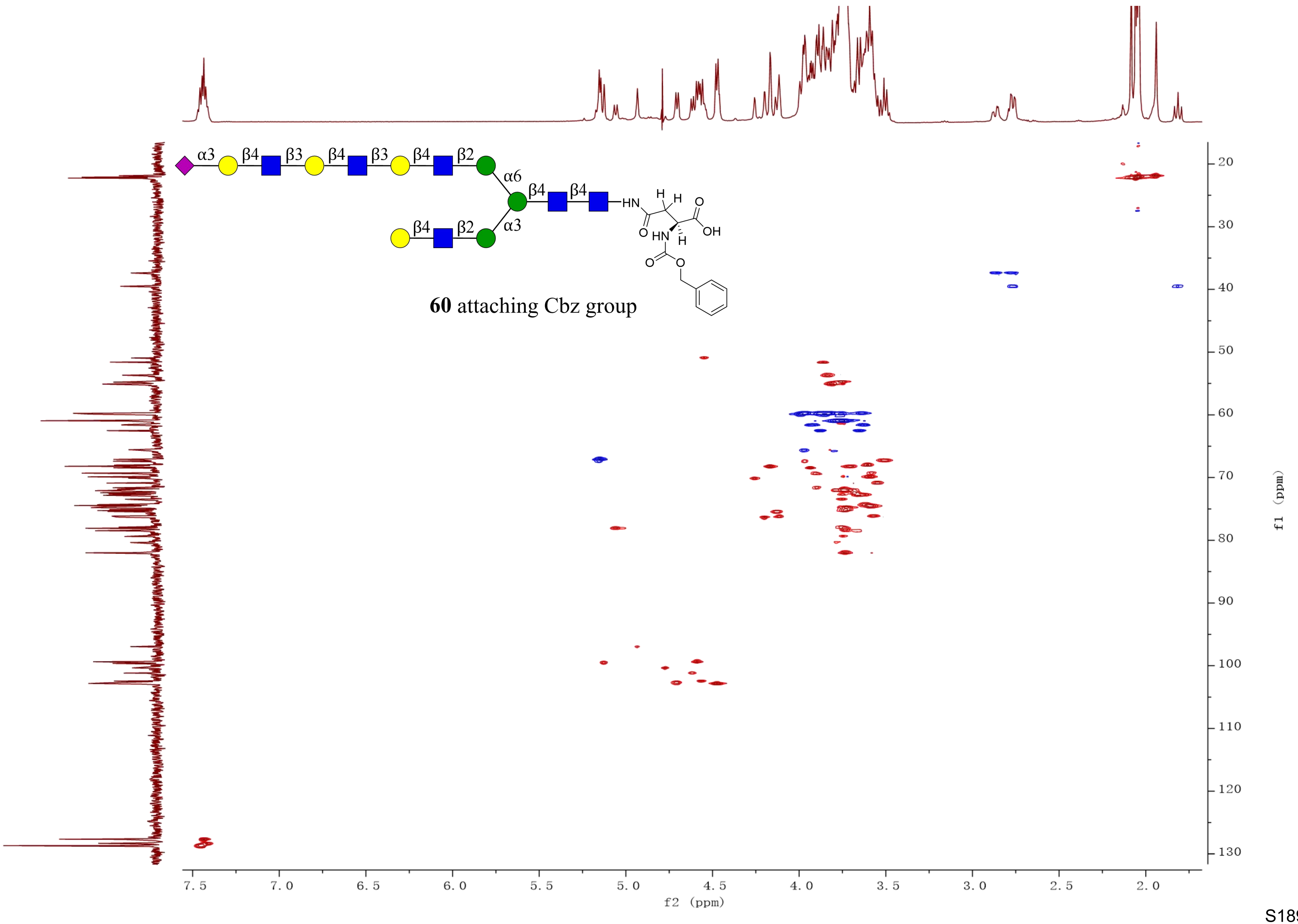
α3
β4
β3
β4
β3
β4
β2
α6
β4
β4
HN
H
H
O
OH
O
HN
H
O
O
β4
β2
α3
**60** attaching Cbz group
20
30
40
50
60
70
80
90
100
110
120
130
f1 (ppm)
7.5
7.0
6.5
6.0
5.5
5.0
4.5
4.0
3.5
3.0
2.5
2.0
f2 (ppm)

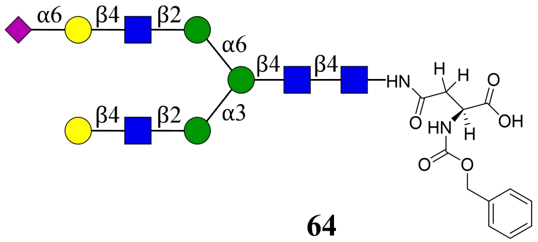
α6
β4
β2
α6
β4
β4
HN
H
H
O
OH
O
HN
H
O
O
β4
β2
α3

**64**

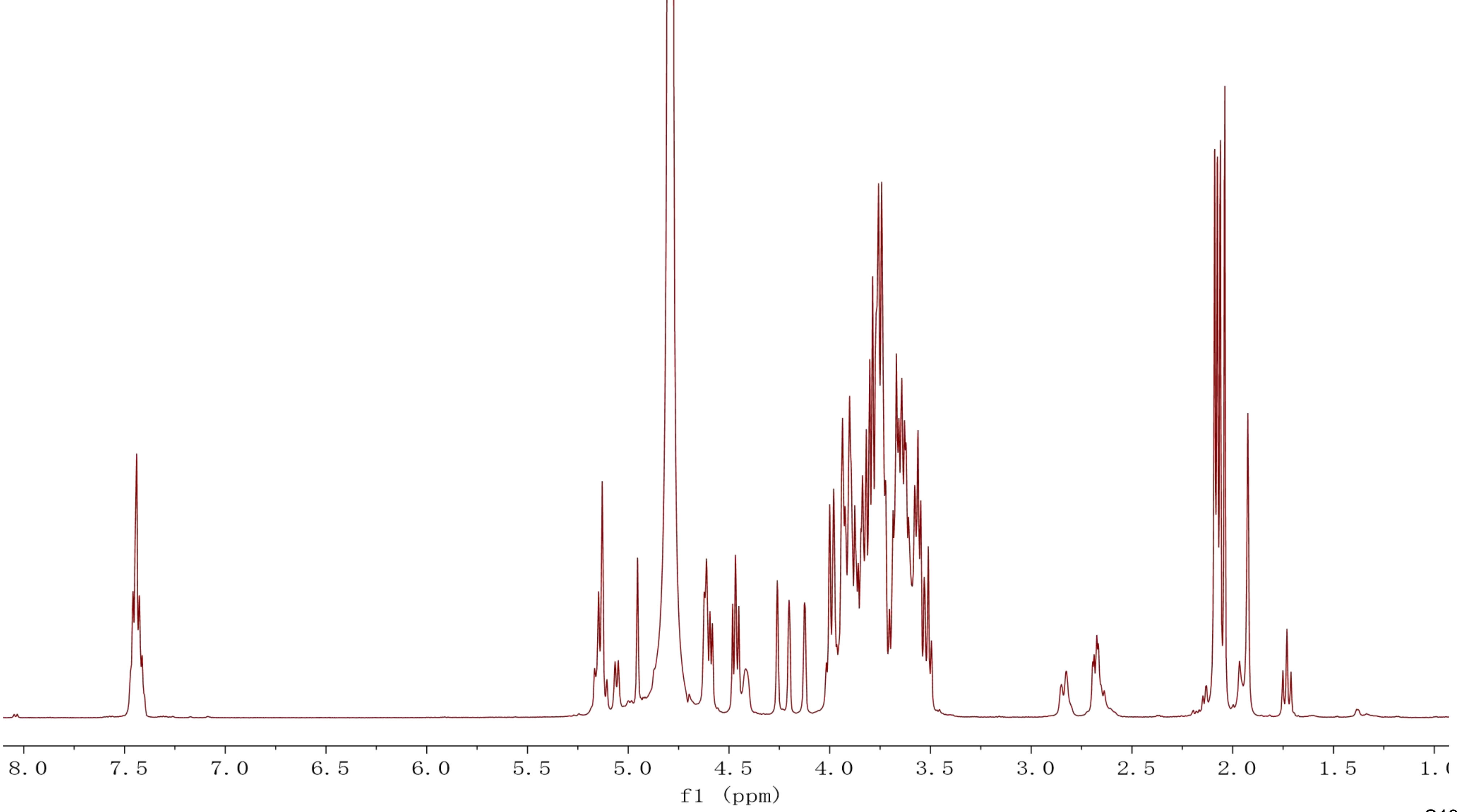
8.0
7.5
7.0
6.5
6.0
5.5
5.0
4.5
4.0
3.5
3.0
2.5
2.0
1.5
f1 (ppm)

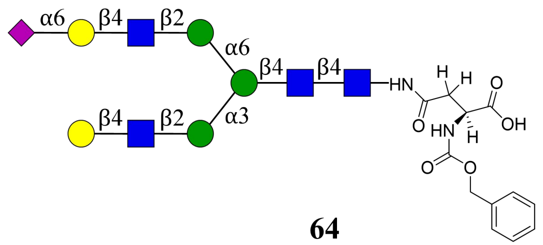

α6
β4
β2
α6
β4
β4
HN
H
H
O
O
OH
HN
H
O
O
β4
β2
α3


**64**

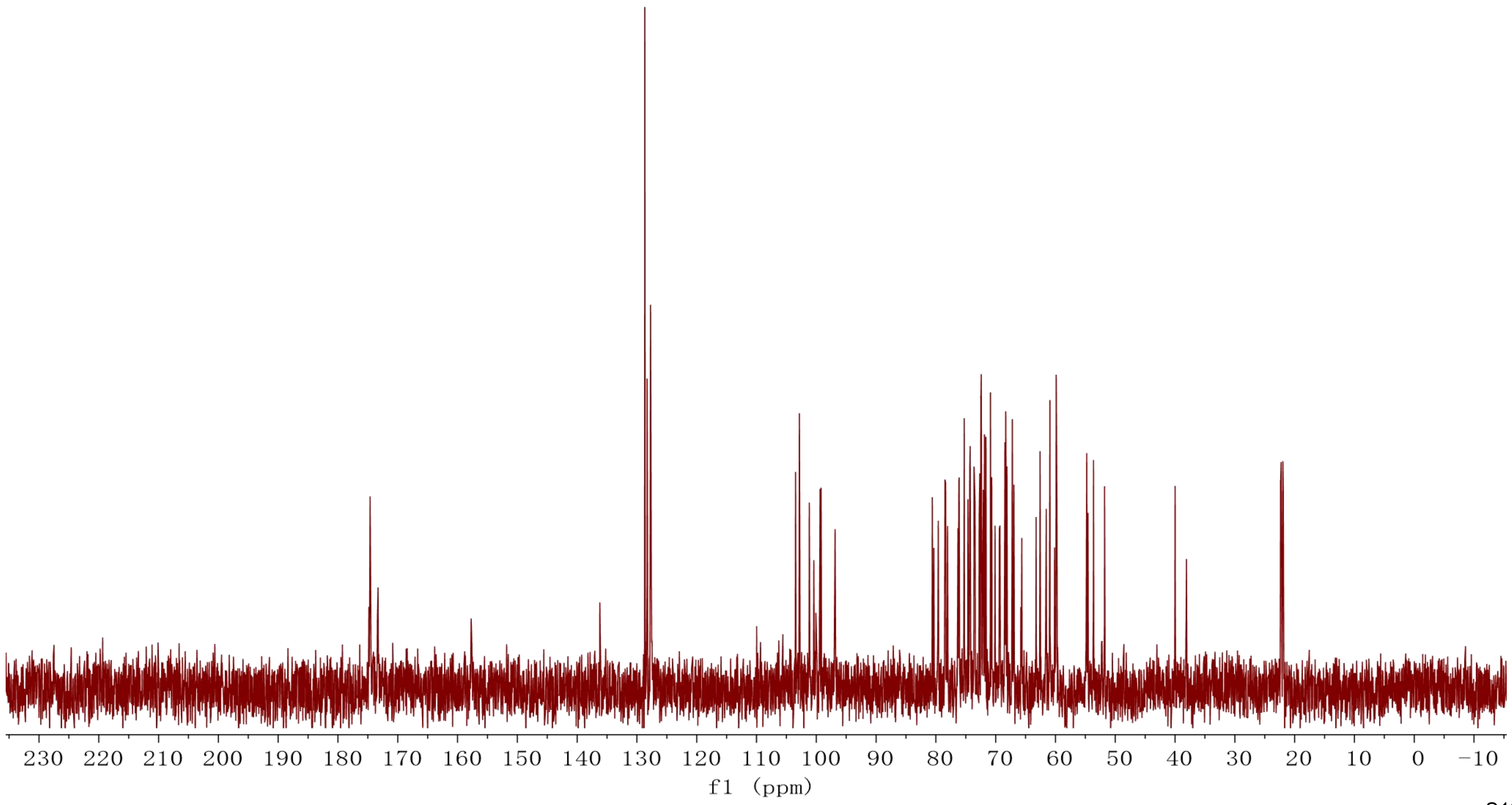

230 220 210 200 190 180 170 160 150 140 130 120 110 100 90 80 70 60 50 40 30 20 10 0 −10
f1 (ppm)

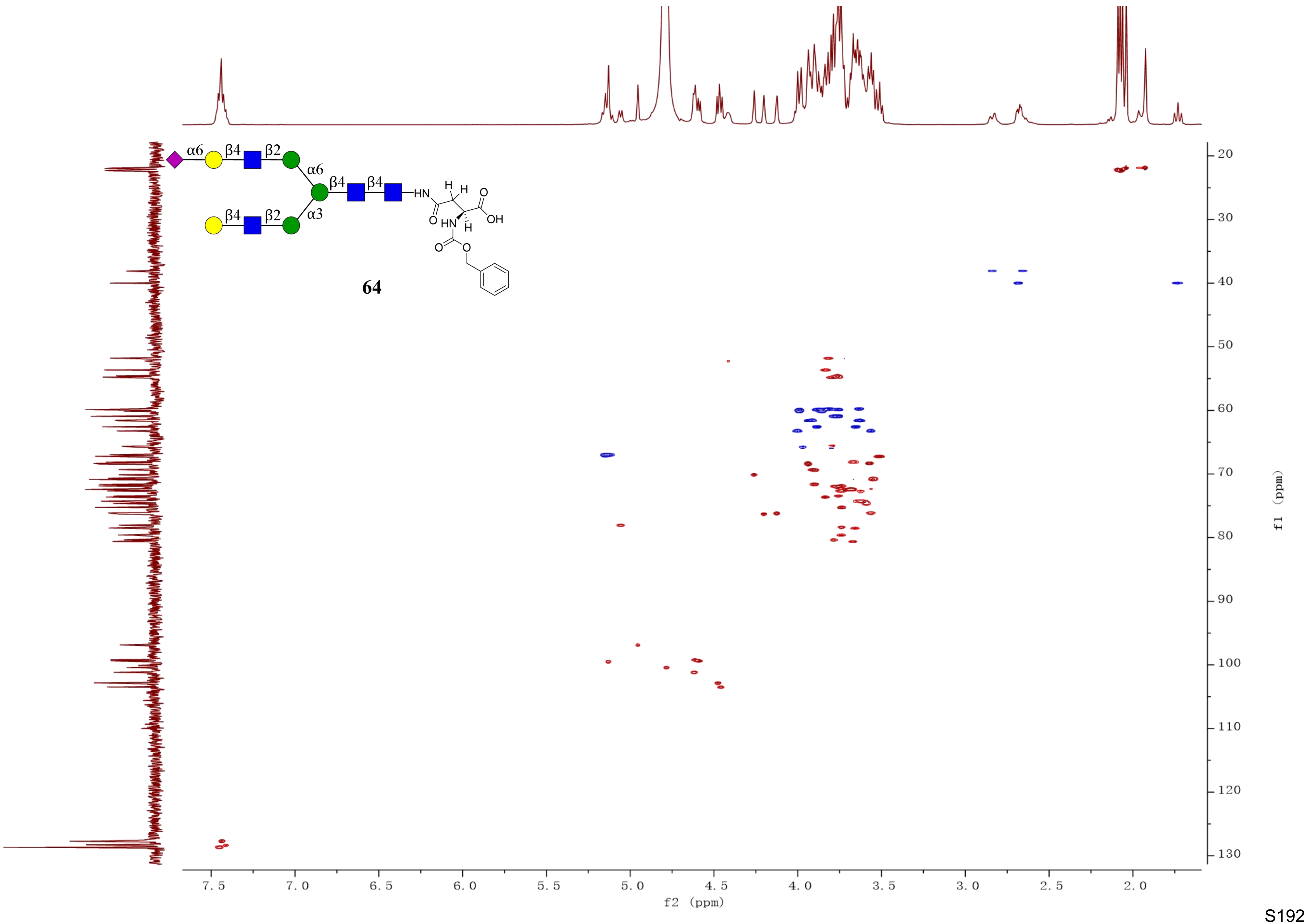

α6
β4
β2
α6
β4
β4
HN
H
H
O
OH
O
HN
H
O
O
β4
β2
α3
64
f2 (ppm)
f1 (ppm)

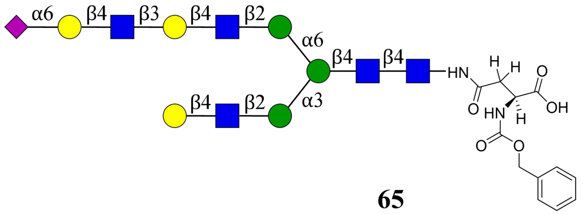
α6
β4
β3
β4
β2
α6
β4
β4
HN
H
H
O
O
OH
HN
H
O
O
β4
β2
α3

**65**

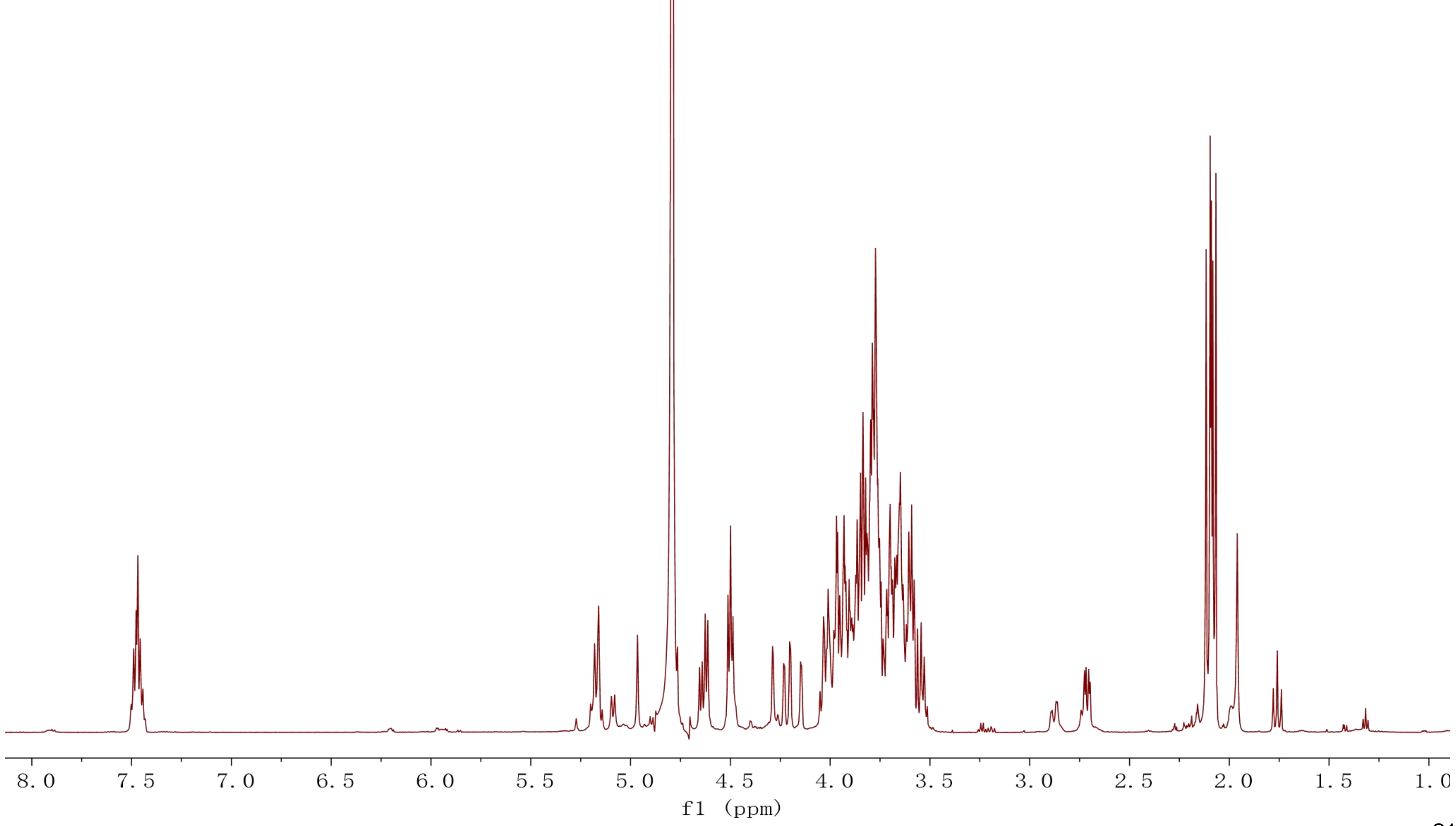
8.0
7.5
7.0
6.5
6.0
5.5
5.0
4.5
4.0
3.5
3.0
2.5
2.0
1.5
1.0
f1 (ppm)

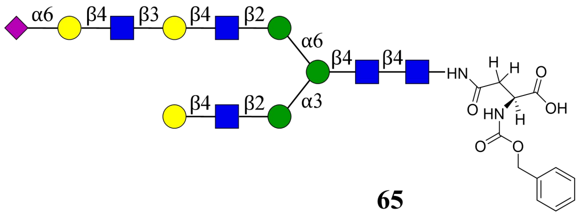

α6
β4
β3
β4
β2
α6
β4
β4
α3
HN
H
H
O
OH
O
HN
H
O
O


**65**

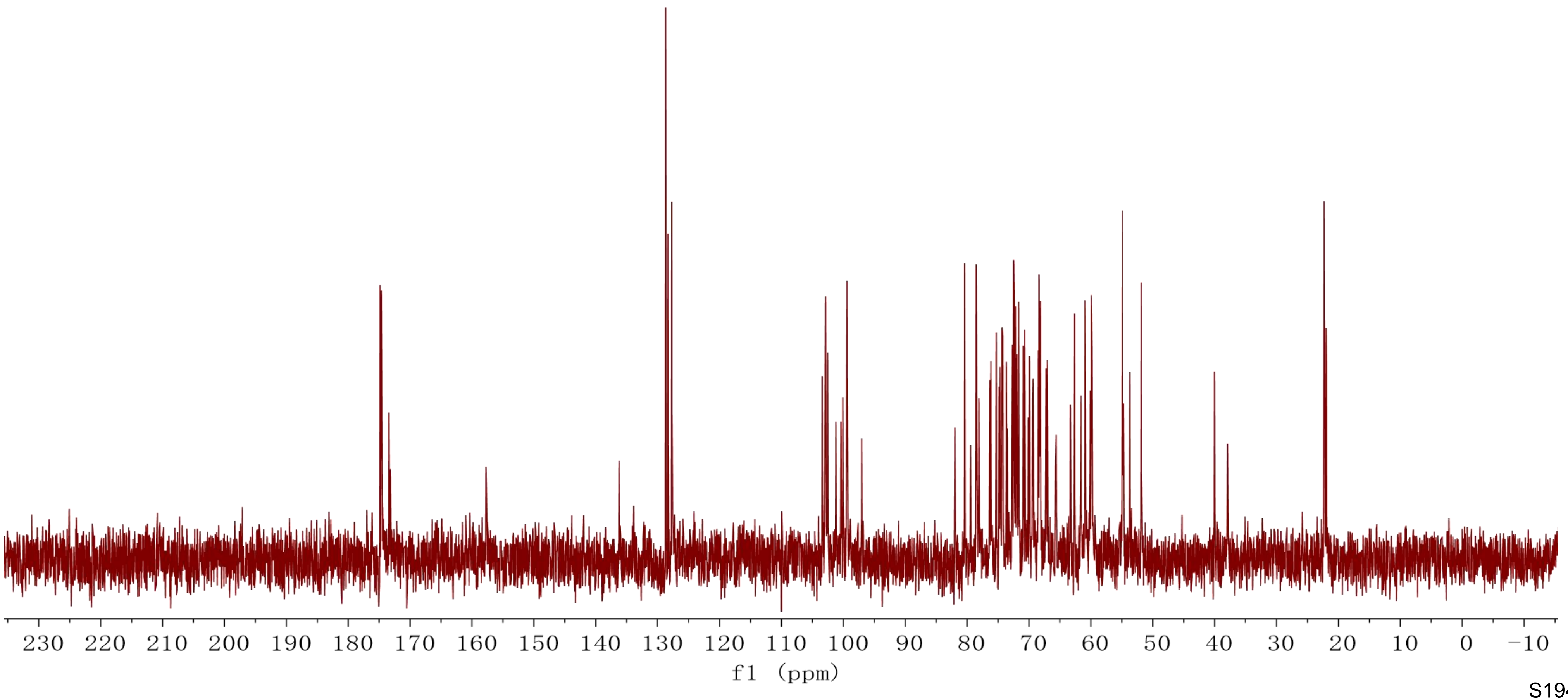

230 220 210 200 190 180 170 160 150 140 130 120 110 100 90 80 70 60 50 40 30 20 10 0 −10
f1 (ppm)

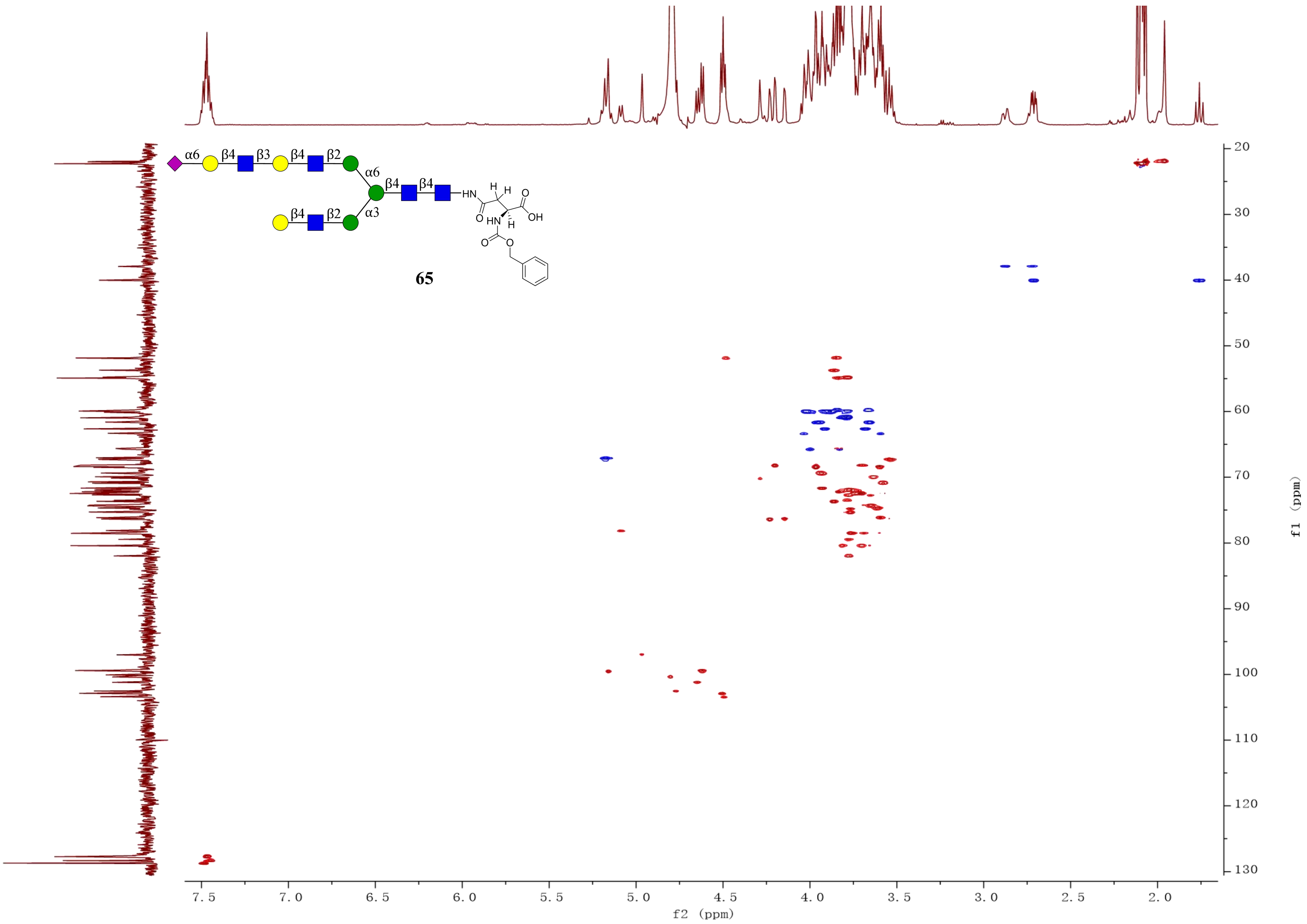
65
f2 (ppm)
f1 (ppm)

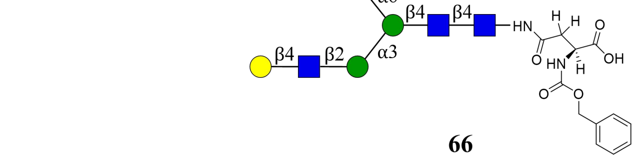
α6
β4
β3
β4
β3
β4
β2
α6
β4
β4
HN
α3
β4
β2
OH

**66**

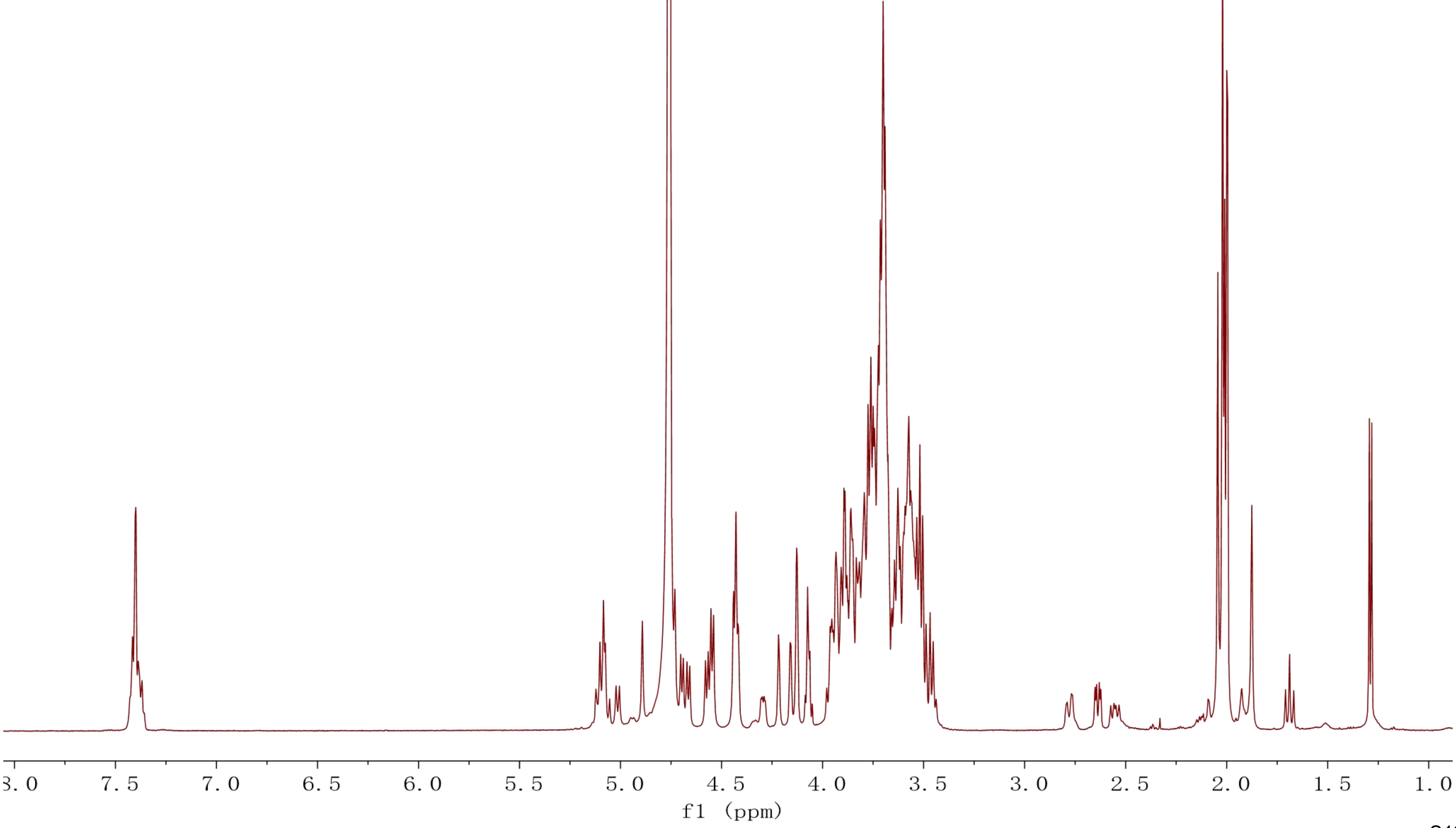
8.0 7.5 7.0 6.5 6.0 5.5 5.0 4.5 4.0 3.5 3.0 2.5 2.0 1.5 1.0
f1 (ppm)

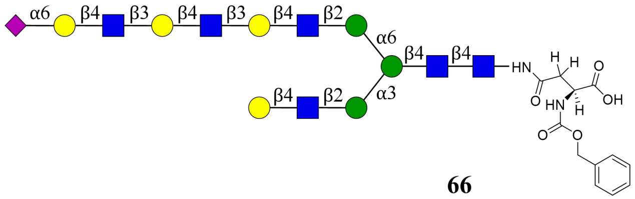
α6
β4
β3
β4
β3
β4
β2
α6
β4
β4
HN
H
H
O
O
HN
H
OH
O
O
β4
β2
α3

**66**

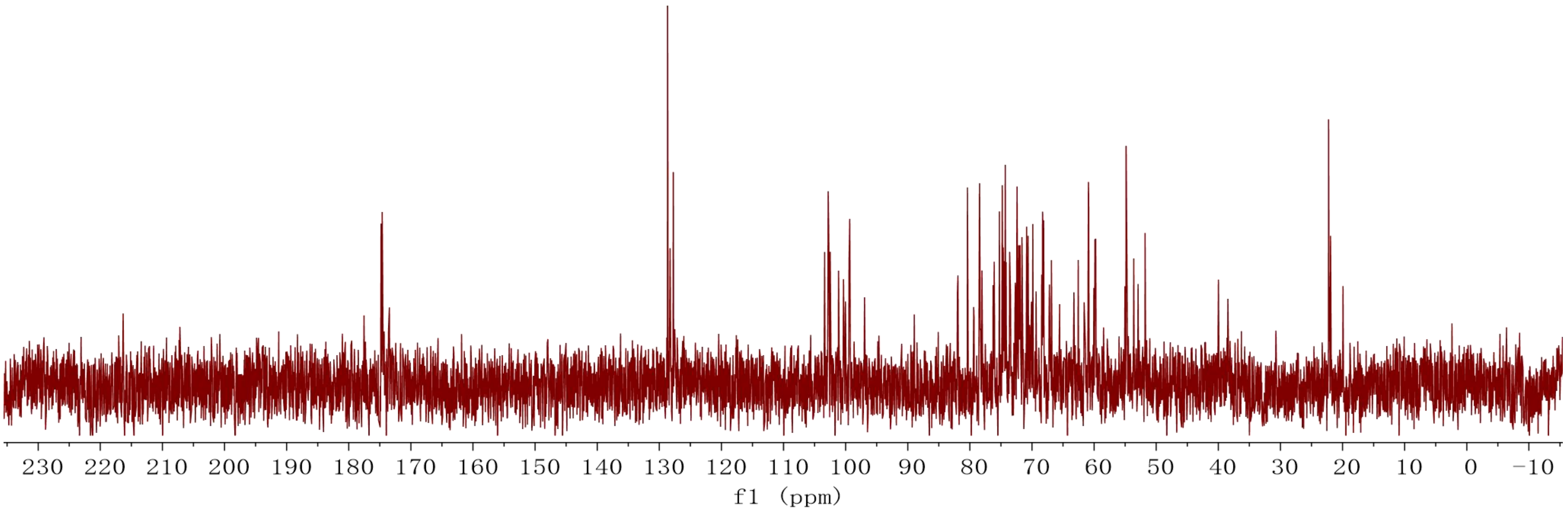
230 220 210 200 190 180 170 160 150 140 130 120 110 100 90 80 70 60 50 40 30 20 10 0 −10
f1 (ppm)

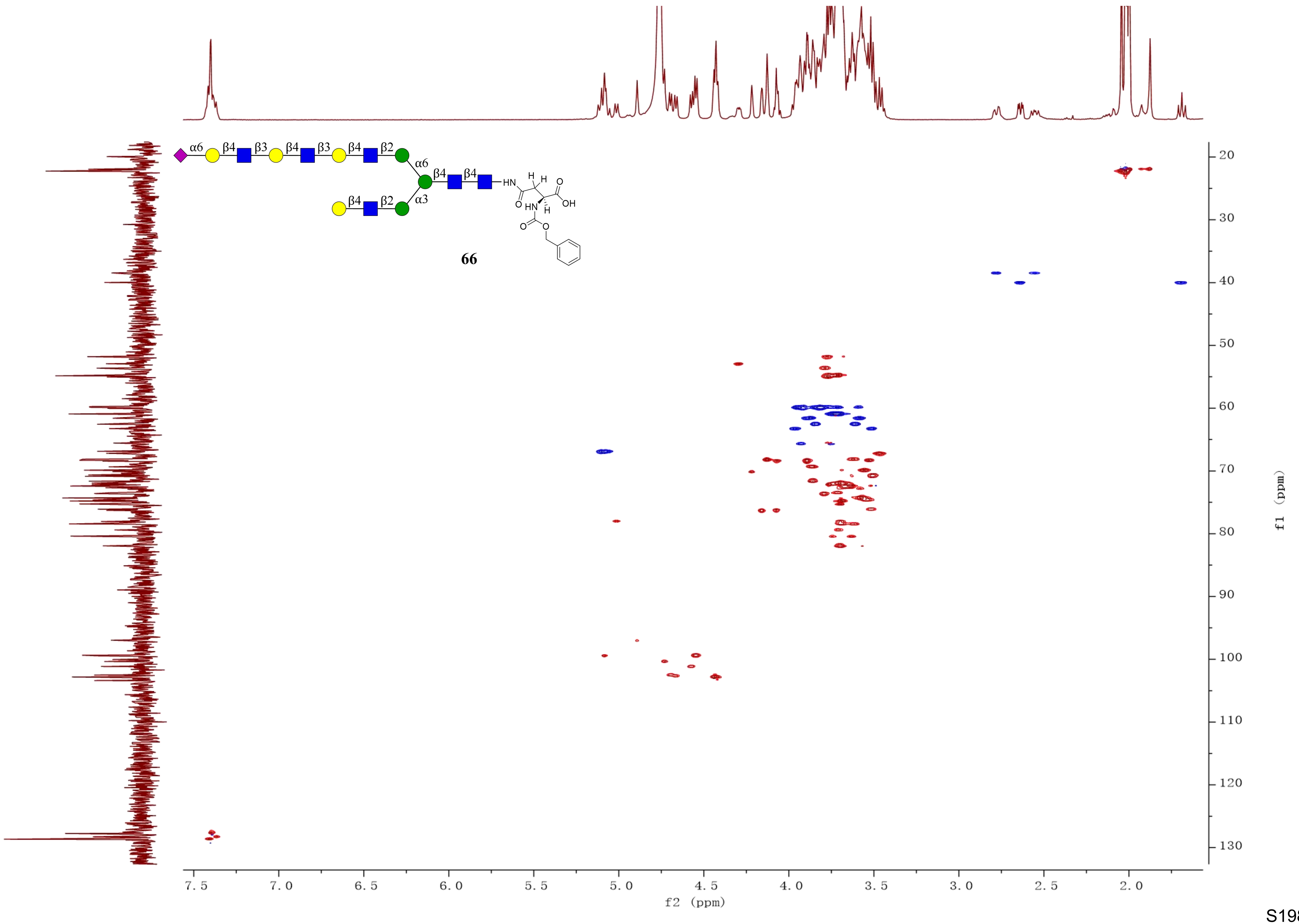

66
f2 (ppm)
f1 (ppm)

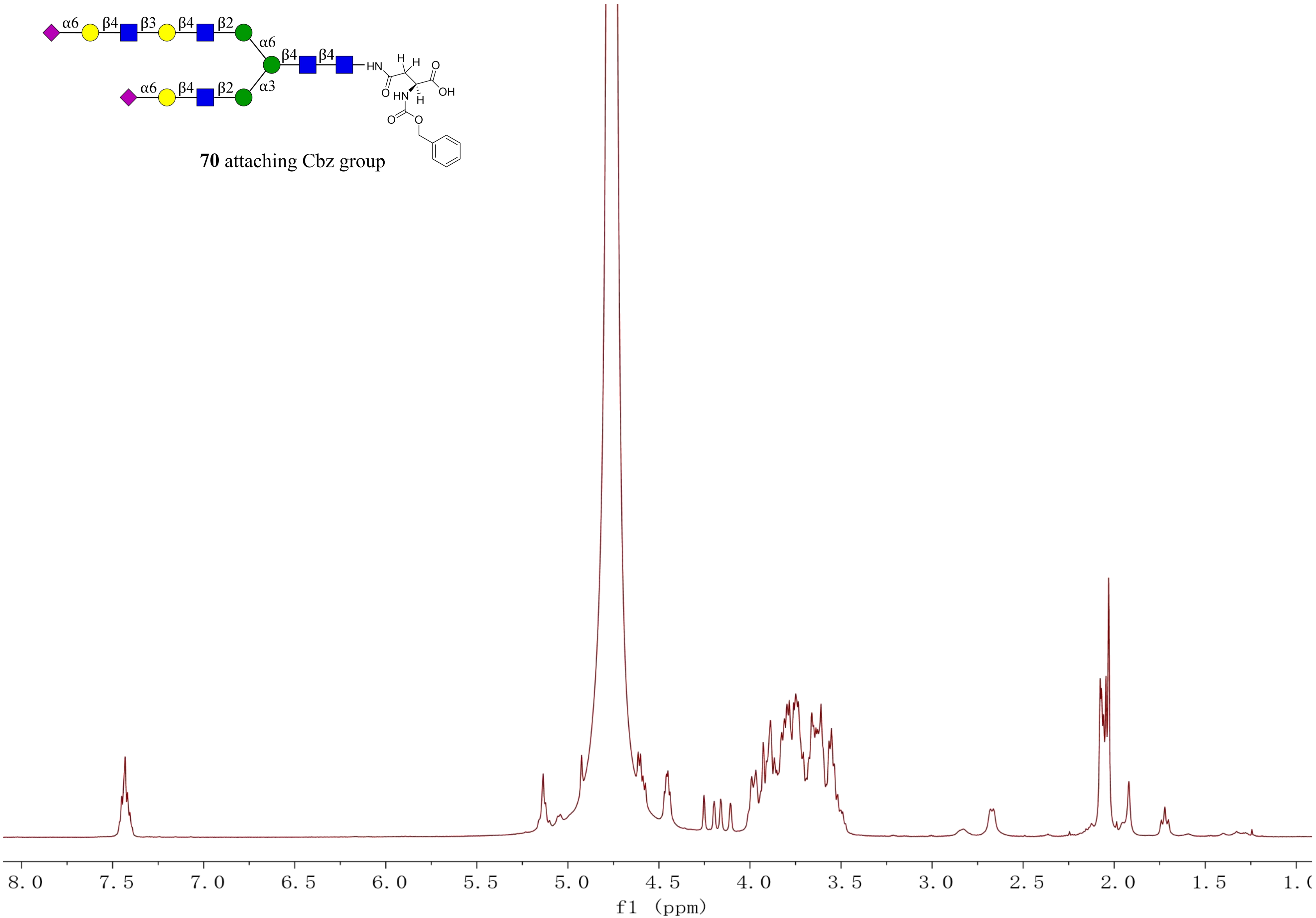
α6
β4
β3
β4
β2
α6
β4
β4
HN
α6
β4
β2
α3
O
OH
HN
H
O
O
**70** attaching Cbz group
8.0
7.5
7.0
6.5
6.0
5.5
5.0
4.5
4.0
3.5
3.0
2.5
2.0
1.5
f1 (ppm)

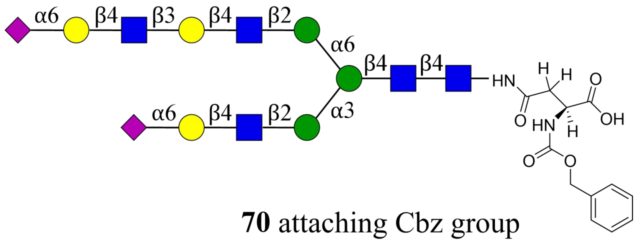

**70** attaching Cbz group

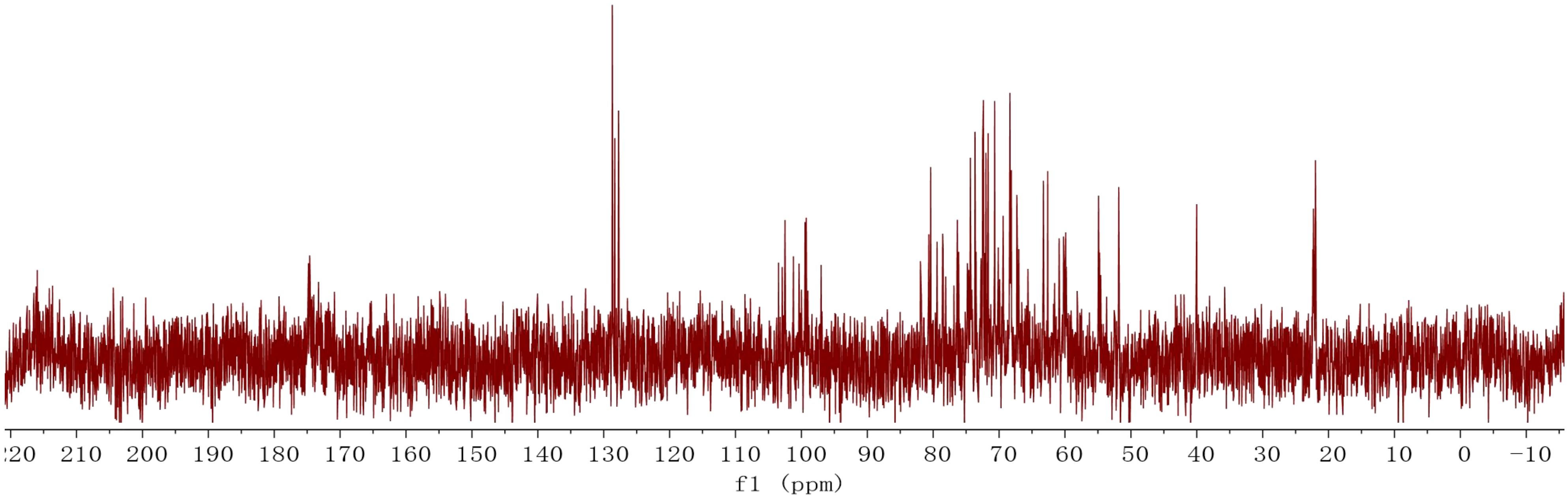

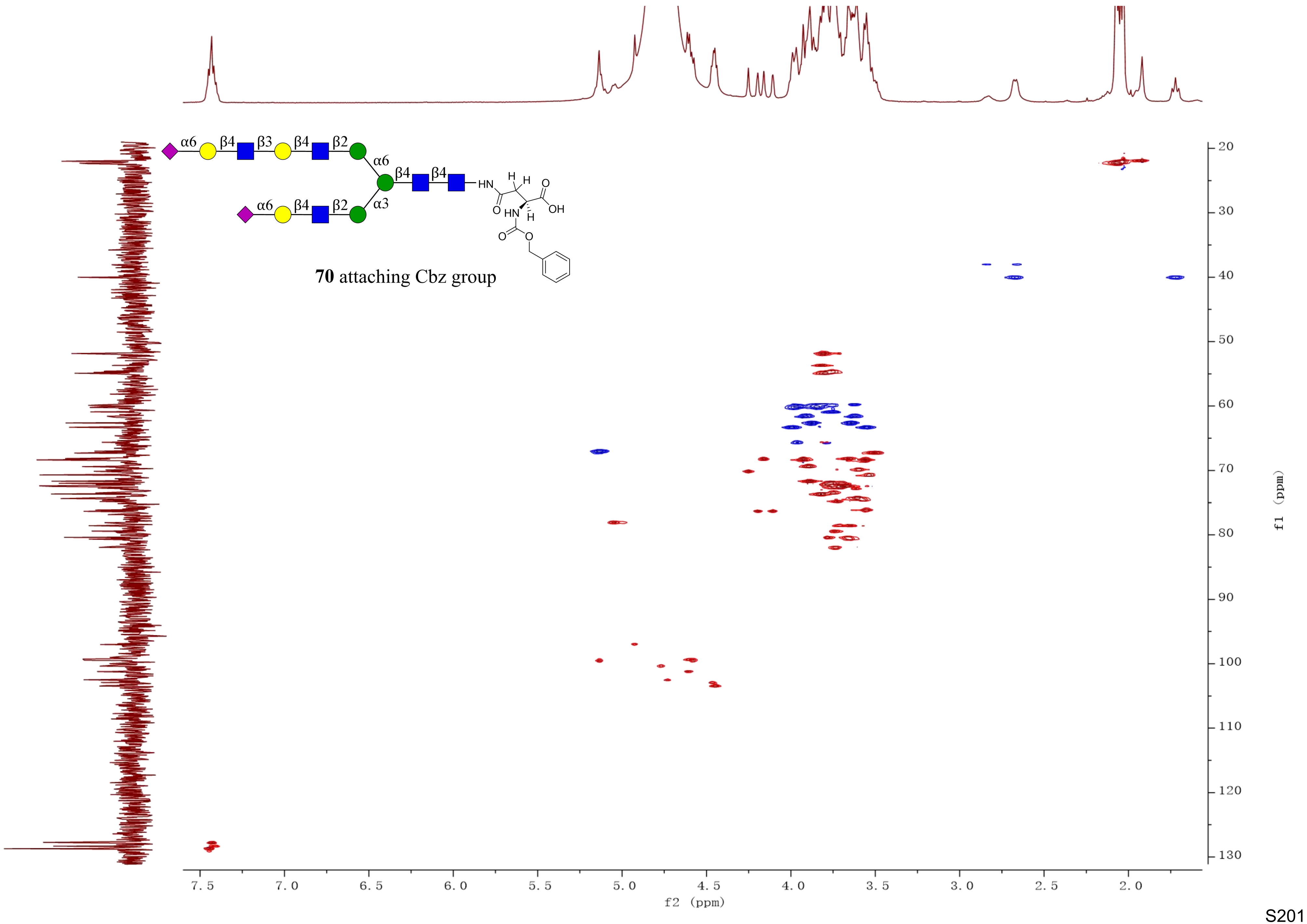

α6
β4
β3
β4
β2
α6
β4
β4
α6
β4
β2
α3
HN
H
H
O
O
OH
HN
H
O
O
70 attaching Cbz group
f2 (ppm)
f1 (ppm)

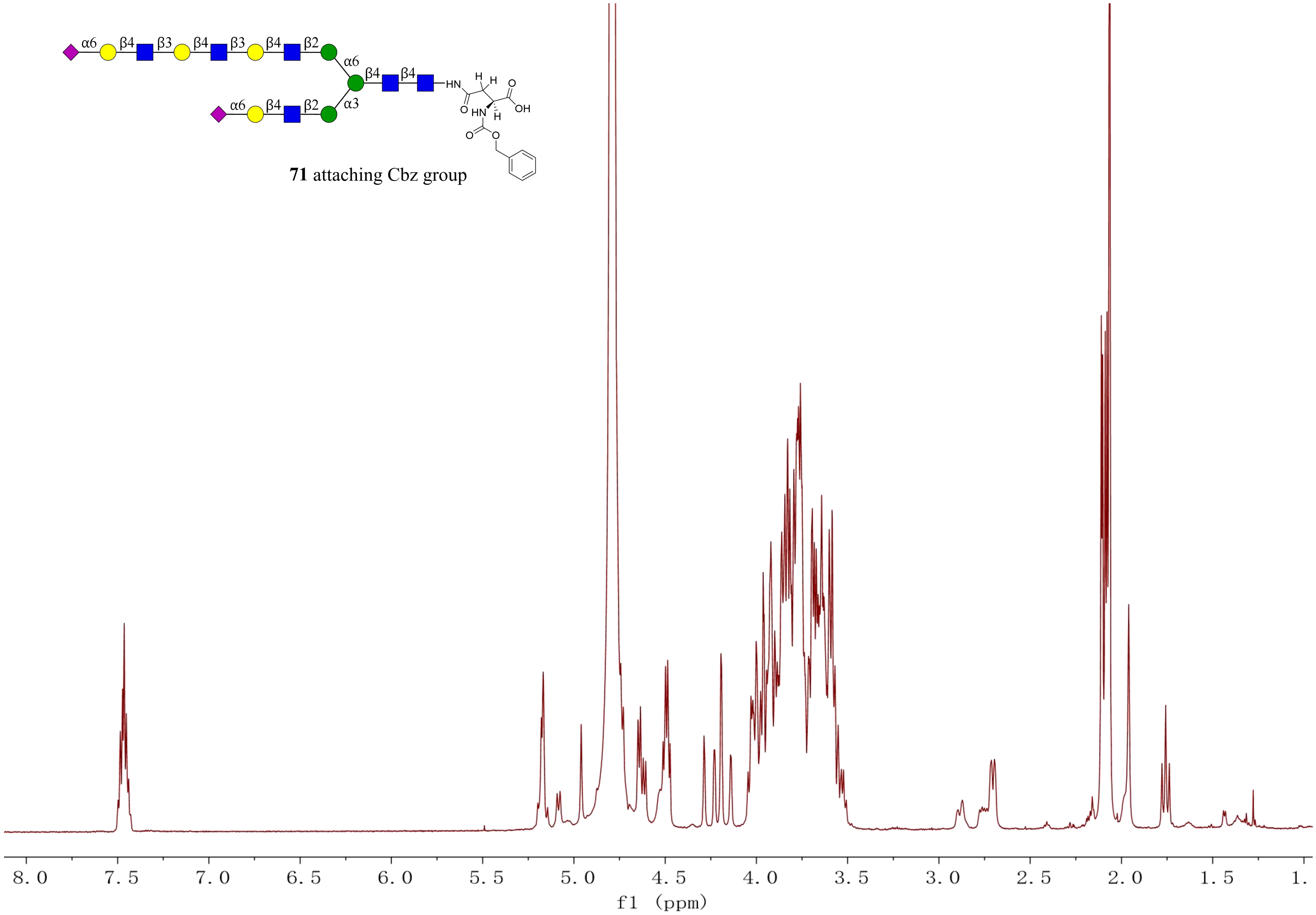


**71** attaching Cbz group

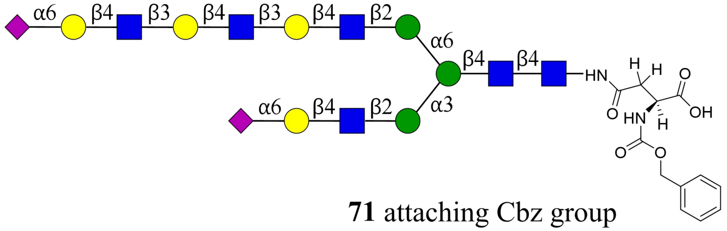

**71** attaching Cbz group

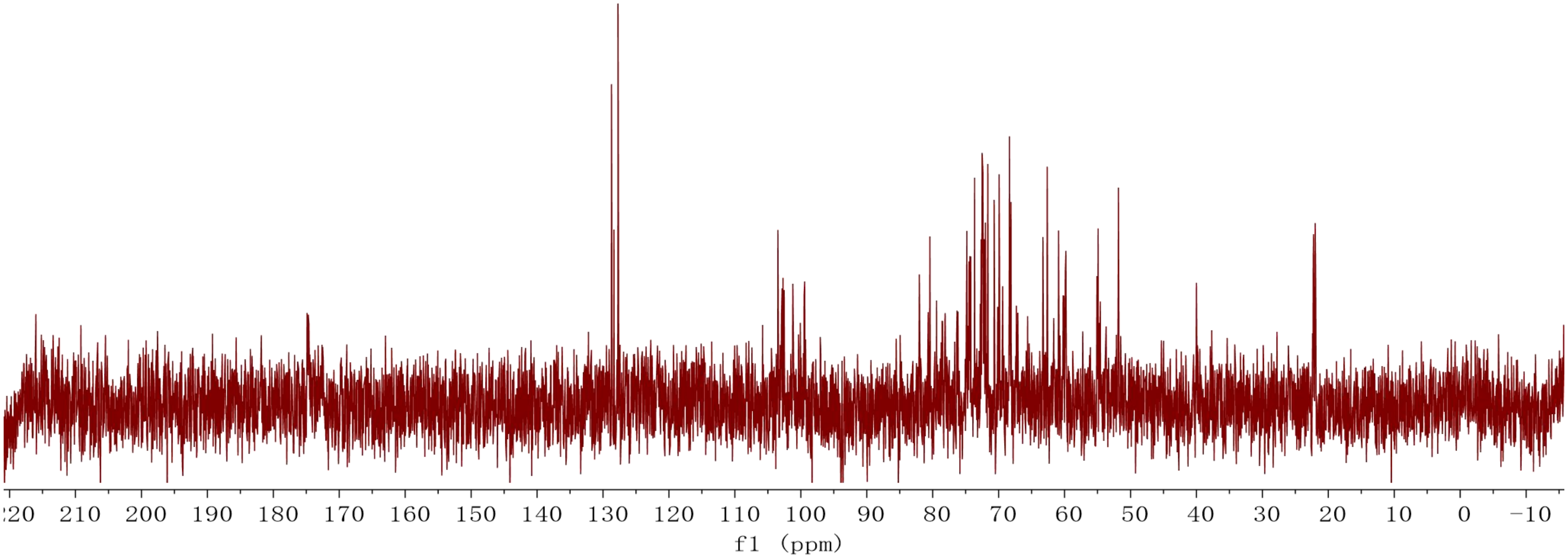

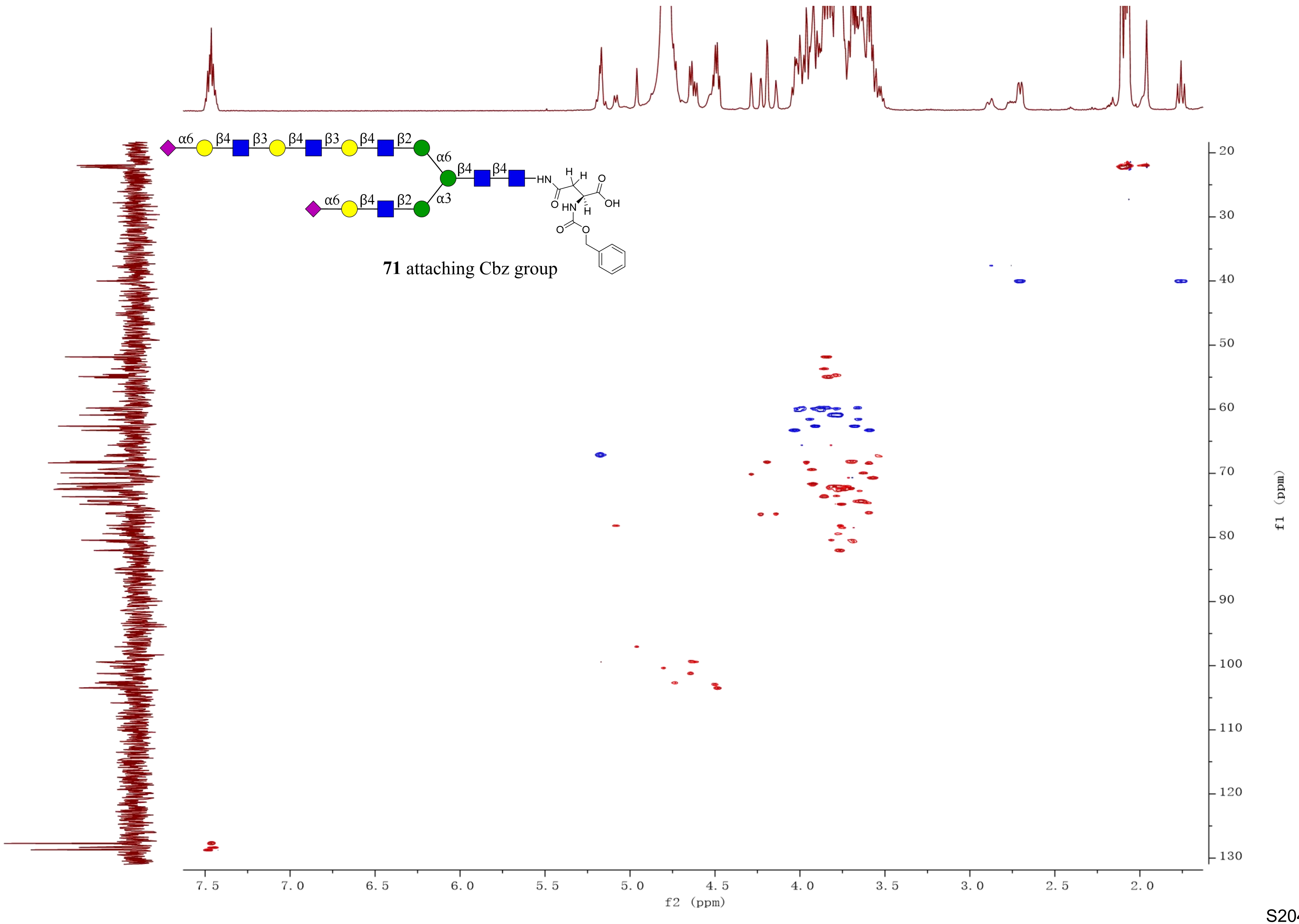
α6
β4
β3
β4
β3
β4
β2
α6
β4
β4
α6
β4
β2
α3
HN
H
H
O
O
OH
HN
H
O
O
**71** attaching Cbz group
7.5
7.0
6.5
6.0
5.5
5.0
4.5
4.0
3.5
3.0
2.5
2.0
f2 (ppm)
20
30
40
50
60
70
80
90
100
110
120
130
f1 (ppm)

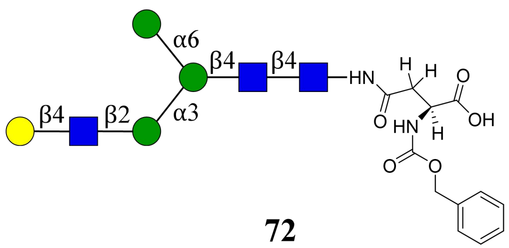


**72**

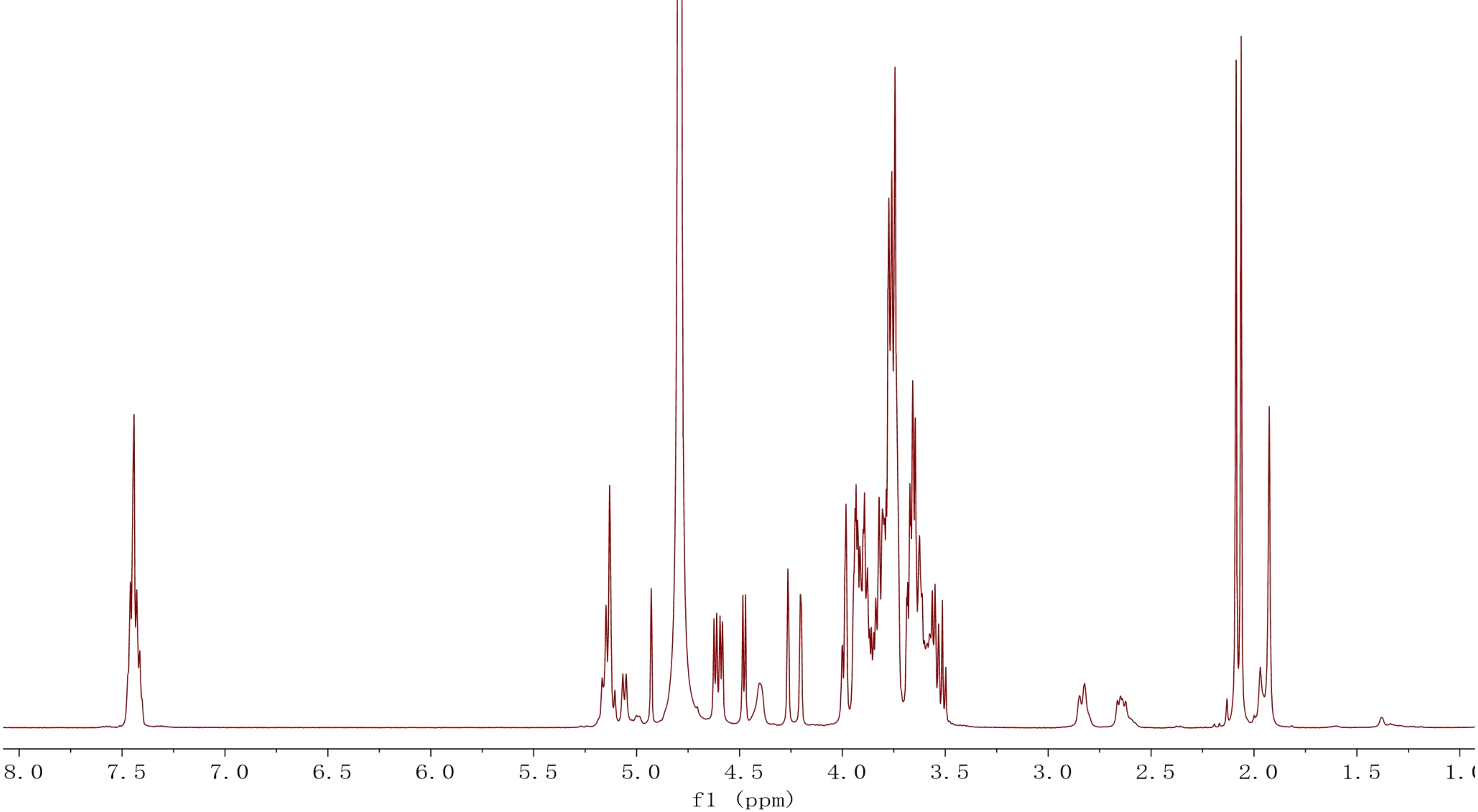

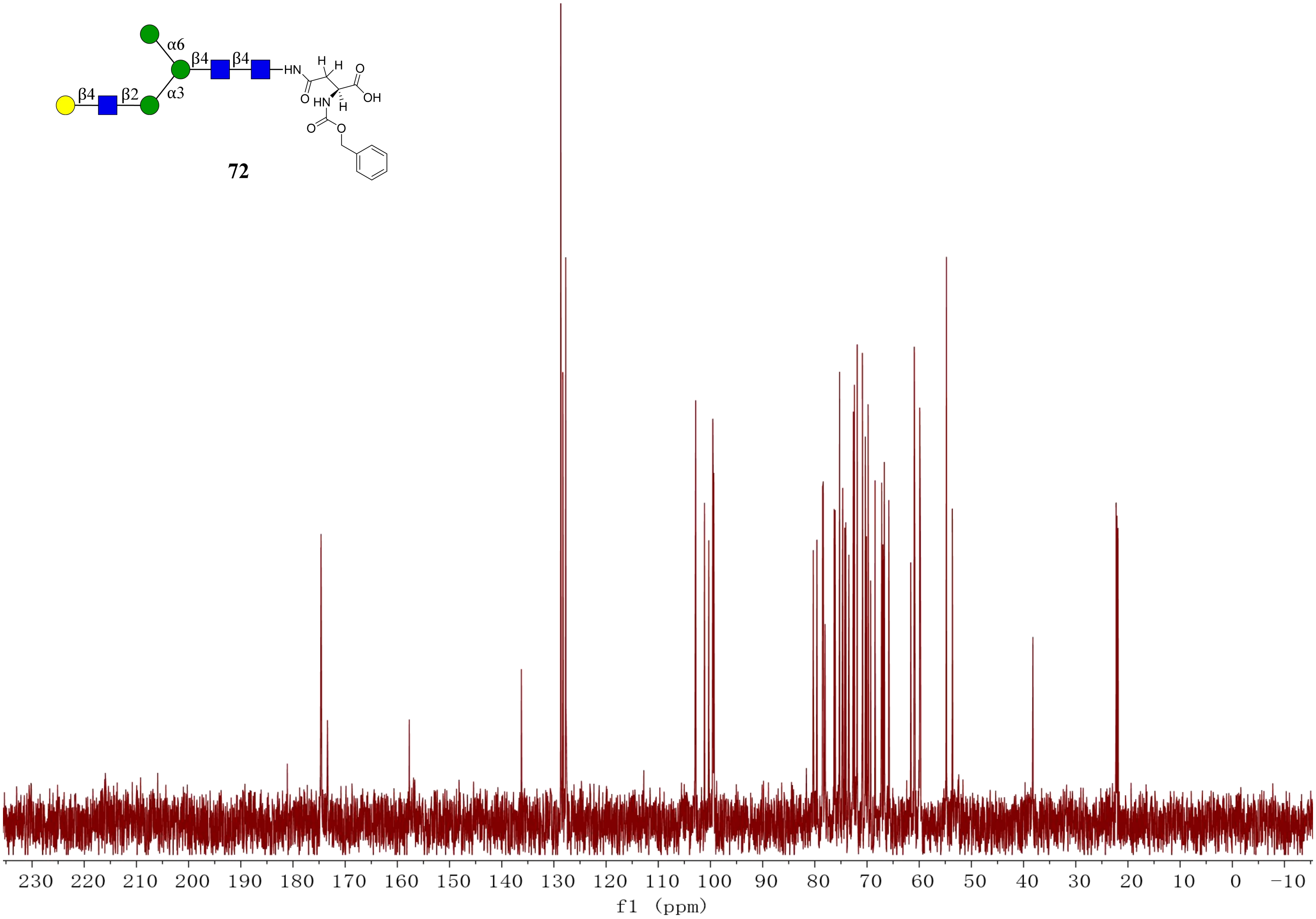
α6
β4
β4
β2
α3
HN
H
H
O
OH
O
HN
H
O
O
72
230 220 210 200 190 180 170 160 150 140 130 120 110 100 90 80 70 60 50 40 30 20 10 0 −10
f1 (ppm)

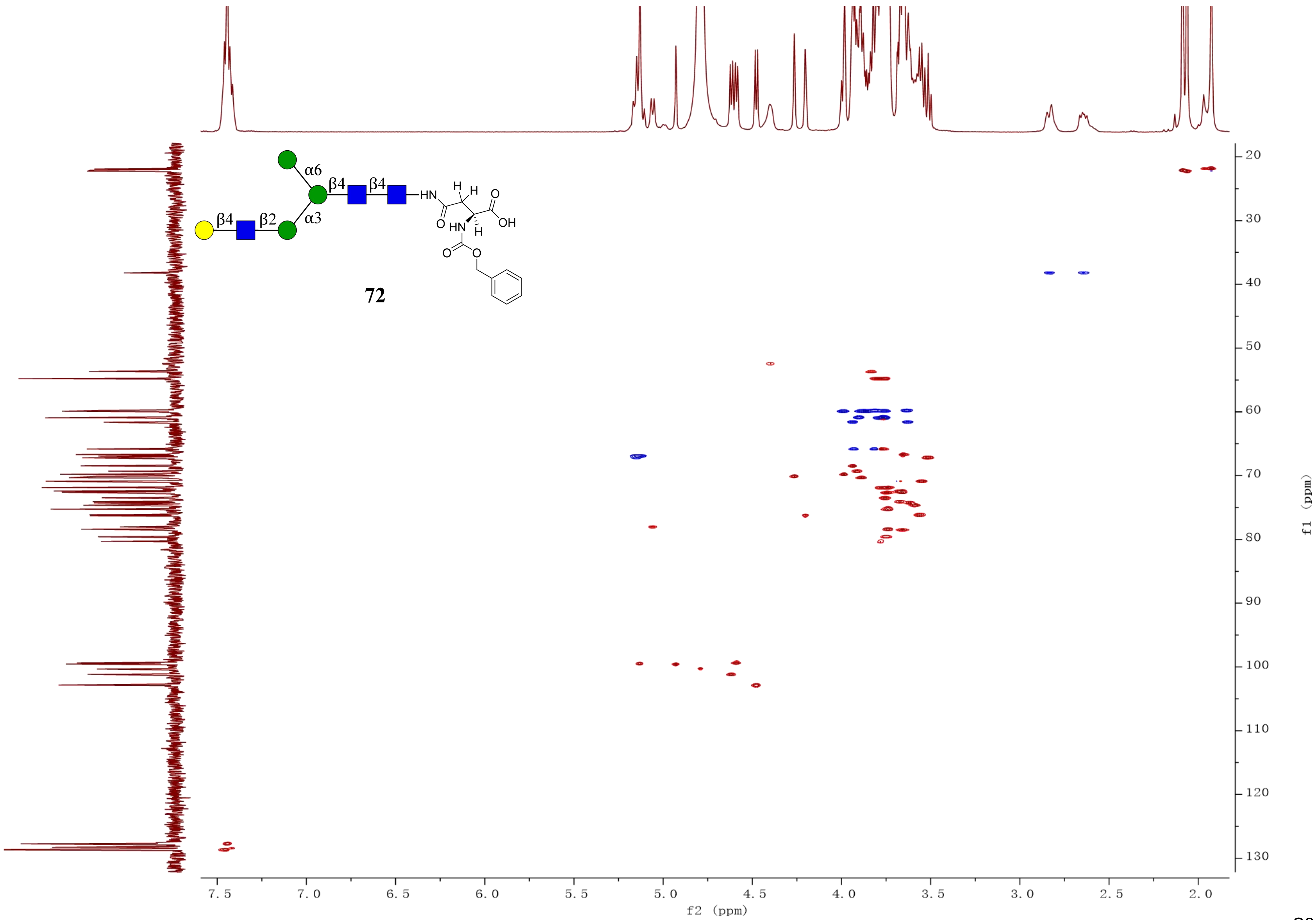
α6
β4
β4
β4
β2
α3
HN
H
H
O
O
OH
HN
H
O
O
72
20
30
40
50
60
70
80
90
100
110
120
130
f1 (ppm)
7.5
7.0
6.5
6.0
5.5
5.0
4.5
4.0
3.5
3.0
2.5
2.0
f2 (ppm)

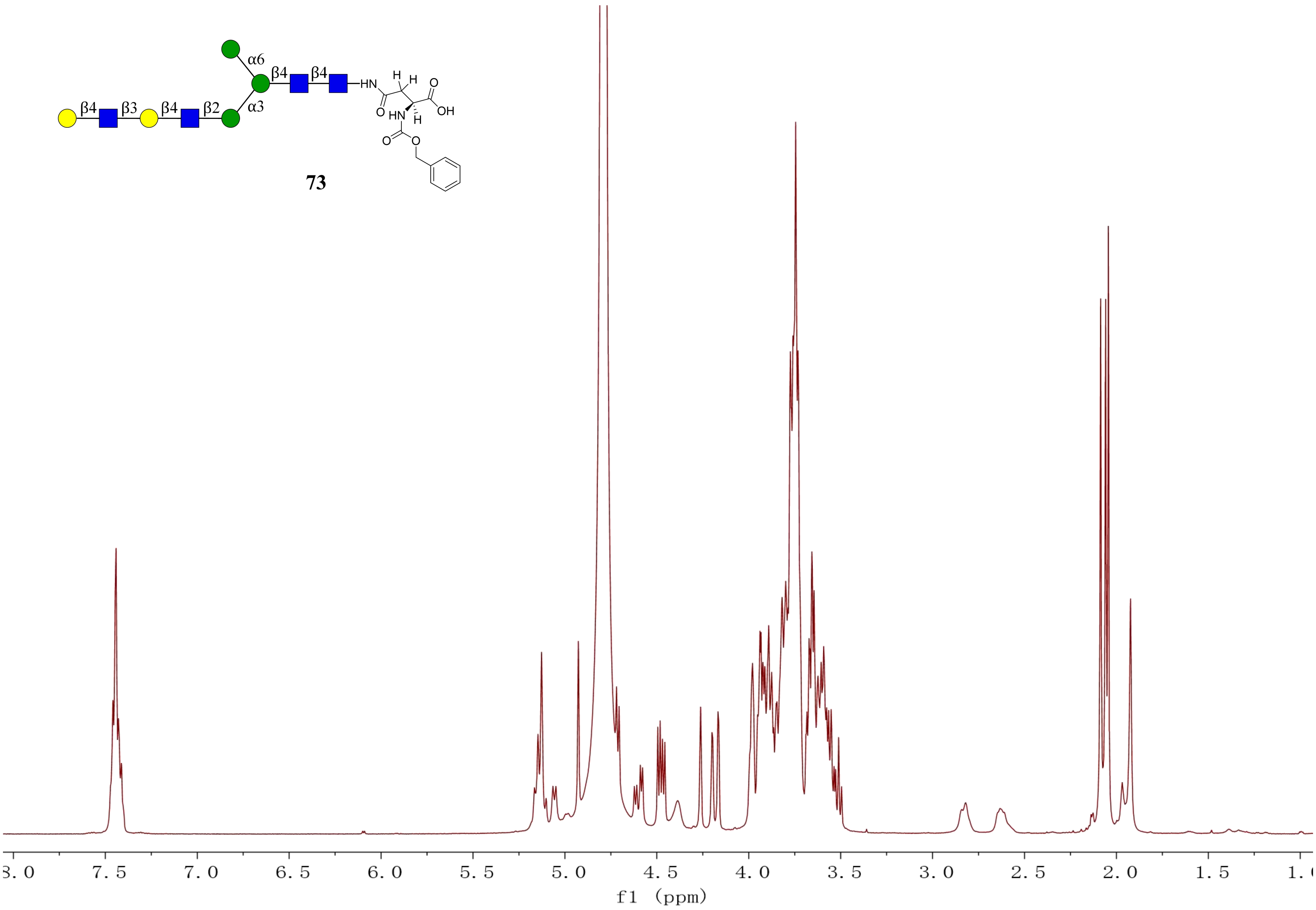
α6
β4
β4
HN
H
H
O
OH
HN
H
O
β4
β3
β4
β2
α3
O
O
73
8.0
7.5
7.0
6.5
6.0
5.5
5.0
4.5
4.0
3.5
3.0
2.5
2.0
1.5
1.0
f1 (ppm)

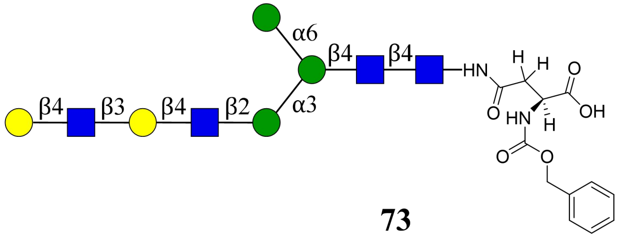
α6
β4
β4
HN
H
H
O
OH
O
HN
H
O
O
β4
β3
β4
β2
α3

**73**

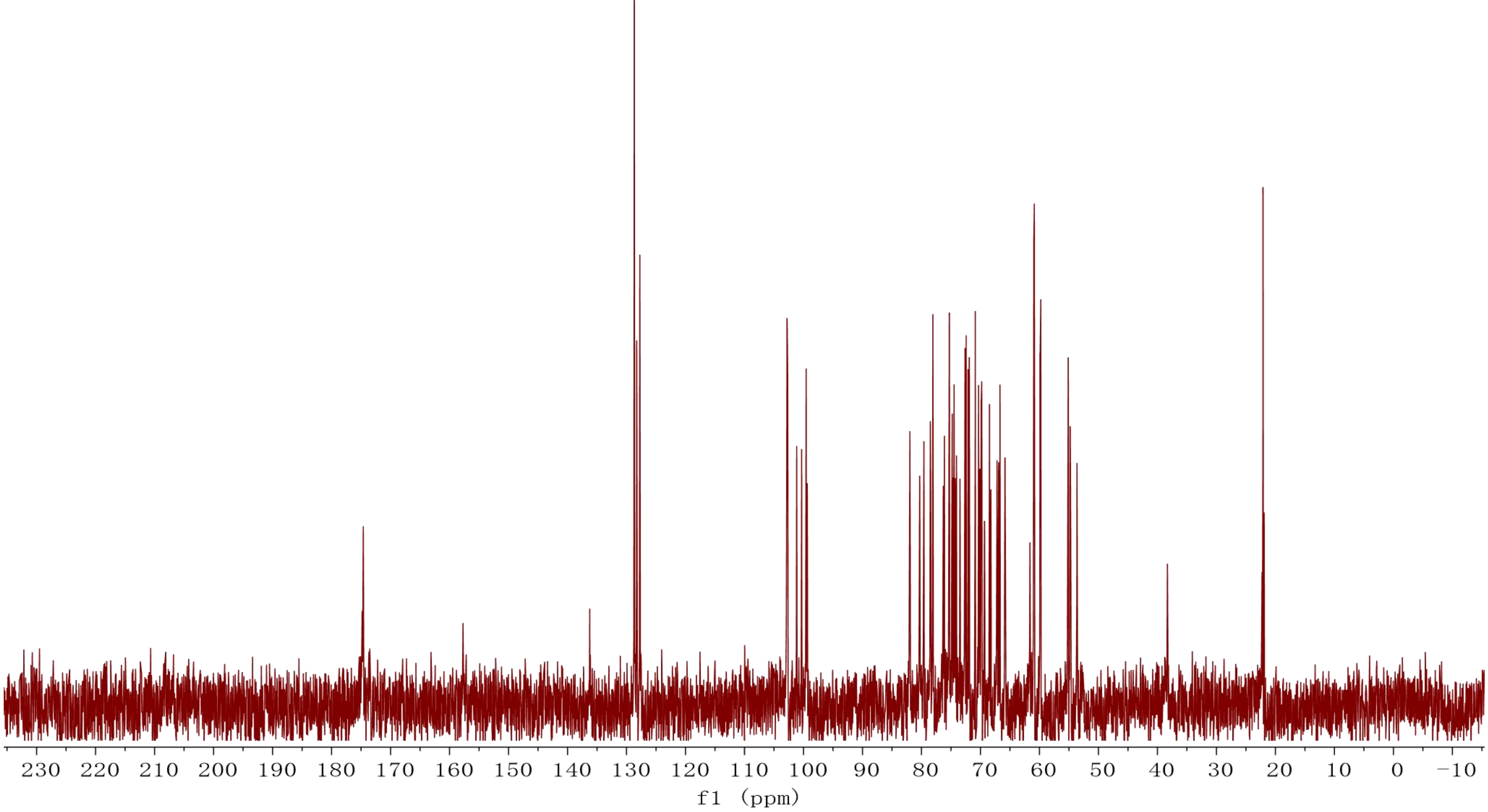
230 220 210 200 190 180 170 160 150 140 130 120 110 100 90 80 70 60 50 40 30 20 10 0 −10
f1 (ppm)

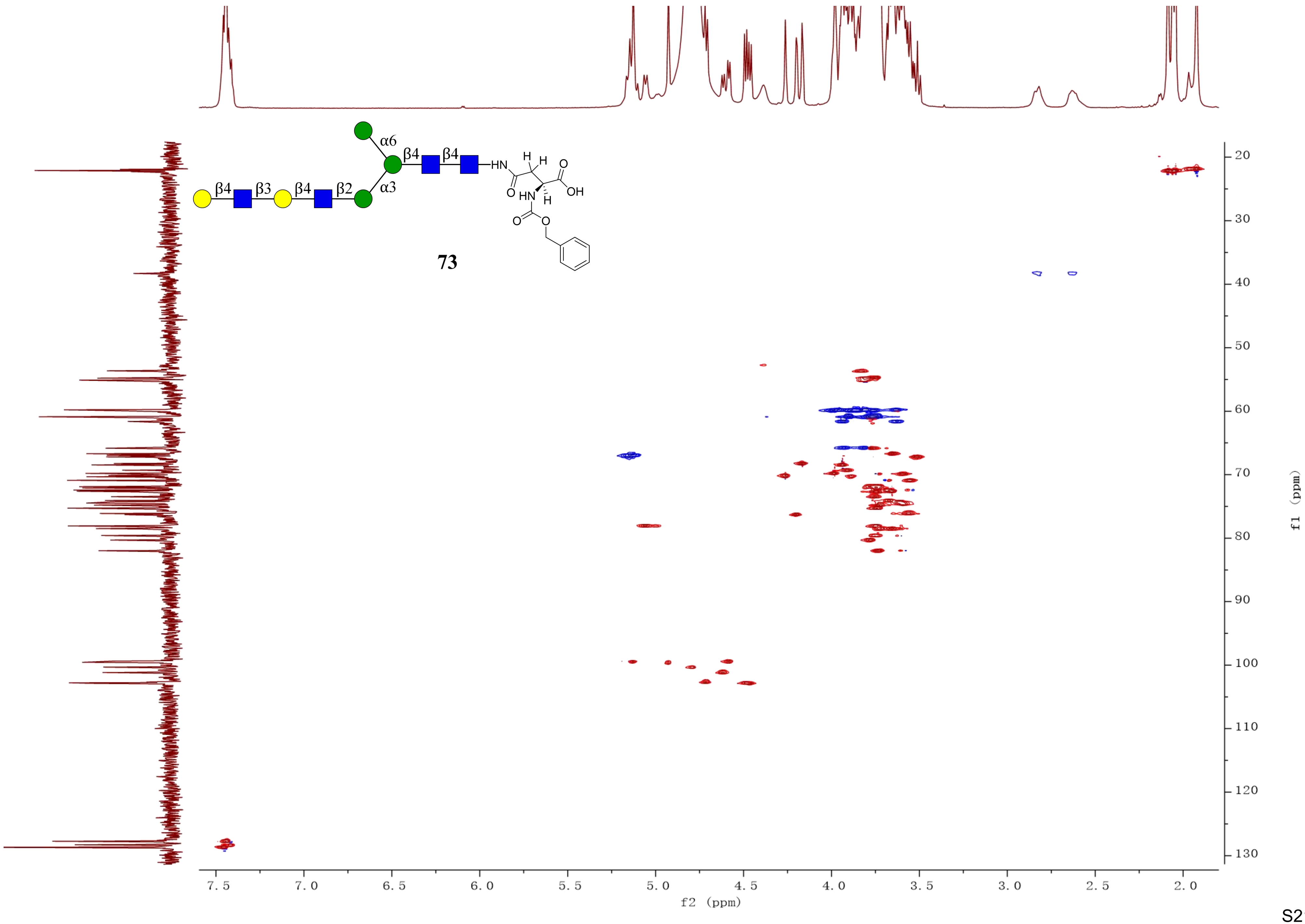

α6
β4
β4
HN
H
H
O
β4
β3
β4
β2
α3
O
HN
H
OH
O
O
73
f1 (ppm)
f2 (ppm)

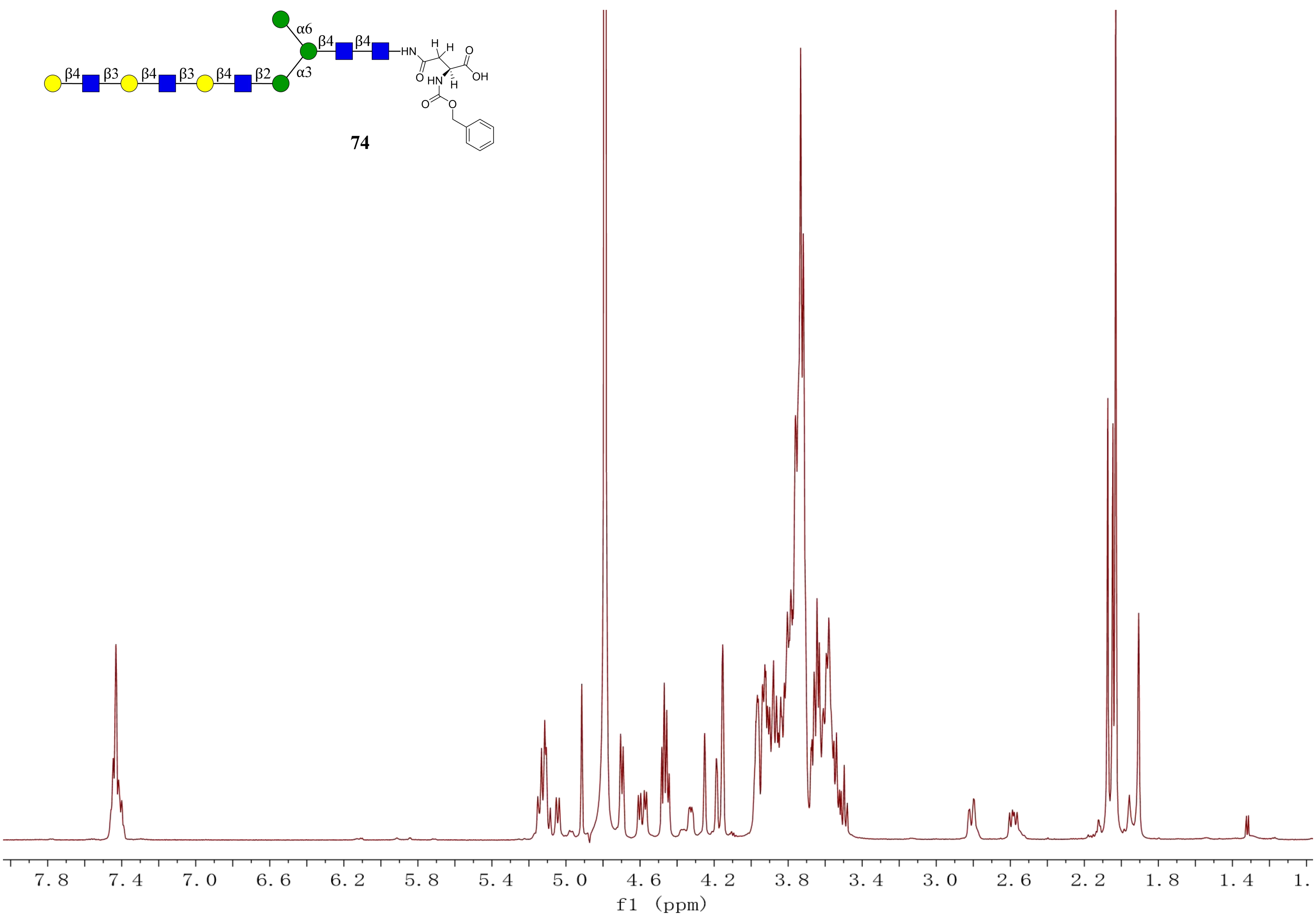

α6
β4
β4
β4
β3
β4
β3
β4
β2
α3
HN
H
H
O
O
OH
HN
H
O
O
7.8
7.4
7.0
6.6
6.2
5.8
5.4
5.0
4.6
4.2
3.8
3.4
3.0
2.6
2.2
1.8
1.4
1.
f1 (ppm)


**74**

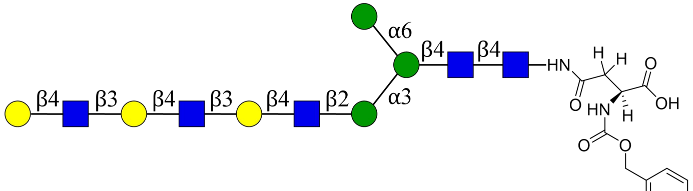
α6
β4
β4
α3
β4
β3
β4
β3
β4
β2
HN
O
OH
HN
O
O

**74**

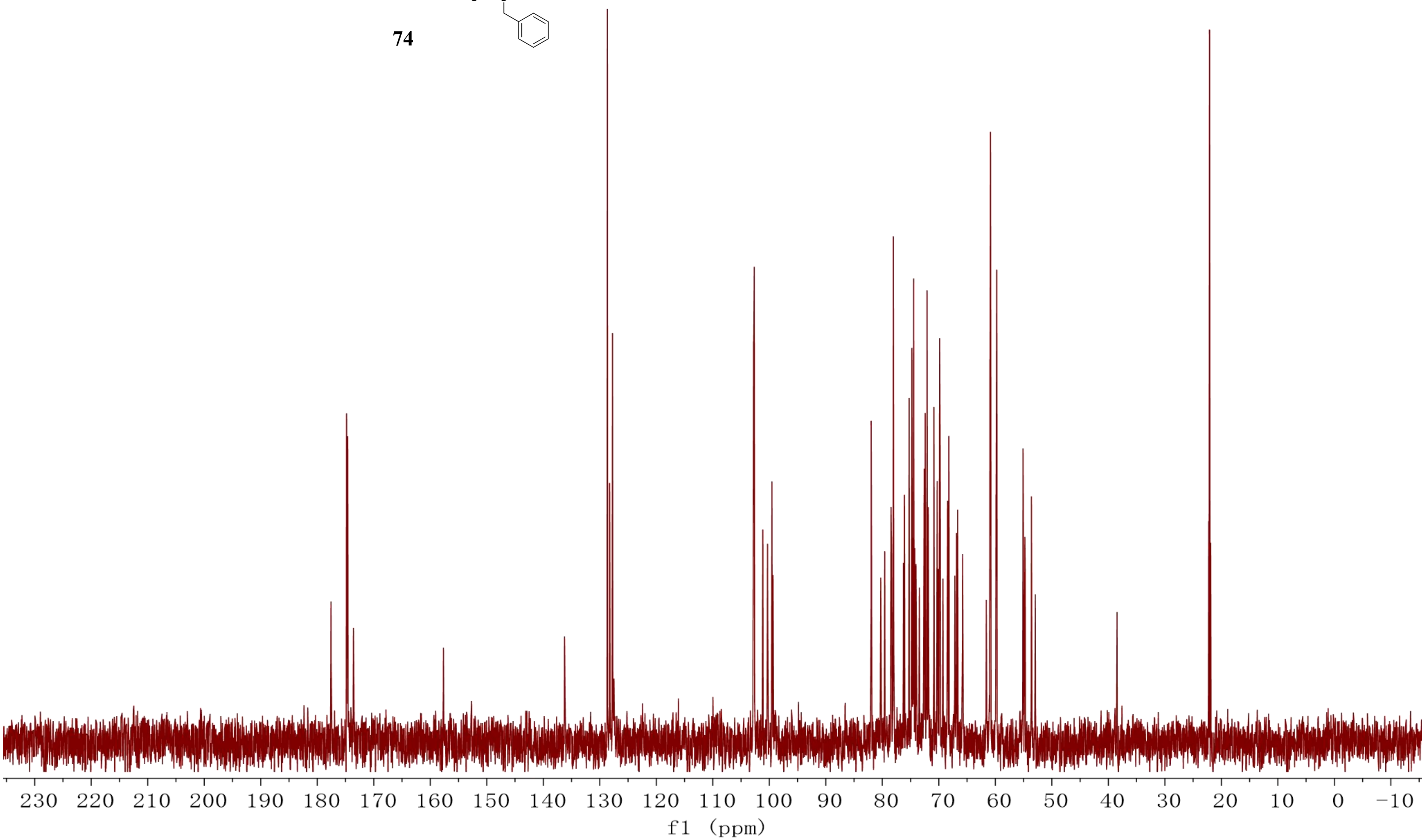
230 220 210 200 190 180 170 160 150 140 130 120 110 100 90 80 70 60 50 40 30 20 10 0 −10
f1 (ppm)

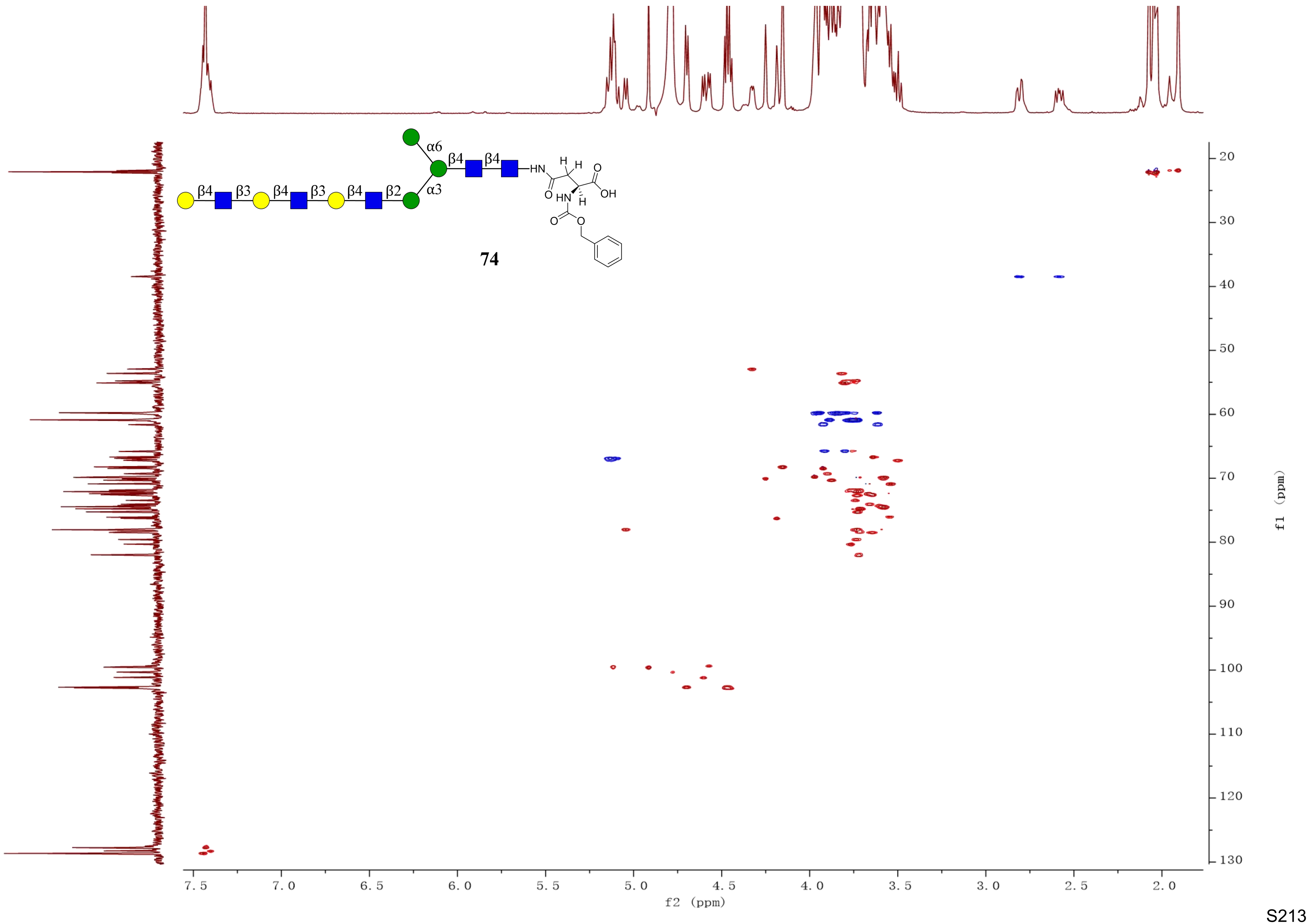

α6
β4
β4
HN
H
H
O
OH
O
HN
H
O
O
β4
β3
β4
β3
β4
β2
α3
74
f2 (ppm)
f1 (ppm)
7.5
7.0
6.5
6.0
5.5
5.0
4.5
4.0
3.5
3.0
2.5
2.0
20
30
40
50
60
70
80
90
100
110
120
130

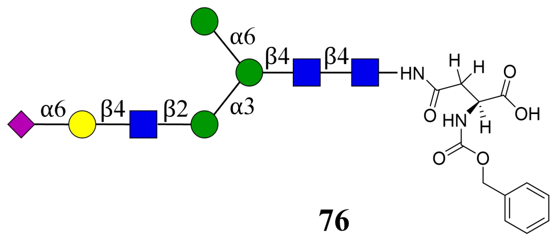
α6
β4
β4
α6
β4
β2
α3
HN
H
H
O
OH
O
HN
H
O
O

**76**

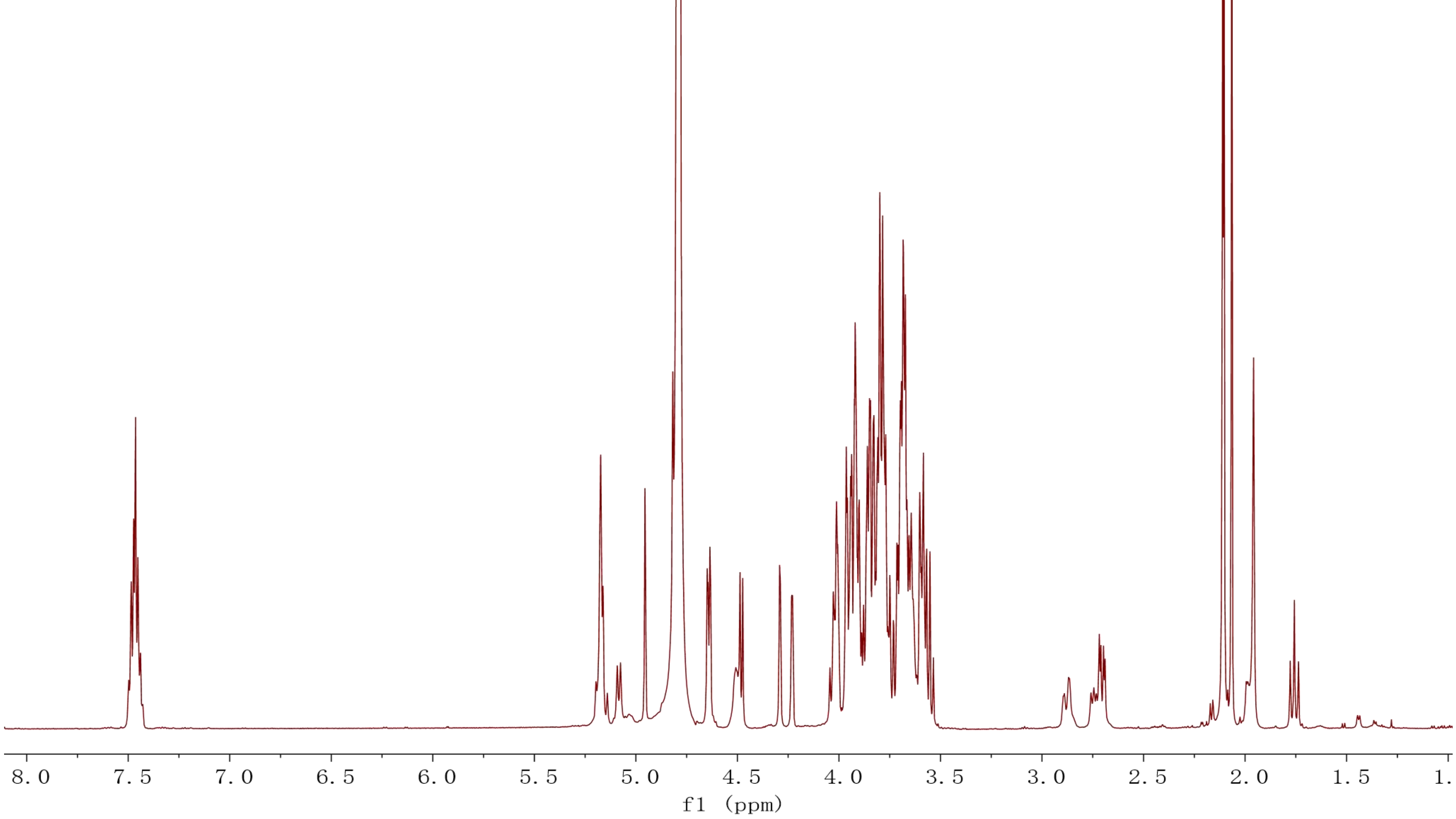
8.0
7.5
7.0
6.5
6.0
5.5
5.0
4.5
4.0
3.5
3.0
2.5
2.0
1.5
1.
f1 (ppm)

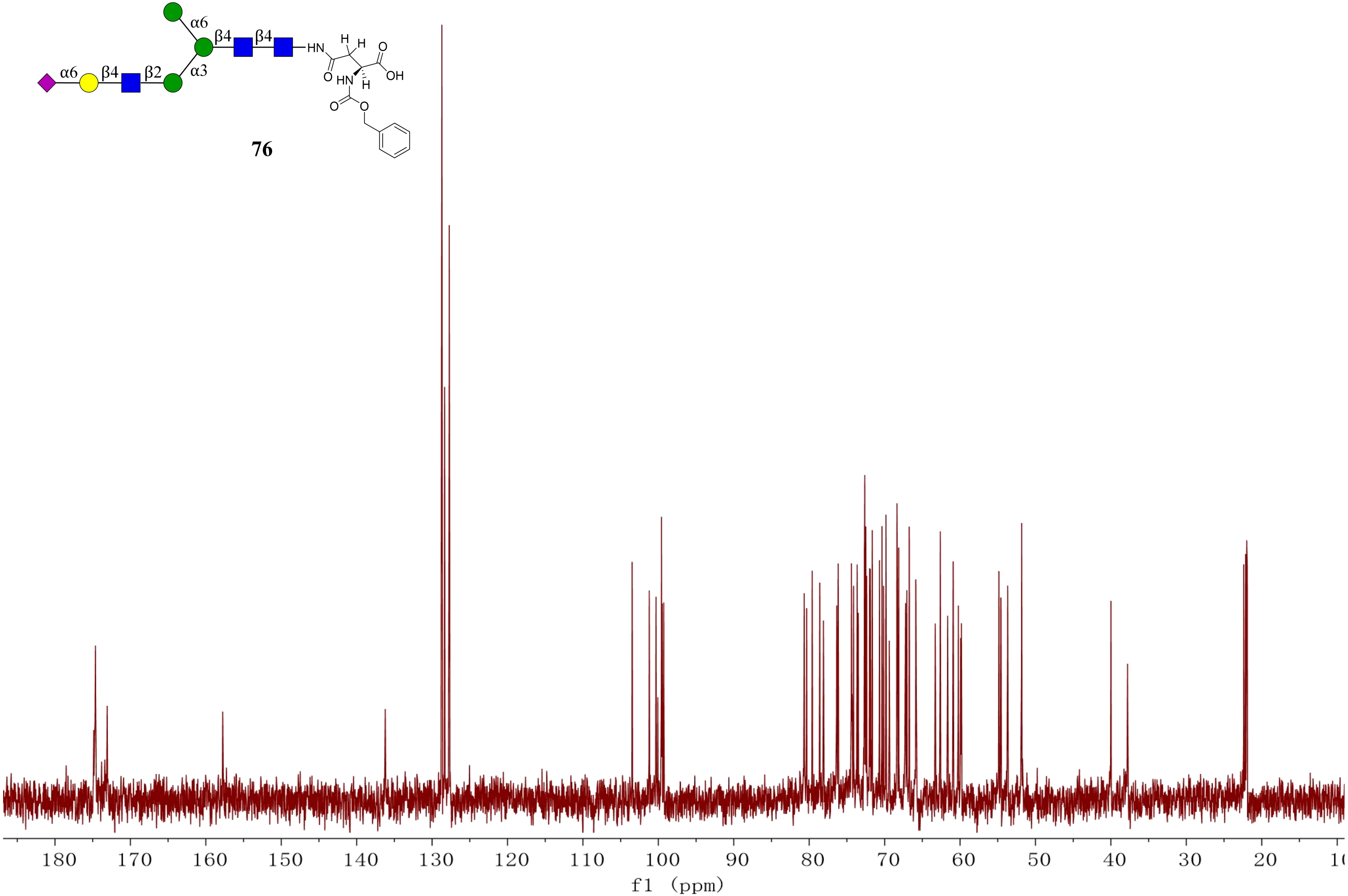
α6
β4
β4
α6
β4
β2
α3
HN
H
H
O
O
OH
HN
H
O
O

**76**
180
170
160
150
140
130
120
110
100
90
80
70
60
50
40
30
20
f1 (ppm)

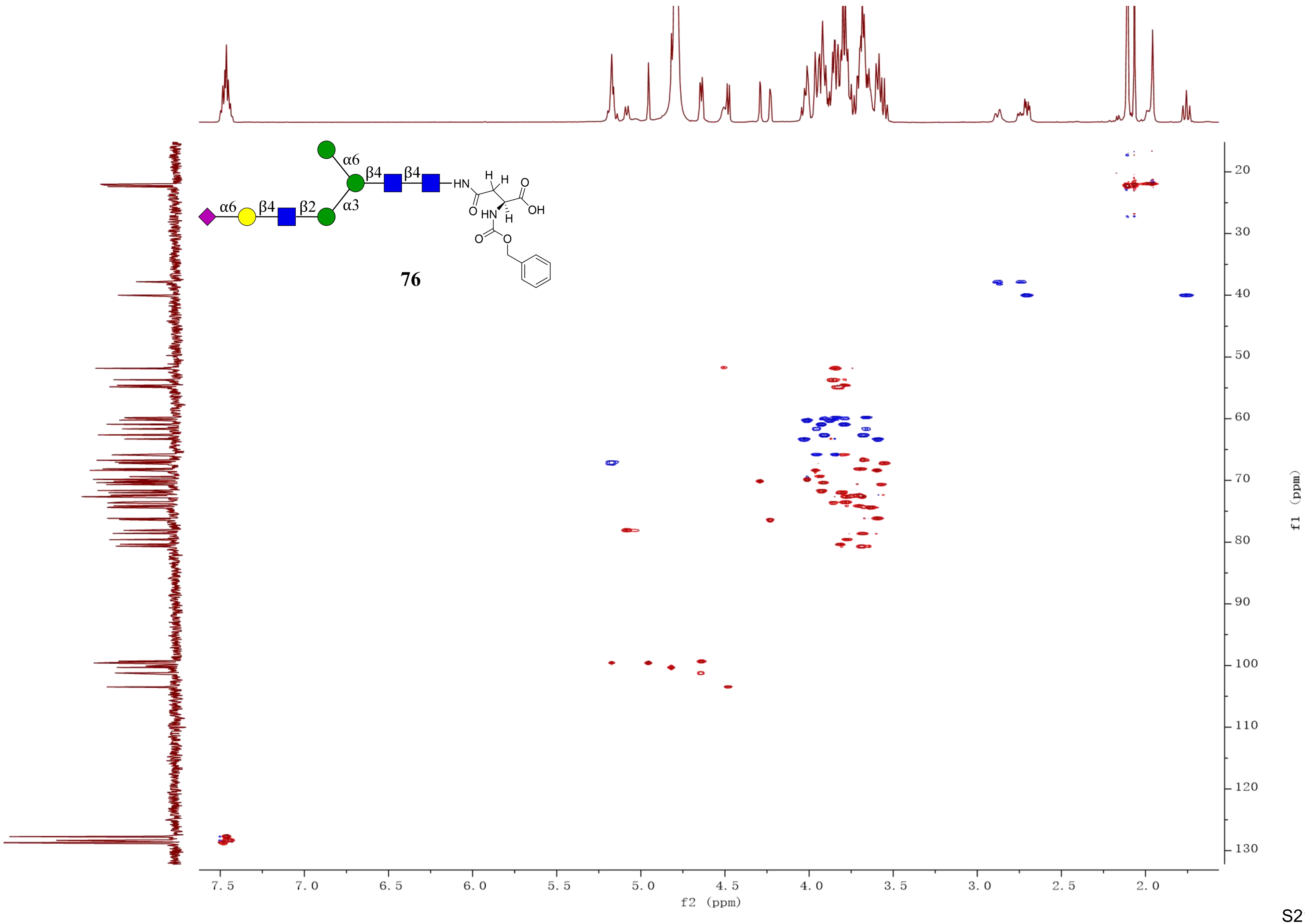


76

**78**

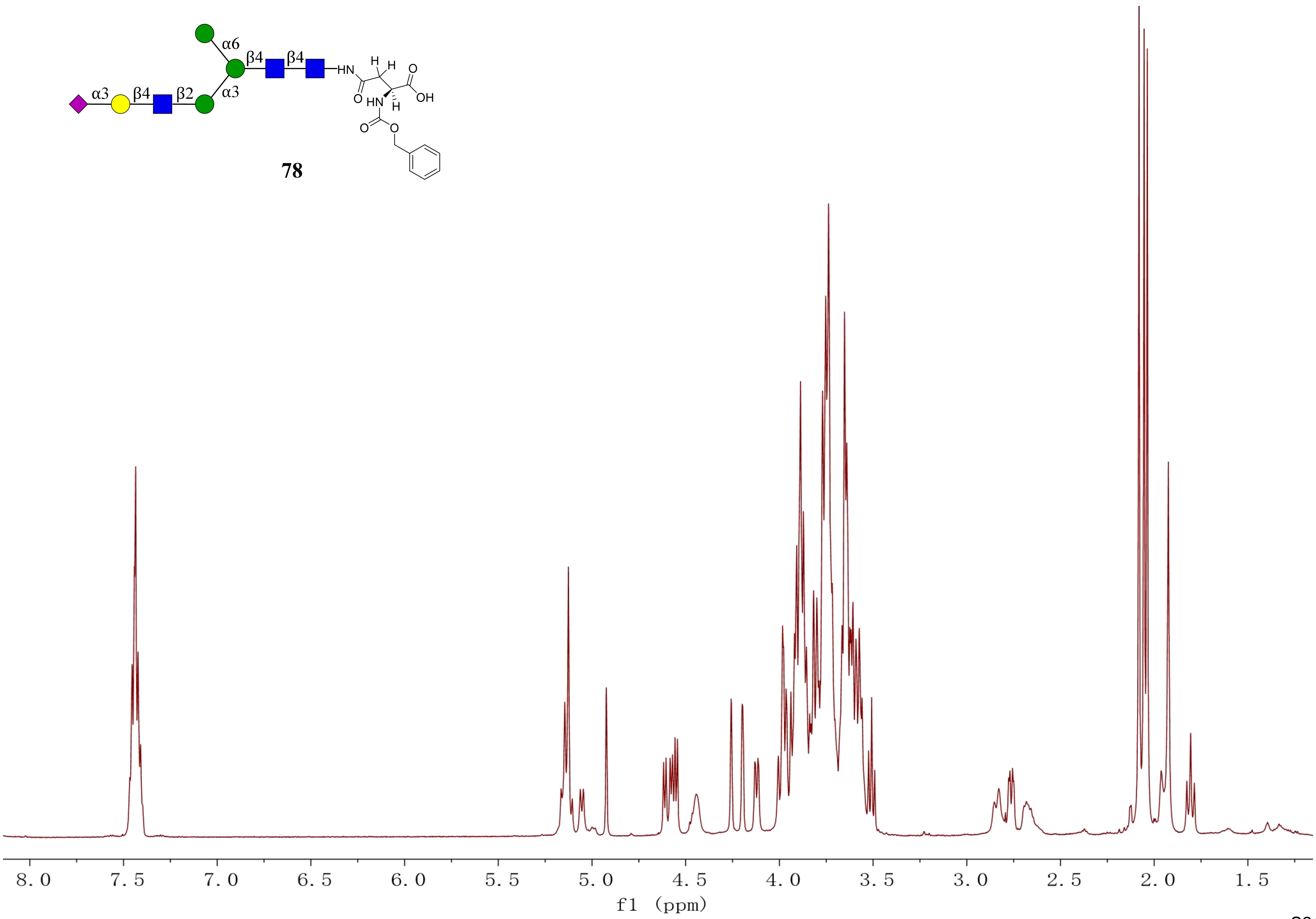
α6
β4
β4
α3
β4
β2
α3
HN
H
H
O
OH
O
HN
H
O
O
8.0
7.5
7.0
6.5
6.0
5.5
5.0
4.5
4.0
3.5
3.0
2.5
2.0
1.5
f1 (ppm)

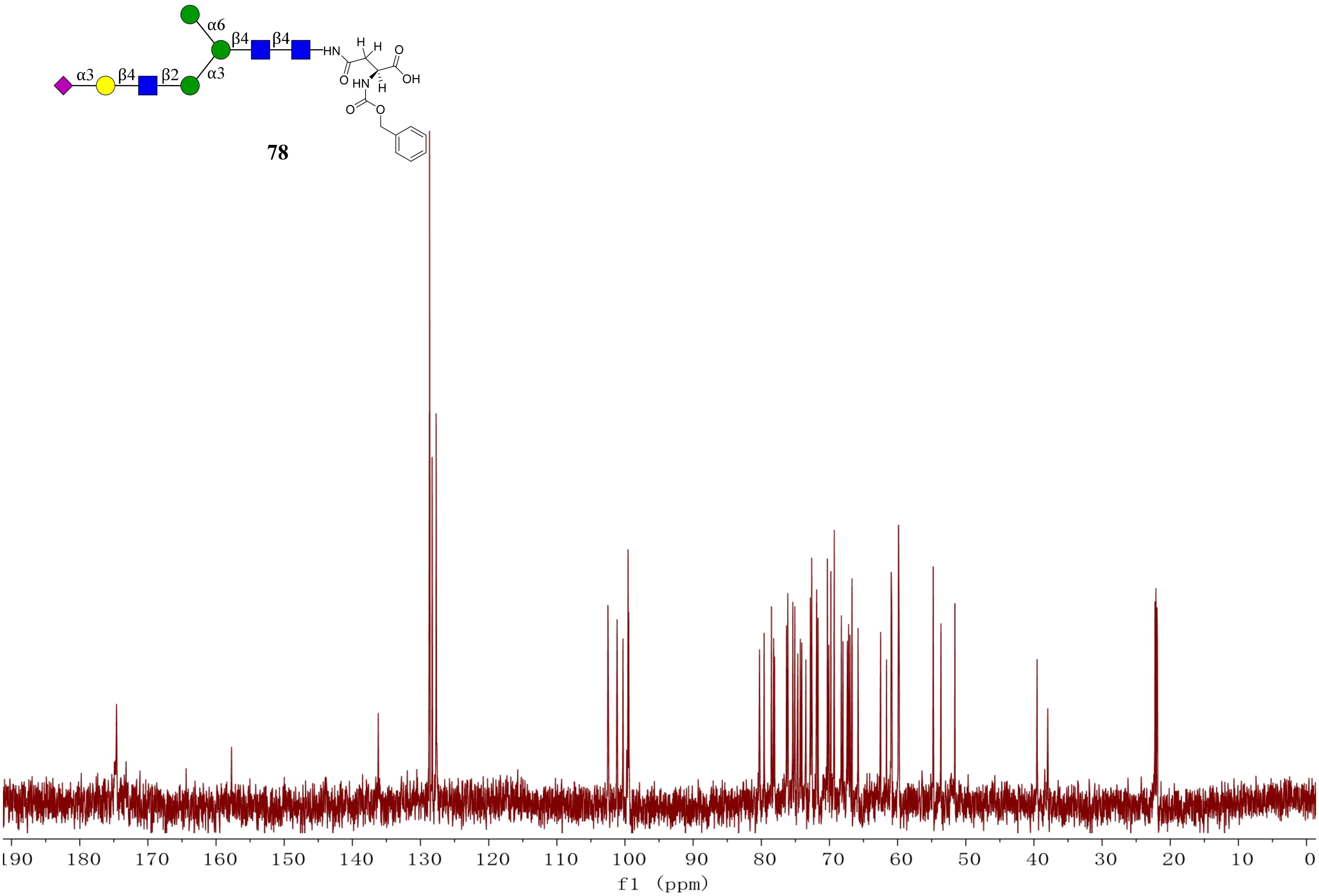
α6
β4
β4
α3
β4
β2
α3
HN
H
H
O
O
OH
HN
H
O
O
78
190
180
170
160
150
140
130
120
110
100
90
80
70
60
50
40
30
20
10
0
f1 (ppm)

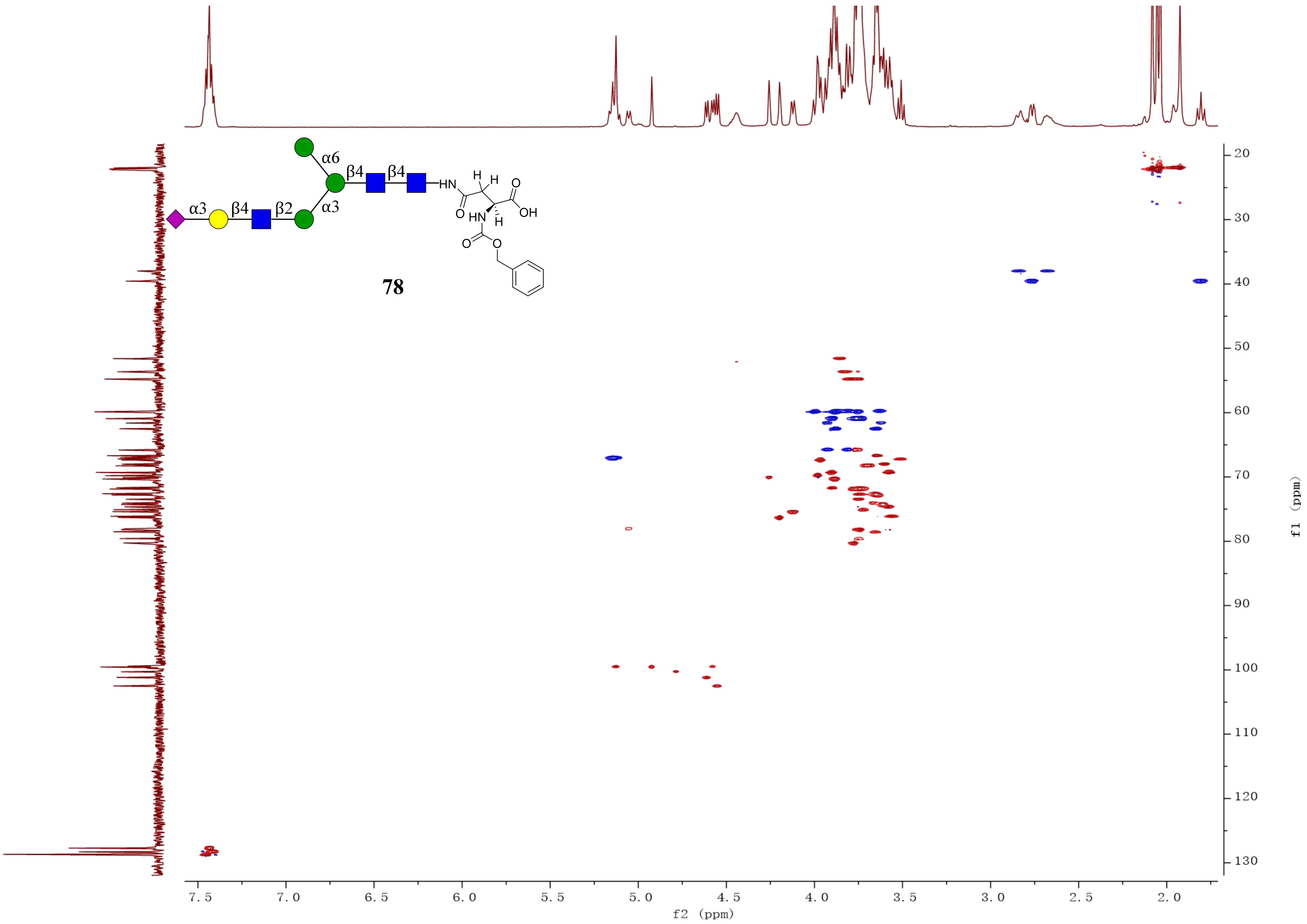

α6
β4
β4
α3
β4
β2
α3
HN
H
H
O
O
HN
H
OH
O
O
78
20
30
40
50
60
70
80
90
100
110
120
130
f1 (ppm)
7.5
7.0
6.5
6.0
5.5
5.0
4.5
4.0
3.5
3.0
2.5
2.0
f2 (ppm)

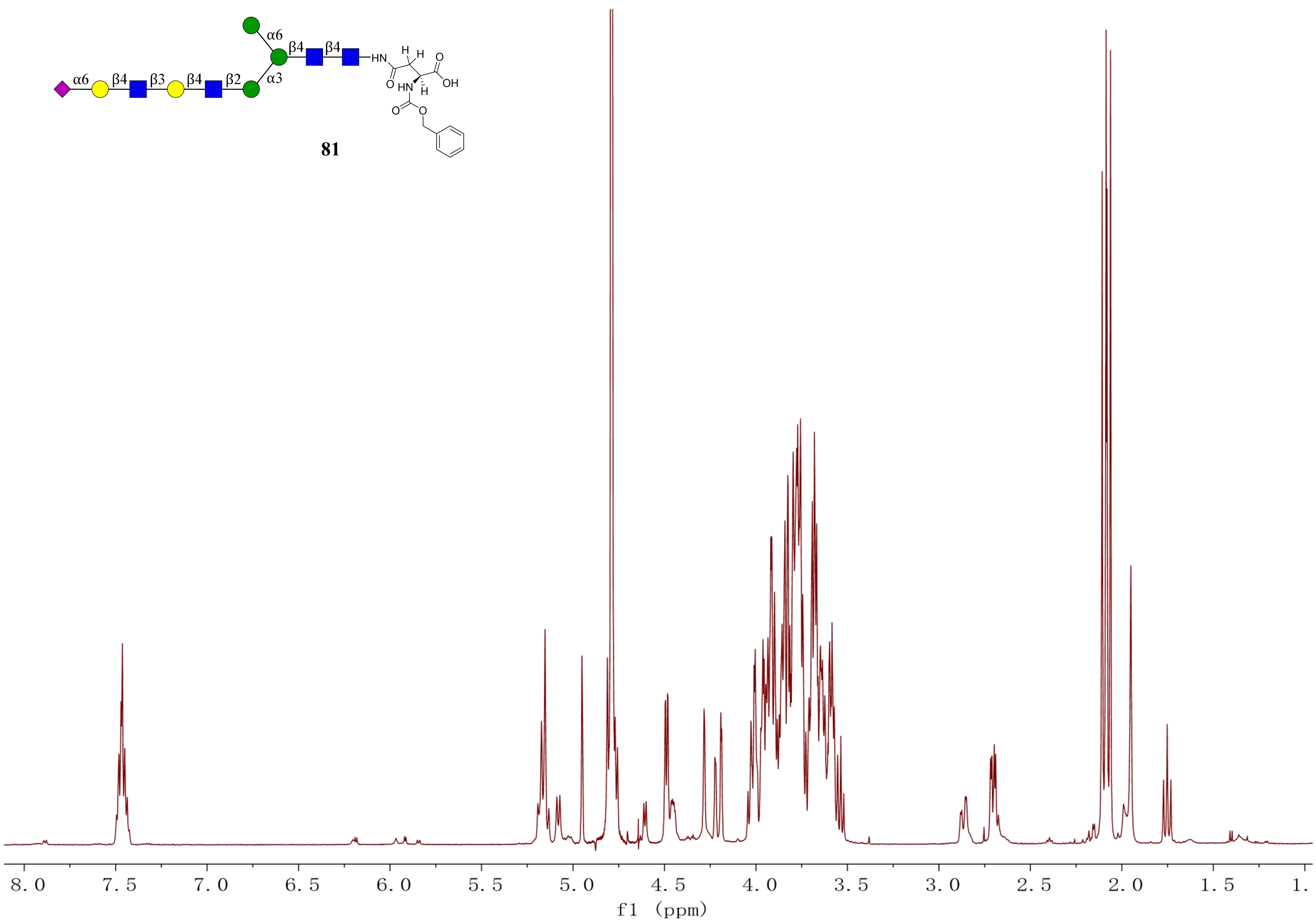
α6
β4
β3
β2
α3
HN
H
O
OH
HN
81
8.0
7.5
7.0
6.5
6.0
5.5
5.0
4.5
4.0
3.5
3.0
2.5
2.0
1.5
1.
f1 (ppm)

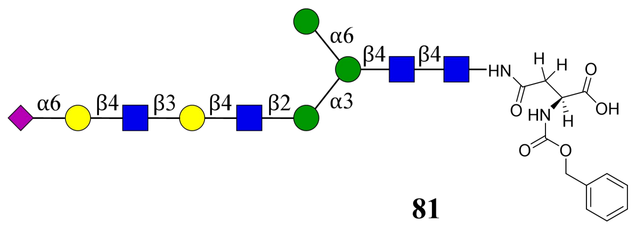
α6
β4
β4
α6
β4
β3
β4
β2
α3
HN
O
H
H
O
OH
HN
H
O
O

**81**

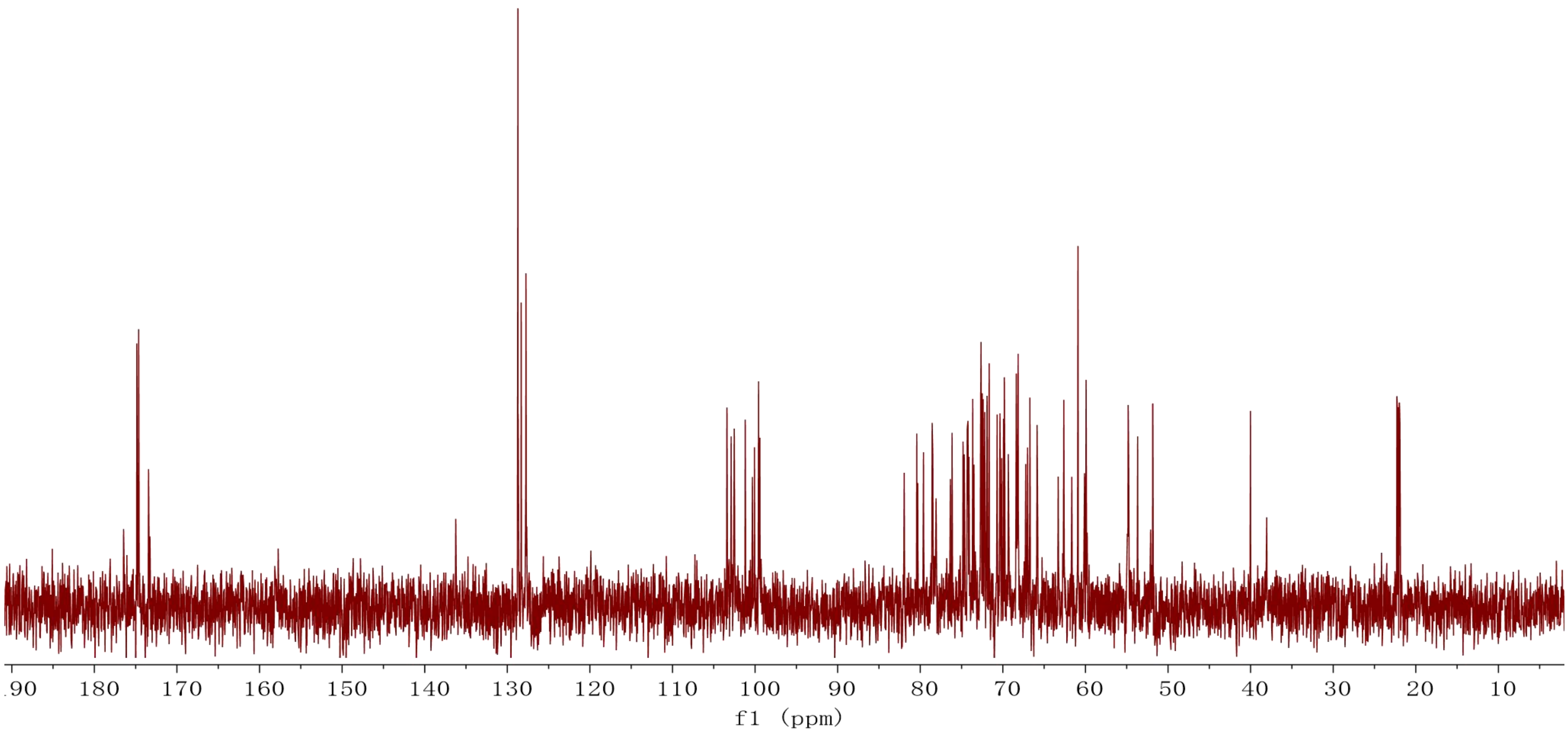
190
180
170
160
150
140
130
120
110
100
90
80
70
60
50
40
30
20
10
f1 (ppm)

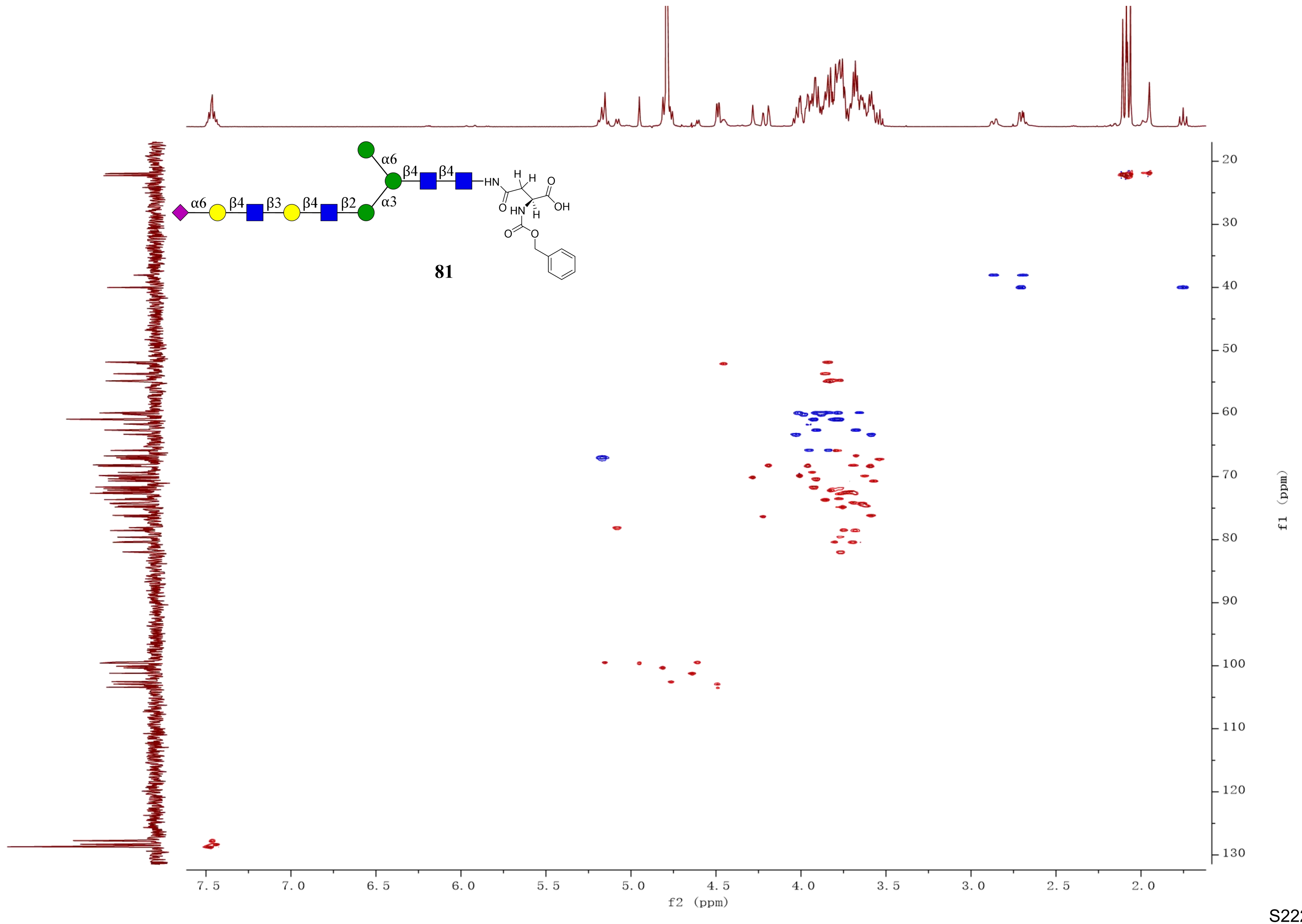
α6
β4
β4
HN
H
H
O
OH
α6
β4
β3
β4
β2
α3
O
HN
H
O
O
81
f1 (ppm)
f2 (ppm)

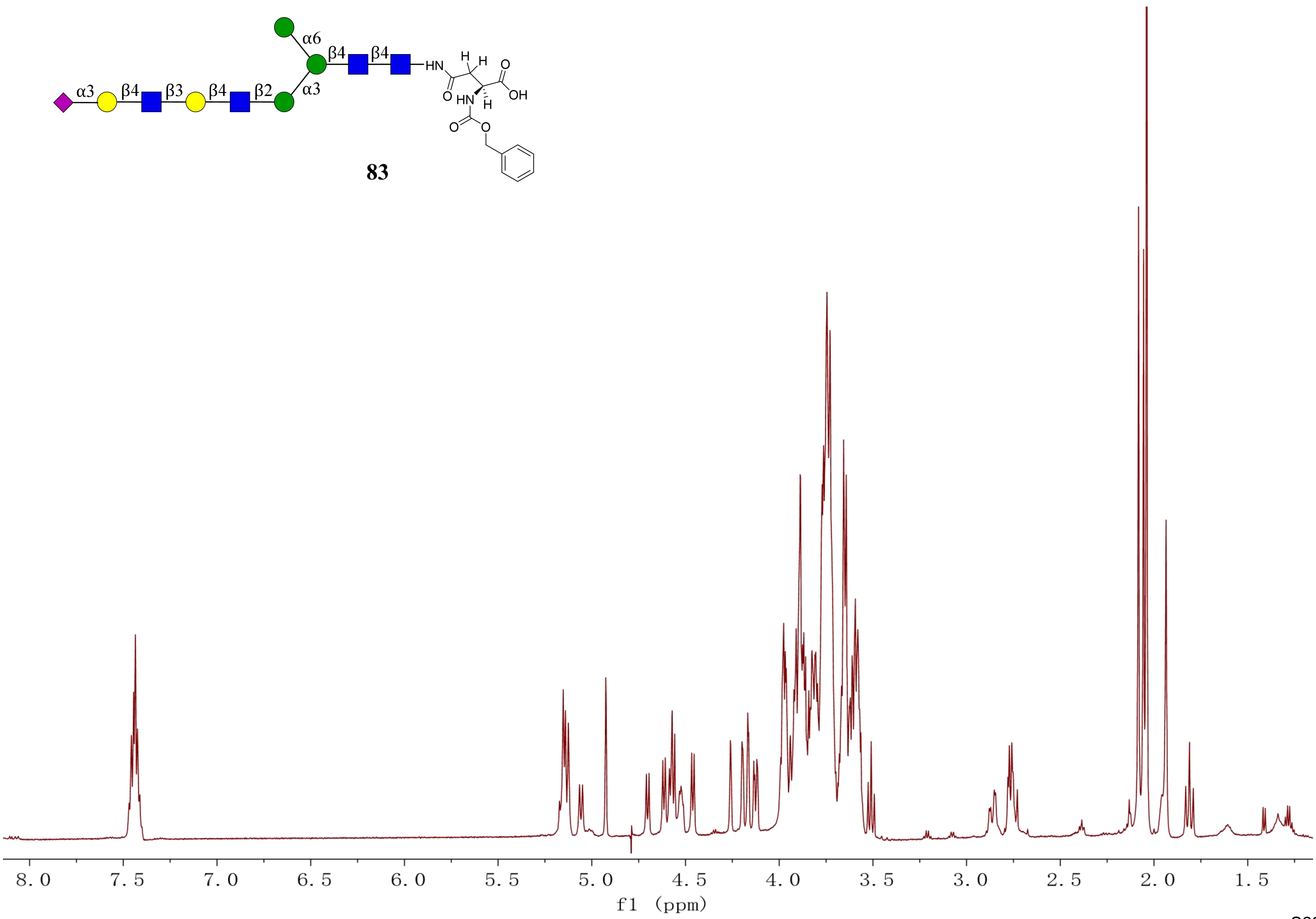
α6
β4
β4
HN
H
H
O
OH
O
HN
H
O
O
α3
β4
β3
β4
β2
α3
83
8.0
7.5
7.0
6.5
6.0
5.5
5.0
4.5
4.0
3.5
3.0
2.5
2.0
1.5
f1 (ppm)

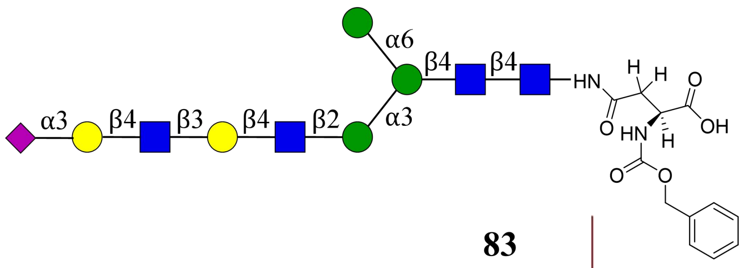
α6
β4
β4
α3
β4
β3
β4
β2
α3
HN
H
H
O
OH
O
HN
H
O
O

**83**

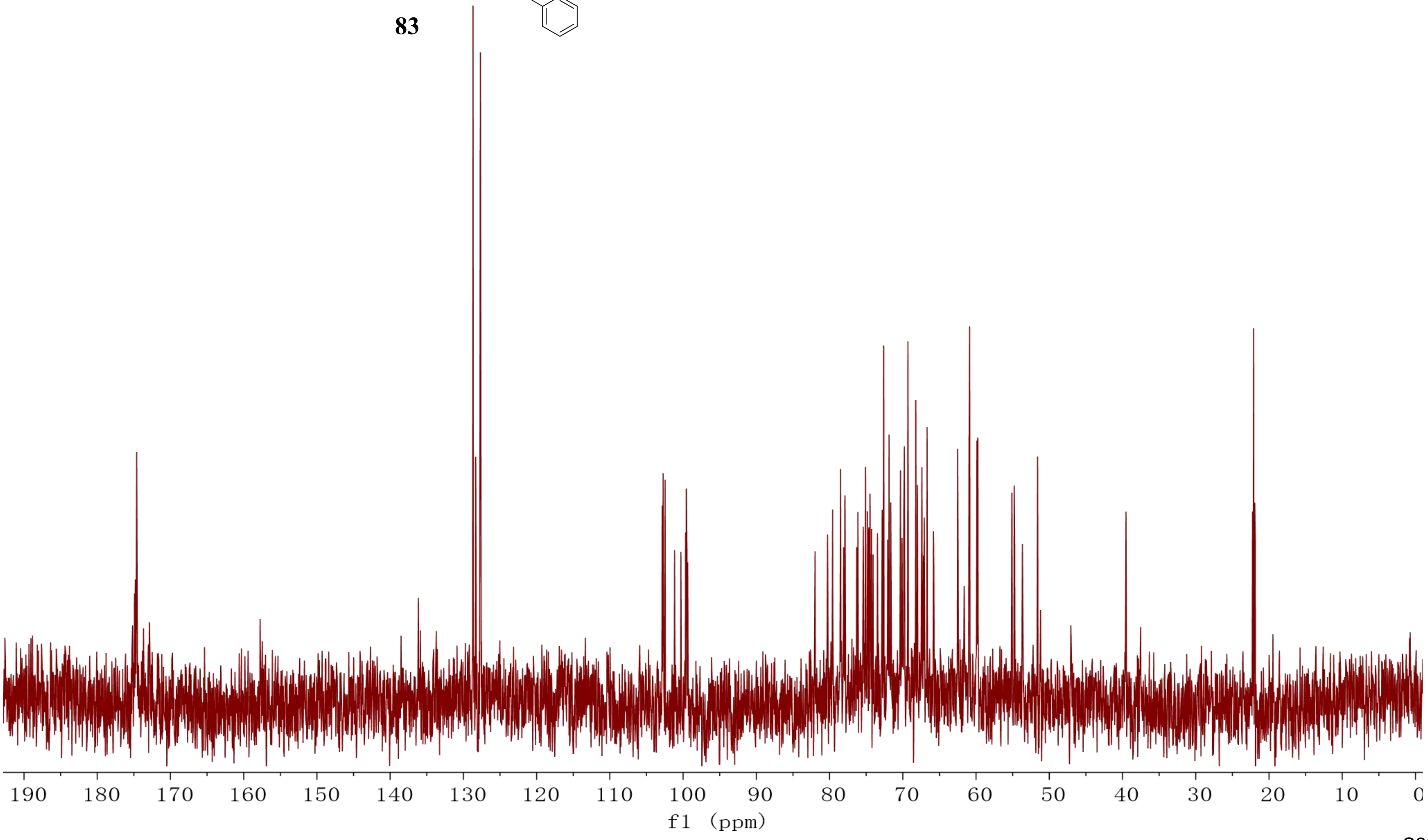
190
180
170
160
150
140
130
120
110
100
90
80
70
60
50
40
30
20
10
0
f1 (ppm)

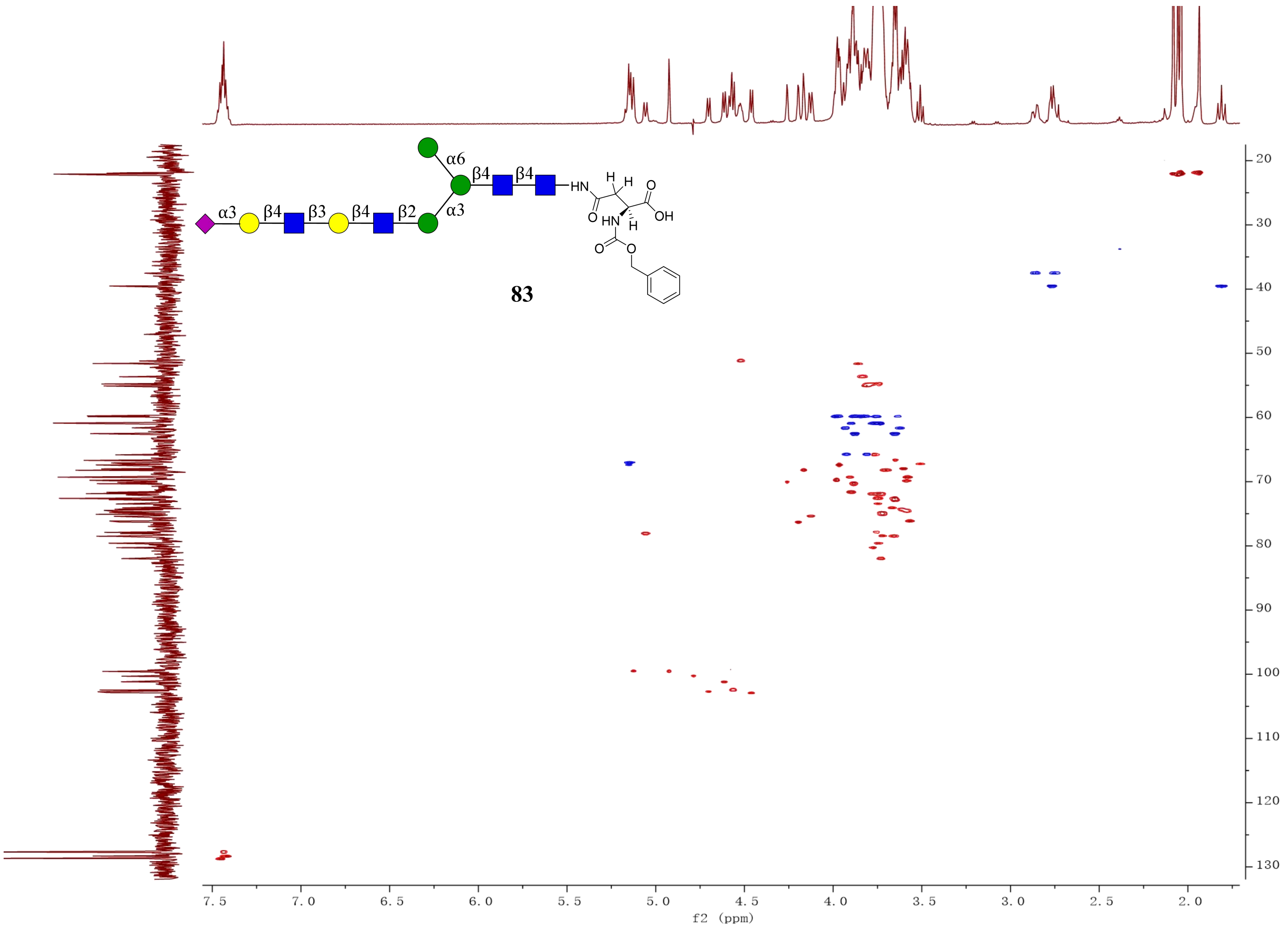


**83**

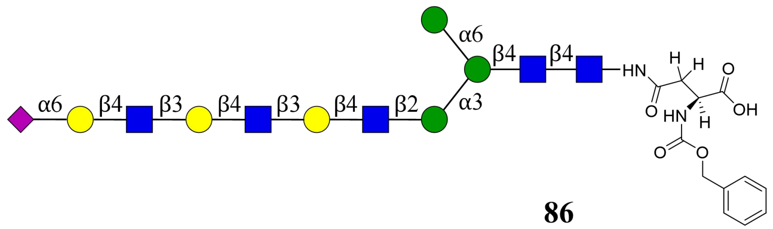
α6
β4
β3
β4
β3
β4
β2
α3
α6
β4
β4
HN
H
H
O
OH
O
HN
H
O
O

**86**

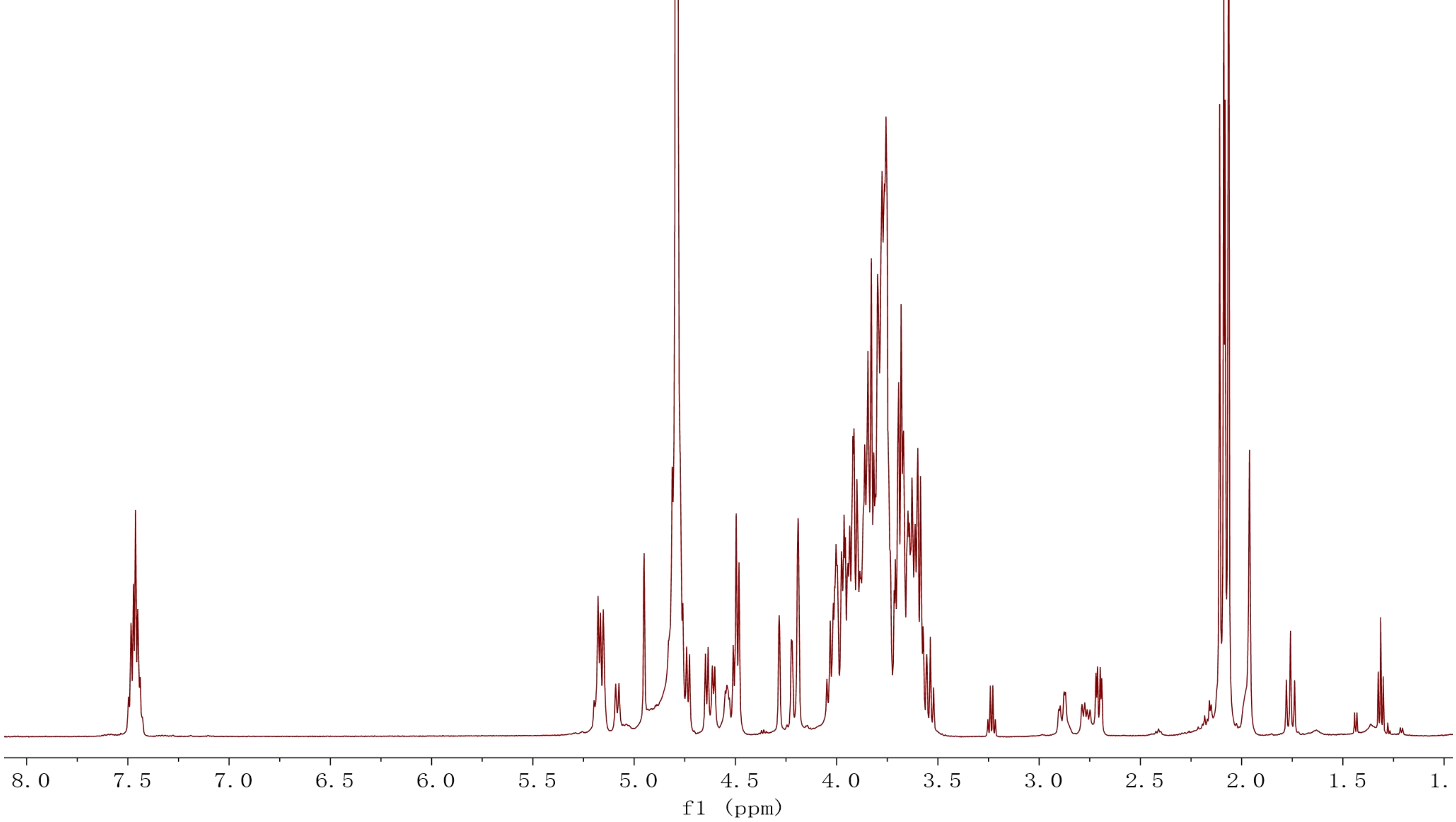
8.0
7.5
7.0
6.5
6.0
5.5
5.0
4.5
4.0
3.5
3.0
2.5
2.0
1.5
1.
f1 (ppm)

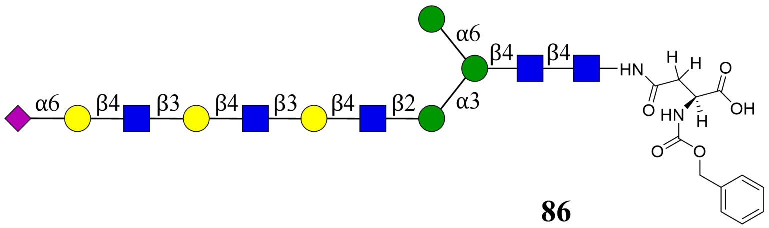

α6
β4
β4
β3
β4
β3
β4
β2
α3
HN
H
O
OH
HN
O


**86**

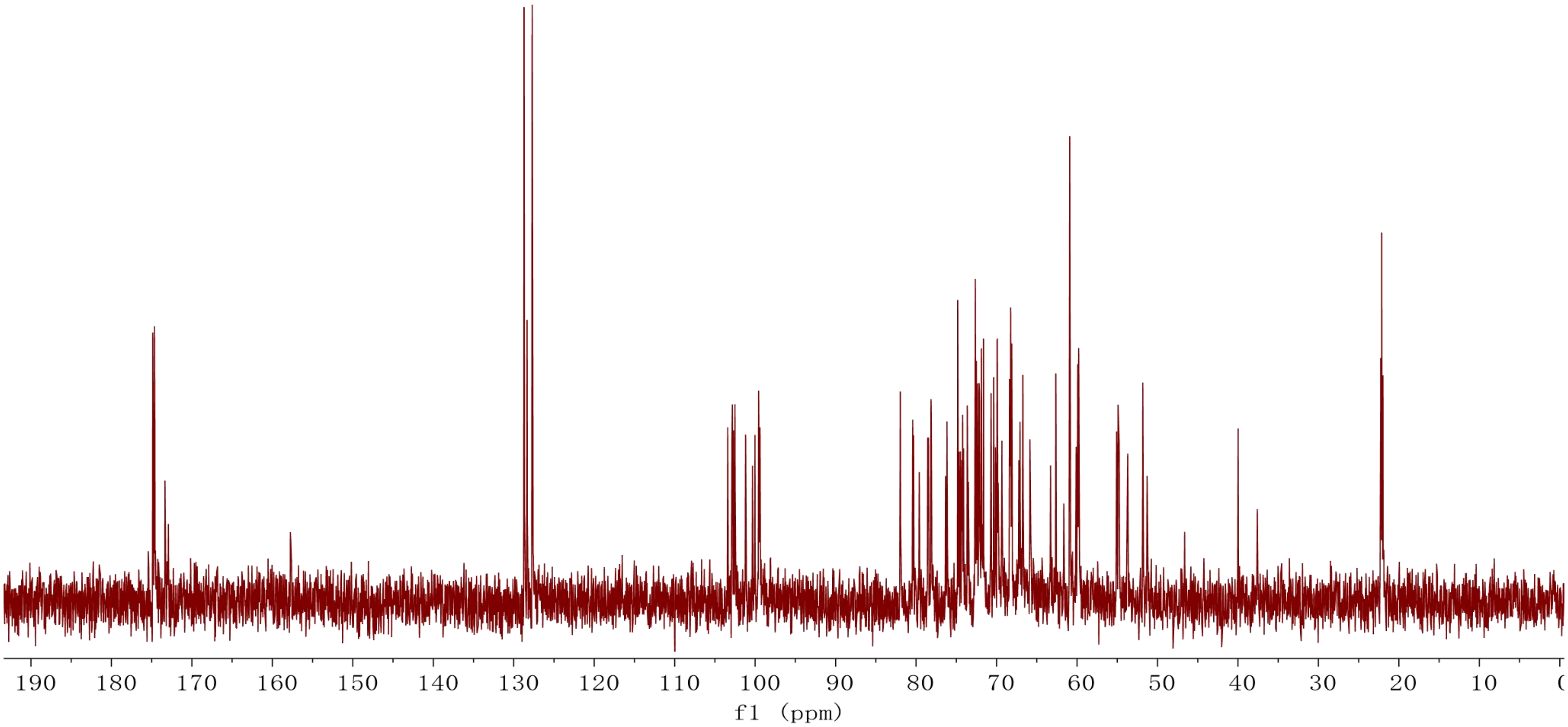

190
180
170
160
150
140
130
120
110
100
90
80
70
60
50
40
30
20
10
f1 (ppm)

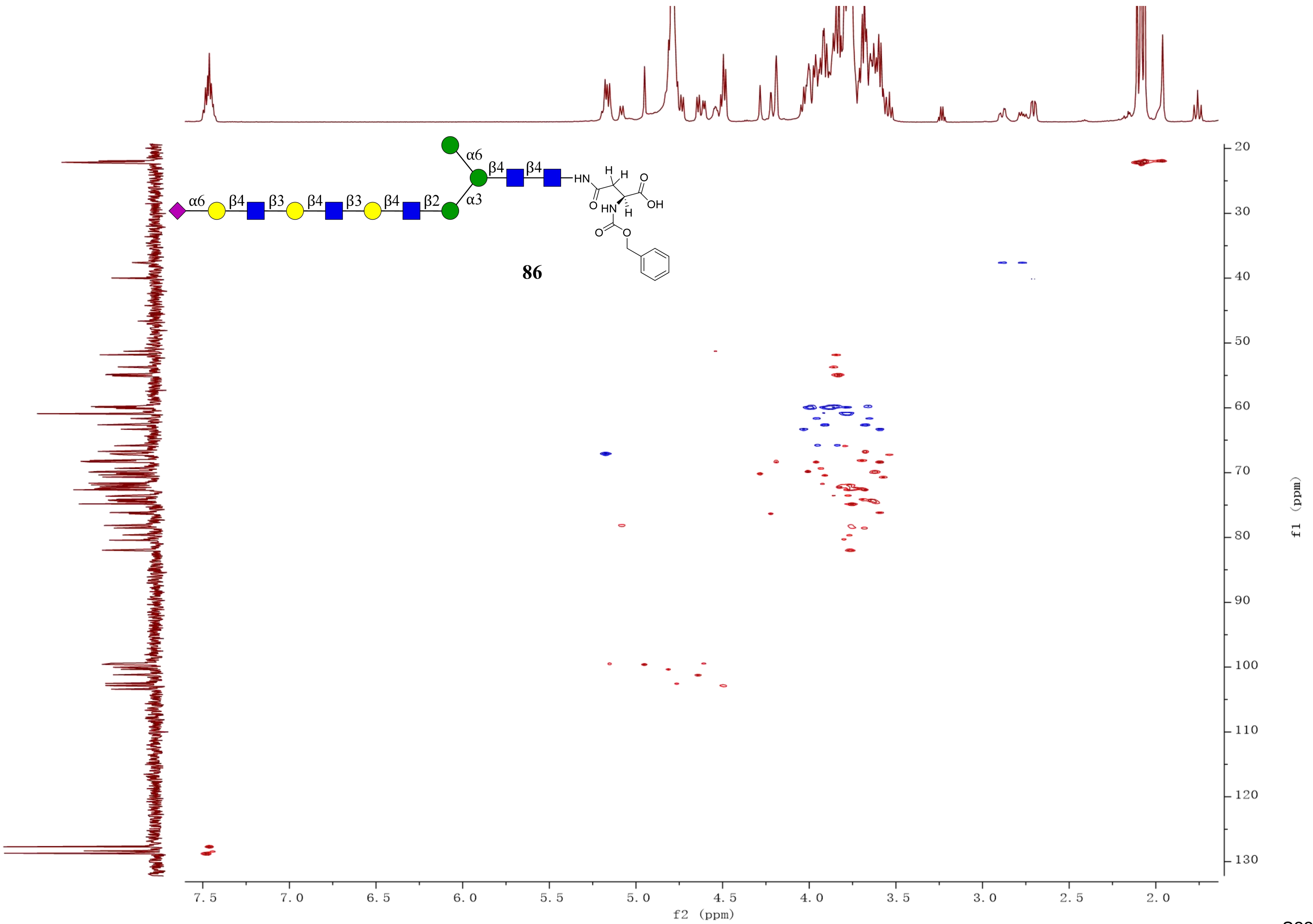
α6
β4
β3
β4
β3
β4
β2
α3
α6
β4
β4
HN
OH
86
f2 (ppm)
f1 (ppm)

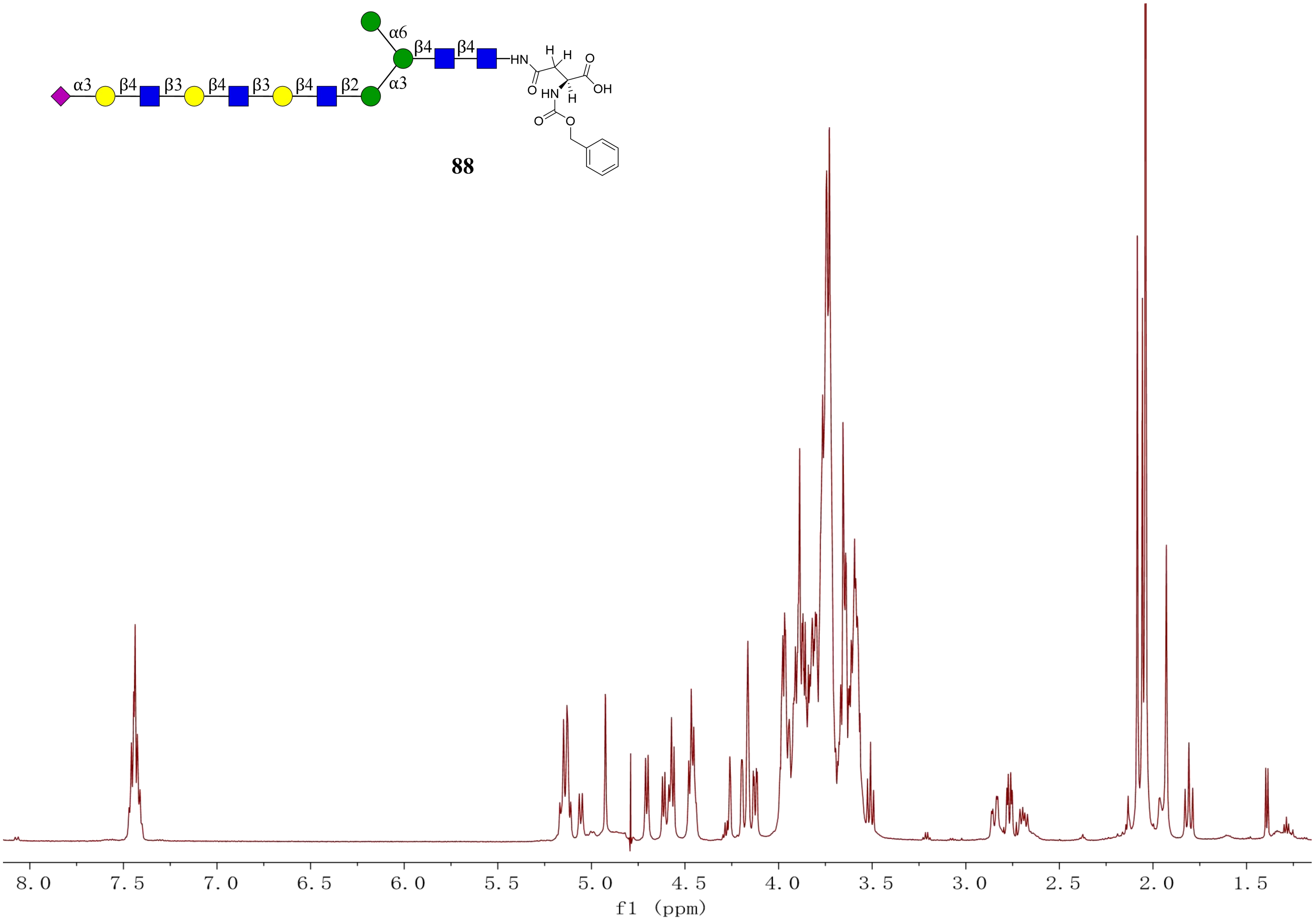
α6
β4
β4
HN
α3
β4
β3
β4
β3
β4
β2
α3
OH
HN
8.0 7.5 7.0 6.5 6.0 5.5 5.0 4.5 4.0 3.5 3.0 2.5 2.0 1.5
f1 (ppm)

**88**

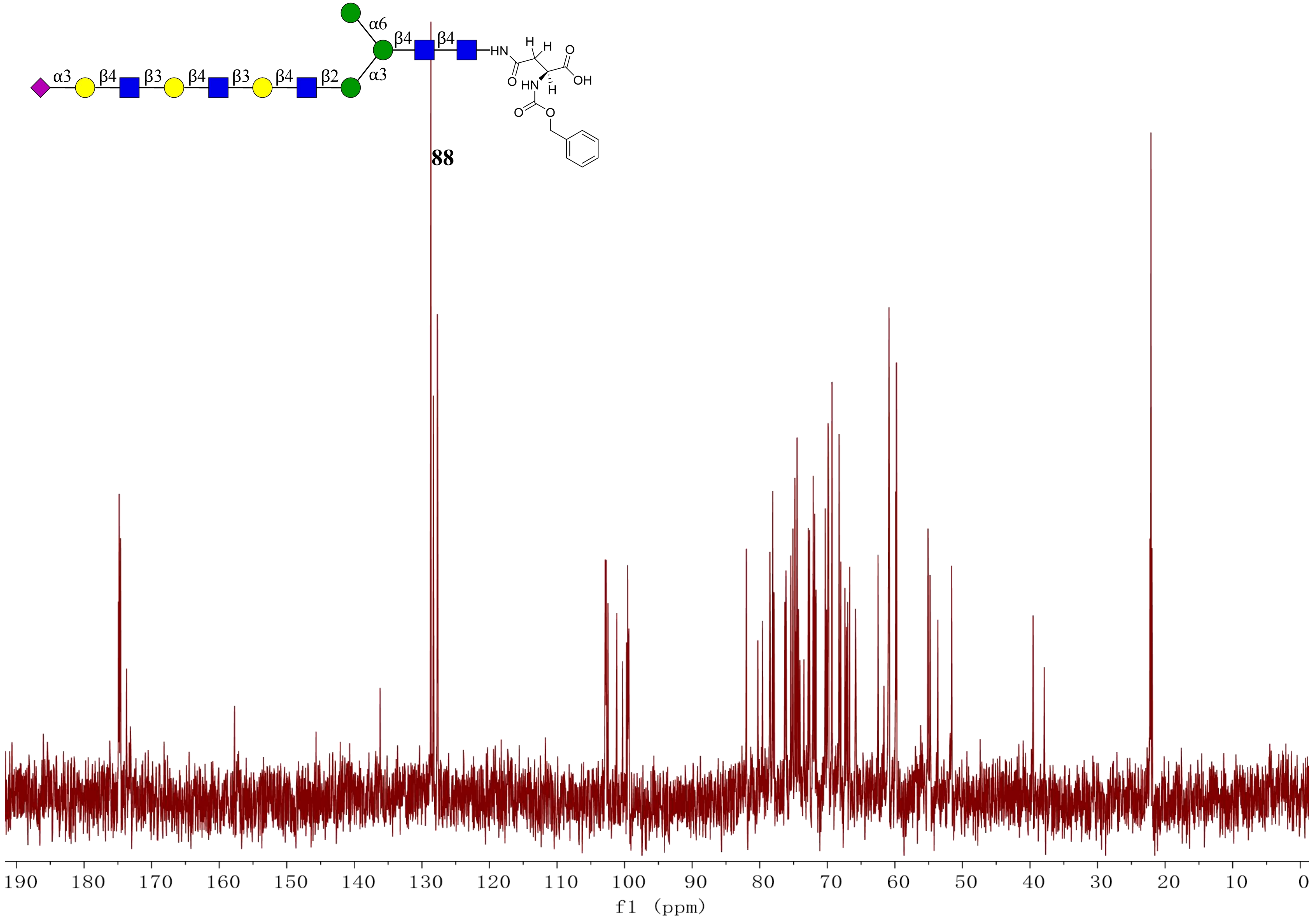
α6
β4
β4
HN
H
H
O
α3
β4
β3
β4
β3
β4
β2
α3
O
HN
H
OH
O
O
88
190 180 170 160 150 140 130 120 110 100 90 80 70 60 50 40 30 20 10 0
f1 (ppm)

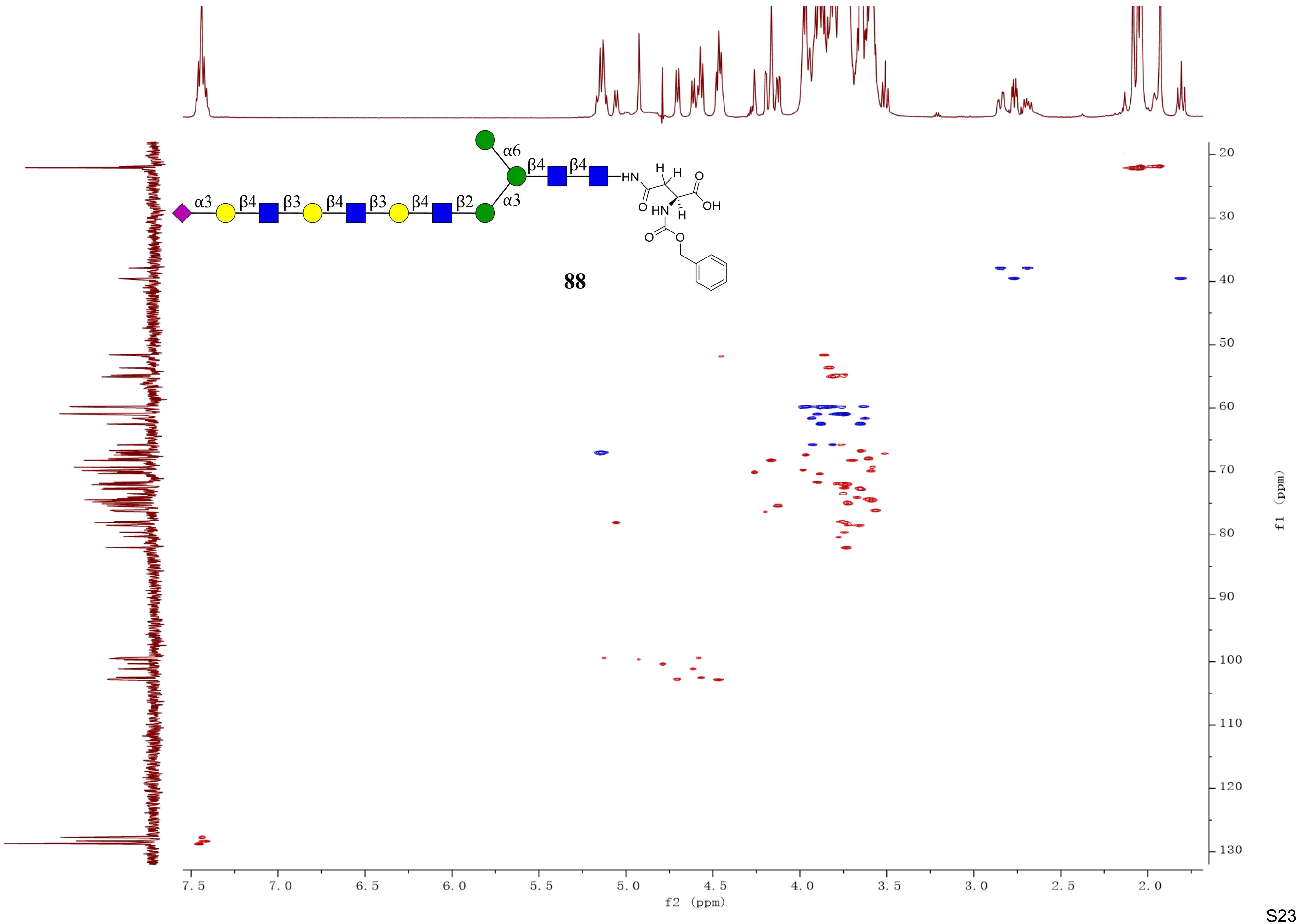
α6
β4
β4
α3
β4
β3
β4
β3
β4
β2
α3
HN
H
H
O
O
OH
HN
H
O
O
88
f2 (ppm)
f1 (ppm)
